# Chemoenzymatic Synthesis of Heparan Sulfate Oligosaccharides having a Domain Structure

**DOI:** 10.1101/2022.07.27.501676

**Authors:** Lifeng Sun, Pradeep Chopra, Geert-Jan Boons

## Abstract

Heparan sulfate (HS) have domain structures in which regions that are substantially modified by epimerization and sulfation (NS domains) are interspersed by unmodified fragments (NA domains). There is data to support that the domain structure of HS can regulate protein binding, however, such a binding mode has been difficult to probe. Here, we report a chemoenzymatic methodology that can provide HS oligosaccharides composed of two or more NS domains separated by NA domains of different length. It is based on the chemical synthesis of a sulfated HS oligosaccharide that enzymatically could be extended by various GlcA-GlcNAc units and terminated in a GlcNAc-6N_3_ moiety. HS oligosaccharides having an azide and alkyne moiety could assembled by copper catalyzed alkyne-azide cycloaddition (CuAAC) to give compounds having various NS domains separated by unsulfated regions. Competition binding studies showed that the length of an NA domain modulates the binding of the chemokines CCL5 and CXCL8.

## Introduction

Heparan sulfate (HS) are highly *O*- and *N*-sulfated carbohydrates that reside on the cell surface and in the extracellular matrix (ECM) of virtually all mammalian cell types where they can interact with chemokines and cytokines, growth factors, blood coagulation factors, proteins of the complement pathways, and cell adhesion proteins.^[1]^ The interaction between HS and these proteins mediates biological processes such as cell–cell and cell–matrix interactions, cell migration and proliferation, growth factor sequestration, chemokine and cytokine activation, and tissue morphogenesis during embryonic development. Alteration in HS expression has been associated with diseases such as cancer, inflammation, neurological disorders, cardiovascular and infectious diseases.^[2]^

HS is biosynthesized on core proteins by the formation of a polymer composed of 1,4-linked repeating disaccharides of D-glucuronic acid (GlcA) and *N*-acetyl-D-glucosamine (GlcNAc)^[1f]^ that are subsequently modified by a series of enzymatic transformations including *N*-deacetylation, *N*-sulfation, epimerization of GlcA to L-iduronic acid (IdoA), and sulfation of the C-2 hydroxyl of IdoA and the C-6 hydroxyl of GlcNAc moieties. Occasional sulfation of the C-3 position of GlcN also occurs. The epimerization and sulfation proceed only partially resulting in substantial structural diversity. This diversity is not random and cells have an ability to create specific HS-epitopes by regulating the expression of specific isoforms of HS biosynthetic enzymes.^[1d,3]^ The premise of the “*HS sulfate code hypothesis”* is that such epitopes can recruit specific HS-binding proteins, thereby influencing multiple biological and disease processes.

Additional structural complexity in HS arises from domain formation in which highly sulfated (*N*-sulfated, NS) regions are interspersed with regions that have undergone no- or very limited modifications (*N*-acetylated, NA) and consist mainly of GlcA-GlcNAc repeating units (Fig. 1).^[1a,1g]^ It has been proposed that the spacing of NS domains can regulate proteins binding.^[1g,4]^ For example, proteins such as CXCL5, CCL8, interferon (IFN)-γ, PF4, and MIP-1α, occur at physiological concentrations as dimers or higher oligomers,^[1c,1f,4a,4c,5]^ and a binding model has been proposed in which each monomer binds to a distinct NS domain resulting in di- or multivalent binding interactions.^[4a-c]^ Analysis of HS-saccharides obtained by affinity purification using several different chemokines indicate they require NA domains of defined length for optimal binding.^[4a-c,4e]^

**Figure 1.**
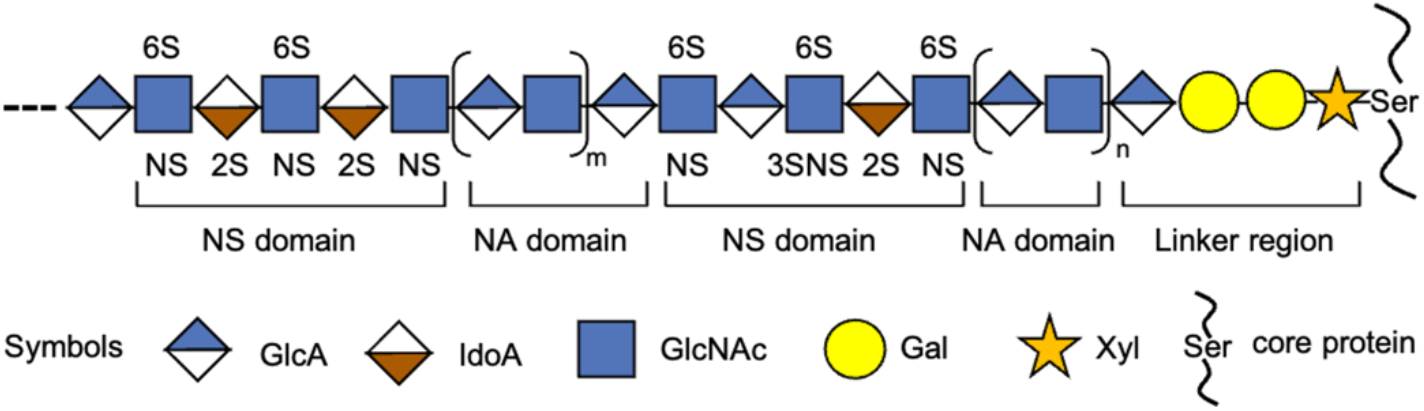
Heparan sulfate domain architectures. HS has a domain structure in which sulfated fragments (NS) are interspersed with unmodified fragments (NA).

Domain organization of HS impacts biological processes, and for example, inactivation of 2-OST changed the domain structure of endothelial HS, which led to a substantial increase in the number of binding sites for CXCL8, which in turn led to acute inflammatory responses in mice.^[4d,6]^ Biochemical data indicate that the bifunctional enzyme *N*-deacetylase/*N*-sulfotransferase isoform 1 (NDST-1) enzyme, which can convert GlcNAc into *N*-sulfoglucosamine (GlcNS) residues, plays a role in domain formation. It appears that *N*-deacetylation is a rate determining step in the modification of HS and because of substrate specificities of enzymes that perform subsequent modification, IdoA and *O*-sulfate residues accumulate in domains of contiguous *N*-sulfated (NS) disaccharide units.^[7]^

Progress in chemical,^[8]^ enzymatic^[9]^ and chemoenzymatic synthesis^[10]^ of HS oligosaccharides has provided collection of compounds that makes it possible to probe the importance of sulfation patterns of NS domains for protein binding. Well-defined synthetic oligosaccharides have also been displayed on dendrimers, polymers and nanoparticles to recapitulate biological activities of heparin.^[11]^ Although these materials demonstrate the importance of multivalent display of HS epitopes for binding and biological activity, they do not recapitulate the NS-NA domain organization of natural heparan sulfate and cannot probe the importance of spacing of NS domains for binding and biological activity. To examine the importance of spacing between two NS domains, a sulfated hexasaccharide having an allyl ether at the anomeric center was chemically synthesized and dimerized by an UV-promoted thiol-ene reaction with α,ω,-bis-(thio)oligo(ethyleneglycol) spacers of different length.^[12]^ The resulting compounds inhibited the binding of IFNγ/heparin in a length dependent manner. To better mimic NA domains of HS, an oligosaccharide was enzymatically assembled composed of regions with GlcNAc and trifluoracetyl glucosamine (GlcNTFA) using the glycosyl transferases KfiA and PmHS2 in combination with UDP-GlcNTFA or UDP-GlcNAc and UDP-GlcA.^[13]^ The TFA moieties could selectively be removed and the resulting amines can be enzymatically sulfated to give a well-defined oligomer having GlcNAc and GlcNS moieties. Further epimerizations and *O*-sulfations resulted, however, in the formation of complex mixtures of products. Thus, the preparation of panels of well-defined HS-oligosaccharides having domain structures is still an unresolved goal.

Here, we report a chemoenzymatic methodology that can give access to HS mimetics composed of two or more well-defined sulfated domains separated by NA domains of defined length (Fig. 2). Competition binding studies by surface plasmon resonance (SPR) showed that the length of an NA domain can modulate the bind of the chemokines CCL5 and CXCL8. The resulting data was rationalized based on structural models of CCL5 and CXCL5. The approach is based on the modular chemical synthesis^[8c,8l]^ of a hexasaccharide that terminates in a GlcA moiety (*e*.*g*. **1**, Fig. 2A), which is an appropriate primer for enzymatic extension by the bi-functional glycosyl transferases *Pasteurella multocida* heparosan synthase (PmHS2)^[10b,14]^ that can act both as α1,4-*N*-acetylglucosaminyltransferase and β1,4-glucuronyltransferase by using UDP-GlcNAc and UDP-GlcA, respectively. Repetitive use of this enzyme module was expected to provide compounds having NA domains of different length. The final enzymatic step exploited the finding that PmHS2 tolerates a GlcNAc moiety having an azido group at the *C*-6 position^[10b]^ to give compounds such as **3**-**8** (n = 0-5) (Fig. 2B). Compound **1** is equipped with an anomeric aminopentyl linker which provided a chemical handle to install an alkyne moiety to afford compound **2** (Fig. 2A). It was envisaged that compounds such as **2** and **3**-**8** could be coupled in a controlled manner by copper catalyzed alkyne-azide cycloaddition (CuAAC)^[15]^ reaction to give derivatives such as **9**-**14** in which sulfated domains are separated by an NA domain of defined length (Fig. 2C). Repeating the process of enzymatic elongation and installation of an azido-containing GlcNAc moiety followed by CuAAC with **2** should then give HS mimetics having multiple NA- and NS-domains.

**Figure 2.**
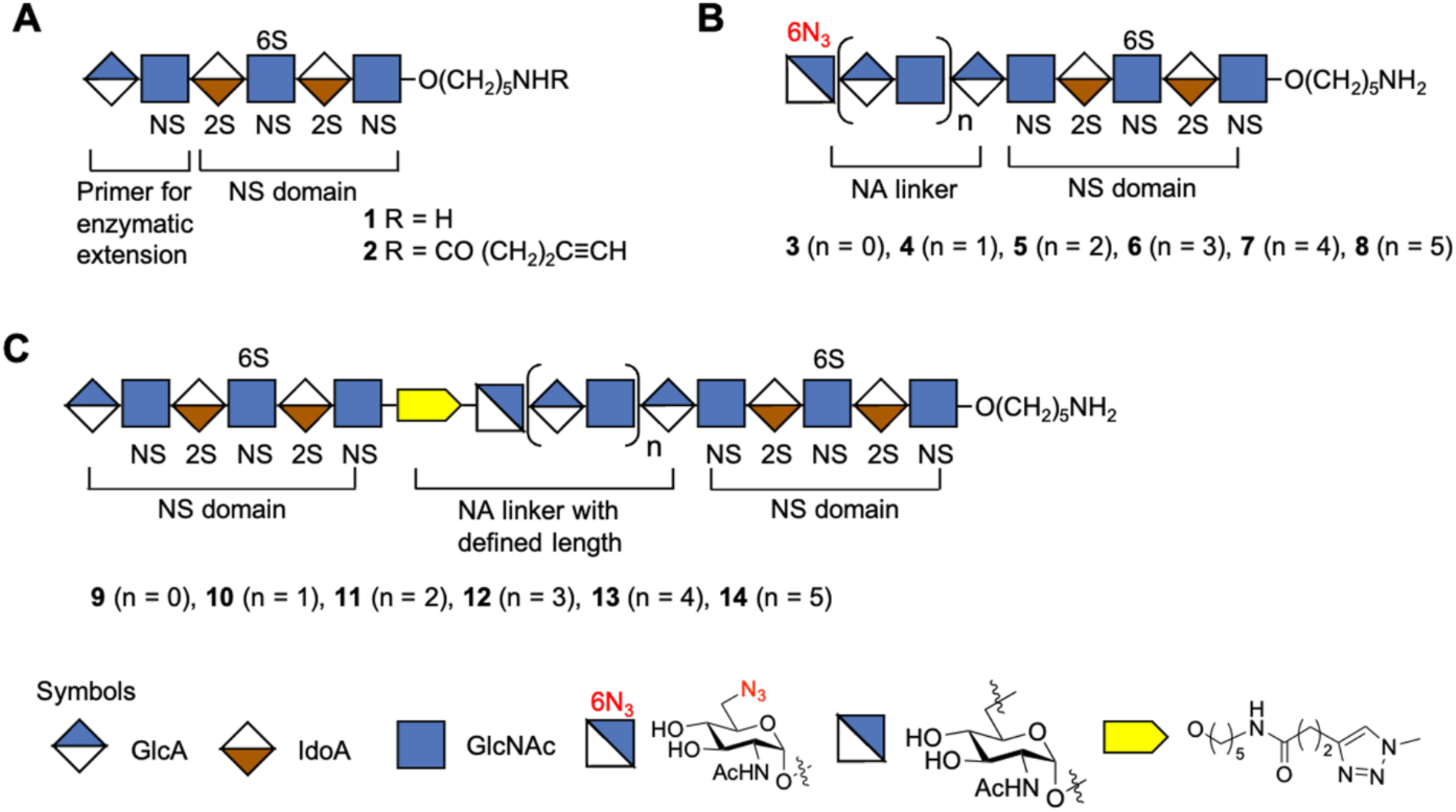
Synthetic strategy for the preparation of HS mimetics having well-defined NA and NS domains. (A) Hexasaccharide **1** that terminates in a GlcA moiety, which is a primer for PmHS2 enzymatic extension. Compound **2** is equipped with an alkyne moiety at the reducing end of **1**. (B) Enzymatically extended compounds **3, 4, 5, 6, 7**, and **8** that terminate in a GlcNAc-6N_3_ moiety and repeated GlcA-GlcNAc moiety. (C) CuAAC click reaction products **9, 10, 11, 12, 13**, and **14** in which NS domains are separated by an NA domain of defined length. Symbol nomenclature for HS backbone monosaccharides, structure of azide modified GlcNAc, and inter-domain triazole linkage is presented. 2S 2-*O*-sulfate, 6S 6-*O*-sulfate and NS *N*-sulfate.

## Results and Discussion

To establish the methodology, hexasaccharide primer **1** was assembled employing modular disaccharide building blocks **15**-**17** (Scheme 1).^[8j,8l]^ Thus, triflic acid (TfOH)-mediated coupling of **15** with **16** gave a tetrasaccharide that was converted into an acceptor by selective removal of the 9-fluorenylmethyl carbonate (Fmoc) followed by further glycosylation with glycosyl donor **17** to give hexasaccharide **18** (Scheme S1). Standard procedures were employed to replace the Fmoc protecting group of **18** by an acetyl ester to give compound **19**, which was treated with hydrazine acetate to remove the levulinoyl esters (Lev) and the resulting hydroxyls were sulfated with sulfur trioxide-pyridine complex (SO_3_·Py) in DMF to give **20**. The latter compound was treated with LiOH/H_2_O_2_ to remove the acetyl and methyl ester (→ **21**), which was followed by reduction of the azido moieties using trimethyl phosphine in THF/H_2_O to give free amines that were subjected to selective *N*-sulfation employing SO_3_·Py complex in MeOH/Et_3_N in the presence of NaOH (pH ∼ 11) to afford **22**. The target hexasaccharide primer **1** was obtained by hydrogenation of **22** over Pd(OH)_2_/C in a mixture of *tert*-butanol/H_2_O (1/1, v/v).

**Scheme 1.**
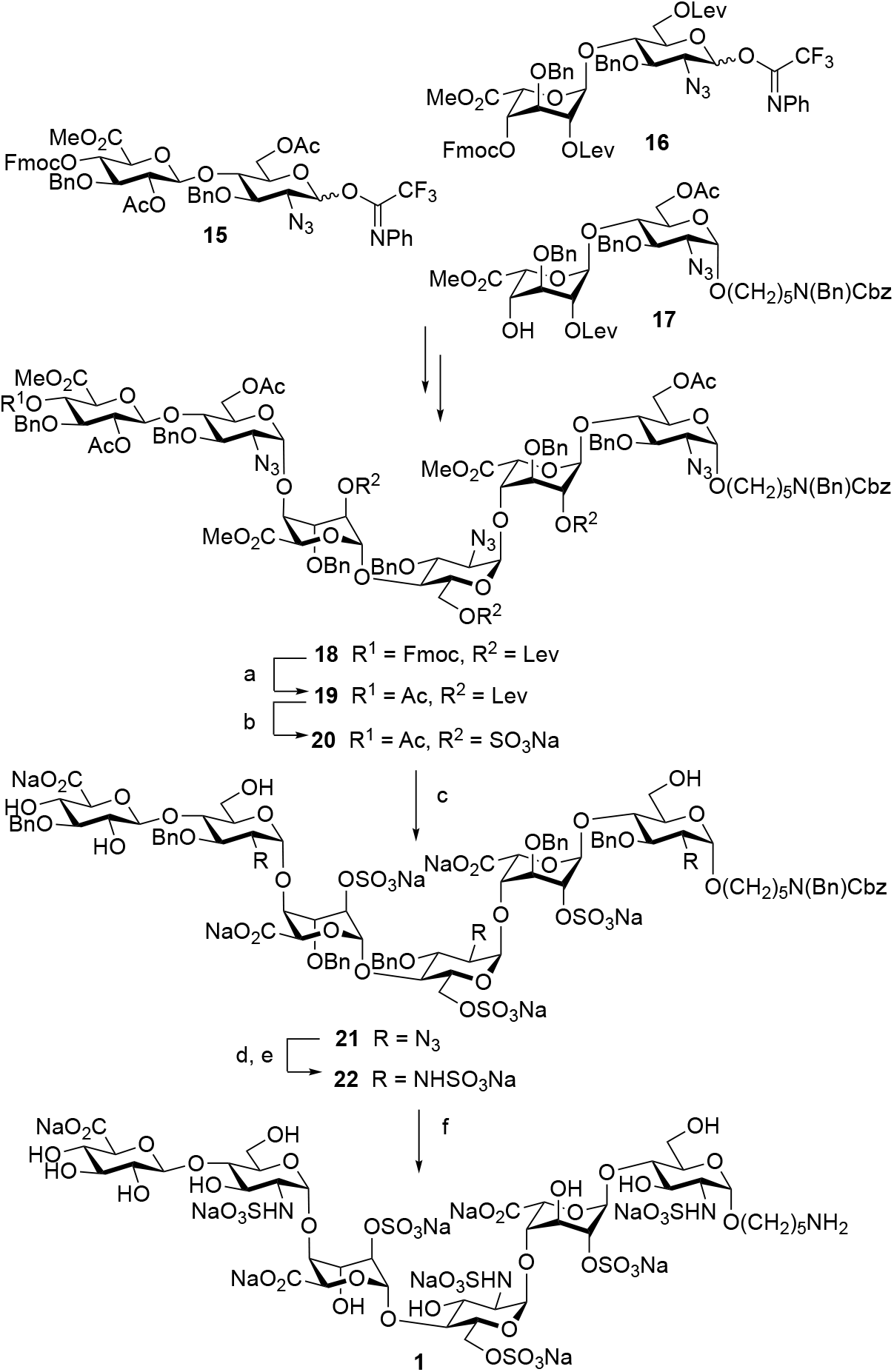
Chemical synthesis of hexasaccharide **1**. Reagents and conditions: (a) DCM/Et_3_N (4/1), 2 h; Ac_2_O, Py., 3 h; (b) NH_2_NH_2_·AcOH, Toluene/EtOH (1/2), 2 h; SO_3_·Py, DMF, 2 h; (c) 1.0 M LiOH, 30% H_2_O_2_, THF/H_2_O (1/1); 51% over five steps; (d) 1.0 M PMe_3_ in THF, 0.1 N NaOH, 2 h; (e) SO_3_·Py, MeOH, Et_3_N, 0.1 N NaOH; 50% over two steps; (f) Pd(OH)_2_/C, H_2_ (atm.), *t*-BuOH/H_2_O (1/1), 48 h; 85%.

Next, attention was focused on the enzymatic extension of **1** to give derivatives **23**-**27** having NA domains of different length (Scheme 2A). Thus, hexasaccharide **1** was converted into octasaccharide **23** (n = 1), first by exposure to UDP-GlcNAc in the presence of the bifunctional enzyme PmHS2 to install a GlcNAc moiety. Upon completion of the reaction as indicated by electrospray ionization mass spectrometry (ESI-MS), the compound was purified by size exclusion chromatography (SEC) over a P6 Biogel column and then re-exposed to PmHS2 in the presence of UDP-GlcA and purification over P6 Biogel was repeated. Of note, intermediate purification was important to remove sugar nucleotides to prevent polymerization. The enzyme module and purification were repeated several times to give compounds **24, 25, 26** and **27**. As anticipated, compounds **1, 23**-**27** could readily be converted into derivatives **3**-**8** having a terminal 6-azido-GlcNAc moiety upon treatment with UDP-GlcNAc-6N_3_ in the presence of PmHS2.^[10b]^ In parallel, the aminopentyl linker of hexasaccharide **1** was functionalized with an alkyne moiety for CuAAC chemistry by treatment of *N*-hydroxysuccinamide (NHS)-activated 4-pentynoic acid in a mixture of 0.3 M NaHCO_3_/acetonitrile/MeOH to give **2** (Scheme 2B).

Various reaction conditions were explored to link compound **2** with **3** by CuAAC to give HS mimetic **9** (Scheme 2C). The reactions were carried out in a 0.1 M ammonium bicarbonate buffer (pH = 8) to ensure that sensitive sulfate moieties stayed intact. It was observed that CuAAC exploiting sodium ascorbate and CuSO_4_ led to by-product formation probably because the initial oxidation product, dehydroascorbate, can hydrolyze to form reactive aldehydes such as 2,3-diketogulonate and glyoxal that can react with the free amino group of the anomeric linker.^[15a,16]^ Previously, it was reported that such side reactions can be avoided by employing aminoguanidine and tris(3-hydroxypropyltriazolylmethyl)amine (THPTA) as stabilizing agents for Cu(I) and sacrificial reductant to protect the biomolecules from oxidation.^[15a]^ When the CuAAC reaction was performed in the presence of these stabilizing agents at room temperature for 1 h, no degradation was observed, however only a trace amount of **9** was formed and mainly starting material was recovered. When the reaction was performed at a higher temperature (37 °C) for an extended period (24 h), it proceeded readily and desired compound **9** was isolated in an isolated yield of 61% after purification by SEC over P6 Biogel and sodium exchange using Dowex® 50 × 8Na^+^ resin. In a similar way, HS mimetics **10**-**14** were synthesized by CuAAC of **2** with **4**-**8**, respectively.

**Scheme 2.**
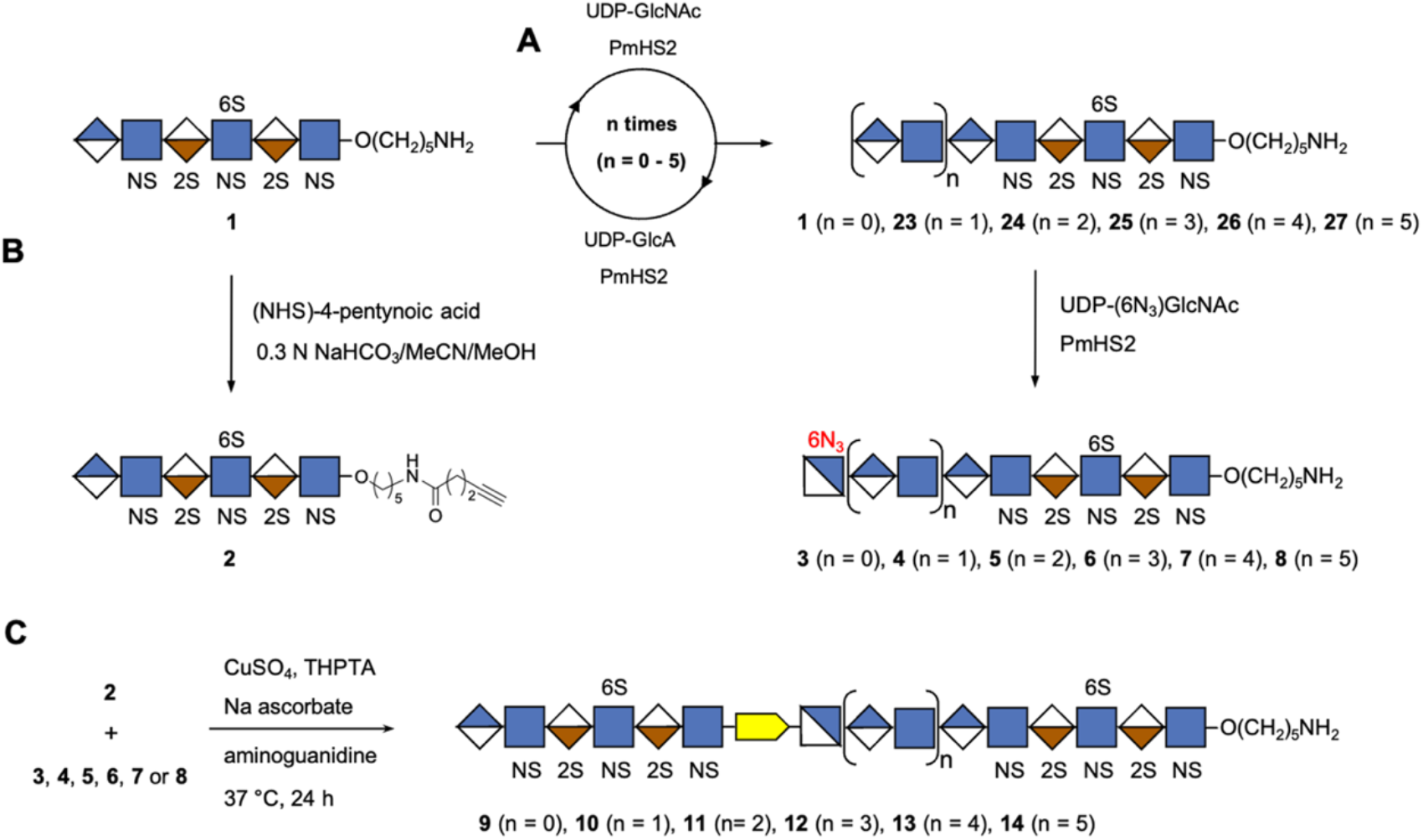
Chemoenzymatic synthesis of HS mimetics having two NS domains separated by an NA domain of different length. (A). Compound **1** was transformed to **23**-**27** by repeated enzymatic extension using PmHS2 in combination with UDP-GlcNAc and UDP-GlcA step by step, resulting compounds **23**-**27** and unmodified **1** were capped with GlcNAc-N_3_ to obtain **3**-**8**. (B). Compound **1** was functionalized with an alkyne moiety by treatment of *N*-hydroxysuccinamide (NHS)-activated 4-pentynoic acid in a mixture of 0.3 M NaHCO_3_/acetonitrile/MeOH (5/5/1) to give **2**. (C). Compounds **9**-**14** with two NS domains separated by an NA domain of defined length were obtained by CuAAC click reaction of **2** with **3**-**8**, respectively.

The HS derivatives **1, 9**-**14** were transformed into the sodium form by treatment with Dowex® 50 × 8Na^+^ resin and then analyzed by ESI-MS and nuclear magnetic resonance spectroscopy (NMR). ^1^H NMR spectra were fully assigned by one-dimensional (1D) and 2D NMR spectroscopy including ^1^H-^1^H correlation spectroscopy (COSY), ^1^H-^13^C heteronuclear single quantum coherence spectroscopy (HSQC), ^1^H-^13^C heteronuclear multiple bond correlation spectroscopy (HMBC), ^1^H-^1^H total correlation spectroscopy (TOCSY), and ^1^H-^1^H nuclear overhauser effect spectroscopy (NOESY). The anomeric configuration was confirmed by ^1^*J*_C1,H1_ coupling constants (^1^*J*_C1,H1_ ∼175 Hz for α linkage) and ^13^C chemical shifts of C1 (< 100 ppm for α linkage). The ^1^H NMR spectra confirmed the formation of a 1,2,3-triazole ring, and for example a unique aromatic proton at ∼7.85 ppm was observed arising from the triazole ring (Fig. 3A and 3D). Further support came from downfield shift of H6 and H5 (Fig. 3C-3D, H6 from ∼3.6 to ∼4.7 ppm, H5 from ∼3.9 to 4.1 ppm) and upfield shift of H4 and H1 (Fig. 3C-3D, H4 from ∼3.5 to ∼2.8 ppm, H1 from 5.44 to 5.35 ppm) of sugar protons connected to the triazole. In addition, the disappearance of alkyne proton and downfield shift of CH_2_ protons that are linked to the triazole (Fig. 3B and 3D and Fig. S1-S2.) confirm product formation.

**Figure 3.**
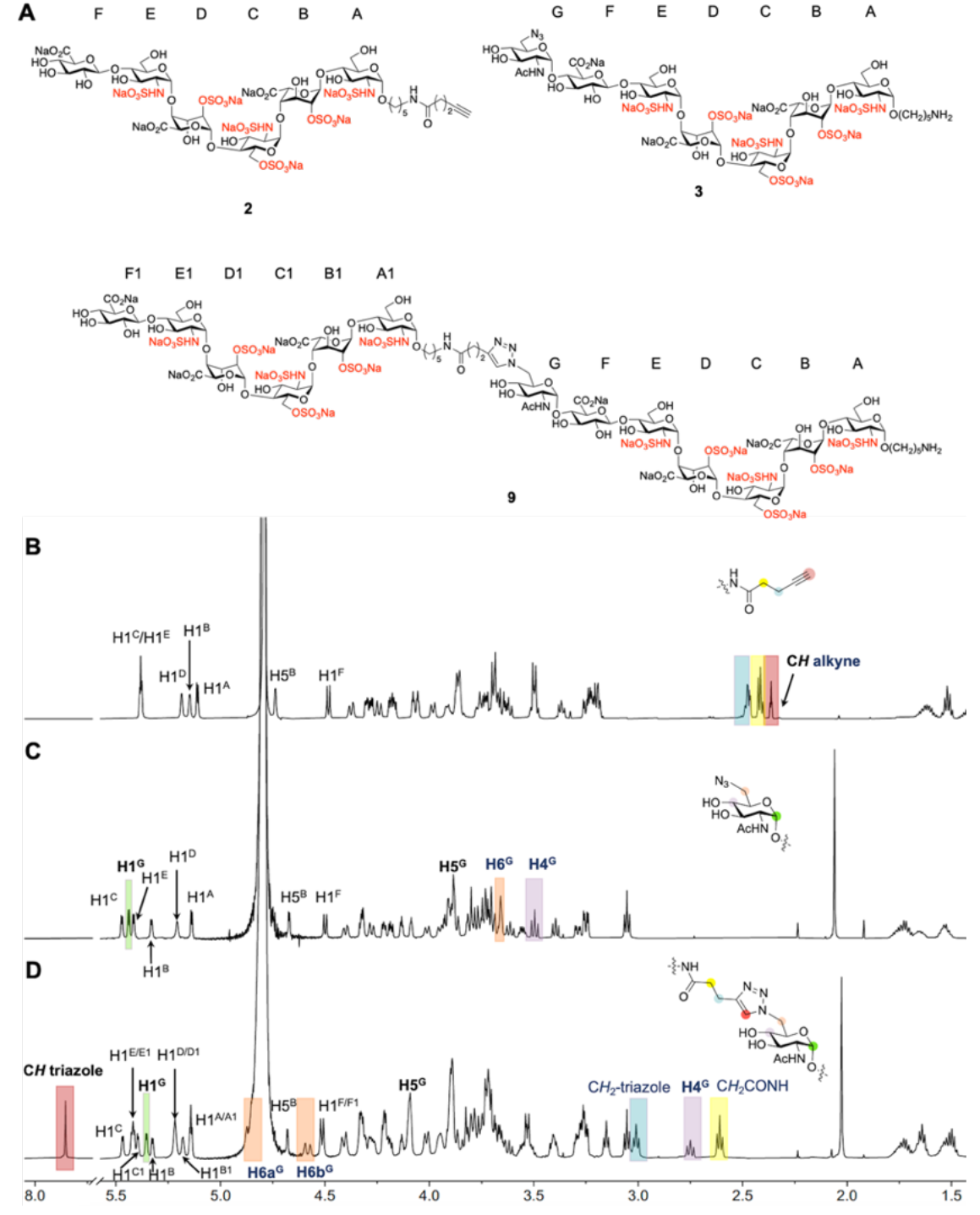
Analysis of synthetic compounds by NMR spectroscopy. ^1^H NMR stacked plots and structures of compounds **2, 3**, and their CuAAC product **9**. (A) Structures of compounds **2, 3**, and **9**, sugar rings are labelled alphabetically, starting from reducing to non-reducing end. (B) ^1^H NMR of hexasaccharide **2** where anomeric linker is extended with alkyne functionality; characteristic protons are annotated. (C) ^1^H NMR of compound **3** with terminal GlcNAc-6N_3_; characteristic protons are annotated. (D) ^1^H NMR of CuAAC product of **2** and **3**, HS mimetic **9**. Blue highlighted area, presence of CH_2_ that is attached to alkyne of compound **2** (B) and presence of CH_2_ that is attached to triazole of compound **9** (D). Yellow highlighted area, presence of CH_2_ that is highlighted in the structure (B and D). Red highlighted area, presence of CH alkyne of compound **2** (B) and presence of CH triazole of compound **9** (D). Orange highlighted area, presence of H6 of GlcNAc-6N_3_ of compound **3** (C) and presence of H6 of sugar with triazole of compound **9** (D). Purple highlighted area, presence of H4 of GlcNAc-6N_3_ of compound **3** (C) and presence of H4 of sugar with triazole of compound **9** (D). Green highlighted area, presence of H1 of GlcNAc-6N_3_ of compound **3** (C) and presence of H1 of sugar with triazole of compound **9** (D).

To synthesize HS mimetics having three NS domains (Scheme 3), compounds **9, 10**, and **11** were transformed into alkyne linker containing derivatives **28, 29**, and **30**, respectively by treatment with NHS-activated 4-pentynoic acid (Scheme 3A). CuAAC of **3** with **28, 4** with **29**, and **5** with **30** gave HS mimetics **31, 32**, and **33**, respectively having three NS domains separated by two NA domains (Scheme 3B-3C). The HS mimetics were purified by SEC over P6 Biogel, converted into their sodium salts by treatment with Dowex® 50 × 8Na^+^ resin and fully characterized by NMR and ESI-MS.

**Scheme 3.**
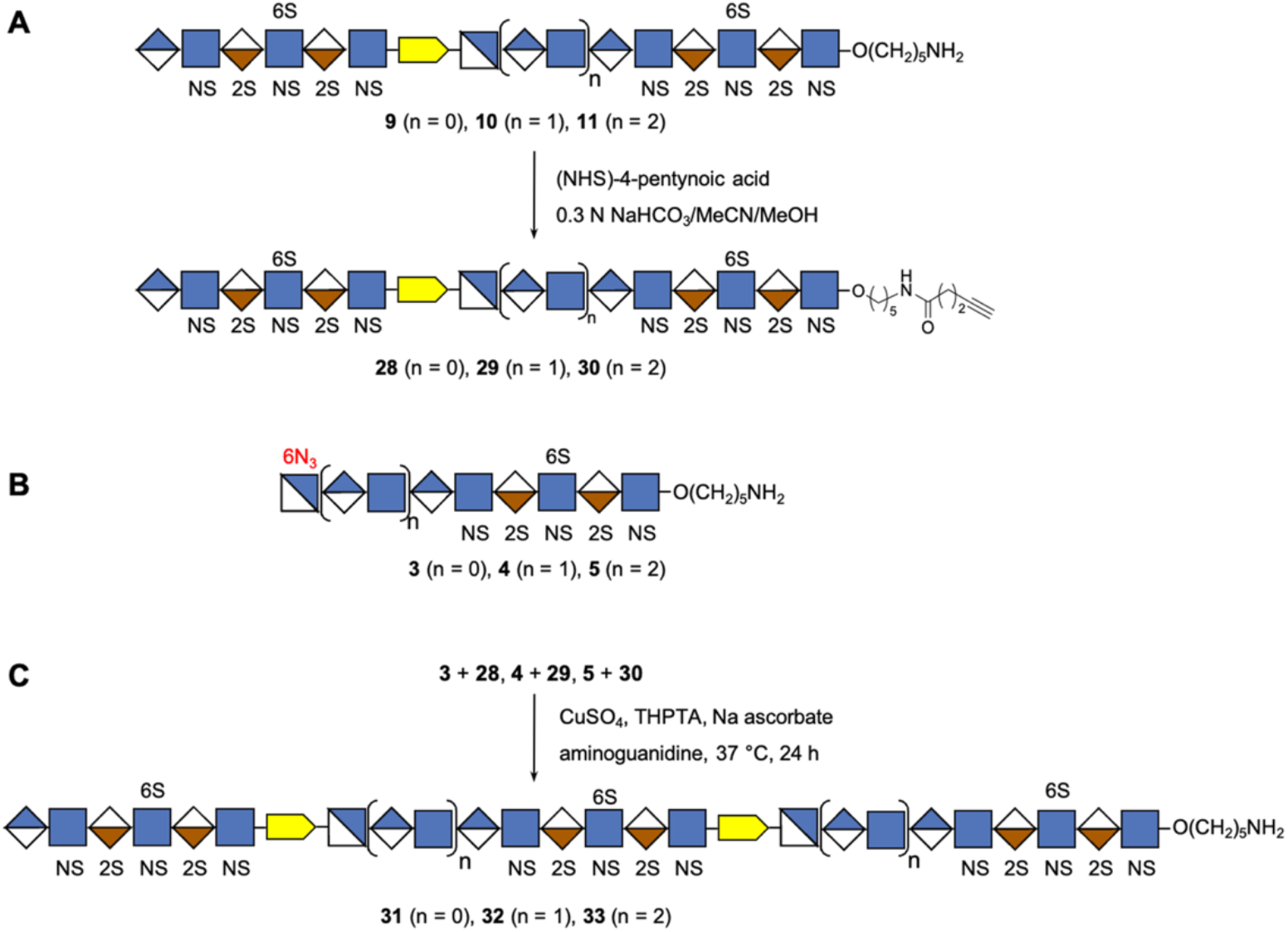
Chemoenzymatic synthesis of HS mimetics bearing three NS domains separated by well-defined NA domains. (A) Preparation of alkyne modified compounds **28, 29**, and **30** with two NS domains separated by a NA domain of defined length. (B) Structure of acceptors **3, 4**, and **5** with terminal azide group. (C) Assembly of **31**-**33** having three NS domains separated by two NA domains via CuAAC click reaction.

An SPR assay^[17]^ was employed to examine the binding of compounds **1, 9**-**14**, and **31**-**33** with the chemokines CXCL8 and CCL5 (Fig. 4). In this assay, biotinylated heparin was immobilized to a streptavidin-coated sensor chip and the binding of proteins of interest was inhibited by employing the synthetic compounds. First, binding experiments were performed using CXCL8 and CCL5 as analyte at different concentrations (Fig. 4B-4D). CXCL8 exhibited fast binding kinetics and therefore the equilibrium dissociation constant (K_D_) was determined by non-linear regression analysis of the steady-state binding responses at different protein concentrations, which gave a moderate affinity of 240 nM (Fig. 4C). CCL5 exhibited much slower binding kinetics and in this case fitting of the binding curves to a 1:1 Langmuir binding model gave a K_D_ of 14.7 nM (Fig. 4D). The measured K_D_ values are in agreement with previously reported data.^[18]^ Next, SPR inhibition experiments were performed by premixing 20 μM of mono-valent **1**, bi-valent **9**-**14**, and trivalent **31**-**33** with CCL5 or CXCL8 followed flow of the mixtures over the heparin modified sensor chip and monitoring of the response units (Fig. 4A-4B). Compounds that exhibited at least 50% inhibition were further evaluated at various concentrations to determine IC_50_. In the case of CXCL8, only compound **9** and **14**, having two NS domains separated by a very short or long NA domain, respectively and derivatives **31** and **32** having three NS domains showed substantially more potent inhibition compared to hexasaccharide **1** (Fig. 4E). CXCL8 is an 8 kDa proinflammatory chemokine that is produced by immune and non-immune cells to establish chemotactic gradients at infected or damaged endothelia.^[19]^ It is biologically active as a monomer but readily forms dimers in which the two binding sites are arranged in an anti-parallel manner. Affinity purification has indicated that the smallest heparin fragment that can bind in solution with appreciable affinity is an 18-mer, and a model was proposed in which two NS domains separated by an NA domain of sufficient length can engages with the two binding sites of the dimeric protein in horseshoe fashion over two antiparallel-oriented helical regions on the dimeric protein (Fig. 4F).^[4c,19]^ Compounds **14, 31** and **32** are expected to be sufficiently long to establish such a binding interaction. More recent NMR and molecular dynamic simulations have indicated that the binding interface of CXCL8 is structurally plastic and additional perpendicular binding modes were identified that require shorter HS oligosaccharides for high avidity binding (Fig. 4F).^[5a]^ It is conceivable that a compound such as **9**, having a very short spacer between the two sulfated domains, can bind in such a mode. Compounds in which sulfated domains are not appropriately spaced are expected to bind in a monovalent manner resulting in lower affinities. (Fig. 4F). HS can promote oligomerization of CXCL8 to form chemotactic gradients and therefore it is conceivable that HS can also promote the bind to two or more dimers.^[18]^

**Figure 4.**
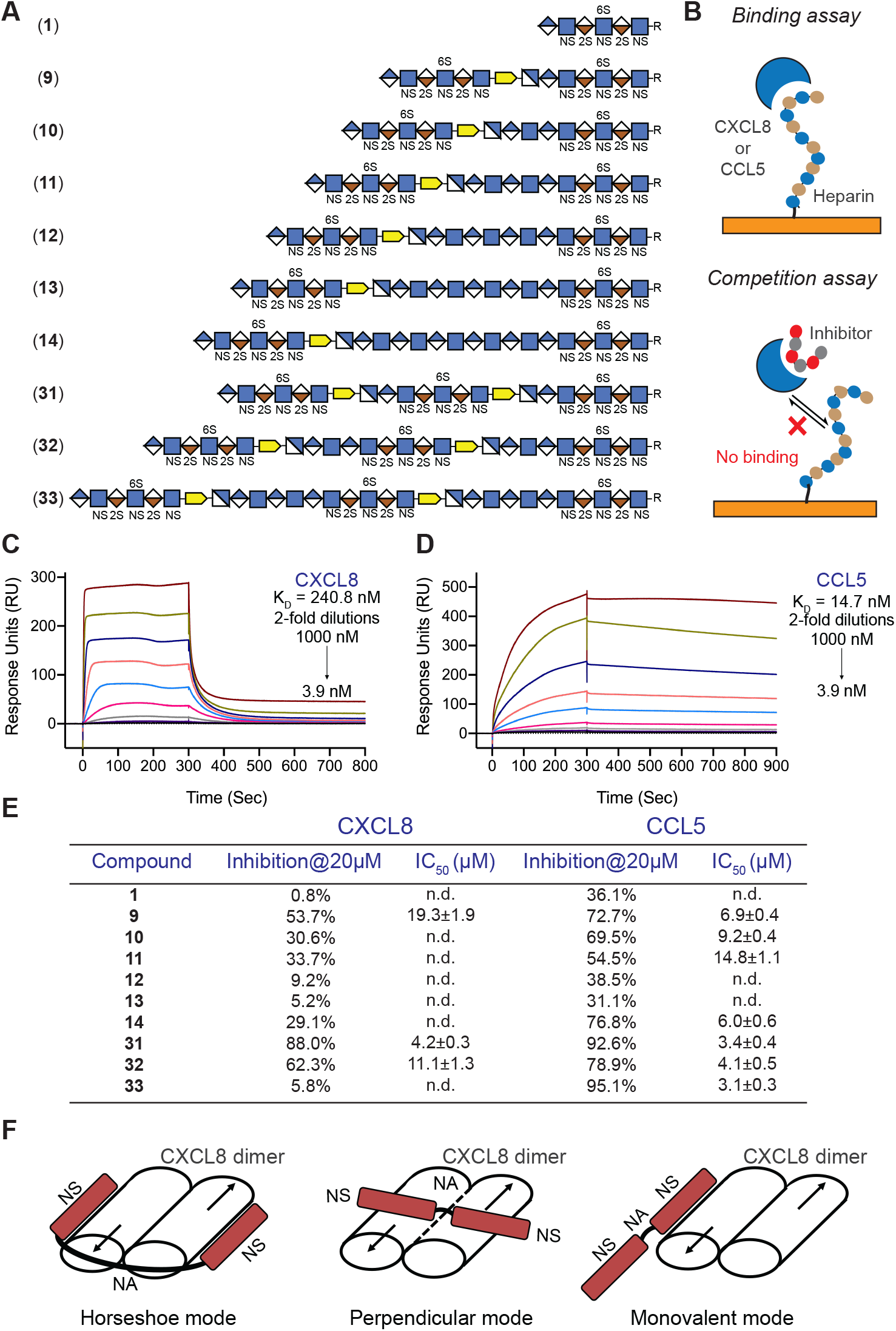
Surface Plasmon Resonance (SPR) binding assay and SPR competition inhibition assay. (A) Symbol structures of HS domain compounds employed in SPR competition inhibition assay, where R = O(CH_2_)_5_NH_2_. (B) Binding assay using a streptavidin-coated sensor chip on which biotinylated heparin was immobilized, CXCL8 or CCL5 as analyte at different concentrations. Competition assay was performed by premixing HS mimetics with CXCL8 or CCL5 followed by flowing the mixtures over the heparin modified sensor chip and monitoring the response units. (C) and (D) SPR sensorgrams representing the concentration-dependent kinetic analysis of the binding of immobilized heparin with CXCL8 (C) and CCL5 (D). Data were analyzed using Biacore T100 evaluation software. For steady state affinity analysis, fitting curves and detailed binding parameters see Supplementary Figs. S3 and S4. (E). SPR-based competition assay. Maximum inhibition observed (at 20 μM) and half-maximal inhibitory concentration (IC_50_) values of domain structures for CXCL8 and CCL5 binding to heparin functionalized surfaces. For individual inhibition curve see Supplementary Figs. S5 and S6. Data are presented as mean ± SEM (n = 3), all experiments were performed three times at the minimum. n.d.: not determined. (F). Different binding modes of CXCL8.

CCL5 is a proinflammatory chemokine that activates leukocytes through binding to the receptor CCR5. Although the monomeric form of CCL5 can induce cell migration *in vitro*, oligomerization is critical for *in vivo* activity. Oligomerization is promoted by interactions with glycosaminoglycans (GAGs), and mutants that cannot interact with GAGs are limited to induce cell migration *in vitro* but not *in vivo*.^[20]^ As can been seen in Fig. 4E, compound **9** which has two sulfated domains separated by a short unsulfated fragment inhibited the binding of CCL5 to the heparin chip more potently that hexasaccharide **1**. Compounds **10** and **11**, which have one or two additional GlcNAc-GlcA units in the unsulfated domain, had slightly lower activities compared to **9** whereas compounds **12** and **13** showed a further reduction in inhibitory activity and had similar responses compared to monovalent compound **1**. Compound **14**, which has the longest NA domain, showed a substantial increase in inhibition. HS mimetics with the shortest and longest NA domain were the most potent inhibitors of CCL5 indicating that the interaction of these chemokines with HS is complex involving different binding modes.

## Conclusion

A chemoenzymatic methodology is described that can provide, for the first time, well-defined HS mimetics that have multiple NS domains separated by NA domains of different length. It is based on the chemical synthesis of a sulfated HS oligosaccharide that is designed in such a way that it can be enzymatically extended by additional GlcA-GlcNAc moieties. In the last step of enzymatic synthesis, UDP-GlcNAc-6N_3_ is employed as glycosyl donor to install a terminal GlcNAc-6N_3_ moiety. The reducing end of compound **1** is equipped with an anomeric aminopentyl linker that made it possible to introduce an alkyne moiety and the resulting compound could be linked by CuAAC chemistry to the GlcNAc-6N_3_ moiety of the afore described oligosaccharides to give compounds having two sulfated domains separated by an unsulfated domain. The process of enzymatic NA introduction and click reaction could be repeated to give mimetics having three sulfated domains. The newly synthesized compounds represent the longest well-defined HS analogs ever prepared having various sulfated domains separated by unsulfated fragments. SPR inhibition studies showed that the length of NA domain influences protein binding in complex manners, which is in agreement with different binding modes of chemokines. It is to be expected that the synthetic methodology presented here in combination with various biophysical, computational, and biological studies will offer opportunities to examine the importance of HS domain structure for biological activity.

## Acknowledgements

This research was supported by the European Union’s Horizon 2020 Research and Innovation Programme (grant number 899687 (HS-SEQ) to G.-J.B.) and the Chinese Scholarship Council (to L.S.).

