## Supplementary material for "Chemoenzymatic Synthesis of Heparan Sulfate Oligosaccharides having a Domain Structure": SI

### 1. Experimental procedures.

**General experimental procedures.** Unless otherwise stated, all chemical reagents were acquired from commercial sources and utilized without additional purification. Molecular sieves (4 Å) were flame-dried prior to use. All moisture-sensitive processes were performed in an argon environment. Unless otherwise stated, all reactions at high temperatures were carried out in a silicon oil bath. Reactions were monitored by thin-layer chromatography (TLC) on silica gel-coated aluminum or glass plates (EMD Chemicals Inc.). Spots were visualized by UV light (254 nm) when applicable and charring with 10 % sulfuric acid in ethanol or a solution of $(NH_4)_6Mo_7O_{24}\cdot 4H_2O$ (24.0 g, 19.4 mmol) and $Ce(NH_4)_2(NO_3)_6$ (0.50 g, 0.9 mmol) in sulfuric acid (5%, 500 mL). Column chromatography was performed on silica gel G60 (Silicycle 60 – 200 µm, 60 Å). Fractions or reaction mixture containing sulfated compound were concentrated under reduced pressure with water bath temperature less than 25 °C. NMR spectra ($^1$H, $^{13}$C, COSY, HSQC, TOCSY, HMBC, NOESY) were recorded on an Agilent 400-MR DD2 or Bruker AVANCE-600 MHz spectrometer at 25 °C. Chemical shifts are reported in parts per million (ppm) relative to tetramethylsilane (TMS δ = 0 ppm) or deuterium oxide ($D_2O$ δ = 4.79 ppm) as the internal standard. NMR data is presented as follows: chemical shift, multiplicity (s = singlet, bs = broad singlet, d = doublet, bd = broad doublet, t = triplet, dd = doublet of doublet, m = multiplet and/or multiple resonances), q = quartet, integration, coupling constant in Hertz (Hz). NMR signals were assigned based on $^1$H NMR, $^{13}$C NMR, COSY, HSQC, HMBC, TOCSY or NOESY experiments. Due to the small sample size, carbon chemical shifts for compounds **1-14**, **21-27** and **31-33** were collected from the F1 dimension in the HSQC spectra. Mass spectra were obtained on a Bruker micrOTOF-Q II (ESI LC-MS). Reported HRMS data was obtained on an Agilent technologies 6560 Ion mobility Q-TOF. $^1$H NMR spectroscopy was used to determine the yield of the final compounds, with acetone as an internal reference. *Pasteurella multocida* heparosan synthase 2 (PmHS2) were expressed in E. coli as previously described.[1]

Abbreviations: Ac, acetyl; AcOH, acetic acid; BAIB, (Diacetoxyiodo)benzene; Bn, benzyl; Bu, butyl; Cbz, benzyloxycabonyl; Bz, benzoyl; CCL5, chemokine (C-C motif) ligand 5; $CDCl_3$, deuterated chloroform; CMPI, 2-Chloro-1-methylpyridinium iodide; CSA,

Camphorsulfonic acid; CuAAC, copper(I)-catalyzed alkyne-azide cycloaddition; CXCL8, C-X-C motif chemokine ligand 8; DABCO, 1,4-Diazabicyclo[2.2.2]octane; DCC, *N*,*N*'-Dicyclohexylcarbodiimide; DCM, dichloromethane; DMF, dimethyl formamide; $D_2O$, deuterated water; Et, ethyl; $Et_3N$, triethylamine; EtOAc, ethyl acetate; EtOH, ethanol; ESI-TOF, electrospray ionizaiton – time of flight; Fmoc, fluorenylmethoxycarbonyl; GlcA, D-glucuronic acid; GlcNAc, *N*-acetyl-D-glucosamine; $H_2O$, water; $H_2O_2$, hydrogen peroxide; HRMS, high resolution mass spectrometry; Lev, levulinoyl; LiOH, lithium hydroxide; Me, methyl; MeCN, acetonitrile; MeOH, methanol; $MgSO_4$, magnesium sulfate; $MnCl_2$, manganese dichloride; $N_3$, azide; $NaHCO_3$, sodium bicarbonate; NaOH, sodium hydroxide; NIS, *N*-Iodosuccinimide; NHS, *N*-Hydroxysuccinimide; $Pd(OH)_2/C$, palladium hydroxide over carbon; Piv, pivaloyl; PmHS2, *Pasteurella multocida* heparosan synthase; TEMPO, 2,2,6,6-Tetramethylpiperidine 1-oxyl; $SO_3$·Py, sulfur trioxide/pyridine complex; SPR, surface plasma resonance; TBAF, tetrabutylammonium fluoride; TDS, dimethyl thexylsilyl; TFA, trifluoroacid; TfOH, trifluoromethanesulfonic acid; THF, tetrahydrofuran; THPTA, Tris(3-hydroxypropyltriazolylmethyl)amine; TLC, thin layer chromatography; $TMSCHN_2$, (Trimethylsilyl)diazomethane; TMSOTf, Trimethylsilyl trifluoromethanesulfonate; Tol, toluene; Tris, Tris(hydroxymethyl)aminomethane; Ts, *p*-Toluenesulfonyl; UDP, Uridine diphosphate; v/v, volume/volume.

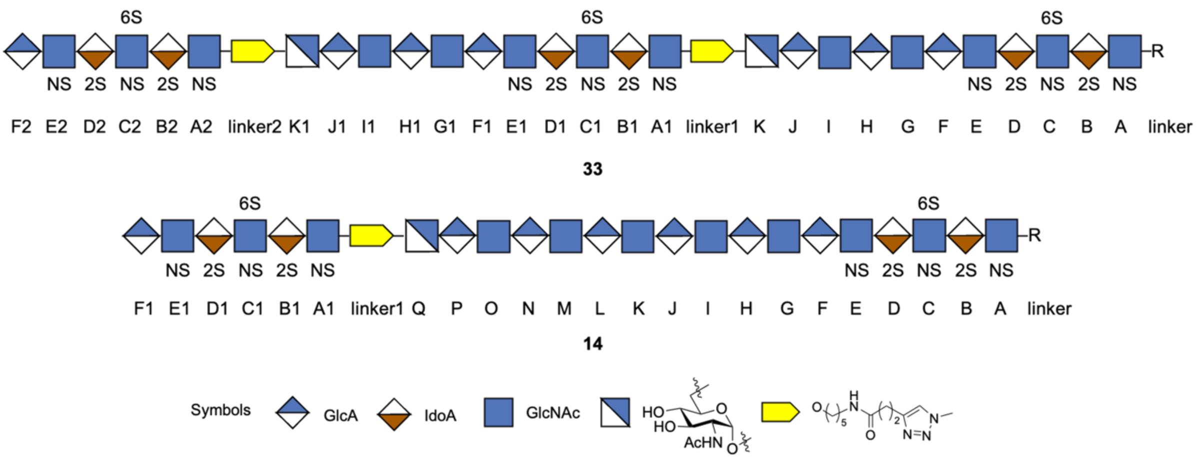

**Figure S1.** Monosaccharide nomenclature system for NMR assignments. For oligosaccharide nomenclature for NMR spectroscopy, the monosaccharide residues of the HS oligosaccharides have been labelled from the reducing end to non-reducing end in an alphabetical order for each domain and labelled together with 1, 2 for the repeated domains (*e.g.* compounds **33** and **14** labelled as above).

**Scheme S1.** Synthesis of hexasaccharide precursor.[a]

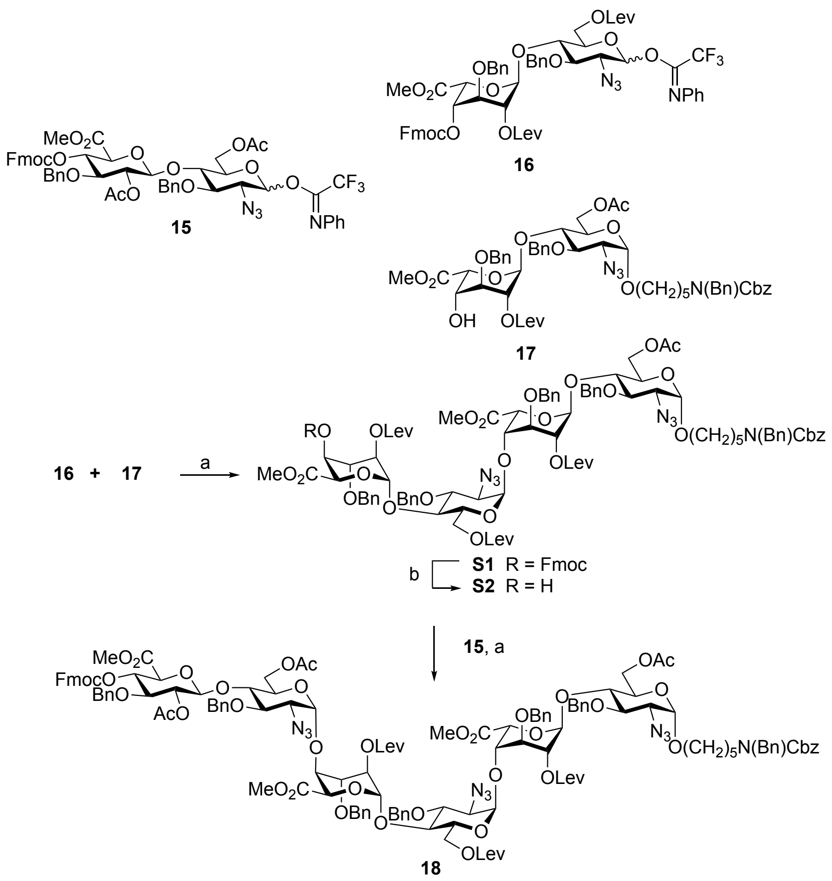

[a]Reagents and conditions: (a) TfOH, DCM, 4Å MS, -40 °C, 1 h, **S1**: 53%; **18**: 62%; (b) DCM/$Et_3N$, 2 h; **S2**: 90%.

**Scheme S2.** Synthesis of 6-Azido-GlcNAc-UDP (**S0**).[a]

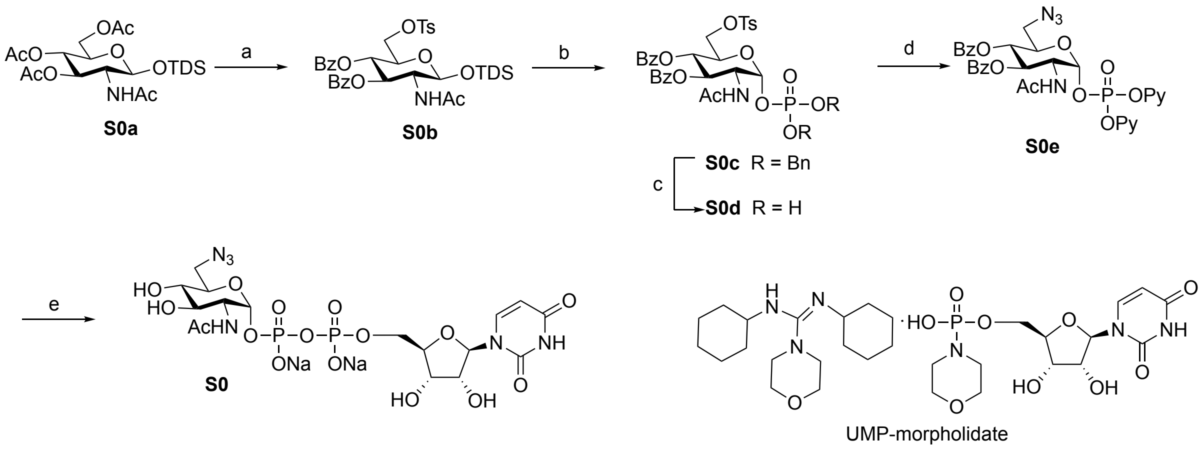

[a]Reagents and conditions: (a) i) NaOMe, MeOH; ii) TsCl, Py; iii) BzCl, Py; (b) i) TBAF·AcOH, THF; ii) tetrazole, dibenzyl *N*, *N* diisopropylphosphoramidite, DCM, 1.5 h; iii) mCPBA, -78 °C to r.t., 2 h; (c) Pd/C, $H_2$, MeOH, 12 h; (d) $Et_3N$; $NaN_3$, DMF, 60 °C, 24 h; (e) i) UMP-morpholidate, tetrazole, Py, 3 days; ii) NaOH ( pH= 12), 1 h.

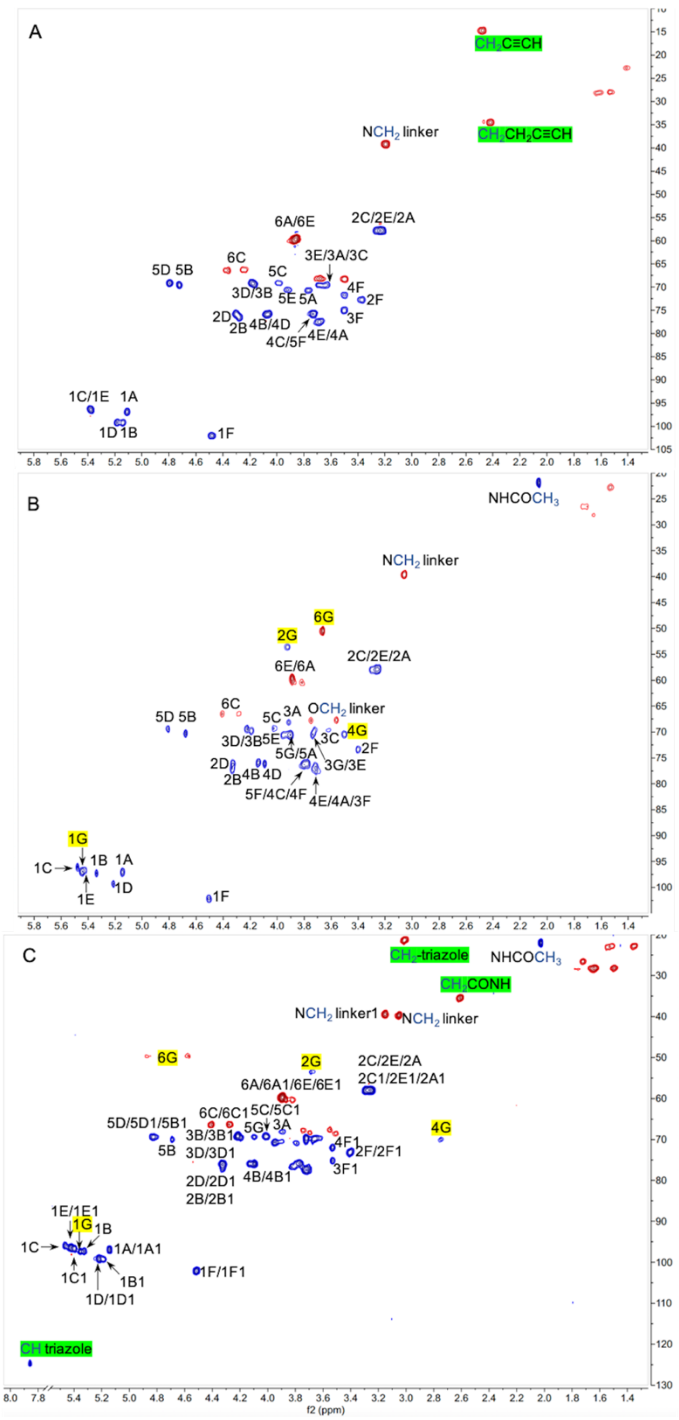

**Figure S2.** An example of structural analysis of synthesized compound by NMR spectroscopy. (A) Annotated HSQC spectra showing proton/carbon assignments of compound **2**. (B) HSQC and assignment of compound **3**. (C) HSQC and assignment of compound **9**. Yellow highlighted area, characteristic shift of sugar protons (H1, H4, H6) connected to the triazole; green highlighted area, characteristic shift of alkyne linker. ($CH_2$-triazole: $CH_2$ linked to triazole; CH triazole: CH of the triazole)

**General procedure for Lev esters removal.**

Hydrazine acetate (5 equiv. per Lev group) was added to a solution of the starting material in a mixture of DCM and MeOH (1/1, v/v, 0.02 M). The reaction mixture was stirred at room temperature for 2 to 4 h until TLC analysis (petroleum ether/EtOAc, 1/1 to 1/2, v/v) indicated completion of the reaction. The reaction mixture was diluted with DCM (30 mL), washed with water (25 mL), saturated bicarbonate solution (2 × 25 mL) and brine (25 mL), dried ($Na_2SO_4$) and filtered. The filtrate was concentrated under reduced pressure, and the residue was purified by silica gel column chromatography using petroleum ether and EtOAc as eluent (1/1 to 1/2, v/v), to give pure product.

**General procedure for *O*-sulfation.** To a solution of the starting material in DMF (0.15 M) was added $SO_3$·Py (10 equiv. per OH). The reaction mixture was stirred at room temperature for 2 to 4 h until TLC (DCM/MeOH, 90/10, v/v) indicated completion of the reaction. A mixture of triethylamine and MeOH (1/1, v/v, 1 mL) was added to the reaction mixture and stirring was continued for another 30 min. The reaction mixture was concentrated under reduced pressure, and the residue was applied to a column of Iatrobeads (5 g), which was eluted with a gradient of DCM and MeOH (from 95/5 to 85/15, v/v). Fractions containing product were concentrated under reduced pressure and the residue was passed through a column of Dowex® 50 × 8 $Na^+$ resin (0.6 cm × 5 cm) using MeOH as eluent, to give pure product.

**General procedure for saponification of methyl esters and de-*O*-acetylation.** A premixed solution of 30% solution of $H_2O_2$ in $H_2O$ (100 equiv. per $CO_2Me$) and 1.0 M LiOH (50 equiv. per $CO_2Me$) was added to a solution of the starting material in THF (0.02 M). The reaction mixture was stirred at room temperature for 12 h. Then, a 4.0 M solution of NaOH was added until pH ~ 14. Stirring was continued until LC-MS indicated completion of the reaction. After adjusting the pH to 8-9 by careful addition of AcOH, the solvents were removed under reduced pressure. The residue was dissolved in water and applied to a Reverse Phase C18 column (1.0 cm × 5 cm), which was eluted with a gradient of $H_2O$ and MeOH (from 90/10 to 40/60, v/v). Fractions containing product were concentrated under reduced pressure and the residue was passed through a column of Dowex® 50 × 8 $Na^+$ resin (0.6 cm × 5 cm) using $H_2O$ as eluent to give pure product.

**General procedure for installation of alkyne moiety.**

Compounds containing linker free amine (1.0 equiv.) were dissolved in aqueous 0.3 M $NaHCO_3$ (29 µL/µmol compound) and MeOH (6 µL/µmol compound), followed by addition of a solution of *N*-succinimido 4-pentynonate (3.0 equiv) in acetonitrile (29 µL/µmol compound). The reaction mixture was stirred at room temperature for 3 h. Bio-Gel P-6 (1.5 cm × 100 cm) was used for purification with 100 mM ammonium bicarbonate as eluent. The fractions containing compound were lyophilized and passed through a short column of Dowex® 50 × 8$Na^+$ resin (1.0 cm × 2 cm) with $H_2O$ as eluent. Appropriate fractions were lyophilized to give the desired product.

**General procedures for installation of α (1→4)-GlcNAc**.

HS glycan (1.0 equiv.) and UDP GlcNAc (1.5 equiv.) were dissolved in Tris buffer (25 mM, pH = 7.5) containing $MnCl_2$ (5 mM). *Pasteurella multocida* heparosan synthase (PmHS2) (final concentration of enzyme 200 µg/mL) was added to form a final concentration of 3.2 mM. The reaction mixture was incubated at 37 °C for 12 h. The progress of the reaction was monitored by ESI-LC-MS, on completion the reaction mixture was lyophilized and the residue was passed through Biogel P6 (1.5 cm × 50 cm or 1.5 cm × 120 cm) with 0.1 M ammonium bicarbonate as eluent. Appropriate fractions as indicated by ESI-LC-MS were lyophilized and passed through a short column of Dowex® 50 × 8$Na^+$ resin (1.0 cm × 2 cm) with $H_2O$ as eluent. Appropriate fractions were lyophilized to give the desired product.

**General procedures for installation of β (1→4)-GlcA.**

HS glycan (1.0 equiv.) and UDP GlcA (1.5 equiv.) were dissolved in Tris buffer (25 mM, pH = 7.5) containing $MnCl_2$ (5 mM). *Pasteurella multocida* heparosan synthase (PmHS2) (final concentration of enzyme 200 µg/mL) was added to form a final concentration of 3.2 mM. The reaction mixture was incubated at 37 °C for 12 h. The progress of the reaction was monitored by ESI-LC-MS, on completion the reaction mixture was lyophilized and the residue was passed through Biogel P6 (1.5 cm × 50 cm or 1.5 cm × 120 cm) with 0.1 M ammonium bicarbonate as eluent. Appropriate fractions as indicated by ESI-LC-MS were lyophilized and passed through a short column of Dowex® 50 × 8$Na^+$ resin (1.0 cm × 2 cm) with $H_2O$ as eluent. Appropriate fractions were lyophilized to give the desired product.

**General procedures for installation of unnatural α (1→4) 6-azido-GlcNAc.**

HS glycan (1.0 equiv.) and UDP 6-azido-GlcNAc (1.5 equiv.) were dissolved in Tris buffer (25 mM, pH = 7.5) containing $MnCl_2$ (5 mM). *Pasteurella multocida* heparosan synthase (PmHS2) (final concentration of enzyme 200 μg/mL) was added to form a final concentration of 3.2 mM. The reaction mixture was incubated at 37 °C for 12 h. The progress of the reaction was monitored by ESI-LC-MS, on completion the reaction mixture was lyophilized and the residue was passed through Biogel P6 (1.5 cm × 50 cm or 1.5 cm × 120 cm) with 0.1 M ammonium bicarbonate as eluent. Appropriate fractions as indicated by ESI-LC-MS were lyophilized and passed through a short column of Dowex® 50 × 8$Na^+$ resin (1.0 cm × 2 cm) with $H_2O$ as eluent. Appropriate fractions were lyophilized to give the desired product.

**General procedure for copper (I)-catalyzed azide-alkyne cycloaddition (CuAAC) reaction.** Compounds containing azide group (1.0 equiv.) and compounds containing alkyne group (1.2 equiv.) were dissolved in 0.1 M ammonium bicarbonate solution (0.4 mM for compounds containing azido) in a 0.5 mL or 1.5 mL centrifugal tube to form solution **A**. A premixed solution of $CuSO_4$ (1.25 equiv.) and THPTA (6.25 equiv.) were added to the solution **A** followed by the addition of aminoguanidine solution (final concentration 10 mM). After adding sodium ascorbate solution (freshly made 20 mg/mL, final concentration 10 mM), the reaction mixture was mixed well, securely closed to prevent more oxygen from diffusing in. The reaction mixture was in 37 °C incubator for 24 h. After lyophilization, the residue was dissolved in 0.1 M ammonium bicarbonate and applied to Biogel P6 (1.5 cm × 120 cm), which was eluted with 0.1 M ammonium bicarbonate. Appropriate fractions as indicated by ESI-LC-MS were lyophilized and passed through a short column of Dowex® 50 × 8$Na^+$ resin (1.0 cm × 2 cm) with $H_2O$ as eluent. Appropriate fractions were lyophilized to give the desired product. All the reactions were performed at a small scale (no more than 1.5 mL scale).

**2. Experimental procedures and analytical data of synthetic compounds.**

**Dimethylthexylsilyl *O*-(3, 4-di-*O*-benzoyl-6-*O*-*p*-toluenesulfonyl-2-acetamido-2-deoxy) -β-D-glucopyranoside (S0b).** To a solution of compound **S0a** (3.00 g, 6.13 mmol) in MeOH

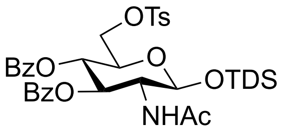

(45 mL) was added sodium methoxide solution (~30% in methanol) (0.2 mL). The reaction mixture was stirred at room temperature for 1 h and neutralized by the addition of Amberlite® IR 120 ($H^+$ form). The resin was filtered, and the filtrate was concentrated under reduced pressure. The residue was dissolved in anhydrous pyridine (13.5 mL) at argon atmosphere followed by the dropwise addition of *p*-toluenesulfonyl chloride (2.05 g, 10.73 mmol) of pyridine solution (0.45 M). The reaction mixture was stirred at 0 °C for 6 h. The reaction mixture was diluted with water (100 mL) and extracted with EtOAc (2 × 100 mL). The combined organic layers were dried ($Na_2SO_4$), filtered and the filtrate was concentrated *in vacuo*. The residue was purified by silica gel column chromatography using DCM and MeOH (30/1, v/v) as the eluent to give **Dimethylthexylsilyl-*O*-(2-acetamido-2-deoxy-6-*O*-*p*-toluenesulfonyl)-β-D-glucopyranoside** as a white amorphous solid (2.32 g, 73% over two steps, $R_f$ = 0.33 (DCM/MeOH, 10/1, v/v) which was dissolved in anhydrous pyridine (22 mL) and cooled down to 0 °C followed by the dropwise addition of benzoyl chloride (1.1 mL, 9.86 mmol). The reaction mixture was stirred at room temperature for 3 h and extracted with EtOAc (2 × 150 mL). The combined organic layers were dried ($Na_2SO_4$), filtered and the filtrate was concentrated *in vacuo*. The residue was purified by silica gel column chromatography using petroleum ether and EtOAc (3/1, v/v) as the eluent to give compound **S0b** as a white amorphous solid (3.07 g, 69% over three steps). $R_f$ = 0.28 (petroleum ether/EtOAc, 3/1, v/v). $^1H$ NMR (400 MHz, $CDCl_3$) δ 7.93 – 7.77 (m, 4H, C*H* aromatic), 7.71 (d, *J* = 8.4 Hz, 2H, C*H* aromatic Ts), 7.51 – 7.28 (m, 6H, C*H* aromatic), 7.22 (d, *J* = 8.2 Hz, 2H, C*H* aromatic Ts), 6.00 (d, *J* = 9.0 Hz, 1H, N*H*Ac), 5.72 (dd, *J* = 10.5, 9.6 Hz, 1H, H3), 5.33 – 5.26 (m, 1H, H4), 5.01 (d, *J* = 7.9 Hz, 1H, H1), 5.43 (t, *J* = 9.9 Hz, 1H, H4), 4.21 – 4.00 (m, 4H, H6a, H6b, H5, H2), 2.37 (s, 3H, $CH_3$ Ts), 1.78 (s, 3H, $CH_3$ Ac), 1.66 – 1.55 [m, 1H, $CH(CH_3)_2$], 0.88 – 0.81 [m, 12H,

$C(CH_3)_2$, $CH(CH_3)_2$], 0.15 & 0.12 [2s, 6H, $Si(CH_3)_2$]. $^{13}$C NMR (101 MHz, $CDCl_3$) δ 170.1 (<u>C</u>O Ac), 166.6 (<u>C</u>O Bz), 165.4 (<u>C</u>O Bz), 145.1, 133.6, 133.5, 132.4, 129.93, 129.90, 129.8, 129.0, 128.7, 128.5, 128.1, 96.1 (C1), 72.8 (C3), 72.1 (C5), 69.8 (C4), 68.5 (C6), 56.8 (C2), 34.1 [<u>C</u>H($CH_3$)$_2$], 24.9 [<u>C</u>($CH_3$)$_2$], 23.2 (<u>C</u>$H_3$ Ac), 21.7 (<u>C</u>$H_3$ Ts), 20.1 [d, $J$ = 2.9 Hz, 2C, CH(<u>C</u>$H_3$)$_2$], 18.6 [d, $J$ = 1.9 Hz, 2C, C(<u>C</u>$H_3$)$_2$], -2.7 [d, $J$ = 176.6 Hz, 2C, Si(<u>C</u>$H_3$)$_2$].

**Dibenzyl phosphate *O*-(3, 4-di-*O*-benzoyl-6-*O*-*p*-toluenesulfonyl-2-acetamido-2-deoxy) -β-D-glucopyranoside (S0c).** Compound **S0b** (1.72 g, 2.37 mmol) was dissolved in anhydrous

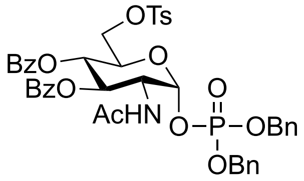

THF (47 mL) then cooled down to 0 °C, and sequentially added AcOH (170 µL, 2.84 mmol) and 1 M TBAF in THF (2.84 mL, 2.84 mmol). The reaction mixture was warmed up to room temperature and stirred for 2 h. The progress of the reaction was monitored by TLC until completion. After the addition of $Et_2O$ (100 mL), the organic layer was washed with brine (2 × 100 mL) and dried ($Na_2SO_4$), filtered and the filtrate was concentrated *in vacuo*. The residue was purified by silica gel column chromatography using petroleum ether and EtOAc (1/3, v/v) as the eluent to give lactol compound as a white amorphous solid (1.30 g, 94%). To a solution of the hemiacetal (1.30 g, 2.23 mmol) in anhydrous DCM (22.3 mL) was added tetrazole (469 mg, 6.69 mmol). Dibenzyl N, N diisopropyl phosphoramidite (1.1 mL, 3.35 mmol) was added dropwise to the solution and the reaction mixture was stirred at room temperature for 1.5 h followed by cooled down to -78 °C and mCPBA (70% wet, 1.38 g, 5.58 mmol) was added. The reaction mixture was allowed to warm up to room temperature and stirred for 2 h. The reaction mixture was diluted with 60 mL of diethyl ether, washed with ice cold saturated aqueous $Na_2S_2O_3$, saturated aqueous $NaHCO_3$, $H_2O$ and brine. The organic layers were dried ($Na_2SO_4$), filtered and the filtrate was concentrated *in vacuo*. The residue was purified by silica gel column chromatography using petroleum ether and EtOAc (1/1 to 1/4, v/v) as the eluent to give compound **S0c** as a white amorphous solid (1.70 g, 90% over two steps). $R_f$ = 0.54 (petroleum ether/EtOAc, 1/4, v/v). $^1$H NMR (400 MHz, $CDCl_3$) δ 7.90 – 7.80 (m, 4H, C*H* aromatic), 7.68 (d, $J$ = 8.2 Hz, 2H, C*H* aromatic Ts), 7.30 – 7.57 (m, 16H, C*H* aromatic), 7.18 (d, $J$ = 8.2 Hz, 2H, C*H* aromatic Ts), 5.91 (d, $J$ = 9.0 Hz, 1H, N*H*Ac), 5.71 (dd, $J$ = 6.2, 3.2 Hz, 1H, H1), 5.55 (dd, $J$ = 10.9, 9.6 Hz, 1H, H3), 5.43 (t, $J$ = 9.9 Hz, 1H, H4), 5.18

– 5.06 (m, 4H, 2 × C$H_2$Bn), 4.52 – 4.46 (m, 1H, H2), 4.30 (ddd, *J* = 10.3, 5.3, 2.2 Hz, 1H, H5), 4.10 (dd, *J* = 11.2, 2.3 Hz, 1H, H6a), 4.00 (dd, *J* = 11.2, 5.3 Hz, 1H, H6b), 2.33 (s, 3H, C$H_3$ Ts), 1.63 (s, 3H, C$H_3$ Ac). $^{13}$C NMR (101 MHz, $CDCl_3$) δ 170.4 (CO Ac), 166.9 (CO Bz), 164.9 (CO, Bz), 145.1 , 135.5 (d, *J* = 6.3 Hz, C Bn), 135.4 (d, *J* = 6.3 Hz, C Bn), 133.72, 133.71, 132.3, 130.0, 129.89, 129.87, 129.04, 129.02, 128.95, 128.91, 128.6, 128.5, 128.34, 128.27, 128.2, 96.1 (d, *J* = 6.6 Hz, C1), 70.4 (C3), 70.22 (d, *J* = 5.5 Hz, $CH_2$Bn), 70.15 (d, *J* = 5.5 Hz, $CH_2$Bn), 70.0 (C5), 68.0 (C4), 67.2 (C6), 52.2 (d, *J* = 7.8 Hz, C2), 22.8 ($CH_3$ Ac), 21.7 ($CH_3$ Ts). $^{31}$P NMR (162 MHz, $CDCl_3$) δ -2.51. HRMS (ESI-MS): m/z: calculated for $C_{43}H_{42}KNO_{13}PS$ $[M+K]^+$: 882.1746; found: 882.1719.

**Uridine diphosphate *O*-(6-azido-6-deoxy-2-acetamido-2-deoxy-)-α-D-glucopyranoside sodium salts (S0).** To a solution of compound **S0c** (1.00 g, 1.19 mmol) in MeOH and 10 % Pd/C (20% w/w) was added. The reaction was stirred at room temperature for 5 h under hydrogen atmosphere. The progress of the reaction was monitored by TLC until completion. The reaction mixture was filtered through celite and concentrated under reduced pressure. The residue was co-evaporated with $Et_3N$ to yield crude bis(trietylyammonium)phosphate compound which was dissolved in anhydrous DMF (10 mL) followed by the addition of $NaN_3$ (773 mg, 11.9 mmol). The reaction mixture was stirred at 60 °C for 24 h. The reaction mixture was filtered, and the filtrate was concentrated under reduced pressure. The residue was co-evaporated with anhydrous pyridine twice (5 mL) and dissolved in anhydrous pyridine (5.0 mL) again, UMP-morpholidate (800 mg, 1.31 mmol) and tetrazole (267 mg, 3.81 mmol) were added and the solvent was removed under reduced pressure. The residue was dissolved in anhydrous pyridine (6.0 mL) and stirred at room temperature for 3 days. The progress of the reaction was monitored by TLC and MALDI until completion. The solvent was removed under reduced pressure and the residue was purified by silica gel column chromatography using DCM/MeOH/$H_2O$ (70/30/5, v/v) as the eluent to give benzoylated 6 Azido GlcNAc-UDP as a white amorphous solid which was dissolved in aqueous NaOH (pH = 12) and stirred for 2 h. The solution was adjusted pH to 8 by careful addition of AcOH. The

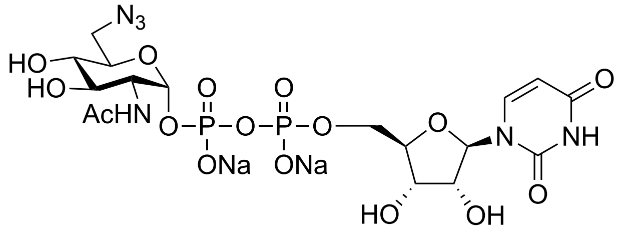

solution was lyophilized, and the residue was purified by Biogel P2 (2.0 cm × 100 cm) using $H_2O$ as the eluent, the fractions containing compound were lyophilized to give compound **S0** (289 mg, 36% over four steps) as a white amorphous solid. $^1$H NMR (600 MHz, $D_2O$) δ 7.96 (d, $J$ = 8.2 Hz, 1H, CO-C*H*=CH-N), 6.04 – 5.91 (m, 2H, CO-CH=C*H*-N, H1$^{UDP}$), 5.50 (dd, $J$ = 7.2, 3.4 Hz, 1H, H1$^{GlcNAc}$), 4.38 – 4.34 (m, 2H, H2$^{UDP}$, H3$^{UDP}$), 4.30 – 4.26 (m, 1H, H4$^{UDP}$), 4.26 – 4.17 (m, 2H, H5a$^{UDP}$, H5b$^{UDP}$), 4.05 (dt, $J$ = 10.1, 3.3 Hz, 1H, H5$^{GlcNAc}$), 4.01 (dt, $J$ = 10.5, 3.1 Hz, 1H, H2$^{GlcNAc}$), 3.79 (dd, $J$ = 10.5, 9.1 Hz, 1H, H3$^{GlcNAc}$), 3.72 (dd, $J$ = 13.7, 2.7 Hz, 1H, H6a$^{GlcNAc}$), 3.62 (dd, $J$ = 13.7, 4.0 Hz, 1H, H6b$^{GlcNAc}$), 3.58 (t, $J$ = 9.6 Hz, 1H, H4$^{GlcNAc}$), 2.07 (s, 3H, C*H*$_3$ Ac). $^{13}$C NMR from HSQC (151 MHz, $D_2O$) δ 141.6 (CO-$\underline{C}$H=CH-N), 102.7 (CO-CH=$\underline{C}$H-N), 94.3 (C1$^{GlcNAc}$), 88.4 (C1$^{UDP}$), 83.2 (C4$^{UDP}$), 73.7 (C3$^{UDP}$), 71.6 (C5$^{GlcNAc}$), 70.7 (C3$^{GlcNAc}$), 70.2 (C4$^{GlcNAc}$), 69.5 (C2$^{UDP}$), 64.9 (C5$^{UDP}$), 53.6 (C2$^{GlcNAc}$), 50.5 (C6$^{GlcNAc}$), 22.04 ($\underline{C}$H$_3$ Ac). The spectral data matches with previously reported compound.[2]

***N*-(Benzyl)-benzyloxycarbonyl-5-aminopentyl *O*-[methyl-2-*O*-levulinoyl-3-*O*-benzyl-4-*O*-(9-fluorenylmethyloxycarbonyl)-α-L-idopyranosyluronate]-(1→4)-*O*-(2-azido-3-*O*-benzyl-6-*O*-levulinoyl-2-deoxy-α-D-glucopyranosyl)-(1→4)-*O*-(methyl-2-*O*-levulinoyl-3-*O*-benzyl-α-L-idopyranosyluronate)-(1→4)-*O*-(2-azido-3-*O*-benzy-6-*O*-acetyl-2-deoxy-α-D-glucopyranoside (S1).** Freshly activated 4 Å molecular sieves was added to a solution of

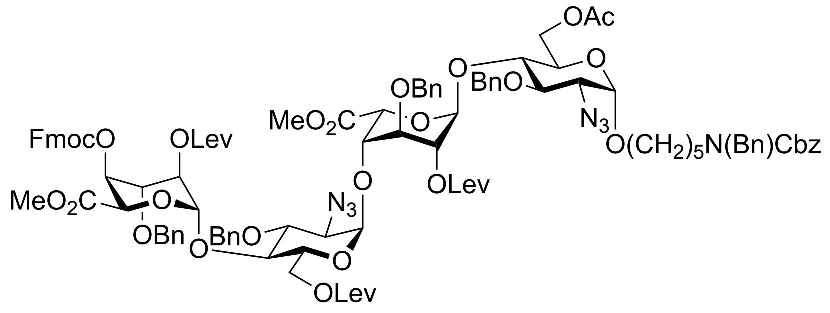

glycosyl donor **16**[3] (956 mg, 0.82 mmol) and glycosyl acceptor **17**[3] (701 mg, 0.68 mmol) in anhydrous DCM (13.7 mL, 0.05 M based on glycosyl acceptor). After stirring for 30 min at room temperature, the solution was cooled to -40 °C followed by the addition of TfOH (30 μL, 0.34 mmol). The reaction mixture was stirred for 1 h followed by quenching with pyridine. The reaction mixture was filtered, and the filtrate was concentrated *in vacuo*. The residue was purified by silica gel column chromatography using a stepwise gradient of petroleum ether and EtOAc (3/2 to 1/1, v/v) as the eluent to give disaccharide **S1** (725 mg, 53%). $R_f$ = 0.24 (petroleum ether/EtOAc, 1/1, v/v). $^1$H NMR (400

MHz, $CDCl_3$) δ 7.82 – 7.15 (m, 38H, CH aromatic), 5.25 (d, *J* = 3.9 Hz, 1H, $H1^{B}$), 5.17 (bd, *J* = 9.5 Hz, 2H, $C\mathit{H}_2$Cbz), 5.09 – 5.04 (m, 2H, $H1^{C}$, $H1^{D}$), 4.98 – 4.93 (m, 2H, $H4^{D}$, $H2^{B}$), 4.89 – 4.68 (m, 10H, $H5^{D}$, $H2^{D}$, $H1^{A}$, $H5^{B}$, 2 × $C\mathit{H}_2$Bn, 2 × C*H*HBn), 4.66 (d, *J* = 10.7 Hz, 1H, C*H*HBn), 4.60 (d, *J* = 10.7 Hz, 1H, C*H*HBn), 4.53 – 4.35 (m, 6H, N$C\mathit{H}_2$Bn, $H6a^{C}$, $H6a^{A}$, $C\mathit{H}_2$ Fmoc), 4.29 (d, *J* = 12.0 Hz, 1H, $H6b^{A}$), 4.23 – 4.14 (m, 2H, C*H* Fmoc, $H6b^{C}$), 4.03 (t, *J* = 4.6 Hz, 1H, $H4^{B}$), 3.98 – 3.79 (m, 7H, $H3^{B}$, $H4^{C}$, $H5^{A}$, $H4^{A}$, $H3^{D}$, $H3^{A}$, $H5^{C}$), 3.68 – 3.59 (m, 2H, $H3^{C}$, OCH*H* linker), 3.47 (s, 3H, $CO_2C\mathit{H}_3$), 3.44 (s, 3H, $CO_2C\mathit{H}_3$), 3.42 – 3.16 (m, 5H, OCH*H* linker, $H2^{A}$, $H2^{C}$, $C\mathit{H}_2$N linker), 2.86 – 2.46 (m, 12H, 6 × $C\mathit{H}_2$ Lev), 2.17 (s, 3H, $C\mathit{H}_3$ Lev), 2.11 (s, 6H, $C\mathit{H}_3$ Ac, $C\mathit{H}_3$ Lev), 2.04 (s, 3H, $C\mathit{H}_3$ Lev), 1.70 – 1.20 (m, 6H, 3 × $C\mathit{H}_2$ linker). $^{13}C$ NMR (151 MHz, $CDCl_3$) δ 207.0 ($CH_3\underline{C}O$ Lev), 206.4 ($CH_3\underline{C}O$ Lev), 206.2 ($CH_3\underline{C}O$ Lev), 172.3 ($CH_2\underline{C}O$ Lev), 172.2 ($CH_2CO$ Lev), 171.8 ($CH_2\underline{C}O$ Lev), 170.9 ($\underline{C}O$ Ac), 169.5 ($\underline{C}O_2CH_3$), 168.6 ($\underline{C}O_2CH_3$), 154.4, 143.3, 143.2, 141.43, 141.41, 138.2, 138.1, 137.7, 137.6, 137.3, 128.7, 128.62, 128.59, 128.35, 128.33, 128.27, 128.1, 128.02, 127.98, 127.9, 127.64, 127.61, 127.29, 127.27, 125.18, 125.13, 120.2, 98.2 ($C1^{B}$), 97.7 ($C1^{A}$), 97.4 ($C1^{D}$), 96.9 ($C1^{C}$), 78.4 ($C3^{A}$), 78.2 ($C3^{C}$), 76.0 ($C4^{A}$), 74.9 ($O\underline{C}H_2$Bn), 74.8 ($O\underline{C}H_2$Bn), 74.3 ($C3^{B}$, $C4^{C}$), 73.6 ($O\underline{C}H_2$Bn), 73.51 ($O\underline{C}H_2$Bn), 73.45 ($C3^{D}$), 72.0 ($C4^{B}$), 71.5 ($C4^{D}$), 70.3 ($\underline{C}H_2$ Fmoc), 69.7 ($C5^{C}$, $C5^{B}$, $C2^{B}$), 69.1 ($C5^{A}$), 68.3 ($O\underline{C}H_2$ linker), 68.2 ($C2^{D}$), 67.3 ($\underline{C}H_2$Cbz), 67.2 ($C5^{D}$), 63.4 ($C2^{A}$), 63.3 ($C2^{C}$), 62.5 ($C6^{A}$), 62.1 ($C6^{C}$), 52.3 ($CO_2\underline{C}H_3$), 51.9 ($CO_2\underline{C}H_3$), 50.7 ($N\underline{C}H_2$Bn), 50.4 ($N\underline{C}H_2$Bn), 47.2 ($\underline{C}H_2$N linker), 46.8 ($\underline{C}H$ Fmoc), 46.3 ($\underline{C}H_2$N linker), 38.1 ($\underline{C}H_2$ Lev), 37.8 ($\underline{C}H_2$ Lev), 37.7 ($\underline{C}H_2$ Lev), 29.9 ($\underline{C}H_3$ Lev), 29.8 ($\underline{C}H_3$ Lev), 29.6 ($\underline{C}H_3$ Lev), 29.1 ($\underline{C}H_2$ linker), 28.3 ($\underline{C}H_2$ Lev), 28.0 ($\underline{C}H_2$ Lev), 27.8 ($\underline{C}H_2$ Lev), 27.6 ($\underline{C}H_2$ linker), 23.4 ($\underline{C}H_2$ linker), 21.0 ($\underline{C}H_3$ Ac). HRMS (ESI-MS): m/z calculated for $C_{106}H_{117}N_7NaO_{32}$ $[M+Na]^+$: 2022.7635; found:.2022.7706.

***N*-(Benzyl)-benzyloxycarbonyl-5-aminopentyl *O*-(methyl-2-*O*-levulinoyl-3-*O*-benzyl-α-L-idopyranosyluronate)-(1→4)-*O*-(2-azido-3-*O*-benzyl-6-*O*-levulinoyl-2-deoxy-α-D-glucopyranosyl)-(1→4)-*O*-(methyl-2-*O*-levulinoyl-3-*O*-benzyl-α-L-idopyranosyluronate)-(1→4)-*O*-(2-azido-3-*O*-benzy-6-*O*-acetyl-2-deoxy-α-D-glucopyranoside (S2).** A solution of tetrasaccharide **S1** (725 mg, 0.36 mmol) in a mixture of DCM and $Et_3N$ (31.9 mL, 4/1, v/v) was stirred at ambient temperature for 2 h until TLC

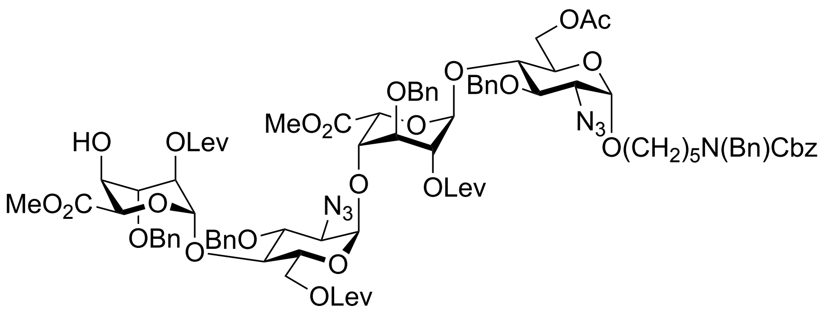

analysis indicated completion of the reaction. The mixture was concentrated under reduced pressure, and the residue was purified by silica gel column chromatography with petroleum ether and EtOAc (1/1, v/v) as the eluent to give pure product **S2** as a white amorphous solid (580 mg, 90%). $R_f$ = 0.36 (petroleum ether/EtOAc, 1/3, v/v). $^1$H NMR (400 MHz, $CDCl_3$) δ 7.77 – 6.56 (m, 30H, CH aromatic), 5.24 (d, *J* = 3.5 Hz, 1H, $H1^B$), 5.17 (bd, *J* = 10.5 Hz, 2H, $CH_2$Cbz), 5.05 (d, *J* = 3.5 Hz, 1H, $H1^C$), 4.99 (bs, 1H, $H1^D$), 4.95 (t, *J* = 3.5 Hz, 1H, $H2^B$), 4.90 – 4.55 (m, 12H, $H2^D$, $H1^A$, $H5^D$, $H5^B$, 4 × $CH_2$ Bn), 4.49 (bs, 2H, N$CH_2$Bn), 4.39 (d, *J* = 12.5 Hz, 2H, $H6a^A$, $H6a^C$), 4.28 (d, *J* = 12.2 Hz, 1H, $H6b^A$), 4.19 (d, *J* = 12.5 Hz, 1H, $H6b^C$), 4.73 (t, *J* = 3.3 Hz, 1H, $H3^D$), 4.04 – 3.77 (m, 8H, $H4^B$, $H4^D$, $H3^B$, $H5^A$, $H4^C$, $H4^A$, $H3^A$, $H5^C$), 3.68 – 3.60 (m, 2H, $H3^C$, OCH*H* linker), 3.48 (s, 3H, $CO_2CH_3$), 3.44 (s, 3H, $CO_2CH_3$), 3.41 – 3.17 (m, 5H, $H2^A$, $H2^C$, OCH*H* linker, $CH_2$N linker), 2.78 – 2.46 (m, 12H, 6 × $CH_2$ Lev), 2.17 (s, 3H, $CH_3$ Lev), 2.15 (s, 3H, $CH_3$ Lev), 2.10 (s, 6H, $CH_3$ Lev, $CH_3$ Ac), 1.65 – 1.26 (m, 6H, 3 × $CH_2$ linker). $^{13}$C NMR (101 MHz, $CDCl_3$) δ 206.8 ($CH_3$CO Lev), 206.5 ($CH_3$CO Lev), 206.1 ($CH_3$CO Lev), 172.3 ($CH_2$CO Lev), 172.2 ($CH_2$CO Lev), 171.5 ($CH_2$CO Lev), 170.9 (CO Ac), 169.6 ($CO_2CH_3$), 169.4 ($CO_2CH_3$), 138.1, 138.0, 137.8, 137.5, 137.3, 128.69, 128.65, 128.6, 128.29, 128.27, 128.10, 127.97, 127.95, 127.9, 127.6, 127.5, 127.4, 127.3, 98.2 ($C1^B$), 97.9 ($C1^D$), 97.7 ($C1^A$), 96.9 ($C1^C$), 78.3 ($C3^A$), 78.2 ($C3^C$), 75.9 ($C4^A$), 75.3 ($C3^D$), 74.8 (O$CH_2$Bn), 74.6 (O$CH_2$Bn), 74.4 ($C5^A$), 73.8 ($C3^B$), 73.5 (O$CH_2$Bn), 72.8 (O$CH_2$Bn), 72.1 ($C4^B$), 69.7 ($C5^C$), 69.6 ($C5^B$), 69.5 ($C2^B$), 69.1 ($C4^C$), 69.0 ($C5^D$), 68.3 (O$CH_2$ linker, $C2^D$), 67.8 ($C4^D$), 67.3 ($CH_2$Cbz), 63.4 ($C2^A$), 63.3 ($C2^C$), 62.5 ($C6^A$), 62.2 ($C6^C$), 52.1 ($CO_2CH_3$), 51.9 ($CO_2CH_3$), 50.7 (N$CH_2$Bn), 50.4 (N$CH_2$Bn), 47.2 ($CH_2$N linker), 46.3 ($CH_2$N linker), 38.0 ($CH_2$ Lev), 37.9 ($CH_2$ Lev), 37.8 ($CH_2$ Lev), 29.9 ($CH_3$ Lev), 29.8 (2 × $CH_3$ Lev), 29.1 ($CH_2$ linker), 28.08 ($CH_2$ Lev), 28.05 ($CH_2$ Lev), 27.8 ($CH_2$ linker), 23.4 ($CH_2$ linker), 21.0 ($CH_3$ Ac). HRMS (ESI-MS): m/z calculated for $C_{91}H_{107}N_7NaO_{30}$ $[M+Na]^+$: 1800.6955; found: 1800.7026.

***N*-(Benzyl)-benzyloxycarbonyl-5-aminopentyl *O*-[methyl-2-*O*-acetyl-3-*O*-benzyl-4-*O*-(9-fluorenylmethyloxycarbonyl)-β-D-glucopyranosyluronate]-(1→4)-*O*-(2-azido-3-*O*-benzyl-6-*O*-acetyl-2-deoxy-α-D-glucopyranosyl)-(1→4)-*O*-(methyl-2-*O*-levulinoyl-3-*O*-benzyl-α-L-idopyranosyluronate)-(1→4)-*O*-(2-azido-3-*O*-benzyl-6-*O*-levulinoyl-2-deoxy-α-D-glucopyranosyl)-(1→4)-*O*-(methyl-2-*O*-levulinoyl-3-*O*-benzyl-α-L-idopyranosyluronate)-(1→4)-*O*-2-azido-3-*O*-benzy-6-*O*-acetyl-2-deoxy-α-D-glucopyranoside (18).**

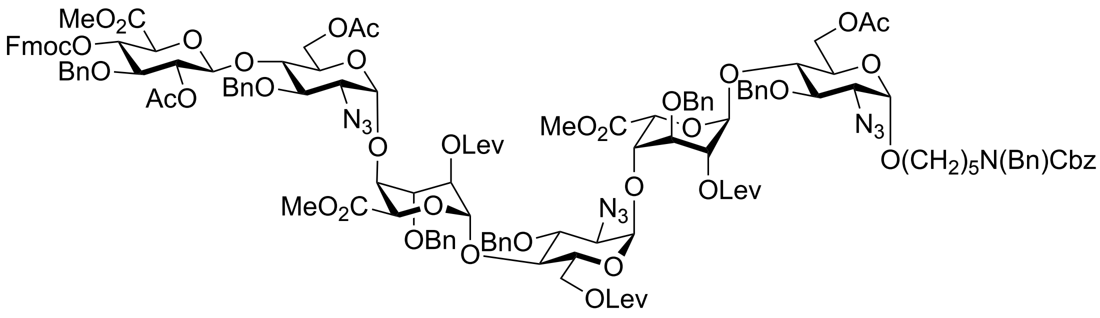

Freshly activated 4 Å molecular sieves was added to a solution of glycosyl donor **15**[3] (400 mg, 0.38 mmol) and glycosyl acceptor **S2** (557 mg, 0.31 mmol) in anhydrous DCM (6.3 mL). After stirring for 30 min at room temperature, the solution was cooled to -40 °C followed by the addition of TfOH (14 μL, 0.16 mmol). The reaction mixture was stirred for 1 h, after which it was quenching with pyridine. The reaction mixture was filtered, and the filtrate was concentrated *in vacuo*. The residue was purified by silica gel column chromatography using a stepwise gradient of petroleum ether and EtOAc (3/1 to 1/1, v/v) as the eluent to give hexasaccharide **18** as a white foam (513 mg, 62%). $R_f$ = 0.54 (petroleum ether/EtOAc, 1/2, v/v). $^1$H NMR (400 MHz, $CDCl_3$) δ 8.26 – 6.64 (m, 48H, CH aromatic), 5.29 (d, *J* = 4.5 Hz, 1H, H1$^B$), 5.27 (d, *J* = 4.9 Hz, 1H, H1$^D$), 5.17 (bd, *J* = 9.5 Hz, 2H, C*H*$_2$Cbz), 5.12 (d, *J* = 11.0 Hz, 1H, C*H*HBn), 5.10 – 5.00 (m, 4H, H2$^F$, H4$^F$, H1$^E$, H1$^C$), 5.00 – 4.52 (m, 17H, H2$^B$, H2$^D$, C*H*HBn, H1$^A$ at 4.82, H5$^B$, 5 × C*H*$_2$ Bn, H5$^D$, H1$^F$ at 4.53), 4.50 (bs, 2H, NC*H*$_2$Bn), 4.47 – 4.12 (m, 9H, H6a$^C$, H6a$^E$, H6a$^A$, H6b$^A$, H6b$^E$, H6b$^C$, C*H*$_2$ Fmoc, C*H* Fmoc), 4.06 (t, *J* = 4.9 Hz, 1H, H4$^B$), 4.03 – 3.64 (m, 15H, H4$^D$, H3$^B$, H3$^D$, H5$^F$, H5$^E$, H5$^C$, H5$^A$, H4$^C$, H4$^E$, H3$^A$, H4$^A$, H3$^F$, H3$^E$, H3$^C$, OCH*H* linker), 3.59 (s, 3H, $CO_2CH_3$), 3.51 (s, 3H, $CO_2CH_3$), 3.46 (s, 3H, $CO_2CH_3$), 3.44 – 3.18 (m, 6H, OCH*H* Linker, C*H*$_2$N linker, H2$^C$, H2$^E$, H2$^A$), 2.82 – 2.42 (m,

12H, 6 × $\mathrm{C}H_2$ Lev), 2.17 (s, 3H, $\mathrm{C}H_3$ Lev), 2.11 (s, 6H, $\mathrm{C}H_3$ Lev, $\mathrm{C}H_3$ Ac), 2.07 (s, 3H, $\mathrm{C}H_3$ Lev), 2.05 (s, 3H, $\mathrm{C}H_3$ Ac), 1.95 (s, 6H, $\mathrm{C}H_3$ Ac), 1.64 – 1.26 (m, 6H, 3 × $\mathrm{C}H_2$ linker). $^{13}$C NMR (101 MHz, $CDCl_3$) δ 206.6 ($CH_3\underline{C}O$ Lev), 206.2 ($CH_3\underline{C}O$ Lev), 206.1 ($CH_3\underline{C}O$ Lev), 172.5 ($CH_2\underline{C}O$ Lev), 172.1 ($CH_2\underline{C}O$ Lev), 172.0 ($CH_2\underline{C}O$ Lev), 170.9 ($\underline{C}O$ Ac), 170.5 ($\underline{C}O$ Ac), 169.8 ($\underline{C}O_2CH_3$), 169.7 ($\underline{C}O_2CH_3$), 169.0 ($\underline{C}O$ Ac), 167.0 ($\underline{C}O_2CH_3$), 154.0 ($\underline{C}O$ Cbz), 143.4, 143.1, 141.43, 141.39, 138.2, 138.04, 137.99, 137.7, 137.63, 137.55, 128.7, 128.59, 128.55, 128.53, 128.4, 128.32, 128.30, 128.11, 128.09, 128.02, 128.00, 127.96, 127.91, 127.89, 127.77, 127.74, 127.68, 127.65, 127.6, 127.3, 125.2, 125.1, 120.2, 100.9 (C1$^{F}$), 98.2 (C1$^{D}$), 98.1 (C1$^{B}$), 97.7 (C1$^{A}$), 97.5 (C1$^{E}$), 97.0 (C1$^{C}$), 79.5 (C3$^{F}$), 78.4, 78.1, 77.9, 77.6, 76.2, 75.7, 75.3 ($O\underline{C}H_2Bn$), 75.2 (C4$^{F}$), 75.0 (C3$^{B}$, C3$^{D}$, $O\underline{C}H_2Bn$), 74.6 ($O\underline{C}H_2Bn$), 74.1 ($O\underline{C}H_2Bn$), 73.7 ($O\underline{C}H_2Bn$), 72.9, 72.8 (C4$^{D}$), 72.4 (C2$^{F}$), 72.2 (C4$^{B}$), 70.6 (C5$^{D}$, $\underline{C}H_2$ Fmoc), 70.5 (C2$^{D}$), 70.0 (C2$^{B}$, C5$^{B}$), 69.7, 69.3, 69.1, 68.3 ($O\underline{C}H_2$ linker), 67.3 ($\underline{C}H_2Cbz$), 63.3 (C2$^{C}$), 63.1 (C2$^{A}$), 62.8 (C2$^{E}$), 62.5 (C6$^{A}$), 62.1 (C6$^{E}$), 61.7 (C6$^{C}$), 52.8 ($CO_2\underline{C}H_3$), 52.3 ($CO_2\underline{C}H_3$), 52.0 ($CO_2\underline{C}H_3$), 50.7 ($N\underline{C}H_2Bn$), 50.4 ($N\underline{C}H_2Bn$), 47.2 ($\underline{C}H_2N$ Linker), 46.7 ($\underline{C}H$ Fmoc), 46.3 ($\underline{C}H_2N$ linker), 38.0 ($\underline{C}H_2$ Lev), 37.8 ($\underline{C}H_2$ Lev), 37.7 ($\underline{C}H_2$ Lev), 29.9 ($\underline{C}H_3$ Lev), 29.8 ($\underline{C}H_3$ Lev), 29.6 ($\underline{C}H_3$ Lev), 29.1 ($\underline{C}H_2$ linker), 28.0 ($\underline{C}H_2$ Lev), 27.8 ($\underline{C}H_2$ Lev), 27.6 ($\underline{C}H_2$ linker), 23.4 ($\underline{C}H_2$ linker), 21.0 ($\underline{C}H_3$ Ac), 21.0 ($\underline{C}H_3$ Ac), 20.7 ($\underline{C}H_3$ Ac). HRMS (ESI-MS): m/z calculated for $C_{137}H_{160}N_{12}O_{44}$ $[M+2NH_4]^{2+}$: 1339.0337; found: 1339.0395.

***N*-(Benzyl)-benzyloxycarbonyl-5-aminopentyl *O*-(3-*O*-benzyl-β-D-glucopyranosyluronate)-(1→4)-*O*-(2-azido-3-*O*-benzyl-2-deoxy-α-D-glucopyranosyl)-(1→4)-*O*-(2-*O*-sulfate-3-*O*-benzyl-α-L-idopyranosyluronate)-(1→4)-*O*-(2-azido-3-*O*-benzyl-6-*O*-sulfate-2-deoxy-α-D-glucopyranosyl)-(1→4)-*O*-(2-*O*-sulfate-3-*O*-benzyl-α-L-idopyranosyluronate)-(1→4)-*O*-2-azido-3-*O*-benzy-2-deoxy-α-D-glucopyranoside, sodium salt (21).**

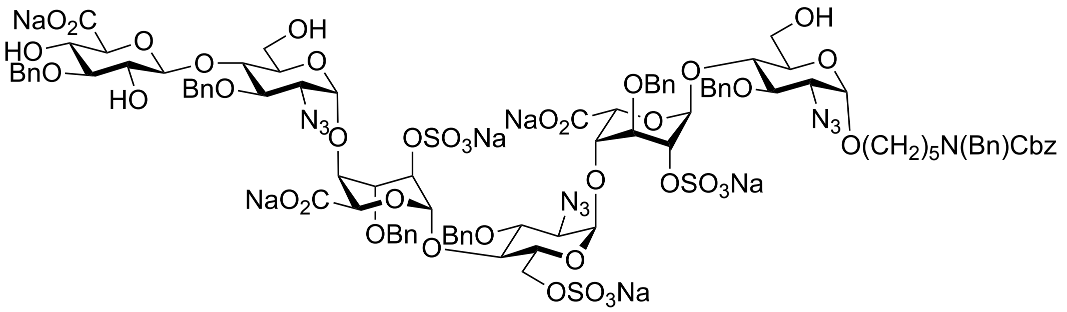

A solution of a hexasaccharide **18** (385 mg, 0.15 mmol) in a mixture of DCM and $Et_3N$ (12.8 mL, 4/1, v/v) was stirred at ambient temperature for 2 h. The reaction mixture was concentrated under reduced pressure, and the residue was dissolved in pyridine (2.0 mL) followed by the addition of acetic anhydride (0.5 mL). Stirring was continued for 3 h until TLC analysis indicated completion of the reaction. The reaction mixture was concentrated under reduced pressure and the residue was dissolved in EtOAc (25 mL), washed with water (25 mL), 1 N HCl solution (25 mL), water (25 mL), saturated bicarbonate (25 mL) and brine (25 mL), dried ($Na_2SO_4$), filtered, and the filtrate was concentrated under reduced pressure, the residue was subjected to the Lev ester removal, *O*-sulfation, saponification, de-*O*-acetylation reaction according to the general procedures to provide compound **21** as a white powder (159 mg, 47% over five steps). $^1H$ NMR (600 MHz, $D_2O$) δ 7.59 – 7.17 (m, 40H, CH aromatic), 5.30 (s, 2H, $H1^B$, $H1^D$), 5.21 – 5.10 (m, 3H, C*H*2Cbz, $H1^C$), 5.08 (d, $J = 10.8$ Hz, 1H, C*H*HBn), 5.02 – 4.84 (m, 6H, C*H*2Bn, C*H*HBn, C*H*HBn, $H1^A$, $H1^E$), 4.84 – 4.66 (m, 6H, $H5^B$, C*H*HBn, $H5^D$, C*H*HBn, C*H*HBn, C*H*HBn), 4.65 – 4.40 (m, 8H, C*H*HBn, $H1^F$ at 4.58, C*H*HBn, $H2^B$, $H2^D$, C*H*HBn, NC*H*2Bn), 4.39 – 4.31 (m, 2H, $H6a^C$, $H6b^C$), 4.28 – 4.13 (m, 4H, $H4^B$, $H3^B$, $H4^D$, $H3^D$), 4.10 – 4.06 (m, 1H, $H5^C$), 4.02 – 3.70 (m, 14H, $H3^E$, $H4^C$, $H4^A$, $H4^E$, $H5^E$, $H3^C$, $H6a^A$, $H6b^A$, $H6a^E$, $H6b^E$, $H3^A$, $H4^F$, $H5^F$, $H5^A$), 3.70 – 3.24 (m, 9H, OCH*H* linker, $H3^F$, $H2^C$, $H2^F$, $H2^A$, $H2^E$, OCH*H* linker, C*H*2N linker), 1.63 – 1.29 (m, 6H, 3 × C*H*2 linker). $^{13}C$ NMR from HSQC (151 MHz, $D_2O$) δ 129.0, 128.7, 129.6, 128.5, 128.4, 102.7 ($C1^F$), 98.3 ($C1^B$, $C1^D$), 96.9 ($C1^A$), 94.4 ($C1^E$), 94.0 ($C1^C$), 83.6 ($C3^F$), 78.7 ($C3^C$, $C3^E$), 77.6 ($C3^A$), 76.8 ($C5^F$), 75.7 ($C4^E$), 75.5 (3 × O$\underline{C}$H2Bn), 74.6 (O$\underline{C}$H2Bn), 74.5 ($C4^A$), 73.3 ($C2^F$), 72.8 ($C4^C$), 72.3 (2 × O$\underline{C}$H2Bn), 71.6 ($C5^A$, $C4^F$), 71.4 ($C2^D$), 71.2 ($C5^E$), 71.0 ($C2^B$), 70.6 ($C4^D$), 70.2 ($C3^B$, $C3^D$), 69.7 ($C5^C$), 69.3 ($C4^B$), 67.9 ($C5^D$), 67.8 (O$\underline{C}$H2 linker), 67.74 ($C5^B$), 67.73 ($\underline{C}$H2Cbz), 66.5 ($C6^C$), 63.2 ($C2^C$), 62.9 ($C2^A$, $C2^E$), 59.6 ($C6^A$, $C6^E$), 50.5 (N$\underline{C}$H2Bn), 47.1 (N$\underline{C}$H2 linker), 27.3 ($\underline{C}$H2 linker), 23.0 ($\underline{C}$H2 linker). HRMS (ESI-MS): m/z calculated for $C_{98}H_{112}N_{10}O_{42}S_3$ $[M-6Na+4H]^{2-}$: 1097.7993; found: 1097.7996.

***N*-(Benzyl)-benzyloxycarbonyl-5-aminopentyl *O*-(3-*O*-benzyl-β-D-glucopyranosyluronate)-(1→4)-*O*-(2-amino-3-*O*-benzyl-2-deoxy-α-D-glucopyranosyl)-(1→4)-*O*-(2-*O*-sulfate-3-*O*-benzyl-α-L-idopyranosyluronate)-(1→4)-*O*-(2-amino-3-*O*-benzyl-6-*O*-sulfate-2-deoxy-α-D-glucopyranosyl)-(1→4)-*O*-(2-*O*-sulfate-3-*O*-benzyl-α-L-idopyranosyluronate)-(1→4)-*O*-2-amino-3-*O*-benzy-2-deoxy-α-D-glucopyranoside, sodium salt (S3).**

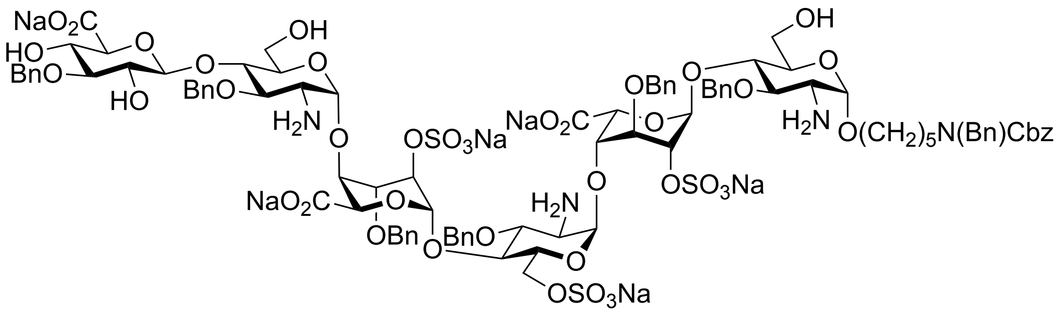

To a solution of compound **21** (154 mg, 66 μmol) in THF (5.1 mL) was added 0.1 M NaOH (19.8 mL). Then, 1.0 M solution of $PMe_3$ in THF (1.6 mL) was added to the solution. The reaction mixture was stirred at room temperature for 1 h until LC-MS indicated completion of the reaction. After adjusting the pH to 8-9 by careful addition of AcOH, the solvents were removed under reduced pressure. The residue was dissolved in water and applied to a Reverse Phase C18 column (1.0 cm × 5 cm), which was eluted with a gradient of $H_2O$ and MeOH (from 90/10 to 40/60, v/v). Fractions containing product were concentrated under reduced pressure and the residue was passed through a column of Dowex® 50 × 8 $Na^+$ resin (0.6 cm × 5 cm) using $H_2O$ as eluent, to give the desired product **S3** as a white powder (120 mg, 81%). $^1H$ NMR (600 MHz, $D_2O$) δ 7.55 – 6.88 (m, 40H, CH aromatic), 5.40 and 5.39 (2s, 2H, $H1^D$, $H1^B$), 5.13 – 4.96 (m, 4H, C*H*HBn, C$H_2$Cbz, $H1^E$), 4.92 (d, $J$ = 3.2 Hz, 1H, $H1^C$), 4.94– 4.50 (m, 15H, $H5^D$, $H5^B$, $H1^A$ at 4.63, 4 × C$H_2$ Bn, C*H*HBn, $H1^F$ at 4.58, $H2^D$, $H2^B$), 4.50 – 4.18 (m, 8H, C$H_2$Bn, NC$H_2$Bn, $H6a^C$, $H6b^C$, $H4^D$, $H4^B$), 4.16 – 4.10 (m, 1H, $H5^E$), 4.09 – 3.66 (m, 14H, $H3^B$, $H3^D$, $H4^E$, $H6a^E$, $H5^C$, $H4^C$, $H4^A$, $H6a^A$, $H6b^A$, $H3^C$, $H6b^E$, $H4^F$, $H5^F$, $H3^E$), 3.61 (t, $J$ =

9.2 Hz, 1H, H3$^F$), 3.63 – 3.23 (m, 4H, H5$^A$, H2$^F$, OCH*H* linker, H3$^A$), 3.23 – 2.87 (m, 3H, OCH*H* linker, C*H*$_2$N linker), 2.78 (d, *J* = 10.7 Hz, 1H, H2$^E$), 2.72 (d, *J* = 10.7 Hz, 1H, H2$^C$), 2.56 – 2.49 (m, 1H, H2$^A$), 1.40 – 0.90 (m, 6H, 3 × C*H*$_2$ linker). $^{13}$C NMR from HSQC (151 MHz, $D_2O$) δ 129.7, 129.1, 128.6, 128.43, 128.40, 128.37, 128.2, 128.0, 127.8, 102.5 (C1$^F$), 98.4 (C1$^A$), 97.4 (C1$^B$, C1$^D$), 97.2 (C1$^E$), 96.4 (C1$^C$), 83.5 (C3$^F$), 81.7 (C3$^E$), 81.2 (C3$^C$), 79.5 (C3$^A$), 76.5 (C5$^F$), 75.8 (C4$^A$), 75.57 (O<u>C</u>H$_2$Bn), 75.56 (3 × O<u>C</u>H$_2$Bn), 74.52 (O<u>C</u>H$_2$Bn), 73.4 (C4$^C$), 73.3 (C2$^F$), 72.8 (C3$^D$), 72.4 (C5$^A$), 72.20 (O<u>C</u>H$_2$Bn), 72.19 (C4$^E$), 72.0 (O<u>C</u>H$_2$Bn), 71.52 (C4$^D$), 71.48 (C5$^C$), 71.4 (C4$^F$), 71.2 (C3$^B$), 70.9 (C2$^B$), 70.8 (C2$^D$), 70.4 (C4$^B$), 70.2 (C5$^E$), 67.7 (C5$^B$), 67.50 (C5$^D$), 67.47 (O<u>C</u>H$_2$ linker), 67.3 (<u>C</u>H$_2$Cbz), 66.7 (C6$^C$), 59.7 (C6$^A$), 59.4 (C6$^E$), 54.7 (C2$^E$), 54.5 (C2$^C$), 54.2 (C2$^A$), 50.2 (N<u>C</u>H$_2$Bn), 47.1 (<u>C</u>H$_2$N linker), 46.2 (<u>C</u>H$_2$N linker), 28.6 (<u>C</u>H$_2$ linker), 27.3 (<u>C</u>H$_2$ linker), 23.0 (<u>C</u>H$_2$ linker). HRMS (ESI-MS): m/z calculated for $C_{98}H_{116}N_4O_{42}S_3$ [M-6Na+4H]$^{2-}$: 1058.8135; found: 1058.8149.

***N*-(Benzyl)-benzyloxycarbonyl-5-aminopentyl *O*-(3-*O*-benzyl-β-D-glucopyranosyluronate)-(1→4)-*O*-(2-sulfamino-3-*O*-benzyl-2-deoxy-α-D-glucopyranosyl)-(1→4)-*O*-(2-*O*-sulfate-3-*O*-benzyl-α-L-idopyranosyluronate)-(1→4)-*O*-(2-sulfamino-3-*O*-benzyl-6-*O*-sulfate-2-deoxy-α-D-glucopyranosyl)-(1→4)-*O*-(2-*O*-sulfate-3-*O*-benzyl-α-L-idopyranosyluronate)-(1→4)-*O*-2-sulfamino-3-*O*-benzy-2-deoxy-α-D-glucopyranoside, sodium salt (22).**

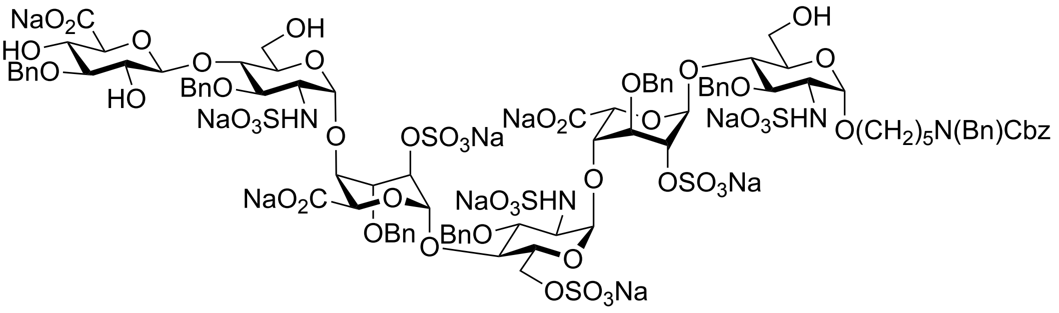

$SO_3$·Py (127 mg, 0.80 mmol) was added to the solution of compound **S3** (120 mg, 53 μmol) in a mixture of MeOH (8.8 mL) and $Et_3N$ (2.7 mL). 1 M NaOH was added to adjust the pH to 11. Three additional portions of $SO_3$·Py (127 mg) was added to the solution after 30 min, 1 h, 2 h followed by adjusting the pH to 11 by careful addition of 1 M NaOH respectively. The progress of the reaction was monitored by TLC (silica gel TLC, EtOAc/Pyridine/$H_2O$/AcOH, 8/5/3/1, v/v/v/v). After stirring for an additional 12 h, the reaction mixture was co-evaporated with water and the residue passed through a short column of Dowex® 50 × 8$Na^+$ resin (1.0 cm × 5 cm) with $H_2O$ as eluent. Fractions containing product were lyophilized, and the residue was dissolved in water and applied to Reverse Phase C18 silica gel column (1.0 cm × 5 cm), which was eluted with a gradient of $H_2O$ and acetonitrile (from 98/2 to 85/15, v/v). Appropriate fractions were lyophilized to give the desired compound **22** as a white powder (84 mg, 62%). $^1$H NMR (600 MHz, $D_2O$) δ 7.58 – 7.14 (m, 40H, CH aromatic), 5.49 (s, 1H, H1$^B$), 5.39 (s, 1H, H1$^D$), 5.32 – 5.27 (m, 2H, H1$^C$, H1$^E$), 5.20 – 5.09 (m, 2H, C*H*2Cbz), 5.00 (bd, 1H, H1$^A$), 4.91 – 4.64 (m, 11H, H5$^B$, H5$^D$, 4 x C*H*2 Bn, C*H*HBn), 4.60 (s, 2H, H2$^B$, H2$^D$), 4.56 – 4.42 (m, 7H, H1$^F$ at 4.54, NC*H*2Bn, C*H*2Bn, C*H*HBn, H6a$^C$), 4.36 – 4.22 (m, 4H, H6b$^C$, H3$^B$, H4$^B$, H3$^D$), 4.18 (s, 1H, H4$^D$), 4.12 – 4.08 (m, 1H, H5$^C$), 3.97 – 3.49 (m, 16H, H4$^C$, H5$^E$, H4$^E$, H4$^A$, H6a$^A$, H6b$^A$, H3$^E$, H6a$^E$, H6b$^E$, H5$^A$, H3$^C$, H4$^F$, H5$^F$, H3$^A$, H3$^F$, OCH*H* linker), 3.44 – 3.30 (m, 4H, OCH*H* linker, H2$^C$, H2$^E$, H2$^F$), 3.30 – 3.20 (m, 3H, C*H*2N linker, H2$^A$), 1.60 – 1.18 (m, 6H, 3 × C*H*2 linker). $^{13}$C NMR from HSQC (151 MHz, $D_2O$) δ 129.3, 129.2, 128.79, 128.75, 128.7, 128.60, 128.3, 102.7 (C1$^F$), 98.5 (C1$^E$, C1$^C$), 97.9 (C1$^B$), 97.4 (C1$^D$), 97.1 (C1$^A$), 83.6 (C3$^F$), 78.0 (C3$^E$), 77.7 (C3$^C$), 76.5 (C5$^F$, C3$^A$), 75.5 (C4$^A$, C4$^E$), 75.4 (O<u>C</u>H2Bn), 75.14 (C3$^D$), 75.06 (O<u>C</u>H2Bn, C4$^D$), 75.0 (C3$^B$, C4$^B$), 74.5 (O<u>C</u>H2Bn), 73.3, 72.62 (C4$^C$), 72.56 (O<u>C</u>H2Bn), 72.3 (C2$^B$, C2$^D$), 71.5 (C5$^A$, C4$^F$), 71.2 (C5$^E$), 69.8 (C5$^C$), 68.6 (C5$^D$), 68.3 (C5$^B$), 68.1 (O<u>C</u>H2 linker), 67.7 (<u>C</u>H2Cbz), 66.9 (C6$^C$), 60.2 (C6$^A$ or C6$^E$), 59.9 (C6$^A$ or C6$^E$), 58.1 (C2$^C$, C2$^E$), 57.4 (C2$^A$), 50.6 (N<u>C</u>H2Bn), 47.2 (N<u>C</u>H2 linker), 27.3 (<u>C</u>H2 linker), 23.0 (<u>C</u>H2 linker). HRMS (ESI-MS): m/z calculated for $C_{98}H_{116}N_4O_{51}S_6$ $[M-9Na+7H]^{2-}$: 1178.7488; found: 1178.7486.

**5-Aminopentyl *O*-(β-D-glucopyranosyluronate)-(1→4)-*O*-(2-sulfamino-2-deoxy-α-D-glucopyranosyl)-(1→4)-*O*-(2-*O*-sulfate-α-L-idopyranosyluronate)-(1→4)-*O*-(2-sulfamino-6-*O*-sulfate-2-deoxy-α-D-glucopyranosyl)-(1→4)-*O*-(2-*O*-sulfate-α-L-**

**idopyranosyluronate)-(1→4)-*O*-2-sulfamino-2-deoxy-α-D-glucopyranoside, sodium salt (1).**

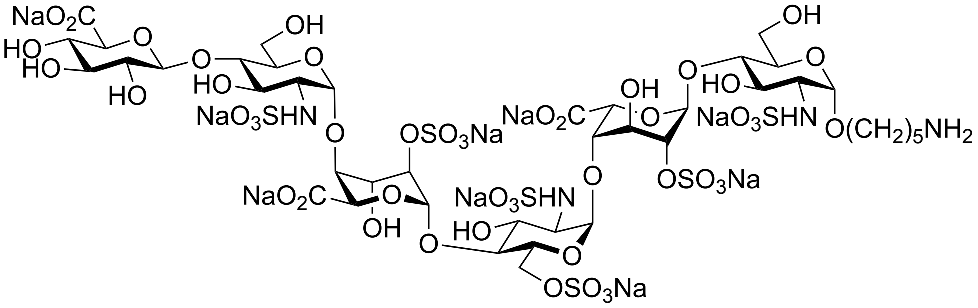

Palladium hydroxide on carbon (Degussa type, 20%, 1.5 times the weight of starting material) was added to a solution of the compound **22** (84 mg, 33 μmol) in *tert*-butanol and $H_2O$ (1/1, v/v, 10 mL). The mixture was placed under an atmosphere of hydrogen until completion of the reaction as indicated by ESI-LC-MS. The mixture was filtered through a PTFE syringe filter (Acrodisc®, 0.2 μm), and the residue was washed with *tert*-butanol and $H_2O$ mixture (1/1, v/v, 3 × 2 mL). The filtrate was lyophilized to give the final product. Biogel P2 (1.5 cm × 50 cm) was used for purification with 0.1 M ammonium bicarbonate as eluent, the fractions containing compound were lyophilized and passed through a short column of Dowex® 50 × 8$Na^+$ resin (1.0 cm × 5 cm) with $H_2O$ as eluent. Appropriate fractions were lyophilized to give the desired compound **1** as a white powder (50 mg, 85%). $^1$H NMR (600 MHz, $D_2O$) δ 5.45 (d, $J$ = 3.6 Hz, 1H, H1$^C$), 5.41 (d, $J$ = 3.5 Hz, 1H, H1$^E$), 5.29 (d, $J$ = 3.6 Hz, 1H, H1$^B$), 5.22 (d, $J$ = 3.0 Hz, 1H, H1$^D$), 5.14 (d, $J$ = 3.5 Hz, 1H, H1$^A$), 4.82 (1H, H5$^D$), 4.70 (d, $J$ = 2.9 Hz, 1H, H5$^B$), 4.52 (d, $J$ = 7.9 Hz, 1H, H1$^F$), 4.40 (dd, $J$ = 11.5, 2.9 Hz, 1H, H6a$^C$), 4.35 – 4.30 (m, 2H, H2$^D$, H2$^B$), 4.28 (dd, $J$ = 11.5, 2.2 Hz, 1H, H6b$^C$), 4.22 (dd, $J$ = 5.8, 3.7 Hz, 1H, H3$^D$), 4.19 (dd, $J$ = 6.6, 3.9 Hz, 1H, H3$^B$), 4.12 (t, $J$ = 3.6 Hz, 1H, H4$^B$), 4.09 (t, $J$ = 3.3 Hz, 1H, H4$^D$), 4.03 – 4.00 (m, 1H, H5$^C$), 3.97 – 3.69 (m, 13H, H6a$^A$, H6b$^A$, H6a$^E$, H6b$^E$, H5$^E$, H5$^A$, H3$^A$, H4$^C$, H5$^F$, H3$^E$, H4$^A$, H4$^E$, OCH*H* linker), 3.66 – 3.61 (m, 1H, H3$^C$), 3.58 – 3.49 (m, 3H, OCH*H* linker, H4$^F$, H3$^F$), 3.44 – 3.38 (m, 1H, H2$^F$), 3.32 – 3.22 (m, 3H, H2$^C$, H2$^E$, H2$^A$), 3.05 (t, $J$ = 7.3 Hz, 2H, C$H_2$N linker), 1.77 – 1.47 (m, 6H, 3 × C$H_2$ linker). $^{13}$C NMR from HSQC (151 MHz, $D_2O$) δ 102.2 (C1$^F$), 99.2 (C1$^D$), 97.8 (C1$^B$), 96.9 (C1$^A$), 96.5 (C1$^E$), 96.1 (C1$^C$), 77.5 (C4$^E$, C4$^A$), 76.7 (C2$^B$), 75.9 (C2$^D$), 75.9 (C4$^B$), 75.8 (C4$^D$), 75.8 (C4$^C$, C5$^F$), 75.1 (C3$^F$), 72.8 (C2$^F$), 71.8 (C4$^F$), 70.6 (C5$^E$), 70.4 (C5$^A$), 69.8 (C5$^B$), 69.6 (C3$^C$), 69.5 (C3$^E$), 69.4 (C3$^B$), 69.3 (C5$^C$), 69.2 (C5$^D$), 69.1 (C3$^D$), 68.5 (C3$^A$), 67.7 (O<u>C</u>H$_2$ linker), 66.4 (C6$^C$), 59.6 (C6$^A$, C6$^E$), 57.9 (C2$^C$, C2$^E$, C2$^A$), 39.5

(N<u>C</u>$H_2$ linker), 28.0 (<u>C</u>$H_2$ linker), 26.4 (<u>C</u>$H_2$ linker), 22.6 (<u>C</u>$H_2$ linker). HRMS (ESI-MS): m/z calculated for $C_{41}H_{68}N_4O_{49}S_6$ [M-9Na+7H]$^{2-}$: 796.0644; found: 796.0635.

$^1$H NMR (600 MHz, $D_2O$)

| | **H1** | **H2** | **H3** | **H4** | **H5** | **H6** |
|---|---|---|---|---|---|---|
| **A** | 5.14 (d, *J* = 3.5 Hz) | 3.25 | 3.84 | 3.73 | 3.87 | 3.88 |
| **B** | 5.29 (d, *J* = 3.6 Hz) | 4.32 | 4.20 | 4.13 | 4.71 | – |
| **C** | 5.45 (d, *J* = 3.6 Hz) | 3.29 | 3.63 | 3.78 | 4.02 | 4.41, 4.28 |
| **D** | 5.22 (d, *J* = 3.0 Hz) | 4.34 | 4.23 | 4.09 | 4.82 | – |
| **E** | 5.41 (d, *J* = 3.5 Hz) | 3.26 | 3.73 | 3.73 | 3.95 | 3.88 |
| **F** | 4.52 (d, *J* = 7.9 Hz) | 3.41 | 3.53 | 3.53 | 3.77 | – |

$^{13}$C NMR from HSQC (151 MHz, $D_2O$)

| | **C1** | **C2** | **C3** | **C4** | **C5** | **C6** |
|---|---|---|---|---|---|---|
| **A** | 96.9 | 57.9 | 68.5 | 77.5 | 70.4 | 59.6 |
| **B** | 97.8 | 76.7 | 69.4 | 75.9 | 69.8 | – |
| **C** | 96.1 | 57.9 | 69.6 | 75.8 | 69.3 | 66.4 |
| **D** | 99.2 | 75.9 | 69.1 | 75.8 | 69.2 | – |
| **E** | 96.5 | 57.9 | 69.5 | 77.5 | 70.6 | 59.6 |
| **F** | 102.2 | 72.8 | 75.1 | 71.8 | 75.8 | – |

**5-(Pent-4-ynamido) pentyl *O*-(β-D-glucopyranosyluronate)-(1 → 4)-*O*-(2-sulfamino-2-deoxy-α-D-glucopyranosyl)-(1 → 4)-*O*-(2-*O*-sulfate-α-L-idopyranosyluronate)-(1 → 4)-*O*-(2-sulfamino-6-*O*-sulfate-2-deoxy-α-D-glucopyranosyl)-(1 → 4)-*O*-(2-*O*-sulfate-α-L-idopyranosyluronate)-(1 → 4)-*O*-2-sulfamino-2-deoxy-α-D-glucopyranoside, sodium salt (2).**

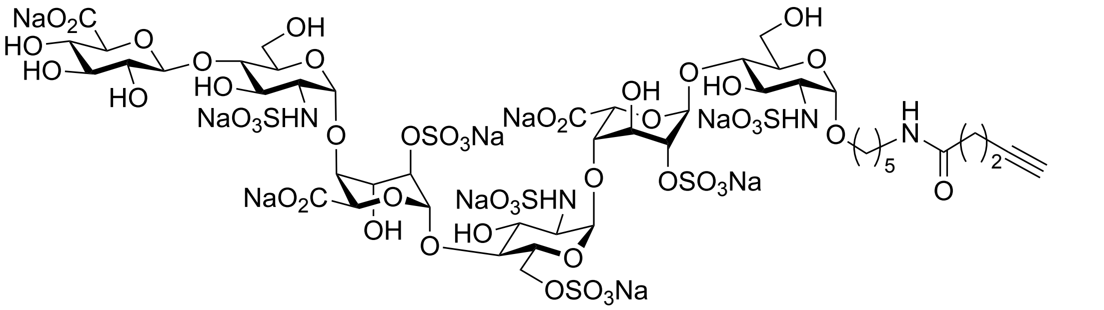

Compound **1** (13.1 mg, 7.3 μmol) was subjected to installation of alkyne moiety according to the general procedure to give the title compound **2** as a white powder (12.1 mg, 88%). $^{1}$H NMR (600 MHz, $D_2O$) δ 5.40 – 5.37 (m, 2H, H1$^{C}$, H1$^{E}$), 5.19 (d, *J* = 1.9 Hz, 1H, H1$^{D}$), 5.15 (d, *J* = 3.2 Hz, 1H, H1$^{B}$), 5.11 (d, *J* = 3.8 Hz, 1H, H1$^{A}$), 4.80 (1H, H5$^{D}$), 4.74 (d, *J* = 2.4 Hz, 1H, H5$^{B}$), 4.48 (d, *J* = 7.9 Hz, 1H, H1$^{F}$), 4.40 – 4.35 (m, 1H, H6a$^{C}$), 4.31 – 4.26 (m, 2H, H2$^{D}$, H2$^{B}$), 4.26 – 4.22 (m, 1H, H6b$^{C}$), 4.20 – 4.15 (m, 2H, H3$^{D}$, H3$^{B}$), 4.08 (t, *J* = 3.2 Hz, 1H, H4$^{B}$), 4.05 (t, *J* = 3.0 Hz, 1H, H4$^{D}$), 4.00 – 3.96 (m, 1H, H5$^{C}$), 3.94 – 3.81 (m, 5H, H5$^{E}$, H6a$^{A}$, H6b$^{A}$, H6a$^{E}$, H6b$^{E}$), 3.79 – 3.58 (m, 9H, H5$^{A}$, H4$^{C}$, H5$^{F}$, H4$^{E}$, H4$^{A}$, H3$^{E}$, H3$^{A}$, H3$^{C}$, OCH*H* linker), 3.53 – 3.46 (m, 3H, OCH*H* linker, H4$^{F}$, H3$^{F}$), 3.40 – 3.34 (m, 1H, H2$^{F}$), 3.27 – 3.16 (m, 5H, H2$^{C}$, H2$^{E}$, H2$^{A}$, C$H_2$N linker), 2.51 – 2.45 (m, 2H, C$H_2$C≡CH), 2.44 – 2.39 (m, 2H, C$H_2$CH$_2$C≡CH), 2.36 (t, *J* = 2.5 Hz, 1H, CH$_2$C≡C*H*), 1.68 – 1.34 (m, 6H, 3 × C$H_2$ linker). $^{13}$C NMR from HSQC (151 MHz, $D_2O$) δ 102.0 (C1$^{F}$), 99.2 (C1$^{B}$, C1$^{D}$), 96.9 (C1$^{A}$), 96.4 (C1$^{C}$, C1$^{E}$), 77.5 (C4$^{E}$, C4$^{A}$), 76.2 (C2$^{B}$), 76.0 (C2$^{D}$), 75.84 (C4$^{B}$, C4$^{D}$), 75.79 (C4$^{C}$, C5$^{F}$), 75.1 (C3$^{F}$), 72.8 (C2$^{F}$), 71.7 (C4$^{F}$), 70.71 (C5$^{A}$), 70.70 (CH$_2$C≡$\underline{C}$H), 70.6 (C5$^{E}$), 69.6 (C3$^{E}$, C3$^{A}$, C3$^{C}$), 69.5 (C5$^{B}$), 69.21 (C3$^{D}$, C3$^{B}$), 69.15 (C5$^{D}$), 69.13 (C5$^{C}$), 68.3 (O$\underline{C}$H$_2$ linker), 66.4 (C6$^{C}$), 59.6 (C6$^{A}$, C6$^{E}$), 57.8 (C2$^{C}$, C2$^{E}$, C2$^{A}$), 39.2 (N$\underline{C}$H$_2$ linker), 34.5 ($\underline{C}$H$_2$CH$_2$C≡CH), 28.0 (2 × $\underline{C}$H$_2$ linker), 22.8 ($\underline{C}$H$_2$ linker), 14.7 ($\underline{C}$H$_2$C≡CH). HRMS (ESI-MS): m/z calculated for $C_{46}H_{72}N_4O_{50}S_6$ [M-9Na+7H]$^{2-}$: 836.0775; found: 836.0782.

$^{1}H$ NMR (600 MHz, $D_2O$)

| | H1 | H2 | H3 | H4 | H5 | H6 |
|---|---|---|---|---|---|---|
| **A** | 5.11 (d, *J* = 3.8 Hz) | 3.23 | 3.65 | 3.69 | 3.76 | 3.85 |
| **B** | 5.15 (d, *J* = 3.2 Hz) | 4.28 | 4.16 | 4.07 | 4.74 (d, *J* = 2.4 Hz) | – |
| **C** | 5.39 | 3.26 | 3.63 | 3.75 | 3.99 | 4.38, 4.24 |
| **D** | 5.19 (d, *J* = 1.9 Hz) | 4.30 | 4.18 | 4.05 | 4.80 | – |
| **E** | 5.38 | 3.22 | 3.69 | 3.69 | 3.91 | 3.85 |
| **F** | 4.48 (d, *J* = 7.9 Hz) | 3.36 | 3.50 | 3.50 | 3.74 | – |

$^{13}C$ NMR from HSQC (151 MHz, $D_2O$)

| | C1 | C2 | C3 | C4 | C5 | C6 |
|---|---|---|---|---|---|---|
| **A** | 96.9 | 57.8 | 69.6 | 77.5 | 70.7 | 59.6 |
| **B** | 99.2 | 76.2 | 69.2 | 75.8 | 69.5 | – |
| **C** | 96.4 | 57.8 | 69.6 | 75.8 | 69.1 | 66.4 |
| **D** | 99.2 | 76.0 | 69.2 | 75.8 | 69.2 | – |
| **E** | 96.4 | 57.8 | 69.6 | 77.5 | 70.6 | 59.6 |
| **F** | 102.0 | 72.8 | 75.1 | 71.7 | 75.8 | – |

**5-Aminopentyl *O*-(2-acetamido-6-azido-2-deoxy-α-D-glucopyranosyl)-(1→4)-*O*-(β-D-glucopyranosyluronate)-(1→4)-*O*-(2-sulfamino-2-deoxy-α-D-glucopyranosyl)-(1→4)-*O*-(2-*O*-sulfate-α-L-idopyranosyluronate)-(1→4)-*O*-(2-sulfamino-6-*O*-sulfate-2-deoxy-α-D-glucopyranosyl)-(1→4)-*O*-(2-*O*-sulfate-α-L-idopyranosyluronate)-(1→4)-*O*-2-sulfamino-2-deoxy-α-D-glucopyranoside, sodium salt (3).**

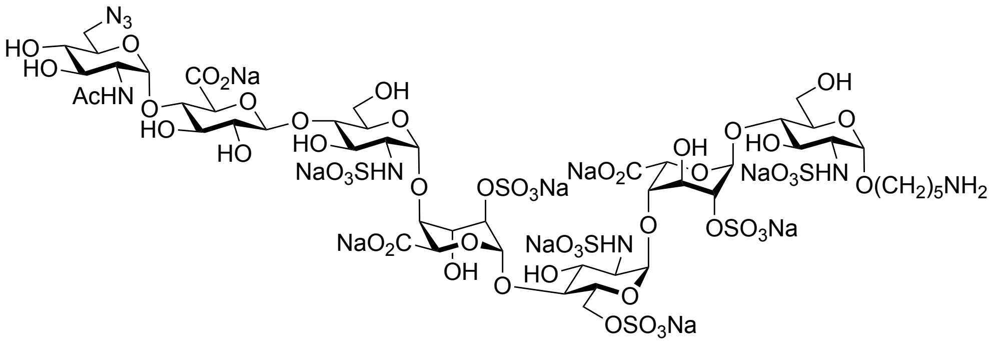

Compound **1** (4.1 mg, 2.3 μmol) was subjected to installation of unnatural α (1→4) 6-azido-GlcNAc according to the general procedure to give the title compound **3** as a white powder (3.9 mg, 84%). $^{1}$H NMR (600 MHz, $D_2O$) δ 5.47 (d, $J$ = 3.5 Hz, 1H, H1$^{C}$), 5.44 (d, $J$ = 3.7 Hz, 1H, H1$^{G}$), 5.42 (d, $J$ = 3.5 Hz, 1H, H1$^{E}$), 5.33 (d, $J$ = 3.4 Hz, 1H, H1$^{B}$), 5.21 (bs, 1H, H1$^{D}$), 5.14 (d, $J$ = 3.4 Hz, 1H, H1$^{A}$), 4.80 (1H, H5$^{D}$), 4.67 (d, $J$ = 2.9 Hz, 1H, H5$^{B}$), 4.50 (d, $J$ = 7.9 Hz, 1H, H1$^{F}$), 4.42 – 4.38 (m, 1H, H6a$^{C}$), 4.34 – 4.30 (m, 2H, H2$^{D}$, H2$^{B}$), 4.29 – 4.26 (m, 1H, H6b$^{C}$), 4.23 – 4.17 (m, 2H, H3$^{D}$, H3$^{B}$ ), 4.15 – 4.12 (m, 1H, H4$^{B}$), 4.10 – 4.07 (m, 1H, H4$^{D}$), 4.03 – 3.99 (m, 1H, H5$^{C}$), 3.96 – 3.64 (m, 20H, H5$^{E}$,H2$^{G}$, H6a$^{A}$, H6b$^{A}$, H6a$^{E}$, H6b$^{E}$, H3$^{A}$, H5$^{G}$, H5$^{A}$, H5$^{F}$, H4$^{C}$, H4$^{F}$, H4$^{A}$, H4$^{E}$, H3$^{F}$, H3$^{G}$, H3$^{E}$, OCH*H* linker, H6a$^{G}$, H6b$^{G}$), 3.63 – 3.59 (m, 1H, H3$^{C}$), 3.58 – 3.54 (m, 1H, OCH*H* linker), 3.50 (t, $J$ = 9.5 Hz, 1H, H4$^{G}$), 3.41 – 3.38 (m, 1H, H2$^{F}$), 3.29 (dd, $J$ = 10.4, 3.5 Hz, 1H, H2$^{C}$), 3.25 (dd, $J$ = 10.1, 3.4 Hz, 2H, H2$^{E}$, H2$^{A}$), 3.05 (t, $J$ = 7.3 Hz, 2H, C$H_2$N linker), 2.06 (s, 3H, NHCOC$H_3$), 1.79 – 1.50 (m, 6H, 3 × C$H_2$ linker). $^{13}$C NMR from HSQC (151 MHz, $D_2O$) δ 102.2 (C1$^{F}$), 99.3 (C1$^{D}$), 97.3 (C1$^{B}$), 97.0 (C1$^{A}$), 96.9 (C1$^{E}$, C1$^{G}$), 96.0 (C1$^{C}$), 77.1 (C2$^{B}$, C4$^{E}$, C4$^{A}$, C3$^{F}$), 76.1 (C2$^{D}$, C4$^{B}$, C4$^{D}$, C5$^{F}$, C4$^{C}$, C4$^{F}$), 73.4 (C2$^{F}$), 70.5 (C5$^{E}$, C5$^{G}$, C5$^{A}$, C3$^{G}$, C4$^{G}$), 70.2 (C5$^{B}$), 69.8 (C3$^{B}$, C3$^{E}$, C3$^{C}$), 69.4 (C5$^{D}$, C3$^{D}$, C5$^{C}$), 68.2 (C3$^{A}$), 67.7 (O$\underline{C}H_2$ linker), 66.5 (C6$^{C}$), 59.7 (C6$^{A}$, C6$^{E}$), 57.9 (C2$^{C}$, C2$^{E}$, C2$^{A}$), 53.6 (C2$^{G}$), 50.5 (C6$^{G}$), 39.6 (N$\underline{C}H_2$ linker), 28.2 ($\underline{C}H_2$ linker), 26.5 ($\underline{C}H_2$ linker), 22.7 ($\underline{C}H_2$ linker), 21.9 (NHCO$\underline{C}H_3$). HRMS (ESI-MS): m/z calculated for $C_{49}H_{80}N_8O_{53}S_6$ [M-9Na+7H]$^{2-}$: 910.1073; found: 910.1073.

$^{1}$H NMR (600 MHz, $D_2O$)

| | H1 | H2 | H3 | H4 | H5 | H6 |
|---|---|---|---|---|---|---|
| **A** | 5.14<br>(d, $J$ = 3.4 Hz) | 3.25<br>(dd, $J$ = 10.1, 3.4 Hz) | 3.91 | 3.73 | 3.91 | 3.88 |

| | | | | | | |
|---|---|---|---|---|---|---|
| **B** | 5.33 (d, *J* = 3.4 Hz) | 4.32 | 4.18 | 4.13 | 4.67 (d, *J* = 2.9 Hz) | – |
| **C** | 5.47 (d, *J* = 3.5 Hz) | 3.29 (dd, *J* = 10.4, 3.5 Hz) | 3.62 | 3.78 | 4.01 | 4.40, 4.27 |
| **D** | 5.21 | 4.32 | 4.21 | 4.09 | 4.80 | – |
| **E** | 5.42 (d, *J* = 3.5 Hz) | 3.25 (dd, *J* = 10.1, 3.4 Hz) | 3.73 | 3.70 | 3.94 | 3.88 |
| **F** | 4.50 (d, *J* = 7.9 Hz) | 3.37 | 3.70 | 3.77 | 3.81 | – |
| **G** | 5.44 (d, *J* = 3.7 Hz) | 3.91 | 3.73 | 3.50 | 3.90 | 3.66 |

$^{13}$C NMR from HSQC (151 MHz, $D_2O$)

| | **C1** | **C2** | **C3** | **C4** | **C5** | **C6** |
|---|---|---|---|---|---|---|
| **A** | 97.0 | 57.9 | 68.2 | 77.1 | 70.5 | 59.7 |
| **B** | 97.3 | 77.1 | 69.8 | 76.1 | 70.2 | – |
| **C** | 96.0 | 57.9 | 69.8 | 76.1 | 69.4 | 66.5 |
| **D** | 99.3 | 76.1 | 69.4 | 76.1 | 69.4 | – |
| **E** | 96.9 | 57.9 | 69.8 | 77.1 | 70.5 | 59.7 |
| **F** | 102.2 | 73.4 | 77.1 | 76.1 | 76.1 | – |
| **G** | 96.9 | 53.6 | 70.5 | 70.5 | 70.5 | 50.5 |

**5-Aminopentyl *O*-(2-acetamido-2-deoxy-α-D-glucopyranosyl)-(1→4)-*O*-(β-D-glucopyranosyluronate)-(1→4)-*O*-(2-sulfamino-2-deoxy-α-D-glucopyranosyl)-(1→4)-*O*-(2-*O*-sulfate-α-L-idopyranosyluronate)-(1→4)-*O*-(2-sulfamino-6-*O*-sulfate-2-deoxy-α-D-**

**glucopyranosyl)-(1→4)-*O*-(2-*O*-sulfate-α-L-idopyranosyluronate)-(1→4)-*O*-2-sulfamino-2-deoxy-α-D-glucopyranoside, sodium salt (S4).**

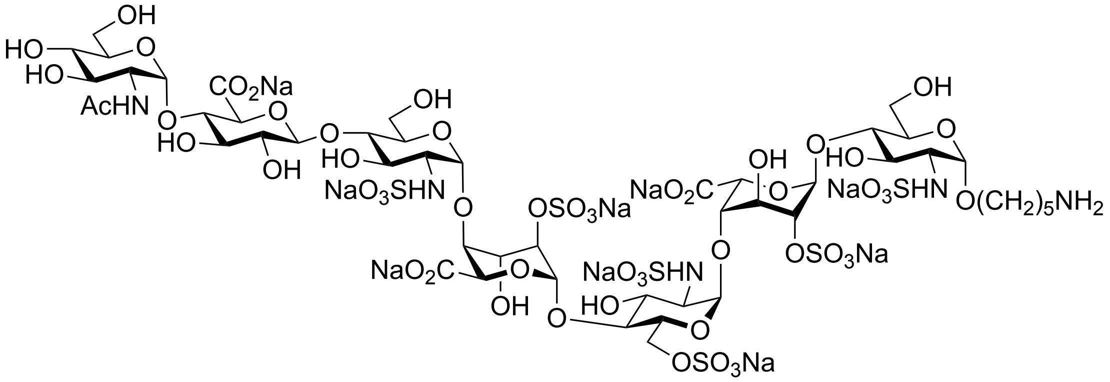

Compound **1** (24.0 mg, 13.4 μmol) was subjected to installation of α (1→4)-GlcNAc according to the general procedure to give the title compound **S4** as a white powder (24.0 mg, 90%). $^{1}$H NMR (600 MHz, $D_2O$) δ 5.41 (d, *J* = 3.5 Hz, 1H, $H1^{C}$), 5.40 (d, *J* = 3.7 Hz, 1H, $H1^{G}$), 5.38 (d, *J* = 3.5 Hz, 1H, $H1^{E}$), 5.26 (d, *J* = 3.4 Hz, 1H, $H1^{B}$), 5.18 (bs, 1H, $H1^{D}$), 5.11 (d, *J* = 3.4 Hz, 1H, $H1^{A}$), 4.78 (1H, $H5^{D}$), 4.67 (d, *J* = 2.7 Hz, 1H, $H5^{B}$), 4.47 (d, *J* = 7.9 Hz, 1H, $H1^{F}$), 4.39 – 4.35 (m, 1H, $H6a^{C}$), 4.31 – 4.27 (m, 2H, $H2^{D}$, $H2^{B}$), 4.27 – 4.22 (m, 1H, $H6b^{C}$), 4.20 – 4.15 (m, 2H, $H3^{D}$, $H3^{B}$ ), 4.10 – 4.07 (m, 1H, $H4^{B}$), 4.07 – 4.04 (m, 1H, $H4^{D}$), 4.00 – 3.96 (m, 1H, $H5^{C}$), 3.93 – 3.89 (m, 1H, $H5^{E}$), 3.88 – 3.63 (m, 189H, $H2^{G}$, $H6a^{A}$, $H6b^{A}$, $H6a^{E}$, $H6b^{E}$, $H6a^{G}$, $H6b^{G}$, $H3^{A}$, $H5^{F}$, $H4^{A}$, $H5^{A}$, $H4^{C}$, $H3^{G}$, $H5^{G}$, $H4^{F}$, $H3^{F}$, $H3^{E}$, $H4^{E}$, OCH*H* linker), 3.60 (t, *J* = 9.6 Hz, 1H, $H3^{C}$), 3.54 – 3.49 (m, 1H, OCH*H* linker), 3.45 (t, *J* = 9.6 Hz, 1H, $H4^{G}$), 3.36 (t, *J* = 8.9 Hz, 1H, $H2^{F}$), 3.25 (dd, *J* = 10.5, 3.3 Hz, 1H, $H2^{C}$), 3.22 (dd, *J* = 10.5, 3.3 Hz, 2H, $H2^{E}$, $H2^{A}$), 3.01 (t, *J* = 7.3 Hz, 2H, $CH_2$N linker), 2.03 (s, 3H, NHCOC$H_3$), 1.76 – 1.43 (m, 6H, 3 × $CH_2$ linker). $^{13}$C NMR from HSQC (151 MHz, $D_2O$) δ 102.1 ($C1^{F}$), 99.2 ($C1^{D}$), 97.8 ($C1^{B}$), 96.9 ($C1^{A}$), 96.6 ($C1^{E}$, $C1^{G}$), 96.1 ($C1^{C}$), 77.5 ($C3^{F}$), 76.9 ($C2^{B}$, $C4^{A}$), 76.1 ($C5^{F}$, $C4^{F}$, $C4^{E}$, $C4^{C}$), 75.8 ($C2^{D}$, $C4^{B}$, $C4^{D}$), 73.2 ($C2^{F}$), 71.6 ($C5^{G}$, $C3^{G}$), 70.5 ($C5^{E}$, $C5^{A}$), 69.8 ($C5^{B}$, $C4^{G}$), 69.5 ($C3^{E}$, $C3^{C}$), 69.2 ($C5^{D}$, $C3^{D}$, $C3^{B}$, $C5^{C}$), 68.5 ($C3^{A}$), 67.7 (O$\underline{C}H_2$ linker), 66.4 ($C6^{C}$), 60.0 ($C6^{G}$), 59.6 ($C6^{A}$, $C6^{E}$), 57.8 ($C2^{C}$, $C2^{E}$, $C2^{A}$), 53.6 ($C2^{G}$), 39.5 (N$\underline{C}H_2$ linker), 28.0 ($\underline{C}H_2$ linker), 26.4 ($\underline{C}H_2$ linker), 22.6 ($\underline{C}H_2$ linker), 21.7 (NHCO$\underline{C}H_3$). HRMS (ESI-MS): m/z calculated for $C_{49}H_{81}N_5O_{54}S_6$ $[M-9Na+7H]^{2-}$: 897.6041; found: 897.6010.

$^{1}$H NMR (600 MHz, $D_2O$)

| | H1 | H2 | H3 | H4 | H5 | H6 |
|---|---|---|---|---|---|---|
| **A** | 5.11 (d, *J* = 3.4 Hz) | 3.22 (dd, *J* = 10.5, 3.3 Hz) | 3.80 | 3.69 | 3.84 | 3.85 |
| **B** | 5.26 (d, *J* = 3.4 Hz) | 4.29 | 4.16 | 4.09 | 4.67 (d, *J* = 2.7 Hz) | – |
| **C** | 5.41 (d, *J* = 3.5 Hz) | 3.25 (dd, *J* = 10.5, 3.3 Hz) | 3.61 | 3.76 | 3.99 | 4.38, 4.25 |
| **D** | 5.18 | 4.30 | 4.19 | 4.05 | 4.79 | – |
| **E** | 5.38 (d, *J* = 3.5 Hz) | 3.22 (dd, *J* = 10.5, 3.3 Hz) | 3.69 | 3.78 | 3.92 | 3.85 |
| **F** | 4.47 (d, *J* = 7.9 Hz) | 3.37 | 3.68 | 3.74 | 3.78 | – |
| **G** | 5.40 (d, *J* = 3.7 Hz) | 3.87 | 3.71 | 3.45 | 3.77 | 3.79 |

$^{13}$C NMR from HSQC (151 MHz, $D_2O$)

| | C1 | C2 | C3 | C4 | C5 | C6 |
|---|---|---|---|---|---|---|
| **A** | 96.9 | 57.8 | 68.5 | 76.9 | 70.5 | 59.6 |
| **B** | 97.8 | 76.9 | 69.2 | 75.8 | 69.8 | – |
| **C** | 96.1 | 57.8 | 69.5 | 76.1 | 69.2 | 66.4 |
| **D** | 99.2 | 75.8 | 69.2 | 75.8 | 69.2 | – |
| **E** | 96.6 | 57.8 | 69.5 | 76.1 | 70.5 | 59.6 |
| **F** | 102.1 | 73.2 | 77.5 | 76.1 | 76.1 | – |
| **G** | 96.6 | 53.6 | 71.6 | 69.8 | 71.6 | 60.0 |

**5-Aminopentyl *O*-(β-D-glucopyranosyluronate)-(1→4)-*O*-(2-acetamido-2-deoxy-α-D-glucopyranosyl)-(1→4)-*O*-(β-D-glucopyranosyluronate)-(1→4)-*O*-(2-sulfamino-2-deoxy-α-D-glucopyranosyl)-(1→4)-*O*-(2-*O*-sulfate-α-L-idopyranosyluronate)-(1→4)-*O*-(2-sulfamino-6-*O*-sulfate-2-deoxy-α-D-glucopyranosyl)-(1→4)-*O*-(2-*O*-sulfate-α-L-idopyranosyluronate)-(1→4)-*O*-2-sulfamino-2-deoxy-α-D-glucopyranoside, sodium salt (23).**

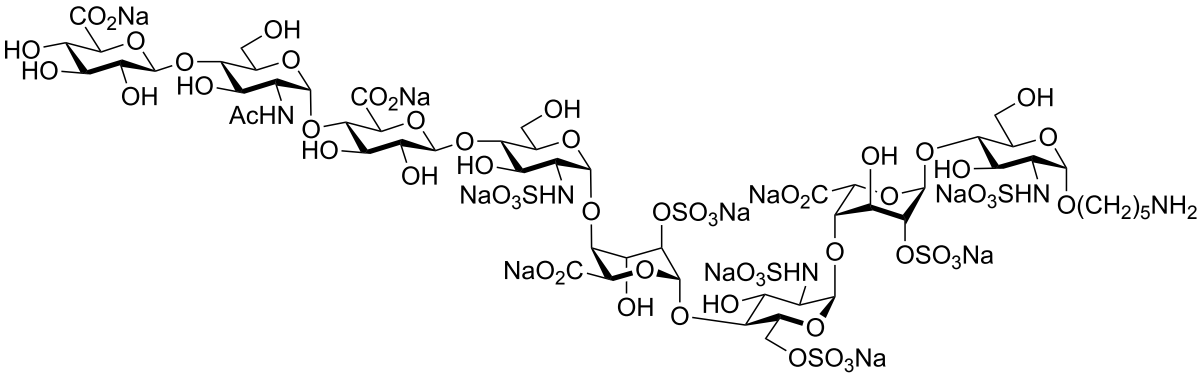

Compound **S4** (23.6 mg, 11.8 μmol) was subjected to installation of β (1→4)-GlcA according to the general procedure to give the title compound **23** as a white powder (23.7 mg, 91%). $^{1}$H NMR (600 MHz, $D_2O$) δ 5.41 (d, *J* = 3.4 Hz, 1H, H1$^{C}$), 5.39 – 5.36 (m, 2H, H1$^{G}$, H1$^{E}$), 5.26 (d, *J* = 3.4 Hz, 1H, H1$^{B}$), 5.18 (d, *J* = 2.5 Hz, 1H, H1$^{D}$), 5.11 (d, *J* = 3.4 Hz, 1H, H1$^{A}$), 4.79 (1H, H5$^{D}$), 4.68 (d, *J* = 2.8 Hz, 1H, H5$^{B}$), 4.50 – 4.46 (m, 2H, H1$^{F}$, H1$^{H}$), 4.37 (dd, *J* = 11.5, 2.4 Hz, 1H, H6a$^{C}$), 4.31 – 4.27 (m, 2H, H2$^{D}$, H2$^{B}$), 4.27 – 4.22 (m, 1H, H6b$^{C}$), 4.20 – 4.15 (m, 2H, H3$^{D}$, H3$^{B}$ ), 4.09 (t, *J* = 3.3 Hz, 1H, H4$^{B}$), 4.06 (t, *J* = 3.1 Hz, 1H, H4$^{D}$), 4.00 – 3.96 (m, 1H, H5$^{C}$), 3.93 – 3.63 (m, 22H, H5$^{E}$, H5$^{G}$, H5$^{A}$, H3$^{G}$, H2$^{G}$, H6a$^{A}$, H6b$^{A}$, H6a$^{E}$, H6b$^{E}$, H6a$^{G}$, H6b$^{G}$, H3$^{A}$, H5$^{F}$, H4$^{F}$, H4$^{E}$, H4$^{G}$, H4$^{C}$, H4$^{A}$, H3$^{F}$, H3$^{E}$, H4$^{E}$, OCH*H* linker), 3.60 (t, *J* = 9.6 Hz, 1H, H3$^{C}$), 3.54 – 3.47 (m, 3H, OCH*H* linker, H4$^{H}$, H3$^{H}$), 3.38 – 3.33 (m, 2H, H2$^{F}$, H2$^{H}$), 3.26 (dd, *J* = 10.5, 3.5 Hz, 1H, H2$^{C}$), 3.24 – 3.20 (m, 2H, H2$^{E}$, H2$^{A}$), 3.01 (t, *J* = 7.3 Hz, 2H, C$H_2$N linker), 2.03 (s, 3H, NHCOC$H_3$), 1.75 – 1.42 (m, 6H, 3 × C$H_2$ linker). $^{13}$C NMR from HSQC (151 MHz, $D_2O$) δ 102.3 (C1$^{F}$, C1$^{H}$), 99.2 (C1$^{D}$), 97.7 (C1$^{B}$), 96.9 (C1$^{A}$), 96.7 (C1$^{E}$, C1$^{G}$), 96.1 (C1$^{C}$), 78.1 (C3$^{F}$), 76.9 (C2$^{B}$, C4$^{A}$), 75.9 (C2$^{D}$, C4$^{B}$, C4$^{D}$, C5$^{F}$, C4$^{F}$, C5$^{H}$, C4$^{E}$, C4$^{G}$, C4$^{C}$), 75.1 (C3$^{H}$), 73.2 (C2$^{F}$, C2$^{H}$), 71.8 (C4$^{H}$), 70.6 (C5$^{E}$, C5$^{A}$, C5$^{G}$, C3$^{G}$), 69.9 (C5$^{B}$), 69.5 (C3$^{E}$, C3$^{C}$), 69.2 (C5$^{D}$, C3$^{D}$, C3$^{B}$, C5$^{C}$), 68.6 (C3$^{A}$), 67.7 (O<u>C</u>H$_2$ linker), 66.4 (C6$^{C}$), 59.6 (C6$^{A}$, C6$^{E}$, C6$^{G}$), 57.9 (C2$^{C}$, C2$^{E}$, C2$^{A}$), 53.3 (C2$^{G}$), 39.6 (N<u>C</u>H$_2$ linker), 28.0 (<u>C</u>H$_2$ linker), 26.5 (<u>C</u>H$_2$ linker),

22.7 ($\underline{C}H_2$ linker), 21.9 (NHCO$\underline{C}H_3$). HRMS (ESI-MS): m/z calculated for $C_{55}H_{89}N_5O_{60}S_6$ [M-10Na+8H]$^{2-}$: 985.6201; found: 985.6208.

$^1$H NMR (600 MHz, $D_2O$)

| | **H1** | **H2** | **H3** | **H4** | **H5** | **H6** |
|---|---|---|---|---|---|---|
| **A** | 5.11 (d, *J* = 3.4 Hz) | 3.22 | 3.81 | 3.69 | 3.84 | 3.85 |
| **B** | 5.26 (d, *J* = 3.4 Hz) | 4.29 | 4.16 | 4.09 (t, *J* = 3.3 Hz) | 4.68 (d, *J* = 2.8 Hz) | – |
| **C** | 5.41 (d, *J* = 3.5 Hz) | 3.25 (dd, *J* = 10.5, 3.5 Hz) | 3.60 | 3.74 | 3.98 | 4.37, 4.24 |
| **D** | 5.18 | 4.30 | 4.19 | 4.05 (t, *J* = 3.1 Hz) | 4.79 | – |
| **E** | 5.38 | 3.22 | 3.67 | 3.78 | 3.90 | 3.85 |
| **F** | 4.47 | 3.37 | 3.66 | 3.74 | 3.78 | – |
| **G** | 5.38 | 3.87 | 3.85 | 3.73 | 3.85 | 3.85 |
| **H** | 4.48 | 3.37 | 3.50 | 3.50 | 3.73 | – |

$^{13}$C NMR from HSQC (151 MHz, $D_2O$)

| | **C1** | **C2** | **C3** | **C4** | **C5** | **C6** |
|---|---|---|---|---|---|---|
| **A** | 96.9 | 57.9 | 68.6 | 76.9 | 70.6 | 59.6 |
| **B** | 97.7 | 76.9 | 69.2 | 75.9 | 69.9 | – |
| **C** | 96.1 | 57.9 | 69.5 | 75.9 | 69.2 | 66.4 |
| **D** | 99.2 | 75.9 | 69.2 | 75.9 | 69.2 | – |
| **E** | 96.7 | 57.9 | 69.5 | 75.9 | 70.6 | 59.6 |
| **F** | 102.3 | 73.2 | 78.1 | 75.9 | 75.9 | – |
| **G** | 96.7 | 53.3 | 70.6 | 75.9 | 70.6 | 59.6 |
| **H** | 102.3 | 73.2 | 75.1 | 71.8 | 75.9 | – |

**5-Aminopentyl *O*-(2-acetamido-6-azido-2-deoxy-α-D-glucopyranosyl)-(1→4)-*O*-(β-D-glucopyranosyluronate)-(1→4)-*O*-(2-acetamido-2-deoxy-α-D-glucopyranosyl)-(1→4)-*O*-(β-D-glucopyranosyluronate)-(1→4)-*O*-(2-sulfamino-2-deoxy-α-D-glucopyranosyl)-(1→4)-*O*-(2-*O*-sulfate-α-L-idopyranosyluronate)-(1→4)-*O*-(2-sulfamino-6-*O*-sulfate-2-deoxy-α-D-glucopyranosyl)-(1→4)-*O*-(2-*O*-sulfate-α-L-idopyranosyluronate)-(1→4)-*O*-2-sulfamino-2-deoxy-α-D-glucopyranoside, sodium salt (4).**

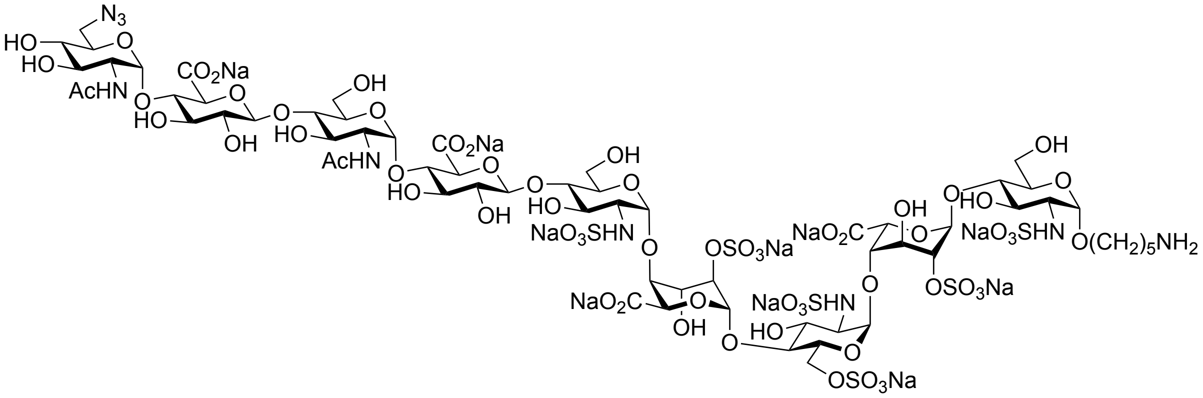

Compound **23** (2.7 mg, 1.2 μmol) was subjected to installation of unnatural α (1→4) 6-azido-GlcNAc according to the general procedure to give the title compound **4** as a white powder (2.6 mg, 87%). $^{1}$H NMR (600 MHz, $D_2O$) δ 5.46 (d, $J$ = 3.3 Hz, 1H, H1$^{C}$), 5.43 (d, $J$ = 3.7 Hz, 1H, H1$^{I}$), 5.41 (d, $J$ = 3.2 Hz, 1H, H1$^{E}$), 5.40 (d, $J$ = 3.5 Hz, 1H, H1$^{G}$), 5.32 (bs, 1H, H1$^{B}$), 5.22 (bs, 1H, H1$^{D}$), 5.14 (d, $J$ = 3.4 Hz, 1H, H1$^{A}$), 4.82 (1H, H5$^{D}$), 4.69 (bs, 1H, H5$^{B}$), 4.49 (d, $J$ = 7.9 Hz, 2H, H1$^{F}$, H1$^{H}$), 4.43 – 4.37 (m, 1H, H6a$^{C}$), 4.35 – 4.30 (m, 2H, H2$^{D}$, H2$^{B}$), 4.30 – 4.26 (m, 1H, H6b$^{C}$), 4.24 – 4.17 (m, 2H, H3$^{D}$, H3$^{B}$ ), 4.15 – 4.11 (m, 1H, H4$^{B}$), 4.10 – 4.07 (m, 1H, H4$^{D}$), 4.04 – 3.99 (m, 1H, H5$^{C}$), 3.97 – 3.59 (m, 30H, H2$^{G}$, H2$^{I}$, H5$^{E}$, H5$^{I}$, H5$^{A}$, H5$^{G}$, H3$^{G}$, H3$^{A}$, H6a$^{A}$, H6b$^{A}$, H6a$^{E}$, H6b$^{E}$, H6a$^{G}$, H6b$^{G}$, H5$^{F}$, H5$^{H}$, H4$^{G}$, H4$^{C}$, H4$^{F}$, H4$^{H}$, H4$^{A}$, H4$^{E}$, H3$^{F}$, H3$^{H}$, OCH*H* linker, H3$^{I}$, H3$^{E}$, H3$^{C}$, H6a$^{I}$, H6b$^{I}$), 3.58 – 3.53 (m, 1H, OCH*H* linker), 3.49 (t, $J$ = 9.5 Hz, 1H, H4$^{I}$), 3.41 – 3.35 (m, 2H, H2$^{F}$, H2$^{H}$), 3.29 (dd, $J$ = 10.3, 3.3 Hz, 1H, H2$^{C}$), 3.25 (dd, $J$ = 10.2, 3.3 Hz, 2H, H2$^{E}$, H2$^{A}$), 3.05 (t, $J$ = 7.3 Hz, 2H, C$H_2$N linker), 2.06 (s, 3H, NHCOC$H_3$), 2.05 (s, 3H, NHCOC$H_3$), 1.79 – 1.48 (m, 6H, 3 × C$H_2$ linker). $^{13}$C NMR from HSQC (151 MHz, $D_2O$) δ 102.3 (C1$^{F}$, C1$^{H}$), 99.0 (C1$^{D}$), 97.4 (C1$^{B}$), 97.0 (C1$^{A}$), 96.8 (C1$^{I}$), 96.7 (C1$^{E}$, C1$^{G}$), 96.0 (C1$^{C}$), 78.0 (C3$^{F}$, C3$^{H}$), 77.0 (C2$^{B}$), 76.4 (C2$^{D}$, C5$^{F}$, C5$^{H}$, C4$^{F}$, C4$^{H}$, C4$^{C}$, C4$^{G}$, C4$^{A}$, C4$^{E}$), 75.9 (C4$^{B}$, C4$^{D}$), 73.4 (C2$^{F}$, C2$^{H}$), 70.5 (C5$^{E}$, C5$^{G}$, C5$^{A}$, C4$^{G}$, C5$^{I}$, C3$^{I}$, C4$^{I}$),

70.0 ($C5^B$), 69.6 ($C3^G$, $C3^B$, $C3^E$, $C3^C$), 69.3 ($C5^D$, $C3^D$, $C5^C$), 68.7 ($C3^A$), 67.7 (O<u>C</u>$H_2$ linker), 66.4 ($C6^C$), 66.3 ($C6^G$), 59.5 ($C6^A$, $C6^E$), 57.9 ($C2^C$, $C2^E$, $C2^A$), 53.4 ($C2^G$, $C2^I$), 50.4 ($C6^I$), 39.6 (N<u>C</u>$H_2$ linker), 28.0 (<u>C</u>$H_2$ linker), 26.4 (<u>C</u>$H_2$ linker), 22.7 (<u>C</u>$H_2$ linker), 21.9 (2 × NHCO<u>C</u>$H_3$). ESI-MS: m/z calculated for $C_{63}H_{101}N_9O_{64}S_6$ $[M-10Na+8H]^{2-}$: 1099.6630; found: 1099.6403.

$^1H$ NMR (600 MHz, $D_2O$)

| | H1 | H2 | H3 | H4 | H5 | H6 |
|---|---|---|---|---|---|---|
| **A** | 5.14 (d, *J* = 3.4 Hz) | 3.25 (dd, *J* = 10.2, 3.3 Hz) | 3.88 | 3.73 | 3.89 | 3.88 |
| **B** | 5.32 | 4.33 | 4.19 | 4.13 | 4.69 | – |
| **C** | 5.46 (d, *J* = 3.3 Hz) | 3.29 (dd, *J* = 10.3, 3.3 Hz) | 3.62 | 3.78 | 4.02 | 4.40, 4.28 |
| **D** | 5.22 | 4.33 | 4.21 | 4.09 | 4.82 | – |
| **E** | 5.41 (d, *J* = 3.2 Hz) | 3.25 (dd, *J* = 10.2, 3.3 Hz) | 3.71 | 3.70 | 3.94 | 3.88 |
| **F** | 4.49 (d, *J* = 7.9 Hz) | 3.38 | 3.70 | 3.76 | 3.81 | – |
| **G** | 5.40 (d, *J* = 3.5 Hz) | 3.90 | 3.67 | 3.67 | 3.90 | 3.83 |
| **H** | 4.49 (d, *J* = 7.9 Hz) | 3.38 | 3.70 | 3.76 | 3.81 | – |
| **I** | 5.43 (d, *J* = 3.7 Hz) | 3.92 | 3.72 | 3.49 | 3.90 | 3.66 |

$^{13}C$ NMR from HSQC (151 MHz, $D_2O$)

| | C1 | C2 | C3 | C4 | C5 | C6 |
|---|---|---|---|---|---|---|
| **A** | 97.0 | 57.9 | 68.7 | 76.4 | 70.5 | 59.5 |
| **B** | 97.4 | 77.0 | 69.6 | 75.9 | 70.0 | – |
| **C** | 96.0 | 57.9 | 69.6 | 76.4 | 69.3 | 66.4 |
| **D** | 99.0 | 76.4 | 69.3 | 75.9 | 69.3 | – |
| **E** | 96.7 | 57.9 | 69.6 | 76.4 | 70.5 | 59.5 |
| **F** | 102.3 | 73.4 | 78.0 | 76.4 | 76.4 | – |
| **G** | 96.7 | 53.4 | 69.6 | 76.4 | 70.5 | 66.3 |
| **H** | 102.3 | 73.4 | 78.0 | 76.4 | 76.4 | |

| | | | | | | |
|---|---|---|---|---|---|---|
| **I** | 96.8 | 53.4 | 70.5 | 70.5 | 70.5 | 50.4 |

**5-Aminopentyl *O*-(2-acetamido-2-deoxy-α-D-glucopyranosyl)-(1→4)-*O*-(β-D-glucopyranosyluronate)-(1→4)-*O*-(2-acetamido-2-deoxy-α-D-glucopyranosyl)-(1→4)-*O*-(β-D-glucopyranosyluronate)-(1→4)-*O*-(2-sulfamino-2-deoxy-α-D-glucopyranosyl)-(1→4)-*O*-(2-*O*-sulfate-α-L-idopyranosyluronate)-(1→4)-*O*-(2-sulfamino-6-*O*-sulfate-2-deoxy-α-D-glucopyranosyl)-(1→4)-*O*-(2-*O*-sulfate-α-L-idopyranosyluronate)-(1→4)-*O*-2-sulfamino-2-deoxy-α-D-glucopyranoside, sodium salt (S5).**

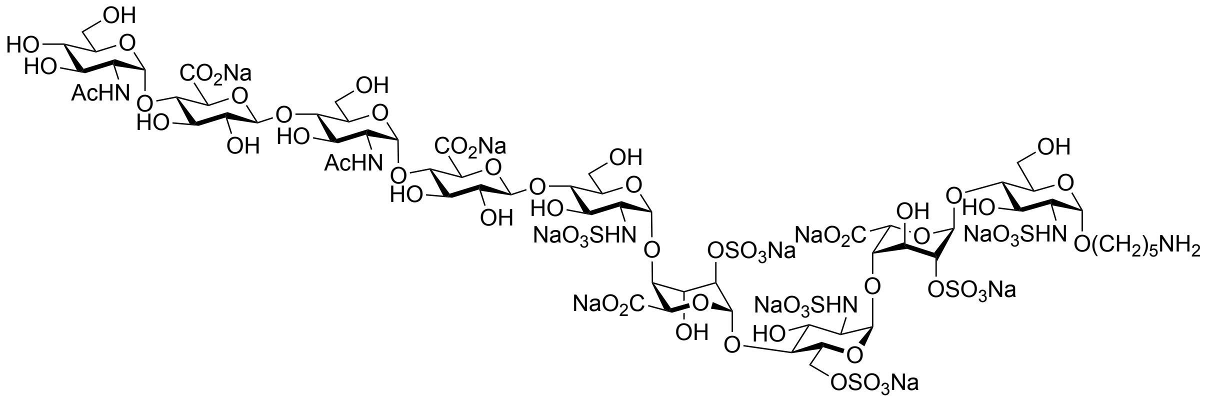

Compound **23** (21.0 mg, 9.6 μmol) was subjected to installation of α (1→4)-GlcNAc according to the general procedure to give the title compound **S5** as a white powder (20.0 mg, 87%). $^{1}$H NMR (600 MHz, $D_2O$) δ 5.46 (d, *J* = 3.4 Hz, 1H, H1$^{C}$), 5.42 (d, *J* = 3.7 Hz, 1H, H1$^{I}$), 5.41 (d, *J* = 3.4 Hz, 1H, H1$^{E}$), 5.40 (d, *J* = 3.6 Hz, 1H, H1$^{G}$), 5.31 (d, *J* = 3.4 Hz, 1H, H1$^{B}$), 5.21 (d, *J* = 2.2 Hz, 1H, H1$^{D}$), 5.14 (d, *J* = 3.4 Hz, 1H, H1$^{A}$), 4.78 (1H, H5$^{D}$), 4.69 (d, *J* = 2.9 Hz, 1H, H5$^{B}$), 4.50 (d, *J* = 7.9 Hz, 2H, H1$^{F}$, H1$^{H}$), 4.40 (dd, *J* = 11.5, 2.4 Hz, 1H, H6a$^{C}$), 4.34 – 4.30 (m, 2H, H2$^{D}$, H2$^{B}$), 4.29 – 4.25 (m, 1H, H6b$^{C}$), 4.23 – 4.17 (m, 2H, H3$^{D}$, H3$^{B}$ ), 4.12 (t, *J* = 3.3 Hz, 1H, H4$^{B}$), 4.08 (t, *J* = 3.1 Hz, 1H, H4$^{D}$), 4.03 – 3.99 (m, 1H, H5$^{C}$), 3.95 – 3.64 (m, 29H, H5$^{E}$, H2$^{G}$, H2$^{I}$, H6a$^{A}$, H6b$^{A}$, H6a$^{E}$, H6b$^{E}$, H6a$^{G}$, H6b$^{G}$, H6a$^{I}$, H6b$^{I}$, H3$^{A}$, H5$^{A}$, H3$^{G}$, H5$^{G}$, H5$^{F}$, H5$^{H}$, H4$^{E}$, H4$^{C}$, H4$^{F}$, H4$^{H}$, H4$^{A}$, H4$^{G}$, H3$^{F}$, H3$^{H}$, H3$^{I}$, H5$^{I}$, H3$^{E}$, OCH*H* linker), 3.65 – 3.60 (m, 1H, H3$^{C}$), 3.57 – 3.53 (m, 1H, OCH*H* linker), 3.40 – 3.36 (m, 2H, H2$^{F}$, H2$^{H}$), 3.48 (t, *J* = 9.6 Hz, 1H, H4$^{I}$), 3.29 (dd, *J* = 10.5, 3.5 Hz, 1H, H2$^{C}$), 3.25 (dd, *J* = 10.3, 3.4 Hz, 2H, H2$^{E}$, H2$^{A}$), 3.05 (t, *J* = 7.2 Hz, 2H, C$H_2$N linker), 2.06 (s, 3H, NHCOC$H_3$), 2.05 (s, 3H, NHCOC$H_3$), 1.79 – 1.48 (m, 6H, 3 × C$H_2$ linker). $^{13}$C NMR from HSQC (151 MHz, $D_2O$) δ 102.3 (C1$^{F}$, C1$^{H}$), 99.2 (C1$^{D}$), 97.5 (C1$^{B}$), 97.0 (C1$^{A}$), 96.8 (C1$^{I}$, C1$^{E}$, C1$^{G}$), 96.0 (C1$^{C}$), 78.1 (C3$^{F}$, C3$^{H}$), 76.7

(C2$^{B}$), 76.2 (C5$^{F}$, C5$^{H}$, C4$^{E}$, C4$^{C}$, C4$^{F}$, C4$^{H}$, C4$^{A}$, C4$^{G}$), 75.9 (C2$^{D}$, C4$^{B}$, C4$^{D}$), 73.4 (C2$^{F}$, C2$^{H}$), 71.5 (C3$^{I}$, C5$^{I}$), 70.6 (C5$^{E}$, C5$^{A}$, C5$^{G}$, C3$^{G}$), 69.9 (C5$^{B}$), 69.7 (C4$^{I}$), 69.5 (C3$^{E}$, C3$^{C}$, C3$^{B}$), 69.2 (C5$^{D}$, C3$^{D}$, C5$^{C}$), 68.7 (C3$^{A}$), 67.7 (O<u>C</u>H$_2$ linker), 66.4 (C6$^{C}$), 60.0 (C6$^{G}$, C6$^{I}$), 59.6 (C6$^{A}$, C6$^{E}$), 57.9 (C2$^{C}$, C2$^{E}$, C2$^{A}$), 53.4 (C2$^{G}$, C2$^{I}$), 39.5 (N<u>C</u>H$_2$ linker), 28.1 (<u>C</u>H$_2$ linker), 26.4 (<u>C</u>H$_2$ linker), 22.7 (<u>C</u>H$_2$ linker), 21.90 (2 × NHCO<u>C</u>H$_3$). HRMS (ESI-MS): m/z calculated for $C_{63}H_{102}N_6O_{65}S_6$ [M-9Na+7H]$^{2-}$: 1087.1598; found: 1087.1601.

$^{1}$H NMR (600 MHz, $D_2O$)

| | **H1** | **H2** | **H3** | **H4** | **H5** | **H6** |
|---|---|---|---|---|---|---|
| **A** | 5.14 (d, *J* = 3.4 Hz) | 3.25 (dd, *J* = 10.3, 3.4 Hz) | 3.87 | 3.73 | 3.88 | 3.88 |
| **B** | 5.31 (d, *J* = 3.4 Hz) | 4.32 | 4.19 | 4.12 (t, *J* = 3.3 Hz) | 4.69 (d, *J* = 2.9 Hz) | – |
| **C** | 5.46 (d, *J* = 3.4 Hz) | 3.28 (dd, *J* = 10.5, 3.5 Hz) | 3.63 | 3.78 | 4.01 | 4.40, 4.28 |
| **D** | 5.21 (d, *J* = 2.2 Hz) | 4.33 | 4.21 | 4.08 (t, *J* = 3.1 Hz) | 4.78 | – |
| **E** | 5.41 (d, *J* = 3.4 Hz) | 3.25 (dd, *J* = 10.3, 3.4 Hz) | 3.71 | 3.70 | 3.94 | 3.88 |
| **F** | 4.50 (d, *J* = 7.9 Hz) | 3.38 | 3.70 | 3.76 | 3.81 | – |
| **G** | 5.40 (d, *J* = 3.6 Hz) | 3.89 | 3.88 | 3.80 | 3.89 | 3.81 |
| **H** | 4.50 (d, *J* = 7.9 Hz) | 3.38 | 3.70 | 3.76 | 3.81 | – |
| **I** | 5.42 (d, *J* = 3.7 Hz) | 3.89 | 3.74 | 3.49 (t, *J* = 9.6 Hz) | 3.74 | 3.81 |

$^{13}$C NMR from HSQC (151 MHz, $D_2O$)

| | **C1** | **C2** | **C3** | **C4** | **C5** | **C6** |
|---|---|---|---|---|---|---|
| **A** | 97.0 | 57.9 | 68.7 | 76.2 | 70.6 | 59.6 |
| **B** | 97.5 | 76.7 | 69.5 | 75.9 | 69.9 | – |
| **C** | 96.0 | 57.9 | 69.5 | 76.2 | 69.2 | 66.4 |
| **D** | 99.2 | 75.9 | 69.2 | 75.9 | 69.2 | – |

| | | | | | | |
|---|---|---|---|---|---|---|
| **E** | 96.8 | 57.9 | 69.5 | 76.2 | 70.6 | 59.6 |
| **F/H** | 102.3 | 73.4 | 78.1 | 76.2 | 76.2 | – |
| **G** | 96.8 | 53.4 | 70.6 | 76.2 | 70.6 | 60.0 |
| **I** | 96.8 | 53.4 | 71.5 | 69.7 | 71.5 | 60.0 |

**5-Aminopentyl *O*-(β-D-glucopyranosyluronate)-(1 → 4)-*O*-(2-acetamido-2-deoxy-α-D-glucopyranosyl)-(1→4)-*O*-(β-D-glucopyranosyluronate)-(1→4)-*O*-(2-acetamido-2-deoxy-α-D-glucopyranosyl)-(1 → 4)-*O*-(β-D-glucopyranosyluronate)-(1 → 4)-*O*-(2-sulfamino-2-deoxy-α-D-glucopyranosyl)-(1 → 4)-*O*-(2-*O*-sulfate-α-L-idopyranosyluronate)-(1 → 4)-*O*-(2-sulfamino-6-*O*-sulfate-2-deoxy-α-D-glucopyranosyl)-(1 → 4)-*O*-(2-*O*-sulfate-α-L-idopyranosyluronate)-(1→4)-*O*-2-sulfamino-2-deoxy-α-D-glucopyranoside, sodium salt (24).**

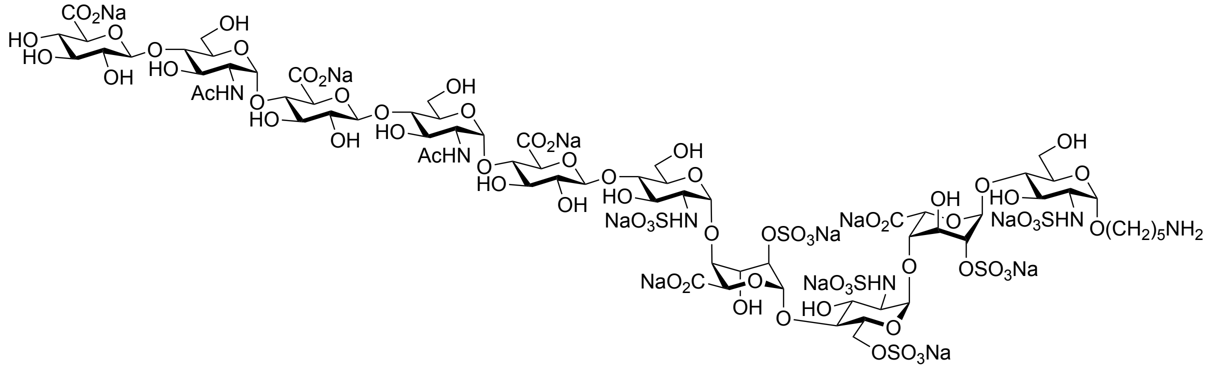

Compound **S5** (20.0 mg, 8.3 μmol) was subjected to installation of β (1→4)-GlcA according to the general procedure to give the title compound **24** as a white powder (19.8 mg, 91%). $^1$H NMR (600 MHz, $D_2O$) δ 5.42 (d, *J* = 2.9 Hz, 1H, H1$^C$), 5.40 – 5.35 (m, 3H, H1$^E$, H1$^G$, H1$^I$), 5.27 (d, *J* = 2.7 Hz, 1H, H1$^B$), 5.19 (bs, 1H, H1$^D$), 5.11 (d, *J* = 2.9 Hz, 1H, H1$^A$), 4.78 (1H, H5$^D$), 4.67 (bs, 1H, H5$^B$), 4.51 – 4.43 (m, 3H, H1$^F$, H1$^H$, H1$^J$), 4.40 – 4.35 (m, 1H, H6a$^C$), 4.32 – 4.27 (m, 2H, H2$^D$, H2$^B$), 4.27 – 4.22 (m, 1H, H6b$^C$), 4.21 – 4.14 (m, 2H, H3$^D$, H3$^B$), 4.12 – 4.07 (m, 1H, H4$^B$), 4.07 – 4.03 (m, 1H, H4$^D$), 4.00 – 3.96 (m, 1H, H5$^C$), 3.93 – 3.57 (m, 32H, H5$^E$, H2$^G$, H2$^I$, H6a$^A$, H6b$^A$, H6a$^E$, H6b$^E$, H6a$^G$, H6b$^G$, H6a$^I$, H6b$^I$, H3$^A$, H5$^A$, H3$^G$, H3$^I$, H5$^G$, H5$^I$, H5$^F$, H5$^H$, H4$^G$, H4$^I$, H4$^C$, H5$^J$, H4$^F$, H4$^H$, H4$^A$, H4$^E$, H3$^F$, H3$^H$, OCH*H* linker, H3$^E$, H3$^C$), 3.55 – 3.47 (m, 3H, OCH*H* linker, H4$^J$, H3$^J$), 3.39 – 3.31 (m, 2H, H2$^F$, H2$^H$, H2$^J$), 3.26 (dd, *J*

= 10.5, 2.9 Hz, 1H, H2$^C$), 3.24 – 3.20 (m, 2H, H2$^E$, H2$^A$), 3.02 (t, *J* = 7.2 Hz, 2H, C*H*$_2$N linker), 2.02 (s, 3H, NHCOC*H*$_3$), 2.02 (s, 3H, NHCOC*H*$_3$), 1.75 – 1.45 (m, 6H, 3 × C*H*$_2$ linker). $^{13}$C NMR from HSQC (151 MHz, $D_2O$) δ 102.3 (C1$^F$, C1$^H$, C1$^J$), 99.2 (C1$^D$), 97.6 (C1$^B$), 96.9 (C1$^A$), 96.6 (C1$^I$, C1$^E$, C1$^G$), 96.0 (C1$^C$), 78.2 (C3$^F$, C3$^H$), 76.9 (C2$^B$), 76.2 (C2$^D$, C5$^F$, C5$^H$, C4$^G$, C4$^I$, C4$^C$, C4$^F$, C4$^H$, C5$^J$, C4$^A$, C4$^E$), 75.8 (C4$^B$, C4$^D$), 75.1 (C3$^J$), 73.2 (C2$^F$, C2$^H$, C2$^J$), 71.8 (C4$^J$), 70.6 (C5$^E$, C5$^A$, C5$^G$, C5$^I$, C3$^G$, C3$^I$), 69.8 (C5$^B$), 69.5 (C3$^E$, C3$^C$, C3$^B$), 69.2 (C5$^D$, C3$^D$, C5$^C$), 68.8 (C3$^A$), 67.7 (O<u>C</u>H$_2$ linker), 66.2 (C6$^C$), 59.5 (C6$^A$, C6$^E$, C6$^G$, C6$^I$), 57.9 (C2$^C$, C2$^E$, C2$^A$), 53.1 (C2$^G$, C2$^I$), 39.6 (N<u>C</u>H$_2$ linker), 28.1 (<u>C</u>H$_2$ linker), 26.5 (<u>C</u>H$_2$ linker), 22.6 (<u>C</u>H$_2$ linker), 21.9 (2 × NHCO<u>C</u>H$_3$). HRMS (ESI-MS): m/z calculated for $C_{69}H_{110}N_6O_{71}S_6$ [M-11Na+9H]$^{2-}$: 1175.1758; found: 1175.1749.

$^1$H NMR (600 MHz, $D_2O$)

| | **H1** | **H2** | **H3** | **H4** | **H5** | **H6** |
|---|---|---|---|---|---|---|
| **A** | 5.11 (d, *J* = 2.9 Hz) | 3.22 | 3.83 | 3.69 | 3.84 | 3.85 |
| **B** | 5.27 (d, *J* = 2.7 Hz) | 4.30 | 4.16 | 4.09 | 4.67 | – |
| **C** | 5.42 (d, *J* = 2.9 Hz) | 3.26 (dd, *J* = 10.5, 2.9 Hz) | 3.60 | 3.75 | 3.98 | 4.37, 4.24 |
| **D** | 5.19 | 4.30 | 4.19 | 4.06 | 4.79 | – |
| **E** | 5.38 | 3.22 | 3.67 | 3.67 | 3.90 | 3.85 |
| **F/H** | 4.47 | 3.35 | 3.66 | 3.74 | 3.78 | – |
| **G/I** | 5.37 | 3.87 | 3.85 | 3.78 | 3.85 | 3.81 |
| **J** | 4.48 | 3.36 | 3.50 | 3.50 | 3.73 | – |

$^{13}$C NMR from HSQC (151 MHz, $D_2O$)

| | **C1** | **C2** | **C3** | **C4** | **C5** | **C6** |
|---|---|---|---|---|---|---|
| **A** | 96.9 | 57.9 | 68.8 | 76.2 | 70.6 | 59.5 |

| | | | | | | |
|---|---|---|---|---|---|---|
| **B** | 97.6 | 76.9 | 69.5 | 75.8 | 69.8 | – |
| **C** | 96.0 | 57.9 | 69.5 | 76.2 | 69.2 | 66.2 |
| **D** | 99.2 | 76.2 | 69.2 | 75.8 | 69.2 | – |
| **E** | 96.6 | 57.9 | 69.5 | 76.2 | 70.6 | 59.5 |
| **F/H** | 102.3 | 73.2 | 78.2 | 76.2 | 76.2 | – |
| **G/I** | 96.6 | 53.1 | 70.6 | 76.2 | 70.6 | 59.5 |
| **J** | 102.3 | 73.2 | 75.1 | 71.8 | 76.2 | – |

**5-Aminopentyl *O*-(2-acetamido-6-azido-2-deoxy-α-D-glucopyranosyl)[-(1→4)-*O*-(β-D-glucopyranosyluronate)-(1→4)-*O*-(2-acetamido-2-deoxy-α-D-glucopyranosyl)]$_2$-(1→4)-*O*-(β-D-glucopyranosyluronate)-(1→4)-*O*-(2-sulfamino-2-deoxy-α-D-glucopyranosyl)-(1→4)-*O*-(2-*O*-sulfate-α-L-idopyranosyluronate)-(1→4)-*O*-(2-sulfamino-6-*O*-sulfate-2-deoxy-α-D-glucopyranosyl)-(1→4)-*O*-(2-*O*-sulfate-α-L-idopyranosyluronate)-(1→4)-*O*-2-sulfamino-2-deoxy-α-D-glucopyranoside, sodium salt (5).**

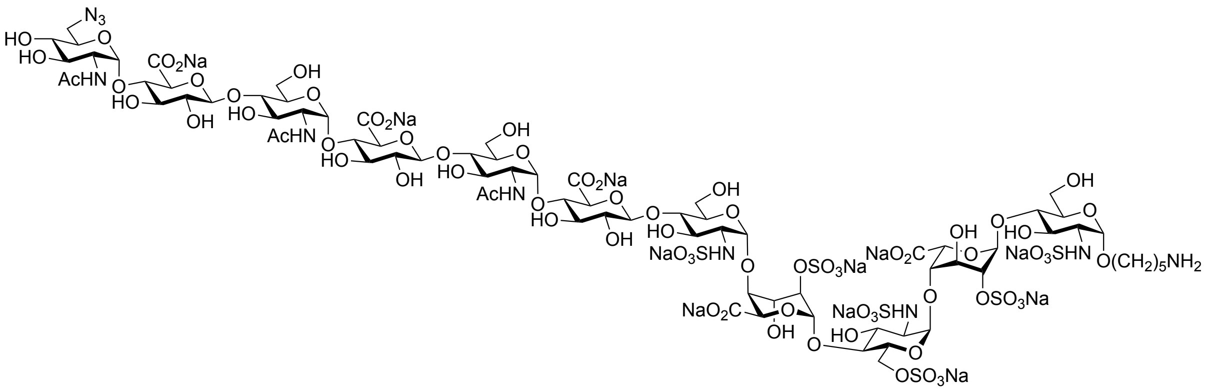

Compound **24** (2.2 mg, 0.85 μmol) was subjected to installation of unnatural α (1→4) 6-azido-GlcNAc according to the general procedure to give the title compound **5** as a white powder (2.0 mg, 84%). $^1$H NMR (600 MHz, $D_2O$) δ 5.47 (d, $J$ = 3.4 Hz, 1H, H1$^C$), 5.43 (d, $J$ = 3.8 Hz, 1H, H1$^K$), 5.42 (d, $J$ = 3.4 Hz, 1H, H1$^E$), 5.40 (d, $J$ = 3.5 Hz, 2H, H1$^G$, H1$^I$), 5.32 (bs, 1H, H1$^B$), 5.22 (bs, 1H, H1$^D$), 5.14 (d, $J$ = 3.4 Hz, 1H, H1$^A$), 4.81 (1H, H5$^D$), 4.68 (bs, 1H, H5$^B$), 4.49 (d, $J$ = 7.8 Hz, 3H, H1$^F$, H1$^H$, H1$^J$), 4.43 – 4.37 (m, 1H, H6a$^C$), 4.35 – 4.30 (m, 2H, H2$^D$, H2$^B$), 4.30 – 4.26 (m, 1H, H6b$^C$), 4.24 – 4.20 (m, 1H, H3$^D$), 4.20 – 4.17 (m, 1H, H3$^B$ ), 4.16 – 4.12 (m, 1H, H4$^B$), 4.09 (bs, 1H, H4$^D$), 4.04 – 3.98 (m, 1H, H5$^C$), 3.97 – 3.58 (m, 39H, H2$^G$, H2$^I$, H2$^K$, H6a$^A$, H6b$^A$, H6a$^E$, H6b$^E$, H6a$^G$, H6b$^G$, H6a$^I$, H6b$^I$, H6a$^K$, H6b$^K$, H5$^E$, H5$^G$, H5$^I$, H5$^K$, H5$^A$, H3$^A$, OCH*H* linker, H3$^E$, H3$^G$, H3$^I$, H3$^K$, H3$^C$, H5$^F$, H5$^H$, H5$^J$, H4$^C$, H4$^F$, H4$^H$, H4$^J$, H4$^A$,

H4$^{E}$, H4$^{G}$, H4$^{I}$, H3$^{F}$, H3$^{H}$, H3$^{J}$), 3.58 – 3.53 (m, 1H, OCH*H* linker), 3.49 (t, *J* = 9.6 Hz, 1H, H4$^{K}$), 3.41 – 3.35 (m, 3H, H2$^{F}$, H2$^{H}$, H2$^{J}$), 3.29 (dd, *J* = 10.3, 3.4 Hz, 1H, H2$^{C}$), 3.25 (dd, *J* = 10.3, 3.4 Hz, 2H, H2$^{E}$, H2$^{A}$), 3.05 (t, *J* = 7.3 Hz, 2H, C$H_2$N linker), 2.06 (s, 3H, NHCOC$H_3$), 2.05 (s, 3H, NHCOC$H_3$), 2.05 (s, 3H, NHCOC$H_3$), 1.79 – 1.48 (m, 6H, 3 × C$H_2$ linker). $^{13}$C NMR from HSQC (151 MHz, $D_2O$) δ 102.3 (C1$^{F}$, C1$^{H}$, C1$^{J}$), 99.2 (C1$^{D}$), 97.3 (C1$^{B}$), 97.0 (C1$^{A}$), 96.8 (C1$^{K}$), 96.7 (C1$^{E}$, C1$^{G}$, C1$^{I}$), 96.0 (C1$^{C}$), 78.0 (C3$^{F}$, C3$^{H}$, C3$^{J}$), 77.0 (C2$^{B}$), 76.4 (C2$^{D}$, C5$^{F}$, C5$^{H}$, C5$^{J}$, C4$^{F}$, C4$^{H}$, C4$^{J}$, C4$^{C}$, C4$^{G}$, C4$^{I}$, C4$^{A}$, C4$^{E}$), 75.8 (C4$^{B}$, C4$^{D}$), 73.4 (C2$^{F}$, C2$^{H}$, C2$^{J}$), 70.5 (C5$^{G}$, C5$^{I}$, C5$^{K}$, C4$^{K}$, C3$^{K}$, C3$^{E}$, C5$^{B}$), 69.5 (C3$^{G}$, C3$^{I}$, C3$^{C}$, C3$^{B}$, C3$^{D}$, C5$^{D}$, C5$^{C}$, C5$^{E}$, C5$^{A}$, C3$^{A}$), 67.7 (O$\underline{C}H_2$ linker), 66.5 (C6$^{C}$), 59.7 (C6$^{A}$, C6$^{E}$, C6$^{G}$, C6$^{I}$), 57.9 (C2$^{C}$, C2$^{E}$, C2$^{A}$), 53.3 (C2$^{G}$, C2$^{I}$, C2$^{K}$), 50.4 (C6$^{K}$), 39.6 (N$\underline{C}H_2$ linker), 28.0 ($\underline{C}H_2$ linker), 26.4 ($\underline{C}H_2$ linker), 22.6 ($\underline{C}H_2$ linker), 21.7 (3 × NHCO$\underline{C}H_3$). HRMS (ESI-MS): m/z calculated for $C_{77}H_{122}N_{10}O_{75}S_6$ $[M-11Na+9H]^{2-}$: 1289.2187; found: 1289.2197.

$^{1}$H NMR (600 MHz, $D_2O$)

| | H1 | H2 | H3 | H4 | H5 | H6 |
|---|---|---|---|---|---|---|
| **A** | 5.14 (d, *J* = 3.4 Hz) | 3.25 (dd, *J* = 10.2, 3.4 Hz) | 3.88 | 3.73 | 3.90 | 3.88 |
| **B** | 5.32 | 4.33 | 4.19 | 4.13 | 4.68 | – |
| **C** | 5.47 (d, *J* = 3.4 Hz) | 3.29 (dd, *J* = 10.3, 3.4 Hz) | 3.62 | 3.78 | 4.00 | 4.39, 4.28 |
| **D** | 5.22 | 4.33 | 4.21 | 4.09 | 4.81 | – |
| **E** | 5.42 (d, *J* = 3.4 Hz) | 3.25 (dd, *J* = 10.2, 3.4 Hz) | 3.70 | 3.70 | 3.91 | 3.86 |
| **F/H/J** | 4.49 (d, *J* = 7.8 Hz) | 3.38 | 3.70 | 3.76 | 3.81 | – |
| **G/I** | 5.40 (d, *J* = 3.5 Hz) | 3.90 | 3.67 | 3.67 | 3.90 | 3.85 |
| **K** | 5.43 (d, *J* = 3.8 Hz) | 3.92 | 3.72 | 3.49 | 3.90 | 3.66 |

$^{13}$C NMR from HSQC (151 MHz, $D_2O$)

| | C1 | C2 | C3 | C4 | C5 | C6 |
|---|---|---|---|---|---|---|
| **A** | 97.0 | 57.9 | 69.9 | 76.4 | 69.9 | 59.7 |

| | | | | | | |
|---|---|---|---|---|---|---|
| **B** | 97.3 | 77.0 | 69.9 | 75.8 | 70.5 | – |
| **C** | 96.0 | 57.9 | 69.9 | 76.4 | 69.9 | 66.5 |
| **D** | 99.2 | 76.4 | 69.9 | 75.8 | 69.9 | – |
| **E** | 96.7 | 57.9 | 70.5 | 76.4 | 69.9 | 59.7 |
| **F/H/J** | 102.3 | 73.4 | 78.0 | 76.4 | 76.4 | – |
| **G/I** | 96.7 | 53.3 | 69.9 | 76.4 | 70.5 | 59.7 |
| **K** | 96.8 | 53.3 | 70.5 | 70.5 | 70.5 | 50.4 |

**5-Aminopentyl [*O*-(β-D-glucopyranosyluronate)-(1→4)-*O*-(2-acetamido-2-deoxy-α-D-glucopyranosyl)-(1→4)]$_3$-*O*-(β-D-glucopyranosyluronate)-(1→4)-*O*-(2-sulfamino-2-deoxy-α-D-glucopyranosyl)-(1→4)-*O*-(2-*O*-sulfate-α-L-idopyranosyluronate)-(1→4)-*O*-(2-sulfamino-6-*O*-sulfate-2-deoxy-α-D-glucopyranosyl)-(1→4)-*O*-(2-*O*-sulfate-α-L-idopyranosyluronate)-(1→4)-*O*-2-sulfamino-2-deoxy-α-D-glucopyranoside, sodium salt (25).**

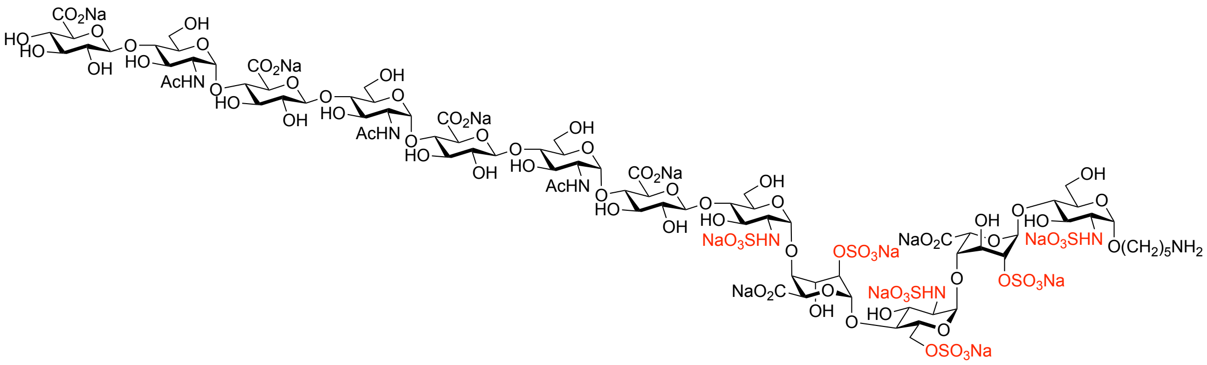

Compound **25** was obtained from compound **24** as a white powder according to the general procedure for installation of α (1→4)-GlcNAc and β (1→4)-GlcA, respectively. $^1$H NMR (600 MHz, $D_2O$) δ 5.43 (bs, 1H, H1$^C$), 5.40 – 5.33 (m, 4H, H1$^E$, H1$^G$, H1$^I$, H1$^K$), 5.29 (bs, 1H, H1$^B$), 5.19 (bs, 1H, H1$^D$), 5.11 (d, $J$ = 3.5 Hz, 1H, H1$^A$), 4.78 (1H, H5$^D$), 4.65 (bs, 1H, H5$^B$), 4.50 – 4.43 (m, 4H, H1$^F$, H1$^H$, H1$^J$, H1$^L$), 4.40 – 4.34 (m, 1H, H6a$^C$), 4.31 – 4.27 (m, 2H, H2$^D$, H2$^B$), 4.26 – 4.22 (m, 1H, H6b$^C$), 4.20 – 4.17 (m, 1H, H3$^D$), 4.17 – 4.14 (m, 1H, H3$^B$ ), 4.11 – 4.08 (m, 1H, H4$^B$), 4.05 (bs, 1H, H4$^D$), 4.00 – 3.95 (m, 1H, H5$^C$), 3.92 – 3.55 (m, 41H, H2$^G$, H2$^I$, H2$^K$, H6a$^A$, H6b$^A$, H6a$^E$, H6b$^E$, H6a$^G$, H6b$^G$, H6a$^I$, H6b$^I$, H6a$^K$, H6b$^K$, H5$^E$, H5$^G$, H5$^I$, H5$^K$, H5$^A$, H3$^A$, OCH*H* linker, H3$^E$, H3$^G$, H3$^I$, H3$^K$, H3$^C$, H5$^F$, H5$^H$, H5$^J$, H5$^L$, H4$^C$, H4$^F$, H4$^H$, H4$^J$,

H4^A^, H4^E^, H4^G^, H4^I^, H4^K^, H3^F^, H3^H^, H3^J^), 3.54 – 3.47 (m, 3H, OCH*H* linker, H4^L^, H3^L^), 3.37 – 3.30 (m, 4H, H2^F^, H2^H^, H2^J^, H2^L^), 3.28 – 3.19 (m, 3H, H2^C^, H2^E^, H2^A^), 3.02 (t, *J* = 7.3 Hz, 2H, C*H*$_2$N linker), 2.01 (s, 9H, 3 × NHCOC*H*$_3$), 1.76 – 1.47 (m, 6H, 3 × C*H*$_2$ linker). $^{13}$C NMR from HSQC (151 MHz, D$_2$O) δ 102.3 (C1^F^, C1^H^, C1^J^, C1^L^), 99.2 (C1^D^), 97.2 (C1^B^), 97.0 (C1^A^), 96.7 (C1^K^, C1^E^, C1^G^, C1^I^), 96.0 (C1^C^), 78.0 (C3^F^, C3^H^, C3^J^), 77.0 (C2^B^), 76.4 (C2^D^, C5^F^, C5^H^, C5^J^, C5^L^, C5^N^, C4^F^, C4^H^, C4^J^, C4^C^, C4^G^, C4^I^, C4^A^, C4^E^, C4^K^), 75.8 (C4^B^, C4^D^), 75.0 (C3^L^), 73.4 (C2^F^, C2^H^, C2^J^, C2^L^), 71.6 (C4^L^), 70.5 (C5^G^, C5^I^, C5^K^, C3^G^, C3^I^, C3^K^), 70.1 (C5^B^), 69.6 (C3^B^), 69.5 (C5^D^, C3^D^, C5^C^, C5^E^, C5^A^, C3^C^, C3^E^), 67.7 (OCH$_2$ linker), 66.5 (C6^C^), 59.7 (C6^A^, C6^E^, C6^G^, C6^I^, C6^K^), 57.9 (C2^C^, C2^E^, C2^A^), 53.3 (C2^G^, C2^I^, C2^K^), 39.6 (NCH$_2$ linker), 28.0 (CH$_2$ linker), 26.4 (CH$_2$ linker), 22.6 (CH$_2$ linker), 21.7 (6 × NHCOCH$_3$). HRMS (ESI-MS): m/z calculated for $C_{83}H_{131}N_7O_{82}S_6$ [M-12Na+10H]$^{2-}$: 1364.7315; found: 1364.7205.

$^1$H NMR (600 MHz, D$_2$O)

| | **H1** | **H2** | **H3** | **H4** | **H5** | **H6** |
|---|---|---|---|---|---|---|
| **A** | 5.11 (d, *J* = 3.5 Hz) | 3.22 | 3.87 | 3.70 | 3.87 | 3.85 |
| **B** | 5.29 | 4.29 | 4.16 | 4.10 | 4.66 | – |
| **C** | 5.43 | 3.26 | 3.59 | 3.76 | 3.98 | 4.37, 4.26 |
| **D** | 5.19 | 4.29 | 4.19 | 4.05 | 4.78 | – |
| **E** | 5.39 | 3.22 | 3.68 | 3.68 | 3.88 | 3.85 |
| **F/H/J** | 4.46 | 3.34 | 3.66 | 3.73 | 3.77 | – |
| **G/I/K** | 5.36 | 3.86 | 3.64 | 3.64 | 3.85 | 3.85 |
| **L** | 4.46 | 3.34 | 3.50 | 3.50 | 3.73 | – |

$^{13}$C NMR from HSQC (151 MHz, D$_2$O)

| | **C1** | **C2** | **C3** | **C4** | **C5** | **C6** |
|---|---|---|---|---|---|---|
| **A** | 97.0 | 57.9 | 68.4 | 76.4 | 69.5 | 59.7 |
| **B** | 97.2 | 77.0 | 69.6 | 75.8 | 70.1 | – |
| **C** | 96.0 | 57.9 | 69.5 | 76.4 | 69.5 | 66.5 |

| | | | | | | |
|---|---|---|---|---|---|---|
| **D** | 99.2 | 76.4 | 69.5 | 75.8 | 69.5 | – |
| **E** | 96.7 | 57.9 | 69.5 | 76.4 | 69.5 | 59.7 |
| **F/H/J** | 102.3 | 73.4 | 78.3 | 76.4 | 76.4 | – |
| **G/I/K** | 96.7 | 53.3 | 70.5 | 76.4 | 70.5 | 59.7 |
| **L** | 102.3 | 73.4 | 75.0 | 71.6 | 76.4 | – |

**5-Aminopentyl *O*-(2-acetamido-6-azido-2-deoxy-α-D-glucopyranosyl)[-(1→4)-*O*-(β-D-glucopyranosyluronate)-(1→4)-*O*-(2-acetamido-2-deoxy-α-D-glucopyranosyl)]$_3$-(1→4)-*O*-(β-D-glucopyranosyluronate)-(1→4)-*O*-(2-sulfamino-2-deoxy-α-D-glucopyranosyl)-(1→4)-*O*-(2-*O*-sulfate-α-L-idopyranosyluronate)-(1→4)-*O*-(2-sulfamino-6-*O*-sulfate-2-deoxy-α-D-glucopyranosyl)-(1→4)-*O*-(2-*O*-sulfate-α-L-idopyranosyluronate)-(1→4)-*O*-2-sulfamino-2-deoxy-α-D-glucopyranoside, sodium salt (6).**

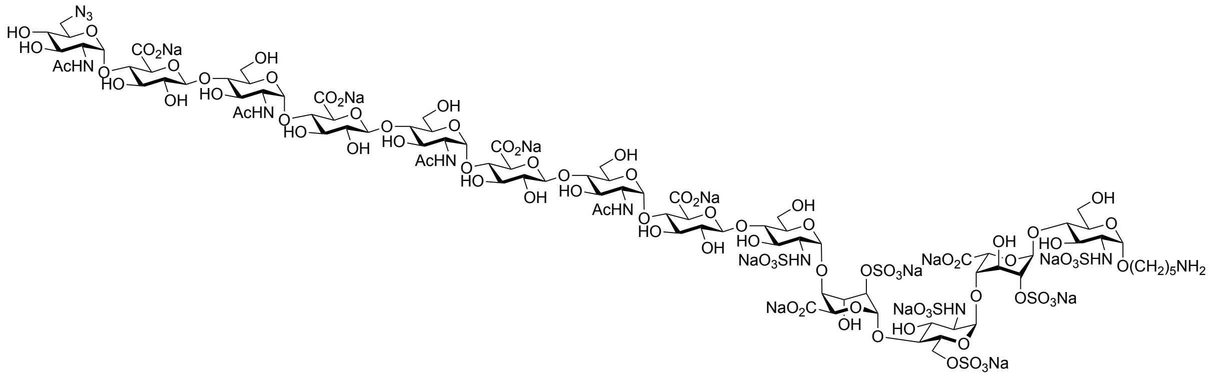

Compound **25** (1.2 mg, 0.40 µmol) was subjected to installation of unnatural α (1→4) 6-azido-GlcNAc according to the general procedure to give the title compound **6** as a white powder (1.1 mg, 85%). $^{1}$H NMR (600 MHz, $D_2O$) δ 5.43 (d, *J* = 3.4 Hz, 1H, H1$^{C}$), 5.40 (d, *J* = 3.8 Hz, 1H, H1$^{M}$), 5.39 (d, *J* = 3.4 Hz, 1H, H1$^{E}$), 5.36 (d, *J* = 3.5 Hz, 3H, H1$^{G}$, H1$^{I}$, H1$^{K}$), 5.30 (d, *J* = 3.0 Hz, 1H, H1$^{B}$), 5.18 (bs, 1H, H1$^{D}$), 5.11 (d, *J* = 3.4 Hz, 1H, H1$^{A}$), 4.78 (1H, H5$^{D}$), 4.66 (bs, 1H, H5$^{B}$), 4.46 (d, *J* = 7.8 Hz, 4H, H1$^{F}$, H1$^{H}$, H1$^{J}$, H1$^{L}$), 4.40 – 4.34 (m, 1H, H6a$^{C}$), 4.31 – 4.27 (m, 2H, H2$^{D}$, H2$^{B}$), 4.27 – 4.23 (m, 1H, H6b$^{C}$), 4.20 – 4.17 (m, 1H, H3$^{D}$), 4.17 – 4.14 (m, 1H, H3$^{B}$ ), 4.11 – 4.08 (m, 1H, H4$^{B}$), 4.05 (bs, 1H, H4$^{D}$), 4.00 – 3.96 (m, 1H, H5$^{C}$), 3.92 – 3.56 (m, 48H, H2$^{G}$, H2$^{I}$, H2$^{K}$, H2$^{M}$, H6a$^{A}$, H6b$^{A}$, H6a$^{E}$, H6b$^{E}$, H6a$^{G}$, H6b$^{G}$, H6a$^{I}$, H6b$^{I}$, H6a$^{K}$, H6b$^{K}$, H6a$^{M}$, H6b$^{M}$, H5$^{E}$, H5$^{G}$, H5$^{I}$, H5$^{K}$, H5$^{M}$, H5$^{A}$, H3$^{A}$, OCH*H* linker, H3$^{E}$, H3$^{G}$, H3$^{I}$, H3$^{K}$, H3$^{M}$, H3$^{C}$, H5$^{F}$, H5$^{H}$, H5$^{J}$, H5$^{L}$, H4$^{C}$, H4$^{F}$, H4$^{H}$, H4$^{J}$, H4$^{L}$, H4$^{A}$, H4$^{E}$, H4$^{G}$, H4$^{I}$, H4$^{K}$, H3$^{F}$, H3$^{H}$, H3$^{J}$,

H3$^{L}$), 3.54 – 3.50 (m, 1H, OCH*H* linker), 3.48 – 3.43 (m, 1H, H4$^{M}$), 3.37 – 3.30 (m, 4H, H2$^{F}$, H2$^{H}$, H2$^{J}$, H2$^{L}$), 3.26 (dd, *J* = 10.3, 3.4 Hz, 1H, H2$^{C}$), 3.22 (dd, *J* = 10.3, 3.4 Hz, 2H, H2$^{E}$, H2$^{A}$), 3.02 (t, *J* = 7.3 Hz, 2H, C*H*$_2$N linker), 2.03 (s, 3H, NHCOC*H*$_3$$^{M}$), 2.01 (s, 9H, 3 × NHCOC*H*$_3$), 1.79 – 1.43 (m, 6H, 3 × C*H*$_2$ linker). $^{13}$C NMR from HSQC (151 MHz, $D_2O$) δ 102.3 (C1$^{F}$, C1$^{H}$, C1$^{J}$, C1$^{L}$), 99.2 (C1$^{D}$), 97.2 (C1$^{B}$), 97.0 (C1$^{A}$), 96.7 (C1$^{K}$, C1$^{M}$, C1$^{E}$, C1$^{G}$, C1$^{I}$), 96.0 (C1$^{C}$), 78.0 (C3$^{F}$, C3$^{H}$, C3$^{J}$, C3$^{L}$), 77.0 (C2$^{B}$), 76.4 (C2$^{D}$, C5$^{F}$, C5$^{H}$, C5$^{J}$, C5$^{L}$, C4$^{F}$, C4$^{H}$, C4$^{J}$, C4$^{L}$, C4$^{C}$, C4$^{G}$, C4$^{I}$, C4$^{A}$, C4$^{E}$, C4$^{K}$), 75.8 (C4$^{B}$, C4$^{D}$), 73.4 (C2$^{F}$, C2$^{H}$, C2$^{J}$, C2$^{L}$), 70.5 (C5$^{G}$, C5$^{I}$, C5$^{K}$, C5$^{M}$, C3$^{G}$, C3$^{I}$, C3$^{K}$, C3$^{M}$), 70.1 (C5$^{B}$, C4$^{M}$), 69.6 (C3$^{B}$), 69.5 (C5$^{D}$, C3$^{D}$, C5$^{C}$, C5$^{E}$, C5$^{A}$, C3$^{C}$, C3$^{E}$), 67.7 (O<u>C</u>H$_2$ linker), 66.5 (C6$^{C}$), 59.7 (C6$^{A}$, C6$^{E}$, C6$^{G}$, C6$^{I}$, C6$^{K}$), 57.9 (C2$^{C}$, C2$^{E}$, C2$^{A}$), 53.3 (C2$^{G}$, C2$^{I}$, C2$^{K}$, C2$^{M}$), 50.4 (C6$^{M}$), 39.6 (N<u>C</u>H$_2$ linker), 28.0 (<u>C</u>H$_2$ linker), 26.4 (<u>C</u>H$_2$ linker), 22.6 (<u>C</u>H$_2$ linker), 21.7 (4 × NHCO<u>C</u>H$_3$). HRMS (ESI-MS): m/z calculated for $C_{91}H_{142}N_{11}O_{86}S_6$ [M-12Na+9H]$^{3-}$: 985.5139; found: 985.5163.

$^{1}$H NMR (600 MHz, $D_2O$)

| | **H1** | **H2** | **H3** | **H4** | **H5** | **H6** |
|---|---|---|---|---|---|---|
| **A** | 5.11 (d, *J* = 3.4 Hz) | 3.22 (dd, *J* = 10.2, 3.4 Hz) | 3.87 | 3.70 | 3.87 | 3.85 |
| **B** | 5.30 (d, *J* = 3.0 Hz) | 4.29 | 4.16 | 4.10 | 4.66 | – |
| **C** | 5.43 (d, *J* = 3.4 Hz) | 3.26 (dd, *J* = 10.3, 3.4 Hz) | 3.59 | 3.76 | 3.98 | 4.37, 4.26 |
| **D** | 5.18 | 4.29 | 4.19 | 4.05 | 4.78 | – |
| **E** | 5.39 (d, *J* = 3.4 Hz) | 3.22 (dd, *J* = 10.2, 3.4 Hz) | 3.68 | 3.68 | 3.88 | 3.85 |
| **F/H/J/L** | 4.46 (d, *J* = 7.8 Hz) | 3.34 | 3.66 | 3.73 | 3.77 | – |
| **G/I/K** | 5.36 (d, *J* = 3.5 Hz) | 3.86 | 3.64 | 3.64 | 3.85 | 3.85 |
| **M** | 5.40 (d, *J* = 3.8 Hz) | 3.85 | 3.68 | 3.47 | 3.88 | 3.62 |

$^{13}$C NMR from HSQC (151 MHz, $D_2O$)

| | **C1** | **C2** | **C3** | **C4** | **C5** | **C6** |
|---|---|---|---|---|---|---|

| | | | | | | |
|---|---|---|---|---|---|---|
| **A** | 97.0 | 57.9 | 68.4 | 76.4 | 69.5 | 59.7 |
| **B** | 97.2 | 77.0 | 69.6 | 75.8 | 70.1 | – |
| **C** | 96.0 | 57.9 | 69.5 | 76.4 | 69.5 | 66.5 |
| **D** | 99.2 | 76.4 | 69.5 | 75.8 | 69.5 | – |
| **E** | 96.7 | 57.9 | 69.5 | 76.4 | 69.5 | 59.7 |
| **F/H/J/L** | 102.3 | 73.4 | 78.3 | 76.4 | 76.4 | – |
| **G/I/K** | 96.7 | 53.3 | 70.5 | 76.4 | 70.5 | 59.7 |
| **M** | 96.7 | 53.3 | 70.5 | 70.1 | 70.5 | 50.4 |

**5-Aminopentyl [*O*-(β-D-glucopyranosyluronate)-(1→4)-*O*-(2-acetamido-2-deoxy-α-D-**

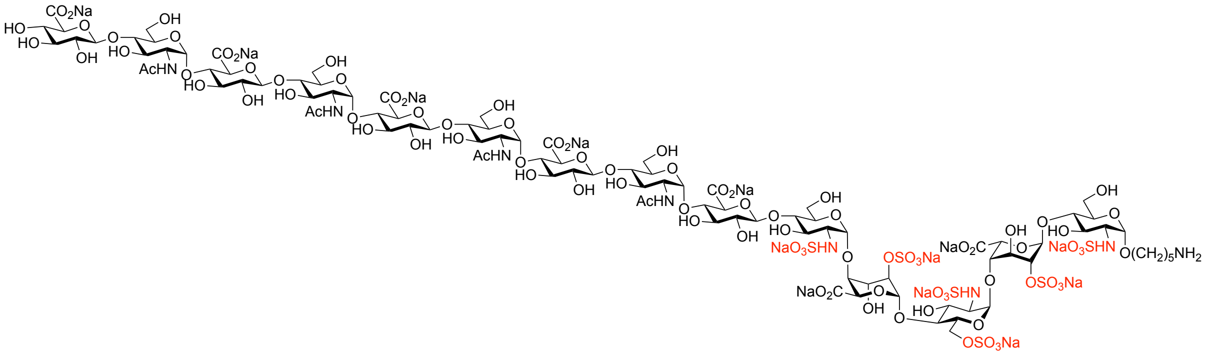

**glucopyranosyl)-(1→4)]$_4$-*O*-(β-D-glucopyranosyluronate)-(1→4)-*O*-(2-sulfamino-2-deoxy-α-D-glucopyranosyl)-(1→4)-*O*-(2-*O*-sulfate-α-L-idopyranosyluronate)-(1→4)-*O*-(2-sulfamino-6-*O*-sulfate-2-deoxy-α-D-glucopyranosyl)-(1→4)-*O*-(2-*O*-sulfate-α-L-idopyranosyluronate)-(1→4)-*O*-2-sulfamino-2-deoxy-α-D-glucopyranoside, sodium salt (26).**

Compound **26** was obtained from compound **25** as a white powder according to the general procedure for installation of α (1→4)-GlcNAc and β (1→4)-GlcA, respectively. $^1$H NMR (600 MHz, D$_2$O) δ 5.43 (d, *J* = 3.4 Hz, 1H, H1$^C$), 5.40 – 5.33 (m, 5H, H1$^E$, H1$^G$, H1$^I$, H1$^K$, H1$^M$), 5.27 (d, *J* = 3.4 Hz, 1H, H1$^B$), 5.19 (bs, 1H, H1$^D$), 5.11 (d, *J* = 3.5 Hz, 1H, H1$^A$), 4.78 (1H, H5$^D$), 4.65 (d, *J* = 2.9 Hz, 1H, H5$^B$), 4.50 – 4.43 (m, 5H, H1$^F$, H1$^H$, H1$^J$, H1$^L$, H1$^N$), 4.40 – 4.34 (m, 1H, H6a$^C$), 4.31 – 4.27 (m, 2H, H2$^D$, H2$^B$), 4.26 – 4.22 (m, 1H, H6b$^C$), 4.20 – 4.17 (m, 1H, H3$^D$), 4.17 – 4.14 (m, 1H, H3$^B$ ), 4.11 – 4.08 (m, 1H, H4$^B$), 4.05 (bs, 1H, H4$^D$), 4.00 – 3.95 (m, 1H, H5$^C$), 3.92 – 3.55 (m, 50H, H2$^G$, H2$^I$, H2$^K$, H2$^M$, H6a$^A$, H6b$^A$, H6a$^E$, H6b$^E$, H6a$^G$, H6b$^G$,

H6a$^{I}$, H6b$^{I}$, H6a$^{K}$, H6b$^{K}$, H6a$^{M}$, H6b$^{M}$, H5$^{E}$, H5$^{G}$, H5$^{I}$, H5$^{K}$, H5$^{M}$, H5$^{A}$, H3$^{A}$, OCH*H* linker, H3$^{E}$, H3$^{G}$, H3$^{I}$, H3$^{K}$, H3$^{M}$, H3$^{C}$, H5$^{F}$, H5$^{H}$, H5$^{J}$, H5$^{L}$, H5$^{N}$, H4$^{C}$, H4$^{F}$, H4$^{H}$, H4$^{J}$, H4$^{L}$, H4$^{A}$, H4$^{E}$, H4$^{G}$, H4$^{I}$, H4$^{K}$, H4$^{M}$, H3$^{F}$, H3$^{H}$, H3$^{J}$, H3$^{L}$), 3.54 – 3.47 (m, 3H, OCH*H* linker, H4$^{N}$, H3$^{N}$), 3.37 – 3.30 (m, 5H, H2$^{F}$, H2$^{H}$, H2$^{J}$, H2$^{L}$, H2$^{N}$), 3.28 – 3.19 (m, 3H, H2$^{C}$, H2$^{E}$, H2$^{A}$), 3.02 (t, $J$ = 7.3 Hz, 2H, C$H_2$N linker), 2.02 & 2.01 (2s, 12H, 4 × NHCOC$H_3$), 1.76 – 1.47 (m, 6H, 3 × C$H_2$ linker). $^{13}$C NMR from HSQC (151 MHz, $D_2O$) δ 102.3 (C1$^{F}$, C1$^{H}$, C1$^{J}$, C1$^{L}$, C1$^{N}$), 99.2 (C1$^{D}$), 97.2 (C1$^{B}$), 97.0 (C1$^{A}$), 96.7 (C1$^{K}$, C1$^{M}$, C1$^{E}$, C1$^{G}$, C1$^{I}$), 96.0 (C1$^{C}$), 78.0 (C3$^{F}$, C3$^{H}$, C3$^{J}$, C3$^{L}$), 77.0 (C2$^{B}$), 76.4 (C2$^{D}$, C5$^{F}$, C5$^{H}$, C5$^{J}$, C5$^{L}$, C5$^{N}$, C4$^{F}$, C4$^{H}$, C4$^{J}$, C4$^{L}$, C4$^{C}$, C4$^{G}$, C4$^{I}$, C4$^{A}$, C4$^{E}$, C4$^{K}$, C4$^{M}$), 75.8 (C4$^{B}$, C4$^{D}$), 75.0 (C3$^{N}$), 73.4 (C2$^{F}$, C2$^{H}$, C2$^{J}$, C2$^{L}$, C2$^{N}$), 71.6 (C4$^{N}$), 70.5 (C5$^{G}$, C5$^{I}$, C5$^{K}$, C5$^{M}$, C3$^{G}$, C3$^{I}$, C3$^{K}$, C3$^{M}$), 70.1 (C5$^{B}$), 69.6 (C3$^{B}$), 69.5 (C5$^{D}$, C3$^{D}$, C5$^{C}$, C5$^{E}$, C5$^{A}$, C3$^{C}$, C3$^{E}$), 67.7 ($O\underline{C}H_2$ linker), 66.5 (C6$^{C}$), 59.7 (C6$^{A}$, C6$^{E}$, C6$^{G}$, C6$^{I}$, C6$^{K}$, C6$^{M}$), 57.9 (C2$^{C}$, C2$^{E}$, C2$^{A}$), 53.3 (C2$^{G}$, C2$^{I}$, C2$^{K}$, C2$^{M}$), 39.6 ($N\underline{C}H_2$ linker), 28.0 ($\underline{C}H_2$ linker), 26.4 ($\underline{C}H_2$ linker), 22.6 ($\underline{C}H_2$ linker), 21.7 (6 × NHCO$\underline{C}H_3$). ESI-MS: m/z calculated for $C_{97}H_{151}N_8O_{93}S_6$ [M-13Na+10H]$^{3-}$: 1036.1902; found: 1036.1767.

$^{1}$H NMR (600 MHz, $D_2O$)

| | **H1** | **H2** | **H3** | **H4** | **H5** | **H6** |
|---|---|---|---|---|---|---|
| **A** | 5.11 (d, $J$ = 3.5 Hz) | 3.22 | 3.87 | 3.70 | 3.87 | 3.85 |
| **B** | 5.27 (d, $J$ = 3.4 Hz) | 4.29 | 4.16 | 4.10 | 4.66 | – |
| **C** | 5.43 (d, $J$ = 3.4 Hz) | 3.26 | 3.59 | 3.76 | 3.98 | 4.37, 4.26 |
| **D** | 5.19 | 4.29 | 4.19 | 4.05 | 4.78 | – |
| **E** | 5.39 | 3.22 | 3.68 | 3.68 | 3.88 | 3.85 |
| **F/H/J/L** | 4.46 | 3.34 | 3.66 | 3.73 | 3.77 | – |
| **G/I/K/M** | 5.36 | 3.86 | 3.64 | 3.64 | 3.85 | 3.85 |
| **N** | 4.46 | 3.34 | 3.50 | 3.50 | 3.73 | – |

$^{13}$C NMR from HSQC (151 MHz, $D_2O$)

| | C1 | C2 | C3 | C4 | C5 | C6 |
|---|---|---|---|---|---|---|
| **A** | 97.0 | 57.9 | 68.4 | 76.4 | 69.5 | 59.7 |
| **B** | 97.2 | 77.0 | 69.6 | 75.8 | 70.1 | – |
| **C** | 96.0 | 57.9 | 69.5 | 76.4 | 69.5 | 66.5 |
| **D** | 99.2 | 76.4 | 69.5 | 75.8 | 69.5 | – |
| **E** | 96.7 | 57.9 | 69.5 | 76.4 | 69.5 | 59.7 |
| **F/H/J/L** | 102.3 | 73.4 | 78.3 | 76.4 | 76.4 | – |
| **G/I/K/M** | 96.7 | 53.3 | 70.5 | 76.4 | 70.5 | 59.7 |
| **N** | 102.3 | 73.4 | 75.0 | 71.6 | 76.4 | – |

**5-Aminopentyl *O*-(2-acetamido-6-azido-2-deoxy-α-D-glucopyranosyl)[-(1→4)-*O*-(β-D-glucopyranosyluronate)-(1→4)-*O*-(2-acetamido-2-deoxy-α-D-glucopyranosyl)]$_4$-(1→4)-*O*-(β-D-glucopyranosyluronate)-(1→4)-*O*-(2-sulfamino-2-deoxy-α-D-glucopyranosyl)-(1→4)-*O*-(2-*O*-sulfate-α-L-idopyranosyluronate)-(1→4)-*O*-(2-sulfamino-6-*O*-sulfate-2-deoxy-α-D-glucopyranosyl)-(1→4)-*O*-(2-*O*-sulfate-α-L-idopyranosyluronate)-(1→4)-*O*-2-sulfamino-2-deoxy-α-D-glucopyranoside, sodium salt (7).**

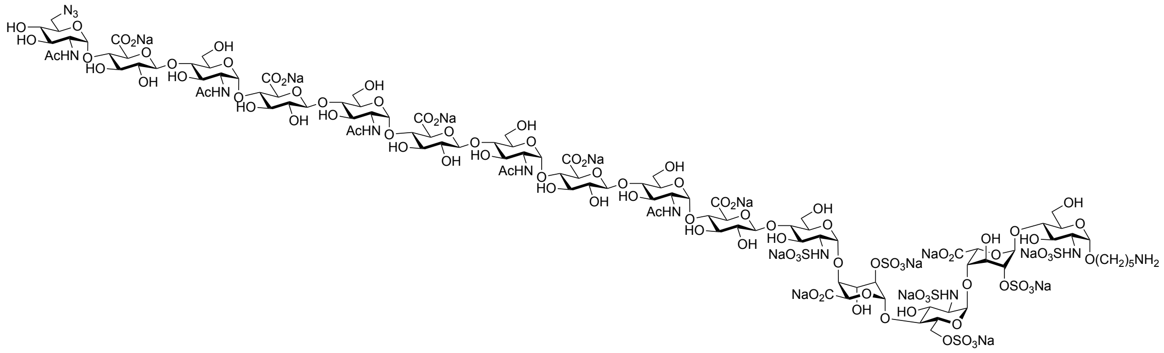

Compound **26** (1.4 mg, 0.41 μmol) was subjected to installation of unnatural α (1→4) 6-azido-GlcNAc according to the general procedure to give the title compound **7** as a white powder (1.2 mg, 80%). $^1$H NMR (600 MHz, $D_2O$) δ 5.44 (d, $J$ = 3.4 Hz, 1H, H1$^C$), 5.42 – 5.38 (m, 2H, H1$^O$, H1$^E$), 5.36 (d, $J$ = 3.2 Hz, 4H, H1$^G$, H1$^I$, H1$^K$, H1$^M$), 5.30 (bs, 1H, H1$^B$), 5.18 (bs, 1H, H1$^D$), 5.11 (d, $J$ = 3.4 Hz, 1H, H1$^A$), 4.78 (1H, H5$^D$), 4.66 (bs, 1H, H5$^B$), 4.46 (d, $J$ = 7.8 Hz, 5H, H1$^F$, H1$^H$, H1$^J$, H1$^L$, H1$^N$), 4.40 – 4.34 (m, 1H, H6a$^C$), 4.31 – 4.27 (m, 2H, H2$^D$, H2$^B$), 4.27 – 4.23 (m, 1H, H6b$^C$), 4.20 – 4.17 (m, 1H, H3$^D$), 4.17 – 4.14 (m, 1H, H3$^B$ ), 4.11 – 4.08 (m,

1H, H4$^{B}$), 4.05 (bs, 1H, H4$^{D}$), 4.00 – 3.95 (m, 1H, H5$^{C}$), 3.92 – 3.55 (m, 57H, H2$^{G}$, H2$^{I}$, H2$^{K}$, H2$^{M}$, H2$^{O}$, H6a$^{A}$, H6b$^{A}$, H6a$^{E}$, H6b$^{E}$, H6a$^{G}$, H6b$^{G}$, H6a$^{I}$, H6b$^{I}$, H6a$^{K}$, H6b$^{K}$, H6a$^{M}$, H6b$^{M}$, H6a$^{O}$, H6b$^{O}$, H5$^{E}$, H5$^{G}$, H5$^{I}$, H5$^{K}$, H5$^{M}$, H5$^{O}$, H5$^{A}$, H3$^{A}$, OCH*H* linker, H3$^{E}$, H3$^{G}$, H3$^{I}$, H3$^{K}$, H3$^{M}$, H3$^{O}$, H3$^{C}$, H5$^{F}$, H5$^{H}$, H5$^{J}$, H5$^{L}$, H5$^{N}$, H4$^{C}$, H4$^{F}$, H4$^{H}$, H4$^{J}$, H4$^{L}$, H4$^{N}$, H4$^{A}$, H4$^{E}$, H4$^{G}$, H4$^{I}$, H4$^{K}$, H4$^{M}$, H3$^{F}$, H3$^{H}$, H3$^{J}$, H3$^{L}$, H3$^{N}$), 3.54 – 3.49 (m, 1H, OCH*H* linker), 3.49 – 3.43 (m, 1H, H4$^{O}$), 3.37 – 3.30 (m, 5H, H2$^{F}$, H2$^{H}$, H2$^{J}$, H2$^{L}$, H2$^{N}$), 3.28 – 3.19 (m, 3H, H2$^{C}$, H2$^{E}$, H2$^{A}$), 3.02 (t, *J* = 7.3 Hz, 2H, C$H_2$N linker), 2.02 (s, 3H, NHCOC$H_3$$^{O}$), 2.01 (s, 12H, 4 × NHCOC$H_3$), 1.76 – 1.47 (m, 6H, 3 × C$H_2$ linker). $^{13}$C NMR from HSQC (151 MHz, $D_2O$) δ 102.3 (C1$^{F}$, C1$^{H}$, C1$^{J}$, C1$^{L}$, C1$^{N}$), 99.2 (C1$^{D}$), 97.2 (C1$^{B}$), 97.0 (C1$^{A}$), 96.7 (C1$^{O}$, C1$^{K}$, C1$^{M}$, C1$^{E}$, C1$^{G}$, C1$^{I}$), 96.0 (C1$^{C}$), 78.0 (C3$^{F}$, C3$^{H}$, C3$^{J}$, C3$^{L}$, C3$^{N}$), 77.0 (C2$^{B}$), 76.4 (C2$^{D}$, C5$^{F}$, C5$^{H}$, C5$^{J}$, C5$^{L}$, C5$^{N}$, C4$^{F}$, C4$^{H}$, C4$^{J}$, C4$^{L}$, C4$^{N}$, C4$^{C}$, C4$^{G}$, C4$^{I}$, C4$^{A}$, C4$^{E}$, C4$^{K}$, C4$^{M}$), 75.8 (C4$^{B}$, C4$^{D}$), 73.4 (C2$^{F}$, C2$^{H}$, C2$^{J}$, C2$^{L}$, C2$^{N}$), 70.5 (C5$^{G}$, C5$^{I}$, C5$^{K}$, C5$^{M}$, C5$^{O}$, C3$^{G}$, C3$^{I}$, C3$^{K}$, C3$^{M}$, C3$^{O}$), 70.1 (C5$^{B}$, C4$^{O}$), 69.6 (C3$^{B}$), 69.5 (C5$^{D}$, C3$^{D}$, C5$^{C}$, C5$^{E}$, C5$^{A}$, C3$^{C}$, C3$^{E}$), 67.7 (O$\underline{C}H_2$ linker), 66.5 (C6$^{C}$), 59.7 (C6$^{A}$, C6$^{E}$, C6$^{G}$, C6$^{I}$, C6$^{K}$, C6$^{M}$), 57.9 (C2$^{C}$, C2$^{E}$, C2$^{A}$), 53.3 (C2$^{G}$, C2$^{I}$, C2$^{K}$, C2$^{M}$, C2$^{O}$), 50.4 (C6$^{O}$), 39.6 (N$\underline{C}H_2$ linker), 28.0 ($\underline{C}H_2$ linker), 26.4 ($\underline{C}H_2$ linker), 22.6 ($\underline{C}H_2$ linker), 21.7 (5 × NHCO$\underline{C}H_3$). HRMS (ESI-MS): m/z calculated for $C_{105}H_{163}N_{12}O_{97}S_6$ [M-13Na+10H]$^{3-}$: 1111.8844; found: 1111.8806.

$^{1}$H NMR (600 MHz, $D_2O$)

| | H1 | H2 | H3 | H4 | H5 | H6 |
|---|---|---|---|---|---|---|
| **A** | 5.11 (d, *J* = 3.4 Hz) | 3.22 | 3.87 | 3.70 | 3.87 | 3.85 |
| **B** | 5.30 | 4.29 | 4.16 | 4.10 | 4.66 | – |
| **C** | 5.44 (d, *J* = 3.4 Hz) | 3.26 | 3.59 | 3.76 | 3.98 | 4.37, 4.26 |
| **D** | 5.18 | 4.29 | 4.19 | 4.05 | 4.78 | – |
| **E** | 5.39 | 3.22 | 3.68 | 3.68 | 3.88 | 3.85 |
| **F/H/J/L/N** | 4.46 (d, *J* = 7.8 Hz) | 3.34 | 3.66 | 3.73 | 3.77 | – |
| **G/I/K/M** | 5.36 (d, *J* = 3.2 Hz) | 3.86 | 3.64 | 3.64 | 3.85 | 3.85 |
| **O** | 5.40 | 3.85 | 3.68 | 3.47 | 3.88 | 3.62 |

$^{13}C$ NMR from HSQC (151 MHz, $D_2O$)

| | **C1** | **C2** | **C3** | **C4** | **C5** | **C6** |
|---|---|---|---|---|---|---|
| **A** | 97.0 | 57.9 | 68.4 | 76.4 | 69.5 | 59.7 |
| **B** | 97.2 | 77.0 | 69.6 | 75.8 | 70.1 | – |
| **C** | 96.0 | 57.9 | 69.5 | 76.4 | 69.5 | 66.5 |
| **D** | 99.2 | 76.4 | 69.5 | 75.8 | 69.5 | – |
| **E** | 96.7 | 57.9 | 69.5 | 76.4 | 69.5 | 59.7 |
| **F/H/J/L/N** | 102.3 | 73.4 | 78.3 | 76.4 | 76.4 | – |
| **G/I/K/M** | 96.7 | 53.3 | 70.5 | 76.4 | 70.5 | 59.7 |
| **O** | 96.7 | 53.3 | 70.5 | 70.1 | 70.5 | 50.4 |

**5-Aminopentyl [*O*-(β-D-glucopyranosyluronate)-(1→4)-*O*-(2-acetamido-2-deoxy-α-D-glucopyranosyl)-(1→4)]$_5$-*O*-(β-D-glucopyranosyluronate)-(1→4)-*O*-(2-sulfamino-2-deoxy-α-D-glucopyranosyl)-(1→4)-*O*-(2-*O*-sulfate-α-L-idopyranosyluronate)-(1→4)-*O*-(2-sulfamino-6-*O*-sulfate-2-deoxy-α-D-glucopyranosyl)-(1→4)-*O*-(2-*O*-sulfate-α-L-idopyranosyluronate)-(1→4)-*O*-2-sulfamino-2-deoxy-α-D-glucopyranoside, sodium salt (27).**

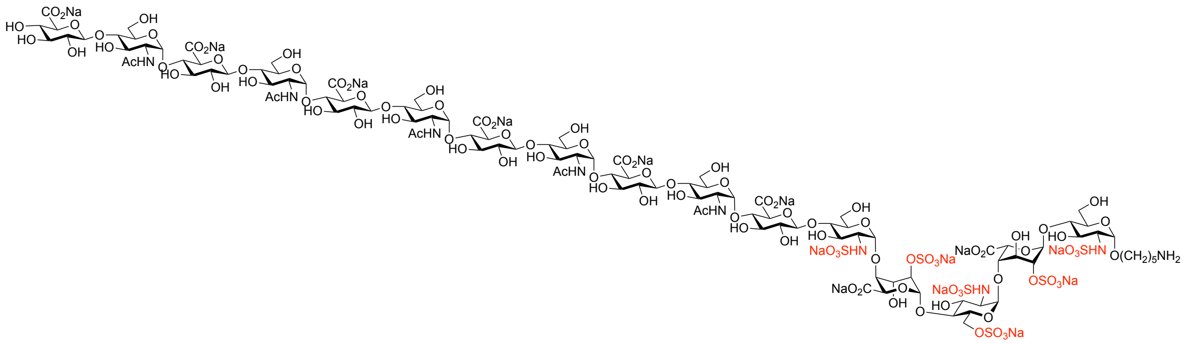

Compound **27** was obtained as a white powder from compound **26** according to the general procedure for installation of α (1→4)-GlcNAc and β (1→4)-GlcA, respectively. $^1$H NMR (600 MHz, $D_2O$) δ 5.43 (d, $J$ = 3.4 Hz, 1H, H1$^C$), 5.40 – 5.33 (m, 6H, H1$^E$, H1$^G$, H1$^I$, H1$^K$, H1$^M$, H1$^O$), 5.29 (d, $J$ = 3.4 Hz, 1H, H1$^B$), 5.18 (bs, 1H, H1$^D$), 5.11 (d, $J$ = 3.5 Hz, 1H, H1$^A$), 4.78 (1H, H5$^D$), 4.65 (d, $J$ = 2.9 Hz, 1H, H5$^B$), 4.50 – 4.43 (m, 6H, H1$^F$, H1$^H$, H1$^J$, H1$^L$, H1$^N$, H1$^P$), 4.40 – 4.34 (m, 1H, H6a$^C$), 4.31 – 4.27 (m, 2H, H2$^D$, H2$^B$), 4.26 – 4.22 (m, 1H, H6b$^C$), 4.20 –

4.17 (m, 1H, H3$^{D}$), 4.17 – 4.14 (m, 1H, H3$^{B}$ ), 4.11 – 4.08 (m, 1H, H4$^{B}$), 4.05 (bs, 1H, H4$^{D}$), 4.00 – 3.95 (m, 1H, H5$^{C}$), 3.92 – 3.55 (m, 59H, H2$^{G}$, H2$^{I}$, H2$^{K}$, H2$^{M}$, H2$^{O}$, H6a$^{A}$, H6b$^{A}$, H6a$^{E}$, H6b$^{E}$, H6a$^{G}$, H6b$^{G}$, H6a$^{I}$, H6b$^{I}$, H6a$^{K}$, H6b$^{K}$, H6a$^{M}$, H6b$^{M}$, H6a$^{O}$, H6b$^{O}$, H5$^{E}$, H5$^{G}$, H5$^{I}$, H5$^{K}$, H5$^{M}$, H5$^{O}$, H5$^{A}$, H3$^{A}$, OCH*H* linker, H3$^{E}$, H3$^{G}$, H3$^{I}$, H3$^{K}$, H3$^{M}$, H3$^{O}$, H3$^{C}$, H5$^{F}$, H5$^{H}$, H5$^{J}$, H5$^{L}$, H5$^{N}$, H5$^{P}$, H4$^{C}$, H4$^{F}$, H4$^{H}$, H4$^{J}$, H4$^{L}$, H4$^{N}$, H4$^{A}$, H4$^{E}$, H4$^{G}$, H4$^{I}$, H4$^{K}$, H4$^{M}$, H4$^{O}$, H3$^{F}$, H3$^{H}$, H3$^{J}$, H3$^{L}$, H3$^{N}$), 3.54 – 3.47 (m, 3H, OCH*H* linker, H4$^{P}$, H3$^{P}$), 3.37 – 3.30 (m, 6H, H2$^{F}$, H2$^{H}$, H2$^{J}$, H2$^{L}$, H2$^{N}$, H2$^{P}$), 3.28 – 3.19 (m, 3H, H2$^{C}$, H2$^{E}$, H2$^{A}$), 3.02 (t, $J$ = 7.3 Hz, 2H, C$H_2$N linker), 2.02 & 2.01 (2s, 15H, 5 × NHCOC$H_3$), 1.76 – 1.47 (m, 6H, 3 × C$H_2$ linker). $^{13}$C NMR from HSQC (151 MHz, $D_2O$) δ 102.3 (C1$^{F}$, C1$^{H}$, C1$^{J}$, C1$^{L}$, C1$^{N}$, C1$^{P}$), 99.2 (C1$^{D}$), 97.2 (C1$^{B}$), 97.0 (C1$^{A}$), 96.7 (C1$^{O}$, C1$^{K}$, C1$^{M}$, C1$^{E}$, C1$^{G}$, C1$^{I}$), 96.0 (C1$^{C}$), 78.0 (C3$^{F}$, C3$^{H}$, C3$^{J}$, C3$^{L}$, C3$^{N}$), 77.0 (C2$^{B}$), 76.4 (C2$^{D}$, C5$^{F}$, C5$^{H}$, C5$^{J}$, C5$^{L}$, C5$^{N}$, C5$^{P}$, C4$^{F}$, C4$^{H}$, C4$^{J}$, C4$^{L}$, C4$^{N}$, C4$^{C}$, C4$^{G}$, C4$^{I}$, C4$^{A}$, C4$^{E}$, C4$^{K}$, C4$^{M}$, C4$^{O}$), 75.8 (C4$^{B}$, C4$^{D}$), 75.0 (C3$^{P}$), 73.4 (C2$^{F}$, C2$^{H}$, C2$^{J}$, C2$^{L}$, C2$^{N}$, C2$^{P}$), 71.6 (C4$^{P}$), 70.5 (C5$^{G}$, C5$^{I}$, C5$^{K}$, C5$^{M}$, C5$^{O}$, C3$^{G}$, C3$^{I}$, C3$^{K}$, C3$^{M}$, C3$^{O}$), 70.1 (C5$^{B}$), 69.6 (C3$^{B}$), 69.5 (C5$^{D}$, C3$^{D}$, C5$^{C}$, C5$^{E}$, C5$^{A}$, C3$^{C}$, C3$^{E}$), 67.7 (O<u>C</u>H2 linker), 66.5 (C6$^{C}$), 59.7 (C6$^{A}$, C6$^{E}$, C6$^{G}$, C6$^{I}$, C6$^{K}$, C6$^{M}$, C6$^{O}$), 57.9 (C2$^{C}$, C2$^{E}$, C2$^{A}$), 53.3 (C2$^{G}$, C2$^{I}$, C2$^{K}$, C2$^{M}$, C2$^{O}$), 39.6 (N<u>C</u>H2 linker), 28.0 (<u>C</u>H2 linker), 26.4 (<u>C</u>H2 linker), 22.6 (<u>C</u>H2 linker), 21.7 (6 × NHCO<u>C</u>H3). ESI-MS: m/z calculated for $C_{111}H_{172}N_9O_{104}S_6$ [M-14Na+11H]$^{3-}$: 1162.5607; found: 1162.5547.

$^1$H NMR (600 MHz, $D_2O$)

| | **H1** | **H2** | **H3** | **H4** | **H5** | **H6** |
|---|---|---|---|---|---|---|
| **A** | 5.11 (d, $J$ = 3.5 Hz) | 3.22 | 3.87 | 3.70 | 3.87 | 3.85 |
| **B** | 5.29 (d, $J$ = 3.4 Hz) | 4.29 | 4.16 | 4.10 | 4.66 | – |
| **C** | 5.43 (d, $J$ = 3.4 Hz) | 3.26 | 3.59 | 3.76 | 3.98 | 4.37, 4.26 |
| **D** | 5.18 | 4.29 | 4.19 | 4.05 | 4.78 | – |
| **E** | 5.39 | 3.22 | 3.68 | 3.68 | 3.88 | 3.85 |
| **F/H/J/L/N** | 4.46 | 3.34 | 3.66 | 3.73 | 3.77 | – |
| **G/I/K/M/O** | 5.36 | 3.86 | 3.64 | 3.64 | 3.85 | 3.85 |
| **P** | 4.46 | 3.34 | 3.50 | 3.50 | 3.73 | – |

$^{13}C$ NMR from HSQC (151 MHz, $D_2O$)

| | C1 | C2 | C3 | C4 | C5 | C6 |
|---|---|---|---|---|---|---|
| **A** | 97.0 | 57.9 | 68.4 | 76.4 | 69.5 | 59.7 |
| **B** | 97.2 | 77.0 | 69.6 | 75.8 | 70.1 | – |
| **C** | 96.0 | 57.9 | 69.5 | 76.4 | 69.5 | 66.5 |
| **D** | 99.2 | 76.4 | 69.5 | 75.8 | 69.5 | – |
| **E** | 96.7 | 57.9 | 69.5 | 76.4 | 69.5 | 59.7 |
| **F/H/J/L/N** | 102.3 | 73.4 | 78.3 | 76.4 | 76.4 | – |
| **G/I/K/M/O** | 96.7 | 53.3 | 70.5 | 76.4 | 70.5 | 59.7 |
| **P** | 102.3 | 73.4 | 75.0 | 71.6 | 76.4 | – |

**5-Aminopentyl *O*-(2-acetamido-6-azido-2-deoxy-α-D-glucopyranosyl)[-(1→4)-*O*-(β-D-glucopyranosyluronate)-(1→4)-*O*-(2-acetamido-2-deoxy-α-D-glucopyranosyl)]$_5$-(1→4)-*O*-(β-D-glucopyranosyluronate)-(1→4)-*O*-(2-sulfamino-2-deoxy-α-D-glucopyranosyl)-(1→4)-*O*-(2-*O*-sulfate-α-L-idopyranosyluronate)-(1→4)-*O*-(2-sulfamino-6-*O*-sulfate-2-deoxy-α-D-glucopyranosyl)-(1→4)-*O*-(2-*O*-sulfate-α-L-idopyranosyluronate)-(1→4)-*O*-2-sulfamino-2-deoxy-α-D-glucopyranoside, sodium salt (8).**

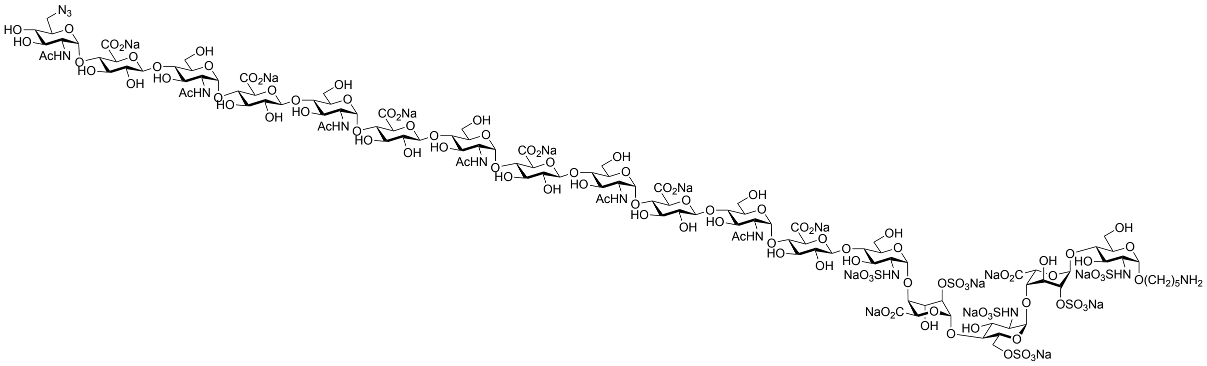

Compound **27** (0.6 mg, 0.16 μmol) was subjected to installation of unnatural α (1→4) 6-azido-GlcNAc according to the general procedure to give the title compound **8** as a white powder (0.5 mg, 79%). $^{1}H$ NMR (600 MHz, $D_2O$) δ 5.44 (d, $J$ = 3.4 Hz, 1H, H1$^{C}$), 5.40 (d, $J$ = 3.8 Hz, 1H, H1$^{Q}$), 5.39 (d, $J$ = 3.4 Hz, 1H, H1$^{E}$), 5.36 (d, $J$ = 3.2 Hz, 5H, H1$^{G}$, H1$^{I}$, H1$^{K}$, H1$^{M}$, H1$^{O}$), 5.30 (bs, 1H, H1$^{B}$), 5.18 (bs, 1H, H1$^{D}$), 5.11 (d, $J$ = 3.4 Hz, 1H, H1$^{A}$), 4.78 (1H, H5$^{D}$), 4.65 (bs, 1H, H5$^{B}$), 4.46 (d, $J$ = 7.8 Hz, 6H, H1$^{F}$, H1$^{H}$, H1$^{J}$, H1$^{L}$, H1$^{N}$, H1$^{P}$), 4.40 – 4.34 (m, 1H, H6a$^{C}$), 4.31 – 4.27 (m, 2H, H2$^{D}$, H2$^{B}$), 4.26 – 4.22 (m, 1H, H6b$^{C}$), 4.20 – 4.17 (m, 1H, H3$^{D}$), 4.17 –

4.14 (m, 1H, $H3^B$ ), 4.11 – 4.08 (m, 1H, $H4^B$), 4.05 (bs, 1H, $H4^D$), 4.00 – 3.95 (m, 1H, $H5^C$), 3.92 – 3.55 (m, 66H, $H2^G$, $H2^I$, $H2^K$, $H2^M$, $H2^O$, $H2^Q$, $H6a^A$, $H6b^A$, $H6a^E$, $H6b^E$, $H6a^G$, $H6b^G$, $H6a^I$, $H6b^I$, $H6a^K$, $H6b^K$, $H6a^M$, $H6b^M$, $H6a^O$, $H6b^O$, $H6a^Q$, $H6b^Q$, $H5^E$, $H5^G$, $H5^I$, $H5^K$, $H5^M$, $H5^O$, $H5^Q$, $H5^A$, $H3^A$, OCH*H* linker, $H3^E$, $H3^G$, $H3^I$, $H3^K$, $H3^M$, $H3^O$, $H3^Q$, $H3^C$, $H5^F$, $H5^H$, $H5^J$, $H5^L$, $H5^N$, $H5^P$, $H4^C$, $H4^F$, $H4^H$, $H4^J$, $H4^L$, $H4^N$, $H4^P$, $H4^A$, $H4^E$, $H4^G$, $H4^I$, $H4^K$, $H4^M$, $H4^O$, $H3^F$, $H3^H$, $H3^J$, $H3^L$, $H3^N$, $H3^P$), 3.54 – 3.49 (m, 1H, OCH*H* linker), 3.49 – 3.43 (m, 1H, $H4^Q$), 3.37 – 3.30 (m, 6H, $H2^F$, $H2^H$, $H2^J$, $H2^L$, $H2^N$, $H2^P$), 3.28 – 3.19 (m, 3H, $H2^C$, $H2^E$, $H2^A$), 3.02 (t, *J* = 7.3 Hz, 2H, $CH_2N$ linker), 2.03 (s, 3H, $NHCOCH_3^Q$), 2.01 (s, 15H, 5 × $NHCOCH_3$), 1.76 – 1.47 (m, 6H, 3 × $CH_2$ linker). $^{13}C$ NMR from HSQC (151 MHz, $D_2O$) δ 102.3 ($C1^F$, $C1^H$, $C1^J$, $C1^L$, $C1^N$, $C1^P$), 99.2 ($C1^D$), 97.2 ($C1^B$), 97.0 ($C1^A$), 96.7 ($C1^Q$, $C1^O$, $C1^K$, $C1^M$, $C1^E$, $C1^G$, $C1^I$), 96.0 ($C1^C$), 78.0 ($C3^F$, $C3^H$, $C3^J$, $C3^L$, $C3^N$, $C3^P$), 77.0 ($C2^B$), 76.4 ($C2^D$, $C5^F$, $C5^H$, $C5^J$, $C5^L$, $C5^N$, $C5^P$, $C4^F$, $C4^H$, $C4^J$, $C4^L$, $C4^N$, $C4^P$, $C4^C$, $C4^G$, $C4^I$, $C4^A$, $C4^E$, $C4^K$, $C4^M$, $C4^O$), 75.8 ($C4^B$, $C4^D$), 73.4 ($C2^F$, $C2^H$, $C2^J$, $C2^L$, $C2^N$, $C2^P$), 70.5 ($C5^G$, $C5^I$, $C5^K$, $C5^M$, $C5^O$, $C5^Q$, $C3^G$, $C3^I$, $C3^K$, $C3^M$, $C3^O$, $C3^Q$), 70.1 ($C5^B$, $C4^Q$), 69.6 ($C3^B$), 69.5 ($C5^D$, $C3^D$, $C5^C$, $C5^E$, $C5^A$, $C3^C$, $C3^E$), 67.7 (O<u>C</u>H$_2$ linker), 66.5 ($C6^C$), 59.7 ($C6^A$, $C6^E$, $C6^G$, $C6^I$, $C6^K$, $C6^M$, $C6^O$), 57.9 ($C2^C$, $C2^E$, $C2^A$), 53.3 ($C2^G$, $C2^I$, $C2^K$, $C2^M$, $C2^O$, $C2^Q$), 50.4 ($C6^Q$), 39.6 (N<u>C</u>H$_2$ linker), 28.0 (<u>C</u>H$_2$ linker), 26.4 (<u>C</u>H$_2$ linker), 22.6 (<u>C</u>H$_2$ linker), 21.7 (6 × NHCO<u>C</u>H$_3$). ESI-MS: m/z calculated for $C_{119}H_{184}N_{13}O_{108}S_6$ $[M-14Na+11H]^{3-}$: 1238.5893; found: 1238.5778.

$^1H$ NMR (600 MHz, $D_2O$)

| | **H1** | **H2** | **H3** | **H4** | **H5** | **H6** |
|---|---|---|---|---|---|---|
| **A** | 5.11<br>(d, *J* = 3.4 Hz) | 3.22 | 3.87 | 3.70 | 3.87 | 3.85 |
| **B** | 5.30 | 4.29 | 4.16 | 4.10 | 4.66 | – |
| **C** | 5.44<br>(d, *J* = 3.4 Hz) | 3.26 | 3.59 | 3.76 | 3.98 | 4.37, 4.26 |
| **D** | 5.18 | 4.29 | 4.19 | 4.05 | 4.78 | – |
| **E** | 5.39<br>(d, *J* = 3.4 Hz) | 3.22 | 3.68 | 3.68 | 3.88 | 3.85 |
| **F/H/J/L/N/P** | 4.46<br>(d, *J* = 7.8 Hz) | 3.34 | 3.66 | 3.73 | 3.77 | – |
| **G/I/K/M/O** | 5.36 | 3.86 | 3.64 | 3.64 | 3.85 | 3.85 |

| | | | | | | |
|---|---|---|---|---|---|---|
| | (d, $J$ = 3.2 Hz) | | | | | |
| **Q** | 5.40<br>(d, $J$ = 3.8 Hz) | 3.85 | 3.68 | 3.47 | 3.88 | 3.62 |

$^{13}$C NMR from HSQC (151 MHz, $D_2O$)

| | **C1** | **C2** | **C3** | **C4** | **C5** | **C6** |
|---|---|---|---|---|---|---|
| **A** | 97.0 | 57.9 | 68.4 | 76.4 | 69.5 | 59.7 |
| **B** | 97.2 | 77.0 | 69.6 | 75.8 | 70.1 | – |
| **C** | 96.0 | 57.9 | 69.5 | 76.4 | 69.5 | 66.5 |
| **D** | 99.2 | 76.4 | 69.5 | 75.8 | 69.5 | – |
| **E** | 96.7 | 57.9 | 69.5 | 76.4 | 69.5 | 59.7 |
| **F/H/J/L/N/P** | 102.3 | 73.4 | 78.3 | 76.4 | 76.4 | – |
| **G/I/K/M/O** | 96.7 | 53.3 | 70.5 | 76.4 | 70.5 | 59.7 |
| **Q** | 96.7 | 53.3 | 70.5 | 70.1 | 70.5 | 50.4 |

**5-Aminopentyl *O*-(β-D-glucopyranosyluronate)-(1→4)-*O*-(2-sulfamino-2-deoxy-α-D-glucopyranosyl)-(1→4)-*O*-(2-*O*-sulfate-α-L-idopyranosyluronate)-(1→4)-*O*-(2-sulfamino-6-*O*-sulfate-2-deoxy-α-D-glucopyranosyl)-(1→4)-*O*-(2-*O*-sulfate-α-L-idopyranosyluronate)-(1→4)-*O*-2-sulfamino-2-deoxy-α-D-glucopyranosyl) -*O*-[*N*-ethyl-3-(1*H*-1,2,3-triazol-4-yl)propanamide]-(1→6)-*N*-(2-acetamido-2-deoxy-α-D-glucopyranosyl)-(1→4)-*O*-(β-D-glucopyranosyluronate)-(1→4)-*O*-(2-sulfamino-2-deoxy-α-D-glucopyranosyl)-(1→4)-*O*-(2-*O*-sulfate-α-L-idopyranosyluronate)- (1→4)-*O*-(2-sulfamino-6-*O*-sulfate-2-deoxy-α-D-glucopyranosyl)-(1→4)-*O*-(2-*O*-sulfate-α-L-**

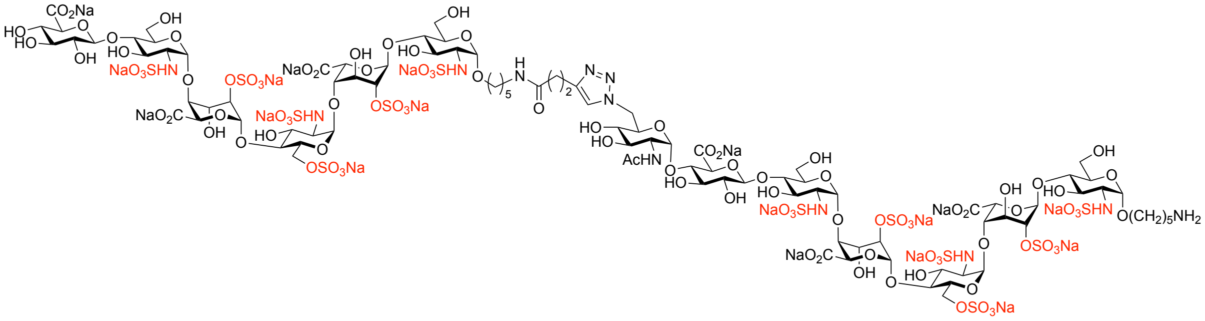

**idopyranosyluronate)-(1→4)-*O*-2-sulfamino-2-deoxy-α-D-glucopyranoside, sodium salt (9).**

Compound **2** (1.1 mg, 0. 54 μmol) and compound **3** (1.1 mg, 0.54 μmol) were subjected to copper (I)-catalyzed azide-alkyne cycloaddition (CuAAC) reaction according to the general procedure to give the title compound **9** as a white powder (1.3 mg, 61%). $^{1}$H NMR (600 MHz, $D_2O$) δ 7.85 (s, 1H, C*H* triazole), 5.47 (d, *J* = 3.5 Hz, 1H, $H1^{C}$), 5.44 – 5.40 (m, 2H, $H1^{E}$, $H1^{E1}$), 5.40 (d, *J* = 3.5 Hz, 1H, $H1^{C1}$), 5.35 (d, *J* = 3.5 Hz, 1H, $H1^{G}$), 5.33 (d, *J* = 3.5 Hz, 1H, $H1^{B}$), 5.22 (bs, 2H, $H1^{D}$, $H1^{D1}$), 5.18 (bs, 1H, $H1^{B1}$), 5.16 – 5.12 (m, 2H, $H1^{A}$, $H1^{A1}$), 4.91 – 4.77 (m, 4H, $H6a^{G}$, $H5^{D}$, $H5^{D1}$, $H5^{B1}$), 4.68 (d, *J* = 2.9 Hz, 1H, $H5^{B}$), 4.61 – 4.56 (m, 1H, $H6b^{G}$), 4.54 – 4.49 (m, 2H, $H1^{F}$, $H1^{F1}$), 4.43 – 4.38 (m, 2H, $H6a^{C}$, $H6a^{C1}$), 4.35 – 4.25 (m, 6H, $H2^{D}$, $H2^{D1}$, $H2^{B}$, $H2^{B1}$, $H6b^{C}$, $H6b^{C1}$), 4.24 – 4.17 (m, 4H, $H3^{D}$, $H3^{D1}$, $H3^{B}$, $H3^{B1}$), 4.15 – 4.12 (m, 1H, $H4^{B}$), 4.12 – 4.07 (m, 4H, $H4^{B1}$, $H4^{D}$, $H4^{D1}$, $H5^{G}$), 4.04 – 3.99 (m, 2H, $H5^{C}$, $H5^{C1}$), 3.97 – 3.60 (m, 32H, $H6a^{A}$, $H6b^{A}$, $H6a^{A1}$, $H6b^{A1}$, $H6a^{E}$, $H6b^{E}$, $H6a^{E1}$, $H6b^{E1}$, $H3^{A}$, $H5^{E}$, $H5^{E1}$, $H5^{A}$, $H5^{A1}$, $H5^{F}$, $H4^{F}$, $H5^{F1}$, $H4^{C}$, $H4^{C1}$, $H4^{A}$, $H4^{A1}$, $H4^{E}$, $H4^{E1}$, $H3^{F}$, $H3^{G}$, $H3^{E}$, $H3^{E1}$, $H3^{C1}$, $H3^{A1}$, $H3^{C}$, $H2^{G}$, OCH*H* linker, OCH*H* linker1), 3.59 – 3.48 (m, 4H, OCH*H* linker, OCH*H* linker1, $H3^{F1}$, $H4^{F1}$), 3.43 – 3.38 (m, 2H, $H2^{F}$, $H2^{F1}$), 3.32 – 3.23 (m, 6H, $H2^{C}$, $H2^{C1}$, $H2^{E}$, $H2^{E1}$, $H2^{A}$, $H2^{A1}$), 3.15 (t, *J* = 7.1 Hz, 2H, N$CH_2$ linker1), 3.04 (t, *J* = 7.2 Hz, 2H, N$CH_2$ linker), 3.00 (t, *J* = 7.5 Hz, 2H, $CH_2$-triazole), 2.79 – 2.72 (m, 1H, $H4^{G}$), 2.61 (t, *J* = 7.6 Hz, 2H, $CH_2$CONH), 2.03 (s, 3H, NHCO$CH_3$), 1.79 – 1.29 (m, 12H, 6 × $CH_2$ linker). $^{13}$C NMR from HSQC (151 MHz, $D_2O$) δ 124.7 (C̲H triazole), 102.1 ($C1^{F}$, $C1^{F1}$), 99.2 ($C1^{D}$, $C1^{D1}$, $C1^{B1}$), 97.3 ($C1^{G}$, $C1^{B}$), 97.0 ($C1^{A}$, $C1^{A1}$), 96.5 ($C1^{E}$, $C1^{E1}$, $C1^{C1}$), 95.9 ($C1^{C}$), 77.5 ($C4^{A}$, $C4^{A1}$, $C4^{E}$, $C4^{E1}$, $C3^{F}$), 76.7 ($C2^{B}$, $C2^{B1}$, $C5^{F}$), 75.9 ($C2^{D}$, $C2^{D1}$, $C4^{B}$, $C4^{B1}$, $C4^{D}$, $C4^{D1}$, $C4^{F}$, $C5^{F1}$, $C4^{C}$, $C4^{C1}$), 75.2 ($C3^{F1}$), 73.2 ($C2^{F}$, $C2^{F1}$), 71.9 ($C4^{F1}$), 70.9 ($C5^{A1}$), 70.7 ($C5^{E}$, $C5^{E1}$, $C5^{A}$), 70.0 ($C5^{B}$, $C4^{G}$), 69.7 ($C3^{G}$, $C3^{E}$, $C3^{E1}$, $C3^{C}$, $C3^{C1}$, $C3^{A1}$), 69.4 ($C5^{D}$, $C5^{D1}$, $C5^{B1}$, $C3^{B}$, $C3^{B1}$, $C3^{D}$, $C3^{D1}$, $C5^{G}$, $C5^{C}$, $C5^{C1}$), 68.4 (OC̲$H_2$ linker1), 68.2 ($C3^{A}$), 67.6 (OC̲$H_2$ linker), 66.4 ($C6^{C}$, $C6^{C1}$), 59.7 ($C6^{A}$, $C6^{A1}$, $C6^{E}$, $C6^{E1}$), 57.9 ($C2^{A}$, $C2^{A1}$, $C2^{C}$, $C2^{C1}$, $C2^{E}$, $C2^{E1}$), 53.5 ($C2^{G}$), 49.6 ($C6^{G}$), 39.5 (2 × NC̲$H_2$ linker), 35.4 (C̲$H_2$CONH), 28.2 (C̲$H_2$ linker, 2 × C̲$H_2$ linker1), 26.5 (C̲$H_2$ linker), 22.7 (C̲$H_2$ linker, C̲$H_2$ Linker1), 22.0 (NHCOC̲$H_3$), 21.4 (C̲$H_2$-triazole). ESI-MS: m/z calculated for $C_{95}H_{153}N_{12}O_{103}S_{12}$ $[M-18Na+15H]^{3-}$: 1164.7934; found: 1164.7493.

$^1H$ NMR (600 MHz, $D_2O$)

| | H1 | H2 | H3 | H4 | H5 | H6 |
|---|---|---|---|---|---|---|
| **A** | 5.14 | 3.25 | 3.89 | 3.74 | 3.90 | 3.90 |
| **A1** | 5.14 | 3.25 | 3.65 | 3.71 | 3.79 | 3.90 |
| **B** | 5.33 (d, $J$ = 3.5 Hz) | 4.33 | 4.18 | 4.13 | 4.68 (d, $J$ = 2.9 Hz) | – |
| **B1** | 5.18 | 4.31 | 4.21 | 4.09 | 4.83 | – |
| **C** | 5.47 (d, $J$ = 3.5 Hz) | 3.29 | 3.62 | 3.78 | 4.01 | 4.40, 4.27 |
| **C1** | 5.40 (d, $J$ = 3.5 Hz) | 3.29 | 3.66 | 3.78 | 4.01 | 4.40, 4.27 |
| **D/D1** | 5.22 | 4.33 | 4.21 | 4.09 | 4.83 | – |
| **E/E1** | 5.42 | 3.26 | 3.73 | 3.71 | 3.94 | 3.90 |
| **F** | 4.51 | 3.39 | 3.70 | 3.75 | 3.82 | – |
| **F1** | 4.51 | 3.39 | 3.53 | 3.53 | 3.75 | – |
| **G** | 5.35 (d, $J$ = 3.5 Hz) | 3.68 | 3.73 | 2.75 | 4.10 | 4.86, 4.58 |

$^{13}C$ NMR from HSQC (151 MHz, $D_2O$)

| | C1 | C2 | C3 | C4 | C5 | C6 |
|---|---|---|---|---|---|---|
| **A** | 97.0 | 57.9 | 68.2 | 77.5 | 70.7 | 59.7 |
| **A1** | 97.0 | 57.9 | 69.7 | 77.5 | 70.9 | 59.7 |
| **B** | 97.3 | 76.7 | 69.4 | 75.9 | 70.0 | – |
| **B1** | 99.2 | 76.7 | 69.4 | 75.9 | 69.4 | – |
| **C** | 95.9 | 57.9 | 69.7 | 75.9 | 69.4 | 66.4 |
| **C1** | 96.5 | 57.9 | 69.7 | 75.9 | 69.4 | 66.4 |
| **D/D1** | 99.2 | 75.9 | 69.4 | 75.9 | 69.4 | – |
| **E/E1** | 96.5 | 57.9 | 69.7 | 77.5 | 70.7 | 59.7 |
| **F** | 102.1 | 73.2 | 77.5 | 75.9 | 76.7 | – |

| | | | | | | |
|---|---|---|---|---|---|---|
| **F1** | 102.1 | 73.2 | 75.2 | 71.9 | 75.9 | – |
| **G** | 97.3 | 53.5 | 69.7 | 70.0 | 69.4 | 49.6 |

**5-Aminopentyl *O*-(β-D-glucopyranosyluronate)-(1→4)-*O*-(2-sulfamino-2-deoxy-α-D-glucopyranosyl)-(1→4)-*O*-(2-*O*-sulfate-α-L-idopyranosyluronate)-(1→4)-*O*-(2-sulfamino-6-*O*-sulfate-2-deoxy-α-D-glucopyranosyl)-(1→4)-*O*-(2-*O*-sulfate-α-L-idopyranosyluronate)-(1→4)-*O*-2-sulfamino-2-deoxy-α-D-glucopyranosyl) -*O*-[*N*-ethyl-3-(1*H*-1,2,3-triazol-4-yl)propanamide]-(1→6)-*N*-(2-acetamido-2-deoxy-α-D-glucopyranosyl)-(1→4)-*O*-(β-D-glucopyranosyluronate)-(1→4)-*O*-(2-acetamido-2-deoxy-α-D-glucopyranosyl)-(1→4)-*O*-(β-D-glucopyranosyluronate)-(1→4)-*O*-(2-sulfamino-2-deoxy-α-D-glucopyranosyl)-(1→4)-*O*-(2-*O*-sulfate-α-L-idopyranosyluronate)-(1→4)-*O*-(2-sulfamino-6-*O*-sulfate-2-deoxy-α-D-glucopyranosyl)-(1→4)-*O*-(2-*O*-sulfate-α-L-idopyranosyluronate)-(1→4)-*O*-2-sulfamino-2-deoxy-α-D-glucopyranoside, sodium salt (10).**

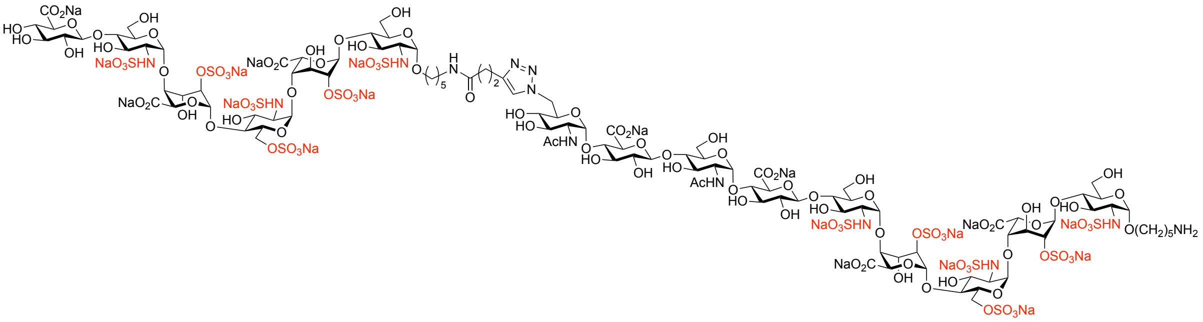

Compound **2** (1.2 mg, 0.66 μmol) and compound **4** (1.6 mg, 0.66 μmol) were subjected to copper (I)-catalyzed azide-alkyne cycloaddition (CuAAC) reaction according to the general procedure to give the title compound **10** as a white powder (1.8 mg, 63%). $^{1}$H NMR (600 MHz, $D_2O$) δ 7.85 (s, 1H, C*H* triazole), 5.46 (d, *J* = 3.4 Hz, 1H, H1$^{C}$), 5.44 – 5.40 (m, 2H, H1$^{E}$, H1$^{E1}$), 5.40 – 5.38 (m, 2H, H1$^{G}$, H1$^{C1}$), 5.35 (d, *J* = 3.3 Hz, 1H, H1$^{I}$), 5.32 (d, *J* = 3.3 Hz, 1H, H1$^{B}$), 5.22 (bs, 2H, H1$^{D}$, H1$^{D1}$), 5.18 (bs, 1H, H1$^{B1}$), 5.16 – 5.12 (m, 2H, H1$^{A}$, H1$^{A1}$), 4.90 – 4.74 (m, 4H, H6a$^{I}$, H5$^{D}$, H5$^{D1}$, H5$^{B1}$), 4.68 (d, *J* = 2.9 Hz, 1H, H5$^{B}$), 4.61 – 4.56 (m, 1H, H6b$^{I}$), 4.54 – 4.47 (m, 3H, H1$^{F}$, H1$^{F1}$, H1$^{H}$), 4.44 – 4.37 (m, 2H, H6a$^{C}$, H6a$^{C1}$), 4.35 – 4.25 (m, 6H, H2$^{D}$, H2$^{D1}$, H2$^{B}$, H2$^{B1}$, H6b$^{C}$, H6b$^{C1}$), 4.23 – 4.17 (m, 4H, H3$^{D}$, H3$^{D1}$, H3$^{B}$, H3$^{B1}$), 4.14 – 4.12 (m, 1H, H4$^{B}$), 4.12 – 4.06 (m, 4H, H4$^{B1}$, H4$^{D}$, H4$^{D1}$, H5$^{I}$), 4.04 – 3.99 (m, 2H, H5$^{C}$, H5$^{C1}$), 3.97 –

3.60 (m, 41H, H2$^G$, H6a$^A$, H6b$^A$, H6a$^{A1}$, H6b$^{A1}$, H6a$^E$, H6b$^E$, H6a$^{E1}$, H6b$^{E1}$, H6a$^G$, H6b$^G$, H3$^A$, H5$^E$, H5$^{E1}$, H5$^A$, H5$^G$, H3$^G$, H5$^{A1}$, H5$^F$, H5$^H$, H4$^G$, H4$^F$, H4$^H$, H5$^{F1}$, H4$^C$, H4$^{C1}$, H4$^A$, H4$^{A1}$, H4$^E$, H4$^{E1}$, H3$^F$, H3$^H$, H3$^I$, H3$^E$, H3$^{E1}$, H3$^{C1}$, H3$^{A1}$, H3$^C$, H2$^I$, OCH*H* linker, OCH*H* linker1), 3.58 – 3.48 (m, 4H, OCH*H* linker, OCH*H* linker1, H3$^{F1}$, H4$^{F1}$), 3.42 – 3.36 (m, 3H, H2$^F$, H2$^{F1}$, H2$^H$), 3.32 – 3.23 (m, 6H, H2$^C$, H2$^{C1}$, H2$^E$, H2$^{E1}$, H2$^A$, H2$^{A1}$), 3.15 (t, $J$ = 7.1 Hz, 2H, N$CH_2$ linker1), 3.05 (t, $J$ = 7.2 Hz, 2H, N$CH_2$ linker), 3.01 (t, $J$ = 7.5 Hz, 2H, $CH_2$-triazole), 2.81 – 2.75 (m, 1H, H4$^I$), 2.61 (t, $J$ = 7.6 Hz, 2H, $CH_2$CONH), 2.05 (s, 3H, NHCO$CH_3$), 2.03 (s, 3H, NHCO$CH_3$), 1.79 – 1.26 (m, 12H, 6 × $CH_2$ linker). $^{13}$C NMR from HSQC (151 MHz, $D_2O$) δ 124.7 (<u>C</u>H triazole), 102.2 (C1$^F$, C1$^{F1}$, C1$^H$), 99.2 (C1$^D$, C1$^{D1}$, C1$^{B1}$), 97.2 (C1$^I$, C1$^B$), 96.9 (C1$^A$, C1$^{A1}$), 96.5 (C1$^E$, C1$^{E1}$, C1$^{C1}$, C1$^G$), 95.9 (C1$^C$), 77.5 (C3$^F$, C3$^H$), 76.5 (C2$^B$, C2$^{B1}$, C5$^F$, C5$^H$, C4$^G$, C4$^F$, C4$^H$, C5$^{F1}$, C4$^C$, C4$^{C1}$, C4$^A$, C4$^{A1}$, C4$^E$, C4$^{E1}$), 75.9 (C2$^D$, C2$^{D1}$, C4$^B$, C4$^{B1}$, C4$^D$, H4$^{D1}$), 75.1 (C3$^{F1}$), 73.2 (C2$^F$, C2$^{F1}$, C2$^H$), 71.8 (C4$^{F1}$), 70.7 (C5$^E$, C5$^{E1}$, C5$^A$, C5$^G$, C3$^G$, C5$^{A1}$), 69.9 (C5$^B$, C4$^I$), 69.7 (C3$^I$, C3$^E$, C3$^{E1}$, C3$^C$, C3$^{C1}$, C3$^{A1}$), 69.3 (C5$^D$, C5$^{D1}$, C5$^{B1}$, C5$^I$, C5$^C$, C5$^{C1}$), 69.1 (C3$^B$, C3$^{B1}$, C3$^D$, C3$^{D1}$), 68.3 (O<u>C</u>$H_2$ linker1), 68.1 (C3$^A$), 67.7 (O<u>C</u>$H_2$ linker), 66.4 (C6$^C$, C6$^{C1}$), 59.5 (C6$^A$, C6$^{A1}$, C6$^E$, C6$^{E1}$, C6$^G$), 57.9 (C2$^A$, C2$^{A1}$, C2$^C$, C2$^{C1}$, C2$^E$, C2$^{E1}$), 53.3 (C2$^G$, C2$^I$), 49.6 (C6$^I$), 39.5 (2 × N<u>C</u>$H_2$ linker), 35.2 (<u>C</u>$H_2$CONH), 28.0 (<u>C</u>$H_2$ linker, 2 × <u>C</u>$H_2$ linker1), 26.3 (<u>C</u>$H_2$ linker), 22.6 (<u>C</u>$H_2$ linker, <u>C</u>$H_2$ linker1), 21.8 (2 × NHCO<u>C</u>$H_3$), 21.1 (<u>C</u>$H_2$-triazole). ESI-MS: m/z calculated for $C_{109}H_{174}N_{13}O_{114}S_{12}$ [M-19Na+16H]$^{3-}$: 1291.1639; found: 1291.1645.

$^1$H NMR (600 MHz, $D_2O$)

| | **H1** | **H2** | **H3** | **H4** | **H5** | **H6** |
|---|---|---|---|---|---|---|
| **A** | 5.14 | 3.25 | 3.89 | 3.74 | 3.89 | 3.89 |
| **A1** | 5.14 | 3.25 | 3.65 | 3.72 | 3.79 | 3.89 |
| **B** | 5.32<br>(d, $J$ = 3.3 Hz) | 4.32 | 4.18 | 4.13 | 4.68<br>(d, $J$ = 2.9 Hz) | – |
| **B1** | 5.18 | 4.31 | 4.21 | 4.09 | 4.79 | – |
| **C** | 5.46<br>(d, $J$ = 3.4 Hz) | 3.29 | 3.62 | 3.78 | 4.01 | 4.41, 4.28 |
| **C1** | 5.40 | 3.29 | 3.66 | 3.78 | 4.01 | 4.41, 4.28 |

| | | | | | | |
|---|---|---|---|---|---|---|
| **D/D1** | 5.22 | 4.33 | 4.21 | 4.09 | 4.82 | – |
| **E/E1** | 5.42 | 3.26 | 3.72 | 3.71 | 3.94 | 3.89 |
| **F/H** | 4.50 | 3.39 | 3.70 | 3.76 | 3.81 | – |
| **F1** | 4.51 | 3.40 | 3.53 | 3.53 | 3.75 | – |
| **G** | 5.40 | 3.89 | 3.89 | 3.79 | 3.89 | 3.89 |
| **I** | 5.35 (d, $J$ = 3.3 Hz) | 3.68 | 3.72 | 2.78 | 4.09 | 4.85, 4.59 |

$^{13}$C NMR from HSQC (151 MHz, $D_2O$)

| | **C1** | **C2** | **C3** | **C4** | **C5** | **C6** |
|---|---|---|---|---|---|---|
| **A** | 96.9 | 57.9 | 68.1 | 76.5 | 70.7 | 59.5 |
| **A1** | 96.9 | 57.9 | 69.7 | 76.5 | 70.7 | 59.5 |
| **B** | 97.2 | 76.5 | 69.1 | 75.9 | 69.9 | – |
| **B1** | 99.2 | 76.5 | 69.1 | 75.9 | 69.3 | – |
| **C** | 95.9 | 57.9 | 69.7 | 76.5 | 69.3 | 66.4 |
| **C1** | 96.5 | 57.9 | 69.7 | 76.5 | 69.3 | 66.4 |
| **D/D1** | 99.2 | 75.9 | 69.1 | 75.9 | 69.3 | – |
| **E/E1** | 96.5 | 57.9 | 69.7 | 76.5 | 70.7 | 59.5 |
| **F/H** | 102.2 | 73.2 | 77.5 | 76.5 | 76.5 | – |
| **F1** | 102.2 | 73.2 | 75.1 | 71.8 | 76.5 | – |
| **G** | 96.5 | 53.3 | 70.7 | 76.5 | 70.7 | 59.5 |
| **I** | 97.2 | 53.3 | 69.7 | 69.9 | 69.3 | 49.6 |

**5-Aminopentyl *O*-(β-D-glucopyranosyluronate)-(1→4)-*O*-(2-sulfamino-2-deoxy-α-D-glucopyranosyl)-(1→4)-*O*-(2-*O*-sulfate-α-L-idopyranosyluronate)-(1→4)-*O*-(2-sulfamino-6-*O*-sulfate-2-deoxy-α-D-glucopyranosyl)-(1→4)-*O*-(2-*O*-sulfate-α-L-idopyranosyluronate)-(1→4)-*O*-2-sulfamino-2-deoxy-α-D-glucopyranosyl) -*O*-[*N*-ethyl-3-(1*H*-1,2,3-triazol-4-yl)propanamide]-(1→6)-*N*-(2-acetamido-2-deoxy-α-D-glucopyranosyl) [-(1→4)-*O*-(β-D-glucopyranosyluronate)-(1→4)-*O*-(2-acetamido-2-**

**deoxy-α-D-glucopyranosyl)]$_2$-(1→4)-*O*-(β-D-glucopyranosyluronate)-(1→4)-*O*-(2-sulfamino-2-deoxy-α-D-glucopyranosyl)-(1→4)-*O*-(2-*O*-sulfate-α-L-idopyranosyluronate)-(1→4)-*O*-(2-sulfamino-6-*O*-sulfate-2-deoxy-α-D-glucopyranosyl)-(1→4)-*O*-(2-*O*-sulfate-α-L-idopyranosyluronate)-(1→4)-*O*-2-sulfamino-2-deoxy-α-D-glucopyranoside, sodium salt (11).**

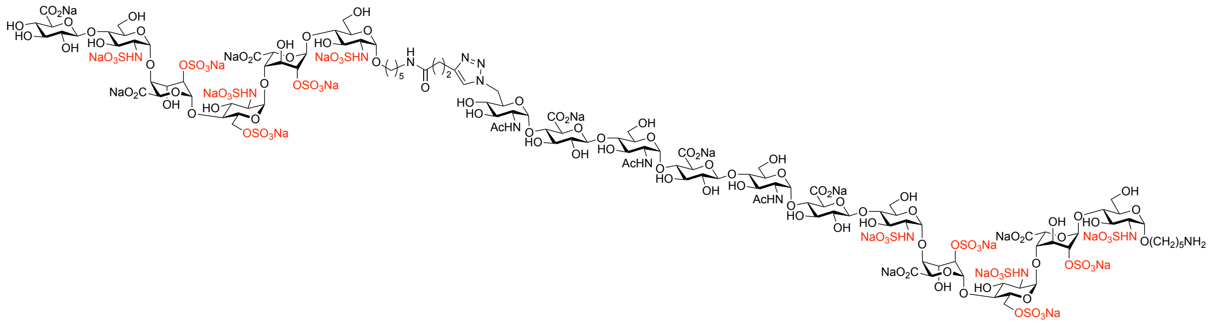

Compound **2** (0.8 mg, 0.43 µmol) and compound **5** (1.2 mg, 0.43 µmol) were subjected to copper (I)-catalyzed azide-alkyne cycloaddition (CuAAC) reaction according to the general procedure to give the title compound **11** as a white powder (1.2 mg, 60%). $^{1}$H NMR (600 MHz, $D_2O$) δ 7.85 (s, 1H, C*H* triazole), 5.46 (d, *J* = 3.4 Hz, 1H, $H1^{C}$), 5.44 – 5.40 (m, 2H, $H1^{E}$, $H1^{E1}$), 5.40 – 5.38 (m, 3H, $H1^{G}$, $H1^{I}$, $H1^{C1}$), 5.35 (d, *J* = 3.3 Hz, 1H, $H1^{K}$), 5.32 (d, *J* = 3.3 Hz, 1H, $H1^{B}$), 5.22 (bs, 2H, $H1^{D}$, $H1^{D1}$), 5.18 (bs, 1H, $H1^{B1}$), 5.16 – 5.12 (m, 2H, $H1^{A}$, $H1^{A1}$), 4.90 – 4.74 (m, 4H, $H6a^{K}$, $H5^{D}$, $H5^{D1}$, $H5^{B1}$), 4.69 (d, *J* = 2.9 Hz, 1H, $H5^{B}$), 4.61 – 4.56 (m, 1H, $H6b^{K}$), 4.54 – 4.47 (m, 4H, $H1^{F1}$, $H1^{F}$, $H1^{H}$, $H1^{J}$), 4.44 – 4.37 (m, 2H, $H6a^{C}$, $H6a^{C1}$), 4.35 – 4.29 (m, 4H, $H2^{D}$, $H2^{D1}$, $H2^{B}$, $H2^{B1}$), 4.29 – 4.25 (m, 2H, $H6b^{C}$, $H6b^{C1}$), 4.23 – 4.17 (m, 4H, $H3^{D}$, $H3^{D1}$, $H3^{B}$, $H3^{B1}$), 4.14 – 4.12 (m, 1H, $H4^{B}$), 4.12 – 4.06 (m, 4H, $H4^{B1}$, $H4^{D}$, $H4^{D1}$, $H5^{K}$), 4.04 – 3.99 (m, 2H, $H5^{C}$, $H5^{C1}$), 3.97 – 3.59 (m, 50H, $H2^{G}$, $H2^{I}$, $H6a^{A}$, $H6b^{A}$, $H6a^{A1}$, $H6b^{A1}$, $H6a^{E}$, $H6b^{E}$, $H6a^{E1}$, $H6b^{E1}$, $H6a^{G}$, $H6b^{G}$, $H6a^{I}$, $H6b^{I}$, $H3^{A}$, $H5^{E}$, $H5^{E1}$, $H5^{A}$, $H5^{G}$, $H3^{G}$, $H5^{I}$, $H3^{I}$, $H5^{A1}$, $H5^{F}$, $H5^{H}$, $H5^{J}$, $H4^{G}$, $H4^{I}$, $H4^{F}$, $H4^{H}$, $H4^{J}$, $H5^{F1}$, $H4^{C}$, $H4^{C1}$, $H4^{A}$, $H4^{A1}$, $H4^{E}$, $H4^{E1}$, $H3^{F}$, $H3^{H}$, $H3^{J}$, $H3^{K}$, $H3^{E}$, $H3^{E1}$, $H3^{C1}$, $H3^{A1}$, $H3^{C}$, $H2^{K}$, OCH*H* linker, OCH*H* linker1), 3.58 – 3.48 (m, 4H, OCH*H* linker, OCH*H* linker1, $H3^{F1}$, $H4^{F1}$), 3.42 – 3.36 (m, 4H, $H2^{F}$, $H2^{F1}$, $H2^{H}$, $H2^{J}$), 3.32 – 3.23 (m, 6H, $H2^{C}$, $H2^{C1}$, $H2^{E}$, $H2^{E1}$, $H2^{A}$, $H2^{A1}$), 3.15 (t, *J* = 7.1 Hz, 2H, NC$H_2$ linker1), 3.05 (t, *J* = 7.2 Hz, 2H, NC$H_2$ linker), 3.01 (t, *J* = 7.5 Hz, 2H, C$H_2$-triazole), 2.81 – 2.75 (m, 1H, $H4^{K}$), 2.61 (t, *J* = 7.6 Hz, 2H, C$H_2$CONH), 2.05 (s, 3H, NHCOC$H_3$), 2.04 (s, 3H, NHCOC$H_3$), 2.02 (s, 3H, NHCOC$H_3$), 1.79 – 1.26 (m, 12H, 6 × C$H_2$ linker). $^{13}$C NMR from HSQC (151

MHz, $D_2O$) δ 124.7 ($\underline{C}$H triazole), 102.2 (C1$^{F}$, C1$^{F1}$, C1$^{H}$, C1$^{J}$), 99.2 (C1$^{D}$, C1$^{D1}$, C1$^{B1}$), 97.2 (C1$^{K}$, C1$^{B}$), 96.9 (C1$^{A}$, C1$^{A1}$), 96.5 (C1$^{E}$, C1$^{E1}$, C1$^{C1}$, C1$^{G}$, C1$^{I}$), 95.9 (C1$^{C}$), 77.5 (C3$^{F}$, C3$^{H}$, C3$^{J}$), 76.5 (C2$^{B}$, C2$^{B1}$, C5$^{F}$, C5$^{H}$, C5$^{J}$, C4$^{G}$, C4$^{I}$, C4$^{F}$, C4$^{H}$, C4$^{J}$, C5$^{F1}$, C4$^{C}$, C4$^{C1}$, C4$^{A}$, C4$^{A1}$, C4$^{E}$, C4$^{E1}$), 75.9 (C2$^{D}$, C2$^{D1}$, C4$^{B}$, C4$^{B1}$, C4$^{D}$, H4$^{D1}$), 75.1 (C3$^{F1}$), 73.2 (C2$^{F}$, C2$^{F1}$, C2$^{H}$, C2$^{J}$), 71.8 (C4$^{F1}$), 70.7 (C5$^{E}$, C5$^{E1}$, C5$^{A}$, C5$^{G}$, C3$^{G}$, C5$^{I}$, C3$^{I}$, C5$^{A1}$), 69.9 (C5$^{B}$, C4$^{K}$), 69.7 (C3$^{K}$, C3$^{E}$, C3$^{E1}$, C3$^{C}$, C3$^{C1}$, C3$^{A1}$), 69.3 (C5$^{D}$, C5$^{D1}$, C5$^{B1}$, C5$^{K}$, C5$^{C}$, C5$^{C1}$), 69.1 (C3$^{B}$, C3$^{B1}$, C3$^{D}$, C3$^{D1}$), 68.3 (O$\underline{C}$H$_2$ linker1), 68.1 (C3$^{A}$), 67.7 (O$\underline{C}$H$_2$ linker), 66.4 (C6$^{C}$, C6$^{C1}$), 59.5 (C6$^{A}$, C6$^{A1}$, C6$^{E}$, C6$^{E1}$, C6$^{G}$, C6$^{I}$), 57.9 (C2$^{A}$, C2$^{A1}$, C2$^{C}$, C2$^{C1}$, C2$^{E}$, C2$^{E1}$), 53.3 (C2$^{G}$, C2$^{I}$, C2$^{K}$), 49.6 (C6$^{K}$), 39.5 (2 × N$\underline{C}$H$_2$ linker), 35.2 ($\underline{C}$H$_2$CONH), 28.0 ($\underline{C}$H$_2$ linker, 2 × $\underline{C}$H$_2$ linker1), 26.3 ($\underline{C}$H$_2$ linker), 22.6 ($\underline{C}$H$_2$ linker, $\underline{C}$H$_2$ linker1), 21.8 (3 × NHCO$\underline{C}$H$_3$), 21.1 ($\underline{C}$H$_2$-triazole). ESI-MS: m/z calculated for $C_{123}H_{194}N_{14}O_{125}S_{12}$ [M-20Na+16H]$^{4-}$: 1062.8989; found: 1062.8737.

$^{1}$H NMR (600 MHz, $D_2O$)

| | **H1** | **H2** | **H3** | **H4** | **H5** | **H6** |
|---|---|---|---|---|---|---|
| **A** | 5.14 | 3.25 | 3.89 | 3.74 | 3.89 | 3.89 |
| **A1** | 5.14 | 3.25 | 3.65 | 3.72 | 3.79 | 3.89 |
| **B** | 5.32 (d, *J* = 3.3 Hz) | 4.32 | 4.18 | 4.13 | 4.68 (d, *J* = 2.9 Hz) | – |
| **B1** | 5.18 | 4.31 | 4.21 | 4.09 | 4.79 | – |
| **C** | 5.46 (d, *J* = 3.4 Hz) | 3.29 | 3.62 | 3.78 | 4.01 | 4.41, 4.28 |
| **C1** | 5.40 | 3.29 | 3.66 | 3.78 | 4.01 | 4.41, 4.28 |
| **D/D1** | 5.22 | 4.33 | 4.21 | 4.09 | 4.82 | – |
| **E/E1** | 5.42 | 3.26 | 3.72 | 3.71 | 3.94 | 3.89 |
| **F/H/J** | 4.50 | 3.39 | 3.70 | 3.76 | 3.81 | – |
| **F1** | 4.51 | 3.40 | 3.53 | 3.53 | 3.75 | – |
| **G/I** | 5.40 | 3.89 | 3.89 | 3.79 | 3.89 | 3.89 |
| **K** | 5.35 (d, *J* = 3.3 Hz) | 3.68 | 3.72 | 2.78 | 4.09 | 4.85, 4.59 |

$^{13}$C NMR from HSQC (151 MHz, $D_2O$)

| | C1 | C2 | C3 | C4 | C5 | C6 |
|---|---|---|---|---|---|---|
| **A** | 96.9 | 57.9 | 68.1 | 76.5 | 70.7 | 59.5 |
| **A1** | 96.9 | 57.9 | 69.7 | 76.5 | 70.7 | 59.5 |
| **B** | 97.2 | 76.5 | 69.1 | 75.9 | 69.9 | – |
| **B1** | 99.2 | 76.5 | 69.1 | 75.9 | 69.3 | – |
| **C** | 95.9 | 57.9 | 69.7 | 76.5 | 69.3 | 66.4 |
| **C1** | 96.5 | 57.9 | 69.7 | 76.5 | 69.3 | 66.4 |
| **D/D1** | 99.2 | 75.9 | 69.1 | 75.9 | 69.3 | – |
| **E/E1** | 96.5 | 57.9 | 69.7 | 76.5 | 70.7 | 59.5 |
| **F/H/J** | 102.2 | 73.2 | 77.5 | 76.5 | 76.5 | – |
| **F1** | 102.2 | 73.2 | 75.1 | 71.8 | 76.5 | – |
| **G/I** | 96.5 | 53.3 | 70.7 | 76.5 | 70.7 | 59.5 |
| **K** | 97.2 | 53.3 | 69.7 | 69.9 | 69.3 | 49.6 |

**5-Aminopentyl *O*-(β-D-glucopyranosyluronate)-(1→4)-*O*-(2-sulfamino-2-deoxy-α-D-glucopyranosyl)-(1→4)-*O*-(2-*O*-sulfate-α-L-idopyranosyluronate)-(1→4)-*O*-(2-sulfamino-6-*O*-sulfate-2-deoxy-α-D-glucopyranosyl)-(1→4)-*O*-(2-*O*-sulfate-α-L-idopyranosyluronate)-(1→4)-*O*-2-sulfamino-2-deoxy-α-D-glucopyranosyl) -*O*-[*N*-ethyl-3-(1*H*-1,2,3-triazol-4-yl)propanamide]-(1→6)-*N*-(2-acetamido-2-deoxy-α-D-glucopyranosyl) [-(1→4)-*O*-(β-D-glucopyranosyluronate)-(1→4)-*O*-(2-acetamido-2-deoxy-α-D-glucopyranosyl)]$_3$-(1→4)-*O*-(β-D-glucopyranosyluronate)-(1→4)-*O*-(2-sulfamino-2-deoxy-α-D-glucopyranosyl)-(1→4)-*O*-(2-*O*-sulfate-α-L-**

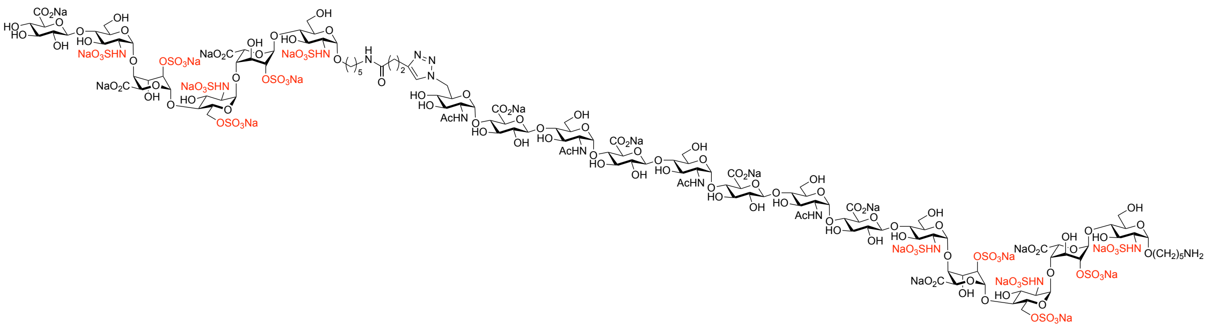

**idopyranosyluronate)-(1→4)-*O*-(2-sulfamino-6-*O*-sulfate-2-deoxy-α-D-glucopyranosyl)-(1→4)-*O*-(2-*O*-sulfate-α-L-idopyranosyluronate)-(1→4)-*O*-2-sulfamino-2-deoxy-α-D-glucopyranoside, sodium salt (12).**

Compound **2** (0.5 mg, 0.27 μmol) and compound **6** (0.8 mg, 0.25 μmol) were subjected to copper (I)-catalyzed azide-alkyne cycloaddition (CuAAC) reaction according to the general procedure to give the title compound **12** as a white powder (0.73 mg, 58%). $^{1}$H NMR (600 MHz, $D_2O$) δ 7.85 (s, 1H, C*H* triazole), 5.46 (bs, 1H, H1$^{C}$), 5.44 – 5.40 (m, 2H, H1$^{E}$, H1$^{E1}$), 5.40 – 5.38 (m, 3H, H1$^{G}$, H1$^{I}$, H1$^{K}$, H1$^{C1}$), 5.35 (bs, 1H, H1$^{M}$), 5.32 (bs, 1H, H1$^{B}$), 5.22 (bs, 2H, H1$^{D}$, H1$^{D1}$), 5.18 (bs, 1H, H1$^{B1}$), 5.16 – 5.12 (m, 2H, H1$^{A}$, H1$^{A1}$), 4.90 – 4.74 (m, 4H, H6a$^{M}$, H5$^{D}$, H5$^{D1}$, H5$^{B1}$), 4.69 (bs, 1H, H5$^{B}$), 4.61 – 4.56 (m, 1H, H6b$^{M}$), 4.54 – 4.47 (m, 5H, H1$^{F1}$, H1$^{F}$, H1$^{H}$, H1$^{J}$, H1$^{L}$), 4.44 – 4.37 (m, 2H, H6a$^{C}$, H6a$^{C1}$), 4.35 – 4.29 (m, 4H, H2$^{D}$, H2$^{D1}$, H2$^{B}$, H2$^{B1}$), 4.29 – 4.25 (m, 2H, H6b$^{C}$, H6b$^{C1}$), 4.23 – 4.17 (m, 4H, H3$^{D}$, H3$^{D1}$, H3$^{B}$, H3$^{B1}$), 4.14 – 4.12 (m, 1H, H4$^{B}$), 4.12 – 4.06 (m, 4H, H4$^{B1}$, H4$^{D}$, H4$^{D1}$, H5$^{M}$), 4.04 – 3.99 (m, 2H, H5$^{C}$, H5$^{C1}$), 3.97 – 3.59 (m, 59H, H2$^{G}$, H2$^{I}$, H2$^{K}$, H6a$^{A}$, H6b$^{A}$, H6a$^{A1}$, H6b$^{A1}$, H6a$^{E}$, H6b$^{E}$, H6a$^{E1}$, H6b$^{E1}$, H6a$^{G}$, H6b$^{G}$, H6a$^{I}$, H6b$^{I}$, H6a$^{K}$, H6b$^{K}$, H3$^{A}$, H5$^{E}$, H5$^{E1}$, H5$^{A}$, H5$^{G}$, H3$^{G}$, H5$^{I}$, H3$^{I}$, H5$^{K}$, H3$^{K}$, H5$^{A1}$, H5$^{F}$, H5$^{H}$, H5$^{J}$, H5$^{L}$, H4$^{G}$, H4$^{I}$, H4$^{K}$, H4$^{F}$, H4$^{H}$, H4$^{J}$, H4$^{L}$, H5$^{F1}$, H4$^{C}$, H4$^{C1}$, H4$^{A}$, H4$^{A1}$, H4$^{E}$, H4$^{E1}$, H3$^{F}$, H3$^{H}$, H3$^{J}$, H3$^{L}$, H3$^{M}$, H3$^{E}$, H3$^{E1}$, H3$^{C1}$, H3$^{A1}$, H3$^{C}$, H2$^{M}$, OCH*H* linker, OCH*H* linker1), 3.58 – 3.48 (m, 4H, OCH*H* linker, OCH*H* linker1, H3$^{F1}$, H4$^{F1}$), 3.42 – 3.36 (m, 5H, H2$^{F}$, H2$^{F1}$, H2$^{H}$, H2$^{J}$, H2$^{L}$), 3.32 – 3.23 (m, 6H, H2$^{C}$, H2$^{C1}$, H2$^{E}$, H2$^{E1}$, H2$^{A}$, H2$^{A1}$), 3.15 (t, *J* = 7.1 Hz, 2H, NC$H_2$ linker1), 3.05 (t, *J* = 7.2 Hz, 2H, NC$H_2$ linker), 3.01 (t, *J* = 7.5 Hz, 2H, C$H_2$-triazole), 2.81 – 2.75 (m, 1H, H4$^{M}$), 2.61 (t, *J* = 7.6 Hz, 2H, C$H_2$CONH), 2.05 (s, 3H, NHCOC$H_3$), 2.04 (s, 3H, NHCOC$H_3$), 2.02 (s, 3H, NHCOC$H_3$), 1.79 – 1.26 (m, 12H, 6 × C$H_2$ linker). $^{13}$C NMR from HSQC (151 MHz, $D_2O$) δ 124.7 (<u>C</u>H triazole), 102.2 (C1$^{F}$, C1$^{F1}$, C1$^{H}$, C1$^{J}$, C1$^{L}$), 99.2 (C1$^{D}$, C1$^{D1}$, C1$^{B1}$), 97.2 (C1$^{M}$, C1$^{B}$), 96.9 (C1$^{A}$, C1$^{A1}$), 96.5 (C1$^{E}$, C1$^{E1}$, C1$^{C1}$, C1$^{G}$, C1$^{I}$, C1$^{K}$), 95.9 (C1$^{C}$), 77.5 (C3$^{F}$, C3$^{H}$, C3$^{J}$, C3$^{L}$), 76.5 (C2$^{B}$, C2$^{B1}$, C5$^{F}$, C5$^{H}$, C5$^{J}$, C5$^{L}$, C4$^{G}$, C4$^{I}$, C4$^{K}$, C4$^{F}$, C4$^{H}$, C4$^{J}$, C4$^{L}$, C5$^{F1}$, C4$^{C}$, C4$^{C1}$, C4$^{A}$, C4$^{A1}$, C4$^{E}$, C4$^{E1}$), 75.9 (C2$^{D}$, C2$^{D1}$, C4$^{B}$, C4$^{B1}$, C4$^{D}$, H4$^{D1}$), 75.1 (C3$^{F1}$), 73.2 (C2$^{F}$, C2$^{F1}$, C2$^{H}$, C2$^{J}$, C2$^{L}$), 71.8 (C4$^{F1}$), 70.7 (C5$^{E}$, C5$^{E1}$, C5$^{A}$, C5$^{G}$, C3$^{G}$, C5$^{I}$, C3$^{I}$, C5$^{K}$, C3$^{K}$, C5$^{A1}$), 69.9 (C5$^{B}$, C4$^{M}$), 69.7 (C3$^{M}$, C3$^{E}$, C3$^{E1}$, C3$^{C}$, C3$^{C1}$, C3$^{A1}$), 69.3 (C5$^{D}$, C5$^{D1}$, C5$^{B1}$, C5$^{M}$, C5$^{C}$, C5$^{C1}$), 69.1 (C3$^{B}$, C3$^{B1}$, C3$^{D}$,

C3$^{D1}$), 68.3 (O<u>C</u>$H_2$ linker1), 68.1 (C3$^{A}$), 67.7 (O<u>C</u>$H_2$ linker), 66.4 (C6$^{C}$, C6$^{C1}$), 59.5 (C6$^{A}$, C6$^{A1}$, C6$^{E}$, C6$^{E1}$, C6$^{G}$, C6$^{I}$, C6$^{K}$), 57.9 (C2$^{A}$, C2$^{A1}$, C2$^{C}$, C2$^{C1}$, C2$^{E}$, C2$^{E1}$), 53.3 (C2$^{G}$, C2$^{I}$, C2$^{K}$, C2$^{M}$), 49.6 (C6$^{M}$), 39.5 (2 × N<u>C</u>$H_2$ linker), 35.2 (<u>C</u>$H_2$CONH), 28.0 (<u>C</u>$H_2$ linker, 2 × <u>C</u>$H_2$ linker1), 26.3 (<u>C</u>$H_2$ linker), 22.6 (<u>C</u>$H_2$ linker, <u>C</u>$H_2$ linker1), 21.8 (3 × NHCO<u>C</u>$H_3$), 21.1 (<u>C</u>$H_2$-triazole). HRMS (ESI-MS): m/z calculated for $C_{137}H_{216}N_{15}O_{136}S_{12}$ [M-21Na+17H]$^{4-}$: 1157.9288; found: 1157.8785.

$^{1}$H NMR (600 MHz, $D_2O$)

| | H1 | H2 | H3 | H4 | H5 | H6 |
|---|---|---|---|---|---|---|
| **A** | 5.14 | 3.25 | 3.89 | 3.74 | 3.89 | 3.89 |
| **A1** | 5.14 | 3.25 | 3.65 | 3.72 | 3.79 | 3.89 |
| **B** | 5.32 | 4.32 | 4.18 | 4.13 | 4.68 | – |
| **B1** | 5.18 | 4.31 | 4.21 | 4.09 | 4.79 | – |
| **C** | 5.46 | 3.29 | 3.62 | 3.78 | 4.01 | 4.41, 4.28 |
| **C1** | 5.40 | 3.29 | 3.66 | 3.78 | 4.01 | 4.41, 4.28 |
| **D/D1** | 5.22 | 4.33 | 4.21 | 4.09 | 4.82 | – |
| **E/E1** | 5.42 | 3.26 | 3.72 | 3.71 | 3.94 | 3.89 |
| **F/H/J/L** | 4.50 | 3.39 | 3.70 | 3.76 | 3.81 | – |
| **F1** | 4.51 | 3.40 | 3.53 | 3.53 | 3.75 | – |
| **G/I/K** | 5.40 | 3.89 | 3.89 | 3.79 | 3.89 | 3.89 |
| **M** | 5.35 | 3.68 | 3.72 | 2.78 | 4.09 | 4.85, 4.59 |

$^{13}$C NMR from HSQC (151 MHz, $D_2O$)

| | C1 | C2 | C3 | C4 | C5 | C6 |
|---|---|---|---|---|---|---|
| **A** | 96.9 | 57.9 | 68.1 | 76.5 | 70.7 | 59.5 |
| **A1** | 96.9 | 57.9 | 69.7 | 76.5 | 70.7 | 59.5 |
| **B** | 97.2 | 76.5 | 69.1 | 75.9 | 69.9 | – |
| **B1** | 99.2 | 76.5 | 69.1 | 75.9 | 69.3 | – |
| **C** | 95.9 | 57.9 | 69.7 | 76.5 | 69.3 | 66.4 |

| | | | | | | |
|---|---|---|---|---|---|---|
| **C1** | 96.5 | 57.9 | 69.7 | 76.5 | 69.3 | 66.4 |
| **D/D1** | 99.2 | 75.9 | 69.1 | 75.9 | 69.3 | – |
| **E/E1** | 96.5 | 57.9 | 69.7 | 76.5 | 70.7 | 59.5 |
| **F/H/J/L** | 102.2 | 73.2 | 77.5 | 76.5 | 76.5 | – |
| **F1** | 102.2 | 73.2 | 75.1 | 71.8 | 76.5 | – |
| **G/I/K** | 96.5 | 53.3 | 70.7 | 76.5 | 70.7 | 59.5 |
| **M** | 97.2 | 53.3 | 69.7 | 69.9 | 69.3 | 49.6 |

**5-Aminopentyl *O*-(β-D-glucopyranosyluronate)-(1→4)-*O*-(2-sulfamino-2-deoxy-α-D-glucopyranosyl)-(1→4)-*O*-(2-*O*-sulfate-α-L-idopyranosyluronate)-(1→4)-*O*-(2-sulfamino-6-*O*-sulfate-2-deoxy-α-D-glucopyranosyl)-(1→4)-*O*-(2-*O*-sulfate-α-L-idopyranosyluronate)-(1→4)-*O*-2-sulfamino-2-deoxy-α-D-glucopyranosyl) -*O*-[*N*-ethyl-3-(1*H*-1,2,3-triazol-4-yl)propanamide]-(1→6)-*N*-(2-acetamido-2-deoxy-α-D-glucopyranosyl) [-(1→4)-*O*-(β-D-glucopyranosyluronate)-(1→4)-*O*-(2-acetamido-2-deoxy-α-D-glucopyranosyl)]$_4$-(1→4)-*O*-(β-D-glucopyranosyluronate)-(1→4)-*O*-(2-sulfamino-2-deoxy-α-D-glucopyranosyl)-(1→4)-*O*-(2-*O*-sulfate-α-L-idopyranosyluronate)-(1→4)-*O*-(2-sulfamino-6-*O*-sulfate-2-deoxy-α-D-glucopyranosyl)-(1→4)-*O*-(2-*O*-sulfate-α-L-idopyranosyluronate)-(1→4)-*O*-2-sulfamino-2-deoxy-α-D-glucopyranoside, sodium salt (13).**

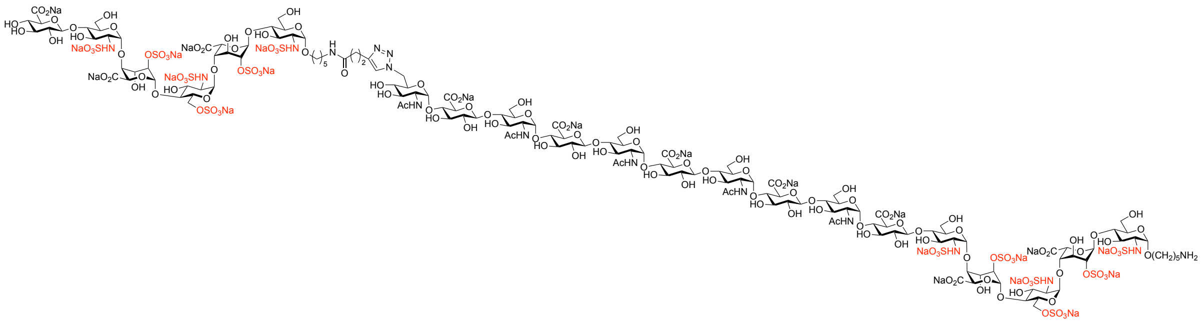

Compound **2** (0.5 mg, 0.27 µmol) and compound **7** (0.9 mg, 0.25 µmol) were subjected to copper (I)-catalyzed azide-alkyne cycloaddition (CuAAC) reaction according to the general procedure to give the title compound **13** as a white powder (0.61 mg, 45%). $^1$H NMR (600 MHz, $D_2O$) δ 7.85 (s, 1H, *CH* triazole), 5.46 (bs, 1H, H1$^C$), 5.44 – 5.40 (m, 2H, H1$^E$, H1$^{E1}$), 5.40 – 5.38 (m, 5H, H1$^G$, H1$^I$, H1$^K$, H1$^M$, H1$^{C1}$), 5.36 – 5.30 (m, 2H, H1$^O$, H1$^B$), 5.22 – 5.20

(m, 2H, H1$^{D}$, H1$^{D1}$), 5.18 (bs, 1H, H1$^{B1}$), 5.16 – 5.12 (m, 2H, H1$^{A}$, H1$^{A1}$), 4.90 – 4.74 (m, 4H, H6a$^{O}$, H5$^{D}$, H5$^{D1}$, H5$^{B1}$), 4.69 (bs, 1H, H5$^{B}$), 4.61 – 4.56 (m, 1H, H6b$^{O}$), 4.54 – 4.47 (m, 6H, H1$^{F1}$, H1$^{F}$, H1$^{H}$, H1$^{J}$, H1$^{L}$, H1$^{N}$), 4.44 – 4.37 (m, 2H, H6a$^{C}$, H6a$^{C1}$), 4.35 – 4.29 (m, 4H, H2$^{D}$, H2$^{D1}$, H2$^{B}$, H2$^{B1}$), 4.29 – 4.25 (m, 2H, H6b$^{C}$, H6b$^{C1}$), 4.23 – 4.17 (m, 4H, H3$^{D}$, H3$^{D1}$, H3$^{B}$, H3$^{B1}$), 4.14 – 4.12 (m, 1H, H4$^{B}$), 4.12 – 4.06 (m, 4H, H4$^{B1}$, H4$^{D}$, H4$^{D1}$, H5$^{O}$), 4.04 – 3.99 (m, 2H, H5$^{C}$, H5$^{C1}$), 3.97 – 3.59 (m, 68H, H2$^{G}$, H2$^{I}$, H2$^{K}$, H2$^{M}$, H6a$^{A}$, H6b$^{A}$, H6a$^{A1}$, H6b$^{A1}$, H6a$^{E}$, H6b$^{E}$, H6a$^{E1}$, H6b$^{E1}$, H6a$^{G}$, H6b$^{G}$, H6a$^{I}$, H6b$^{I}$, H6a$^{K}$, H6b$^{K}$, H6a$^{M}$, H6b$^{M}$, H3$^{A}$, H5$^{E}$, H5$^{E1}$, H5$^{A}$, H5$^{G}$, H3$^{G}$, H5$^{I}$, H3$^{I}$, H5$^{K}$, H3$^{K}$, H5$^{M}$, H3$^{M}$, H5$^{A1}$, H5$^{F}$, H5$^{H}$, H5$^{J}$, H5$^{L}$, H5$^{N}$, H4$^{G}$, H4$^{I}$, H4$^{K}$, H4$^{M}$, H4$^{F}$, H4$^{H}$, H4$^{J}$, H4$^{L}$, H4$^{N}$, H5$^{F1}$, H4$^{C}$, H4$^{C1}$, H4$^{A}$, H4$^{A1}$, H4$^{E}$, H4$^{E1}$, H3$^{F}$, H3$^{H}$, H3$^{J}$, H3$^{L}$, H3$^{N}$, H3$^{O}$, H3$^{E}$, H3$^{E1}$, H3$^{C1}$, H3$^{A1}$, H3$^{C}$, H2$^{O}$, OCH*H* linker, OCH*H* linker1), 3.58 – 3.48 (m, 4H, OCH*H* linker, OCH*H* linker1, H3$^{F1}$, H4$^{F1}$), 3.42 – 3.36 (m, 6H, H2$^{F}$, H2$^{F1}$, H2$^{H}$, H2$^{J}$, H2$^{L}$, H2$^{N}$), 3.32 – 3.23 (m, 6H, H2$^{C}$, H2$^{C1}$, H2$^{E}$, H2$^{E1}$, H2$^{A}$, H2$^{A1}$), 3.15 (t, $J$ = 7.1 Hz, 2H, NC$H_2$ linker1), 3.05 (t, $J$ = 7.2 Hz, 2H, NC$H_2$ linker), 3.01 (t, $J$ = 7.5 Hz, 2H, C$H_2$-triazole), 2.81 – 2.75 (m, 1H, H4$^{O}$), 2.61 (t, $J$ = 7.6 Hz, 2H, C$H_2$CONH), 2.05 (s, 3H, NHCOC$H_3$), 2.05 (s, 9H, 3 × NHCOC$H_3$), 2.02 (s, 3H, NHCOC$H_3$), 1.79 – 1.26 (m, 12H, 6 × C$H_2$ linker). $^{13}$C NMR from HSQC (151 MHz, $D_2O$) δ 124.7 ($\underline{C}$H triazole), 102.2 (C1$^{F}$, C1$^{F1}$, C1$^{H}$, C1$^{J}$, C1$^{L}$, C1$^{N}$), 99.2 (C1$^{D}$, C1$^{D1}$, C1$^{B1}$), 97.2 (C1$^{O}$, C1$^{B}$), 96.9 (C1$^{A}$, C1$^{A1}$), 96.5 (C1$^{E}$, C1$^{E1}$, C1$^{C1}$, C1$^{G}$, C1$^{I}$, C1$^{K}$, C1$^{M}$), 95.9 (C1$^{C}$), 77.5 (C3$^{F}$, C3$^{H}$, C3$^{J}$, C3$^{L}$, C3$^{N}$), 76.5 (C2$^{B}$, C2$^{B1}$, C5$^{F}$, C5$^{H}$, C5$^{J}$, C5$^{L}$, C5$^{N}$, C4$^{G}$, C4$^{I}$, C4$^{K}$, C4$^{M}$, C4$^{F}$, C4$^{H}$, C4$^{J}$, C4$^{L}$, C4$^{N}$, C5$^{F1}$, C4$^{C}$, C4$^{C1}$, C4$^{A}$, C4$^{A1}$, C4$^{E}$, C4$^{E1}$), 75.9 (C2$^{D}$, C2$^{D1}$, C4$^{B}$, C4$^{B1}$, C4$^{D}$, H4$^{D1}$), 75.1 (C3$^{F1}$), 73.2 (C2$^{F}$, C2$^{F1}$, C2$^{H}$, C2$^{J}$, C2$^{L}$, C2$^{N}$), 71.8 (C4$^{F1}$), 70.7 (C5$^{E}$, C5$^{E1}$, C5$^{A}$, C5$^{G}$, C3$^{G}$, C5$^{I}$, C3$^{I}$, C5$^{K}$, C3$^{K}$, C5$^{M}$, C3$^{M}$, C5$^{A1}$), 69.9 (C5$^{B}$, C4$^{O}$), 69.7 (C3$^{O}$, C3$^{E}$, C3$^{E1}$, C3$^{C}$, C3$^{C1}$, C3$^{A1}$), 69.3 (C5$^{D}$, C5$^{D1}$, C5$^{B1}$, C5$^{O}$, C5$^{C}$, C5$^{C1}$), 69.1 (C3$^{B}$, C3$^{B1}$, C3$^{D}$, C3$^{D1}$), 68.3 (O$\underline{C}H_2$ linker1), 68.1 (C3$^{A}$), 67.7 (O$\underline{C}H_2$ linker), 66.4 (C6$^{C}$, C6$^{C1}$), 59.5 (C6$^{A}$, C6$^{A1}$, C6$^{E}$, C6$^{E1}$, C6$^{G}$, C6$^{I}$, C6$^{K}$, C6$^{M}$), 57.9 (C2$^{A}$, C2$^{A1}$, C2$^{C}$, C2$^{C1}$, C2$^{E}$, C2$^{E1}$), 53.3 (C2$^{G}$, C2$^{I}$, C2$^{K}$, C2$^{M}$, C2$^{O}$), 49.6 (C6$^{O}$), 39.5 (2 × N$\underline{C}H_2$ linker), 35.2 ($\underline{C}H_2$CONH), 28.0 ($\underline{C}H_2$ linker, 2 × $\underline{C}H_2$ linker1), 26.3 ($\underline{C}H_2$ linker), 22.6 ($\underline{C}H_2$ linker, $\underline{C}H_2$ linker1), 21.8 (3 × NHCO$\underline{C}H_3$), 21.1 ($\underline{C}H_2$-triazole). ESI-MS: m/z calculated for $C_{151}H_{236}N_{16}O_{147}S_{12}$ [M-22Na+18H]$^{4-}$: 1252.4547; found: 1252.4438.

$^{1}$H NMR (600 MHz, $D_2O$)

| | H1 | H2 | H3 | H4 | H5 | H6 |
|---|---|---|---|---|---|---|
| **A** | 5.14 | 3.25 | 3.89 | 3.74 | 3.89 | 3.89 |
| **A1** | 5.14 | 3.25 | 3.65 | 3.72 | 3.79 | 3.89 |
| **B** | 5.32 | 4.32 | 4.18 | 4.13 | 4.68 | – |
| **B1** | 5.18 | 4.31 | 4.21 | 4.09 | 4.79 | – |
| **C** | 5.46 | 3.29 | 3.62 | 3.78 | 4.01 | 4.41, 4.28 |
| **C1** | 5.40 | 3.29 | 3.66 | 3.78 | 4.01 | 4.41, 4.28 |
| **D/D1** | 5.22 | 4.33 | 4.21 | 4.09 | 4.82 | – |
| **E/E1** | 5.42 | 3.26 | 3.72 | 3.71 | 3.94 | 3.89 |
| **F/H/J/L/N** | 4.50 | 3.39 | 3.70 | 3.76 | 3.81 | – |
| **F1** | 4.51 | 3.40 | 3.53 | 3.53 | 3.75 | – |
| **G/I/K/M** | 5.40 | 3.89 | 3.89 | 3.79 | 3.89 | 3.89 |
| **O** | 5.35 | 3.68 | 3.72 | 2.78 | 4.09 | 4.85, 4.59 |

$^{13}C$ NMR from HSQC (151 MHz, $D_2O$)

| | C1 | C2 | C3 | C4 | C5 | C6 |
|---|---|---|---|---|---|---|
| **A** | 96.9 | 57.9 | 68.1 | 76.5 | 70.7 | 59.5 |
| **A1** | 96.9 | 57.9 | 69.7 | 76.5 | 70.7 | 59.5 |
| **B** | 97.2 | 76.5 | 69.1 | 75.9 | 69.9 | – |
| **B1** | 99.2 | 76.5 | 69.1 | 75.9 | 69.3 | – |
| **C** | 95.9 | 57.9 | 69.7 | 76.5 | 69.3 | 66.4 |
| **C1** | 96.5 | 57.9 | 69.7 | 76.5 | 69.3 | 66.4 |
| **D/D1** | 99.2 | 75.9 | 69.1 | 75.9 | 69.3 | – |
| **E/E1** | 96.5 | 57.9 | 69.7 | 76.5 | 70.7 | 59.5 |
| **F/H/J/L/N** | 102.2 | 73.2 | 77.5 | 76.5 | 76.5 | – |
| **F1** | 102.2 | 73.2 | 75.1 | 71.8 | 76.5 | – |
| **G/I/K/M** | 96.5 | 53.3 | 70.7 | 76.5 | 70.7 | 59.5 |
| **O** | 97.2 | 53.3 | 69.7 | 69.9 | 69.3 | 49.6 |

**5-Aminopentyl *O*-(β-D-glucopyranosyluronate)-(1→4)-*O*-(2-sulfamino-2-deoxy-α-D-glucopyranosyl)-(1→4)-*O*-(2-*O*-sulfate-α-L-idopyranosyluronate)-(1→4)-*O*-(2-sulfamino-6-*O*-sulfate-2-deoxy-α-D-glucopyranosyl)-(1→4)-*O*-(2-*O*-sulfate-α-L-idopyranosyluronate)-(1→4)-*O*-2-sulfamino-2-deoxy-α-D-glucopyranosyl) -*O*-[*N*-ethyl-3-(1*H*-1,2,3-triazol-4-yl)propanamide]-(1→6)-*N*-(2-acetamido-2-deoxy-α-D-glucopyranosyl) [-(1→4)-*O*-(β-D-glucopyranosyluronate)-(1→4)-*O*-(2-acetamido-2-deoxy-α-D-glucopyranosyl)]$_5$-(1→4)-*O*-(β-D-glucopyranosyluronate)-(1→4)-*O*-(2-sulfamino-2-deoxy-α-D-glucopyranosyl)-(1→4)-*O*-(2-*O*-sulfate-α-L-idopyranosyluronate)-(1→4)-*O*-(2-sulfamino-6-*O*-sulfate-2-deoxy-α-D-glucopyranosyl)-(1→4)-*O*-(2-*O*-sulfate-α-L-idopyranosyluronate)-(1→4)-*O*-2-sulfamino-2-deoxy-α-D-glucopyranoside, sodium salt (14).**

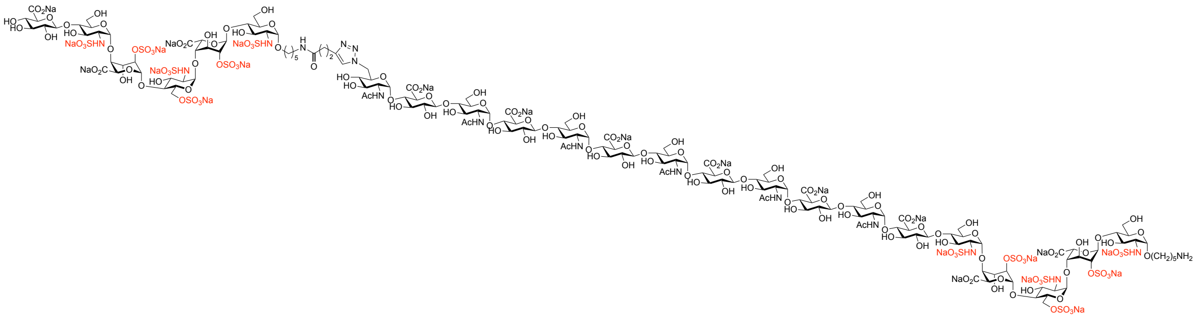

Compound **2** (0.4 mg, 0.2 μmol) and compound **8** (0.4 mg, 0.1 μmol) were subjected to copper (I)-catalyzed azide-alkyne cycloaddition (CuAAC) reaction according to the general procedure to give the title compound **14** as a white powder (0.30 mg, 51%). $^1$H NMR (600 MHz, $D_2O$) δ 7.85 (s, 1H, C*H* triazole), 5.46 (bs, 1H, H1$^C$), 5.44 – 5.40 (m, 2H, H1$^E$, H1$^{E1}$), 5.40 – 5.38 (m, 6H, H1$^G$, H1$^I$, H1$^K$, H1$^M$, H1$^O$, H1$^{C1}$), 5.36 – 5.30 (m, 2H, H1$^Q$, H1$^B$), 5.22 – 5.20 (m, 2H, H1$^D$, H1$^{D1}$), 5.18 (bs, 1H, H1$^{B1}$), 5.16 – 5.12 (m, 2H, H1$^A$, H1$^{A1}$), 4.90 – 4.74 (m, 4H, H6a$^Q$, H5$^D$, H5$^{D1}$, H5$^{B1}$), 4.69 (bs, 1H, H5$^B$), 4.61 – 4.56 (m, 1H, H6b$^Q$), 4.54 – 4.47 (m, 7H, H1$^{F1}$, H1$^F$, H1$^H$, H1$^J$, H1$^L$, H1$^N$, H1$^P$), 4.44 – 4.37 (m, 2H, H6a$^C$, H6a$^{C1}$), 4.35 – 4.29 (m, 4H, H2$^D$, H2$^{D1}$, H2$^B$, H2$^{B1}$), 4.29 – 4.25 (m, 2H, H6b$^C$, H6b$^{C1}$), 4.23 – 4.17 (m, 4H, H3$^D$, H3$^{D1}$, H3$^B$, H3$^{B1}$), 4.14 – 4.12 (m, 1H, H4$^B$), 4.12 – 4.06 (m, 4H, H4$^{B1}$, H4$^D$, H4$^{D1}$, H5$^Q$), 4.04 – 3.99 (m, 2H, H5$^C$, H5$^{C1}$), 3.97 – 3.59 (m, 77H, H2$^G$, H2$^I$, H2$^K$, H2$^M$, H2$^O$, H6a$^A$, H6b$^A$, H6a$^{A1}$, H6b$^{A1}$, H6a$^E$, H6b$^E$, H6a$^{E1}$, H6b$^{E1}$, H6a$^G$, H6b$^G$, H6a$^I$, H6b$^I$, H6a$^K$, H6b$^K$, H6a$^M$, H6b$^M$, H6a$^O$, H6b$^O$, H3$^A$, H5$^E$, H5$^{E1}$, H5$^A$, H5$^G$, H3$^G$, H5$^I$, H3$^I$, H5$^K$, H3$^K$, H5$^M$, H3$^M$, H5$^O$, H3$^O$, H5$^{A1}$, H5$^F$, H5$^H$, H5$^J$, H5$^L$,

$H5^N$, $H5^P$, $H4^G$, $H4^I$, $H4^K$, $H4^M$, $H4^O$, $H4^F$, $H4^H$, $H4^J$, $H4^L$, $H4^N$, $H4^P$, $H5^{F1}$, $H4^C$, $H4^{C1}$, $H4^A$, $H4^{A1}$, $H4^E$, $H4^{E1}$, $H3^F$, $H3^H$, $H3^J$, $H3^L$, $H3^N$, $H3^P$, $H3^Q$, $H3^E$, $H3^{E1}$, $H3^{C1}$, $H3^{A1}$, $H3^C$, $H2^Q$, OCH*H* linker, OCH*H* linker1), 3.58 – 3.48 (m, 4H, OCH*H* linker, OCH*H* linker1, $H3^{F1}$, $H4^{F1}$), 3.42 – 3.36 (m, 7H, $H2^F$, $H2^{F1}$, $H2^H$, $H2^J$, $H2^L$, $H2^N$, $H2^P$), 3.32 – 3.23 (m, 6H, $H2^C$, $H2^{C1}$, $H2^E$, $H2^{E1}$, $H2^A$, $H2^{A1}$), 3.15 (t, $J$ = 7.1 Hz, 2H, $NCH_2$ linker1), 3.05 (t, $J$ = 7.2 Hz, 2H, $NCH_2$ linker), 3.01 (t, $J$ = 7.5 Hz, 2H, $CH_2$-triazole), 2.81 – 2.75 (m, 1H, $H4^Q$), 2.61 (t, $J$ = 7.6 Hz, 2H, $CH_2CONH$), 2.05 (s, 3H, $NHCOCH_3$), 2.05 (s, 6H, 2 × $NHCOCH_3$), 2.02 (s, 3H, $NHCOCH_3$), 1.79 – 1.26 (m, 12H, 6 × $CH_2$ linker). $^{13}C$ NMR from HSQC (151 MHz, $D_2O$) δ 124.7 ($\underline{C}H$ triazole), 102.2 ($C1^F$, $C1^{F1}$, $C1^H$, $C1^J$, $C1^L$, $C1^N$, $C1^P$), 99.2 ($C1^D$, $C1^{D1}$, $C1^{B1}$), 97.2 ($C1^Q$, $C1^B$), 96.9 ($C1^A$, $C1^{A1}$), 96.5 ($C1^E$, $C1^{E1}$, $C1^{C1}$, $C1^G$, $C1^I$, $C1^K$, $C1^M$, $C1^O$), 95.9 ($C1^C$), 77.5 ($C3^F$, $C3^H$, $C3^J$, $C3^L$, $C3^N$, $C3^P$), 76.5 ($C2^B$, $C2^{B1}$, $C5^F$, $C5^H$, $C5^J$, $C5^L$, $C5^N$, $C5^P$, $C4^G$, $C4^I$, $C4^K$, $C4^M$, $C4^O$, $C4^F$, $C4^H$, $C4^J$, $C4^L$, $C4^N$, $C4^P$, $C5^{F1}$, $C4^C$, $C4^{C1}$, $C4^A$, $C4^{A1}$, $C4^E$, $C4^{E1}$), 75.9 ($C2^D$, $C2^{D1}$, $C4^B$, $C4^{B1}$, $C4^D$, $H4^{D1}$), 75.1 ($C3^{F1}$), 73.2 ($C2^F$, $C2^{F1}$, $C2^H$, $C2^J$, $C2^L$, $C2^N$, $C2^P$), 71.8 ($C4^{F1}$), 70.7 ($C5^E$, $C5^{E1}$, $C5^A$, $C5^G$, $C3^G$, $C5^I$, $C3^I$, $C5^K$, $C3^K$, $C5^M$, $C3^M$, $C5^O$, $C3^O$, $C5^{A1}$), 69.9 ($C5^B$, $C4^Q$), 69.7 ($C3^Q$, $C3^E$, $C3^{E1}$, $C3^C$, $C3^{C1}$, $C3^{A1}$), 69.3 ($C5^D$, $C5^{D1}$, $C5^{B1}$, $C5^Q$, $C5^C$, $C5^{C1}$), 69.1 ($C3^B$, $C3^{B1}$, $C3^D$, $C3^{D1}$), 68.3 ($O\underline{C}H_2$ linker1), 68.1 ($C3^A$), 67.7 ($O\underline{C}H_2$ linker), 66.4 ($C6^C$, $C6^{C1}$), 59.5 ($C6^A$, $C6^{A1}$, $C6^E$, $C6^{E1}$, $C6^G$, $C6^I$, $C6^K$, $C6^M$, $C6^O$), 57.9 ($C2^A$, $C2^{A1}$, $C2^C$, $C2^{C1}$, $C2^E$, $C2^{E1}$), 53.3 ($C2^G$, $C2^I$, $C2^K$, $C2^M$, $C2^O$, $C2^Q$), 49.6 ($C6^Q$), 39.5 (2 × $N\underline{C}H_2$ linker), 35.2 ($\underline{C}H_2CONH$), 28.0 ($\underline{C}H_2$ linker, 2 × $\underline{C}H_2$ linker1), 26.3 ($\underline{C}H_2$ linker), 22.6 ($\underline{C}H_2$ linker, $\underline{C}H_2$ linker1), 21.8 (3 × $NHCO\underline{C}H_3$), 21.1 ($\underline{C}H_2$-triazole). ESI-MS: m/z calculated for $C_{165}H_{257}N_{17}O_{158}S_{12}$ [M-23Na+19H]$^{4-}$: 1347.2325; found: 1347.2143.

$^1H$ NMR (600 MHz, $D_2O$)

| | H1 | H2 | H3 | H4 | H5 | H6 |
|---|---|---|---|---|---|---|
| **A** | 5.14 | 3.25 | 3.89 | 3.74 | 3.89 | 3.89 |
| **A1** | 5.14 | 3.25 | 3.65 | 3.72 | 3.79 | 3.89 |
| **B** | 5.32 | 4.32 | 4.18 | 4.13 | 4.68 | – |
| **B1** | 5.18 | 4.31 | 4.21 | 4.09 | 4.79 | – |
| **C** | 5.46 | 3.29 | 3.62 | 3.78 | 4.01 | 4.41, 4.28 |

| | | | | | | |
|---|---|---|---|---|---|---|
| **C1** | 5.40 | 3.29 | 3.66 | 3.78 | 4.01 | 4.41, 4.28 |
| **D/D1** | 5.22 | 4.33 | 4.21 | 4.09 | 4.82 | – |
| **E/E1** | 5.42 | 3.26 | 3.72 | 3.71 | 3.94 | 3.89 |
| **F/H/J/L/N/P** | 4.50 | 3.39 | 3.70 | 3.76 | 3.81 | – |
| **F1** | 4.51 | 3.40 | 3.53 | 3.53 | 3.75 | – |
| **G/I/K/M/O** | 5.40 | 3.89 | 3.89 | 3.79 | 3.89 | 3.89 |
| **Q** | 5.35 | 3.68 | 3.72 | 2.78 | 4.09 | 4.85, 4.59 |

$^{13}C$ NMR from HSQC (151 MHz, $D_2O$)

| | **C1** | **C2** | **C3** | **C4** | **C5** | **C6** |
|---|---|---|---|---|---|---|
| **A** | 96.9 | 57.9 | 68.1 | 76.5 | 70.7 | 59.5 |
| **A1** | 96.9 | 57.9 | 69.7 | 76.5 | 70.7 | 59.5 |
| **B** | 97.2 | 76.5 | 69.1 | 75.9 | 69.9 | – |
| **B1** | 99.2 | 76.5 | 69.1 | 75.9 | 69.3 | – |
| **C** | 95.9 | 57.9 | 69.7 | 76.5 | 69.3 | 66.4 |
| **C1** | 96.5 | 57.9 | 69.7 | 76.5 | 69.3 | 66.4 |
| **D/D1** | 99.2 | 75.9 | 69.1 | 75.9 | 69.3 | – |
| **E/E1** | 96.5 | 57.9 | 69.7 | 76.5 | 70.7 | 59.5 |
| **F/H/J/L/N/P** | 102.2 | 73.2 | 77.5 | 76.5 | 76.5 | – |
| **F1** | 102.2 | 73.2 | 75.1 | 71.8 | 76.5 | – |
| **G/I/K/M/O** | 96.5 | 53.3 | 70.7 | 76.5 | 70.7 | 59.5 |
| **Q** | 97.2 | 53.3 | 69.7 | 69.9 | 69.3 | 49.6 |

**5-Aminopentyl *O*[-(β-D-glucopyranosyluronate)-(1→4)-*O*-(2-sulfamino-2-deoxy-α-D-glucopyranosyl)-(1→4)-*O*-(2-*O*-sulfate-α-L-idopyranosyluronate)-(1→4)-*O*-(2-sulfamino-6-*O*-sulfate-2-deoxy-α-D-glucopyranosyl)-(1→4)-*O*-(2-*O*-sulfate-α-L-idopyranosyluronate)-(1→4)-*O*-2-sulfamino-2-deoxy-α-D-glucopyranosyl) -*O*-[*N*-ethyl-3-(1*H*-1,2,3-triazol-4-yl)propanamide]-(1→6)-*N*-(2-acetamido-2-deoxy-α-D-glucopyranosyl)-(1→4)-*O*]$_2$-(β-D-glucopyranosyluronate)-(1→4)-*O*-(2-sulfamino-2-deoxy-α-D-glucopyranosyl)-(1→4)-*O*-(2-*O*-sulfate-α-L-idopyranosyluronate)- (1→4)-*O*-(2-sulfamino-6-*O*-sulfate-2-deoxy-α-D-glucopyranosyl)-(1→4)-*O*-(2-*O*-sulfate-α-L-idopyranosyluronate)-(1→4)-*O*-2-sulfamino-2-deoxy-α-D-glucopyranoside, sodium salt (31).**

Compound **9** (0.7 mg, 0.2 μmol) was subjected to installation of alkyne moiety reaction according to the general procedure and the resulting compound **28** and compound **3** (0.6 mg,

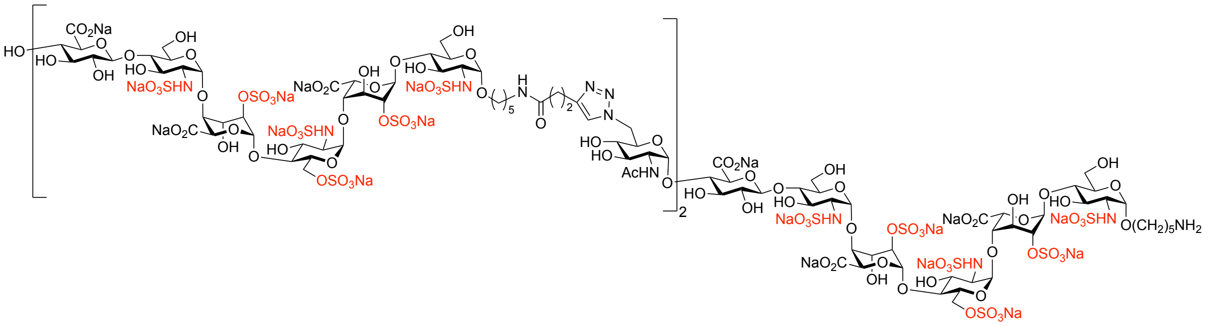

0.3 μmol) were subjected to copper (I)-catalyzed azide-alkyne cycloaddition (CuAAC) reaction according to the general procedure to give the title compound **31** as a white powder (0.53 mg, 50% over two steps). $^1$H NMR (600 MHz, $D_2O$) δ 7.83 (s, 2H, C*H* triazole), 5.46 – 5.26 (m, 9H, $H1^{C}$, $H1^{E}$, $H1^{E1}$, $H1^{E2}$, $H1^{C1}$, $H1^{C2}$, $H1^{G}$, $H1^{G1}$, $H1^{B}$), 5.26 – 5.07 (m, 8H, $H1^{D}$, $H1^{D1}$, $H1^{D2}$, $H1^{B1}$, $H1^{B2}$, $H1^{A}$, $H1^{A1}$, $H1^{A2}$), 4.91 – 4.63 (m, 8H, $H6a^{G}$, $H6a^{G1}$, $H5^{D}$, $H5^{D1}$, $H5^{D2}$, $H5^{B}$, $H5^{B1}$, $H5^{B2}$), 4.58 – 4.51 (m, 1H, $H6b^{G}$, $H6b^{G1}$), 4.51 – 4.44 (m, 2H, $H1^{F}$, $H1^{F1}$, $H1^{F2}$), 4.43 – 4.34 (m, 3H, $H6a^{C}$, $H6a^{C1}$, $H6a^{C2}$), 4.34 – 4.22 (m, 9H, $H2^{D}$, $H2^{D1}$, $H2^{D2}$, $H2^{B}$, $H2^{B1}$, $H2^{B2}$, $H6b^{C}$, $H6b^{C1}$, $H6b^{C2}$), 4.22 – 4.14 (m, 6H, $H3^{D}$, $H3^{D1}$, $H3^{D2}$, $H3^{B}$, $H3^{B1}$, $H3^{B2}$), 4.13 – 4.02 (m, 8H, $H4^{B}$, $H4^{B1}$, $H4^{B2}$, $H4^{D}$, $H4^{D1}$, $H4^{D2}$, $H5^{G}$, $H5^{G1}$), 4.02 – 3.95 (m, 3H, $H5^{C}$, $H5^{C1}$, $H5^{C2}$), 3.95 – 3.56 (m, 50H, $H6a^{A}$, $H6b^{A}$, $H6a^{A1}$, $H6b^{A1}$, $H6a^{A2}$, $H6b^{A2}$, $H6a^{E}$, $H6b^{E}$, $H6a^{E1}$, $H6b^{E1}$, $H6a^{E2}$, $H6b^{E2}$, $H3^{A}$, $H5^{E}$, $H5^{E1}$, $H5^{E2}$, $H5^{A}$, $H5^{A1}$, $H5^{A2}$, $H5^{F}$, $H5^{F1}$, $H4^{F}$, $H4^{F1}$, $H5^{F2}$, $H4^{C}$, $H4^{C1}$, $H4^{C2}$,

$H4^{A}$, $H4^{A1}$, $H4^{A2}$, $H4^{E}$, $H4^{E1}$, $H4^{E2}$, $H3^{F}$, $H3^{F1}$, $H3^{G}$, $H3^{G1}$, $H3^{E}$, $H3^{E1}$, $H3^{E2}$, $H3^{C1}$, $H3^{C2}$, $H3^{A1}$, $H3^{A2}$, $H3^{C}$, $H2^{G}$, $H2^{G1}$, OCH*H* linker, OCH*H* linker1, OCH*H* linker2), 3.56 – 3.44 (m, 5H, OCH*H* linker, OCH*H* linker1, OCH*H* linker2, $H3^{F2}$, $H4^{F2}$), 3.40 – 3.34 (m, 3H, $H2^{F}$, $H2^{F1}$, $H2^{F2}$), 3.28 – 3.18 (m, 9H, $H2^{C}$, $H2^{C1}$, $H2^{C2}$, $H2^{E}$, $H2^{E1}$, $H2^{E2}$, $H2^{A}$, $H2^{A1}$, $H2^{A2}$), 3.12 (t, $J$ = 7.1 Hz, 4H, N$CH_2$ linker1, N$CH_2$ linker2), 3.01 (t, $J$ = 7.2 Hz, 2H, N$CH_2$ linker), 2.97 (t, $J$ = 7.5 Hz, 4H, 2 × $CH_2$-triazole), 2.76 – 2.68 (m, 2H, $H4^{G}$, $H4^{G1}$), 2.58 (t, $J$ = 7.6 Hz, 4H, 2 × $CH_2$CONH), 2.02 (s, 6H, 2 × NHCO$CH_3$), 1.79 – 1.29 (m, 18H, 9 × $CH_2$ linker). $^{13}$C NMR from HSQC (151 MHz, $D_2O$) δ 124.7 (<u>C</u>H triazole), 102.2 ($C1^{F}$, $C1^{F1}$, $C1^{F2}$), 99.2 ($C1^{D}$, $C1^{D1}$, $C1^{D2}$, $C1^{B1}$, $C1^{B2}$), 97.3 ($C1^{G}$, $C1^{G1}$, $C1^{B}$), 97.0 ($C1^{A}$, $C1^{A1}$, $C1^{A2}$), 96.5 ($C1^{E}$, $C1^{E1}$, $C1^{E2}$, $C1^{C1}$, $C1^{C2}$), 95.9 ($C1^{C}$), 77.5 ($C4^{A}$, $C4^{A1}$, $C4^{A2}$, $C4^{E}$, $C4^{E1}$, $C4^{E2}$, $C3^{F}$, $C3^{F1}$), 76.7 ($C2^{B}$, $C2^{B1}$, $C2^{B2}$, $C5^{F}$, $C5^{F1}$, $C5^{F2}$), 75.9 ($C2^{D}$, $C2^{D1}$, $C2^{D2}$, $C4^{B}$, $C4^{B1}$, $C4^{B2}$, $C4^{D}$, $C4^{D1}$, $C4^{D2}$, $C4^{F}$, $C4^{F1}$, $C5^{F1}$, $C5^{F2}$, $C4^{C}$, $C4^{C1}$, $C4^{C2}$), 75.2 ($C3^{F2}$), 73.2 ($C2^{F}$, $C2^{F1}$, $C2^{F2}$), 71.9 ($C4^{F2}$), 70.9 ($C5^{A2}$), 70.7 ($C5^{E}$, $C5^{E1}$, $C5^{E1}$, $C5^{A}$, $C5^{A1}$), 70.0 ($C5^{B}$, $C5^{B1}$, $C4^{G}$, $C4^{G1}$), 69.7 ($C3^{G}$, $C3^{G1}$, $C3^{E}$, $C3^{E1}$, $C3^{E2}$, $C3^{C}$, $C3^{C1}$, $C3^{C2}$, $C3^{A2}$), 69.4 ($C5^{D}$, $C5^{D1}$, $C5^{D2}$, $C5^{B2}$, $C3^{B}$, $C3^{B1}$, $C3^{B2}$, $C3^{D}$, $C3^{D1}$, $C3^{D2}$, $C5^{G}$, $C5^{G1}$, $C5^{C}$, $C5^{C1}$, $C5^{C2}$), 68.4 (O<u>C</u>$H_2$ linker1&2), 68.2 ($C3^{A}$, $C3^{A1}$, $C3^{A2}$), 67.6 (O<u>C</u>$H_2$ linker), 66.4 ($C6^{C}$, $C6^{C1}$, $C6^{C2}$), 59.7 ($C6^{A}$, $C6^{A1}$, $C6^{A2}$, $C6^{E}$, $C6^{E1}$, $C6^{E2}$), 57.9 ($C2^{A}$, $C2^{A1}$, $C2^{A2}$, $C2^{C}$, $C2^{C1}$, $C2^{C2}$, $C2^{E}$, $C2^{E1}$, $C2^{E2}$), 53.5 ($C2^{G}$, $C2^{G1}$), 49.6 ($C6^{G}$, $C6^{G1}$), 39.5 (3 × N<u>C</u>$H_2$ linker), 35.4 (2 × <u>C</u>$H_2$CONH), 28.2 (6 × <u>C</u>$H_2$ linker), 26.3 (<u>C</u>$H_2$ linker), 22.7 (2 × <u>C</u>$H_2$ linker), 22.0 (2 × NHCO<u>C</u>$H_3$), 21.4 (2 × <u>C</u>$H_2$-triazole). ESI-MS: m/z calculated for $C_{149}H_{238}N_{20}O_{157}S_{18}$ $[M-27Na+23H]^{4-}$: 1348.9071; found: 1348.8879.

$^{1}$H NMR (600 MHz, $D_2O$)

| | **H1** | **H2** | **H3** | **H4** | **H5** | **H6** |
|---|---|---|---|---|---|---|
| **A** | 5.11 | 3.22 | 3.86 | 3.71 | 3.87 | 3.87 |
| **A1/A2** | 5.11 | 3.22 | 3.62 | 3.68 | 3.76 | 3.87 |
| **B** | 5.30 | 4.30 | 4.15 | 4.10 | 4.65 | – |
| **B1/B2** | 5.15 | 4.28 | 4.18 | 4.06 | 4.80 | – |
| **C** | 5.44 | 3.26 | 3.59 | 3.75 | 3.98 | 4.37, 4.24 |
| **C1/C2** | 5.37 | 3.26 | 3.63 | 3.75 | 3.98 | 4.37, 4.24 |

| | | | | | | |
|---|---|---|---|---|---|---|
| **D/D1/D2** | 5.19 | 4.30 | 4.18 | 4.06 | 4.80 | – |
| **E/E1/E2** | 5.39 | 3.23 | 3.70 | 3.68 | 3.91 | 3.87 |
| **F/F1** | 4.48 | 3.36 | 3.67 | 3.72 | 3.79 | – |
| **F2** | 4.48 | 3.36 | 3.50 | 3.50 | 3.72 | – |
| **G/G1** | 5.32 | 3.65 | 3.70 | 2.72 | 4.08 | 4.83, 4.55 |

$^{13}C$ NMR from HSQC (151 MHz, $D_2O$)

| | **C1** | **C2** | **C3** | **C4** | **C5** | **C6** |
|---|---|---|---|---|---|---|
| **A** | 97.0 | 57.9 | 68.2 | 77.5 | 70.7 | 59.7 |
| **A1/A2** | 97.0 | 57.9 | 69.7 | 77.5 | 70.9 | 59.7 |
| **B** | 97.3 | 76.7 | 69.4 | 75.9 | 70.0 | – |
| **B1/B2** | 99.2 | 76.7 | 69.4 | 75.9 | 69.4 | – |
| **C** | 95.9 | 57.9 | 69.7 | 75.9 | 69.4 | 66.4 |
| **C1/C2** | 96.5 | 57.9 | 69.7 | 75.9 | 69.4 | 66.4 |
| **D/D1/D2** | 99.2 | 75.9 | 69.4 | 75.9 | 69.4 | – |
| **E/E1/E2** | 96.5 | 57.9 | 69.7 | 77.5 | 70.7 | 59.7 |
| **F/F1** | 102.2 | 73.2 | 77.5 | 75.9 | 76.7 | – |
| **F1** | 102.2 | 73.2 | 75.2 | 71.9 | 75.9 | – |
| **G/G1** | 97.3 | 53.5 | 69.7 | 70.0 | 69.4 | 49.6 |

**5-Aminopentyl *O*[-(β-D-glucopyranosyluronate)-(1→4)-*O*-(2-sulfamino-2-deoxy-α-D-glucopyranosyl)-(1→4)-*O*-(2-*O*-sulfate-α-L-idopyranosyluronate)-(1→4)-*O*-(2-sulfamino-6-*O*-sulfate-2-deoxy-α-D-glucopyranosyl)-(1→4)-*O*-(2-*O*-sulfate-α-L-idopyranosyluronate)-(1→4)-*O*-2-sulfamino-2-deoxy-α-D-glucopyranosyl) -*O*-[*N*-ethyl-3-(1*H*-1,2,3-triazol-4-yl)propanamide]-(1→6)-*N*-(2-acetamido-2-deoxy-α-D-glucopyranosyl)-(1→4)-*O*-(β-D-glucopyranosyluronate)-(1→4)-*O*-(2-acetamido-2-deoxy-α-D-glucopyranosyl)-(1→4)-*O*]$_2$-(β-D-glucopyranosyluronate)-(1→4)-*O*-(2-sulfamino-2-**

**deoxy-α-D-glucopyranosyl)-(1→4)-*O*-(2-*O*-sulfate-α-L-idopyranosyluronate)-(1→4)-*O*-(2-sulfamino-6-*O*-sulfate-2-deoxy-α-D-glucopyranosyl)-(1→4)-*O*-(2-*O*-sulfate-α-L-idopyranosyluronate)-(1→4)-*O*-2-sulfamino-2-deoxy-α-D-glucopyranoside, sodium salt (32).**

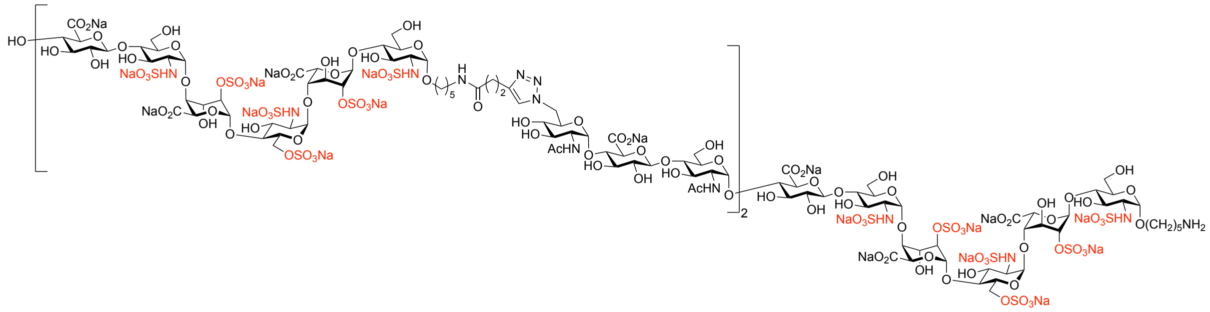

Compound **10** (1.0 mg, 0.2 µmol) was subjected to installation of alkyne moiety reaction according to the general procedure and the resulting compound **29** and compound **4** (0.7 mg, 0.3 µmol) were subjected to copper (I)-catalyzed azide-alkyne cycloaddition (CuAAC) reaction according to the general procedure to give the title compound **32** as a white powder (0.76 mg, 48% over two steps). $^{1}$H NMR (600 MHz, $D_2O$) δ 7.82 (s, 2H, C*H* triazole), 5.43 (bs, 1H, H1$^{C}$), 5.41 – 5.34 (m, 7H, H1$^{E}$, H1$^{E1}$, H1$^{E2}$, H1$^{G}$, H1$^{G1}$, H1$^{C1}$, H1$^{C2}$), 5.31 (d, *J* = 3.3 Hz, 2H, H1$^{I}$, H1$^{I1}$), 5.28 (bs, 1H, H1$^{B}$), 5.20 – 5.16 (m, 3H, H1$^{D}$, H1$^{D1}$, H1$^{D2}$), 5.14 (bs, 2H, H1$^{B1}$, H1$^{B2}$), 5.12 – 5.10 (m, 3H, H1$^{A}$, H1$^{A1}$, H1$^{A2}$), 4.90 – 4.71 (m, 7H, H6a$^{I}$, H6a$^{I1}$, H5$^{D}$, H5$^{D1}$, H5$^{D2}$, H5$^{B1}$, H5$^{B2}$), 4.65 (bs, 1H, H5$^{B}$), 4.60 – 4.52 (m, 2H, H6b$^{I}$, H6b$^{I1}$), 4.50 – 4.44 (m, 5H, H1$^{F}$, H1$^{F1}$, H1$^{F2}$, H1$^{H}$, H1$^{H1}$), 4.40 – 4.35 (m, 3H, H6a$^{C}$, H6a$^{C1}$, H6a$^{C2}$), 4.32 – 4.22 (m, 9H, H2$^{D}$, H2$^{D1}$, H2$^{D2}$, H2$^{B}$, H2$^{B1}$, H2$^{B2}$, H6b$^{C}$, H6b$^{C1}$, H6b$^{C2}$), 4.20 – 4.14 (m, 6H, H3$^{D}$, H3$^{D1}$, H3$^{D2}$, H3$^{B}$, H3$^{B1}$, H3$^{B2}$), 4.11 – 4.03 (m, 8H, H4$^{B}$, H4$^{B1}$, H4$^{B2}$, H4$^{D}$, H4$^{D1}$, H4$^{D2}$, H5$^{I}$, H5$^{I1}$), 4.00 – 3.95 (m, 3H, H5$^{C}$, H5$^{C1}$, H5$^{C2}$), 3.95 – 3.55 (m, 68H, H2$^{G}$, H2$^{G1}$, H6a$^{A}$, H6b$^{A}$, H6a$^{A1}$, H6b$^{A1}$, H6a$^{A2}$, H6b$^{A2}$, H6a$^{E}$, H6b$^{E}$, H6a$^{E1}$, H6b$^{E1}$, H6a$^{E2}$, H6b$^{E2}$, H6a$^{G}$, H6b$^{G}$, H6a$^{G1}$, H6b$^{G1}$, H3$^{A}$, H5$^{E}$, H5$^{E1}$, H5$^{E2}$, H5$^{A}$, H5$^{G}$, H5$^{G1}$, H3$^{G}$, H3$^{G1}$, H5$^{A1}$, H5$^{A2}$, H5$^{F}$, H5$^{F1}$, H5$^{H}$, H5$^{H1}$, H4$^{G}$, H4$^{G1}$, H4$^{F}$, H4$^{F1}$, H4$^{H}$, H4$^{H1}$, H5$^{F2}$, H4$^{C}$, H4$^{C1}$, H4$^{C2}$, H4$^{A}$, H4$^{A1}$, H4$^{A2}$, H4$^{E}$, H4$^{E1}$, H4$^{E2}$, H3$^{F}$, H3$^{F1}$, H3$^{H}$, H3$^{H1}$, H3$^{I}$, H3$^{I1}$, H3$^{E}$, H3$^{E1}$, H3$^{E2}$, H3$^{C1}$, H3$^{C2}$, H3$^{A1}$, H3$^{A2}$, H3$^{C}$, H2$^{I}$, H2$^{I1}$, OCH*H* linker, OCH*H* linker1, OCH*H* linker2), 3.53 – 3.46 (m, 5H, OCH*H* linker, OCH*H* linker1, OCH*H* linker2, H3$^{F2}$, H4$^{F2}$), 3.39 – 3.33 (m, 5H, H2$^{F}$, H2$^{F1}$, H2$^{F2}$, H2$^{H}$, H2$^{H1}$), 3.28 – 3.20 (m, 9H, H2$^{C}$, H2$^{C1}$, H2$^{C2}$, H2$^{E}$, H2$^{E1}$, H2$^{E2}$, H2$^{A}$, H2$^{A1}$, H2$^{A2}$), 3.12 (t, *J* = 7.1 Hz, 4H, NC$H_2$ linker1, NC$H_2$ linker2), 3.03 –

2.96 (m, 6H, N$CH_2$ linker, 2 × $CH_2$-triazole), 2.77 – 2.72 (m, 2H, $H4^{I}$, $H4^{I1}$), 2.58 (t, $J$ = 7.6 Hz, 4H, 2 × $CH_2$CONH), 2.02 (s, 6H, 2 × NHCOC$H_3$), 1.99 (s, 6H, 2 × NHCOC$H_3$), 1.79 – 1.26 (m, 12H, 9 × $CH_2$ linker). $^{13}$C NMR from HSQC (151 MHz, $D_2O$) δ 124.7 (2 × C̲H triazole), 102.2 ($C1^{F}$, $C1^{F1}$, $C1^{F2}$, $C1^{H}$, $C1^{H1}$), 99.2 ($C1^{D}$, $C1^{D1}$, $C1^{D1}$, $C1^{B1}$, $C1^{B2}$), 97.2 ($C1^{I}$, $C1^{I1}$, $C1^{B}$), 96.9 ($C1^{A}$, $C1^{A1}$, $C1^{A2}$), 96.5 ($C1^{E}$, $C1^{E1}$, $C1^{E2}$, $C1^{C1}$, $C1^{C2}$, $C1^{G}$, $C1^{G1}$), 95.9 ($C1^{C}$), 77.5 ($C3^{F}$, $C3^{F1}$, $C3^{H}$, $C3^{H1}$), 76.5 ($C2^{B}$, $C2^{B1}$, $C2^{B2}$, $C5^{F}$, $C5^{F1}$, $C5^{H}$, $C5^{H1}$, $C4^{G}$, $C4^{G1}$, $C4^{F}$, $C4^{F1}$, $C4^{H}$, $C4^{H1}$, $C4^{C}$, $C4^{C1}$, $C4^{C2}$, $C4^{A}$, $C4^{A1}$, $C4^{A2}$, $C4^{E}$, $C4^{E1}$, $C4^{E2}$), 75.9 ($C2^{D}$, $C2^{D1}$, $C2^{D2}$, $C4^{B}$, $C4^{B1}$, $C4^{B2}$, $C4^{D}$, $C4^{D1}$, $C4^{D2}$), 75.1 ($C3^{F2}$), 73.2 ($C2^{F}$, $C2^{F1}$, $C2^{F2}$, $C2^{H}$, $C2^{H1}$), 71.8 ($C4^{F2}$), 70.7 ($C5^{E}$, $C5^{E1}$, $C5^{E2}$, $C5^{A}$, $C5^{G}$, $C5^{G1}$, $C3^{G}$, $C3^{G1}$, $C5^{A1}$, $C5^{A2}$), 69.9 ($C5^{B}$, $C4^{I}$, $C4^{I1}$), 69.7 ($C3^{I}$, $C3^{I1}$, $C3^{E}$, $C3^{E1}$, $C3^{E2}$, $C3^{C}$, $C3^{C1}$, $C3^{C2}$, $C3^{A1}$, $C3^{A2}$), 69.3 ($C5^{D}$, $C5^{D1}$, $C5^{D2}$, $C5^{B1}$, $C5^{B2}$, $C5^{I}$, $C5^{I1}$, $C5^{C}$, $C5^{C1}$, $C5^{C2}$), 69.1 ($C3^{B}$, $C3^{B1}$, $C3^{B2}$, $C3^{D}$, $C3^{D1}$, $C3^{D2}$), 68.3 (OC̲$H_2$ linker1), 68.1 ($C3^{A}$), 67.7 (OC̲$H_2$ linker), 66.4 ($C6^{C}$, $C6^{C1}$, $C6^{C2}$), 59.5 ($C6^{A}$, $C6^{A1}$, $C6^{A2}$, $C6^{E}$, $C6^{E1}$, $C6^{E2}$, $C6^{G}$, $C6^{G1}$), 57.9 ($C2^{A}$, $C2^{A1}$, $C2^{A2}$, $C2^{C}$, $C2^{C1}$, $C2^{C2}$, $C2^{E}$, $C2^{E1}$, $C2^{E2}$), 53.3 ($C2^{G}$, $C2^{G1}$, $C2^{I}$, $C2^{I1}$), 49.6 ($C6^{I}$, $C6^{I1}$), 39.5 (3 × NC̲$H_2$ linker), 35.2 (2 × C̲$H_2$CONH), 28.0 (C̲$H_2$ linker, 2 × C̲$H_2$ linker1, 2 × C̲$H_2$ linker2), 26.3 (C̲$H_2$ linker), 22.6 (C̲$H_2$ linker, C̲$H_2$ linker1, C̲$H_2$ linker2), 21.8 (4 × NHCOC̲$H_3$), 21.1 (2 × C̲$H_2$-triazole). ESI-MS: m/z calculated for $C_{175}H_{296}N_{28}O_{175}S_{18}$ $[M-29Na+8NH_4+17H]^{4-}$: 1541.5038; found: 1541.4449.

$^{1}$H NMR (600 MHz, $D_2O$)

| | **H1** | **H2** | **H3** | **H4** | **H5** | **H6** |
|---|---|---|---|---|---|---|
| **A** | 5.11 | 3.21 | 3.83 | 3.70 | 3.85 | 3.85 |
| **A1/A2** | 5.11 | 3.21 | 3.61 | 3.68 | 3.75 | 3.85 |
| **B** | 5.28 | 4.28 | 4.17 | 4.09 | 4.65 | – |
| **B1/B2** | 5.14 | 4.28 | 4.17 | 4.05 | 4.75 | – |
| **C** | 5.43 | 3.25 | 3.58 | 3.74 | 3.98 | 4.38, 4.23 |
| **C1/C2** | 5.36 | 3.25 | 3.62 | 3.74 | 3.98 | 4.38, 4.23 |
| **D/D1/D2** | 5.18 | 4.28 | 4.17 | 4.05 | 4.79 | – |
| **E/E1/E2** | 5.39 | 3.22 | 3.68 | 3.67 | 3.91 | 3.85 |
| **F/F1/H/H1** | 4.47 | 3.35 | 3.66 | 3.72 | 3.77 | – |

| | | | | | | |
|---|---|---|---|---|---|---|
| **F2** | 4.47 | 3.36 | 3.49 | 3.49 | 3.71 | – |
| **G/G1** | 5.36 | 3.87 | 3.85 | 3.75 | 3.85 | 3.85 |
| **I/I1** | 5.31 | 3.64 | 3.68 | 2.74 | 4.05 | 4.82, 4.54 |

$^{13}C$ NMR from HSQC (151 MHz, $D_2O$)

| | **C1** | **C2** | **C3** | **C4** | **C5** | **C6** |
|---|---|---|---|---|---|---|
| **A** | 96.9 | 57.9 | 68.1 | 76.5 | 70.7 | 59.5 |
| **A1/A2** | 96.9 | 57.9 | 69.7 | 76.5 | 70.7 | 59.5 |
| **B** | 97.2 | 76.5 | 69.1 | 75.9 | 69.9 | – |
| **B1/B2** | 99.2 | 76.5 | 69.1 | 75.9 | 69.3 | – |
| **C** | 95.9 | 57.9 | 69.7 | 76.5 | 69.3 | 66.4 |
| **C1/C2** | 96.5 | 57.9 | 69.7 | 76.5 | 69.3 | 66.4 |
| **D/D1/D2** | 99.2 | 75.9 | 69.1 | 75.9 | 69.3 | – |
| **E/E1/E2** | 96.5 | 57.9 | 69.7 | 76.5 | 70.7 | 59.5 |
| **F/F1/H/H1** | 102.2 | 73.2 | 77.5 | 76.5 | 76.5 | – |
| **F2** | 102.2 | 73.2 | 75.1 | 71.8 | 76.5 | – |
| **G/G1** | 96.5 | 53.3 | 70.7 | 76.5 | 70.7 | 59.5 |
| **I/I1** | 97.2 | 53.3 | 69.7 | 69.9 | 69.3 | 49.6 |

**5-Aminopentyl *O*[-(β-D-glucopyranosyluronate)-(1→4)-*O*-(2-sulfamino-2-deoxy-α-D-glucopyranosyl)-(1→4)-*O*-(2-*O*-sulfate-α-L-idopyranosyluronate)-(1→4)-*O*-(2-sulfamino-6-*O*-sulfate-2-deoxy-α-D-glucopyranosyl)-(1→4)-*O*-(2-*O*-sulfate-α-L-idopyranosyluronate)-(1→4)-*O*-2-sulfamino-2-deoxy-α-D-glucopyranosyl) -*O*-[*N*-ethyl-3-(1*H*-1,2,3-triazol-4-yl)propanamide]-(1→6)-*N*-(2-acetamido-2-deoxy-α-D-glucopyranosyl) [-(1→4)-*O*-(β-D-glucopyranosyluronate)-(1→4)-*O*-(2-acetamido-2-deoxy-α-D-glucopyranosyl)]$_2$-(1→4)-*O*]$_2$-(β-D-glucopyranosyluronate)-(1→4)-*O*-(2-sulfamino-2-deoxy-α-D-glucopyranosyl)-(1→4)-*O*-(2-*O*-sulfate-α-L-idopyranosyluronate)-(1→4)-*O*-(2-sulfamino-6-*O*-sulfate-2-deoxy-α-D-glucopyranosyl)-**

**(1→4)-*O*-(2-*O*-sulfate-α-L-idopyranosyluronate)-(1→4)-*O*-2-sulfamino-2-deoxy-α-D-glucopyranoside, sodium salt (33).**

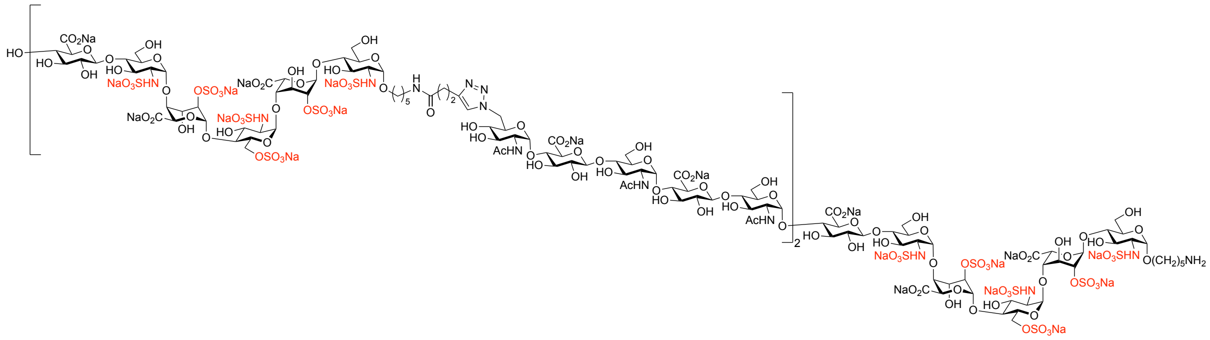

Compound **11** (0.6 mg, 0.1 μmol) was subjected to installation of alkyne moiety reaction according to the general procedure and the resulting compound **30** and compound **5** (0.6 mg, 0.2 μmol) were subjected to copper (I)-catalyzed azide-alkyne cycloaddition (CuAAC) reaction according to the general procedure to give the title compound **33** as a white powder (0.30 mg, 33% over two steps). $^{1}$H NMR (600 MHz, $D_2O$) δ 7.82 (s, 1H, C*H* triazole), 5.43 (bs, 1H, $H1^{C}$), 5.41 – 5.33 (m, 9H, $H1^{E}$, $H1^{E1}$, $H1^{E2}$, $H1^{G}$, $H1^{G1}$, $H1^{I}$, $H1^{I1}$, $H1^{C}$, $H1^{C1}$), 5.33 – 5.28 (m, 3H, $H1^{K}$, $H1^{K1}$, $H1^{B}$), 5.23 – 5.06 (m, 8H, $H1^{D}$, $H1^{D1}$, $H1^{D2}$, $H1^{B1}$, $H1^{B2}$, $H1^{A}$, $H1^{A1}$, $H1^{A2}$), 4.90 – 4.60 (m, 8H, $H6a^{K}$, $H6a^{K1}$, $H5^{D}$, $H5^{D1}$, $H5^{D2}$, $H5^{B1}$, $H5^{B2}$, $H5^{B}$), 4.58 – 4.51 (m, 2H, $H6b^{K}$, $H6b^{K1}$), 4.50 – 4.41 (m, 7H, $H1^{F}$, $H1^{F1}$, $H1^{H}$, $H1^{H1}$, $H1^{J}$, $H1^{J1}$, $H1^{F2}$), 4.41 – 4.34 (m, 3H, $H6a^{C}$, $H6a^{C1}$, $H6a^{C2}$), 4.34 – 4.20 (m, 9H, $H2^{D}$, $H2^{D1}$, $H2^{D2}$, $H2^{B}$, $H2^{B1}$, $H2^{B2}$, $H6b^{C}$, $H6b^{C1}$, $H6b^{C2}$), 4.20 – 4.12 (m, 6H, $H3^{D}$, $H3^{D1}$, $H3^{D2}$, $H3^{B}$, $H3^{B1}$, $H3^{B2}$), 4.12 – 4.01 (m, 8H, $H4^{B}$, $H4^{B1}$, $H4^{B2}$, $H4^{D}$, $H4^{D1}$, $H4^{D2}$, $H5^{K}$, $H5^{K1}$), 4.00 – 3.95 (m, 3H, $H5^{C}$, $H5^{C1}$, $H5^{C2}$), 3.95 – 3.56 (m, 86H, $H2^{G}$, $H2^{G1}$, $H2^{I}$, $H2^{I1}$, $H6a^{A}$, $H6b^{A}$, $H6a^{A1}$, $H6b^{A1}$, $H6a^{A2}$, $H6b^{A2}$, $H6a^{E}$, $H6b^{E}$, $H6a^{E1}$, $H6b^{E1}$, $H6a^{E2}$, $H6b^{E2}$, $H6a^{G}$, $H6b^{G}$, $H6a^{G1}$, $H6b^{G1}$, $H6a^{I}$, $H6b^{I}$, $H6a^{I1}$, $H6b^{I1}$, $H3^{A}$, $H5^{E}$, $H5^{E1}$, $H5^{E2}$, $H5^{A}$, $H5^{G}$, $H5^{G1}$, $H3^{G}$, $H3^{G1}$, $H5^{I}$, $H5^{I1}$, $H3^{I}$, $H3^{I1}$, $H5^{A1}$, $H5^{A2}$, $H5^{F}$, $H5^{F1}$, $H5^{H}$, $H5^{H1}$, $H5^{J}$, $H5^{J1}$, $H4^{G}$, $H4^{G1}$, $H4^{I}$, $H4^{I1}$, $H4^{F}$, $H4^{F1}$, $H4^{H}$, $H4^{H1}$, $H4^{J}$, $H4^{J1}$, $H5^{F2}$, $H4^{C}$, $H4^{C1}$, $H4^{C2}$, $H4^{A}$, $H4^{A1}$, $H4^{A2}$, $H4^{E}$, $H4^{E1}$, $H4^{E2}$, $H3^{F}$, $H3^{F1}$, $H3^{H}$, $H3^{H1}$, $H3^{J}$, $H3^{J1}$, $H3^{K}$, $H3^{K1}$, $H3^{E}$, $H3^{E1}$, $H3^{E2}$, $H3^{C1}$, $H3^{C2}$, $H3^{A1}$, $H3^{A2}$, $H3^{C}$, $H2^{K}$, $H2^{K1}$, OCH*H* linker, OCH*H* linker1, OCH*H* linker2), 3.54 – 3.42 (m, 4H, OCH*H* linker, OCH*H* linker1, OCH*H* linker2, $H3^{F1}$, $H4^{F1}$), 3.41 – 3.29 (m, 7H, $H2^{F}$, $H2^{F1}$, $H2^{F2}$, $H2^{H}$, $H2^{H1}$, $H2^{J}$, $H2^{J1}$), 3.29 – 3.16 (m, 9H, $H2^{C}$, $H2^{C1}$, $H2^{C2}$, $H2^{E}$, $H2^{E1}$, $H2^{E2}$, $H2^{A}$, $H2^{A1}$, $H2^{A2}$), 3.12 (t, $J$ = 7.1 Hz, 4H, $NCH_2$ linker1, $NCH_2$ linker2), 3.02 (t, $J$ = 7.2 Hz, 2H, $NCH_2$

linker), 3.97 (t, $J$ = 7.5 Hz, 4H, 2 × C$H_2$-triazole), 2.76 – 2.68 (m, 2H, H4$^{K}$, H4$^{K1}$), 2.58 (t, $J$ = 7.6 Hz, 4H, 2 × C$H_2$CONH), 2.02 (s, 6H, 2 × NHCOC$H_3$), 2.01 (s, 6H, 2 × NHCOC$H_3$), 1.99 (s, 6H, 2 × NHCOC$H_3$), 1.79 – 1.26 (m, 12H, 6 × C$H_2$ linker). $^{13}$C NMR from HSQC (151 MHz, $D_2O$) δ 124.7 (C̲H triazole), 102.2 (C1$^{F}$, C1$^{F1}$, C1$^{H}$, C1$^{J}$), 99.2 (C1$^{D}$, C1$^{D1}$, C1$^{B1}$), 97.2 (C1$^{K}$, C1$^{B}$), 96.9 (C1$^{A}$, C1$^{A1}$), 96.5 (C1$^{E}$, C1$^{E1}$, C1$^{C1}$, C1$^{G}$, C1$^{I}$), 95.9 (C1$^{C}$), 77.5 (C3$^{F}$, C3$^{H}$, C3$^{J}$), 76.5 (C2$^{B}$, C2$^{B1}$, C5$^{F}$, C5$^{H}$, C5$^{J}$, C4$^{G}$, C4$^{I}$, C4$^{F}$, C4$^{H}$, C4$^{J}$, C5$^{F1}$, C4$^{C}$, C4$^{C1}$, C4$^{A}$, C4$^{A1}$, C4$^{E}$, C4$^{E1}$), 75.9 (C2$^{D}$, C2$^{D1}$, C4$^{B}$, C4$^{B1}$, C4$^{D}$, H4$^{D1}$), 75.1 (C3$^{F1}$), 73.2 (C2$^{F}$, C2$^{F1}$, C2$^{H}$, C2$^{J}$), 71.8 (C4$^{F1}$), 70.7 (C5$^{E}$, C5$^{E1}$, C5$^{A}$, C5$^{G}$, C3$^{G}$, C5$^{I}$, C3$^{I}$, C5$^{A1}$), 69.9 (C5$^{B}$, C4$^{K}$), 69.7 (C3$^{K}$, C3$^{E}$, C3$^{E1}$, C3$^{C}$, C3$^{C1}$, C3$^{A1}$), 69.3 (C5$^{D}$, C5$^{D1}$, C5$^{B1}$, C5$^{K}$, C5$^{C}$, C5$^{C1}$), 69.1 (C3$^{B}$, C3$^{B1}$, C3$^{D}$, C3$^{D1}$), 68.3 (OC̲$H_2$ linker1), 68.1 (C3$^{A}$), 67.7 (OC̲$H_2$ linker), 66.4 (C6$^{C}$, C6$^{C1}$), 59.5 (C6$^{A}$, C6$^{A1}$, C6$^{E}$, C6$^{E1}$, C6$^{G}$, C6$^{I}$), 57.9 (C2$^{A}$, C2$^{A1}$, C2$^{C}$, C2$^{C1}$, C2$^{E}$, C2$^{E1}$), 53.3 (C2$^{G}$, C2$^{I}$, C2$^{K}$), 49.6 (C6$^{K}$), 39.5 (2 × NC̲$H_2$ linker), 35.2 (C̲$H_2$CONH), 28.0 (C̲$H_2$ linker, 2 × C̲$H_2$ linker1), 26.3 (C̲$H_2$ linker), 22.6 (C̲$H_2$ linker, C̲$H_2$ linker1), 21.8 (3 × NHCOC̲$H_3$), 21.1 (C̲$H_2$-triazole). ESI-MS: m/z calculated for $C_{202}H_{311}N_{24}Na_8O_{195}S_{18}$ [M-23Na+19H]$^{4-}$: 1738.4850; found: 1738.4875.

$^{1}$H NMR (600 MHz, $D_2O$)

| | **H1** | **H2** | **H3** | **H4** | **H5** | **H6** |
|---|---|---|---|---|---|---|
| **A** | 5.11 | 3.22 | 3.86 | 3.71 | 3.86 | 3.86 |
| **A1/A2** | 5.11 | 3.22 | 3.62 | 3.69 | 3.76 | 3.86 |
| **B** | 5.29 | 4.28 | 4.15 | 4.10 | 4.65 | – |
| **B1/B1** | 5.15 | 4.28 | 4.18 | 4.06 | 4.76 | – |
| **C** | 5.44 | 3.26 | 3.59 | 3.75 | 3.98 | 4.37, 4.25 |
| **C1/C1** | 5.37 | 3.26 | 3.63 | 3.75 | 3.98 | 4.37, 4.25 |
| **D/D1/D2** | 5.19 | 4.30 | 4.18 | 4.06 | 4.79 | – |
| **E/E1/E2** | 5.39 | 3.23 | 3.68 | 3.68 | 3.91 | 3.86 |
| **F/F1/H/H1/J/J1** | 4.47 | 3.36 | 3.65 | 3.73 | 3.79 | – |
| **F2** | 4.47 | 3.36 | 3.50 | 3.50 | 3.72 | – |
| **G/G1/I/I1** | 5.37 | 3.86 | 3.86 | 3.76 | 3.86 | 3.86 |
| **K/K1** | 5.29 | 3.65 | 3.69 | 2.75 | 4.06 | 4.83, 4.56 |

$^{13}C$ NMR from HSQC (151 MHz, $D_2O$)

| | C1 | C2 | C3 | C4 | C5 | C6 |
|---|---|---|---|---|---|---|
| **A** | 96.9 | 57.9 | 68.1 | 76.5 | 70.7 | 59.5 |
| **A1/A2** | 96.9 | 57.9 | 69.7 | 76.5 | 70.7 | 59.5 |
| **B** | 97.2 | 76.5 | 69.1 | 75.9 | 69.9 | – |
| **B1/B2** | 99.2 | 76.5 | 69.1 | 75.9 | 69.3 | – |
| **C** | 95.9 | 57.9 | 69.7 | 76.5 | 69.3 | 66.4 |
| **C1/C2** | 96.5 | 57.9 | 69.7 | 76.5 | 69.3 | 66.4 |
| **D/D1/D2** | 99.2 | 75.9 | 69.1 | 75.9 | 69.3 | – |
| **E/E1/E2** | 96.5 | 57.9 | 69.7 | 76.5 | 70.7 | 59.5 |
| **F/F1/H/H1/J/J1** | 102.2 | 73.2 | 77.5 | 76.5 | 76.5 | – |
| **F2** | 102.2 | 73.2 | 75.1 | 71.8 | 76.5 | – |
| **G/G1/I/I1** | 96.5 | 53.3 | 70.7 | 76.5 | 70.7 | 59.5 |
| **K/K1** | 97.2 | 53.3 | 69.7 | 69.9 | 69.3 | 49.6 |

**3. Surface plasma resonance (SPR) experiments**

For the preparation of a heparin functionalized sensor chip, CM5 chip (Biacore Inc., GE Healthcare) was first coated with streptavidin by standard amine coupling, followed by immobilization of biotin-heparin (prepared using literature report).[4] Briefly, the surface was activated using freshly mixed N-hydroxysuccinimide (NHS; 100 mM) and 1-(3-dimethylaminopropyl)-ethylcarbodiimide (EDC; 350 mM) (1/1, v/v) in water. Next, streptavidin (50 μg/mL, Invitrogen) in aqueous NaOAc (10 mM, pH 4.5) was passed over the chip surface until a ligand density of approximately 2000 RU was achieved. The remaining NHS-activated esters were quenched by aqueous ethanolamine (1.0 M, pH 8.5). Next, biotin-heparin (50 μg/mL) was passed over one of the flow channels at a flow rate of 10 μL/min for 60 sec resulting in a response of 100 RU. Next, the reference and modified flow cells were washed with three consecutive injections of 60 sec with 2.0 M NaCl. HBS-EP (0.01 M HEPES, 150 mM NaCl, 3 mM EDTA, 0.005% polysorbate 20; pH 7.4) was used as the running buffer for the immobilization, kinetic studies, and inhibition studies. Recombinant human chemokine C-C motif ligand 5 (CCL5, also known as RANTES) was obtained from RayBiotech Life, Inc. Recombinant human chemokine CXC motif ligand 8 (CXCL8, also known as Interleukin-8, IL-8) was obtained from Sino Biological, Inc. For kinetic studies, various concentration of CCL5 or CXCL8 (ranging from 1000 nM to 3.9 nM, 2-fold dilutions) were dissolved in running buffer and flowed over heparin chip (association time: 300 sec and dissociation time: 600 sec) at a flow rate of 30 μL/min and a constant temperature of 25 °C. A 60 sec injection of 2.0 M NaCl at a flow rate of 30 μL/min was used for regeneration and to achieve prior baseline status. To further stabilize the baseline, washing was continued for another 240 sec with running buffer at a flow rate of 30 μL/min. Using Biacore T100 evaluation software, steady-state affinity analysis was performed on the response curves of various CXCL8 concentrations and equilibrium dissociation constant ($K_D$) was calculated (Figure S3). For CCL5, $K_D$ value was determined using fitting response curves of various CCL5 concentrations to a 1:1 Langmuir binding model built in Biacore T100 evaluation software (Figure S4).

For competition assays, CCL5 (100 nM) or CXCL8 (100 nM) alone or in the presence of domain structures (**1**, **9-14**, **31-33**, at a concentration of 20 μM) was flowed over heparin chip. The samples were diluted in running buffer and a flow rate of 30 μL/min was employed for association (150 sec for CCL5 and 100 sec for CXCL8) and dissociation (150 sec for CCL5 and 150 sec for CXCL8) at a constant temperature of 25 °C. A 60 sec injection of 2.0 M NaCl at a flow rate of 30 μL/min was used for regeneration and achieved prior baseline status. In addition, control runs were performed in the presence of unfractionated heparin (UFH) and heparin oligomers (hexamer and octamer). Identified hits (**9**, **31**, **32** for CXCL8; **9-11**, **14**, **31-33** for CCL5) were further screened at various concentrations (ranging from 20 μM to 0.16 μM, 2-fold dilutions) and the $IC_{50}$ values (Figures S5 and S6) were calculated using dose−response equations [nonlinear regression, log(inhibitor) vs response-variable slope (four parameters)] built in Prism software 9 (GraphPad Software, Inc.). Standard error (S.E.) was calculated from 95% CI values using formula, S.D. = [(upper limit-lower limit)/3.92]. All experiments were performed (in duplicate) two times at the minimum.

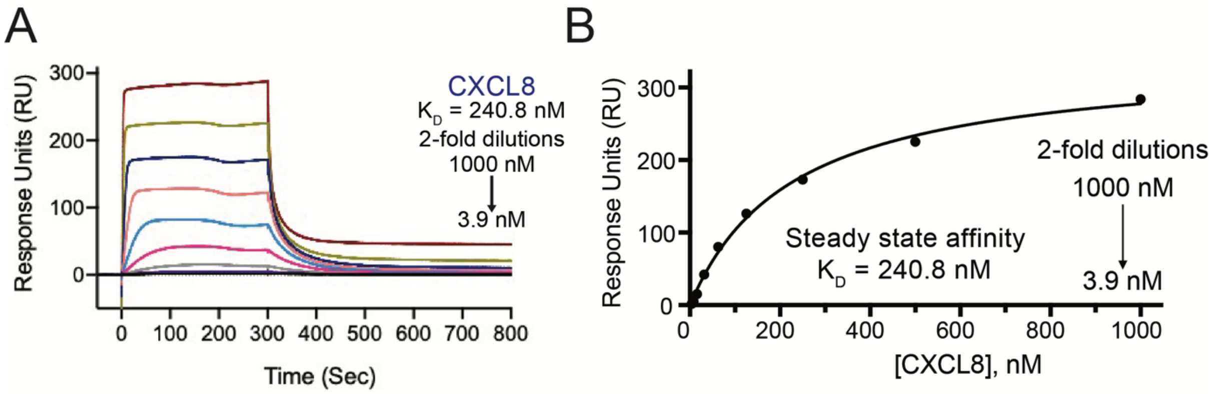

**Figure S3.** SPR data of CXCL8. (A) Sensorgram of CXCL8 showing concentration-dependent binding to immobilized heparin; (B) Steady state affinity curve to calculate equilibrium dissociation constant ($K_D$).

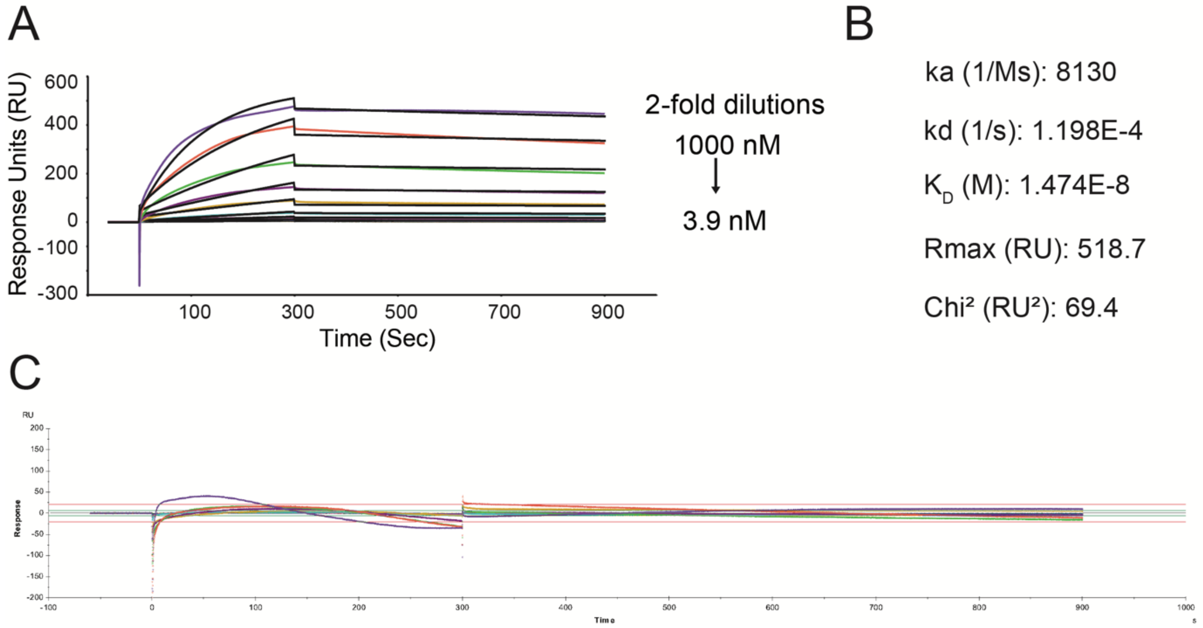

**Figure S4.** SPR data of CCL5. (A) Sensorgram of CCL5 fitted to 1:1 Langmuir binding model; (B) Detailed kinetic parameters obtained from curve fitting; (C) Residue plot of curve fitting various concentrations of CCL5.

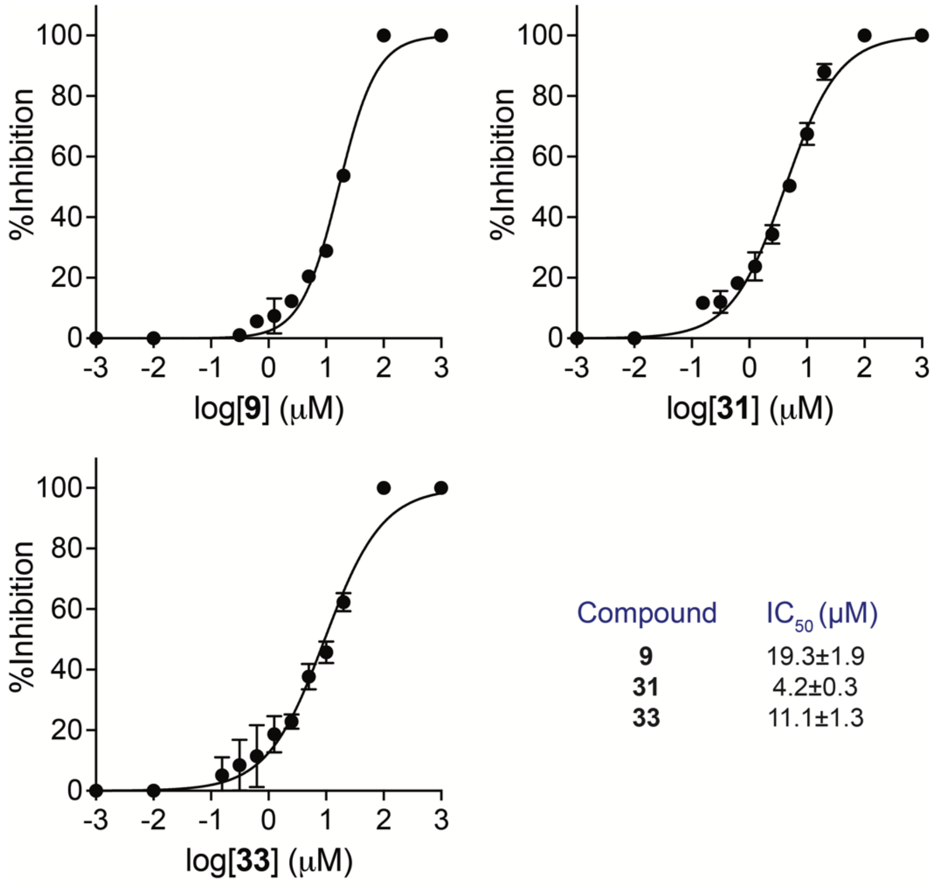

| Compound | $IC_{50}$ (μM) |
|---|---|
| **9** | 19.3±1.9 |
| **31** | 4.2±0.3 |
| **33** | 11.1±1.3 |

**Figure S5**. Inhibition curves of compound **9**, **31** and **33** in a SPR-based competition assays for binding of CXCL8 to heparin-immobilized surface.

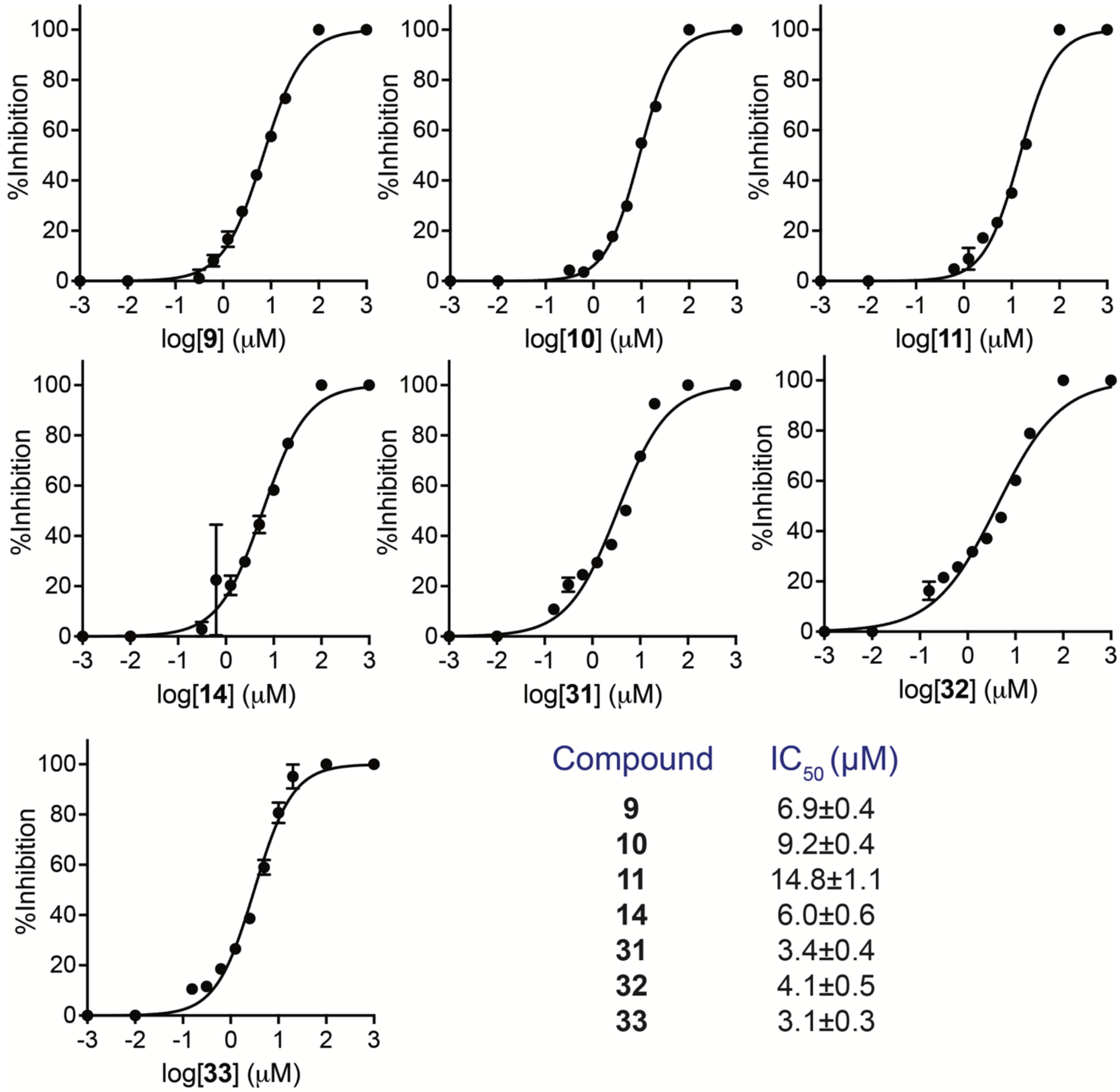

| Compound | $IC_{50}$ (μM) |
|---|---|
| **9** | 6.9±0.4 |
| **10** | 9.2±0.4 |
| **11** | 14.8±1.1 |
| **14** | 6.0±0.6 |
| **31** | 3.4±0.4 |
| **32** | 4.1±0.5 |
| **33** | 3.1±0.3 |

**Figure S6**. Inhibition curves of compound **9**, **10**, **11**, **31**, **32** and **33** in a SPR-based competition assays for binding of CCL5 to heparin-immobilized surface.

### 5. NMR spectra

[1]H NMR spectrum of **S0b** (400 MHz, $CDCl_3$)

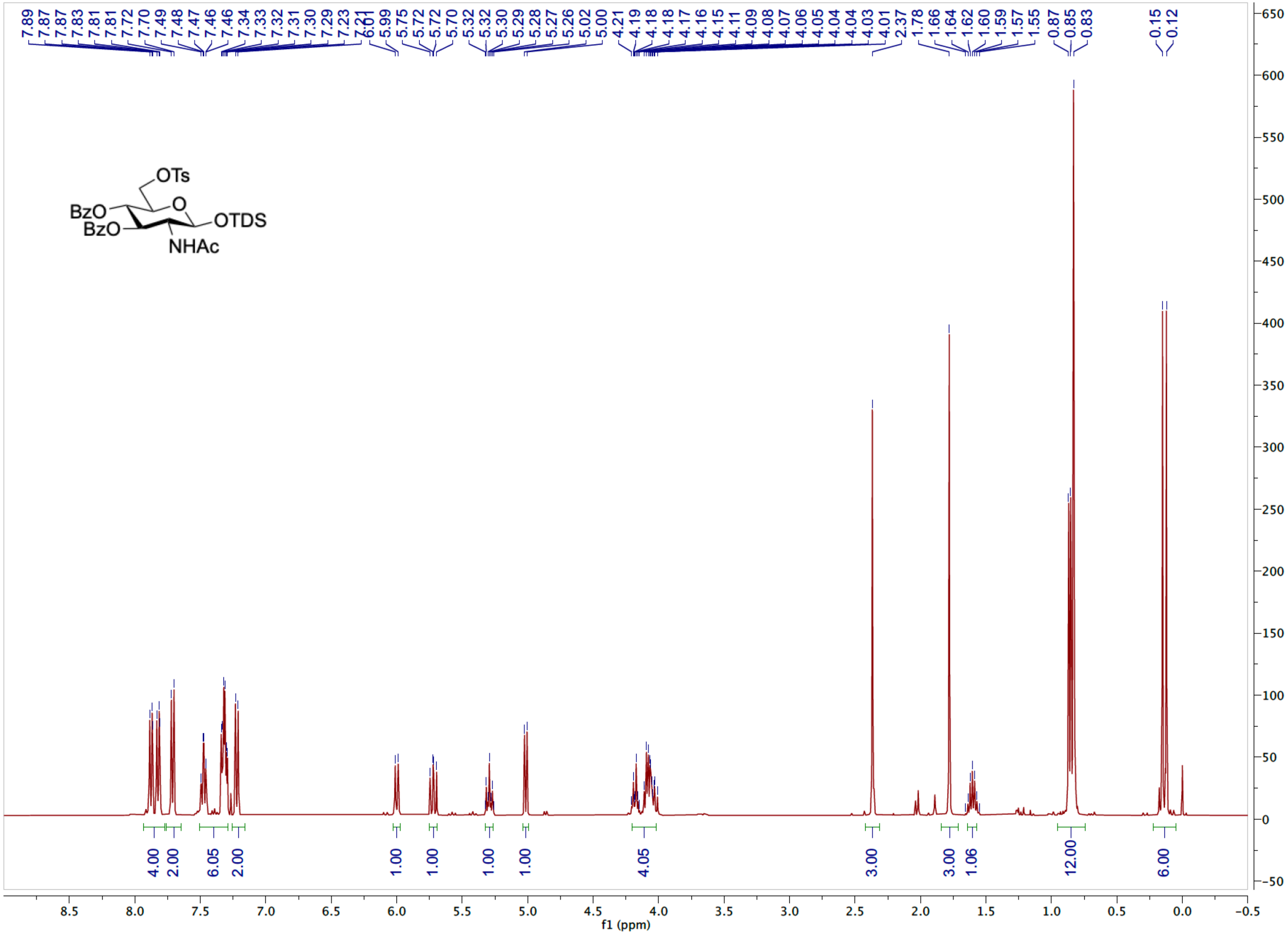

$^{13}C$ NMR spectrum of **S0b** (101 MHz, $CDCl_3$)

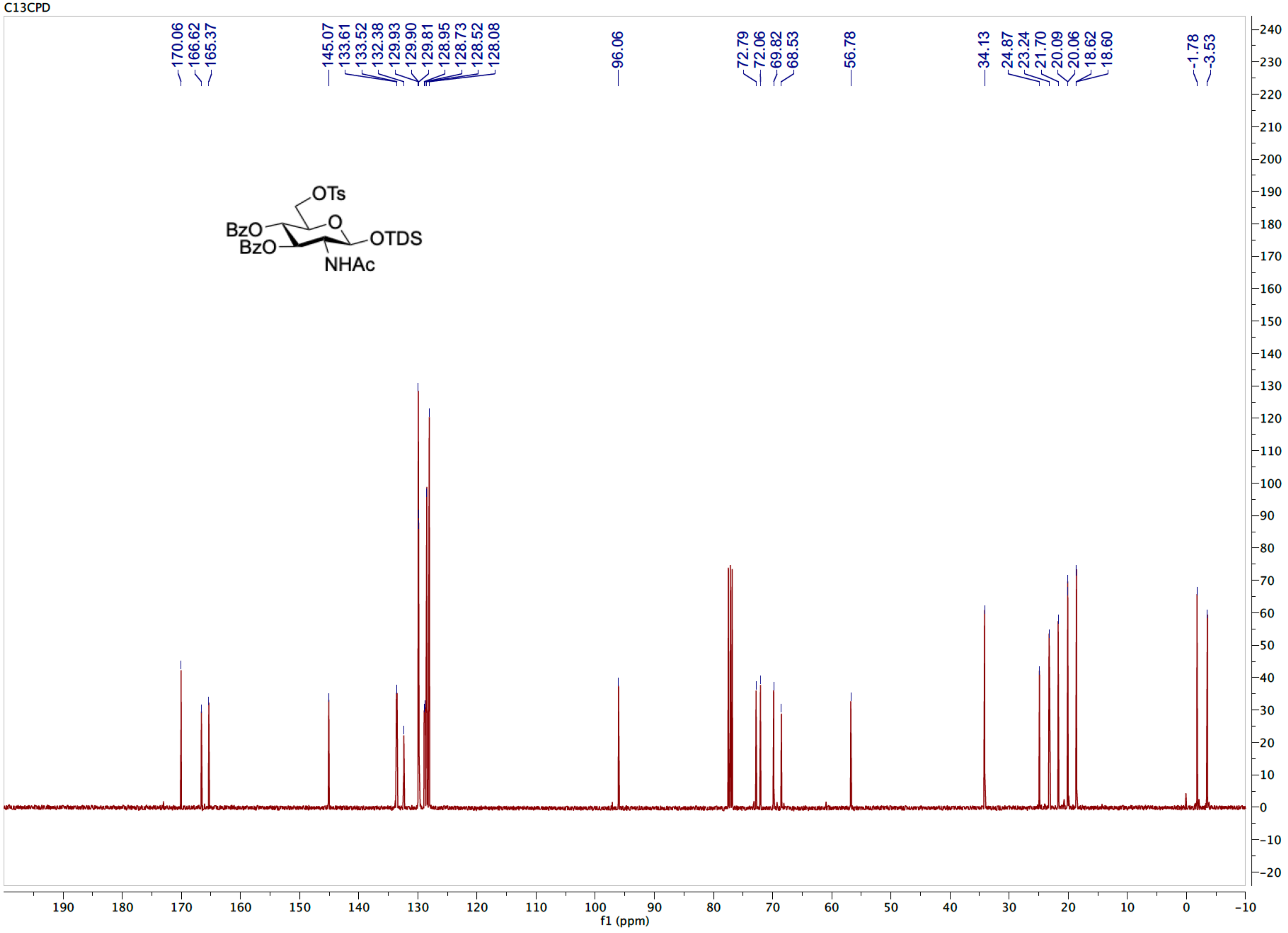

HSQC spectrum of **S0b** ($CDCl_3$)

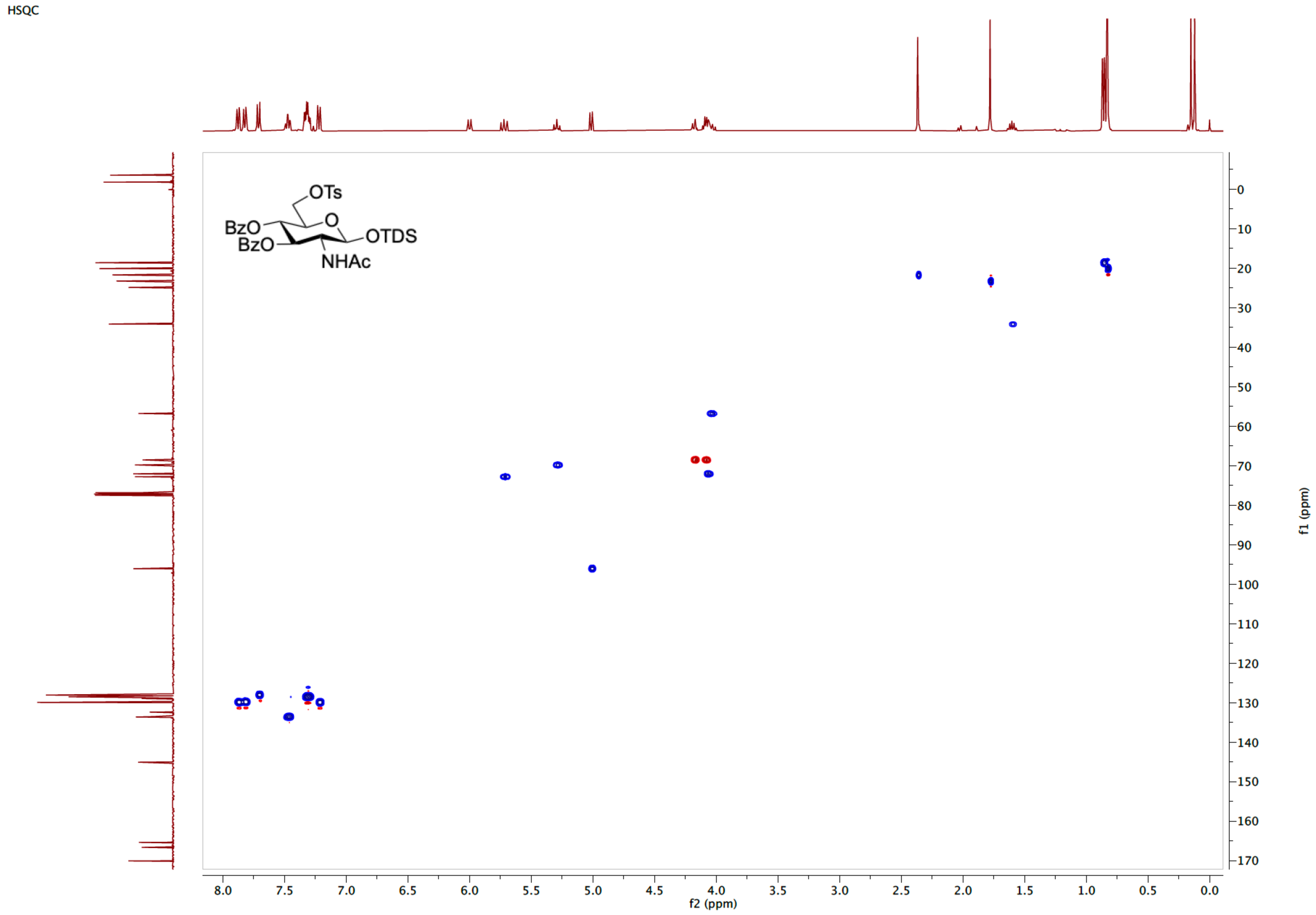

[1]H NMR spectrum of **S0c** (400 MHz, $CDCl_3$)

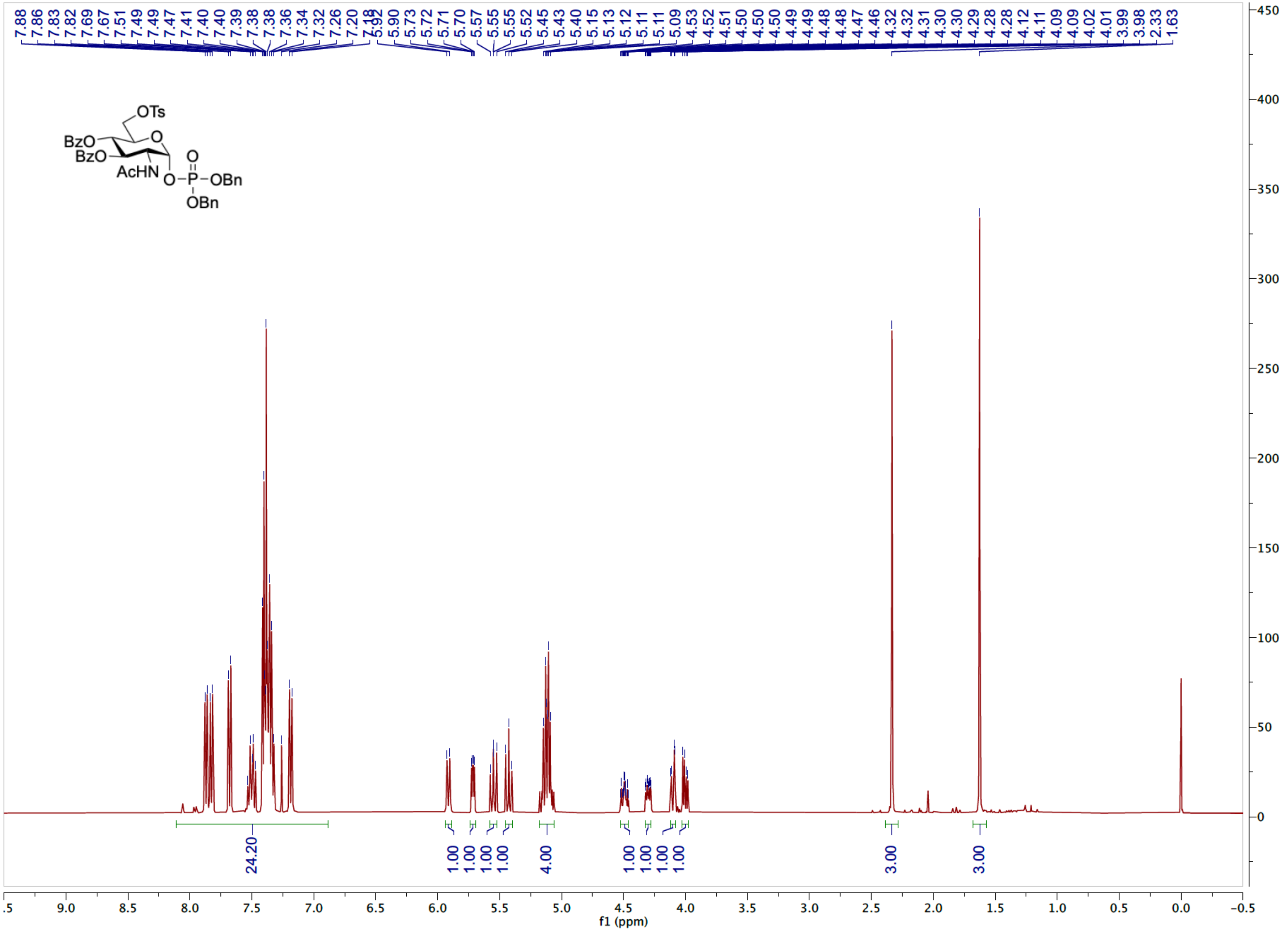

[13]C NMR spectrum of **S0c** (101 MHz, $CDCl_3$)

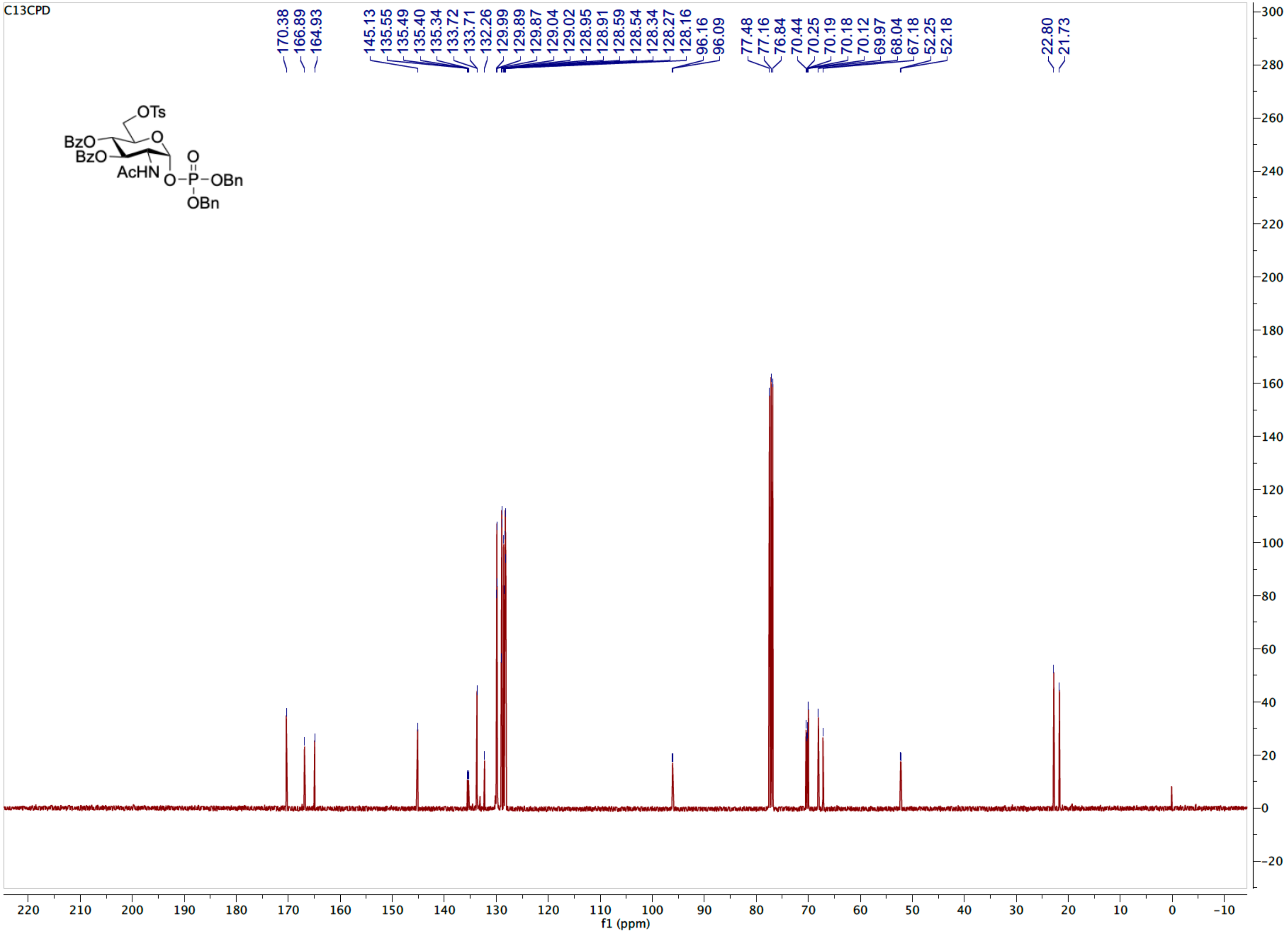

HSQC spectrum of **S0c** ($CDCl_3$)

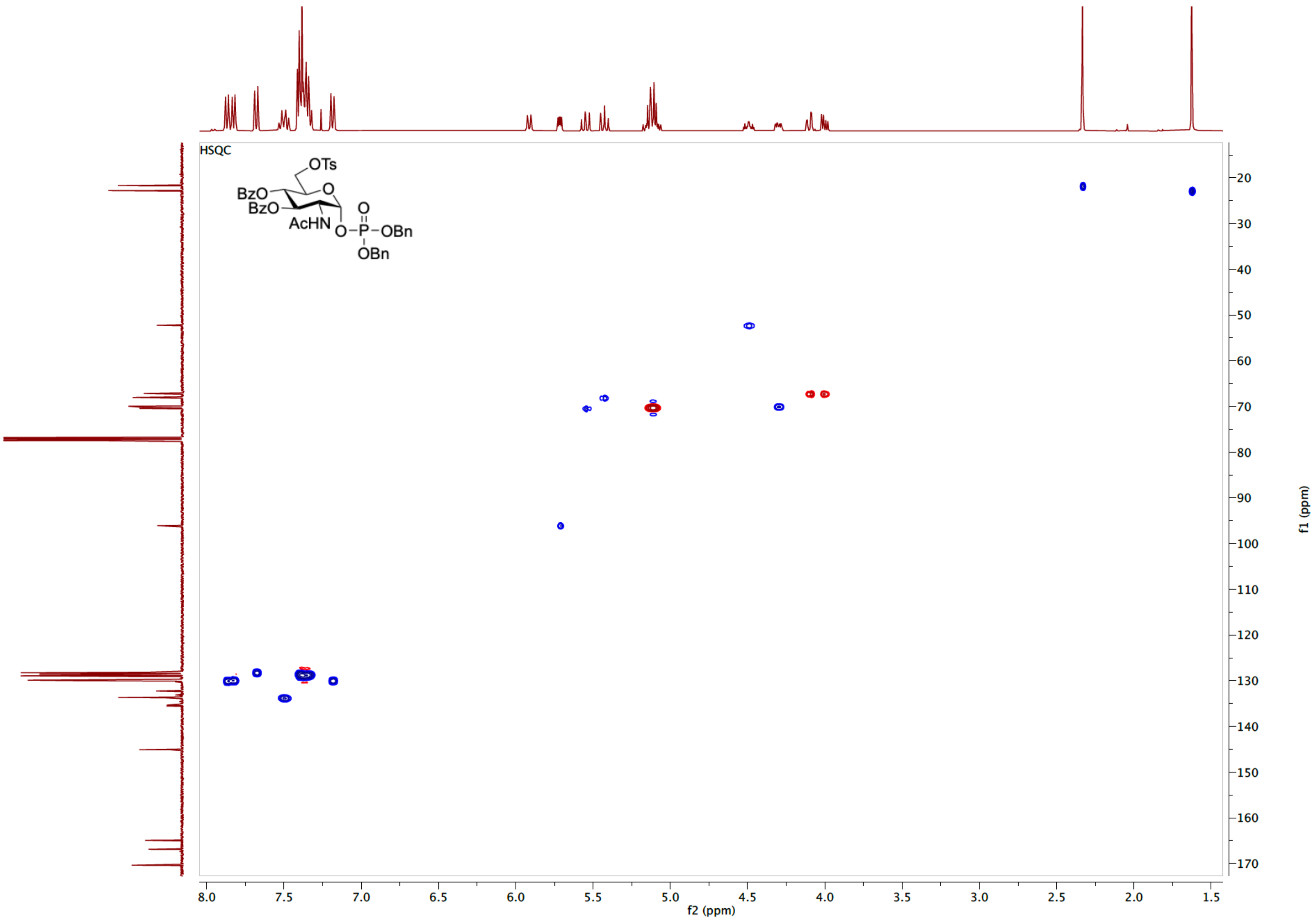

$^{31}$H NMR spectrum of **S0c** (162 MHz, $CDCl_3$)

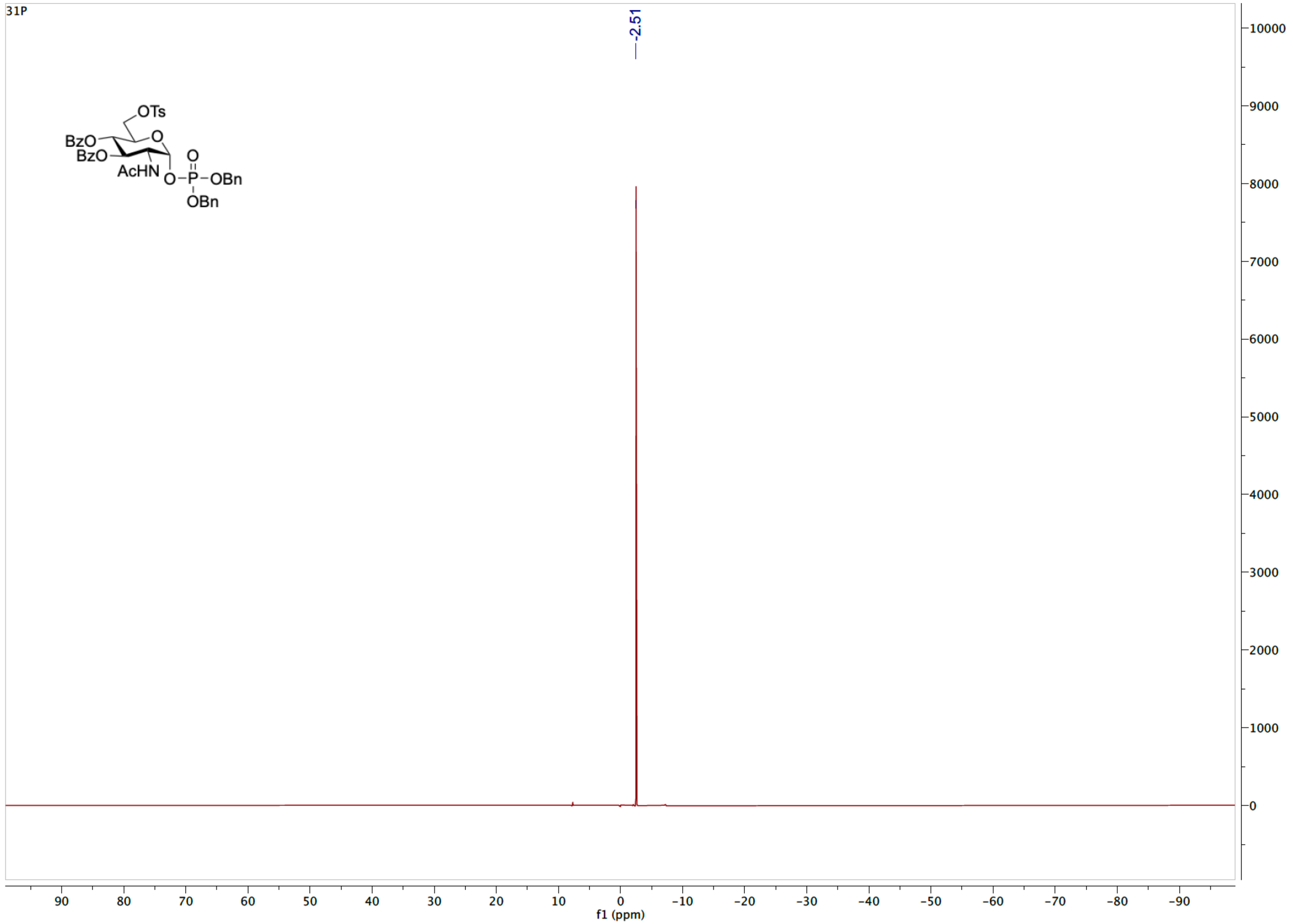

<sup></sup>

[1]H NMR spectrum of **S0** (600 MHz, $D_2O$).

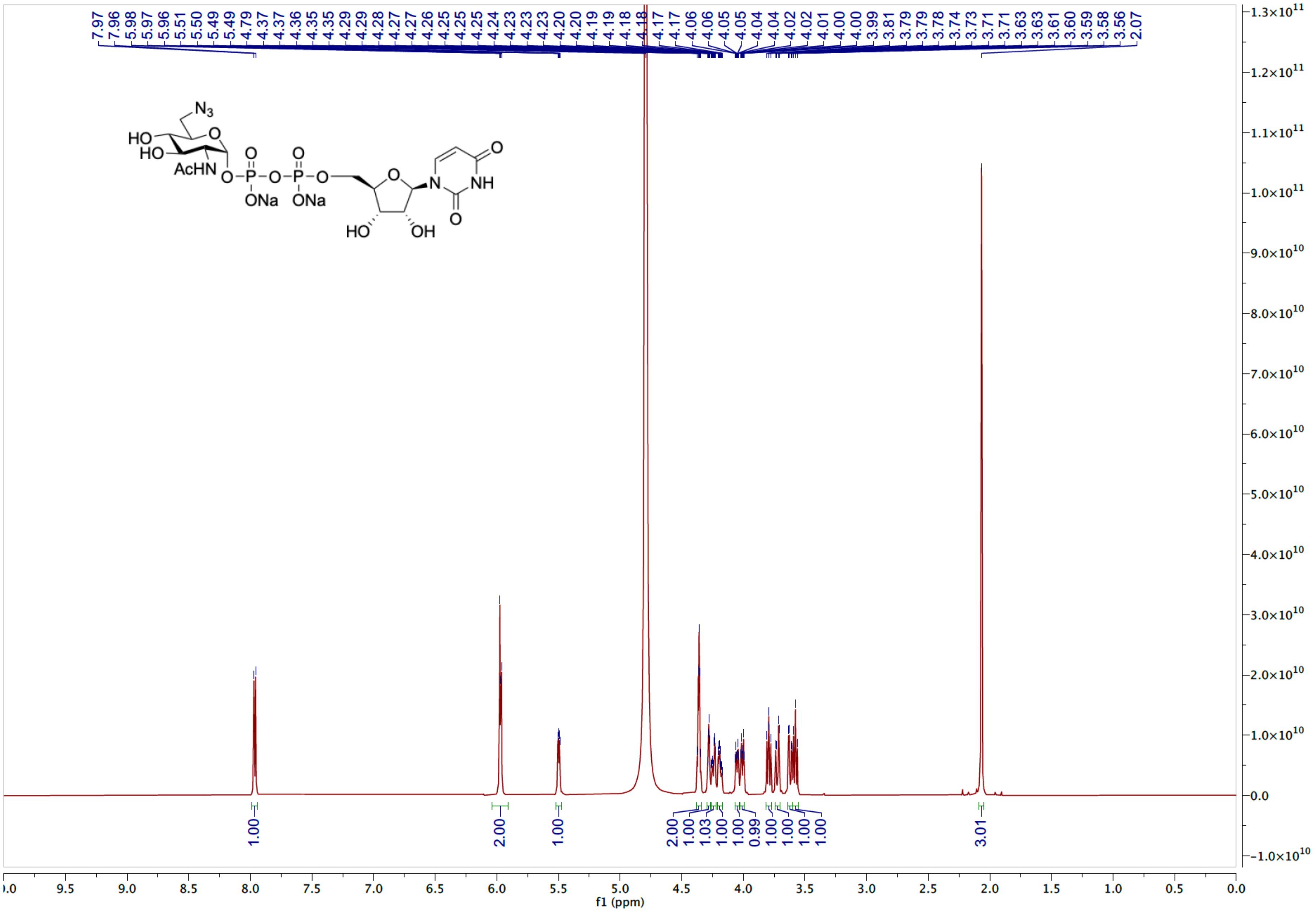

HSQC spectrum of **S0** ($D_2O$)

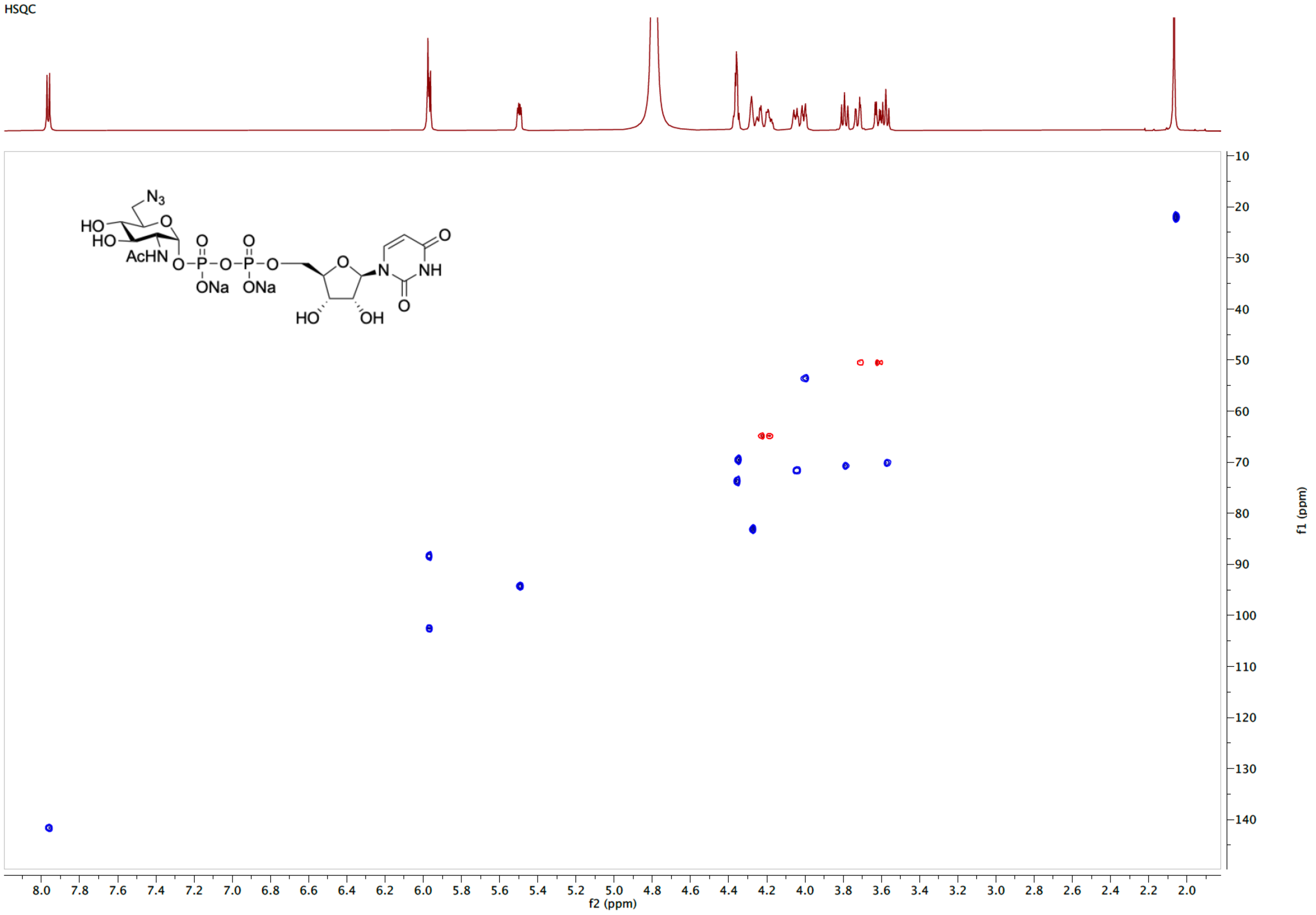

[1]H NMR spectrum of **S1** (600 MHz, $CDCl_3$)

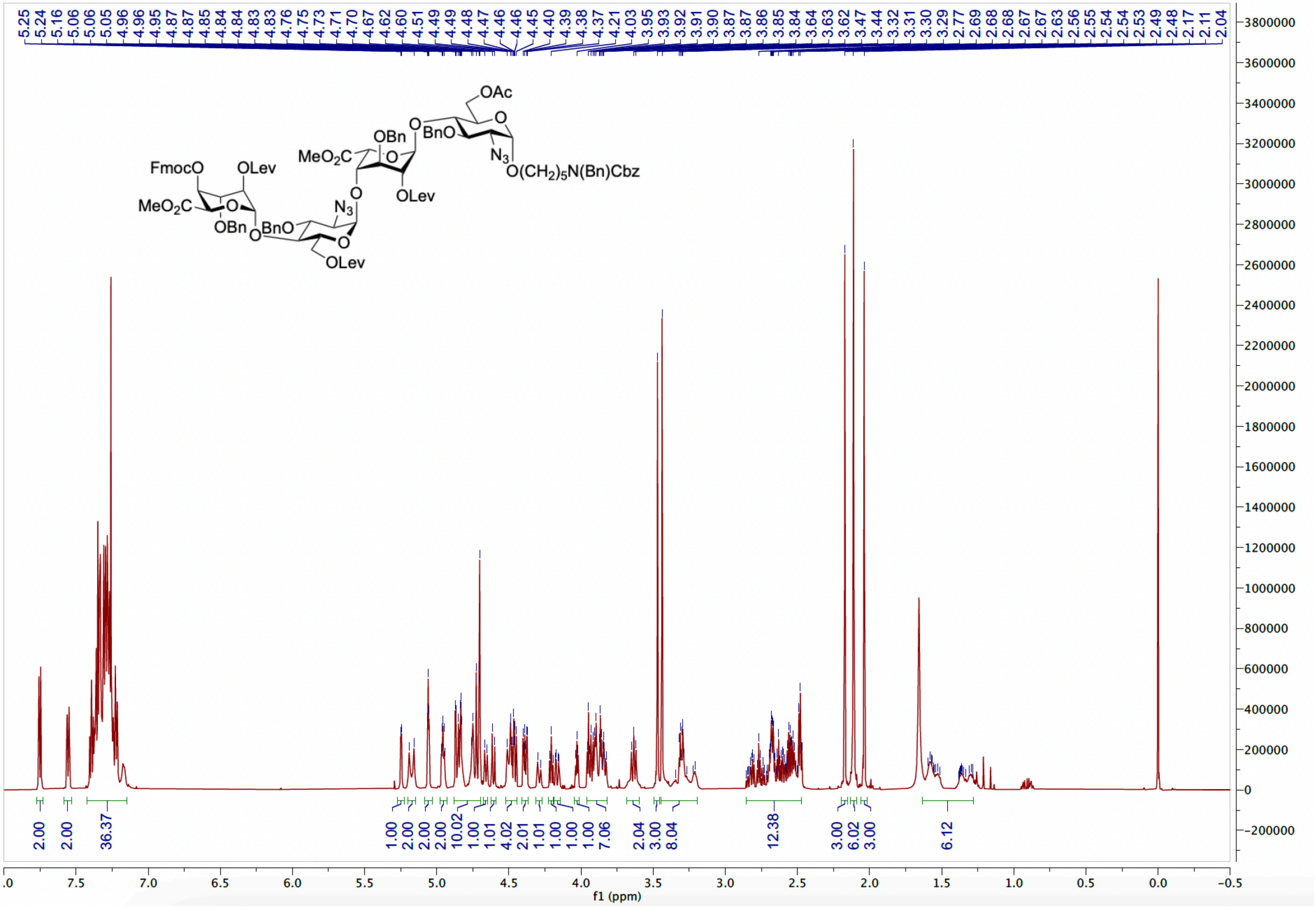

$^{13}$C NMR spectrum of **S1** (151 MHz, $CDCl_3$)

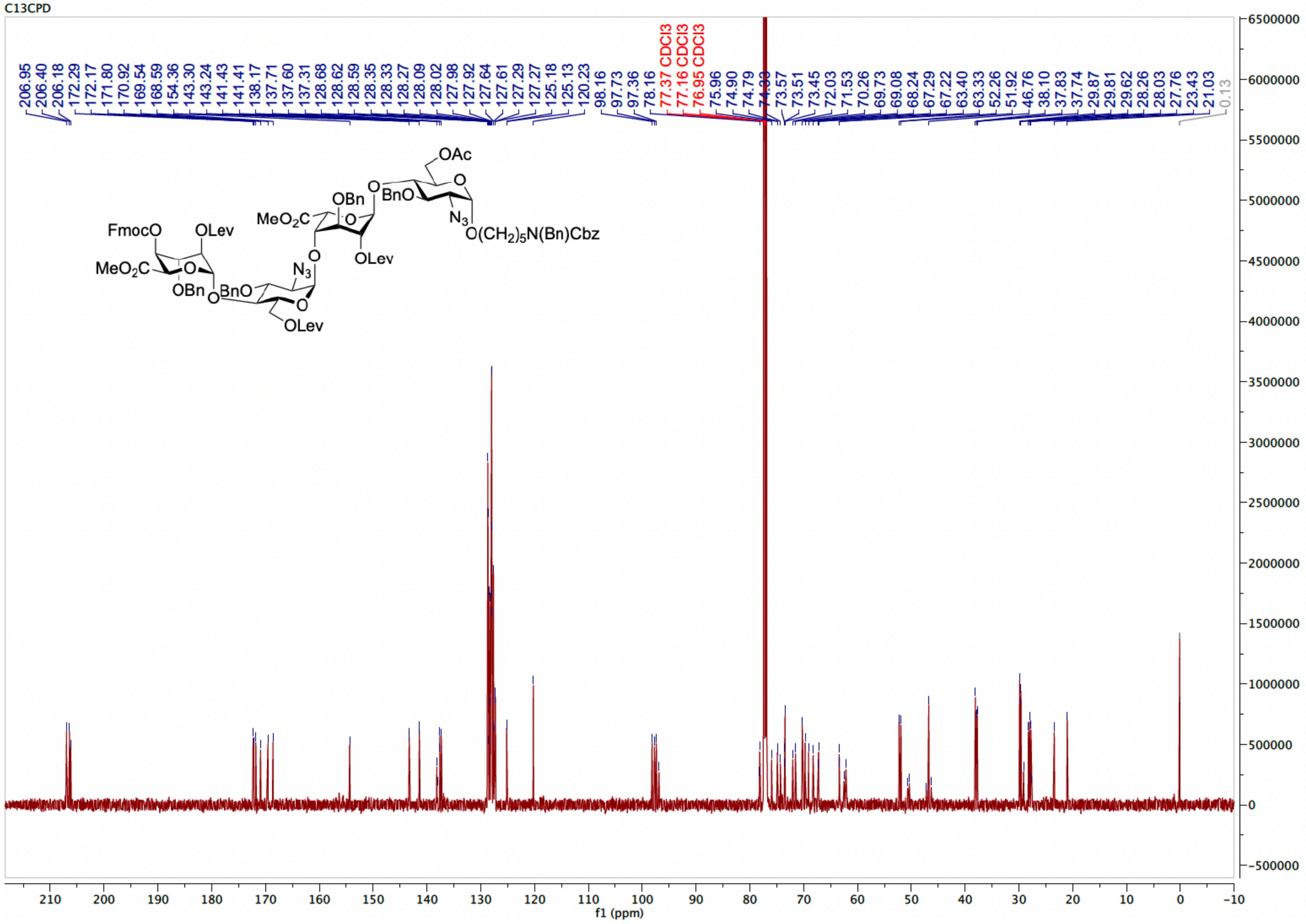

HSQC spectrum of **S1** ($CDCl_3$)

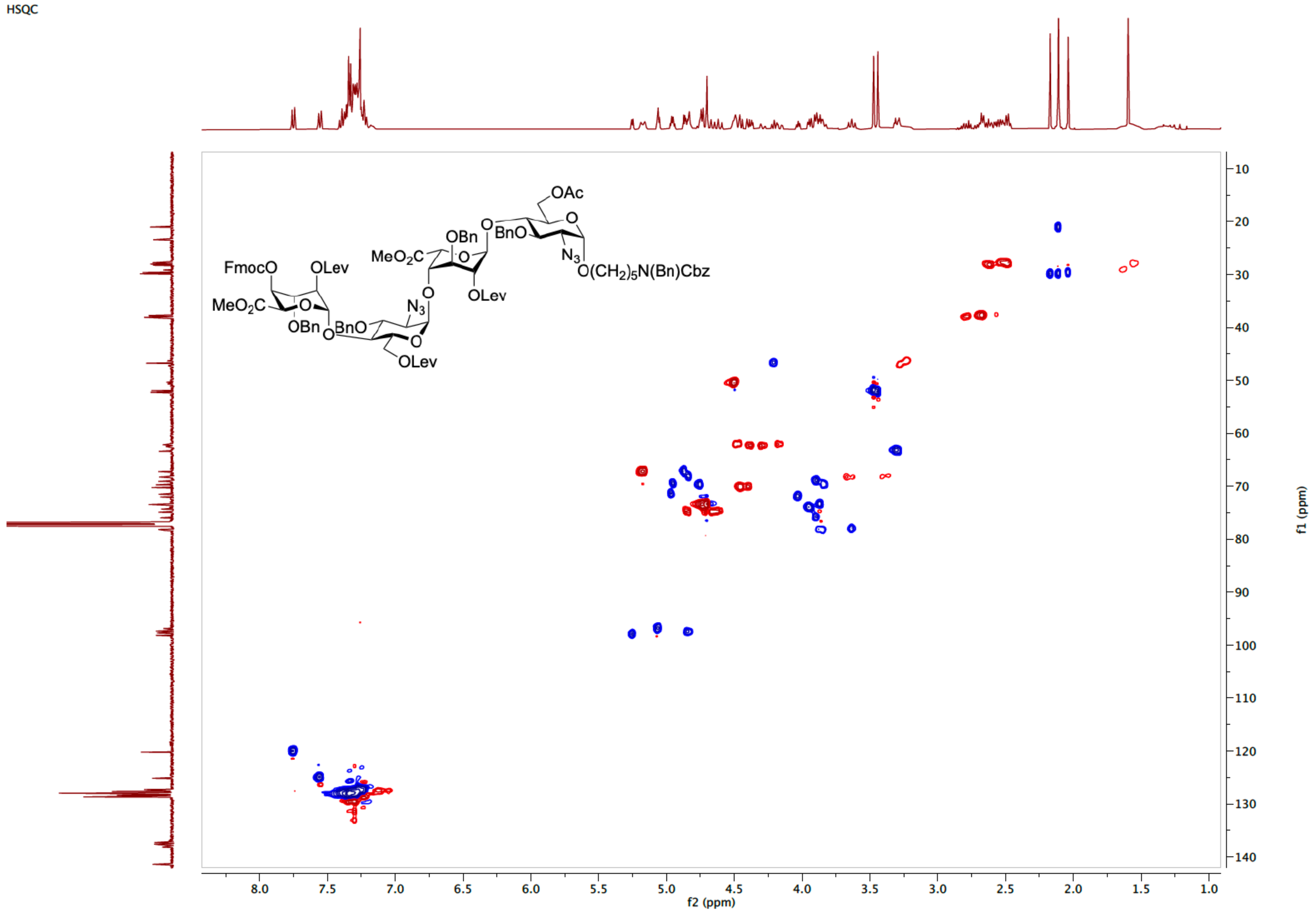

[1]H NMR spectrum of **S2** (400 MHz, $CDCl_3$)

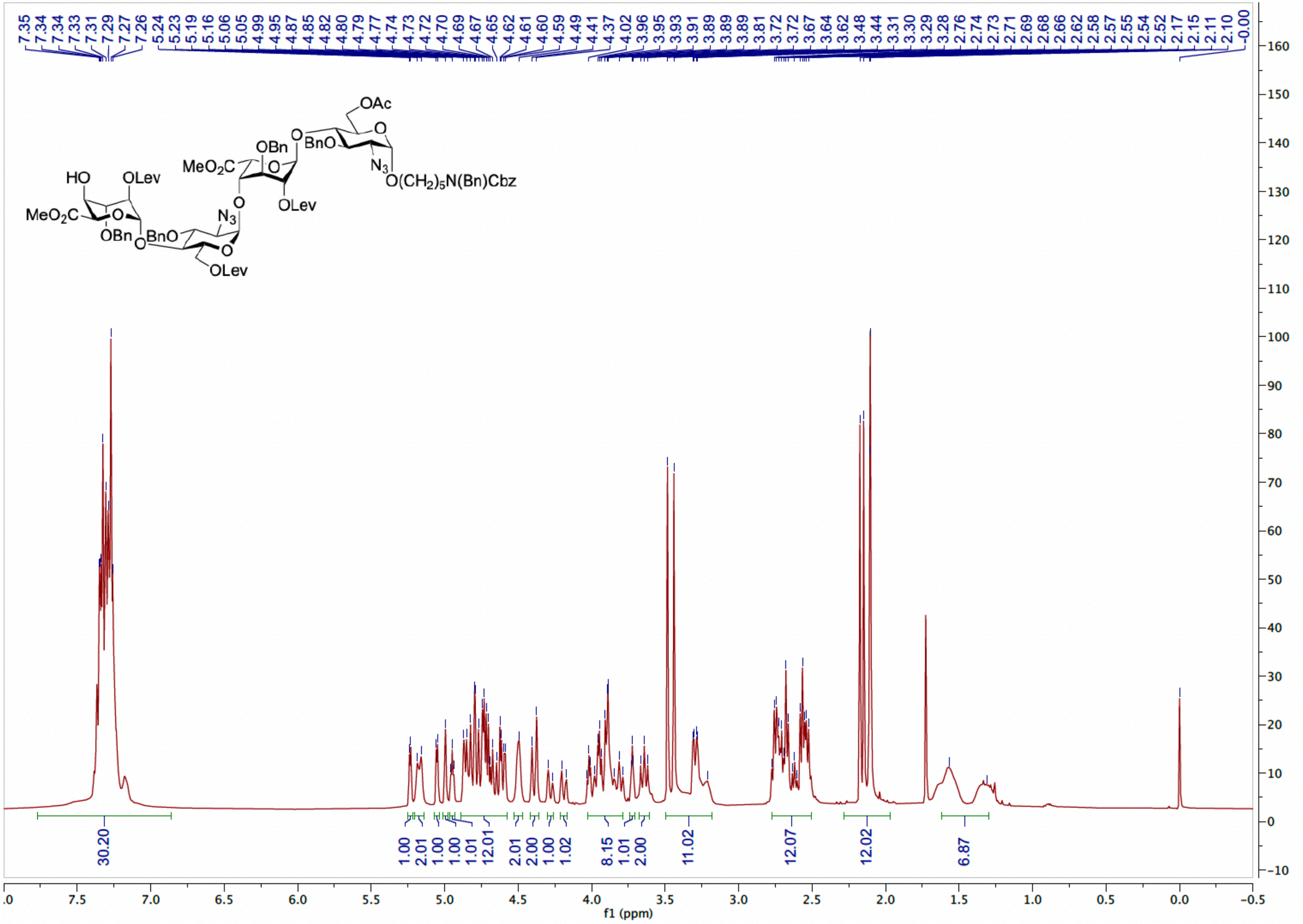

[13]C NMR spectrum of **S2** (101 MHz, $CDCl_3$)

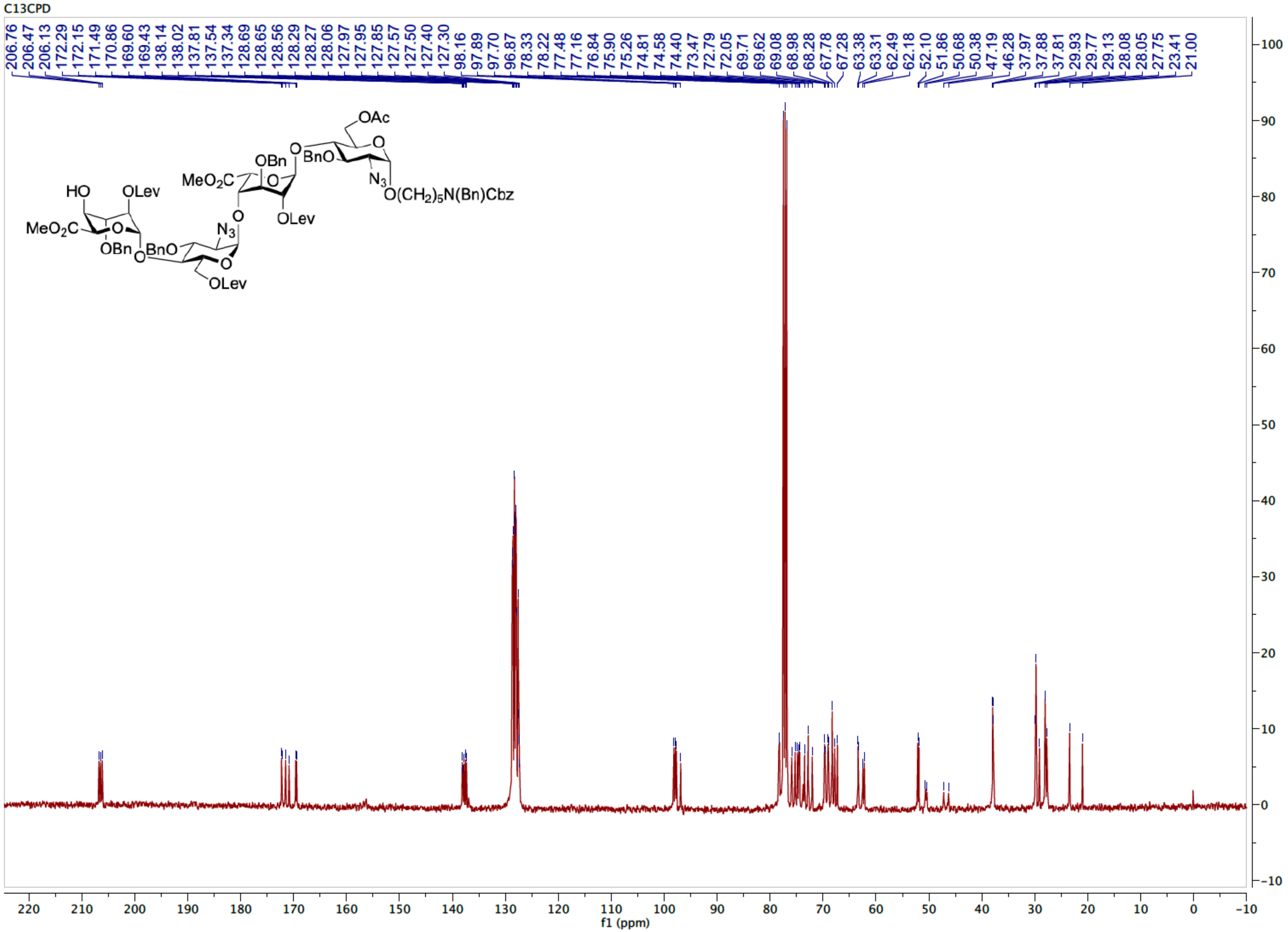

HSQC spectrum of **S2** ($CDCl_3$)

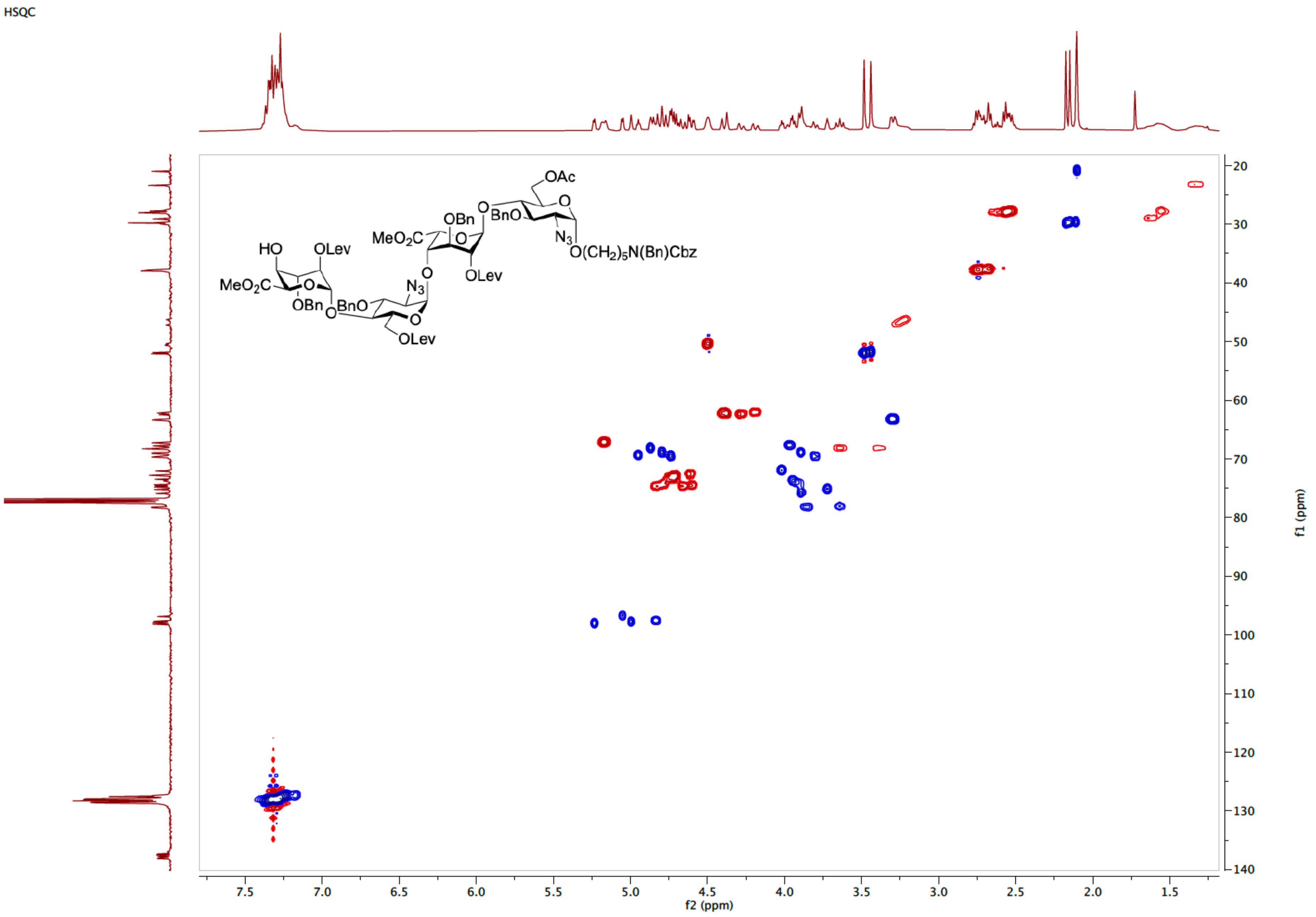

$^1$H NMR spectrum of **18** (400 MHz, $CDCl_3$)

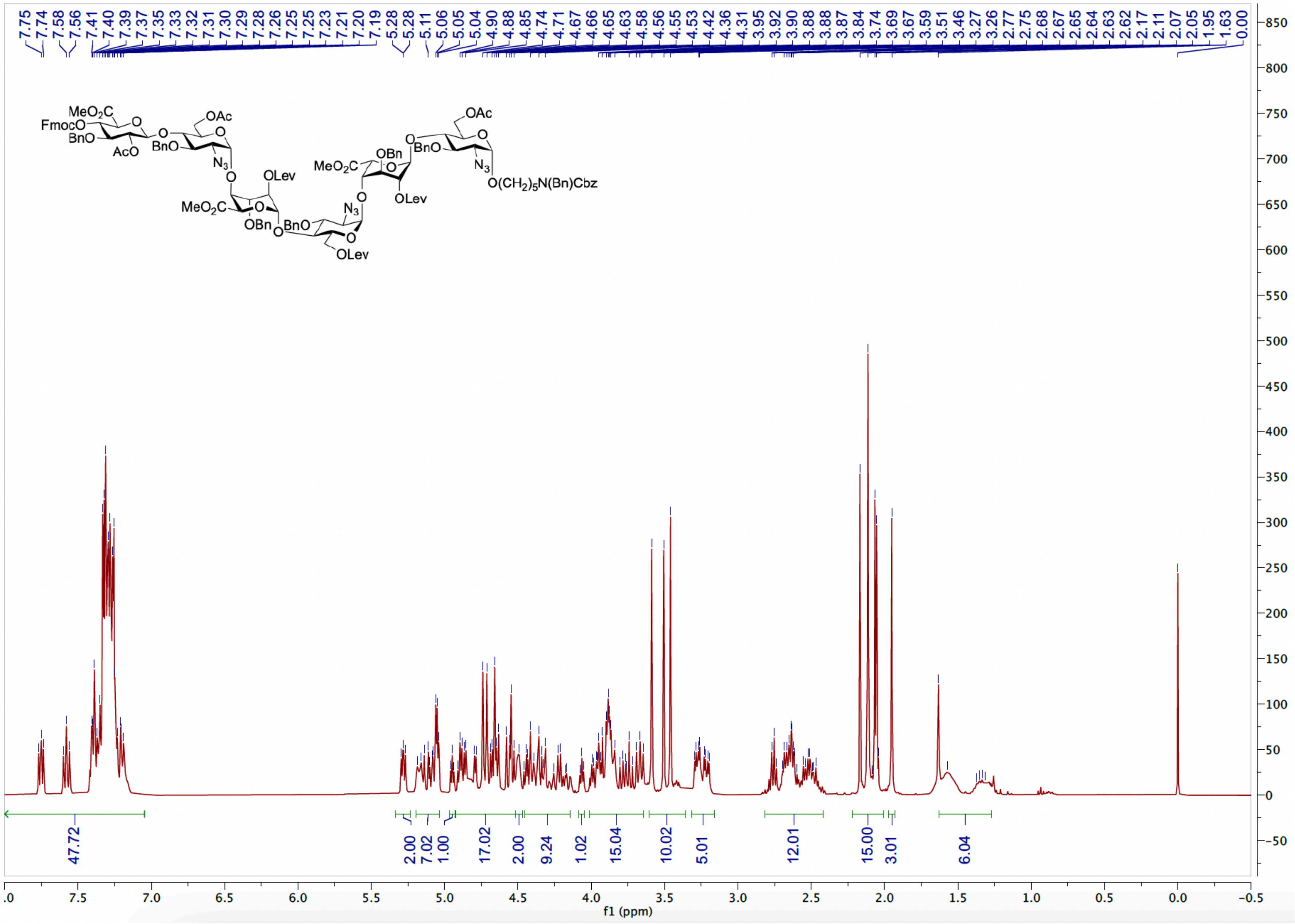

$^{13}C$ APT NMR spectrum of **18** (101 MHz, $CDCl_3$)

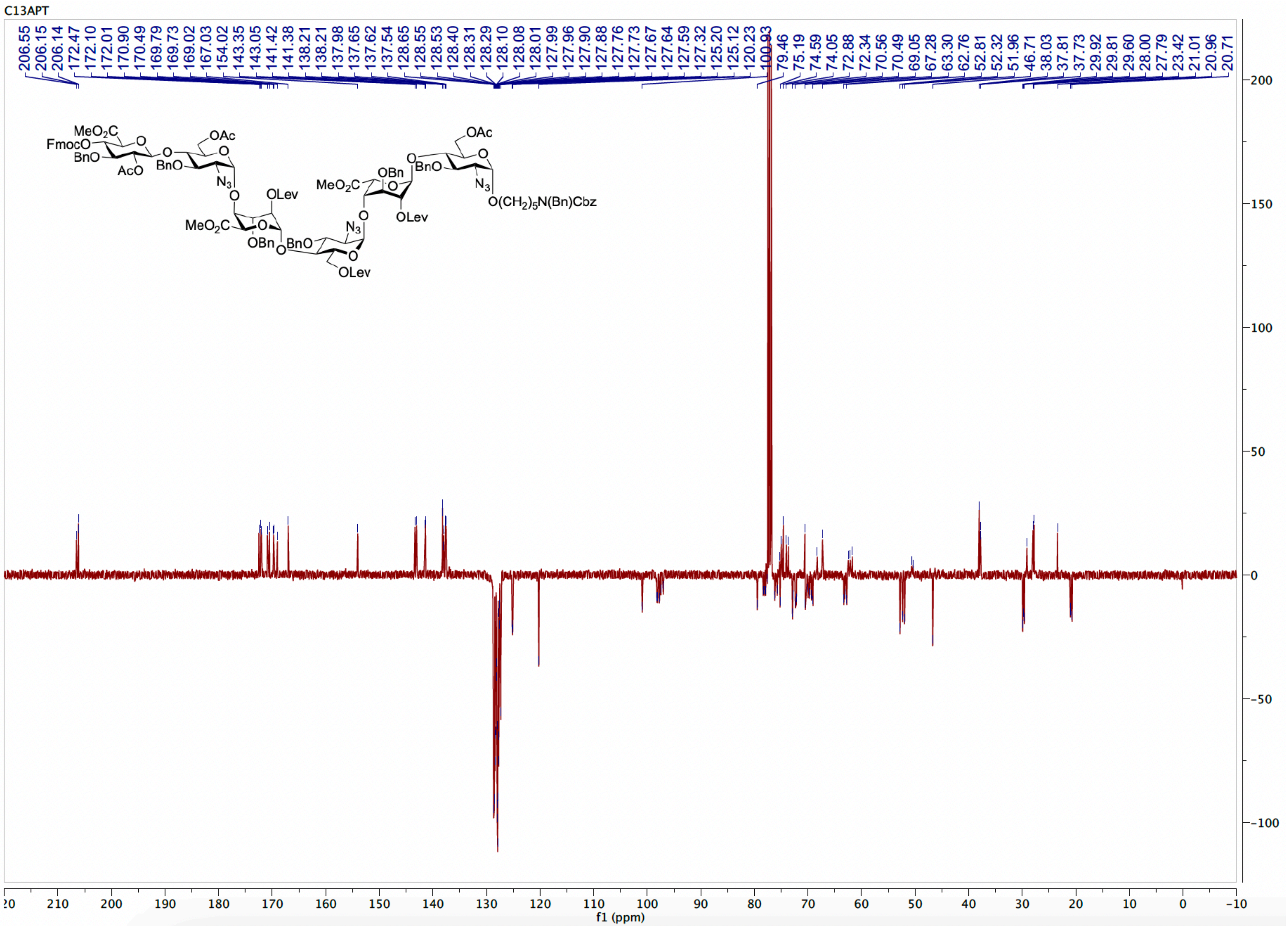

HSQC spectrum of **18** ($CDCl_3$)

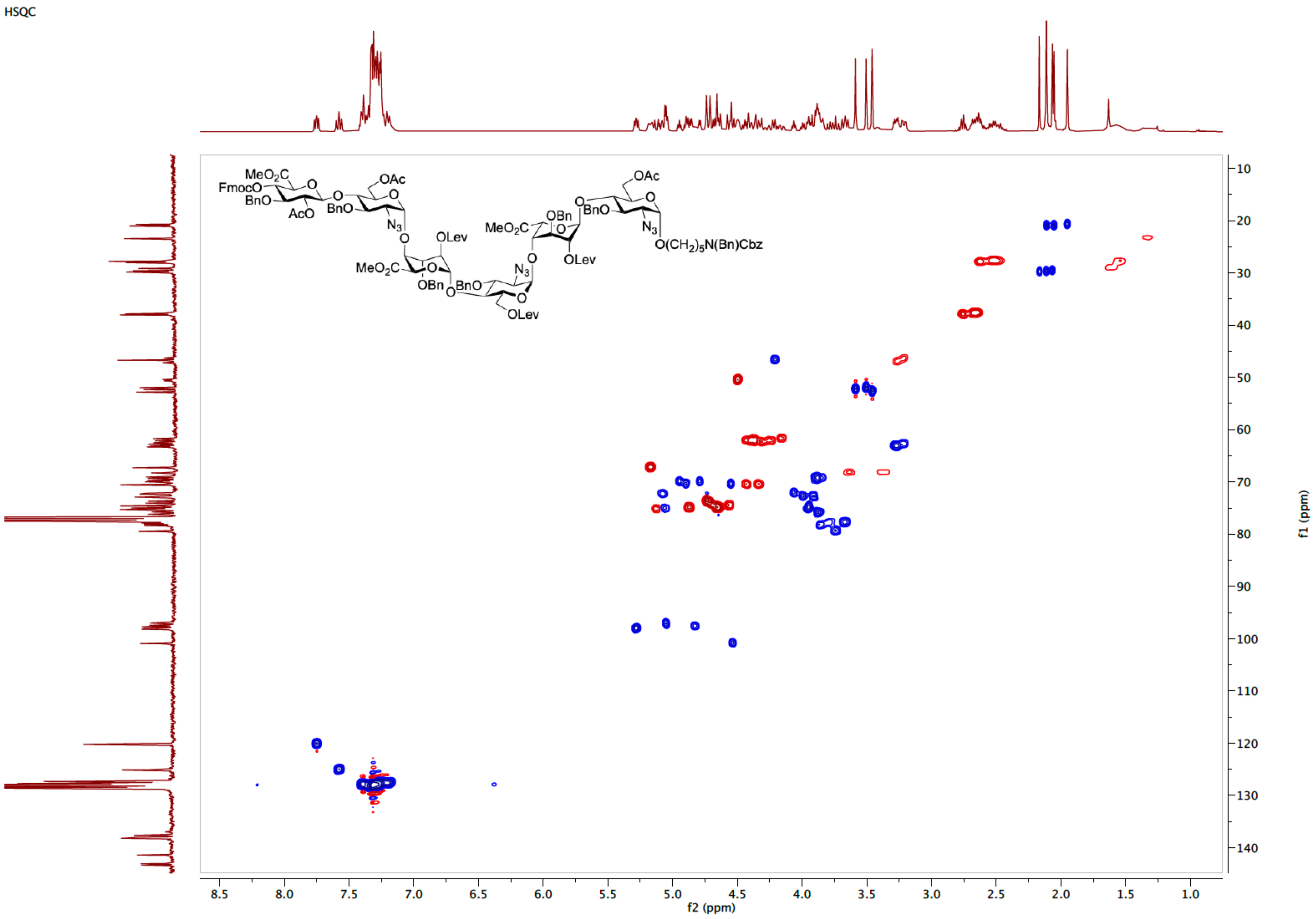

[1]H NMR spectrum of **21** (600 MHz, $D_2O$)

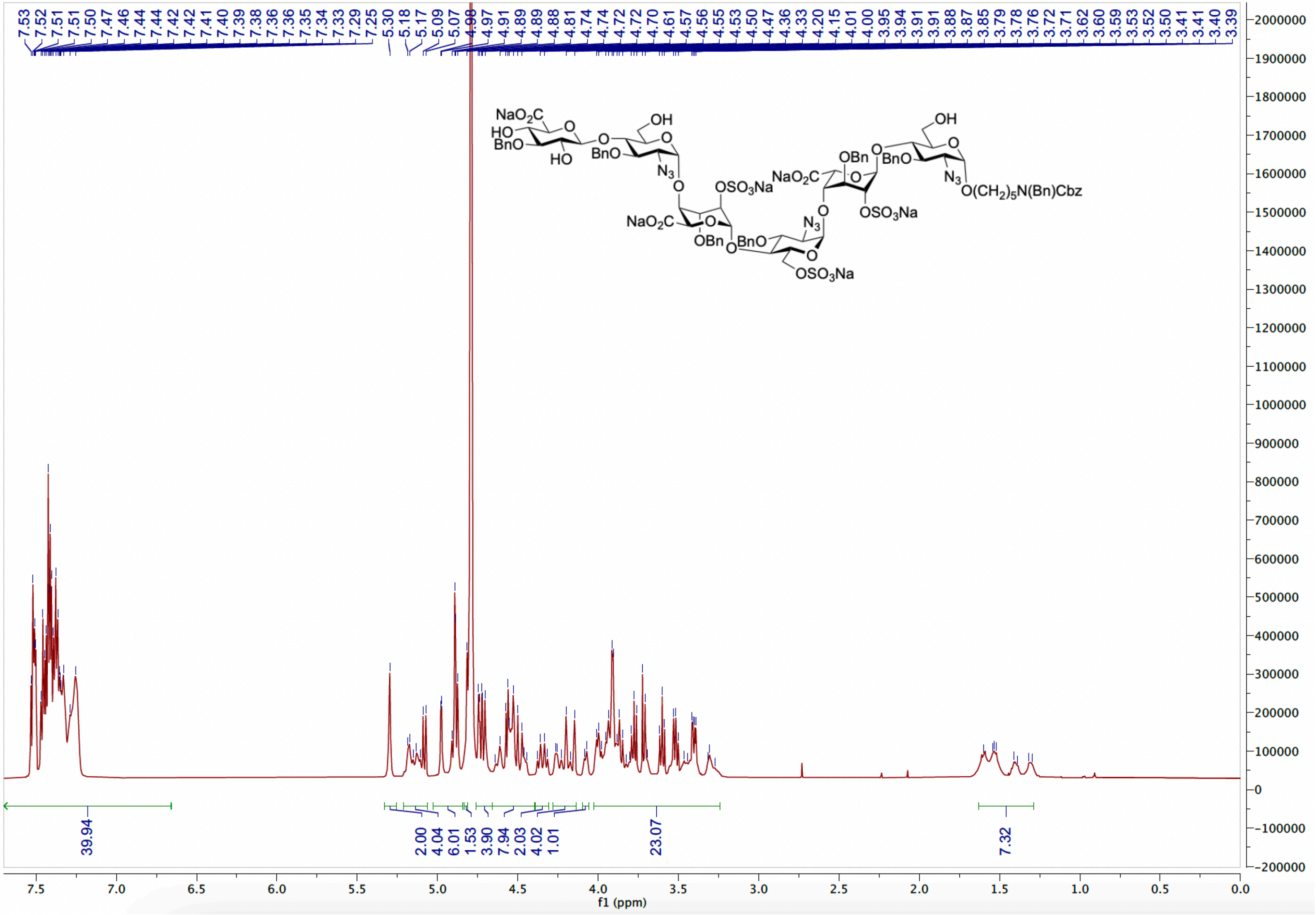

HSQC spectrum of **21** ($D_2O$)

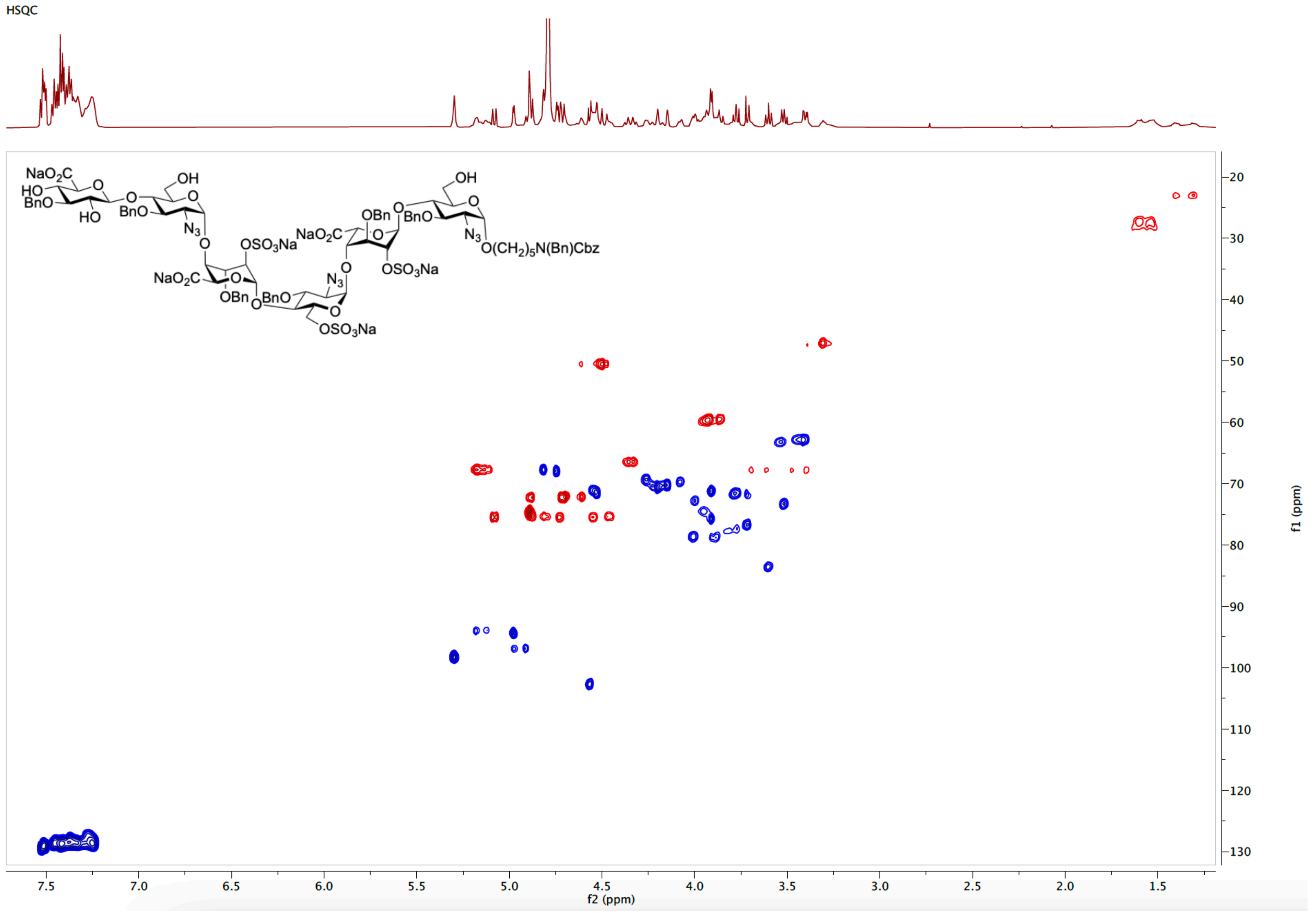

<sup></sup>[1]H NMR spectrum of **22** (600 MHz, $D_2O$)

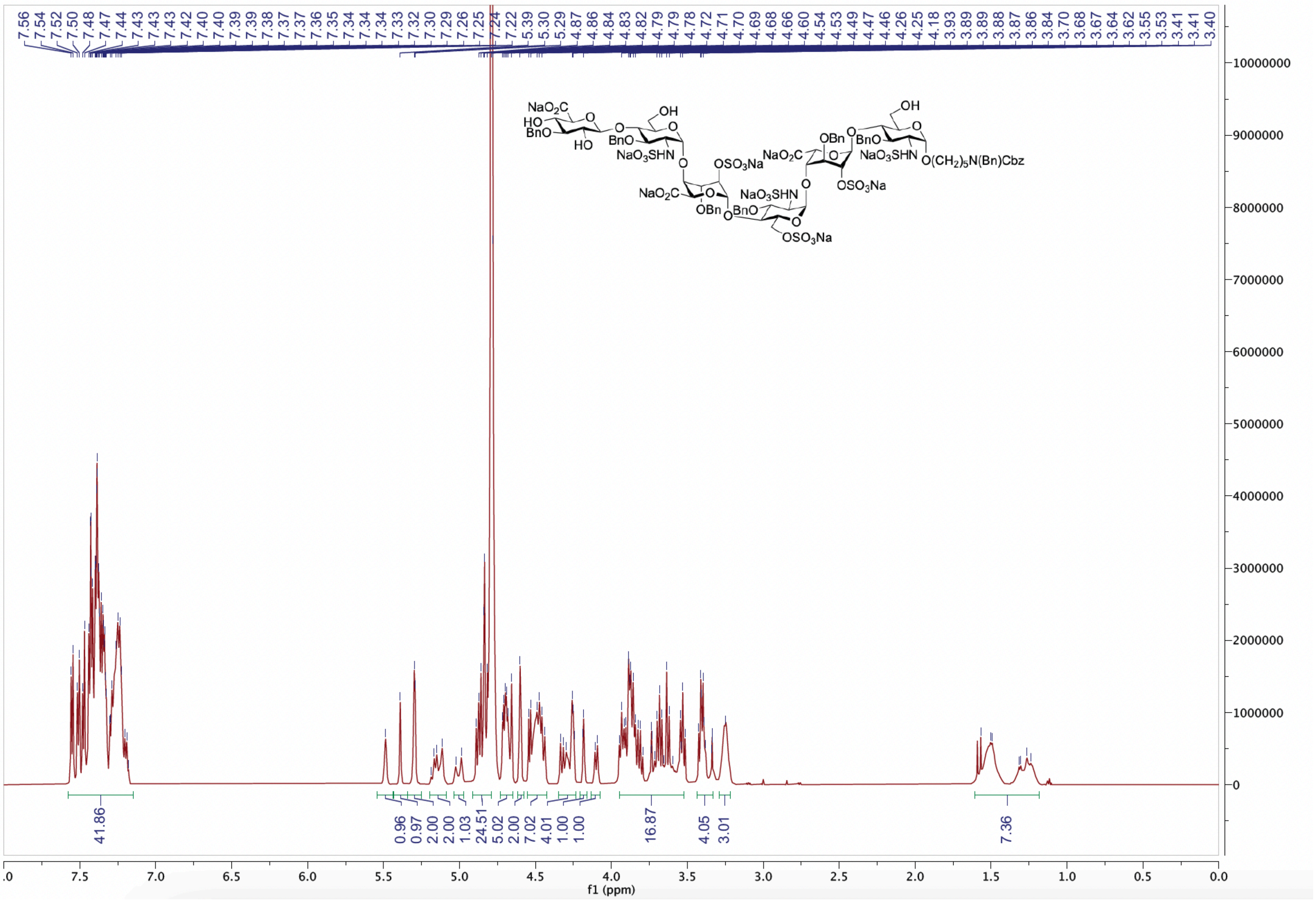

HSQC spectrum of **22** ($D_2O$)

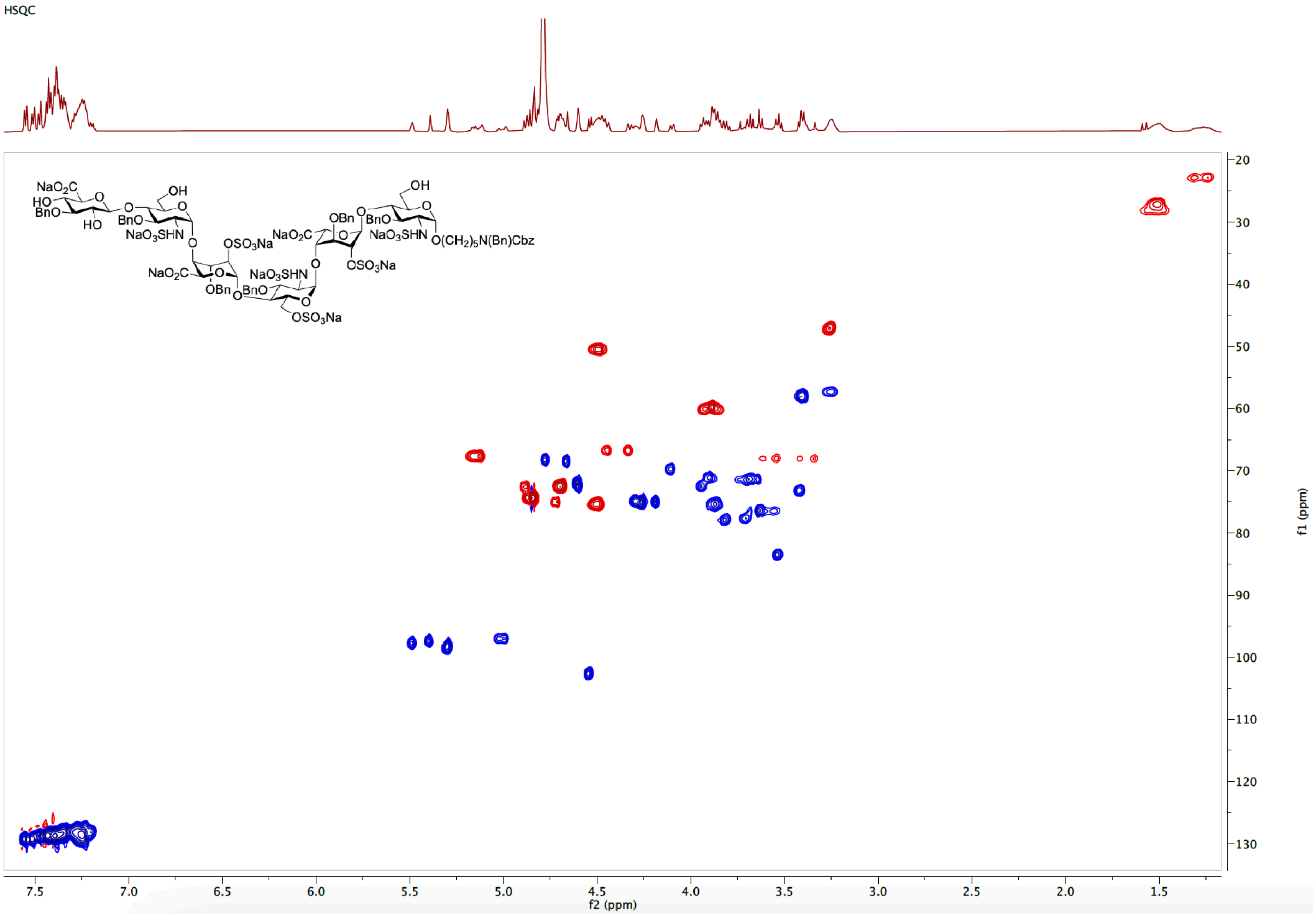

[1]H NMR spectrum of **1** (600 MHz, $D_2O$)

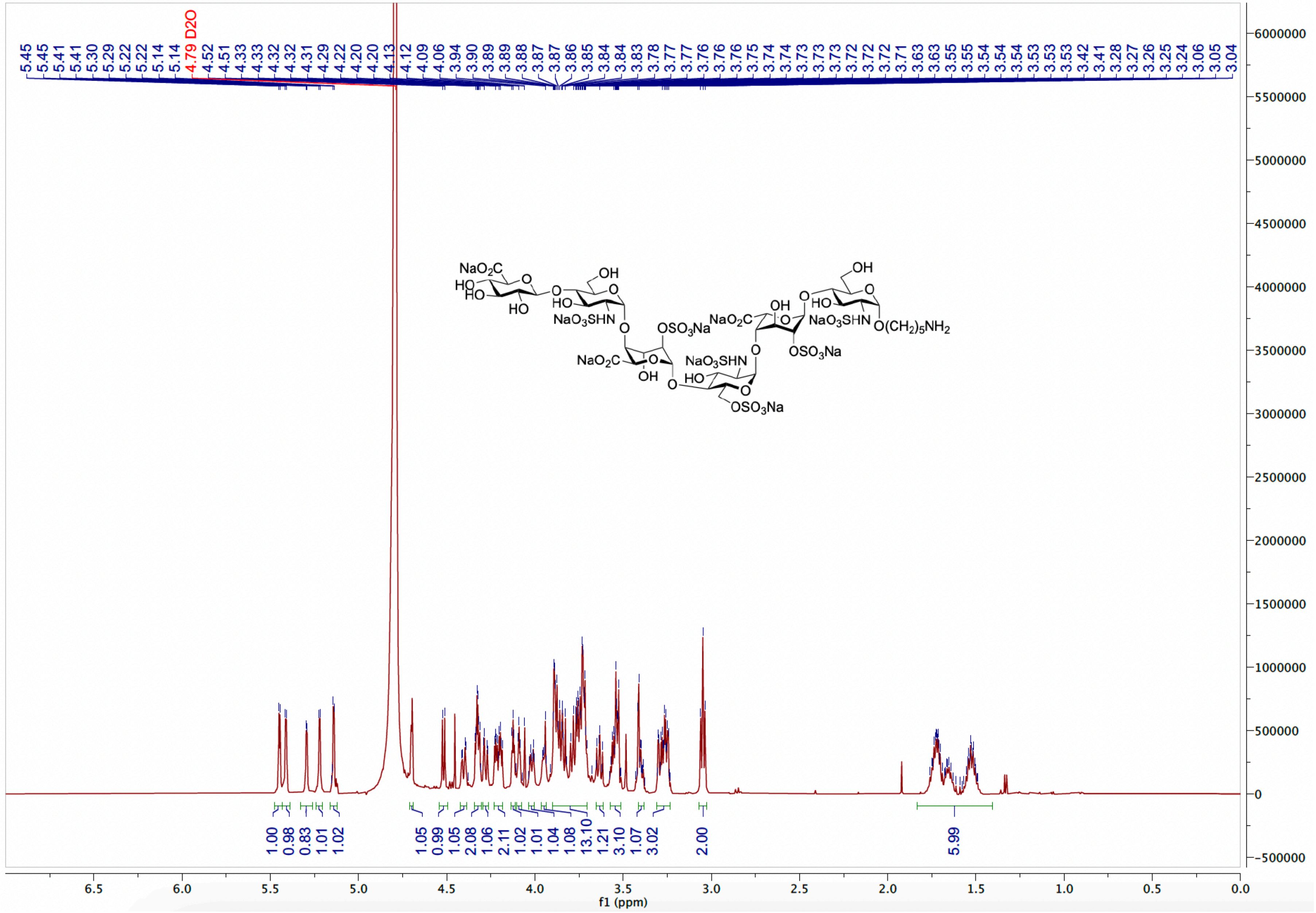

HSQC spectrum of **1** ($D_2O$)

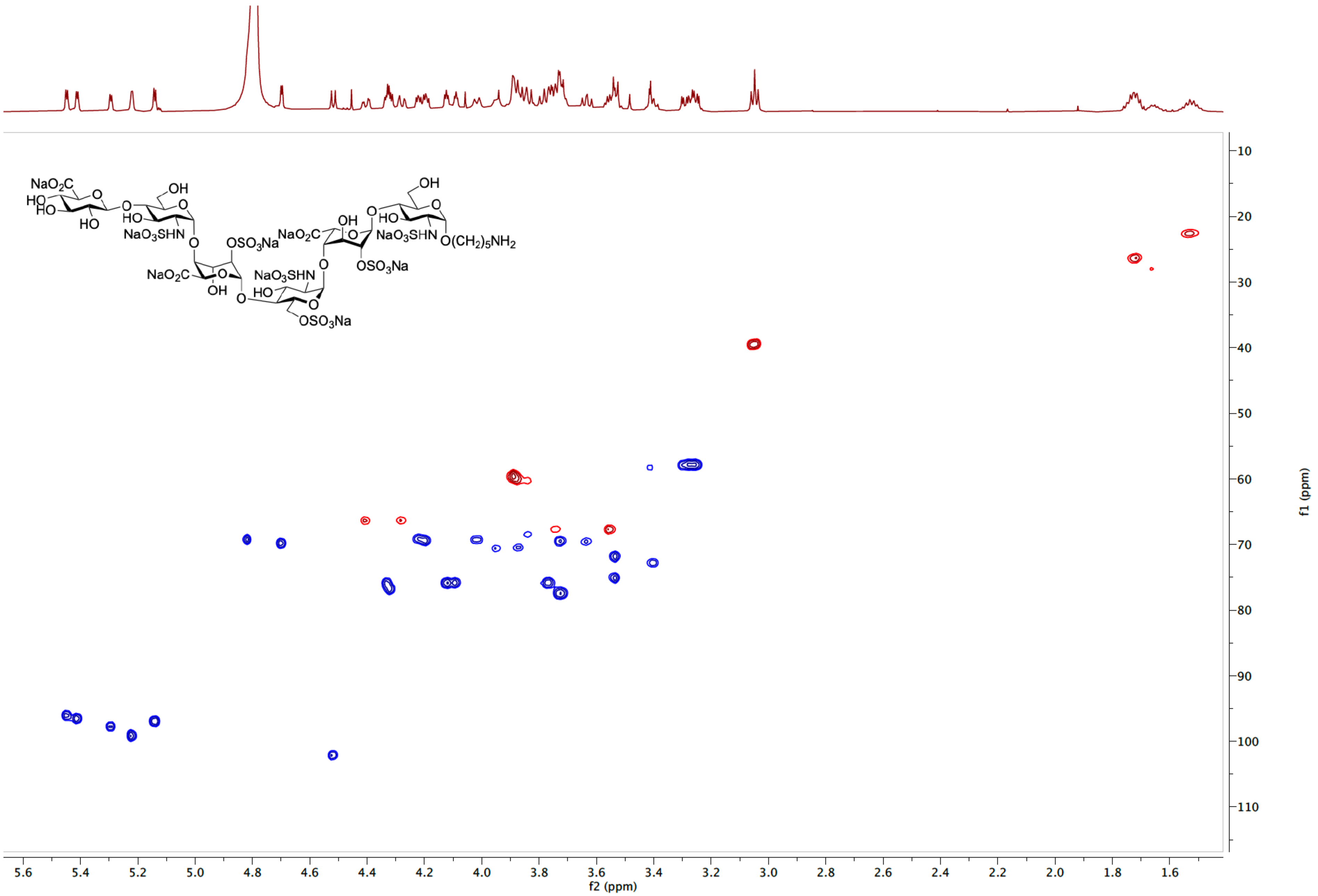

COSY spectrum of **1** ($D_2O$)

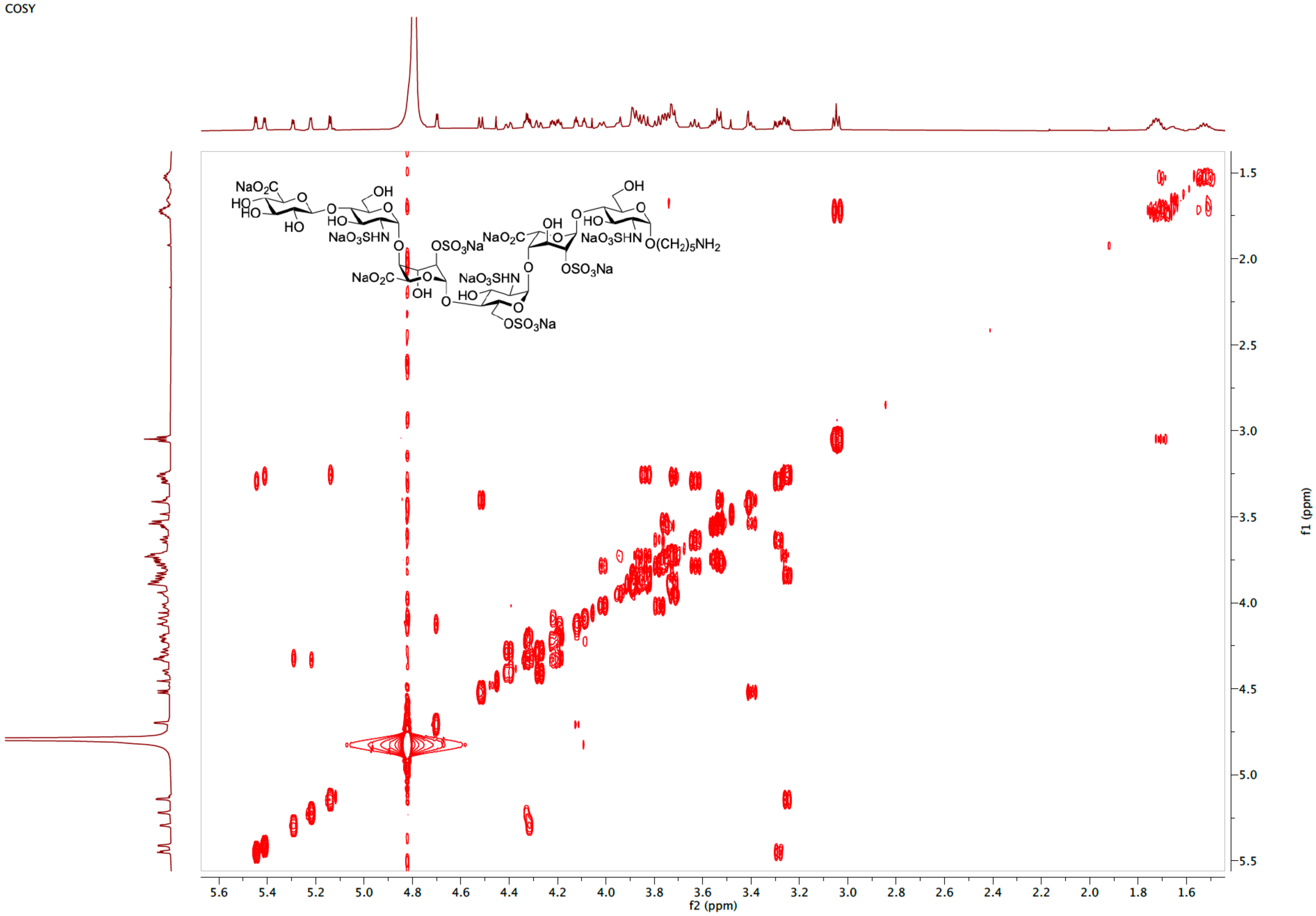

TOCSY spectrum of **1** ($D_2O$)

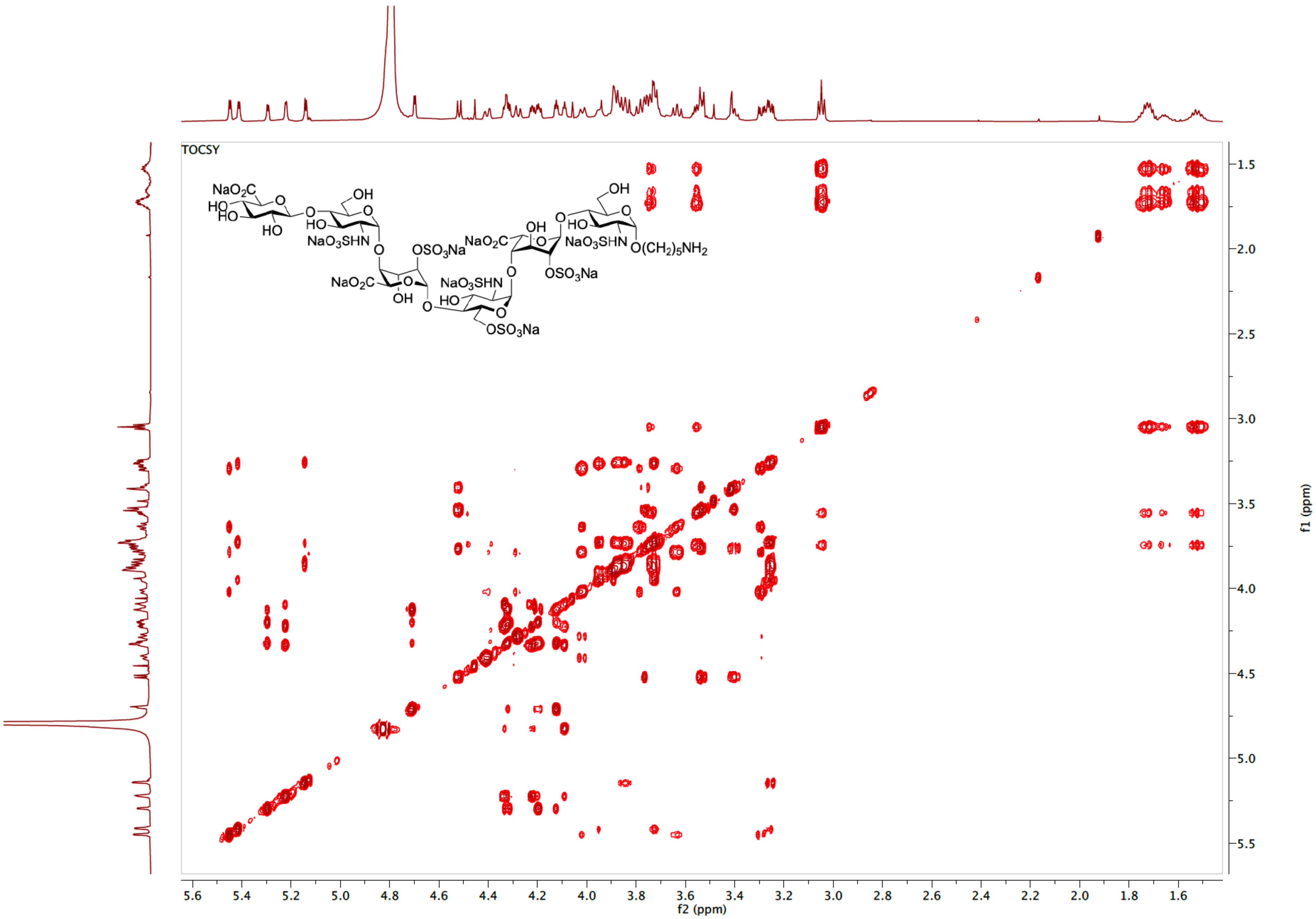

[1]H NMR spectrum of **2** (600 MHz, $D_2O$)

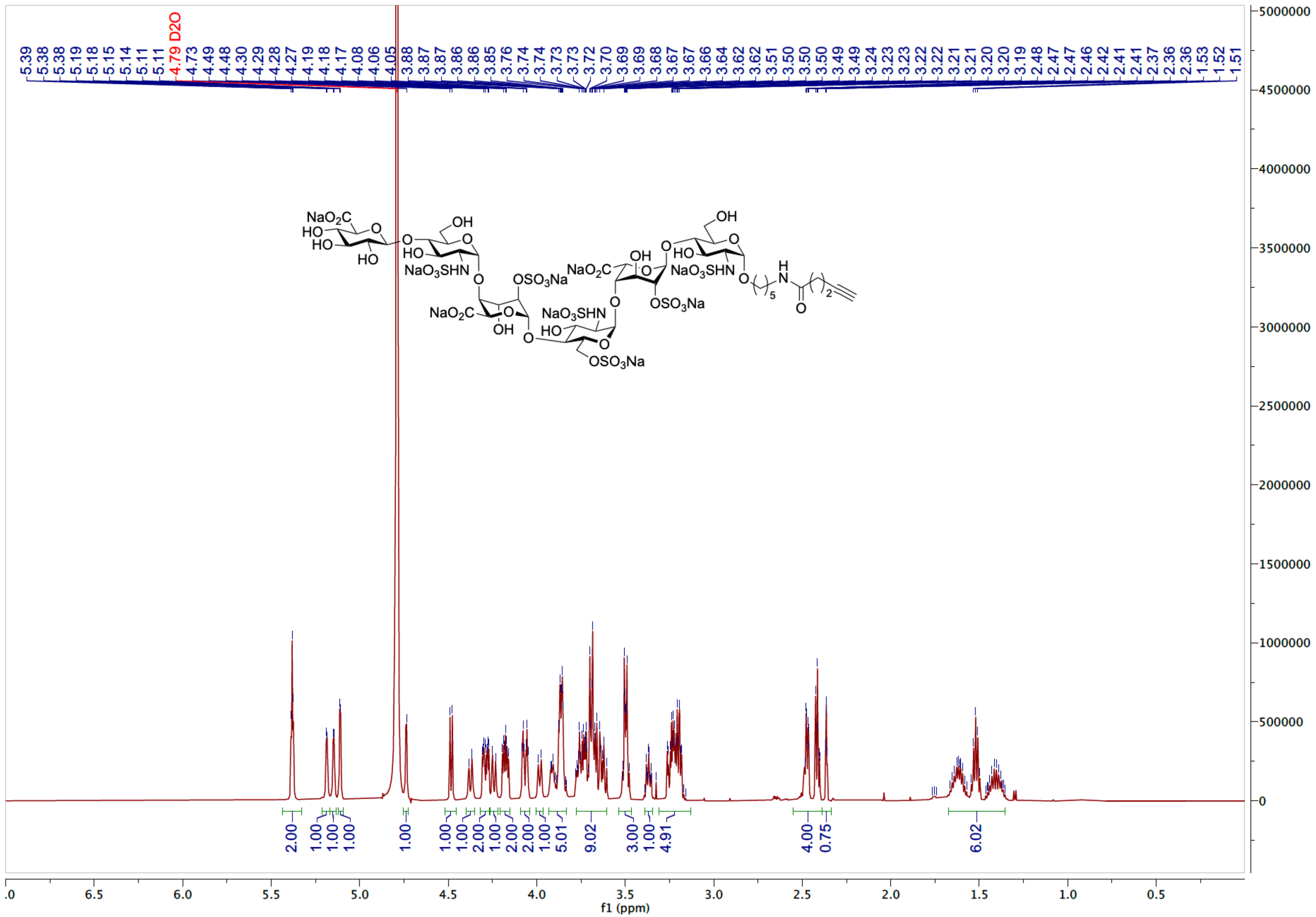

HSQC spectrum of **2** ($D_2O$)

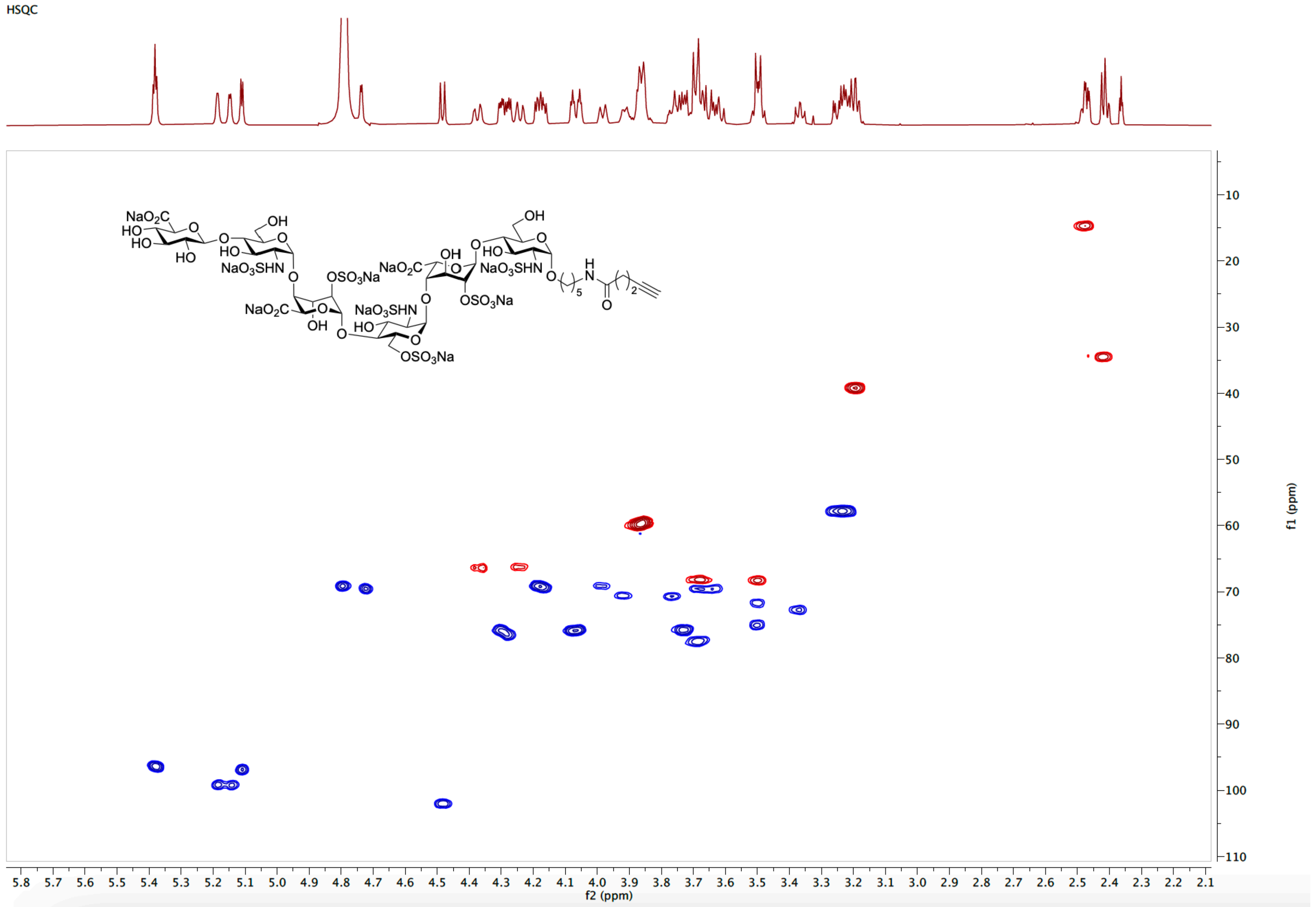

COSY spectrum of **2** ($D_2O$)

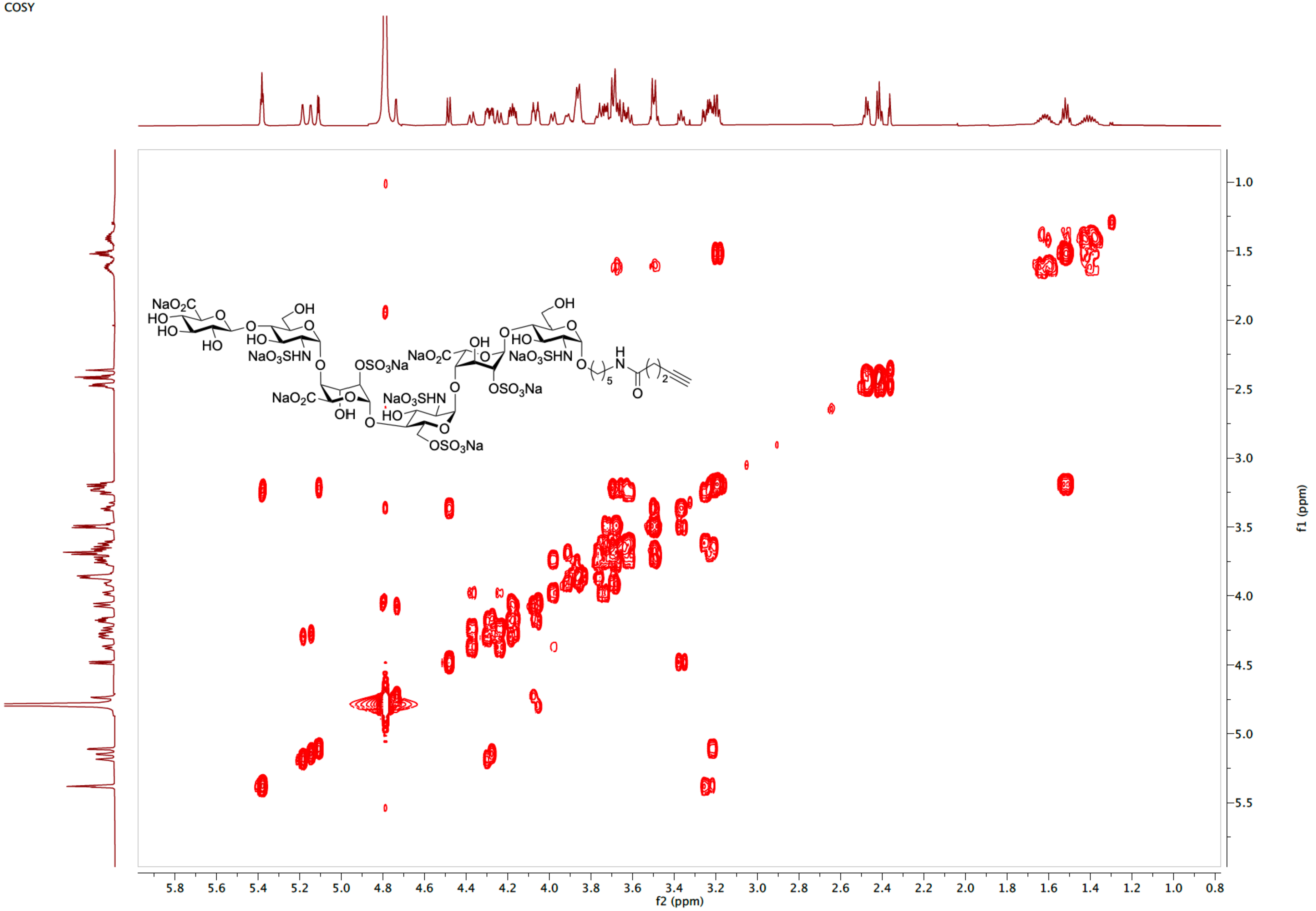

TOCSY spectrum of **2** ($D_2O$)

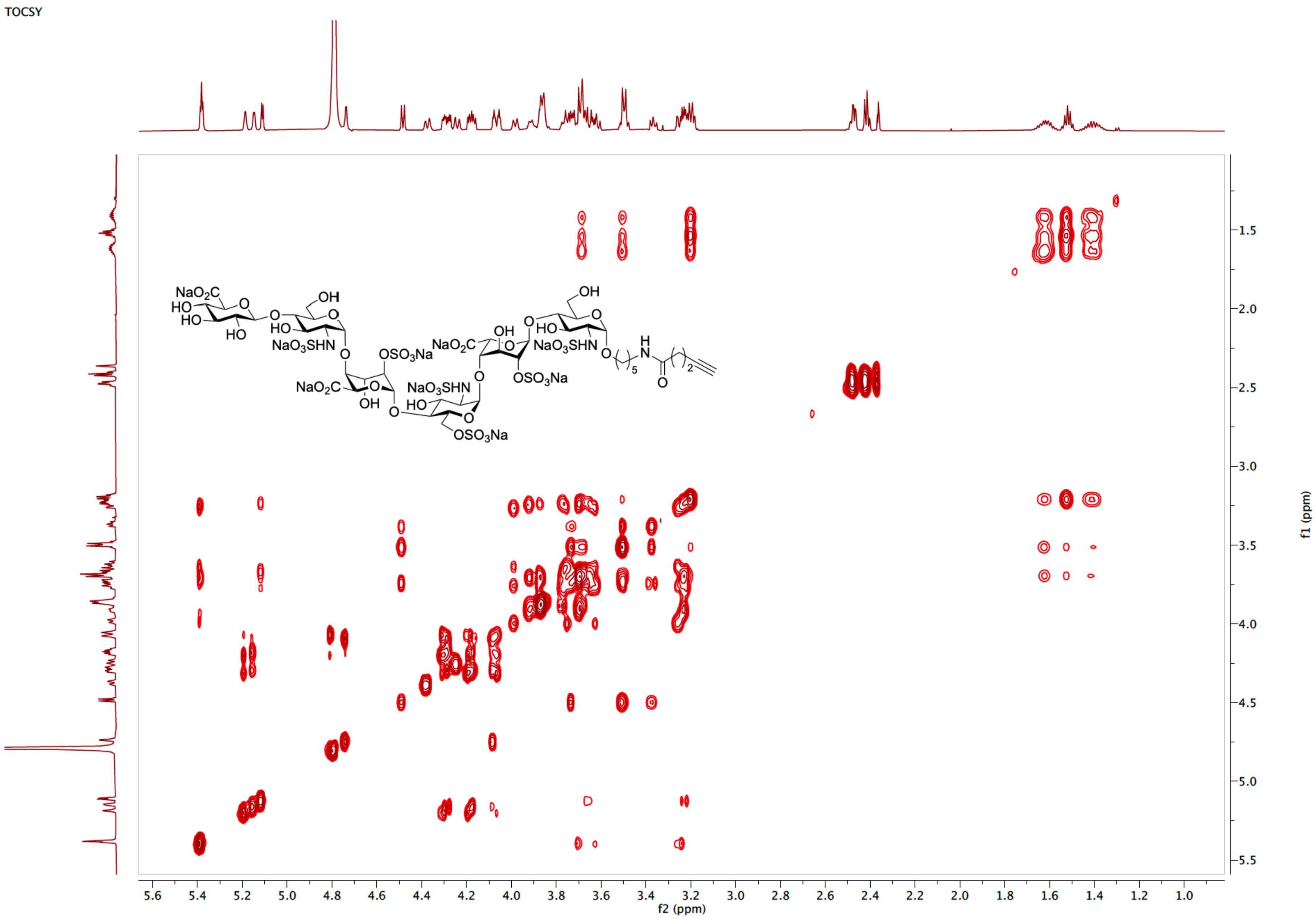

$^{1}H$ NMR spectrum of **S4** (600 MHz, $D_2O$)

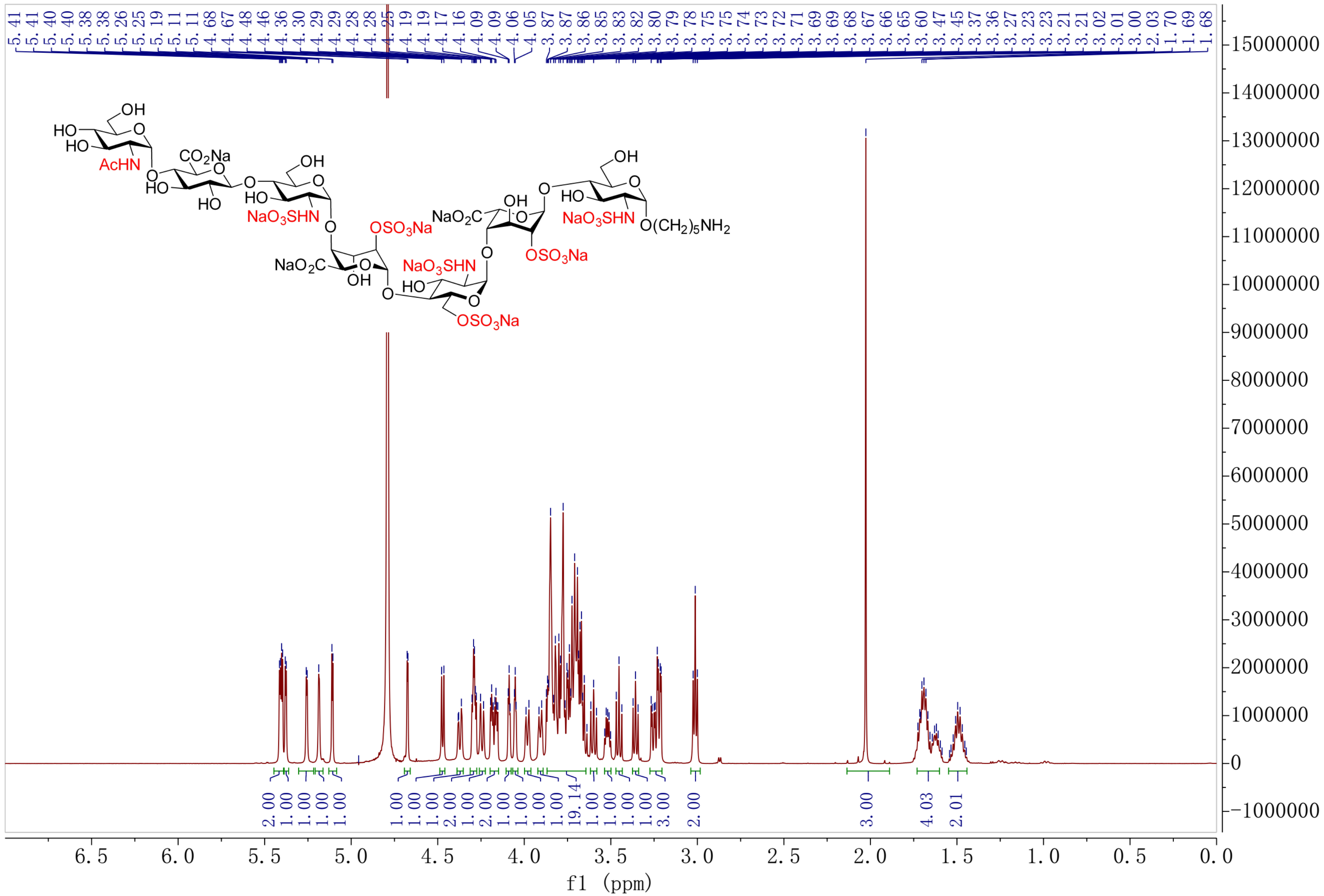

HSQC spectrum of **S4** ($D_2O$)

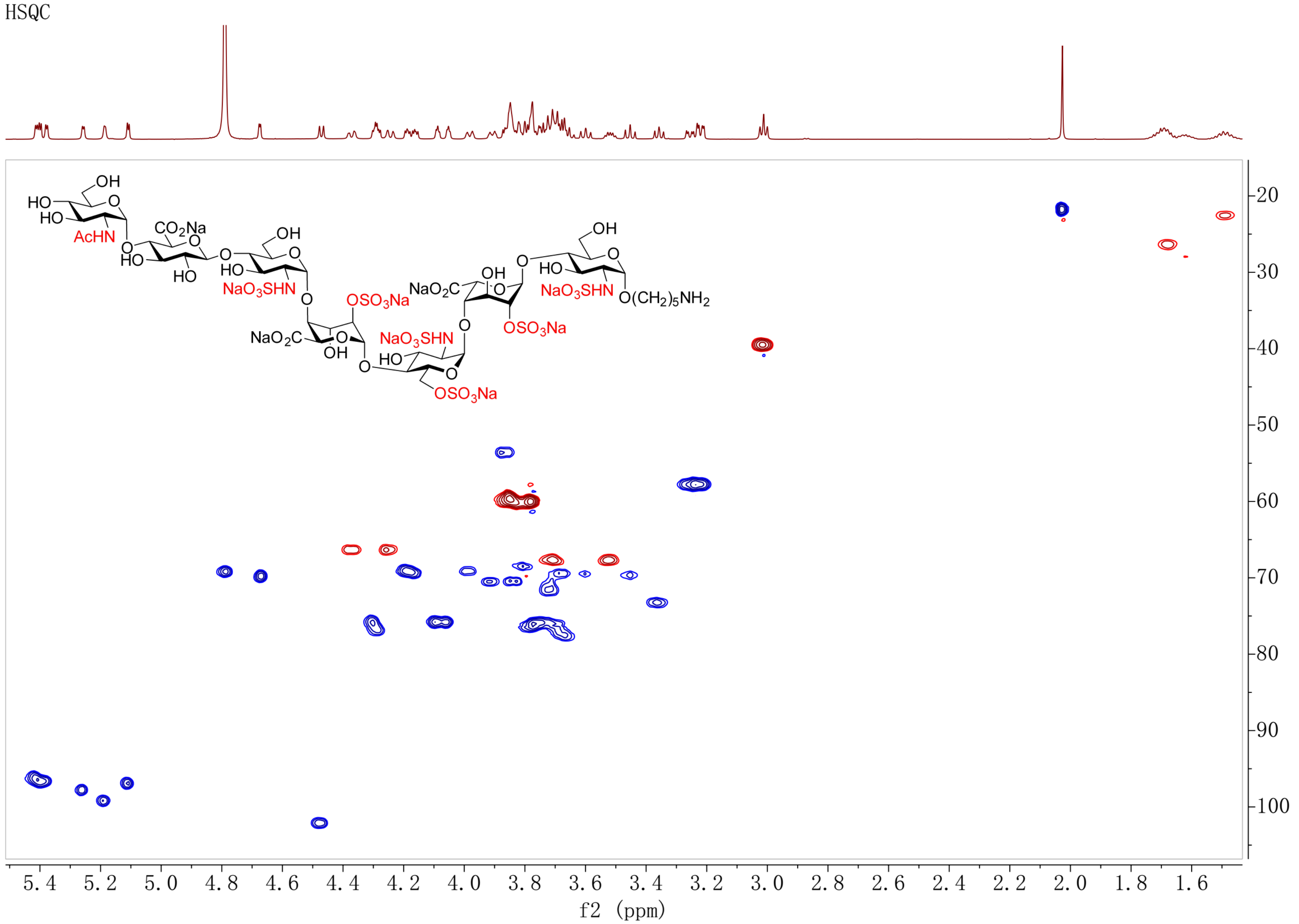

COSY spectrum of **S4** ($D_2O$)

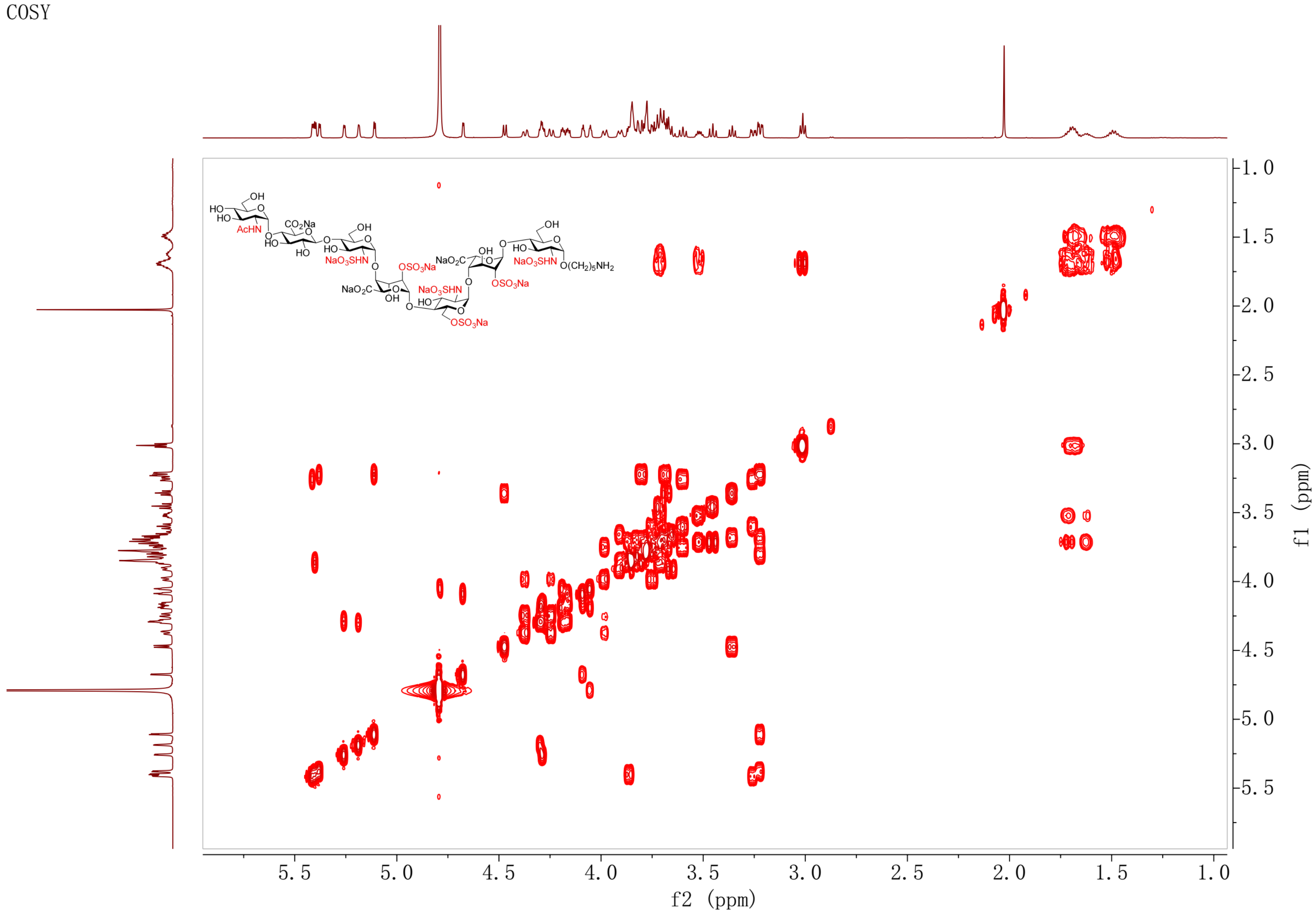

TOCSY spectrum of **S4** ($D_2O$)

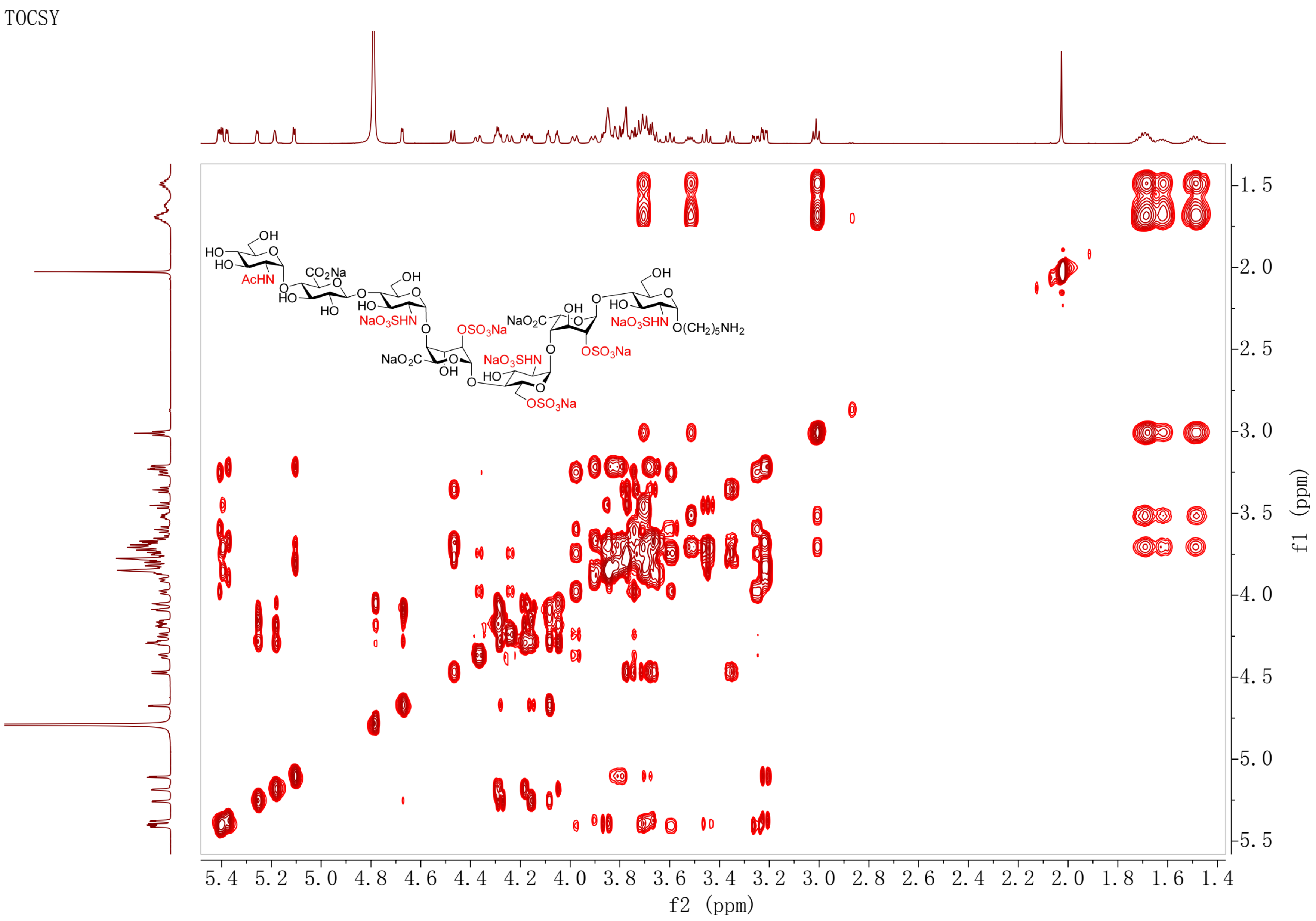

NOESY spectrum of **S4** ($D_2O$)

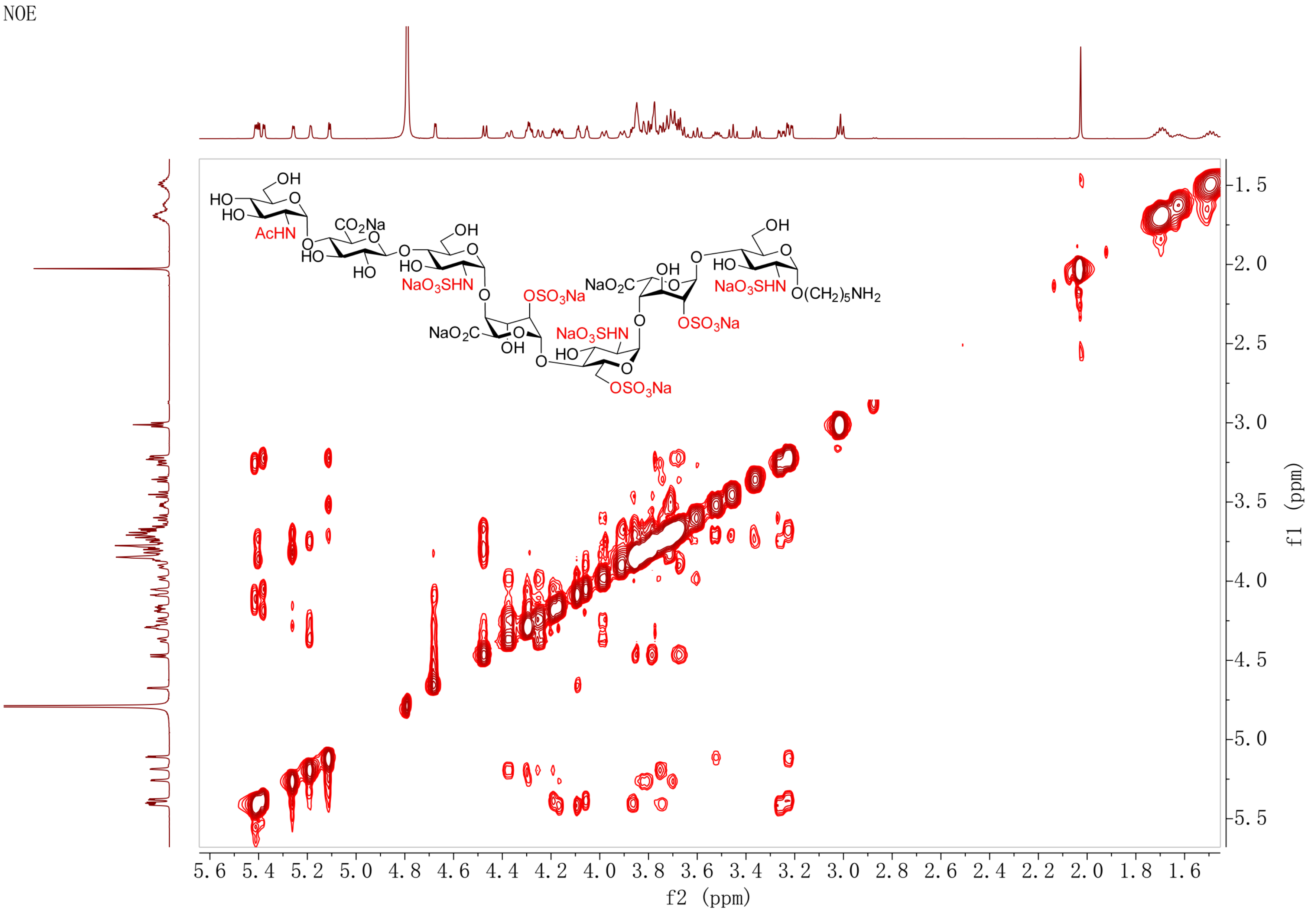

HMBC spectrum of **S4** ($D_2O$)

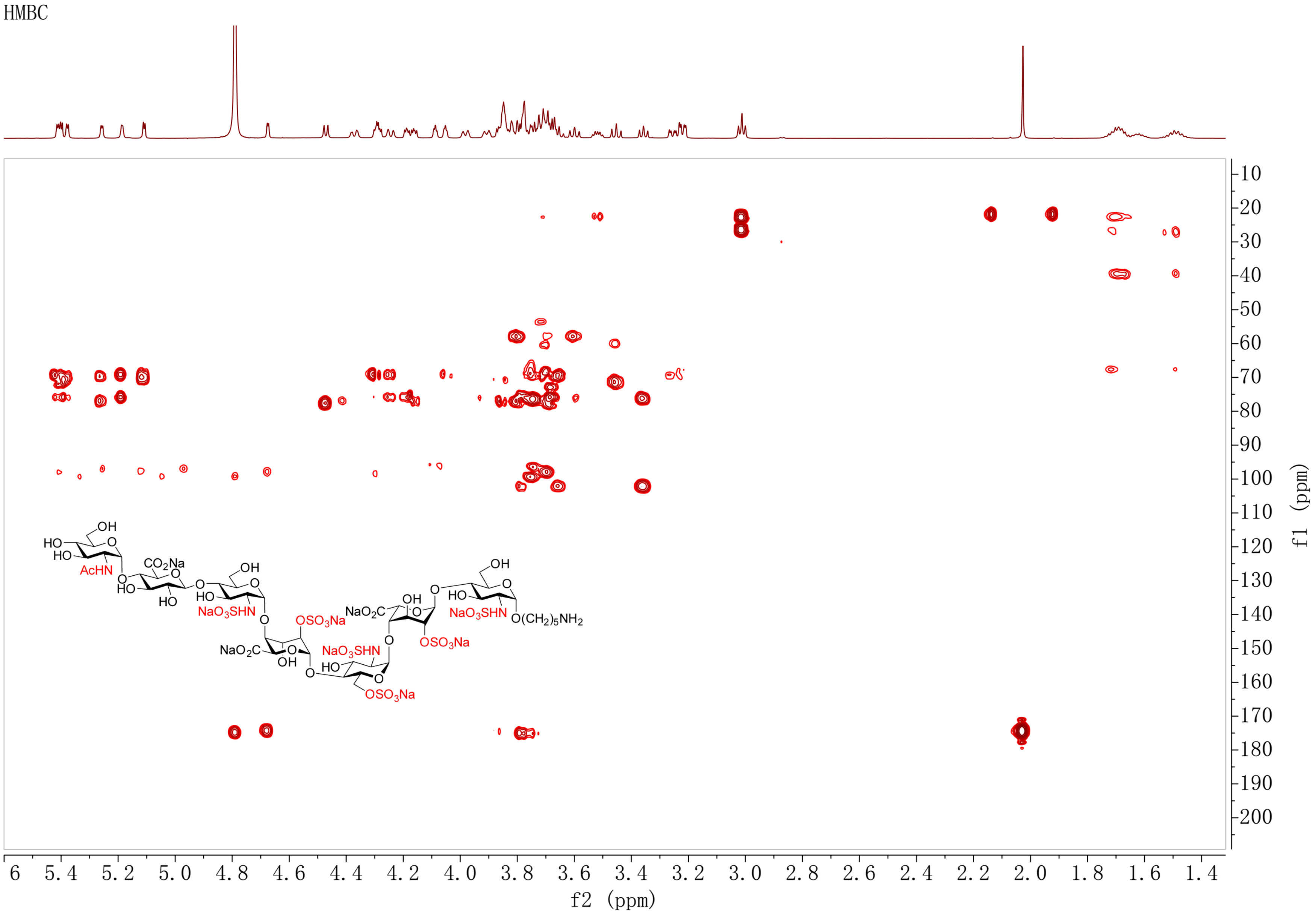

[1]H NMR spectrum of **23** (600 MHz, $D_2O$)

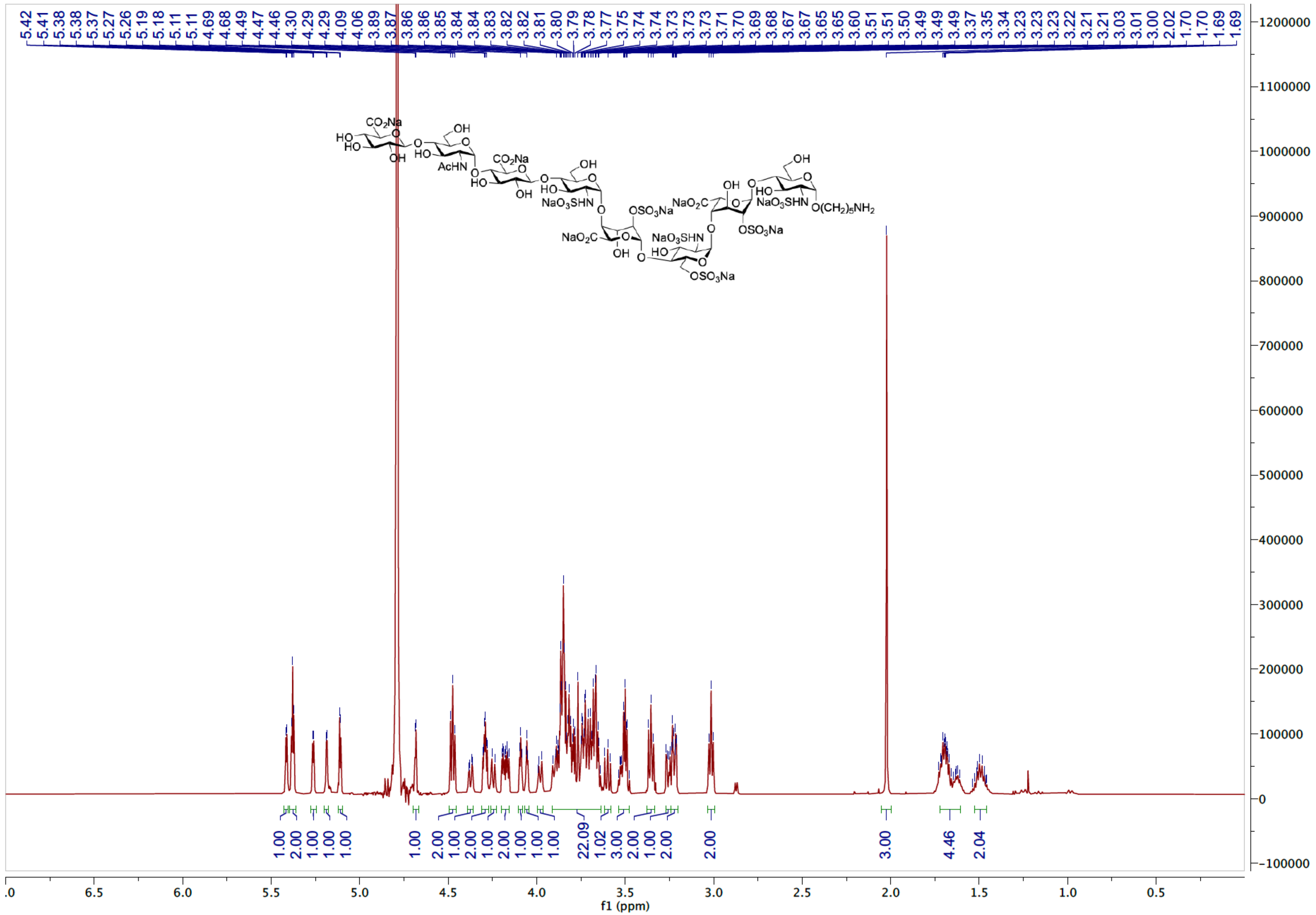

HSQC spectrum of **23** ($D_2O$)

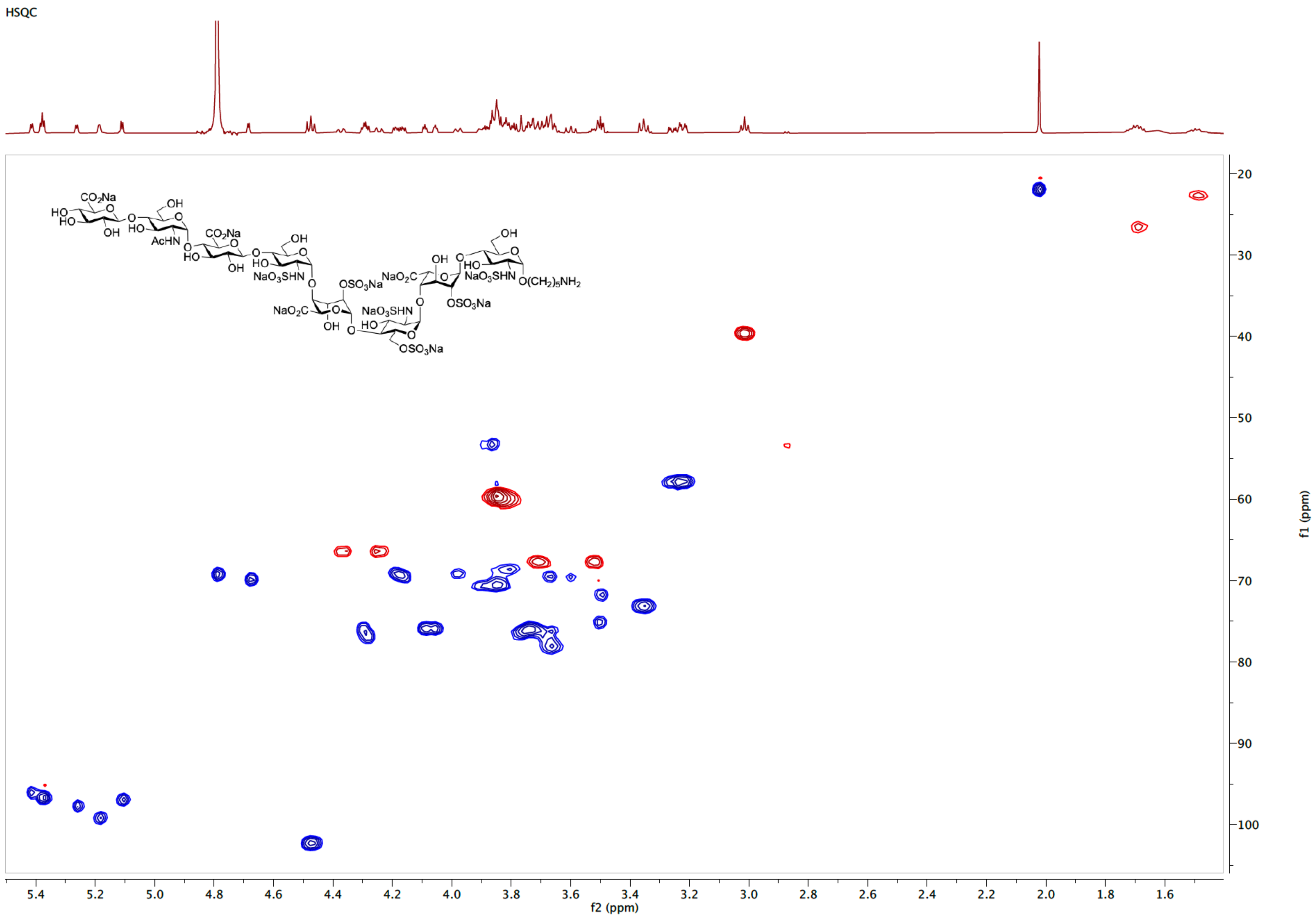

COSY spectrum of **23** ($D_2O$)

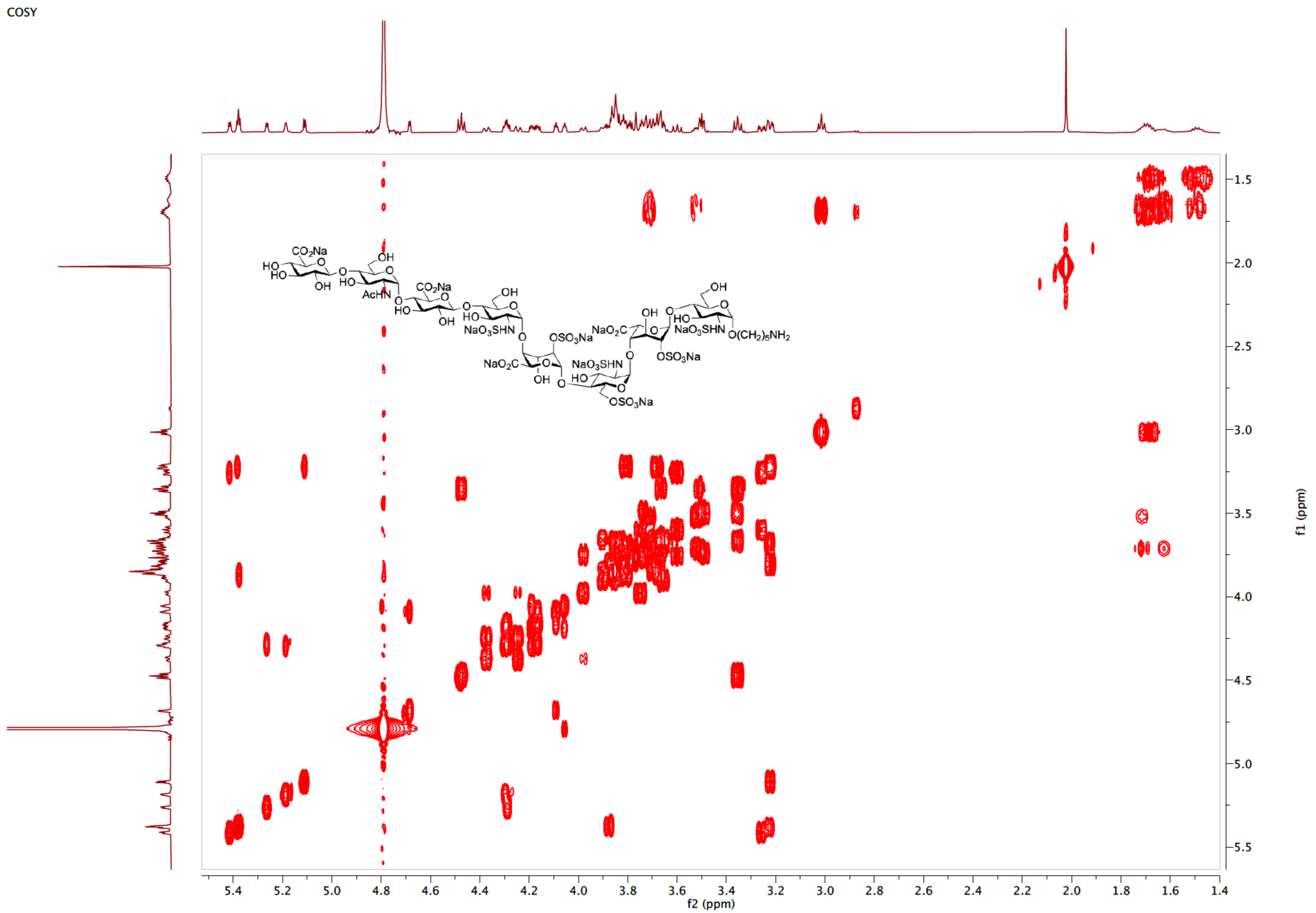

TOCSY spectrum of **23** ($D_2O$)

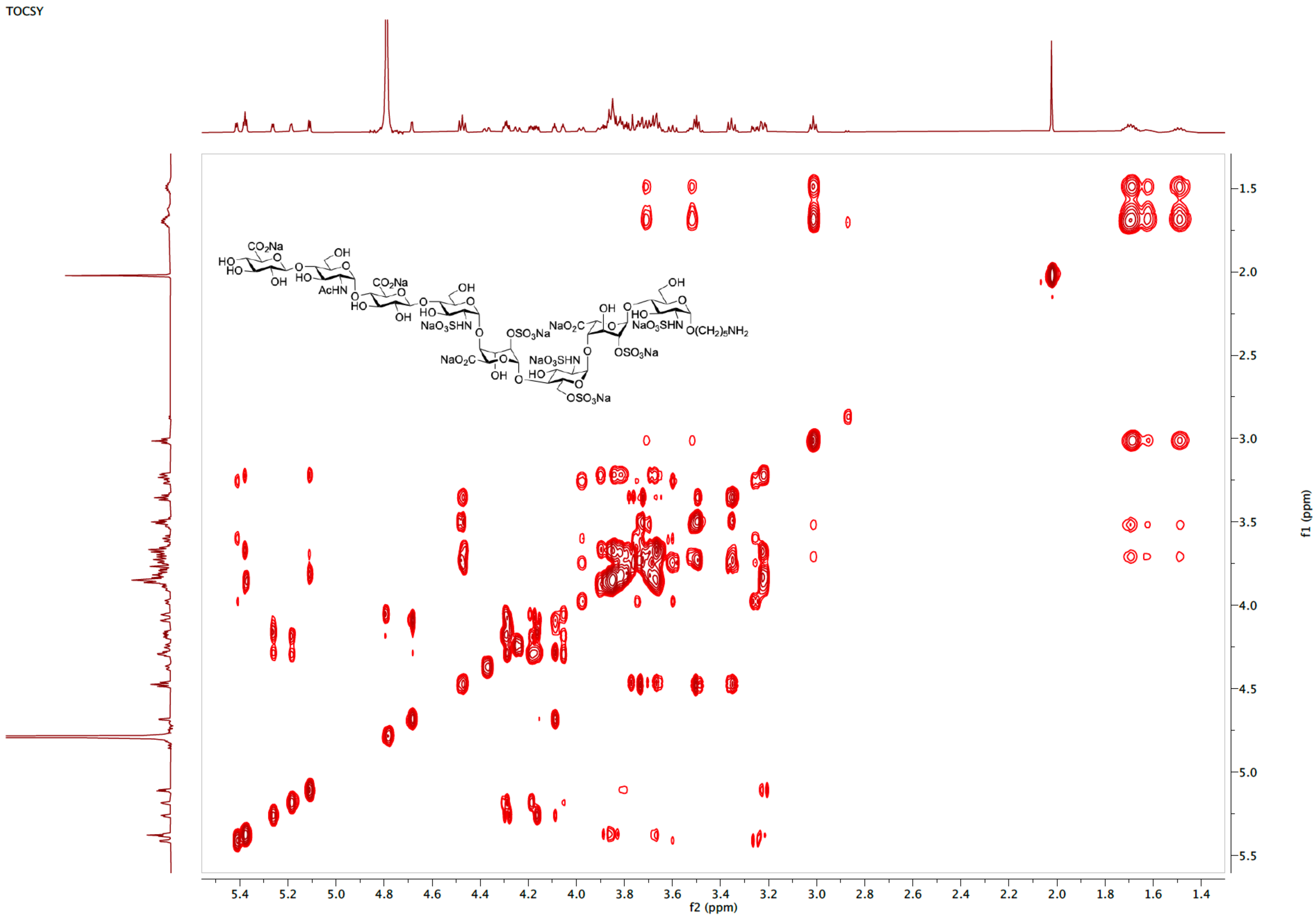

NOESY spectrum of **23** ($D_2O$)

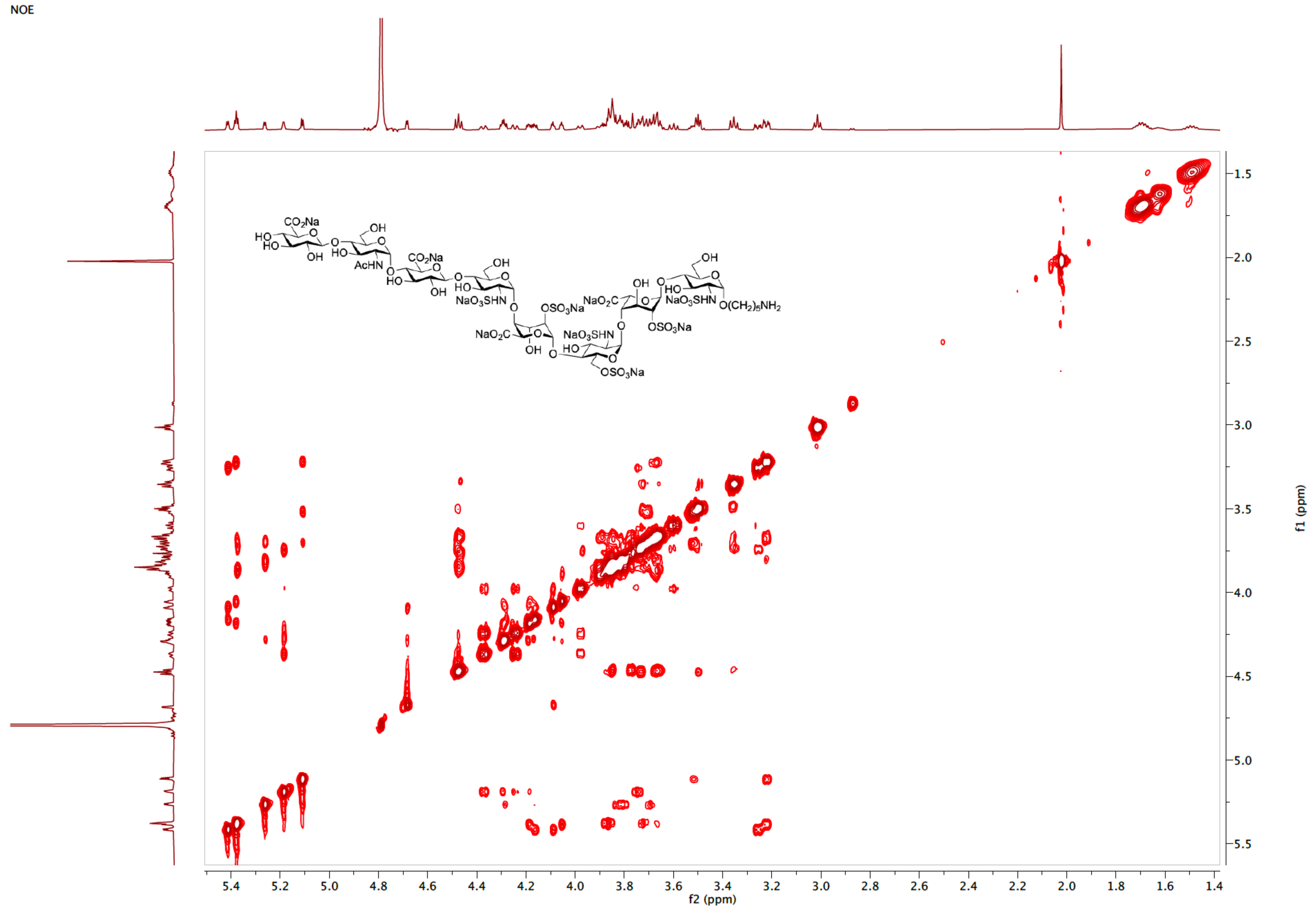

HMBC spectrum of **23** ($D_2O$)

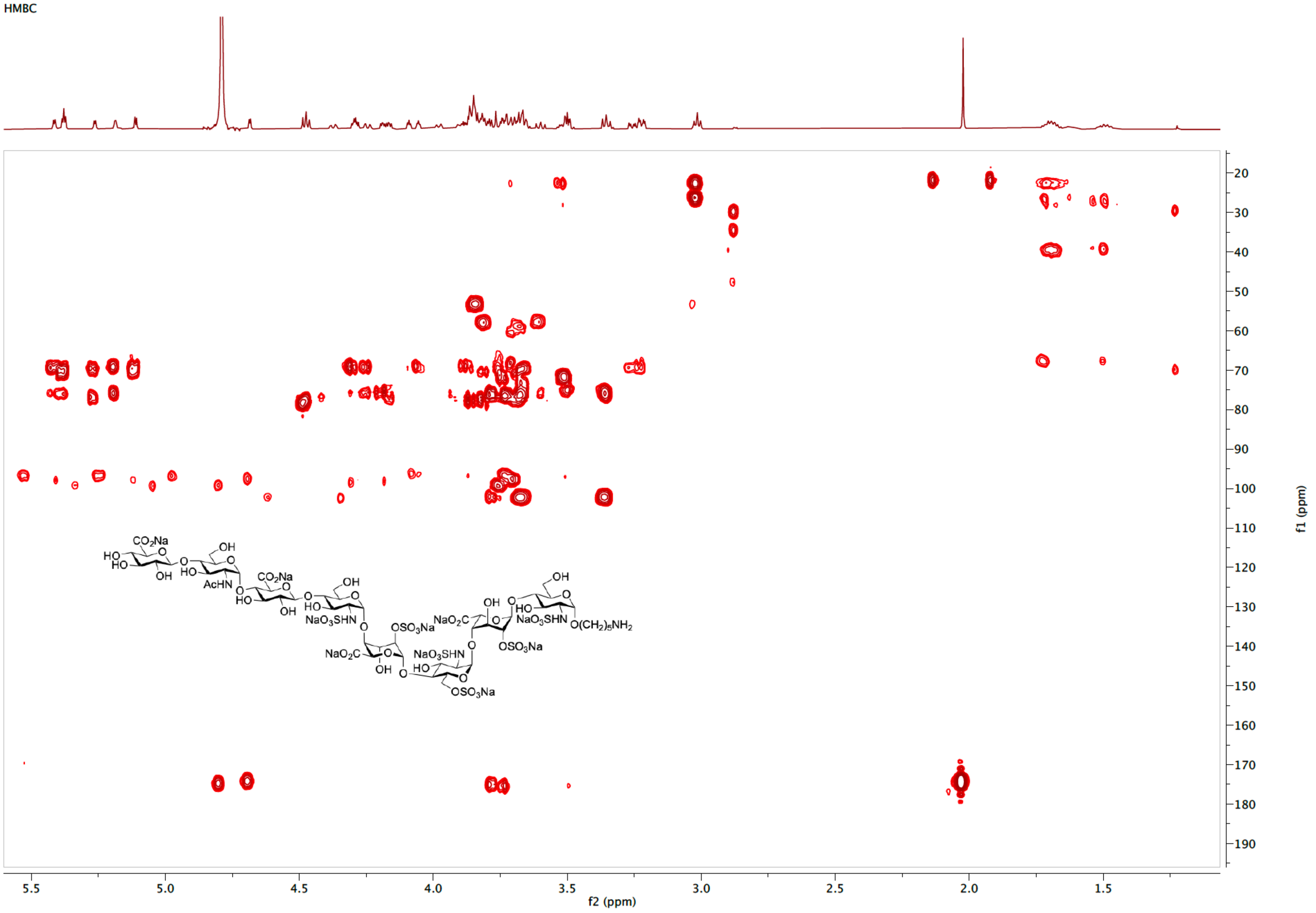

[1]H NMR spectrum of **S5** (600 MHz, $D_2O$)

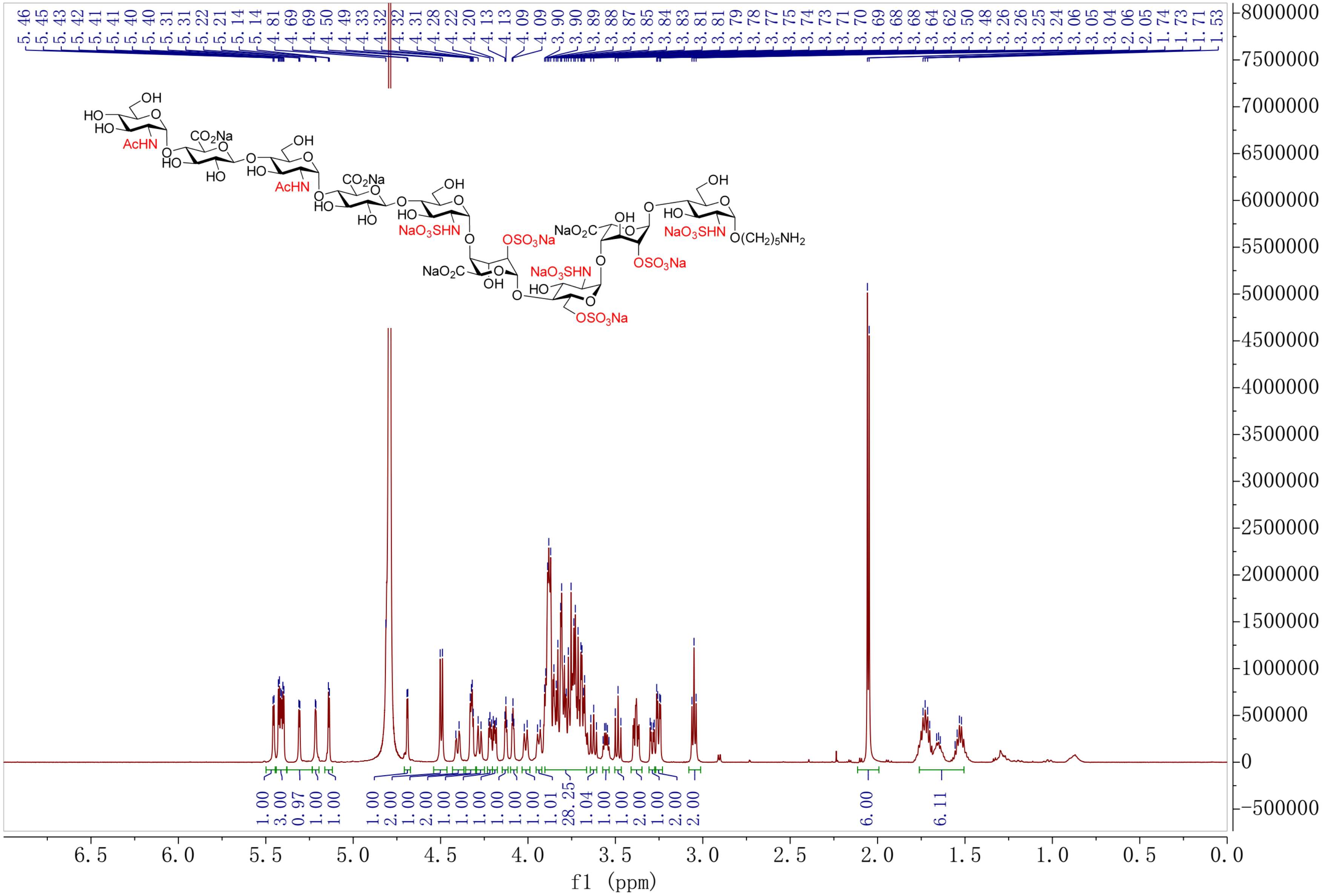

HSQC spectrum of **S5** ($D_2O$)

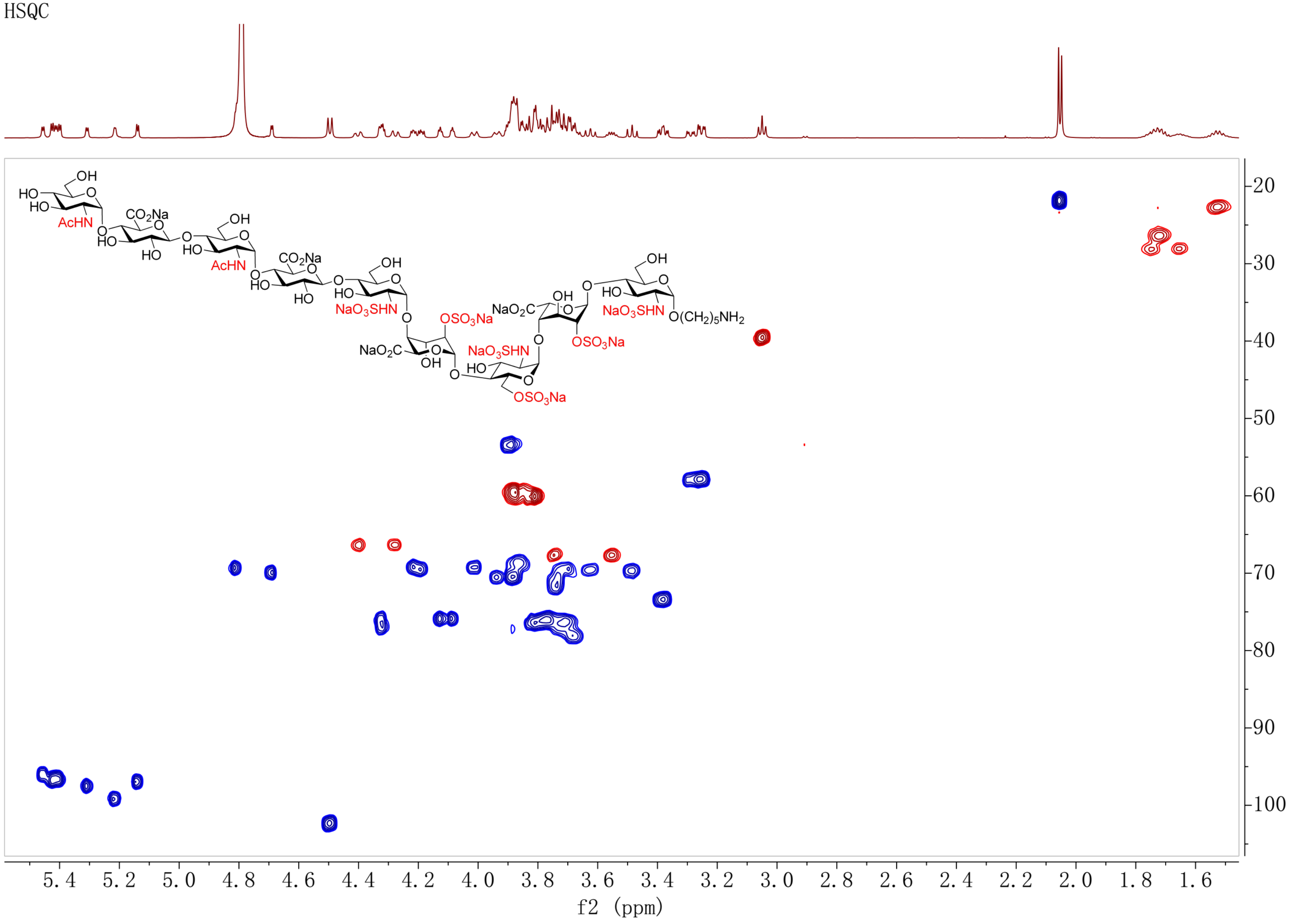

COSY spectrum of **S5** ($D_2O$)

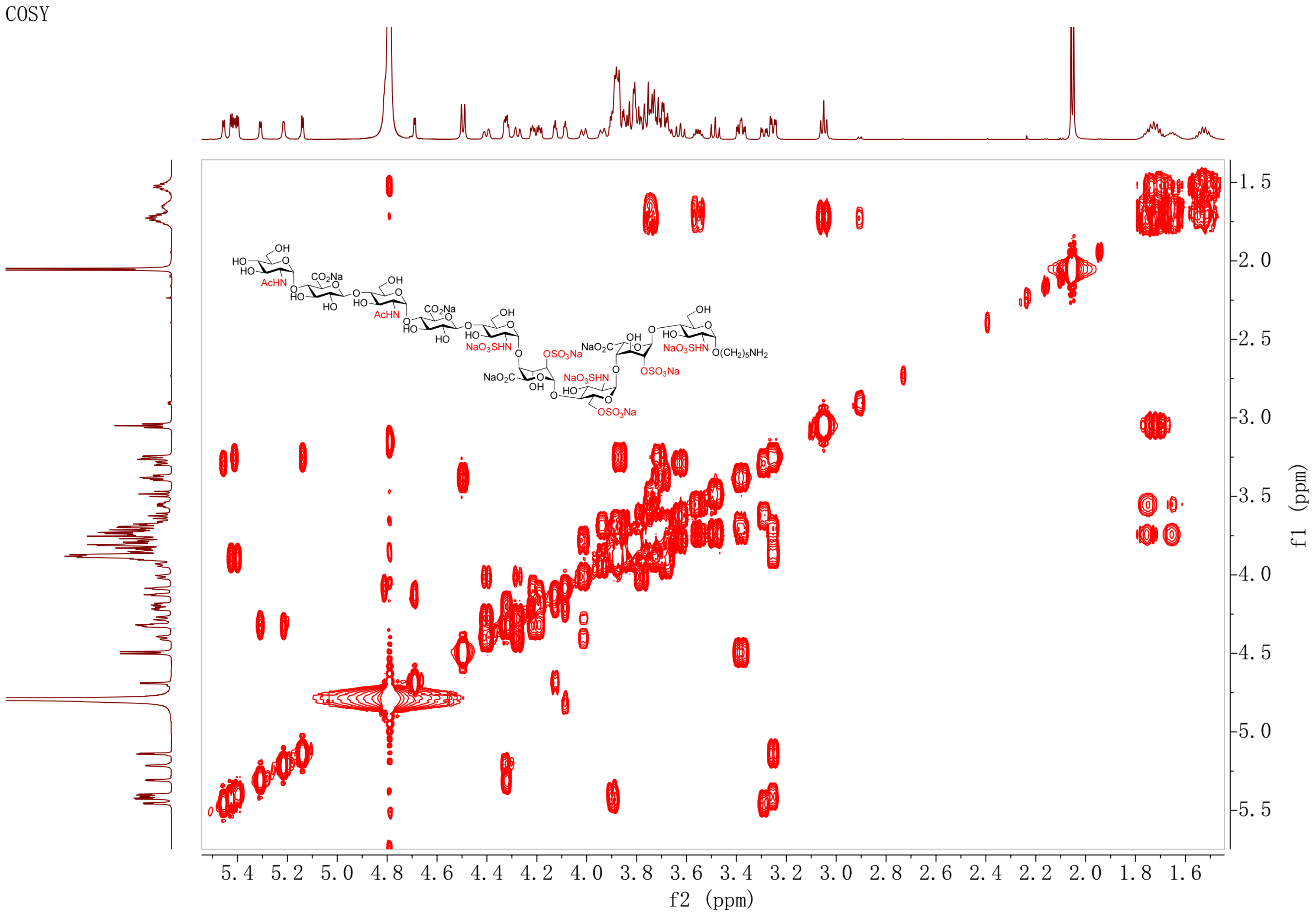

TOCSY spectrum of **S5** ($D_2O$)

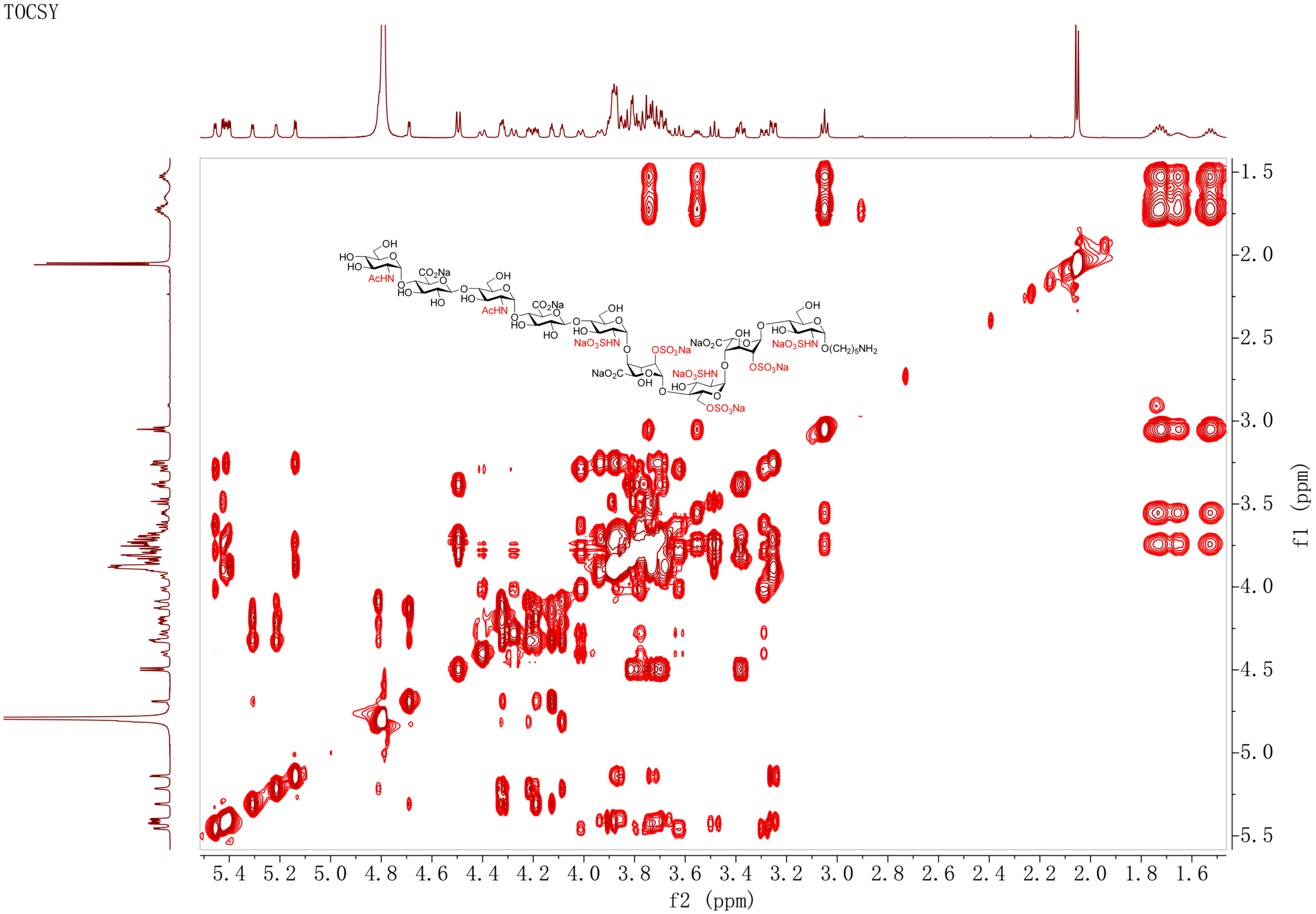

NOESY spectrum of **S5** ($D_2O$)

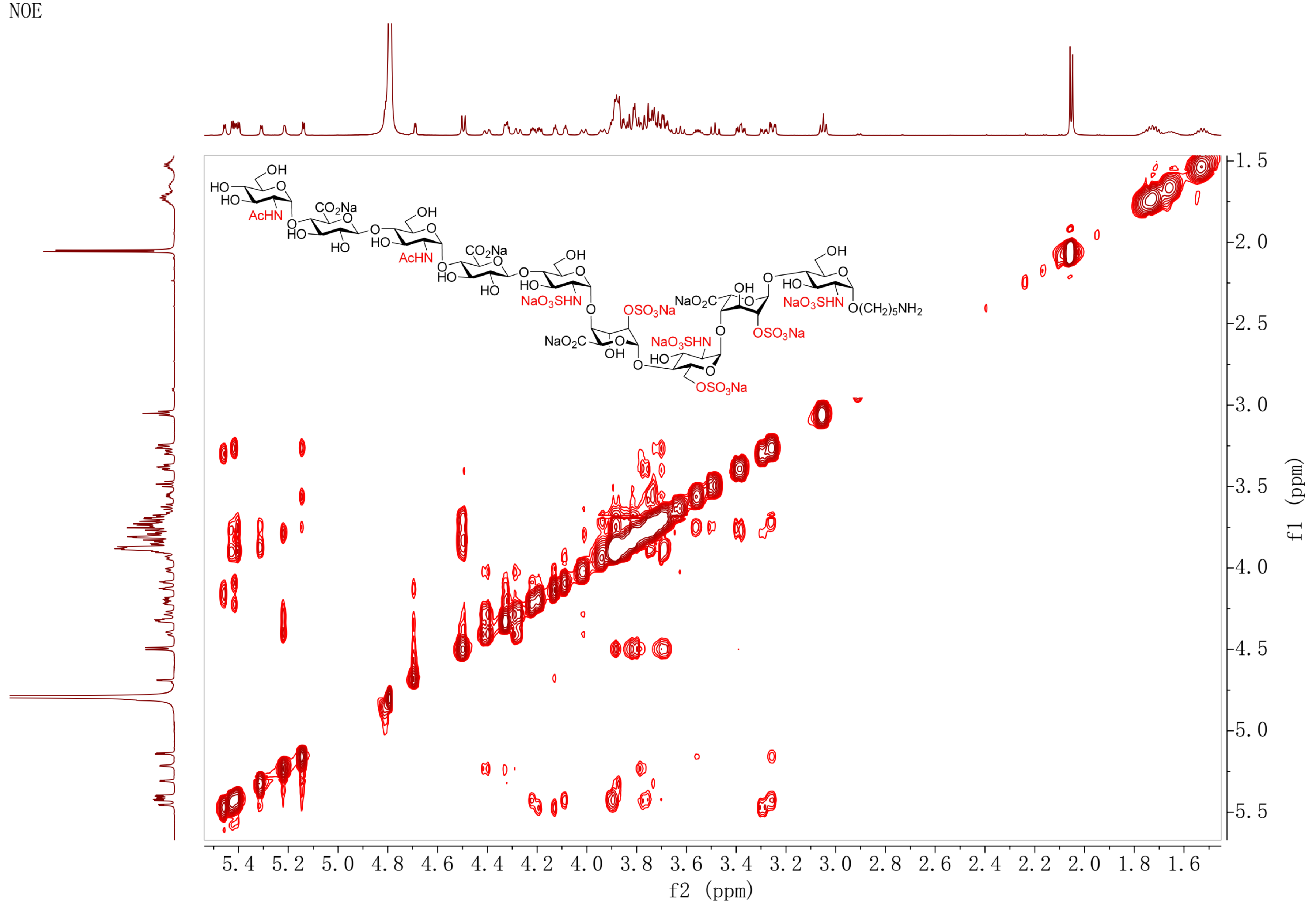

HMBC spectrum of **S5** ($D_2O$)

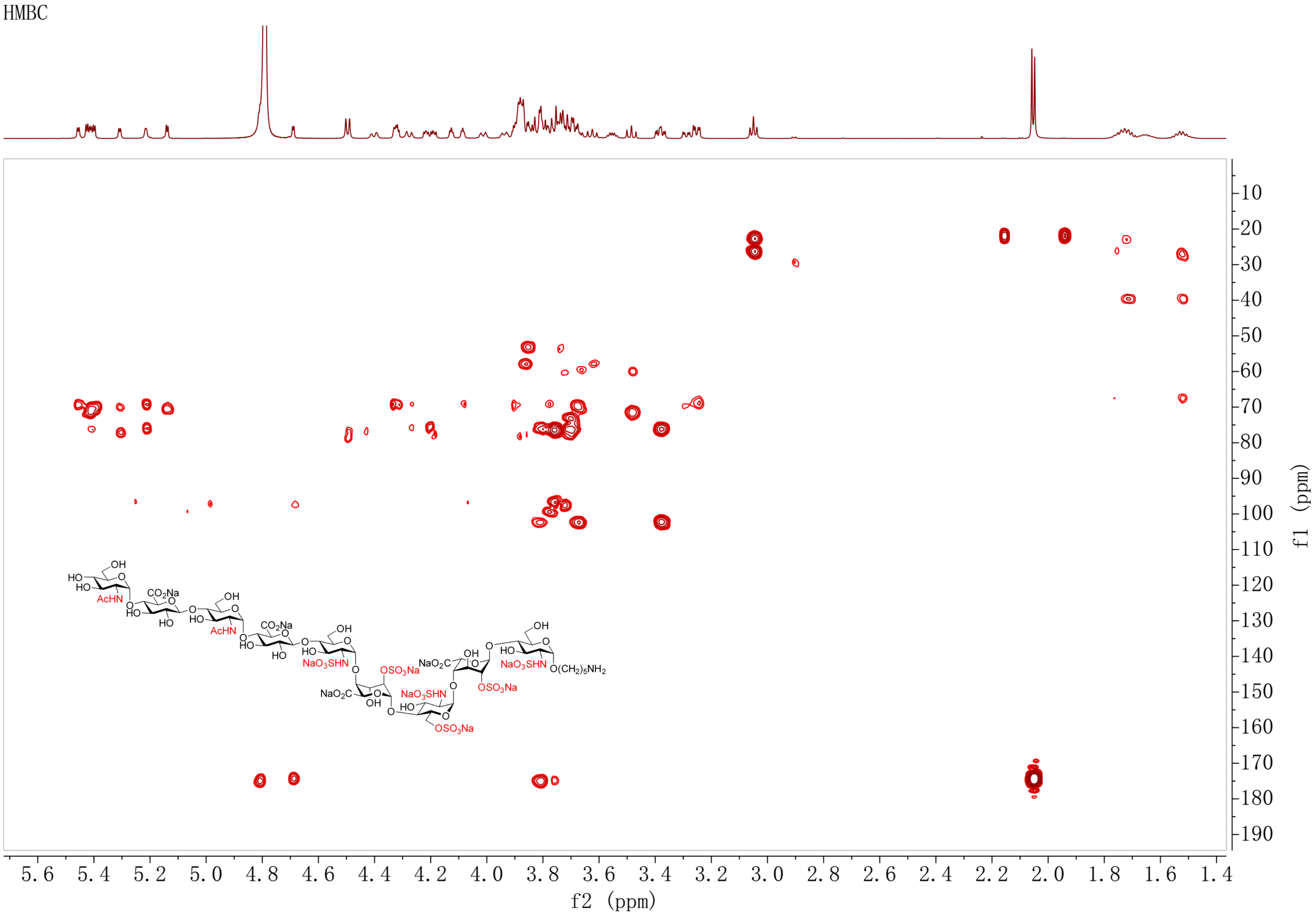

[1]H NMR spectrum of **24** (600 MHz, $D_2O$)

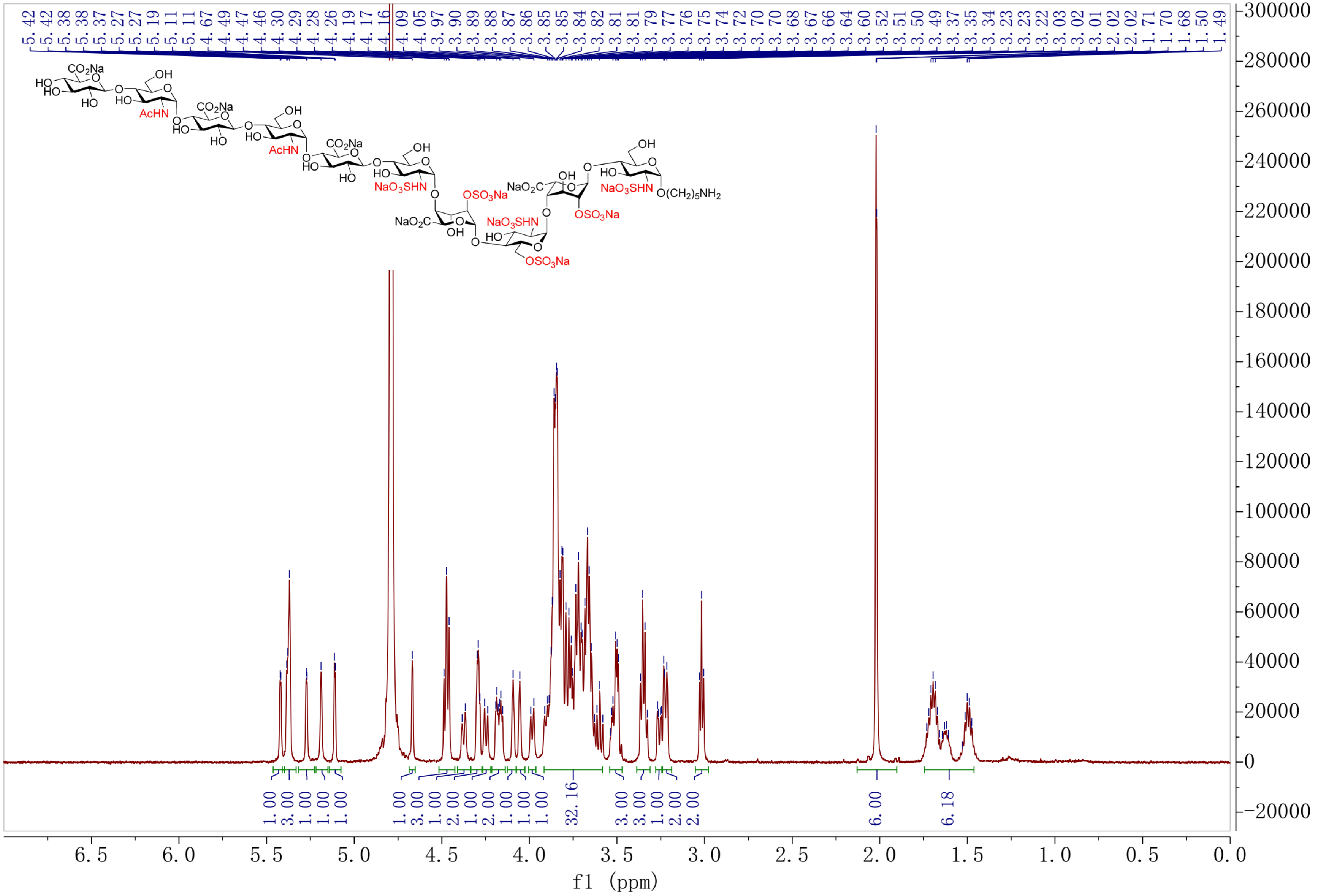

HSQC spectrum of **24** ($D_2O$)

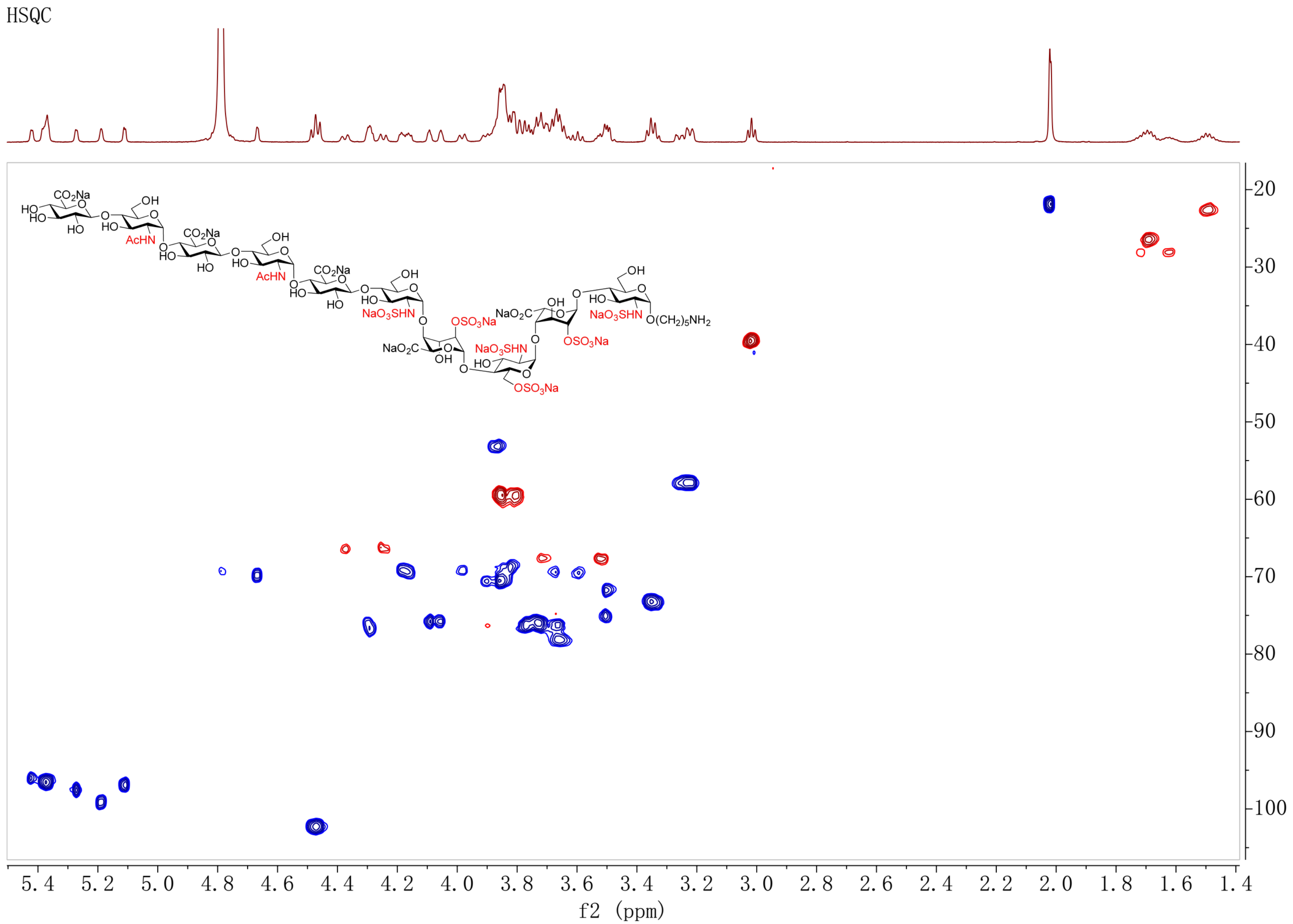

COSY spectrum of **24** ($D_2O$)

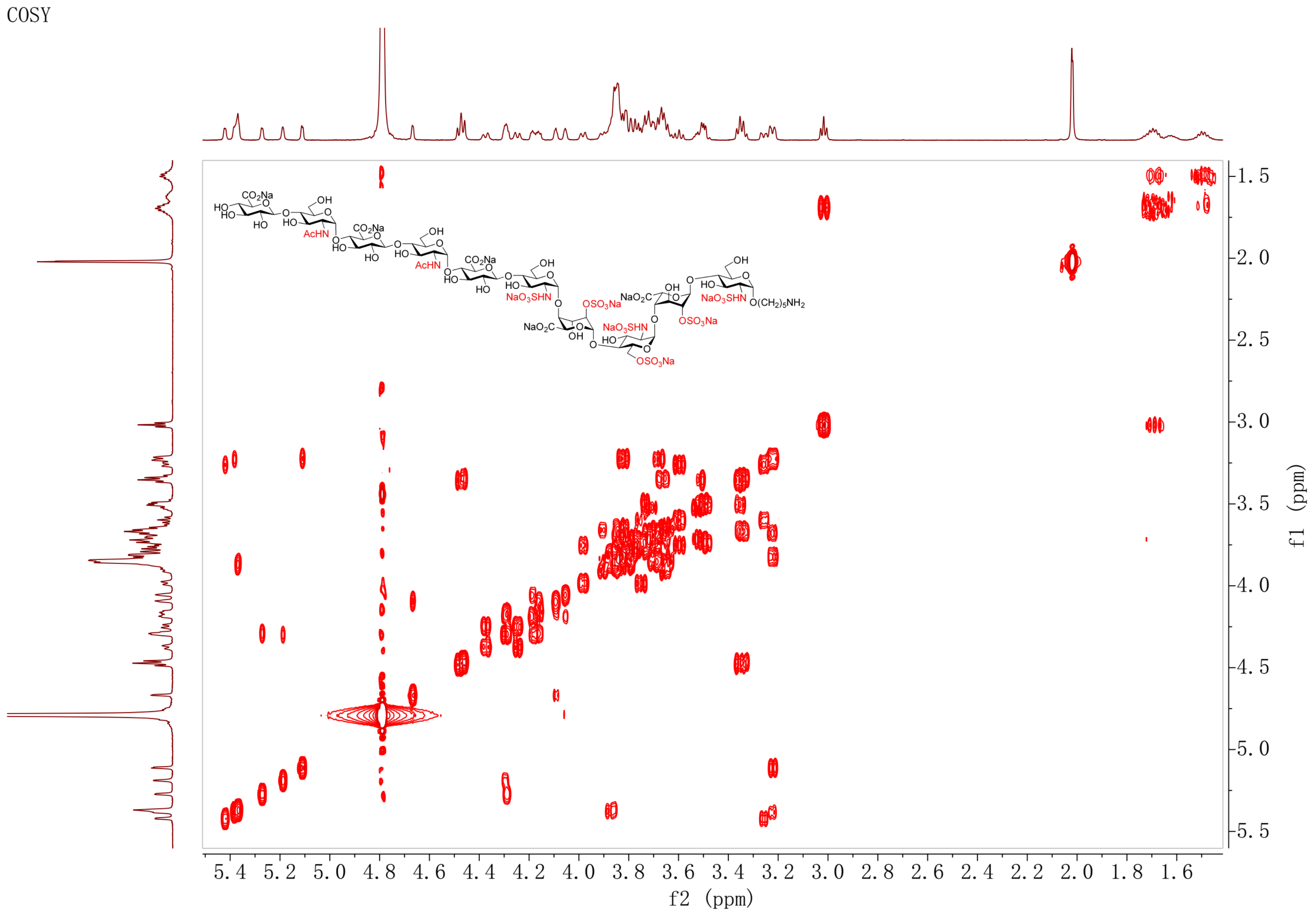

TOCSY spectrum of **24** ($D_2O$)

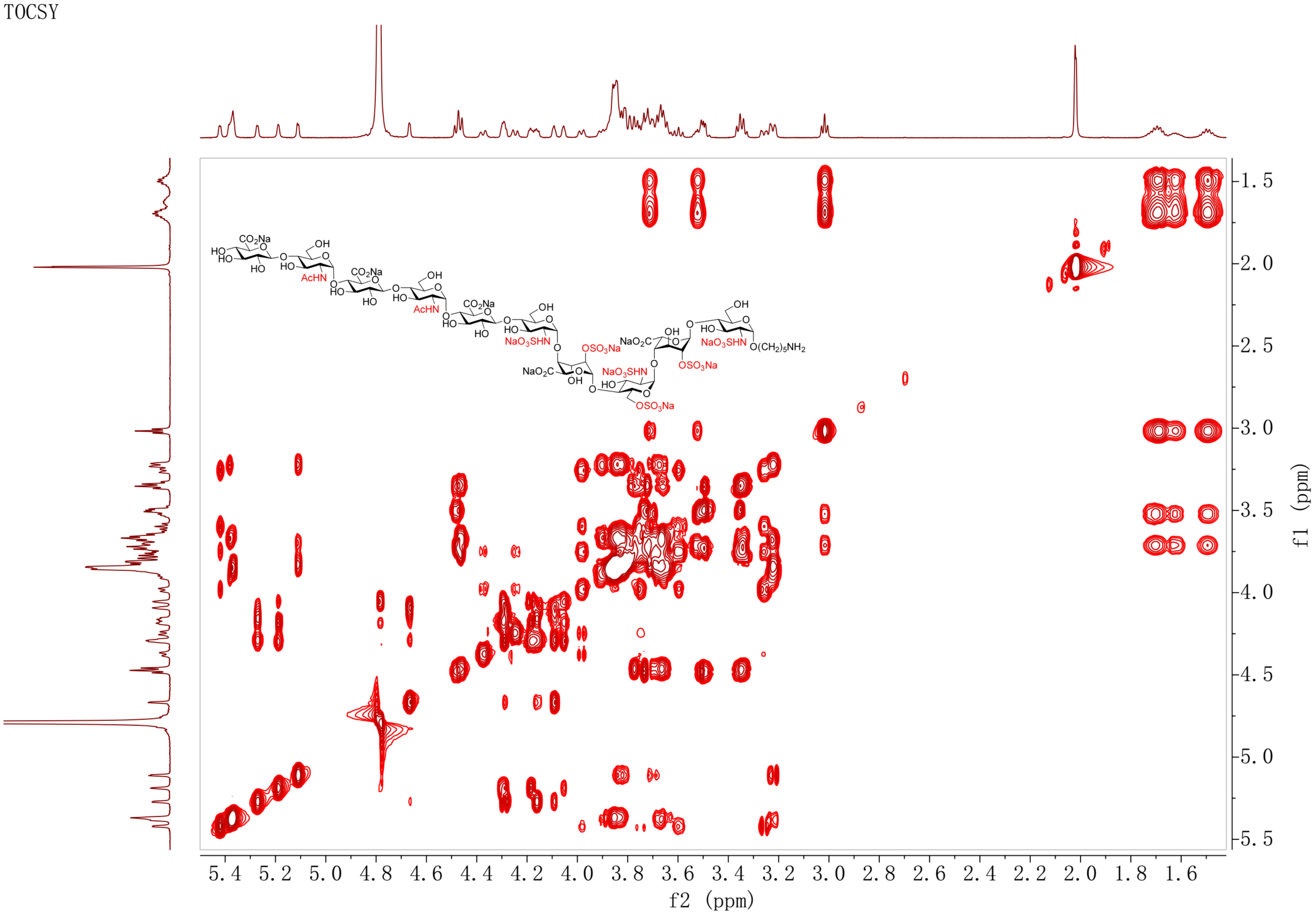

NOESY spectrum of **24** ($D_2O$)

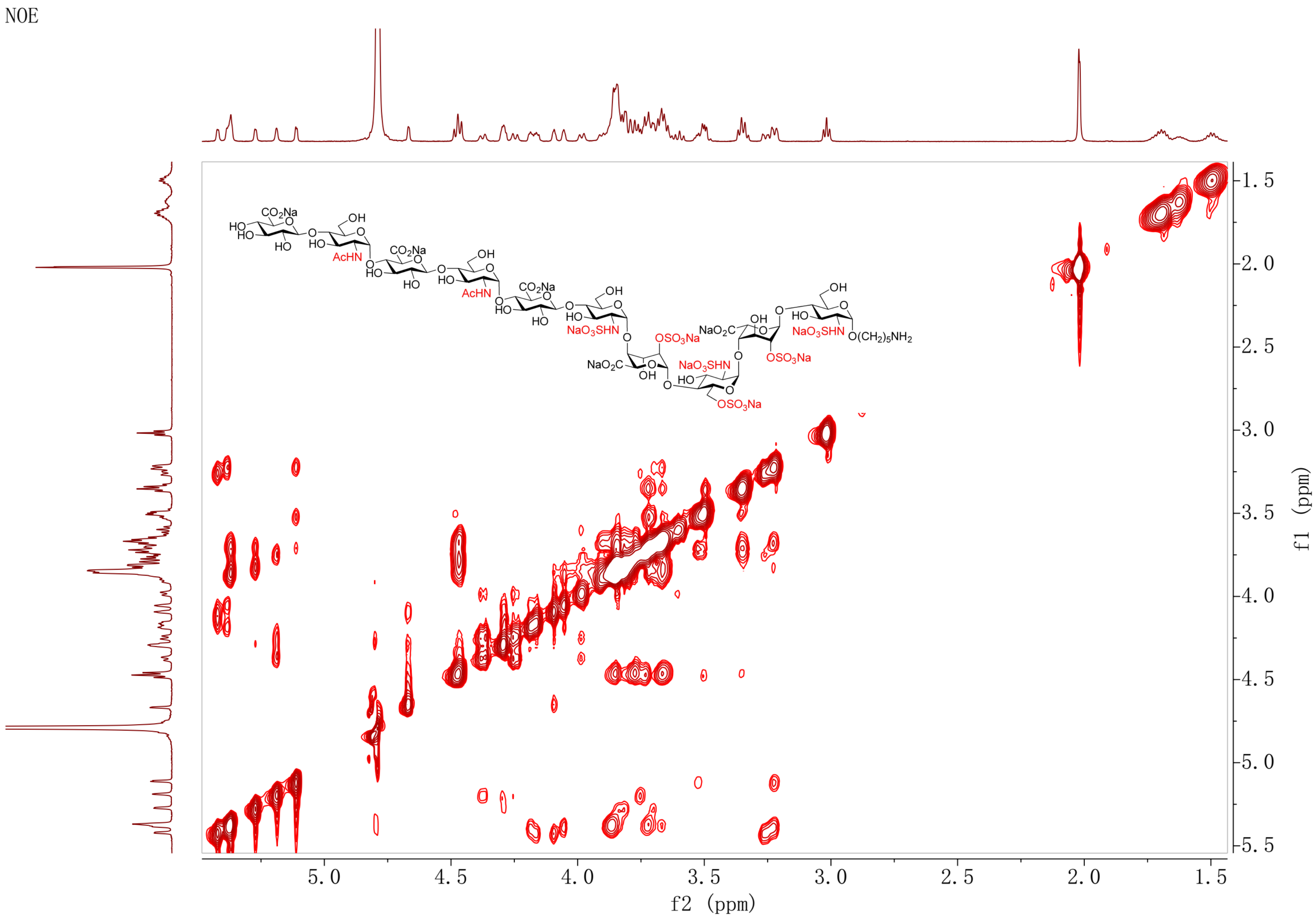

[1]H NMR spectrum of **25** (600 MHz, $D_2O$)

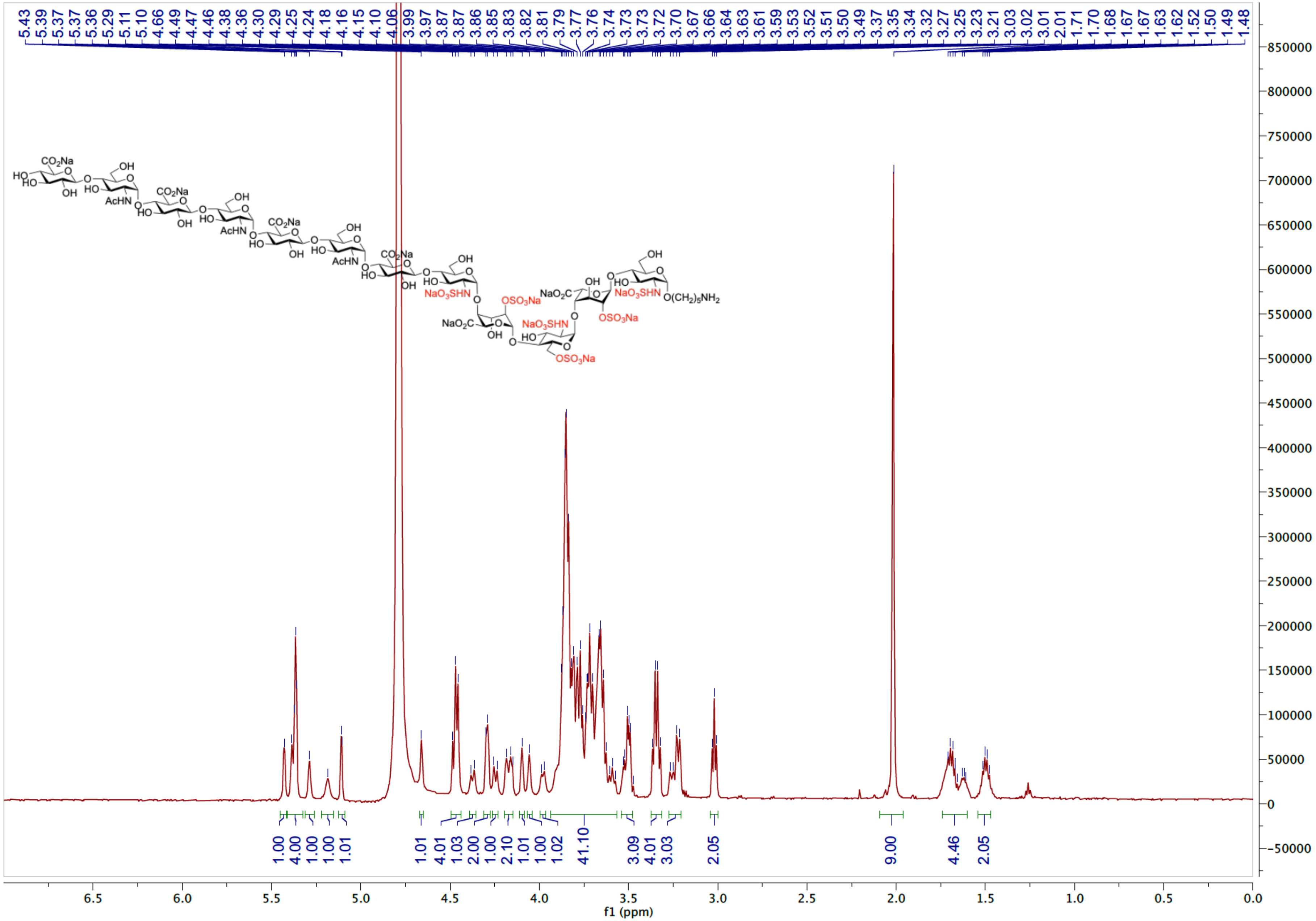

HSQC spectrum of **25** ($D_2O$)

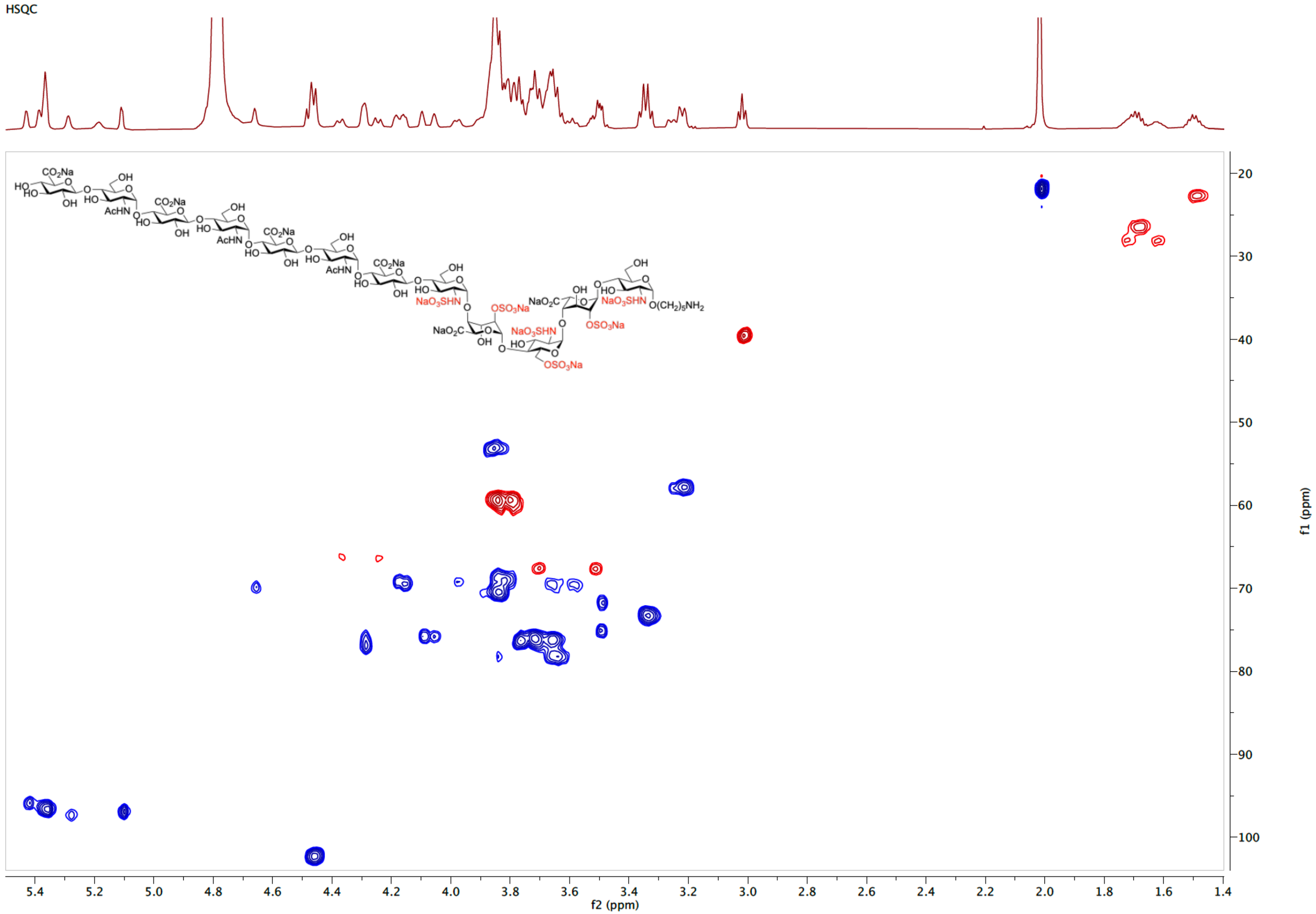

COSY spectrum of **25** ($D_2O$)

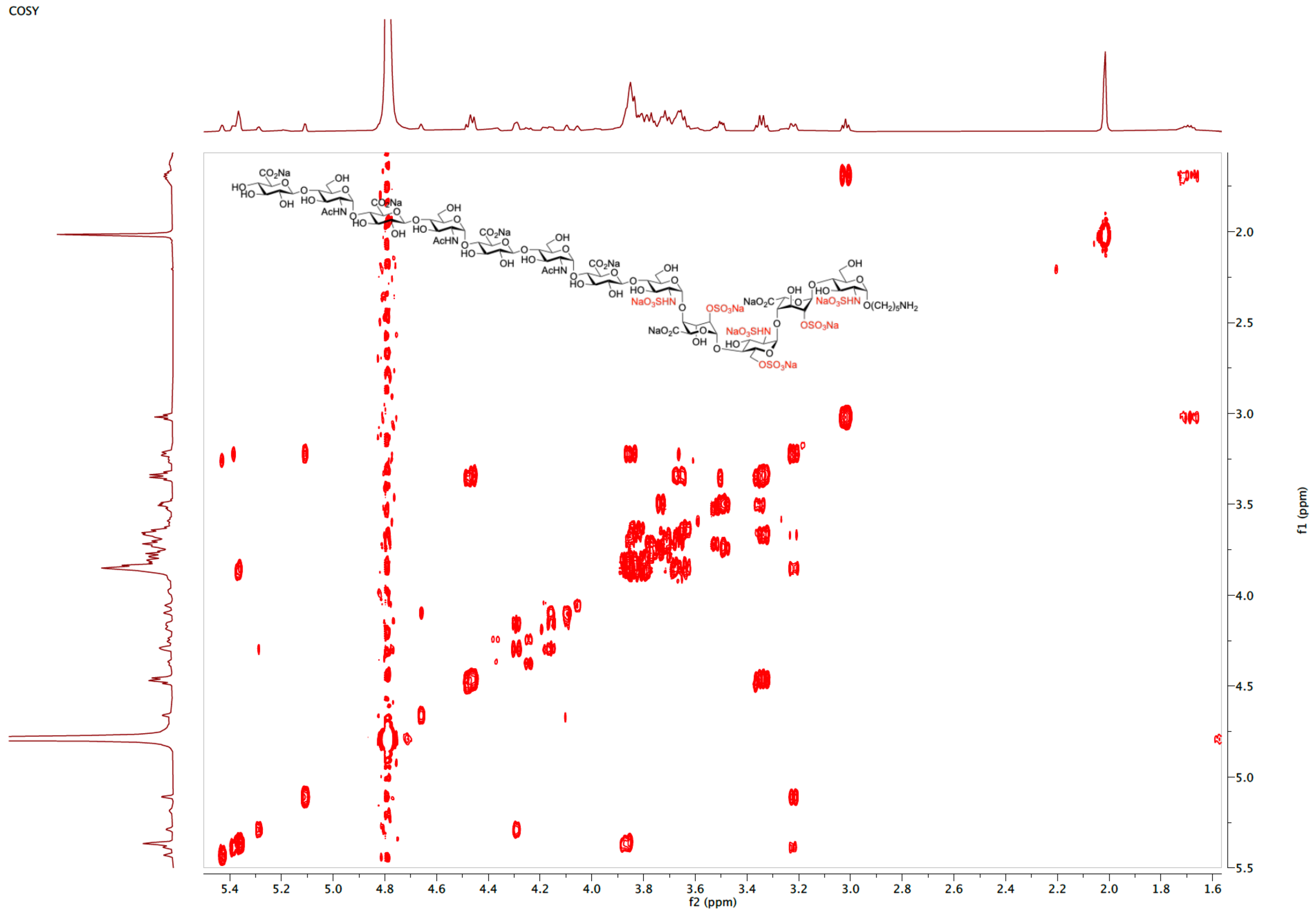

TOCSY spectrum of **25** ($D_2O$)

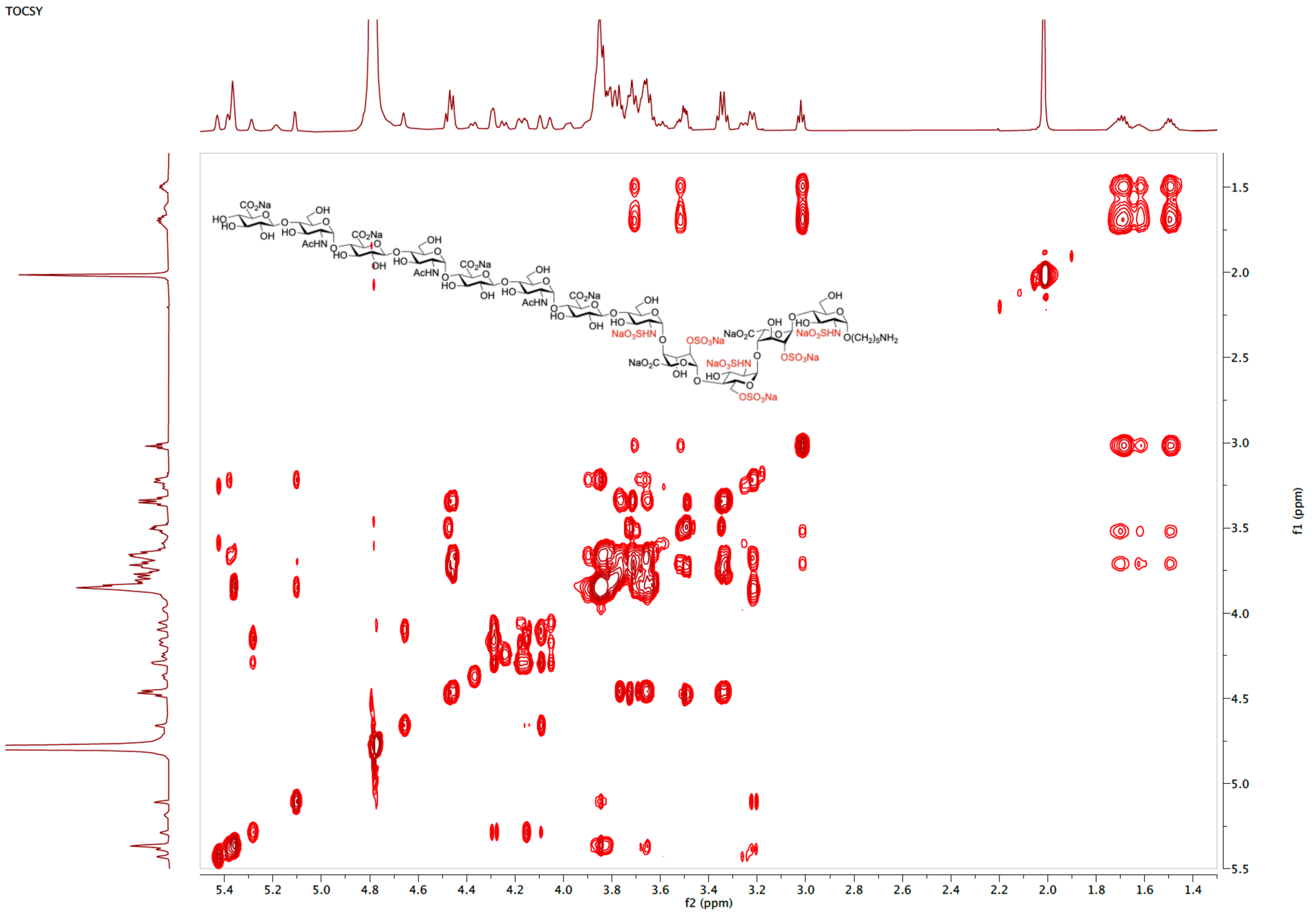

NOESY spectrum of **25** ($D_2O$)

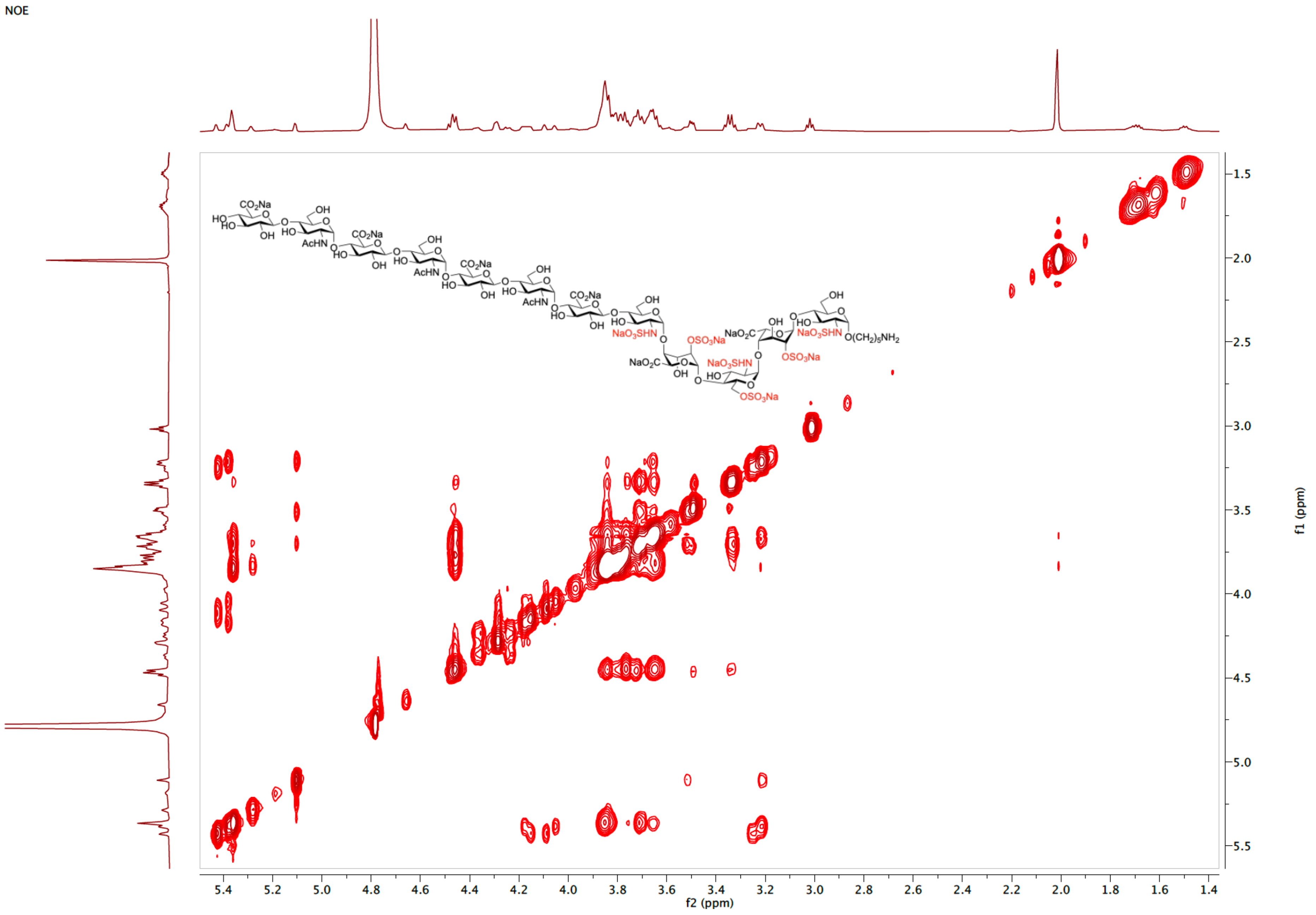

[1]H NMR spectrum of **26** (600 MHz, $D_2O$)

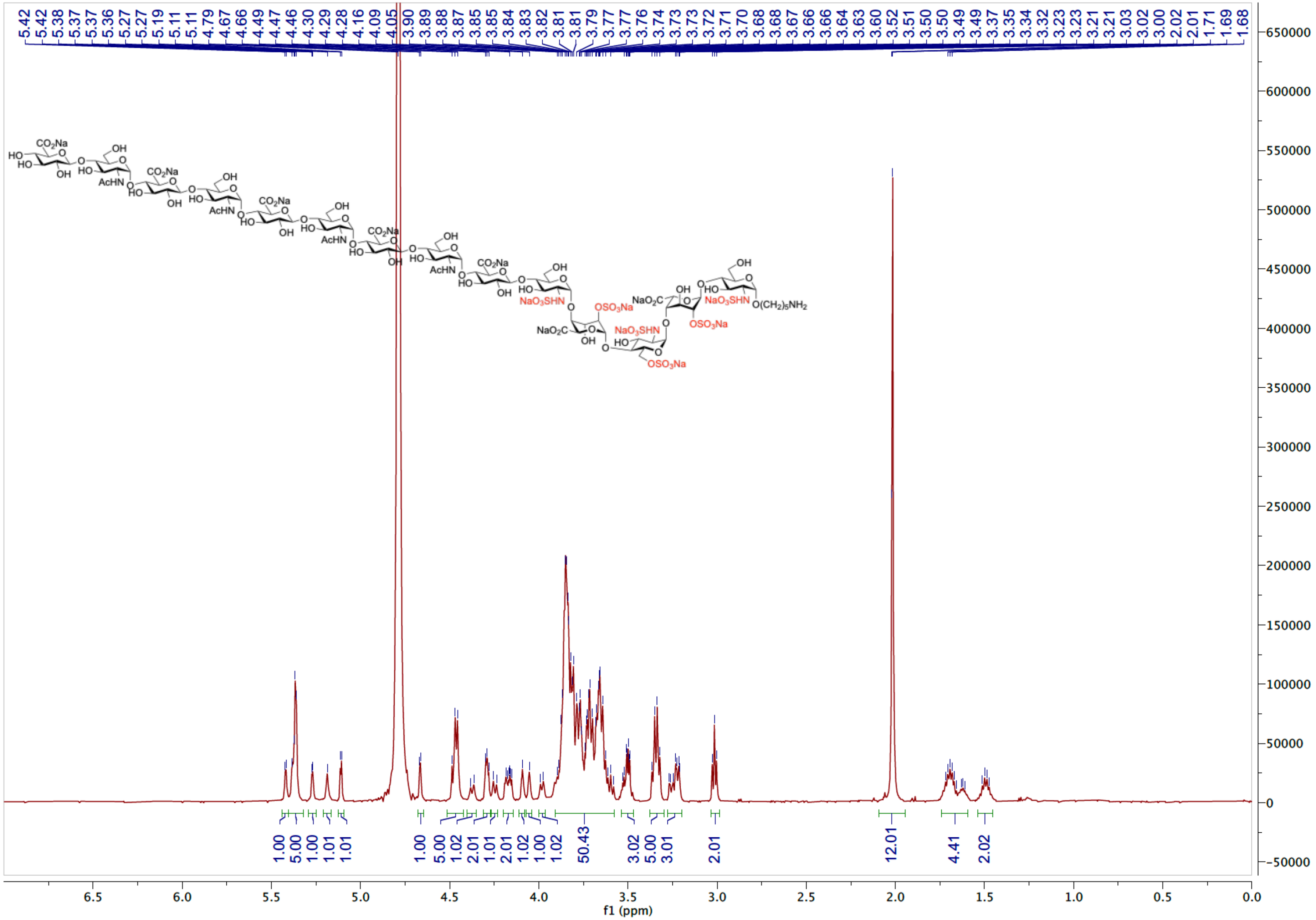

HSQC spectrum of **26** ($D_2O$)

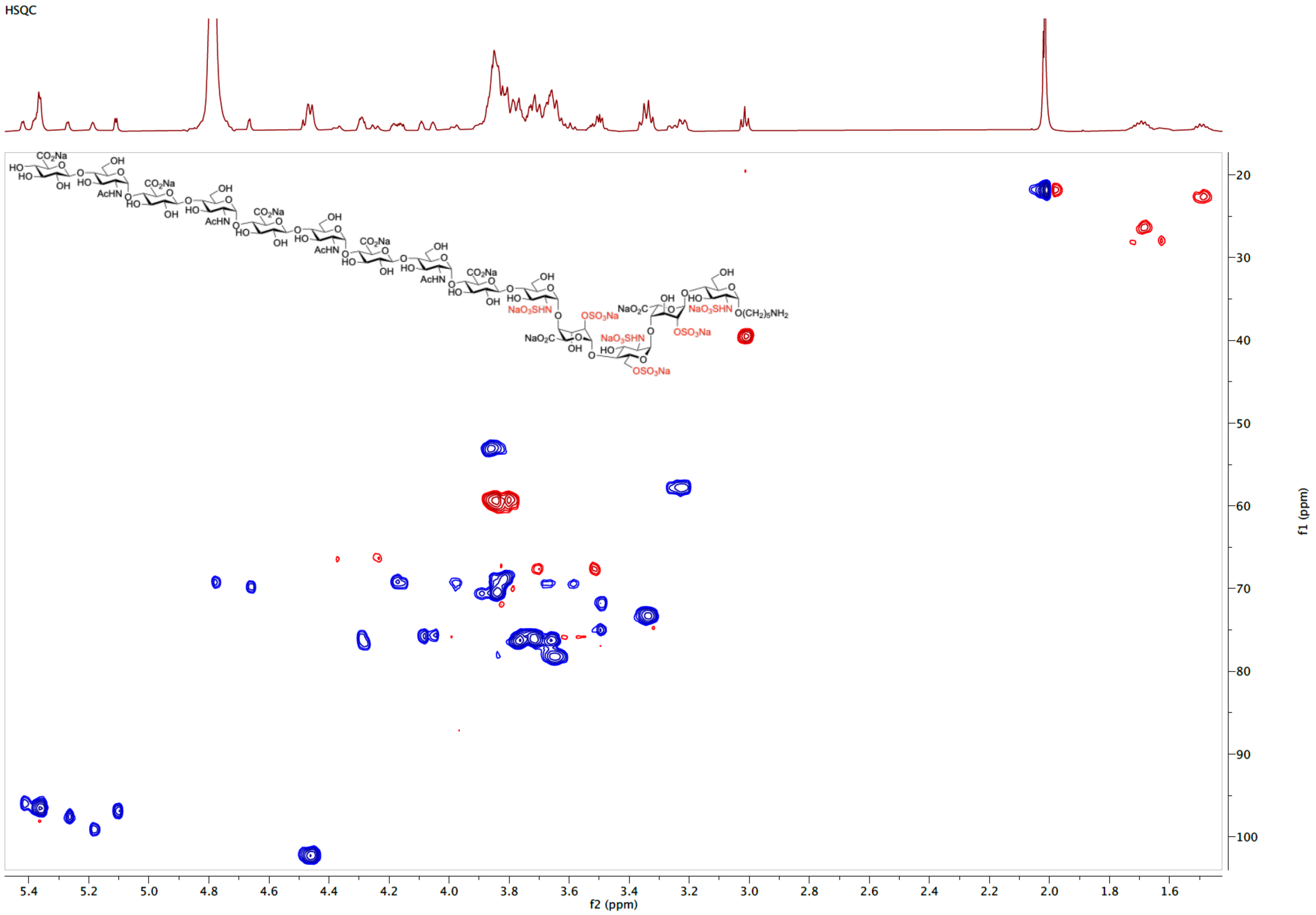

COSY spectrum of **26** ($D_2O$)

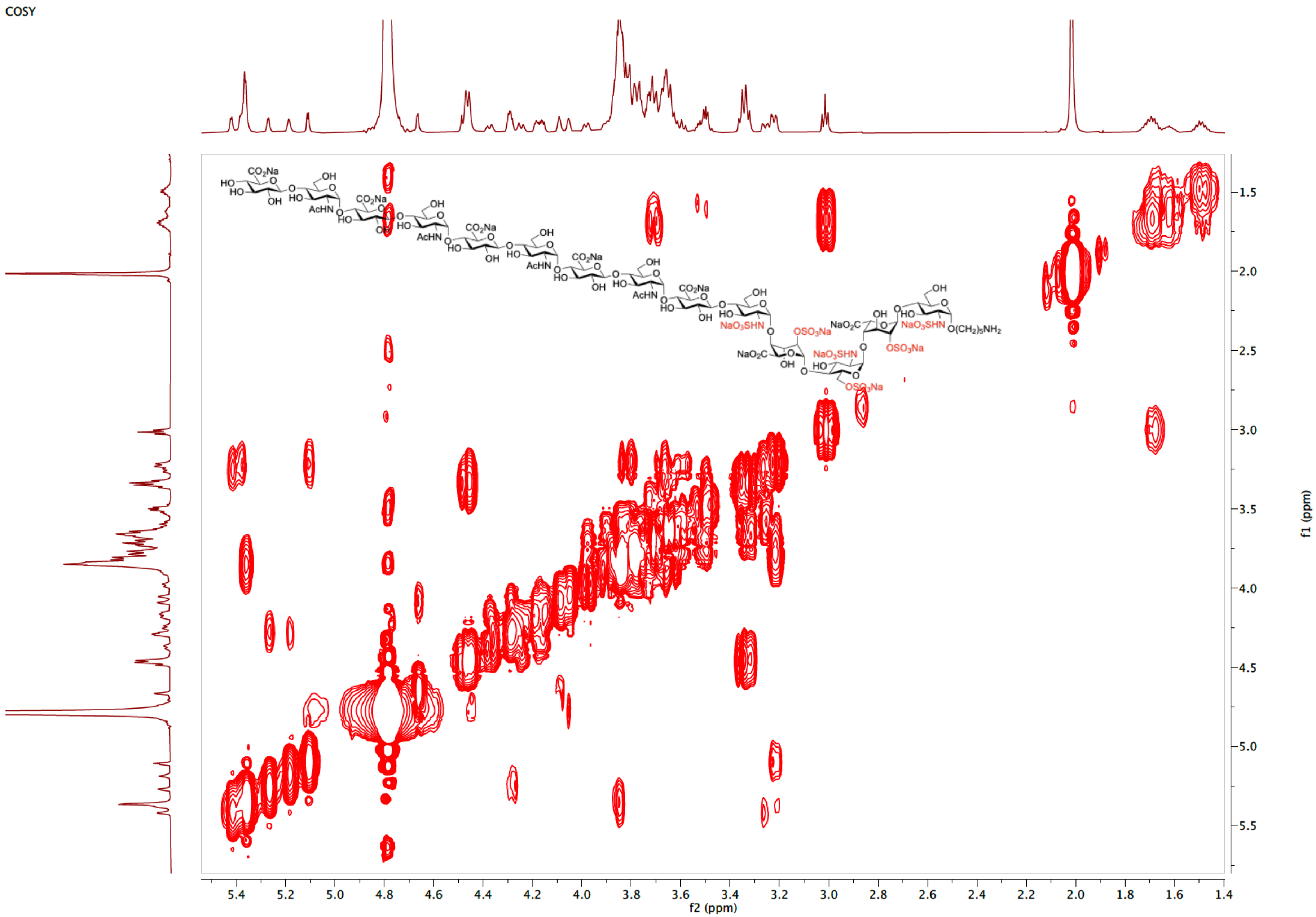

TOCSY spectrum of **26** ($D_2O$)

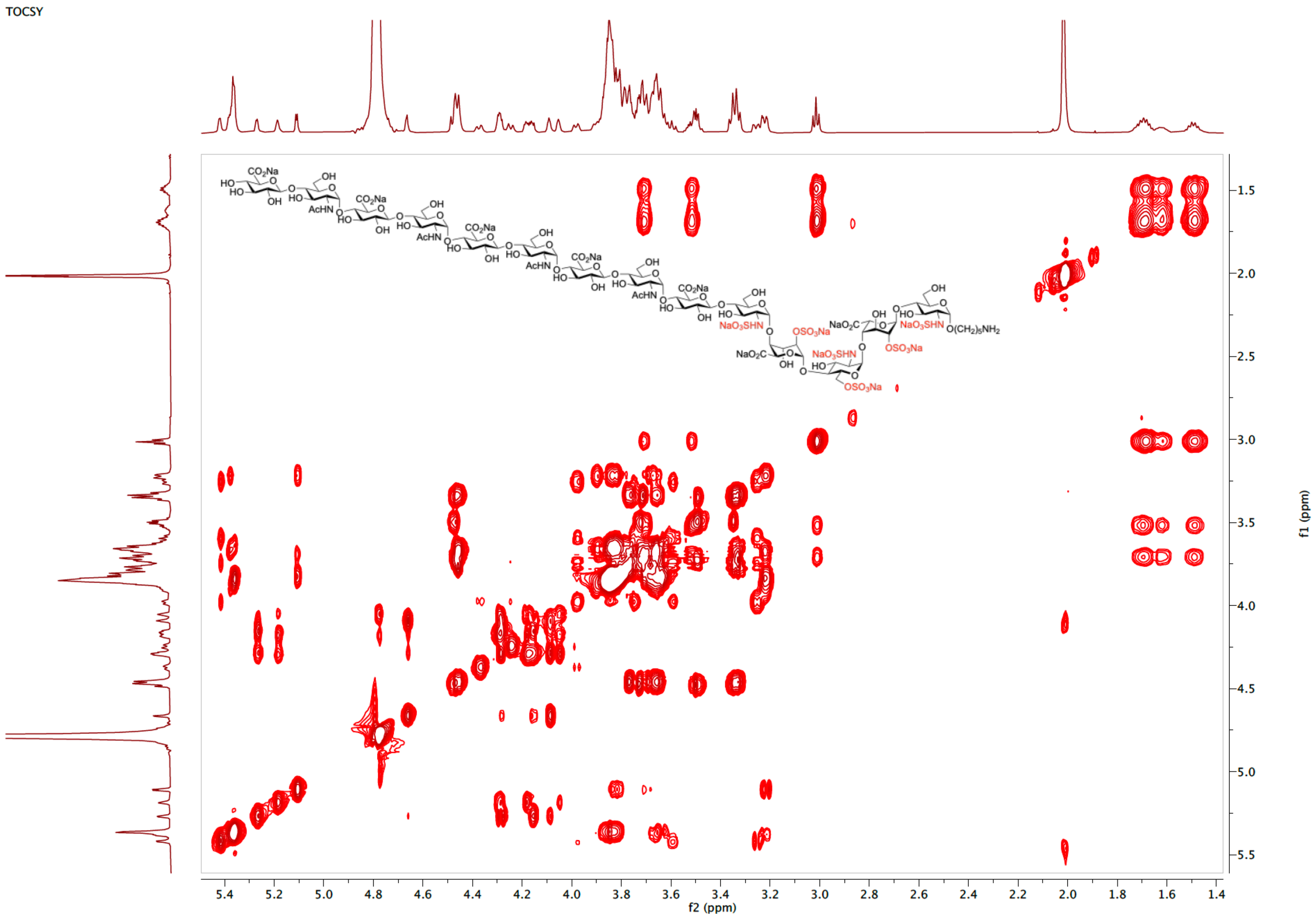

$^{1}$H NMR spectrum of **27** (600 MHz, $D_2O$)

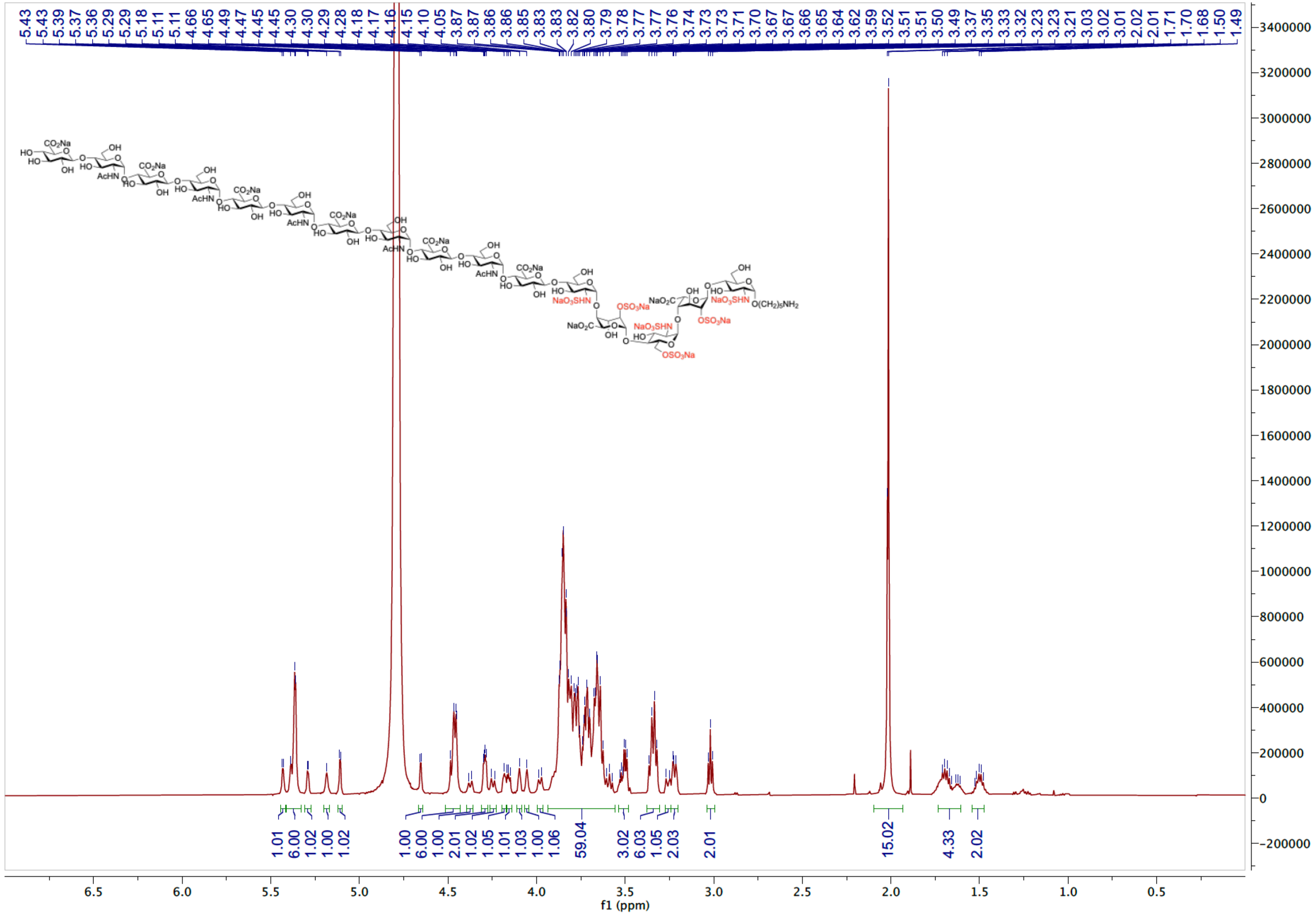

HSQC spectrum of **27** ($D_2O$)

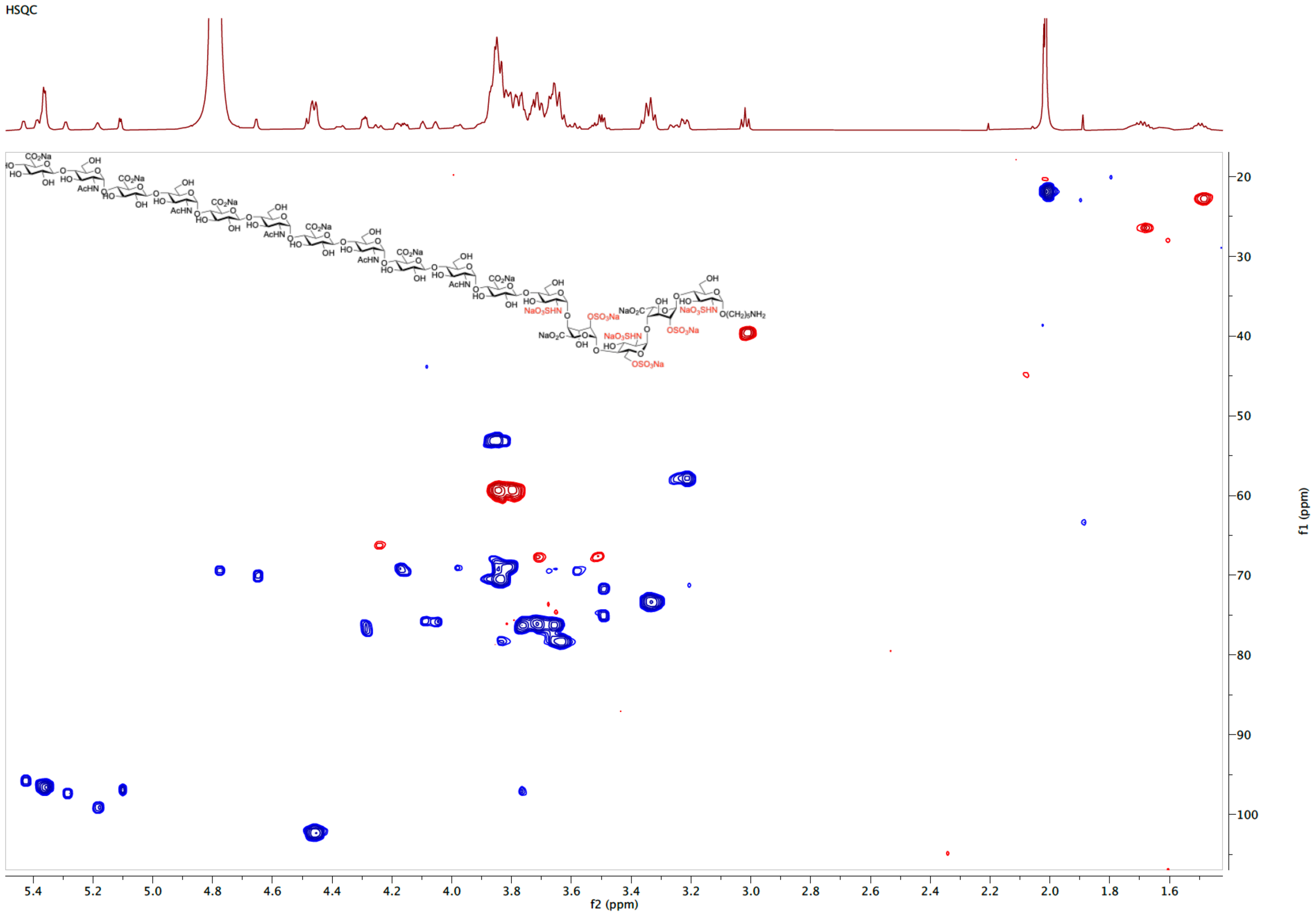

COSY spectrum of **27** ($D_2O$)

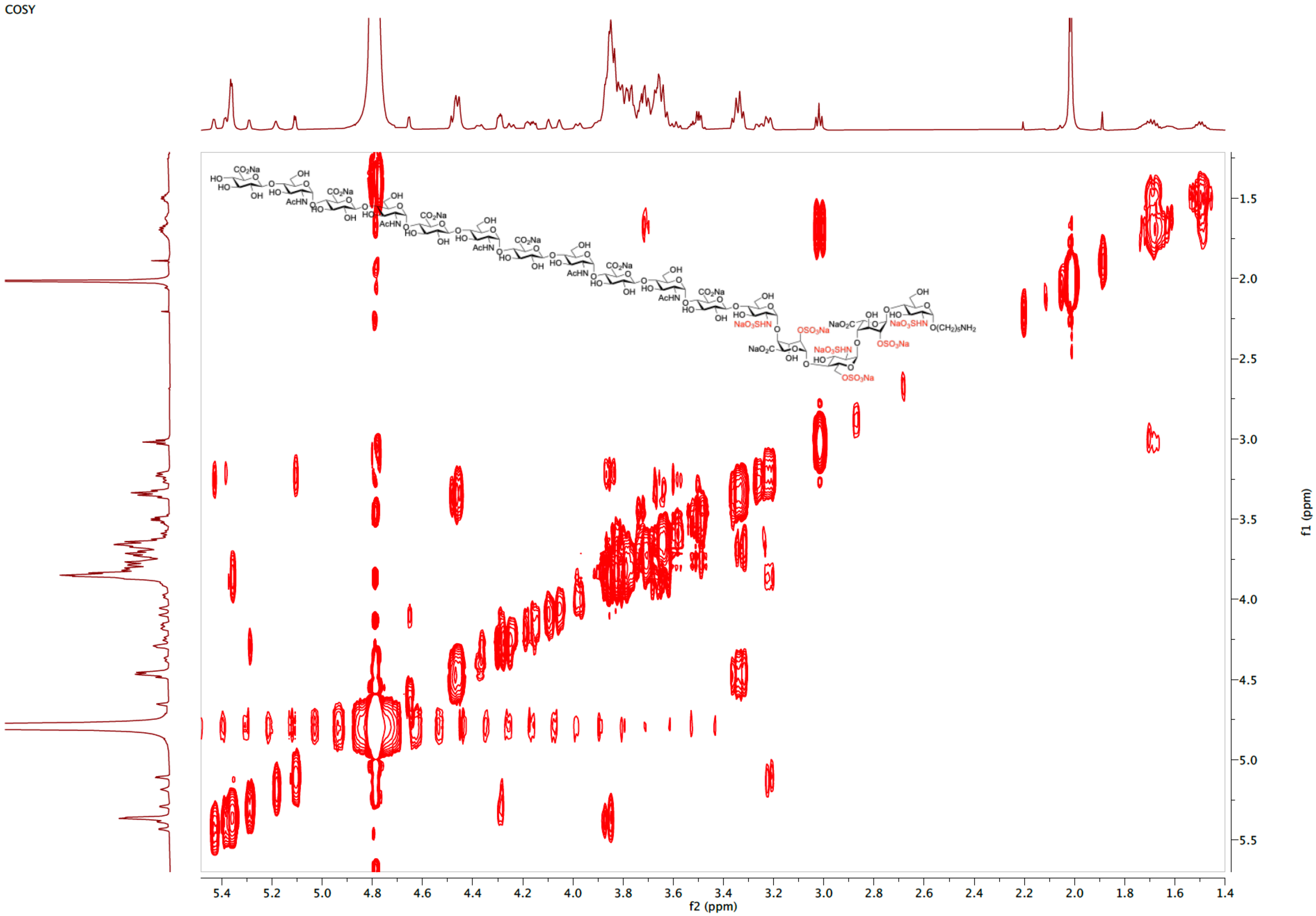

TOCSY spectrum of **27** ($D_2O$)

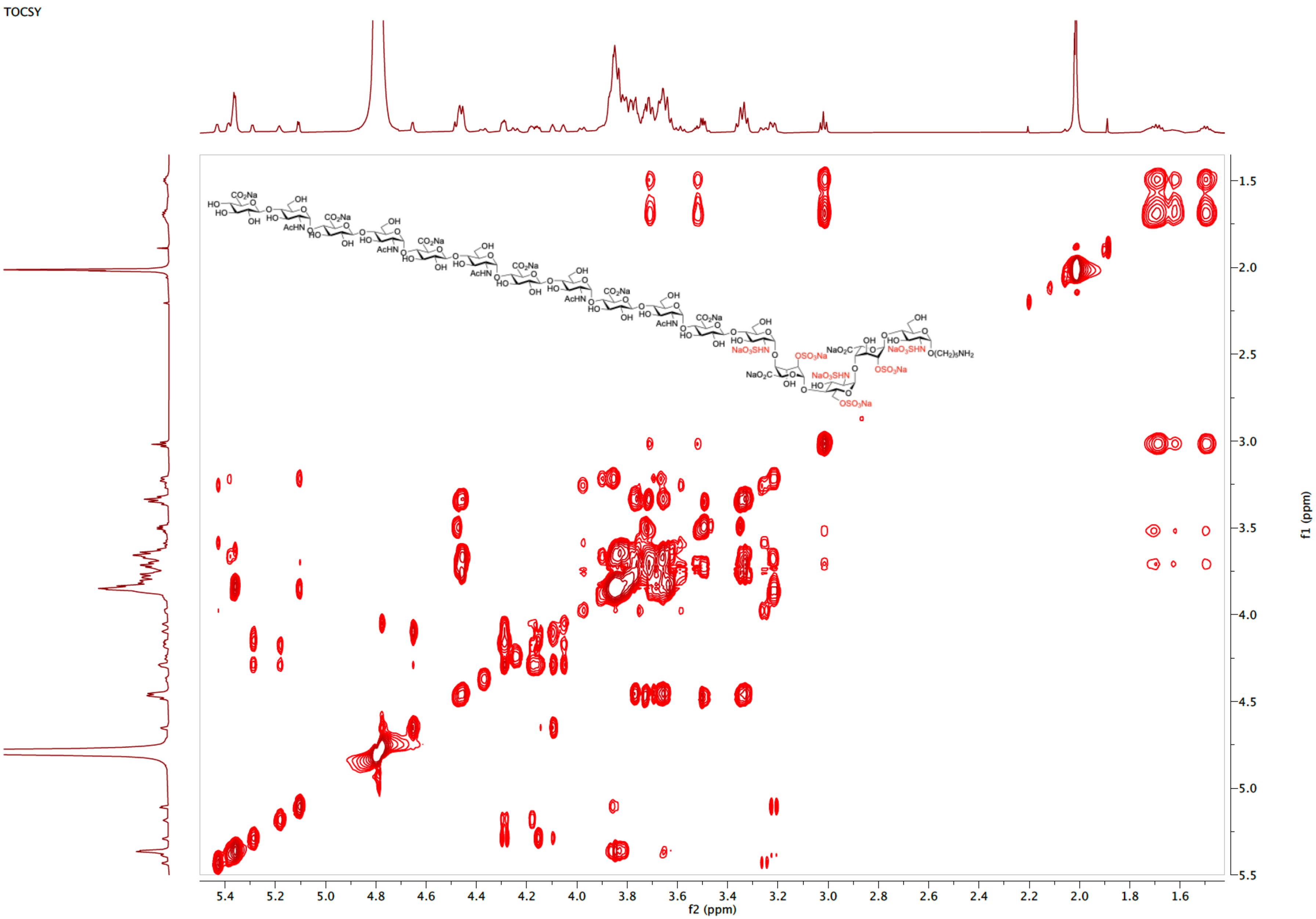

[1]H NMR spectrum of **3** (600 MHz, $D_2O$)

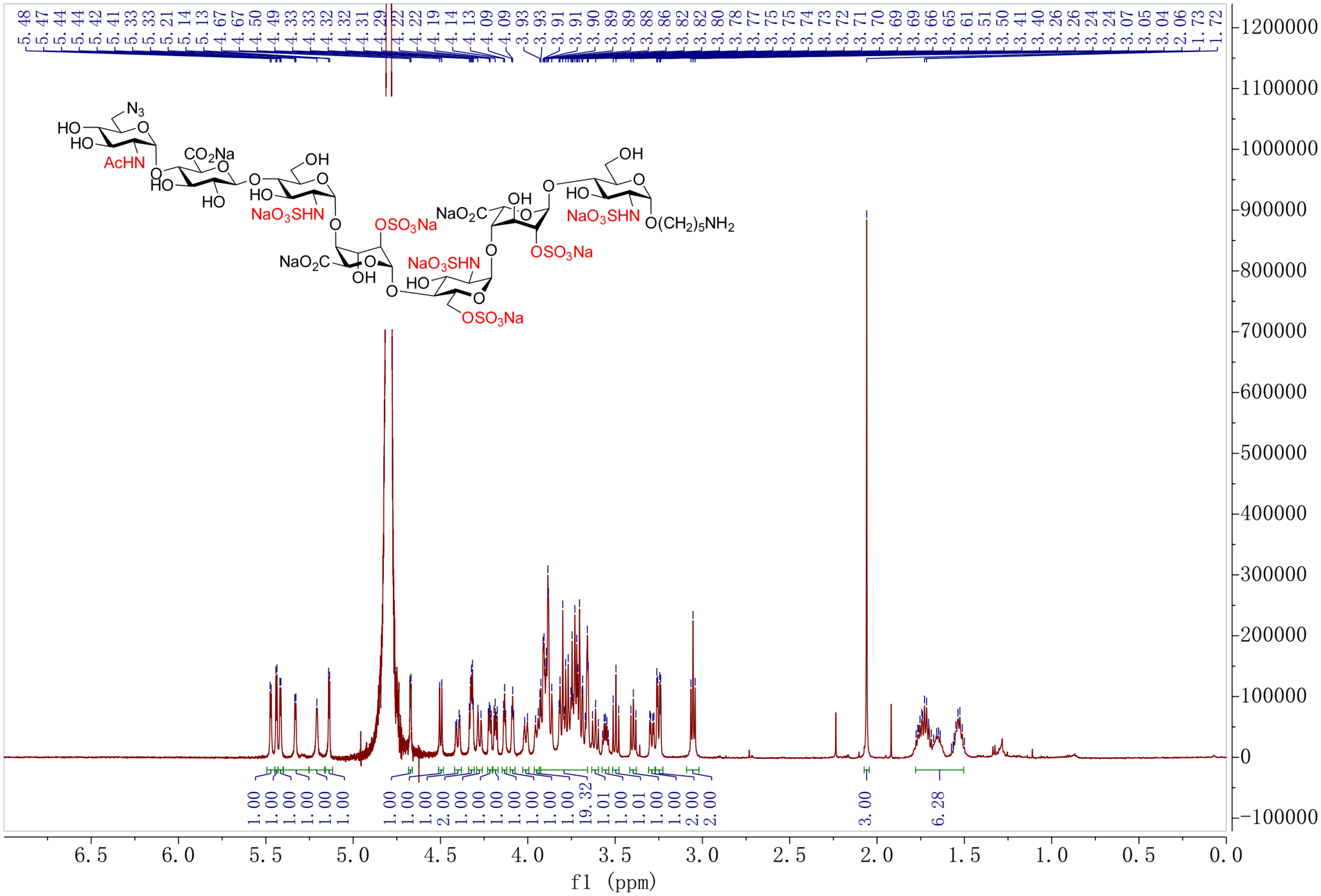

HSQC spectrum of **3** ($D_2O$)

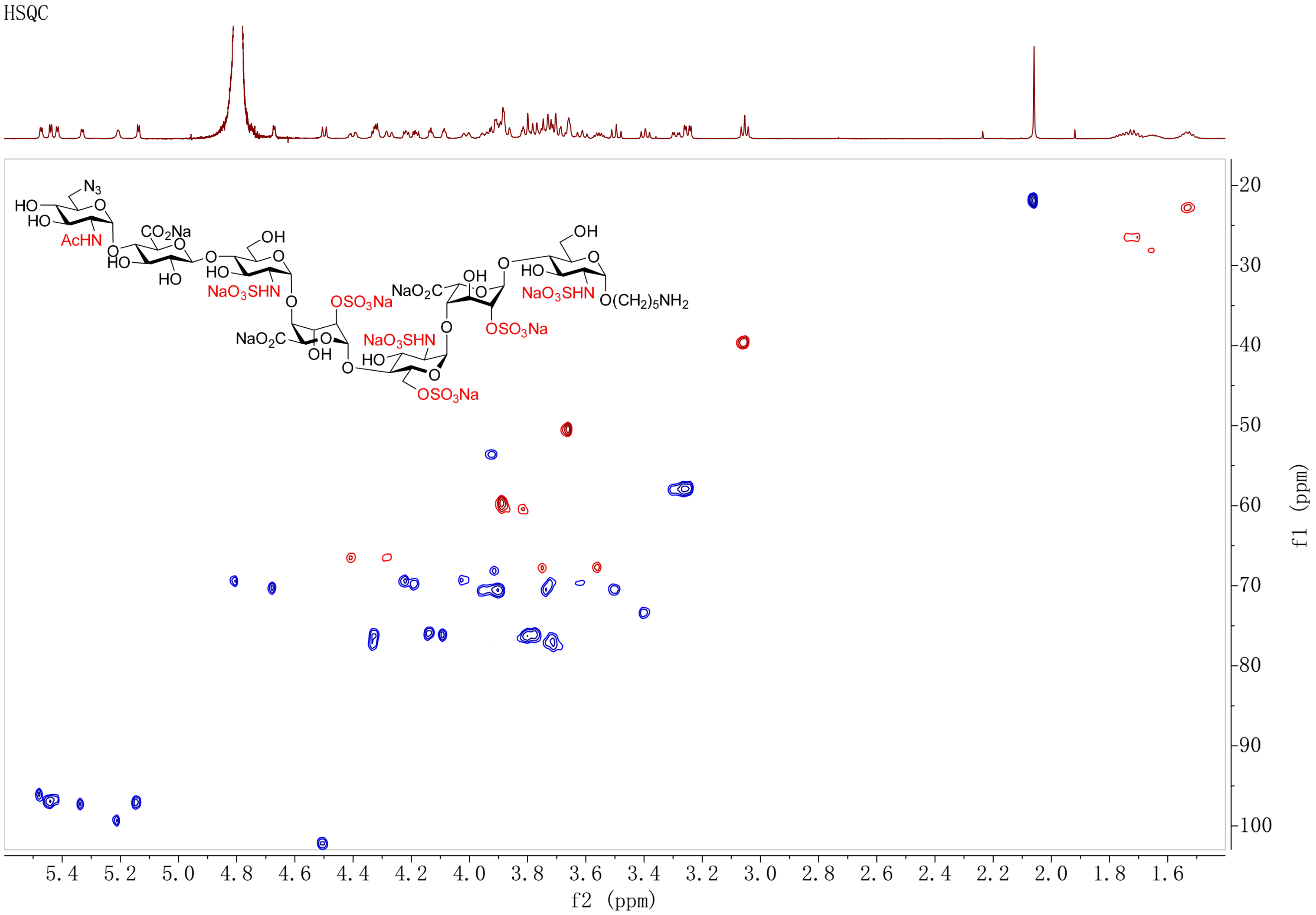

COSY spectrum of **3** ($D_2O$)

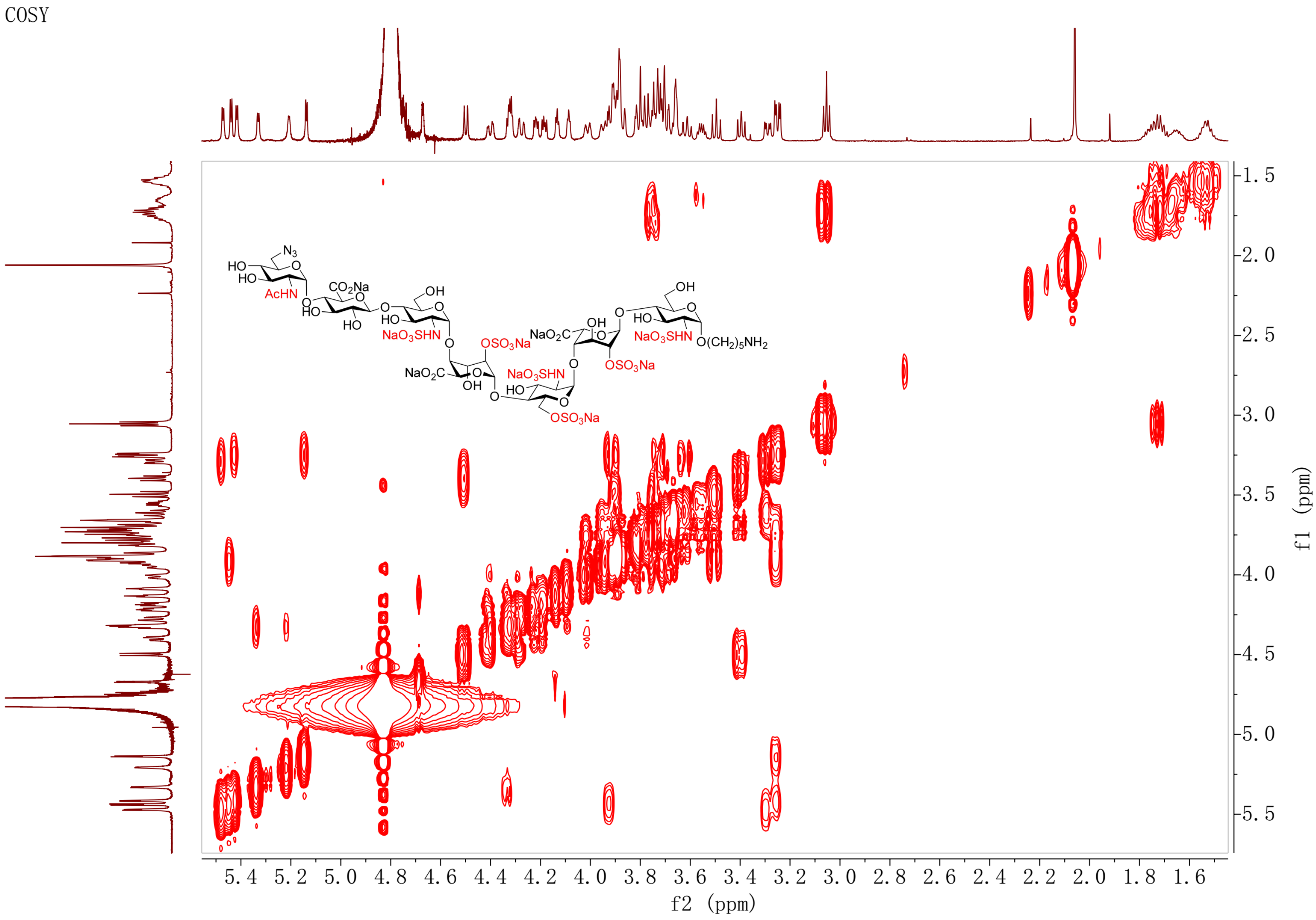

TOCSY spectrum of **3** ($D_2O$)

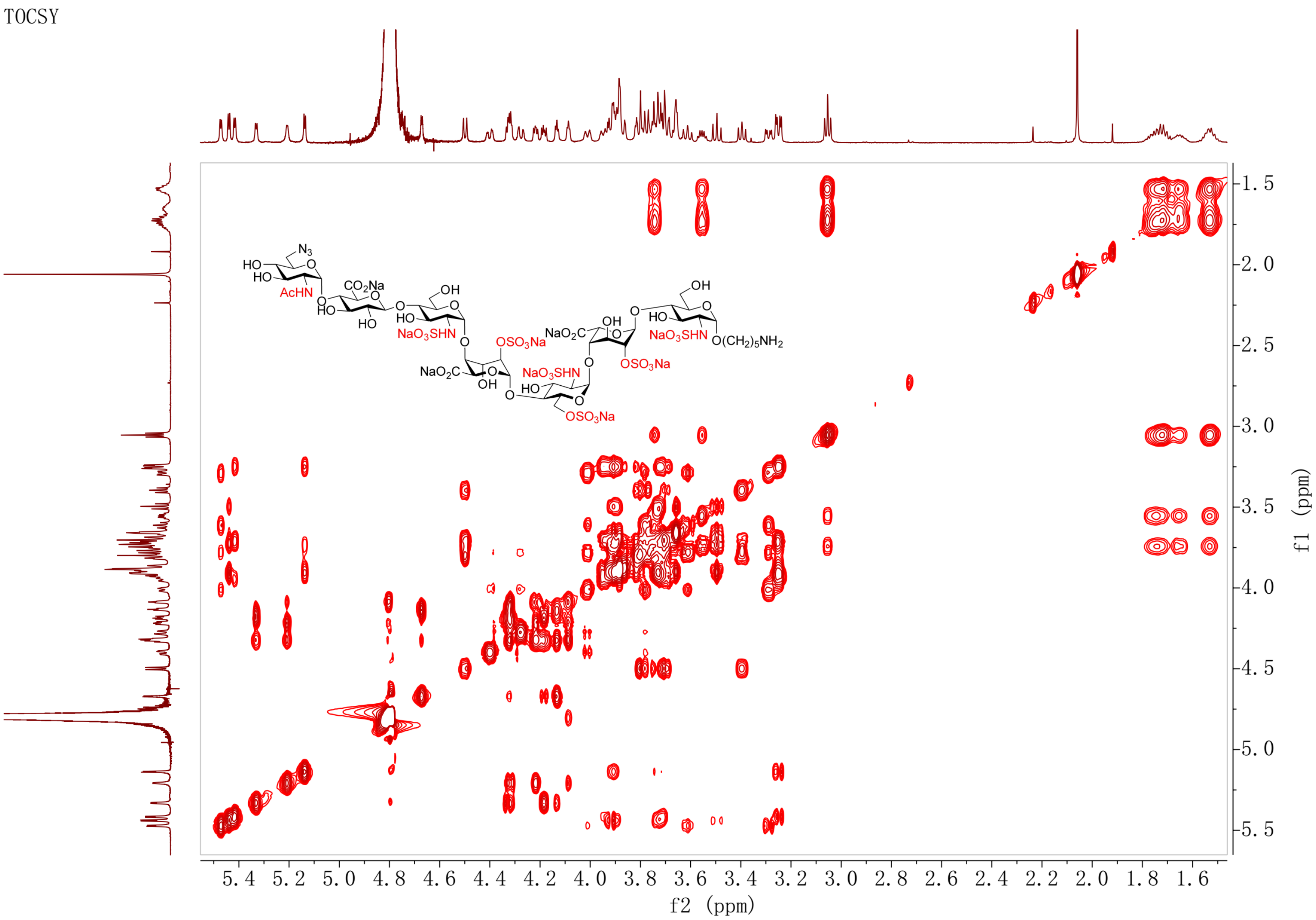

NOESY spectrum of **3** ($D_2O$)

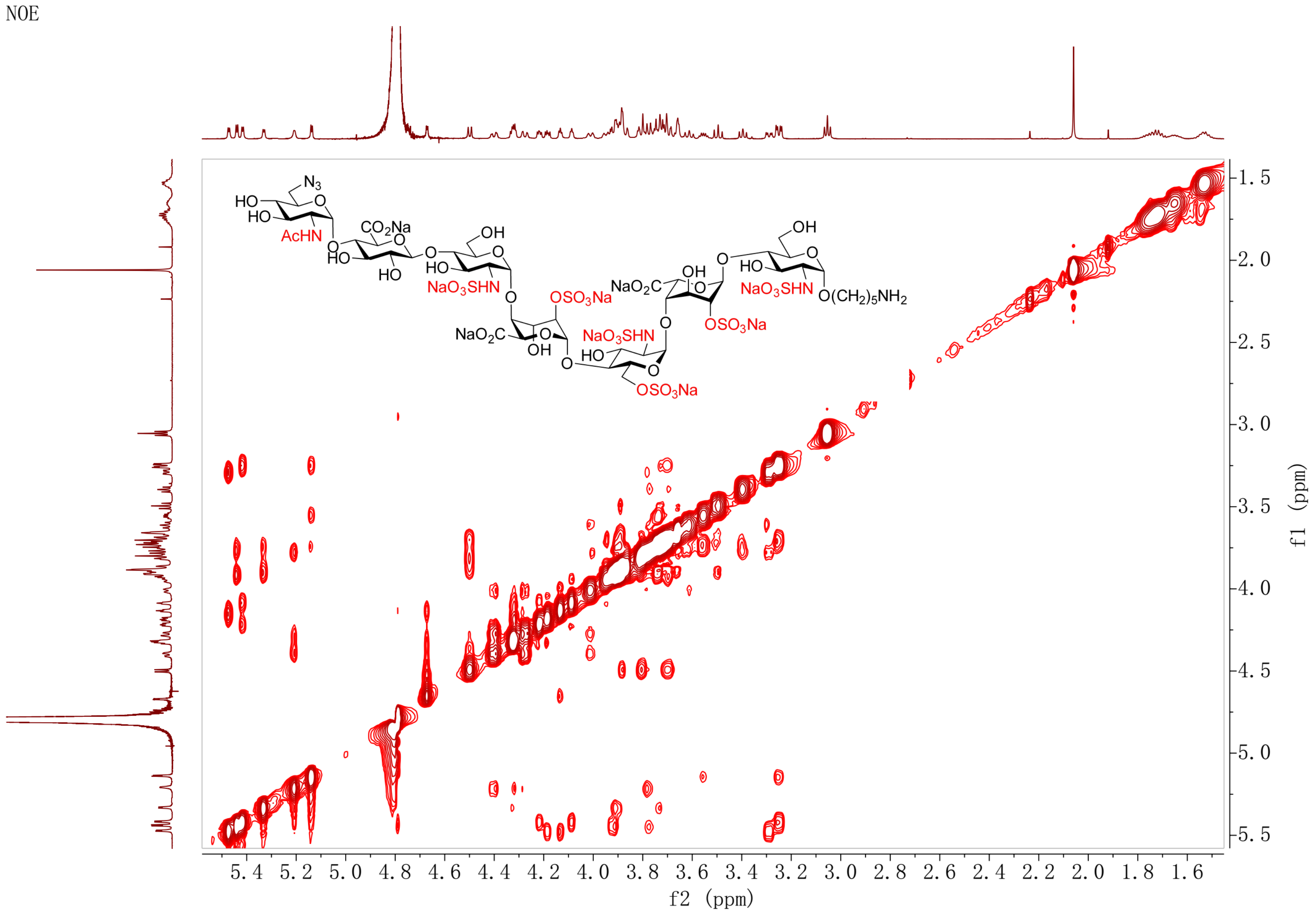

[1]H NMR spectrum of **4** (600 MHz, $D_2O$)

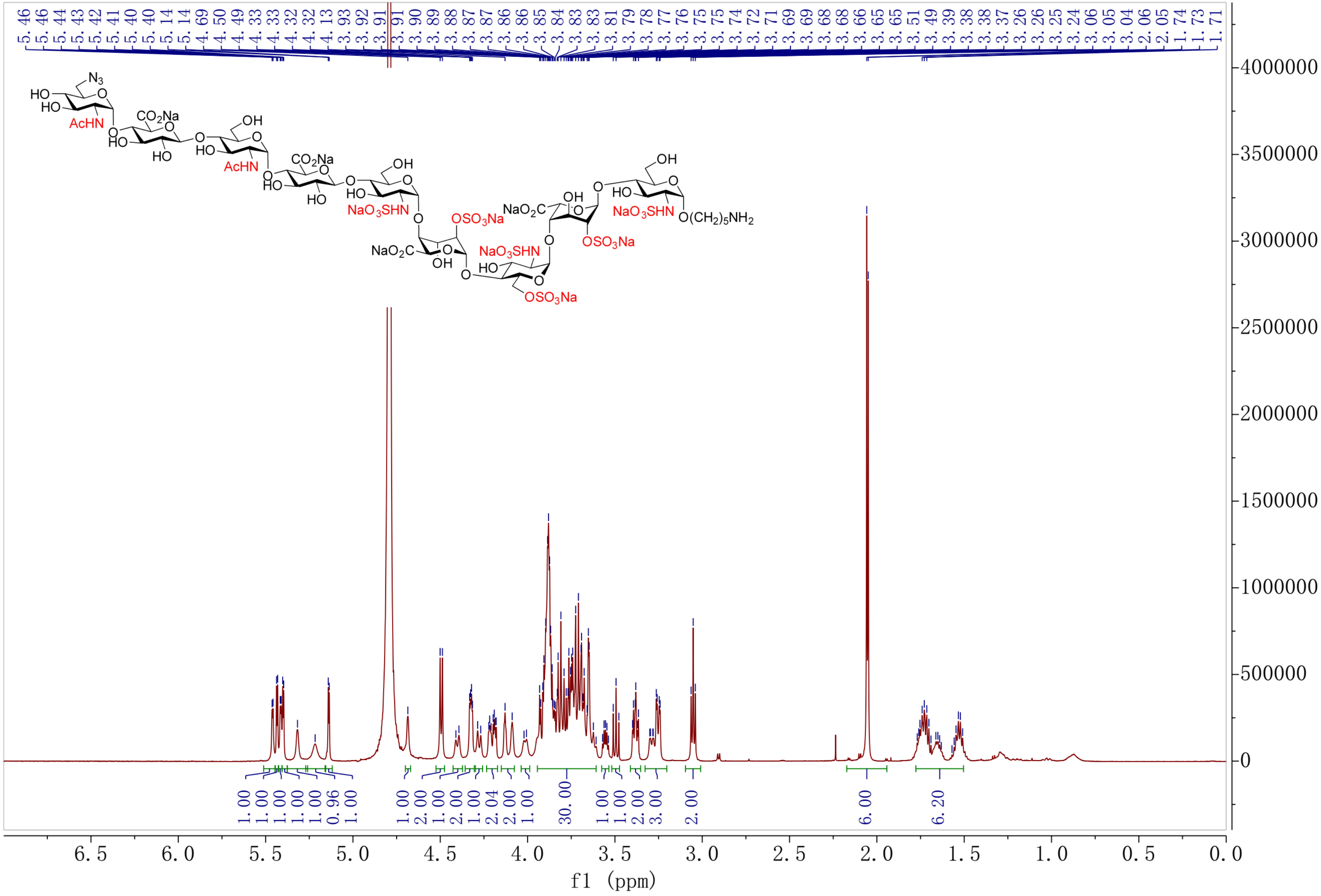

HSQC spectrum of **4** ($D_2O$)

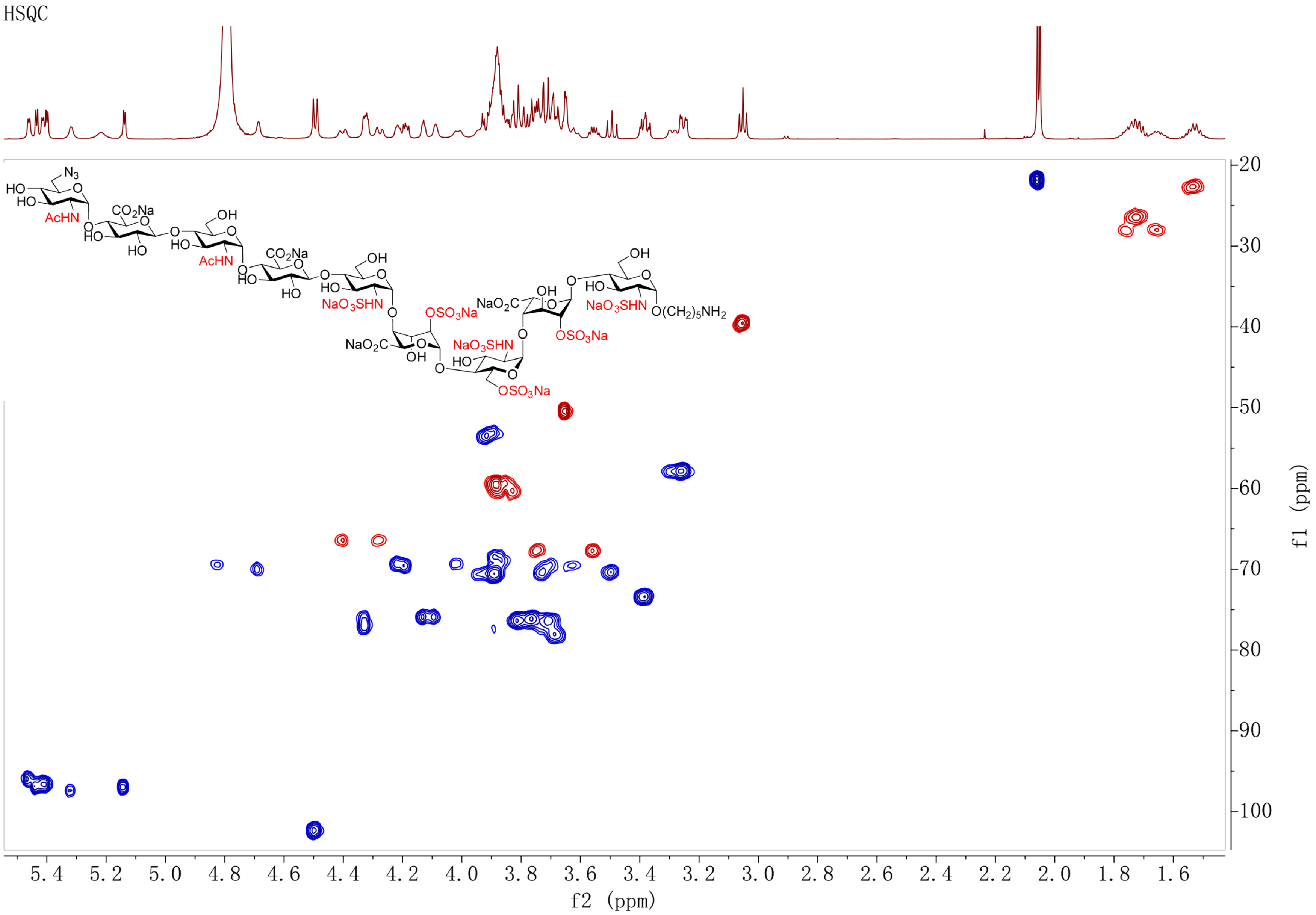

COSY spectrum of **4** ($D_2O$)

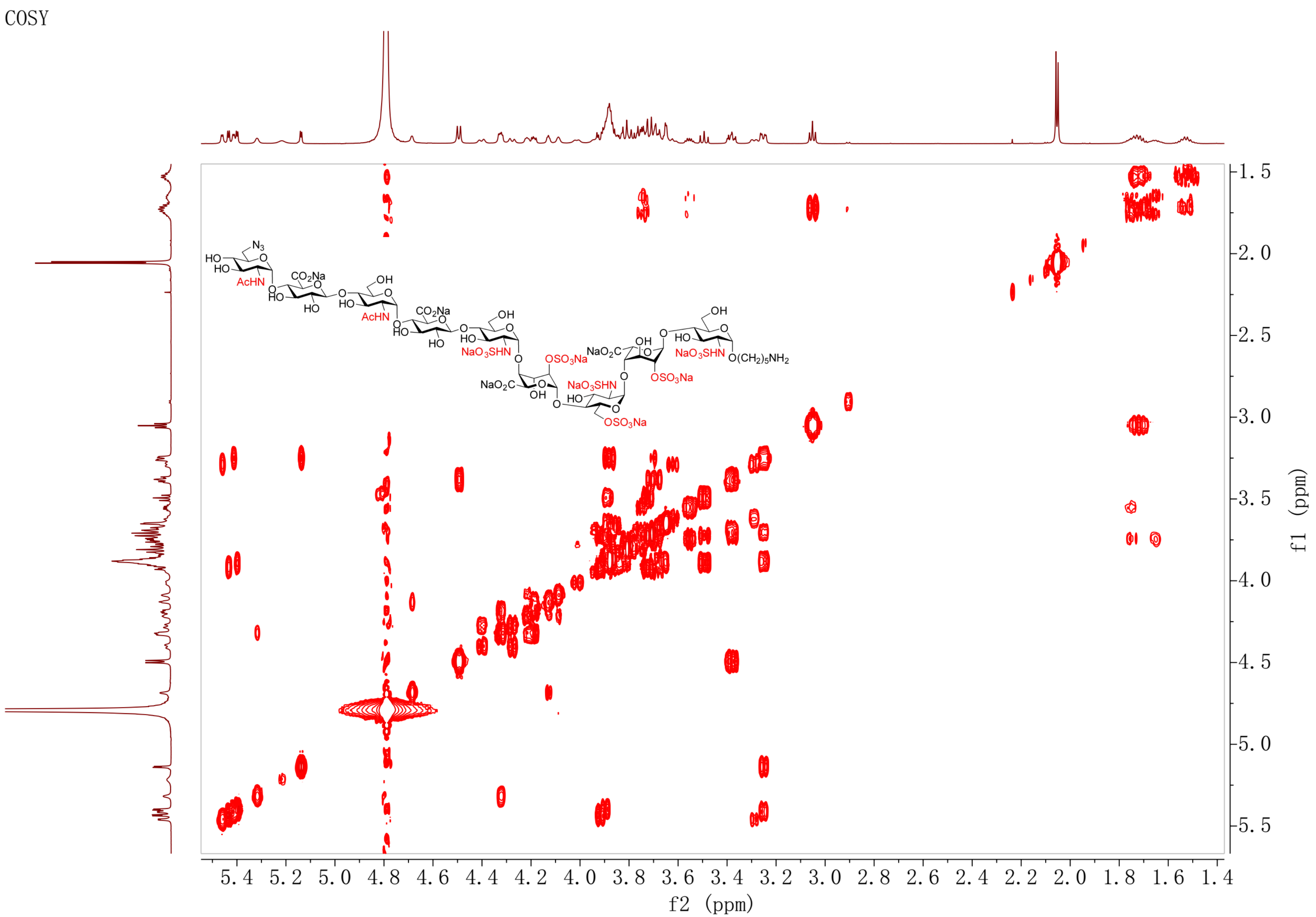

TOCSY spectrum of **4** ($D_2O$)

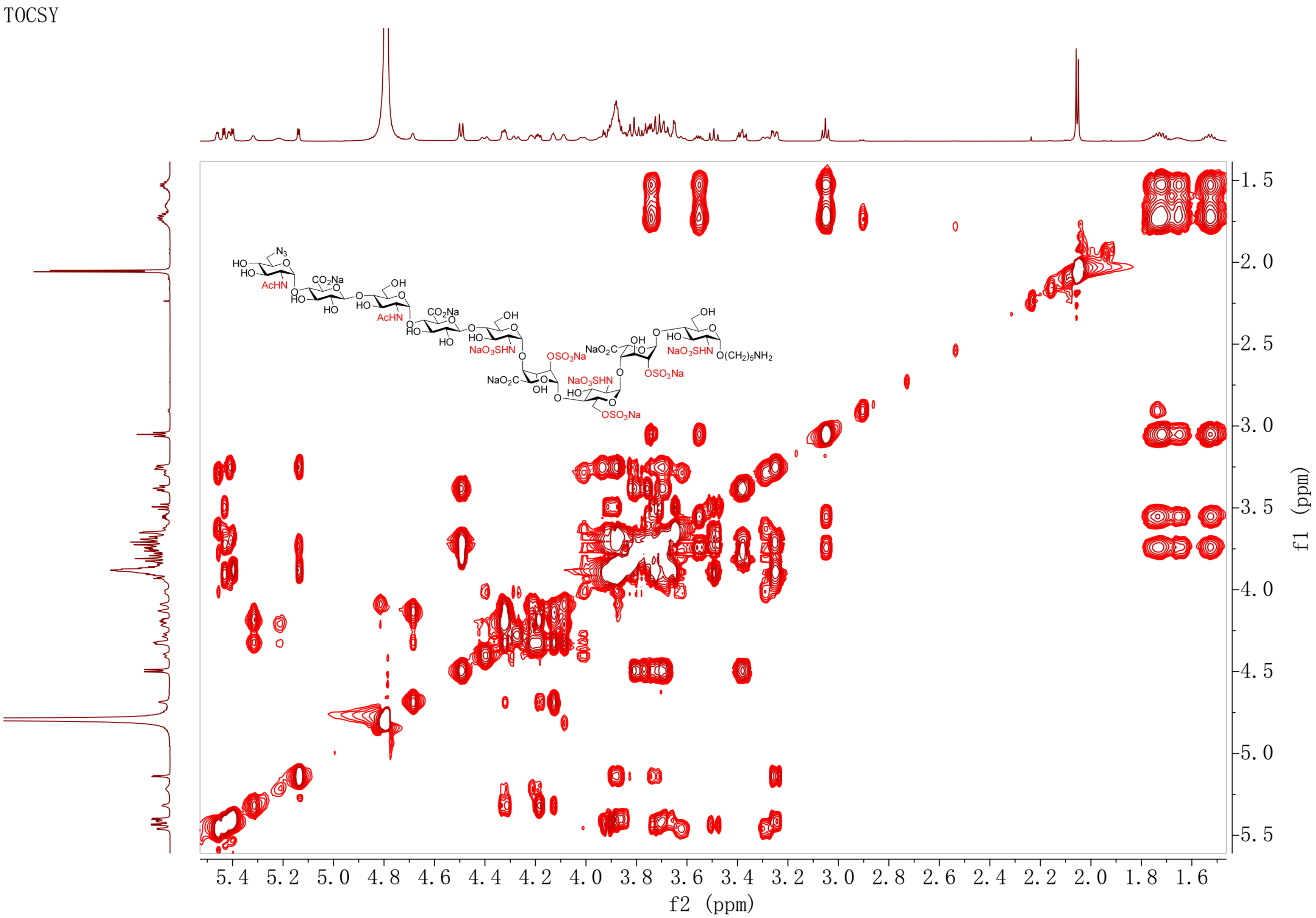

NOESY spectrum of **4** ($D_2O$)

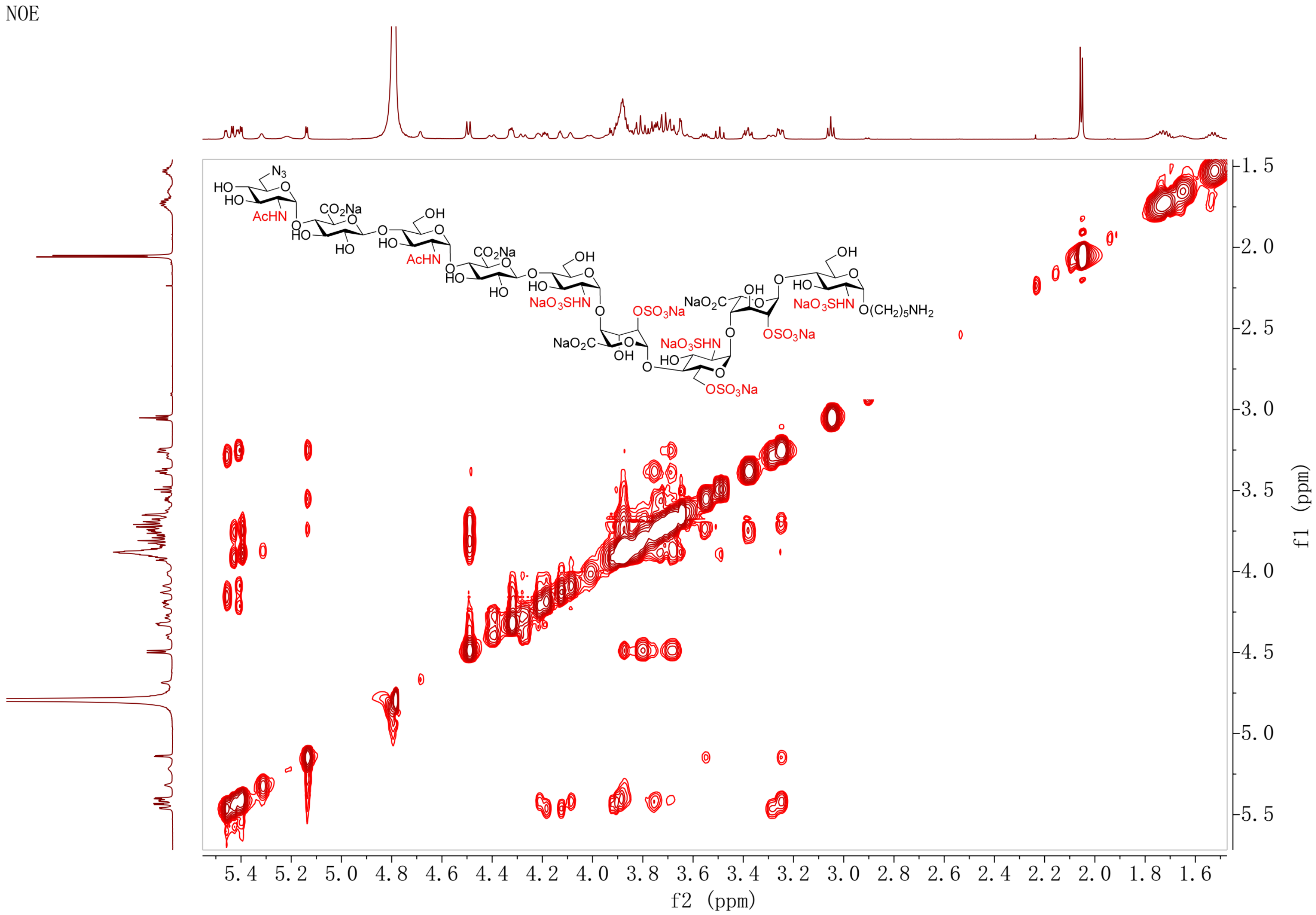

[1]H NMR spectrum of **5** (600 MHz, $D_2O$)

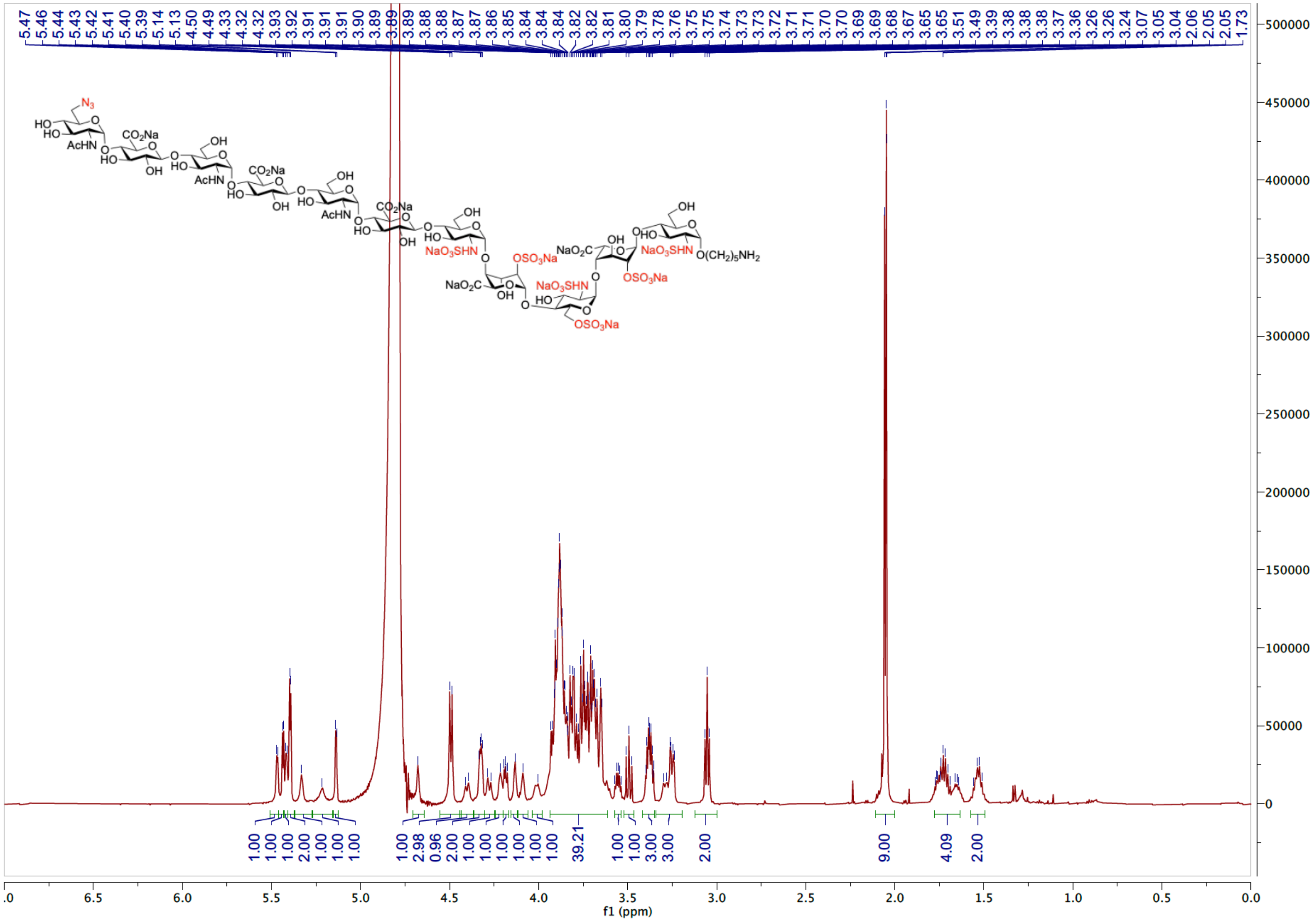

HSQC spectrum of **5** ($D_2O$)

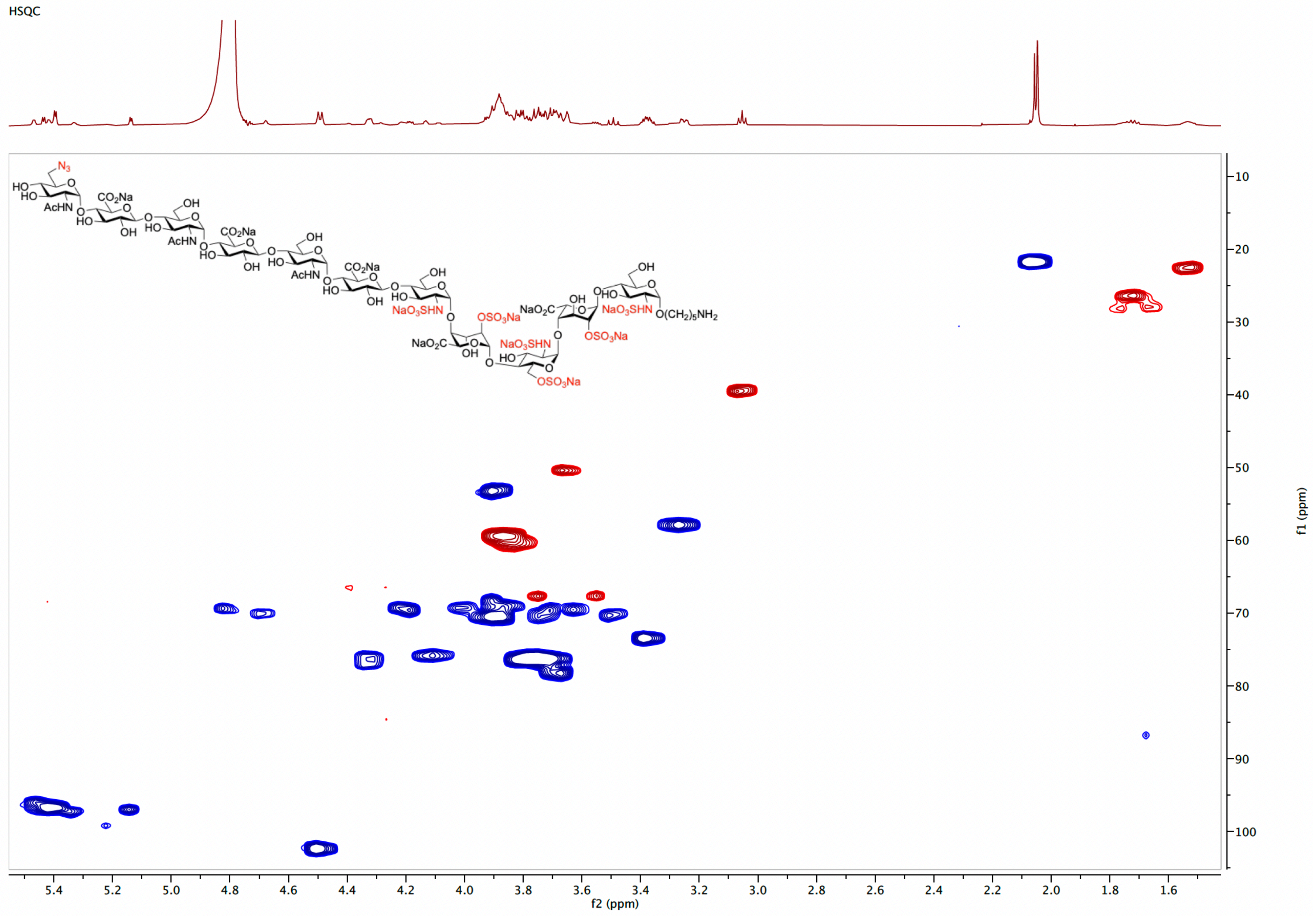

TOCSY spectrum of **5** ($D_2O$)

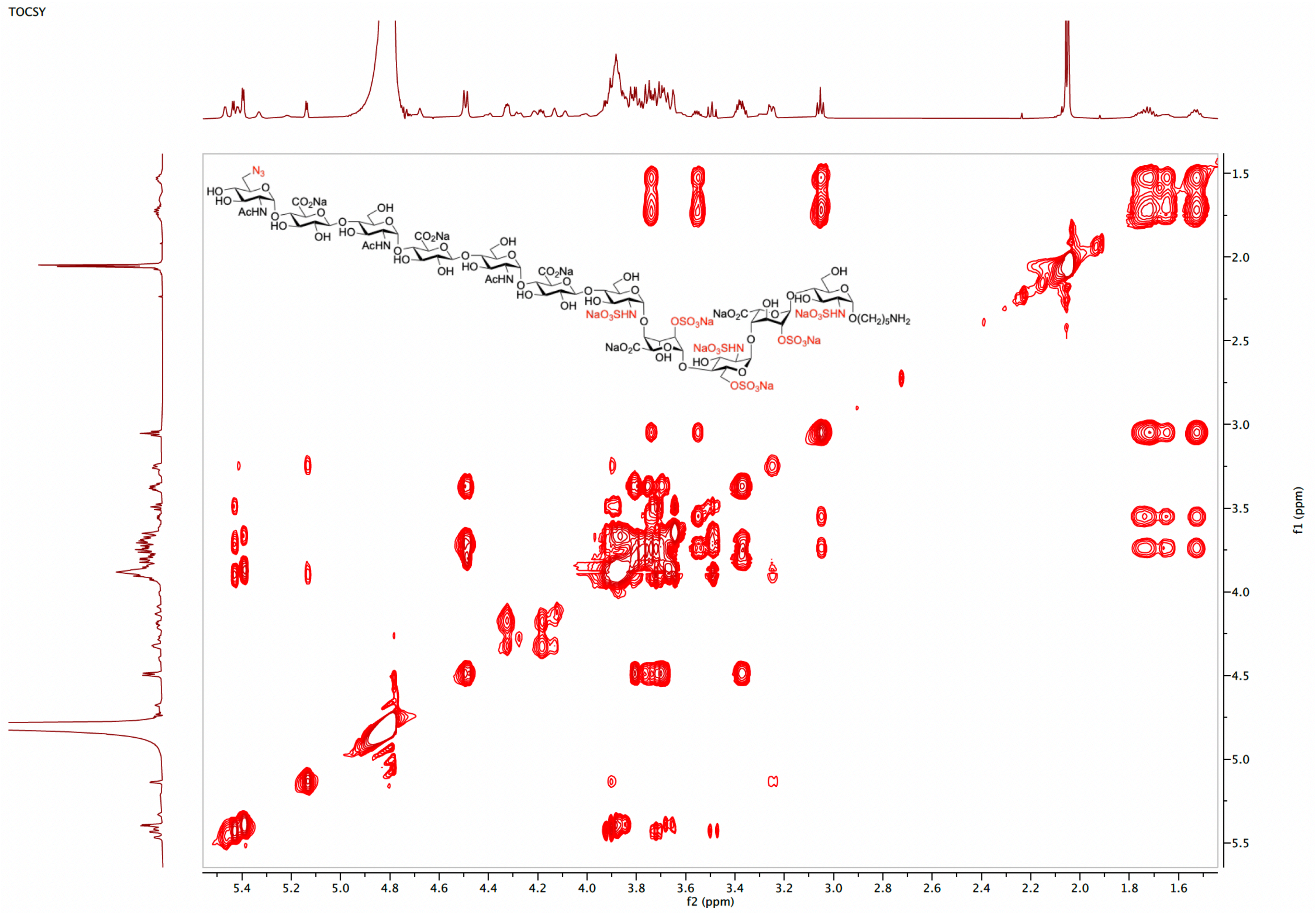

[1]H NMR spectrum of **6** (600 MHz, $D_2O$)

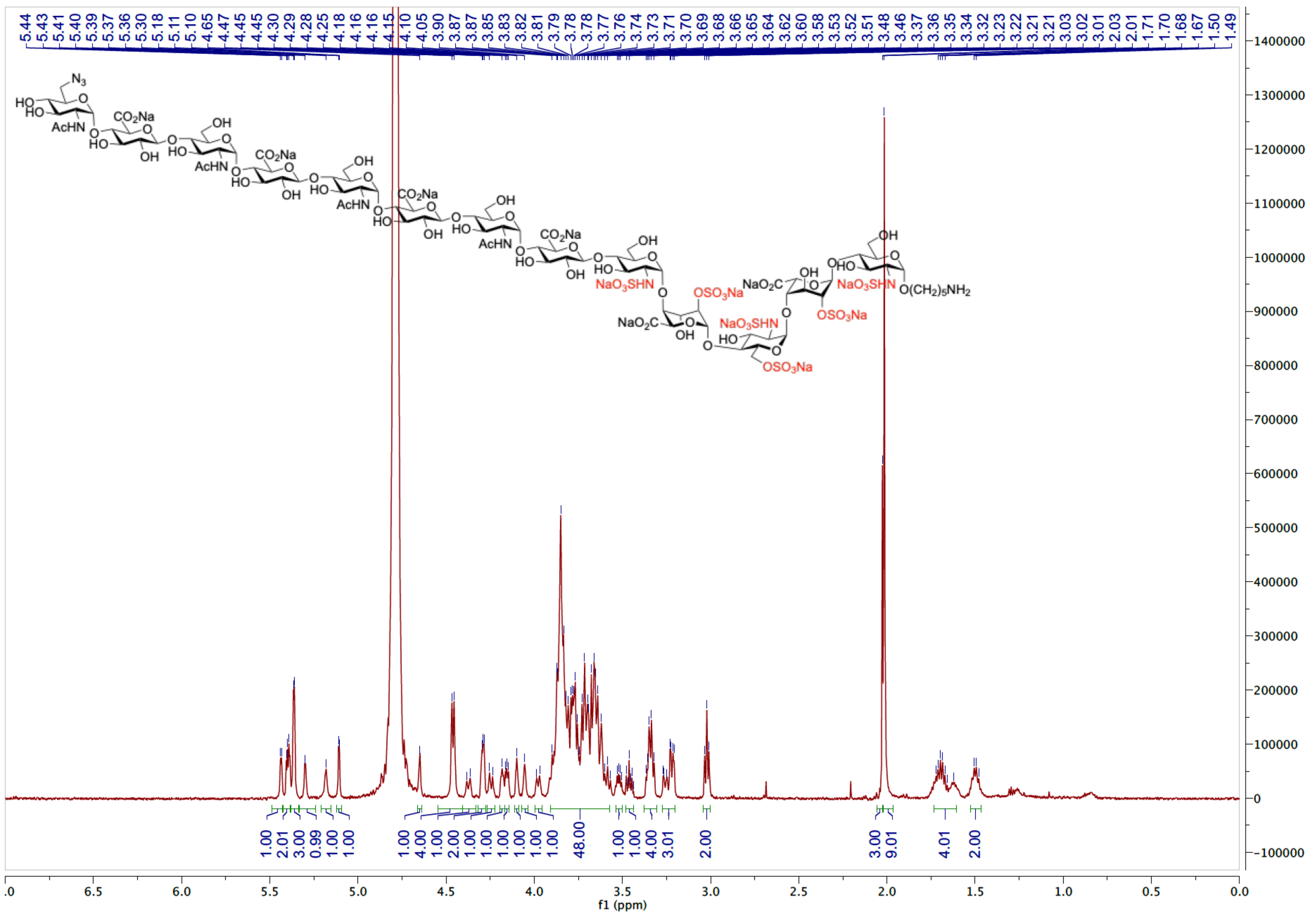

HSQC spectrum of **6** ($D_2O$)

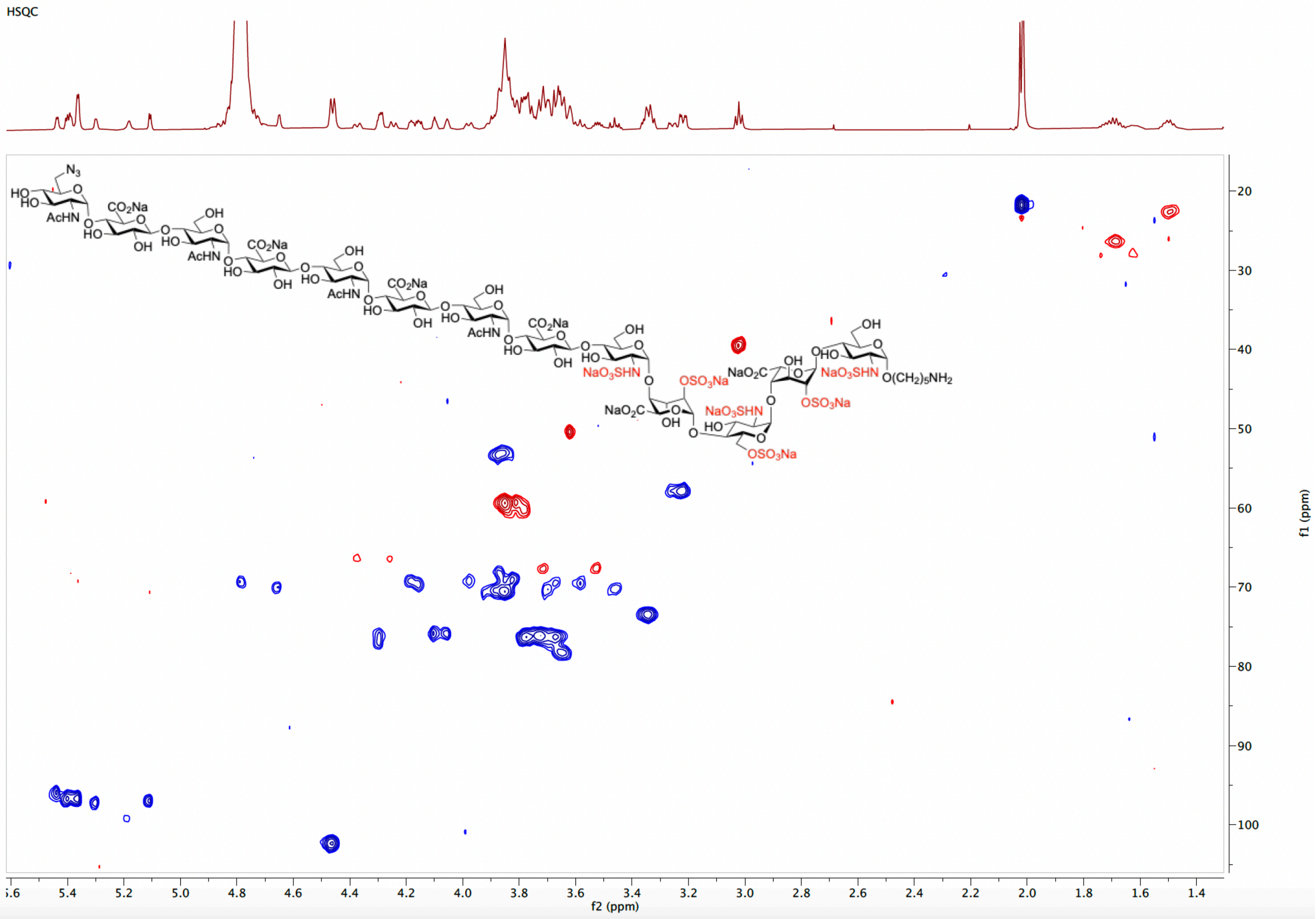

COSY spectrum of **6** ($D_2O$)

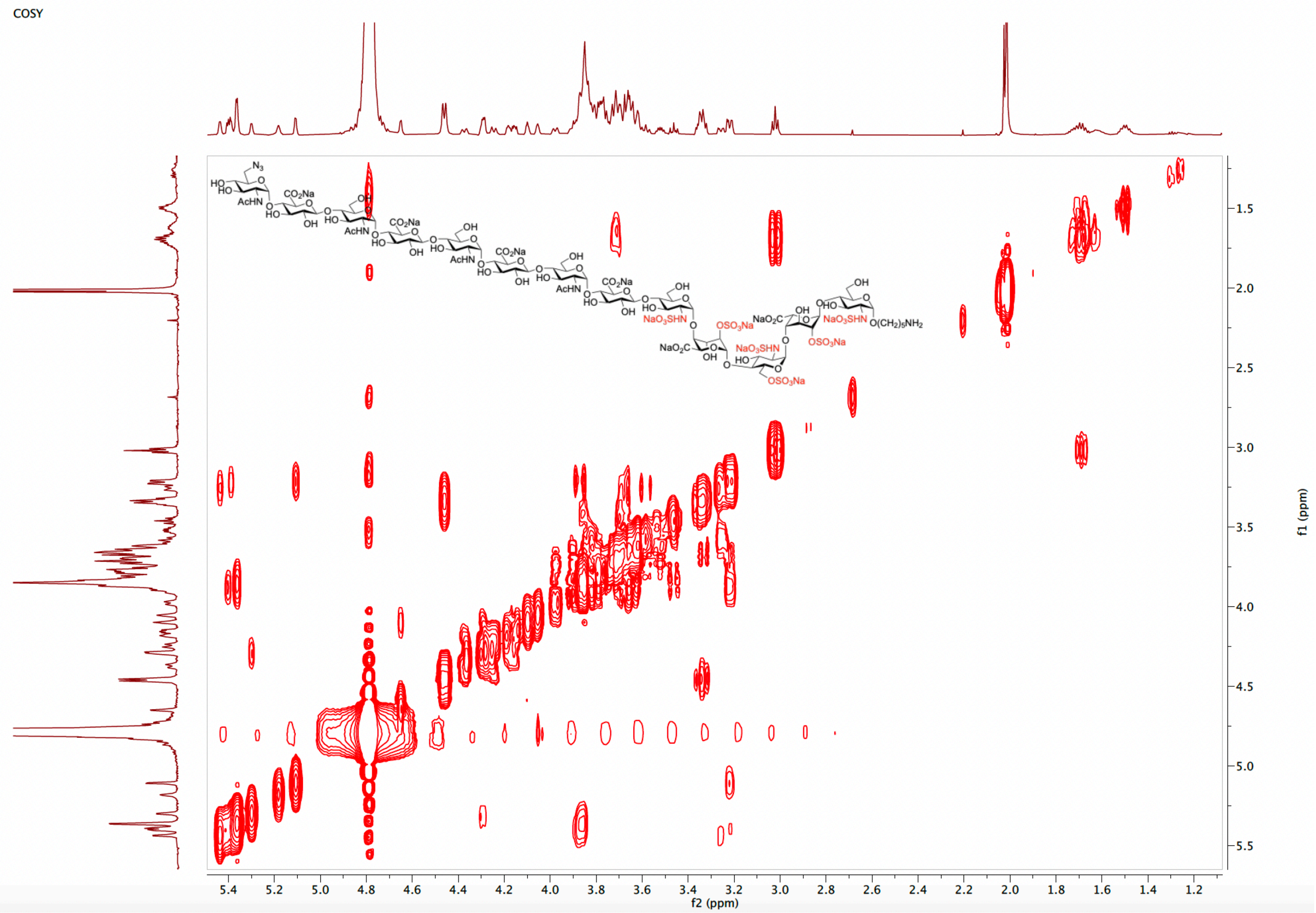

TOCSY spectrum of **6** ($D_2O$)

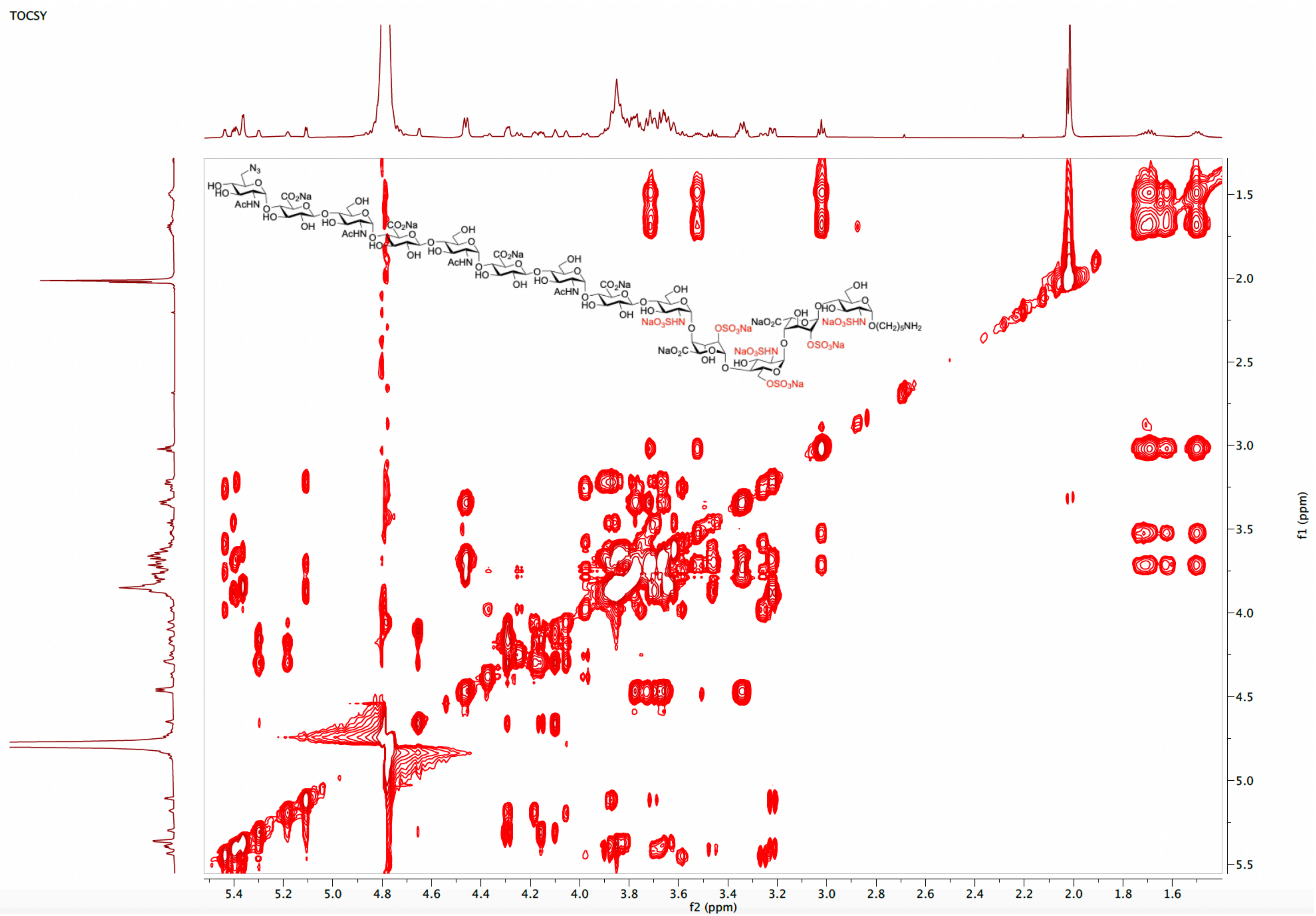

NOESY spectrum of **6** ($D_2O$)

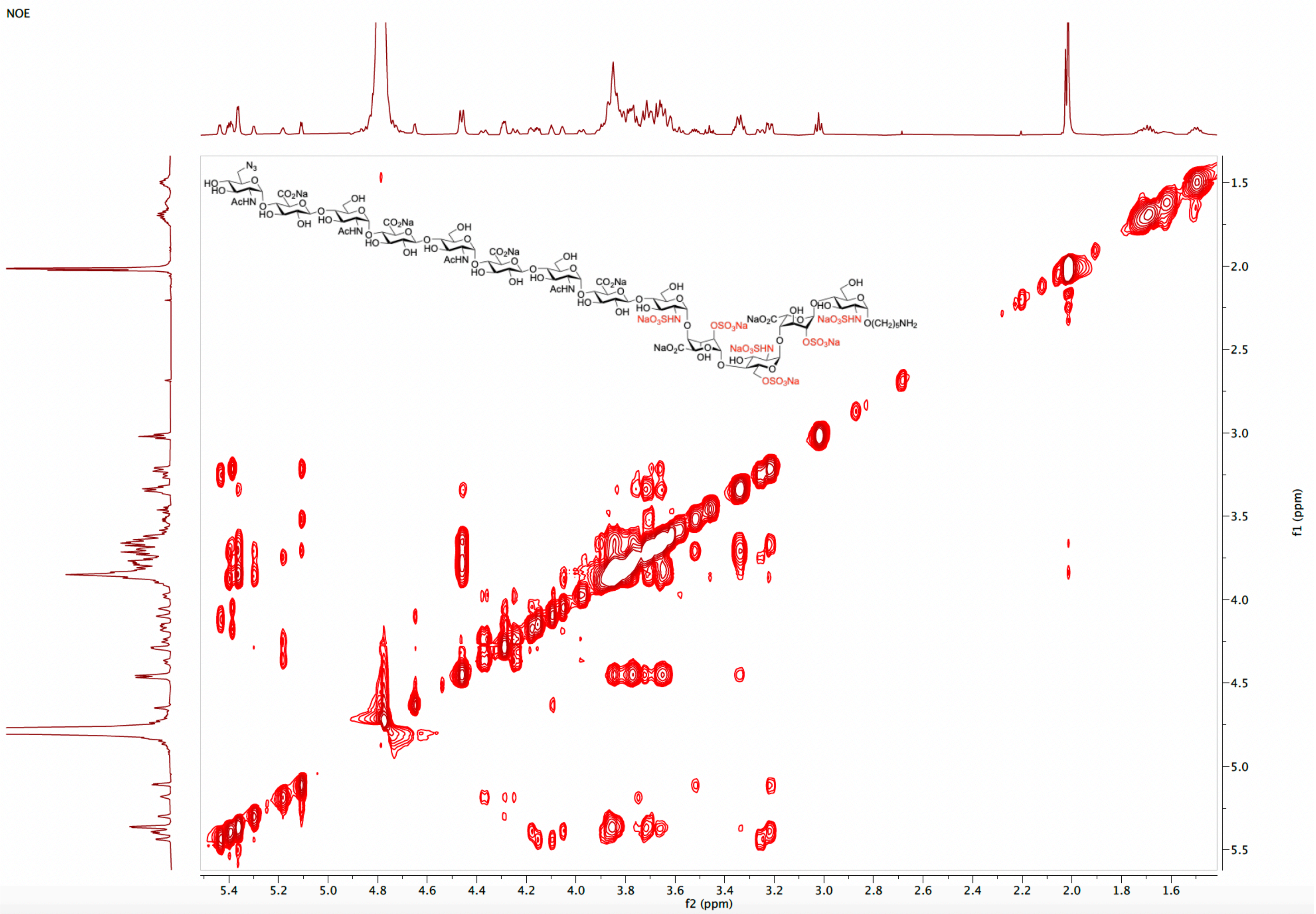

[1]H NMR spectrum of **7** (600 MHz, $D_2O$)

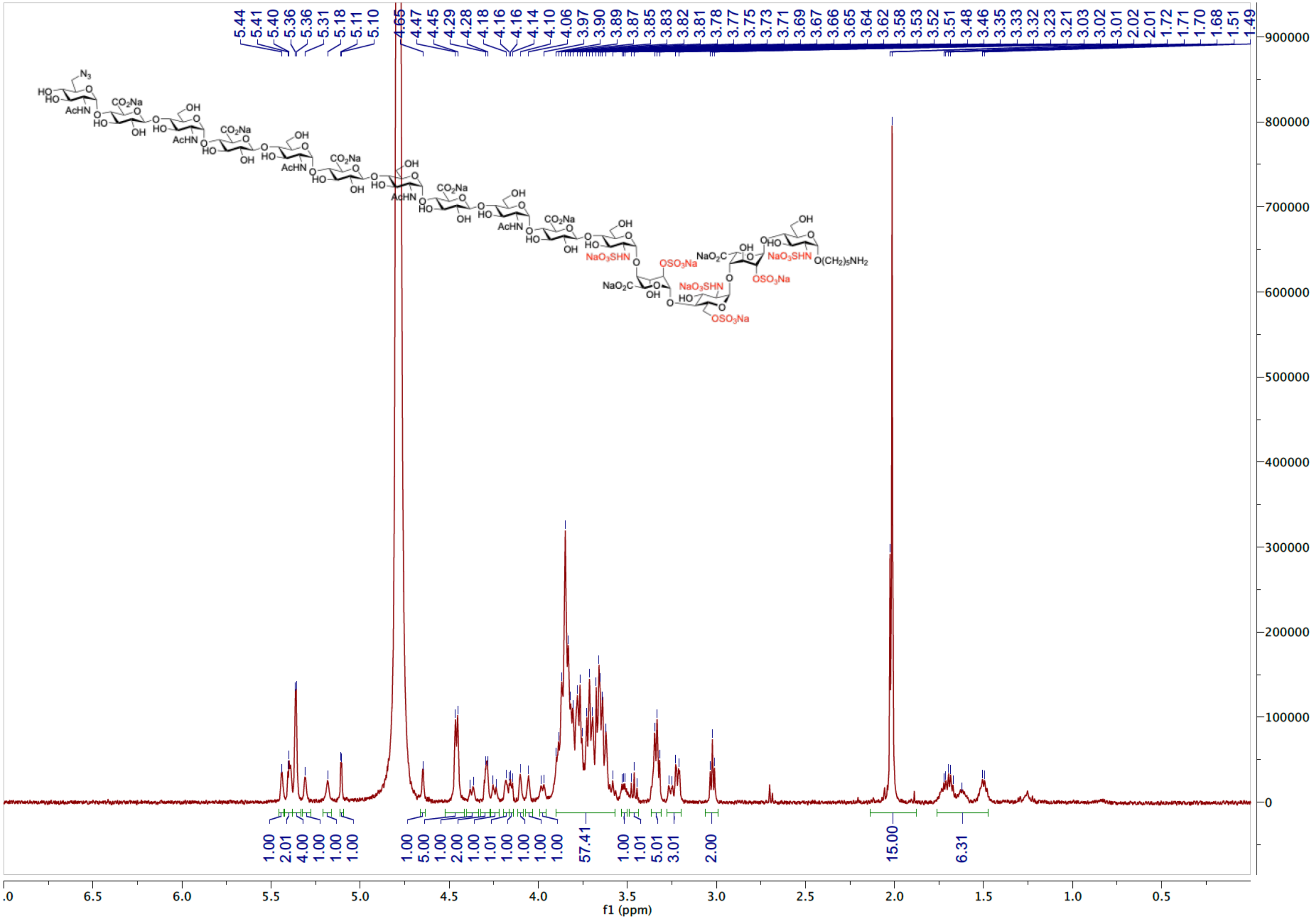

HSQC spectrum of **7** ($D_2O$)

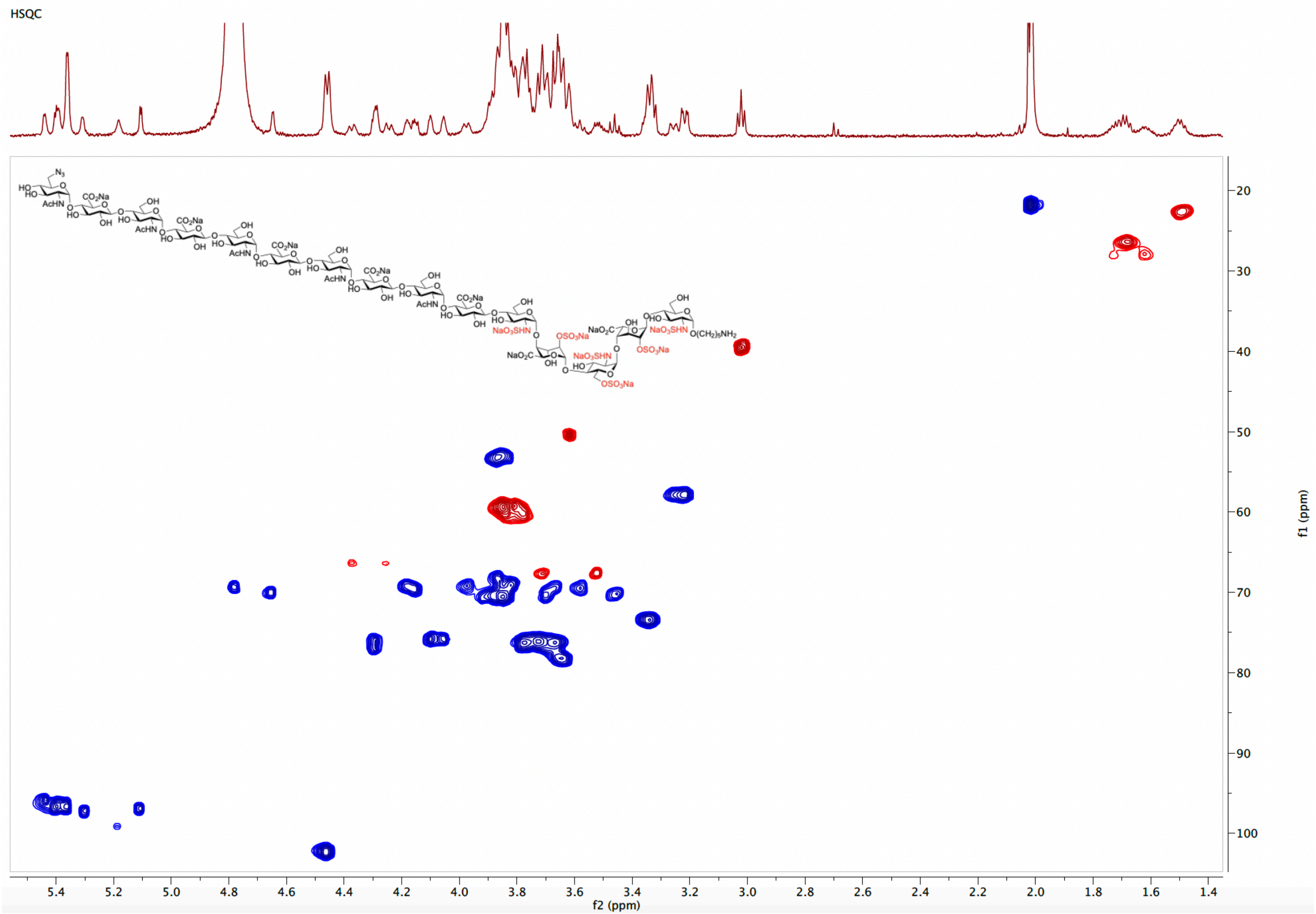

TOCSY spectrum of **7** ($D_2O$)

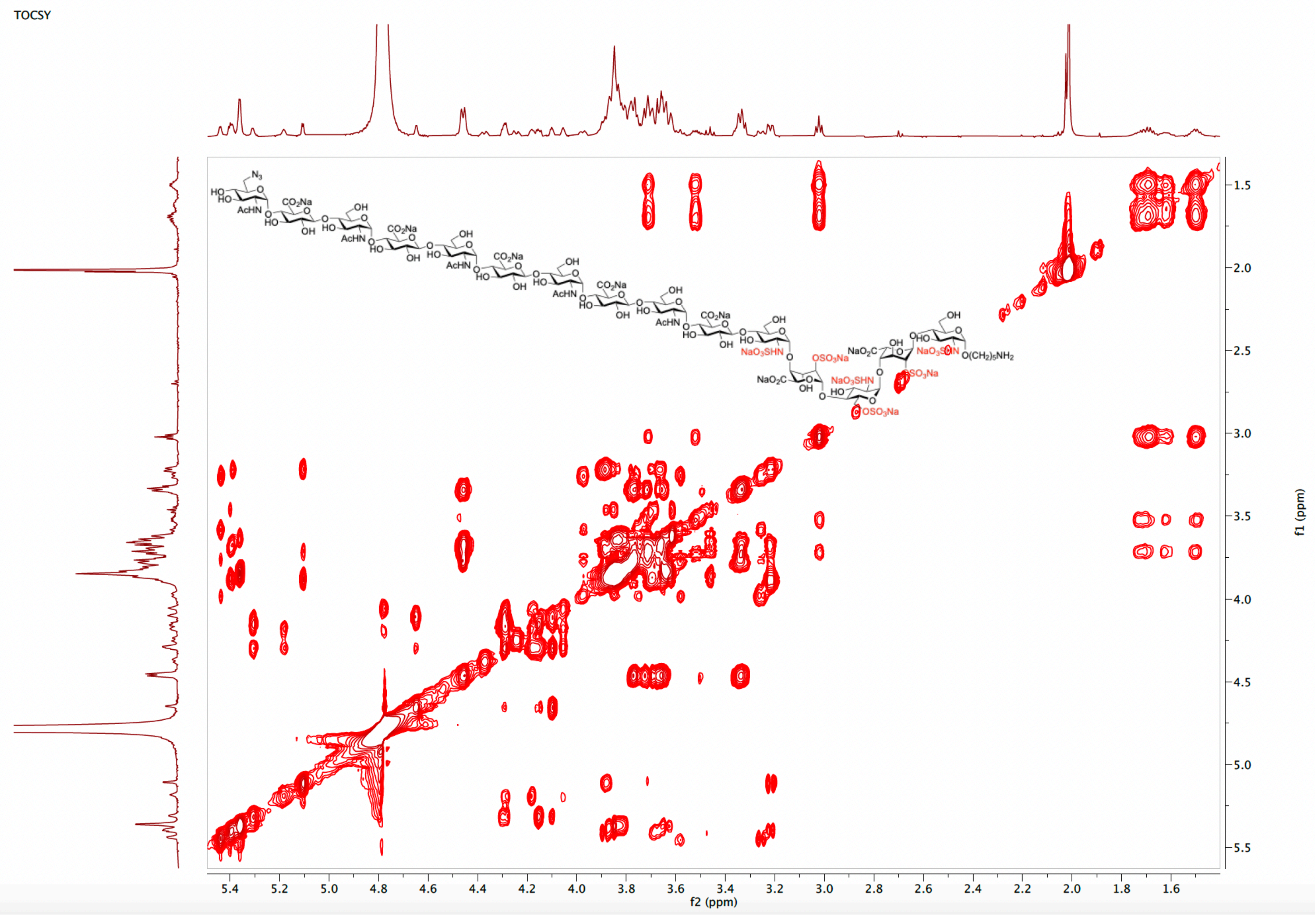

NOE spectrum of **7** ($D_2O$)

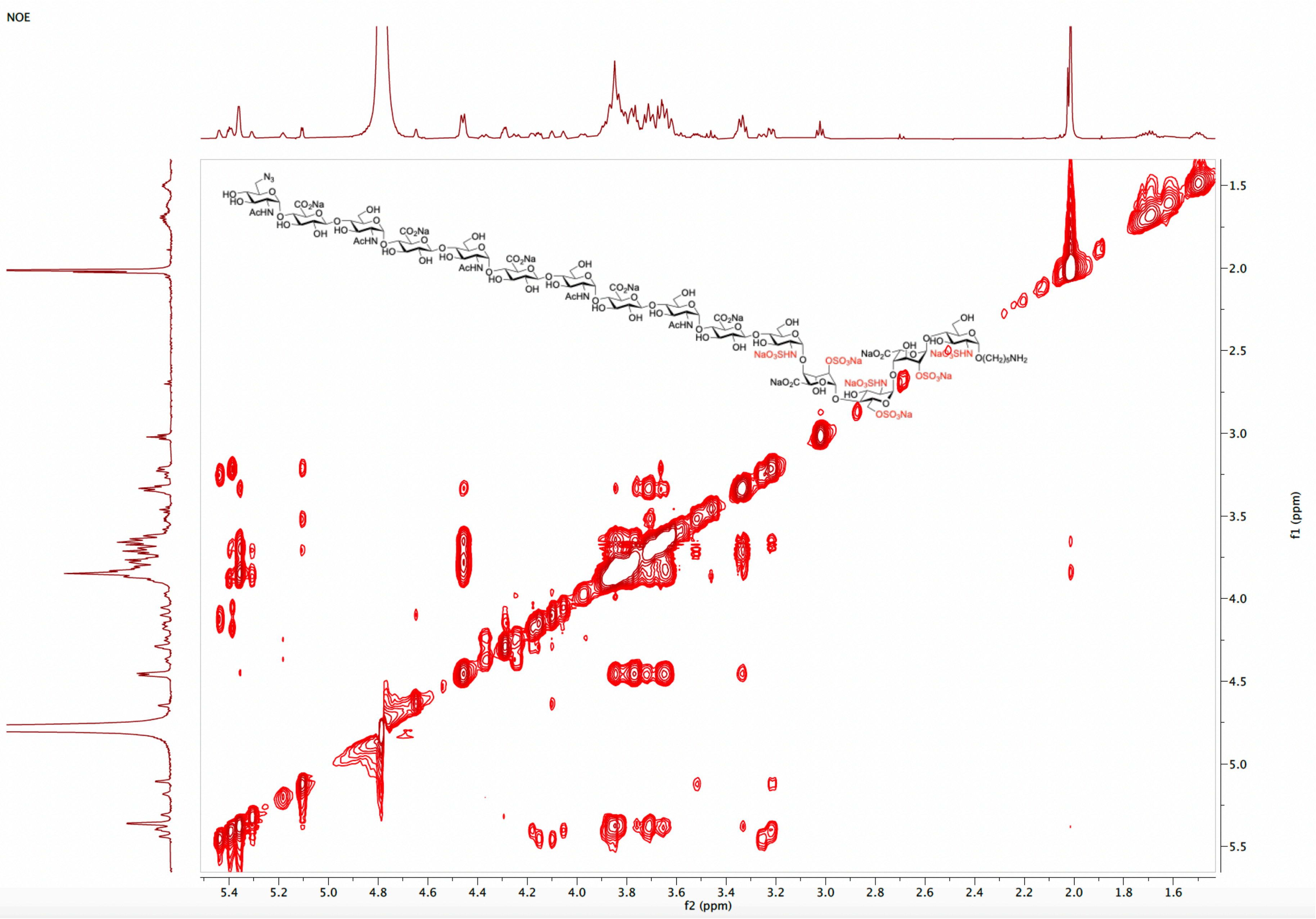

[1]H NMR spectrum of **8** (600 MHz, $D_2O$)

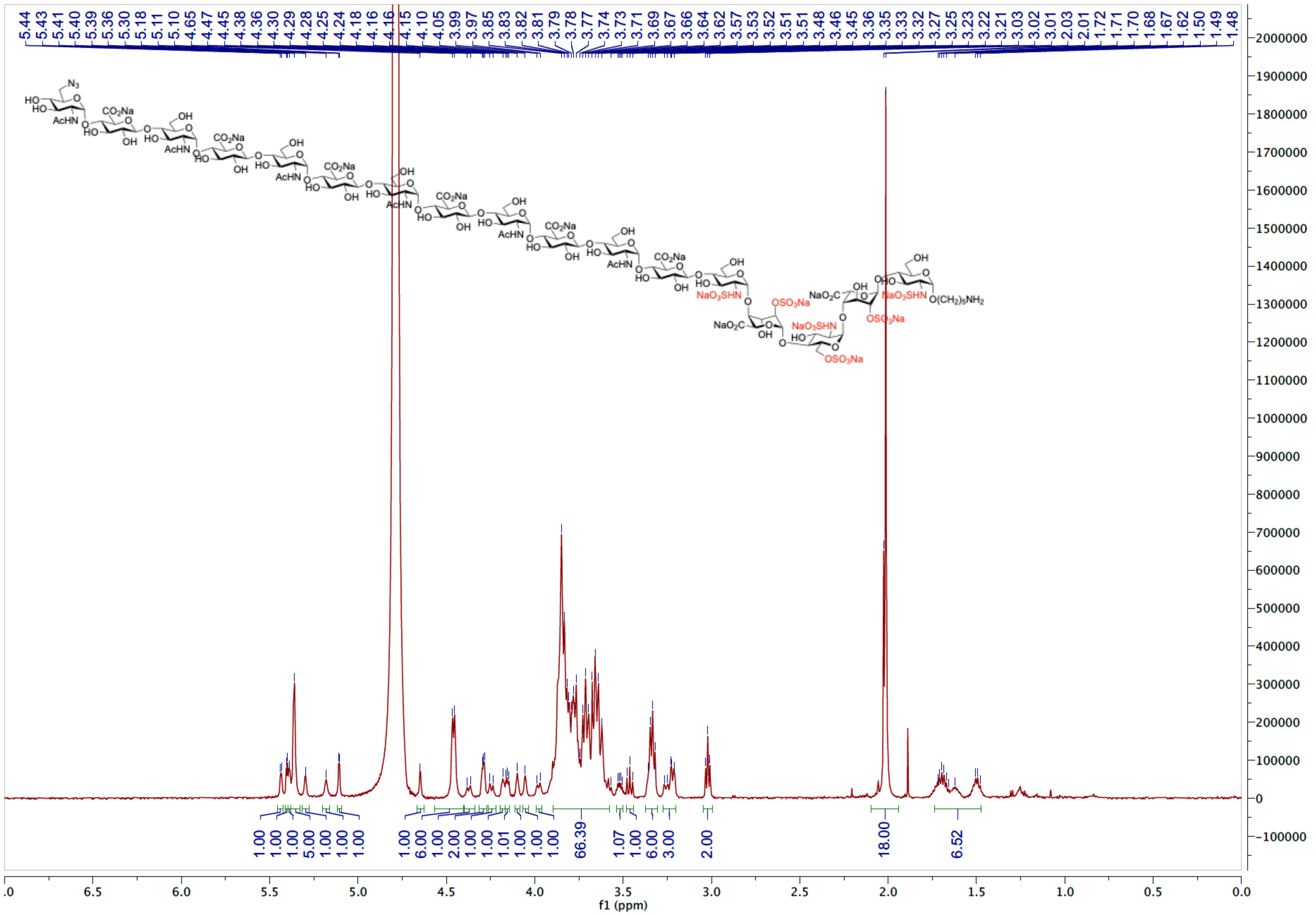

HSQC spectrum of **8** ($D_2O$)

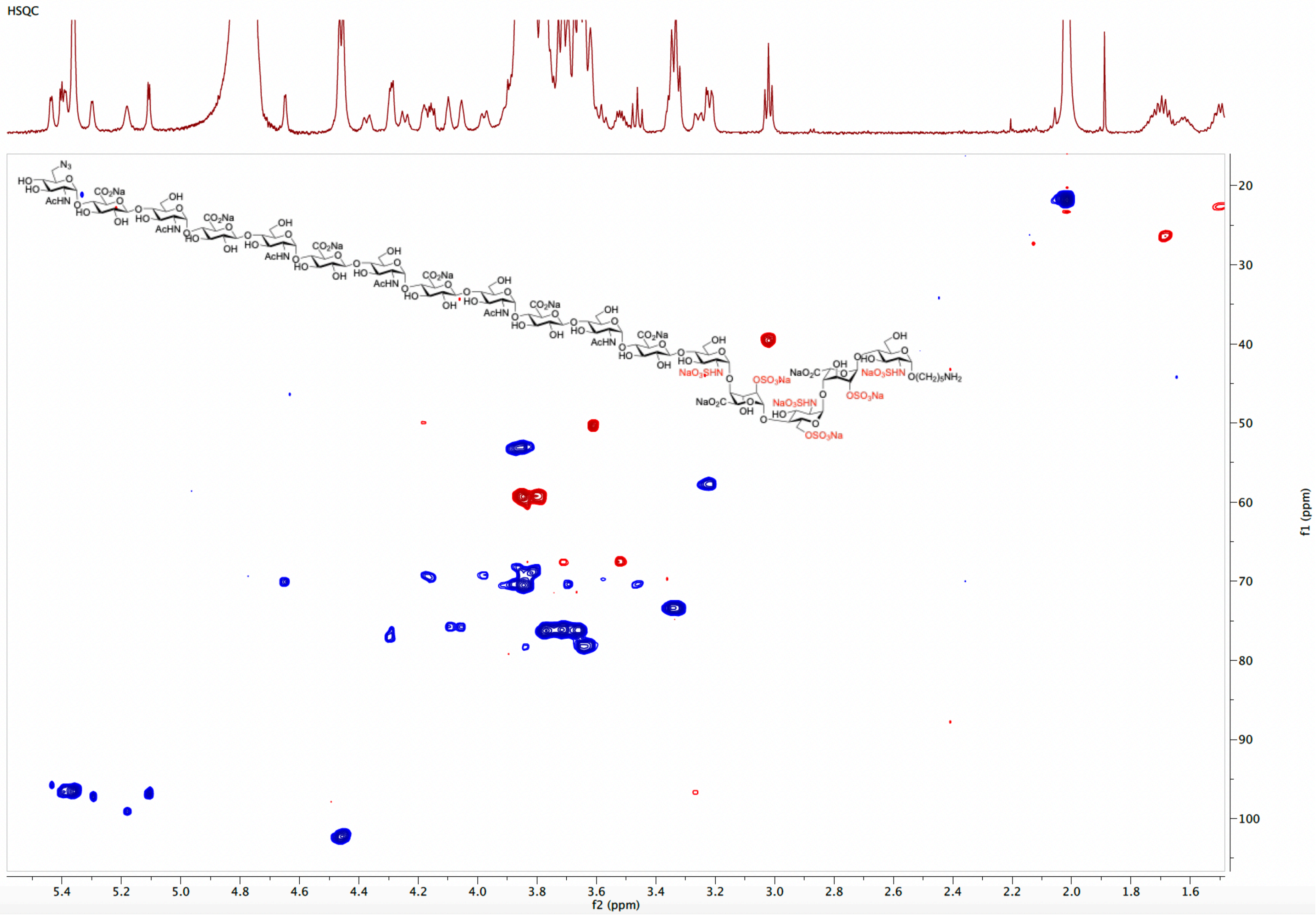

TOCSY spectrum of **8** ($D_2O$)

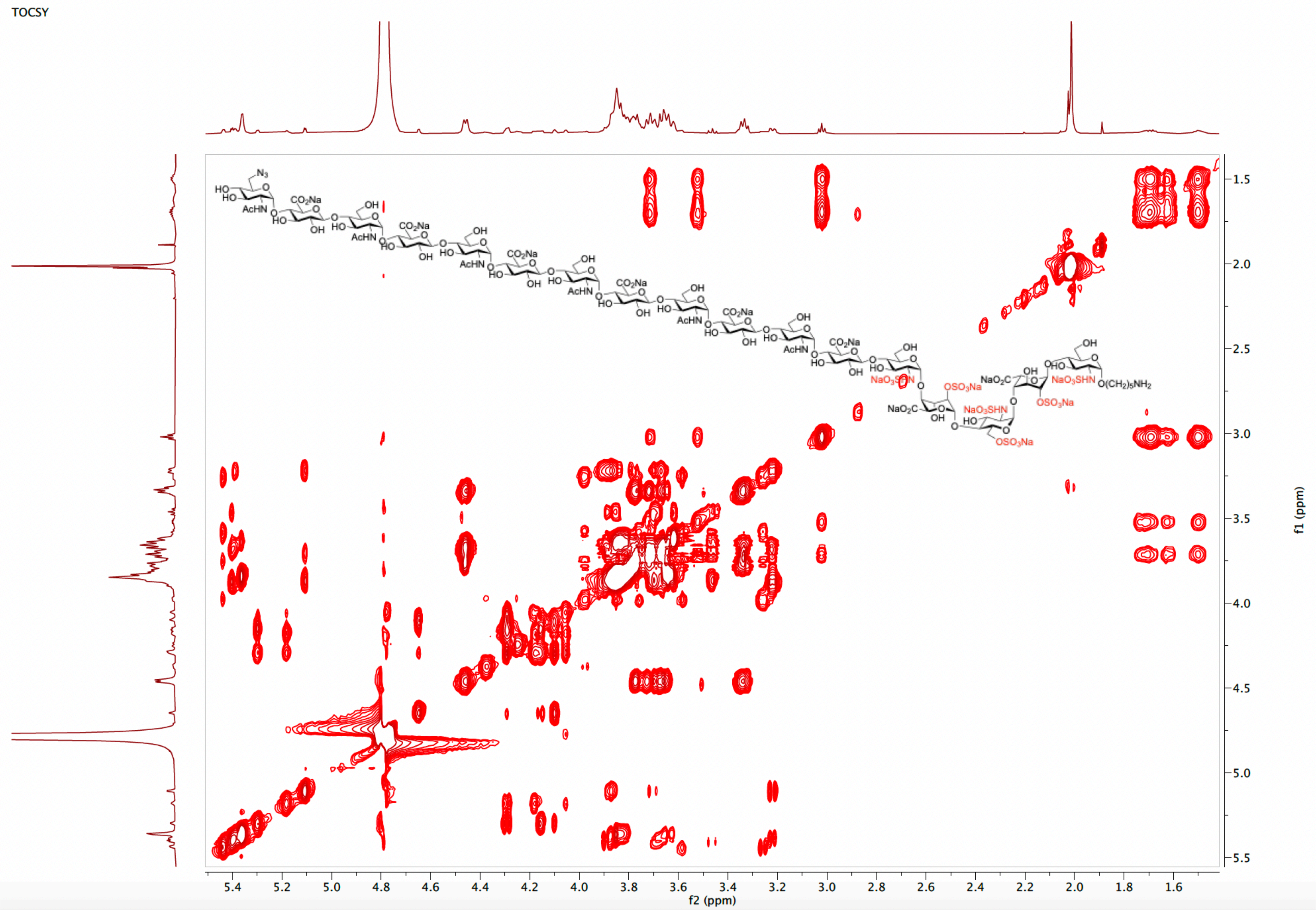

[1]H NMR spectrum of **9** (600 MHz, $D_2O$)

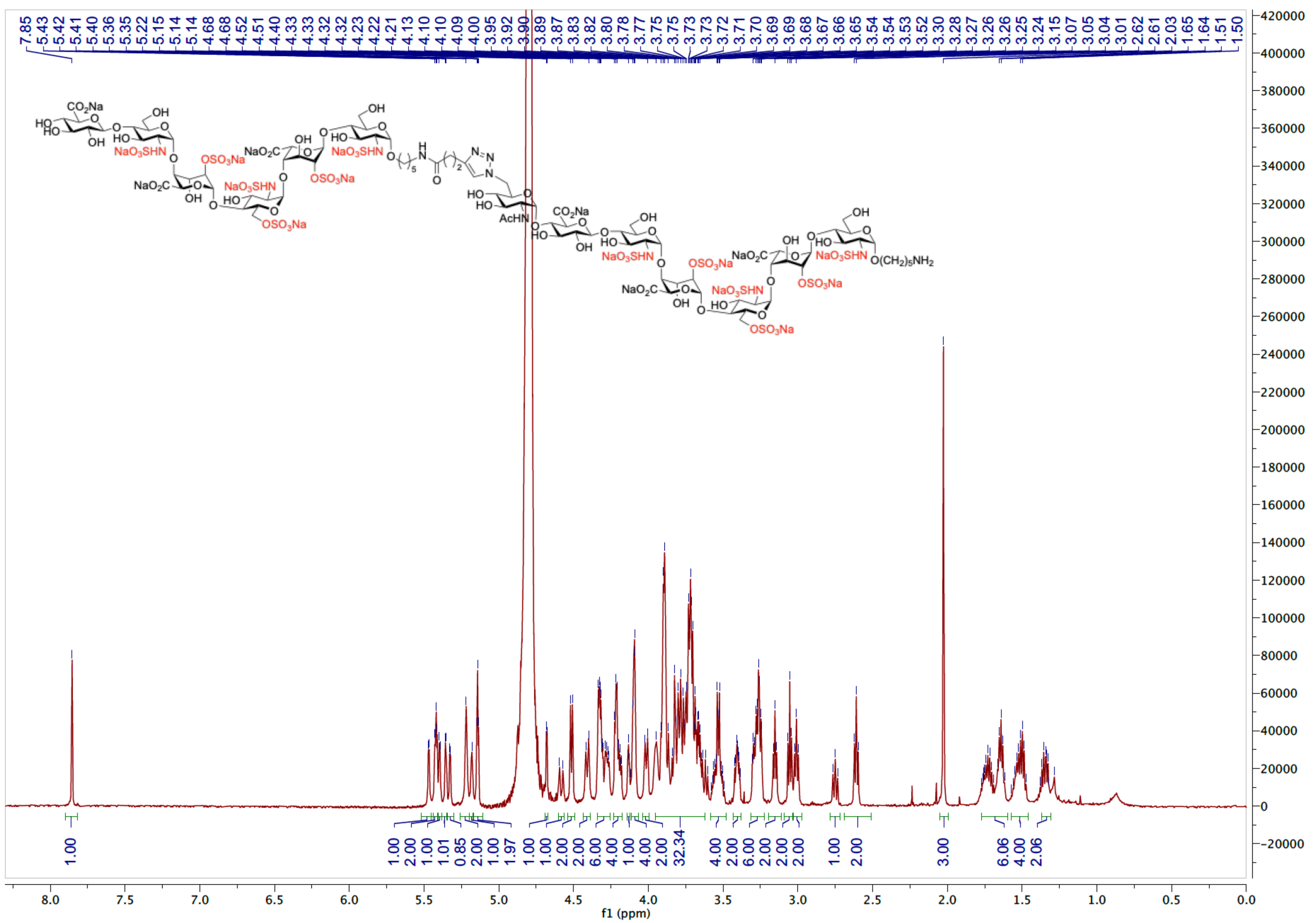

HSQC spectrum of **9** ($D_2O$)

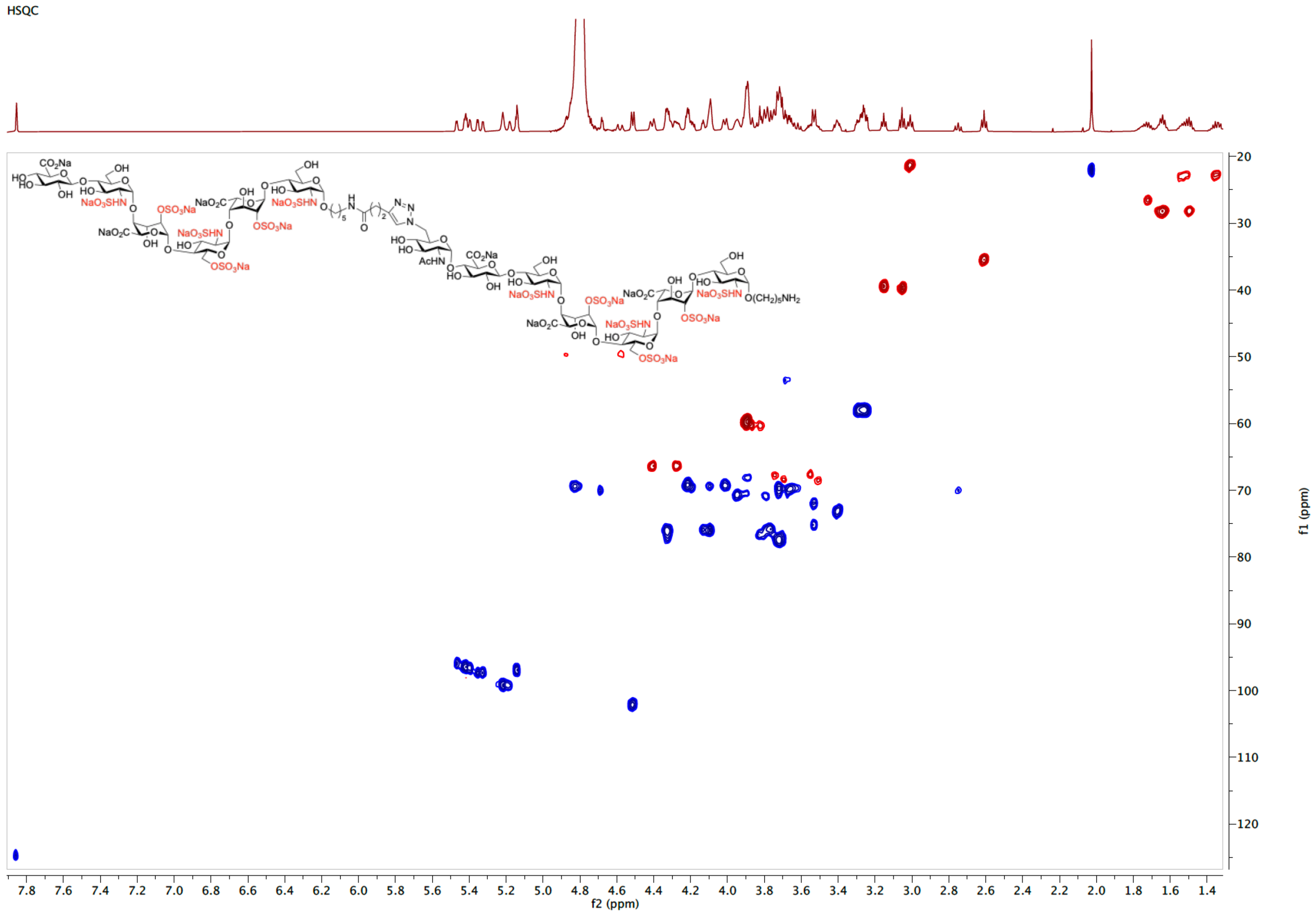

COSY spectrum of **9** ($D_2O$)

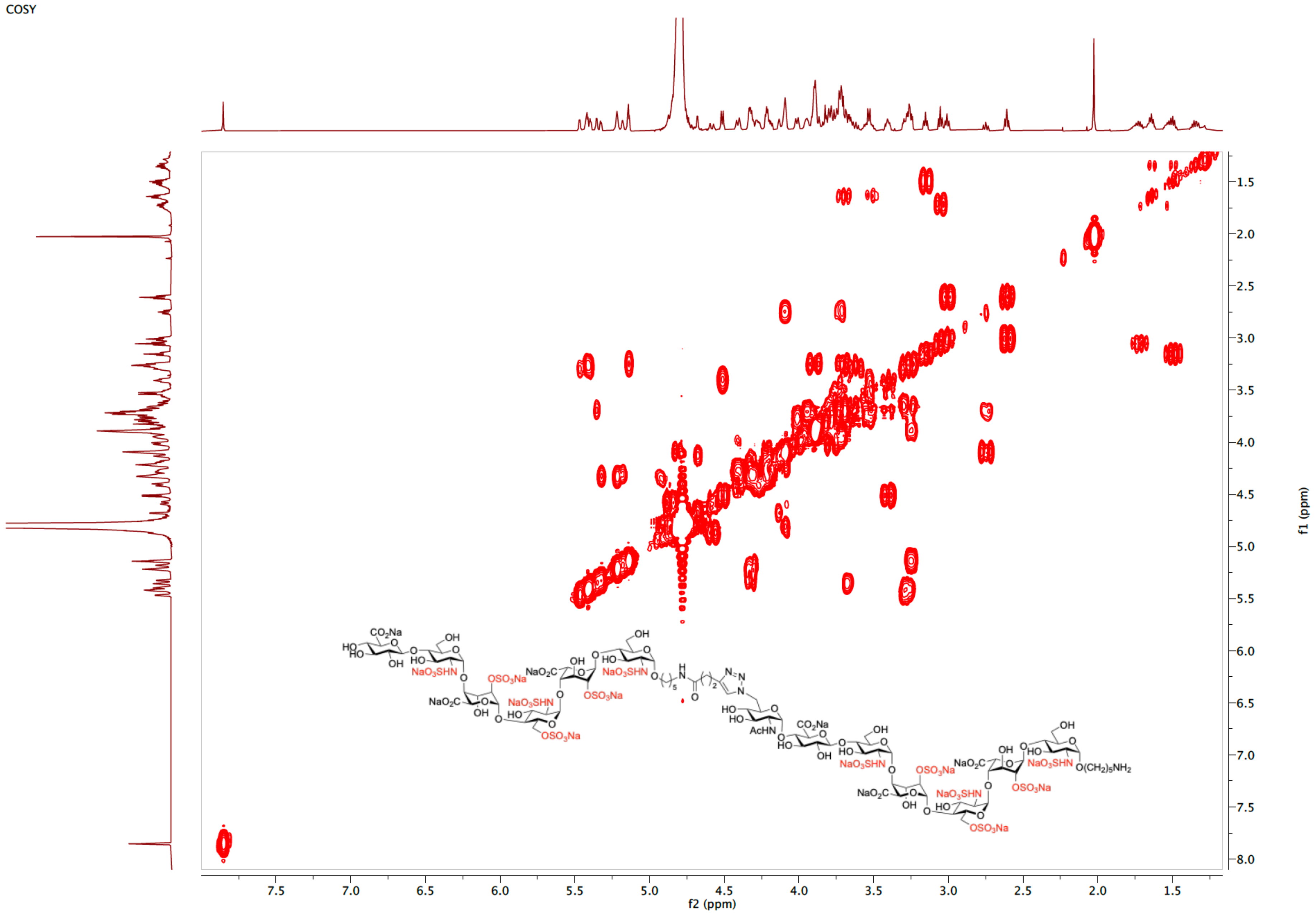

TOCSY spectrum of **9** ($D_2O$)

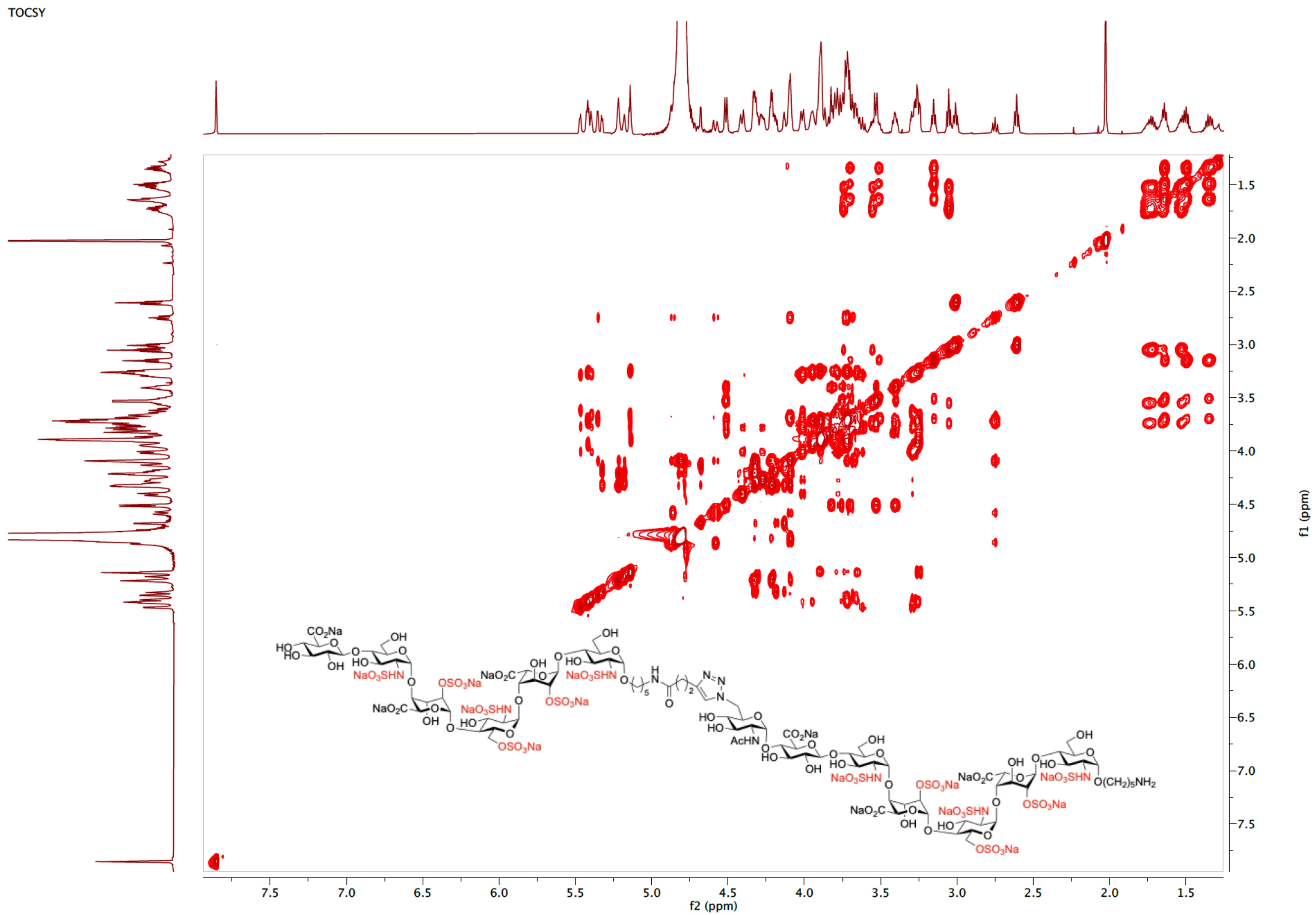

NOESY spectrum of **9** ($D_2O$)

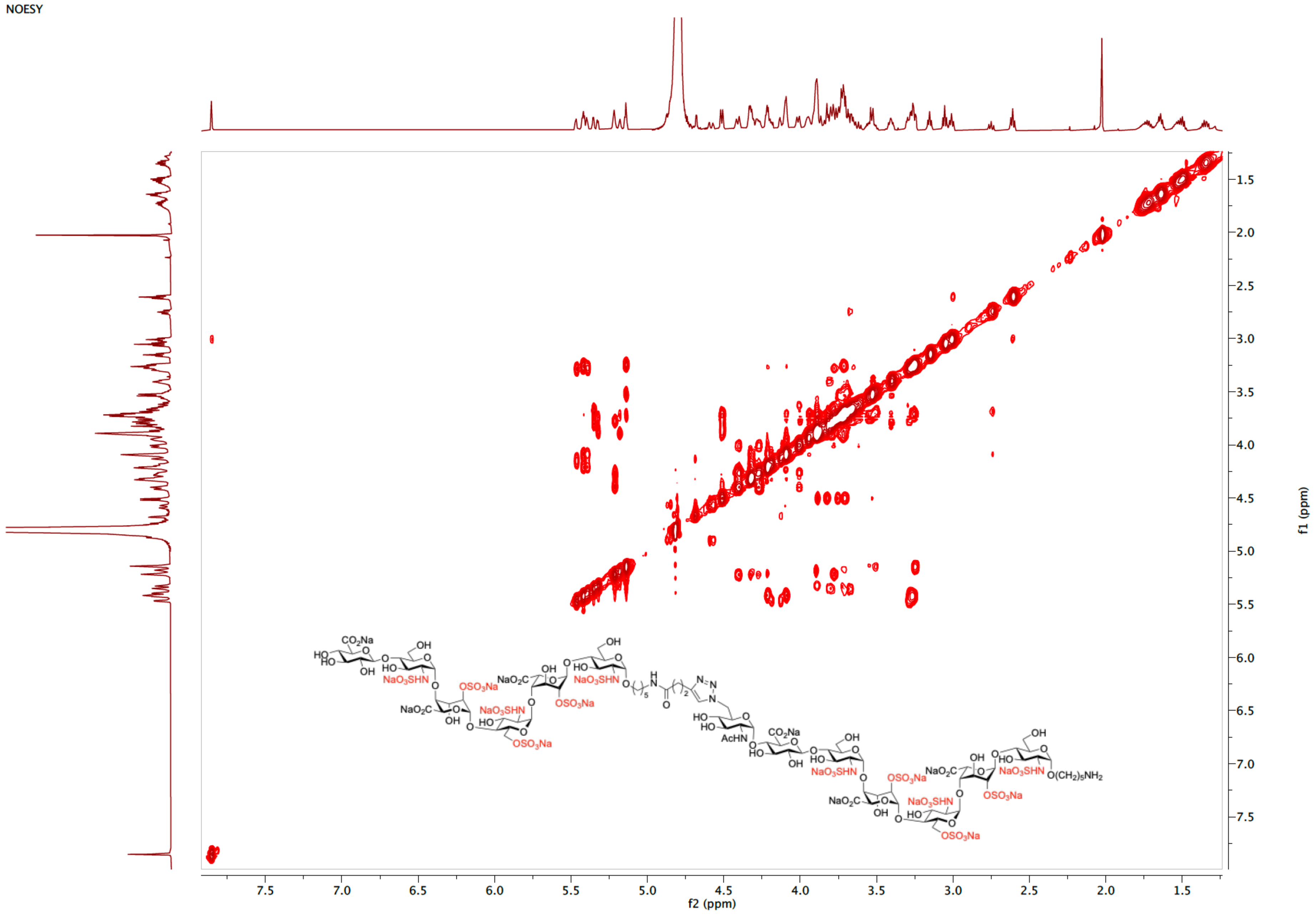

ESI-MS (negative) of **9**

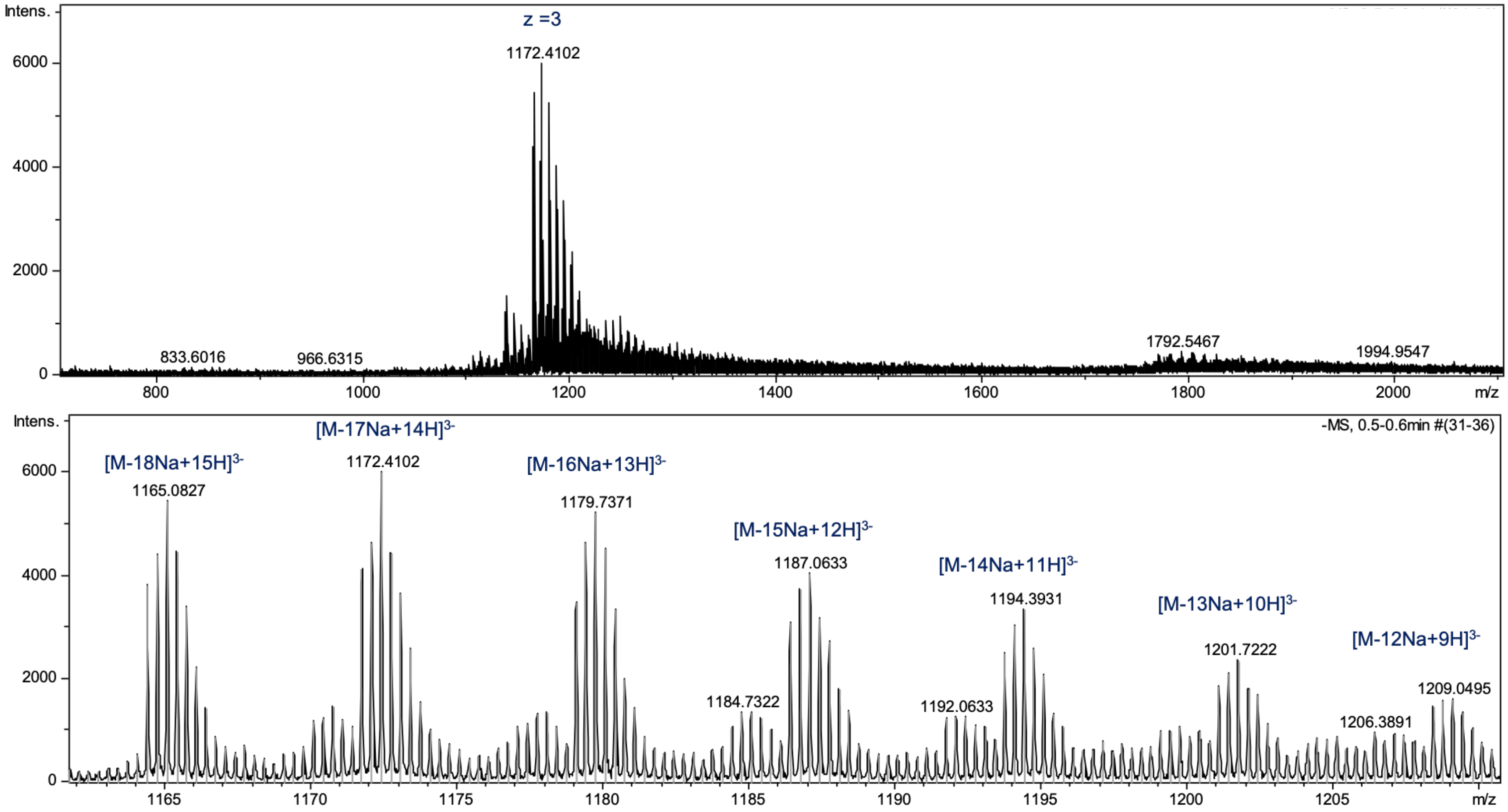

[1]H NMR spectrum of **10** (600 MHz, $D_2O$)

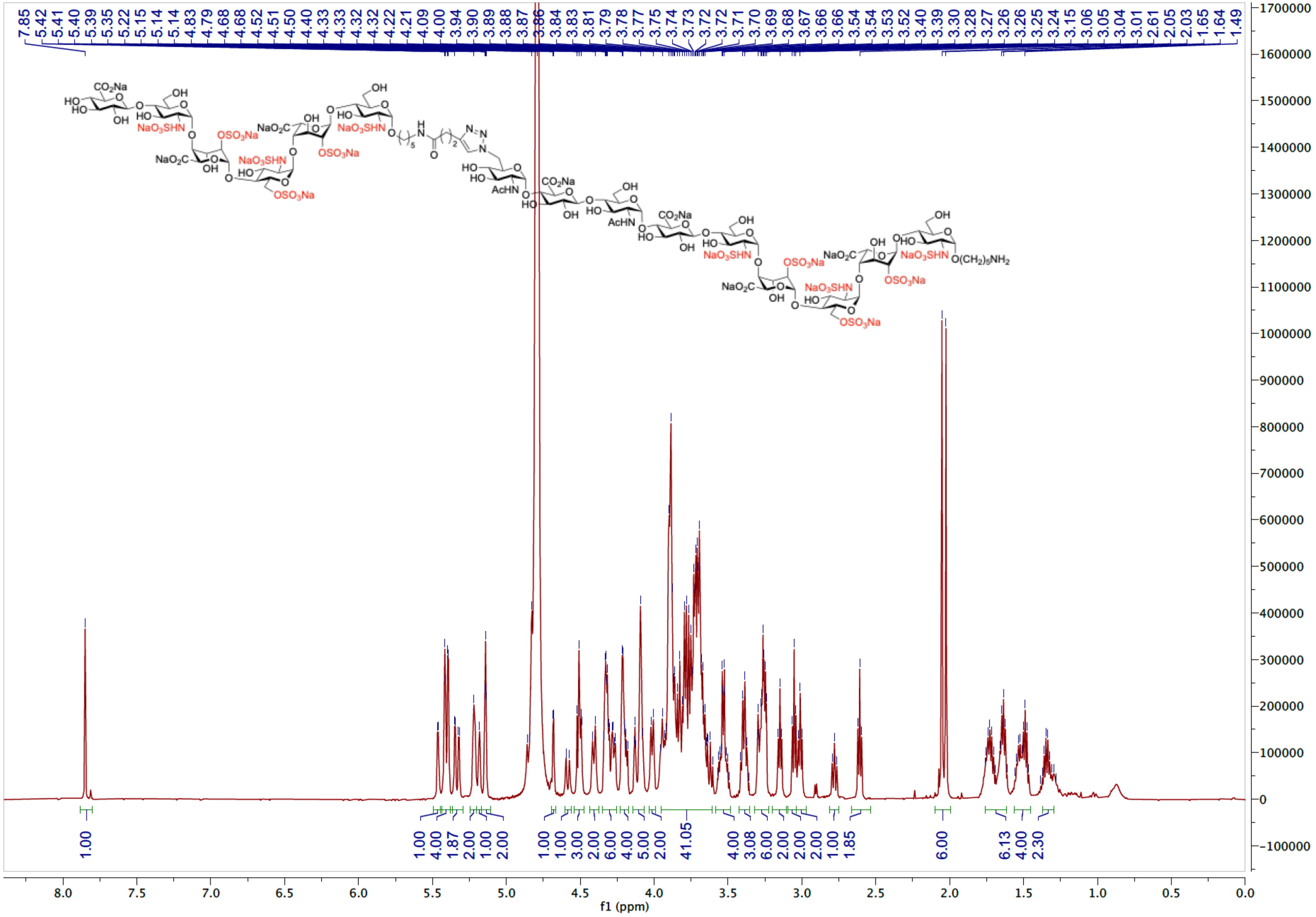

HSQC spectrum of **10** ($D_2O$)

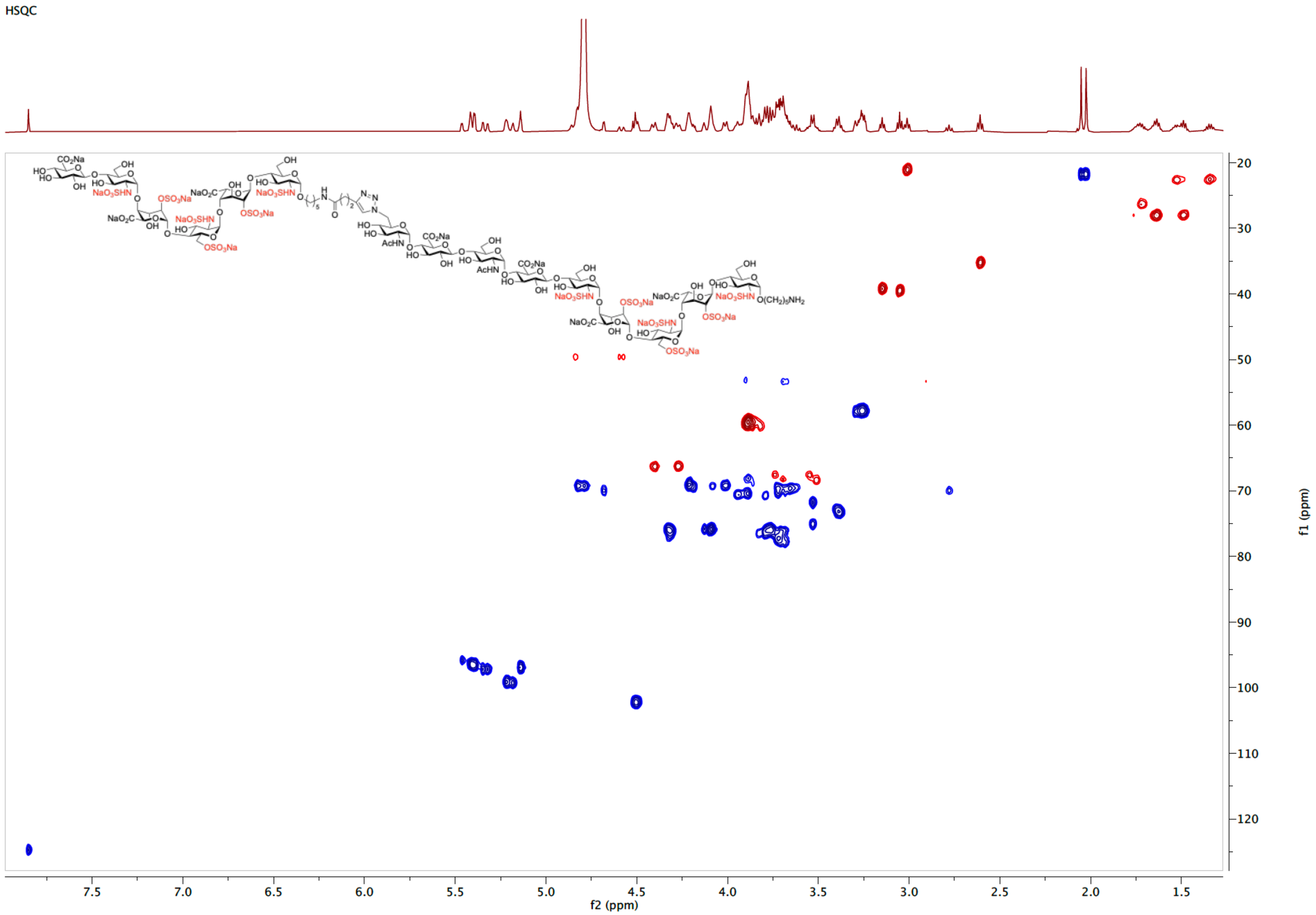

COSY spectrum of **10** ($D_2O$)

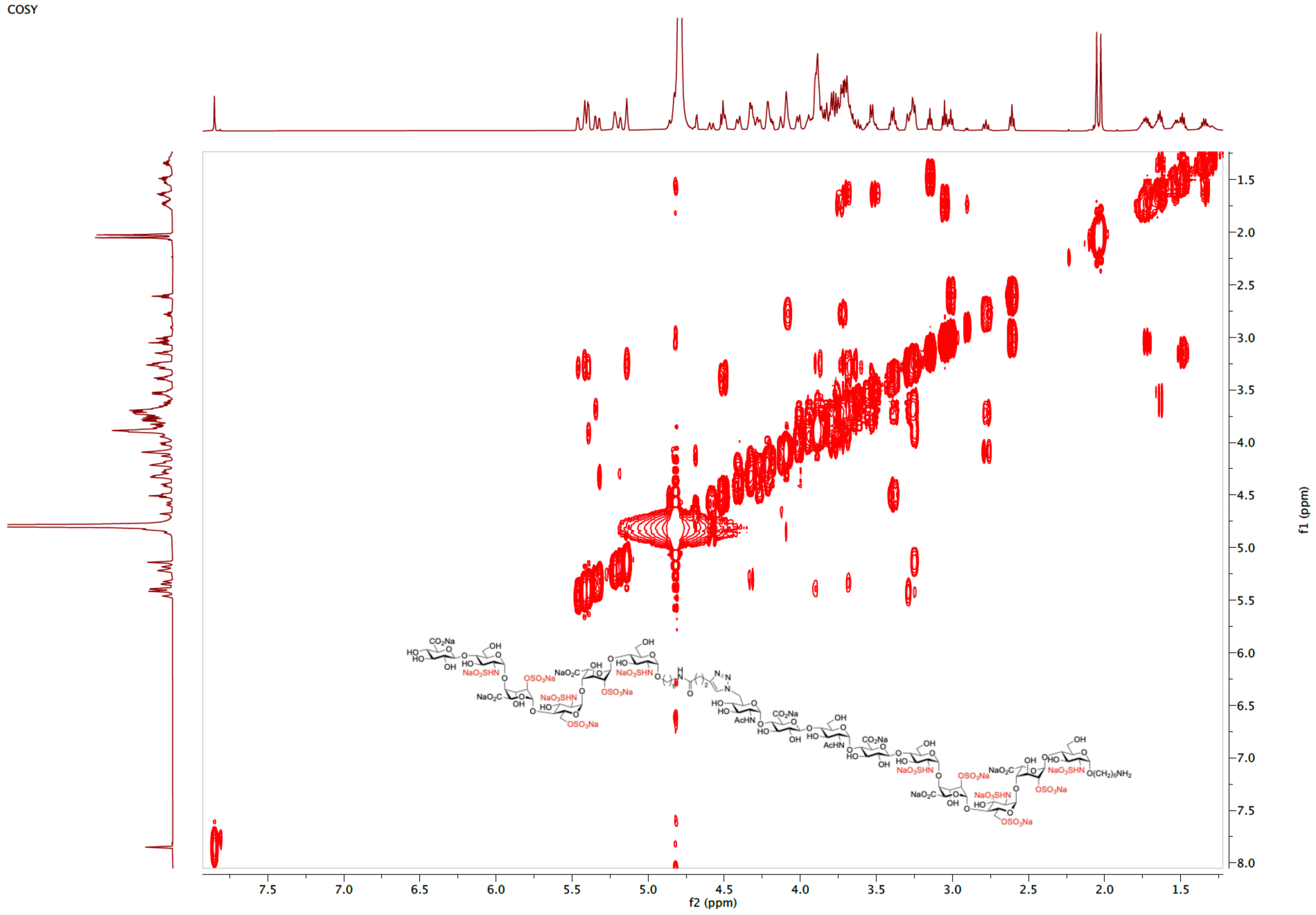

TOCSY spectrum of **10** ($D_2O$)

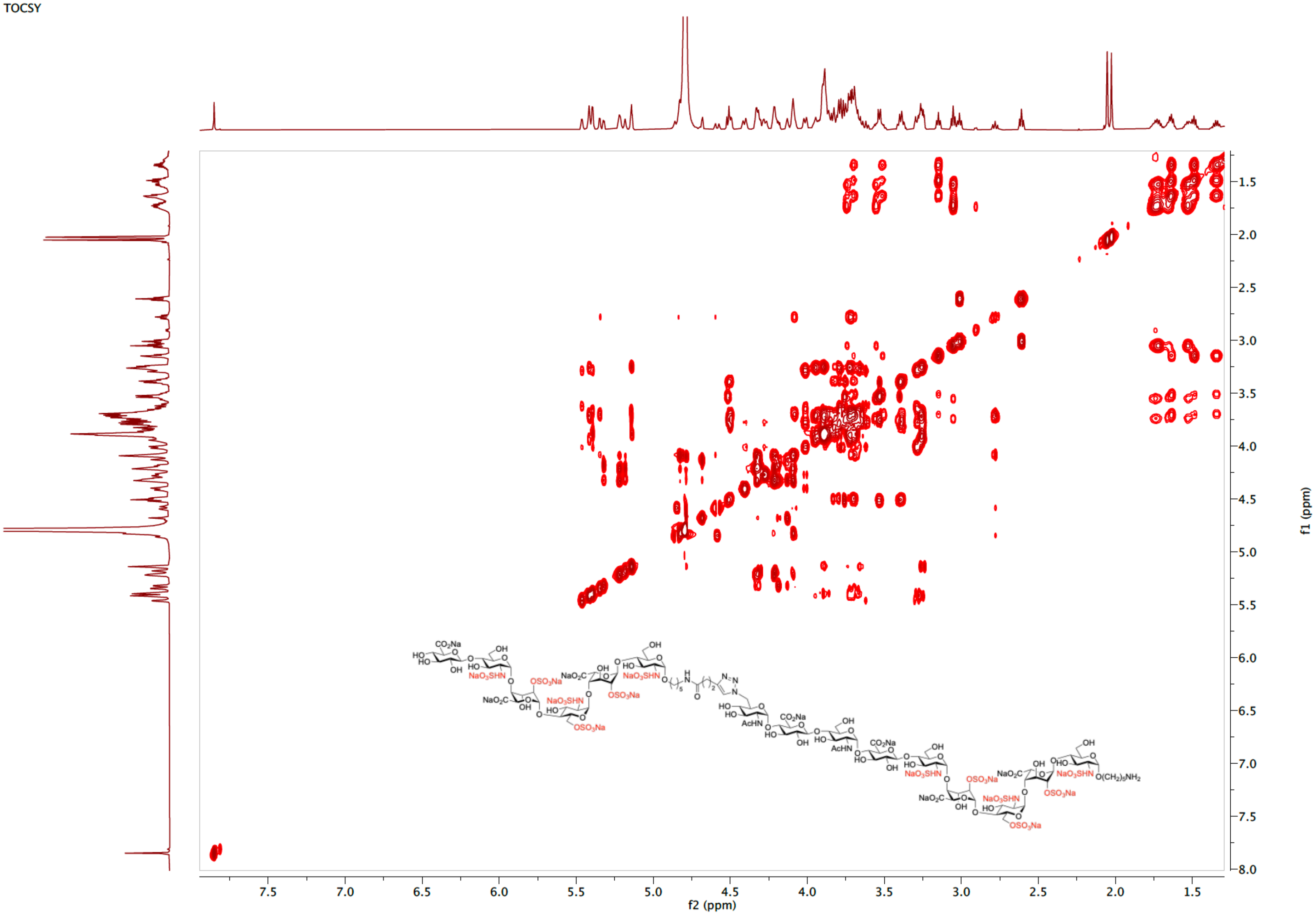

NOESY spectrum of **10** ($D_2O$)

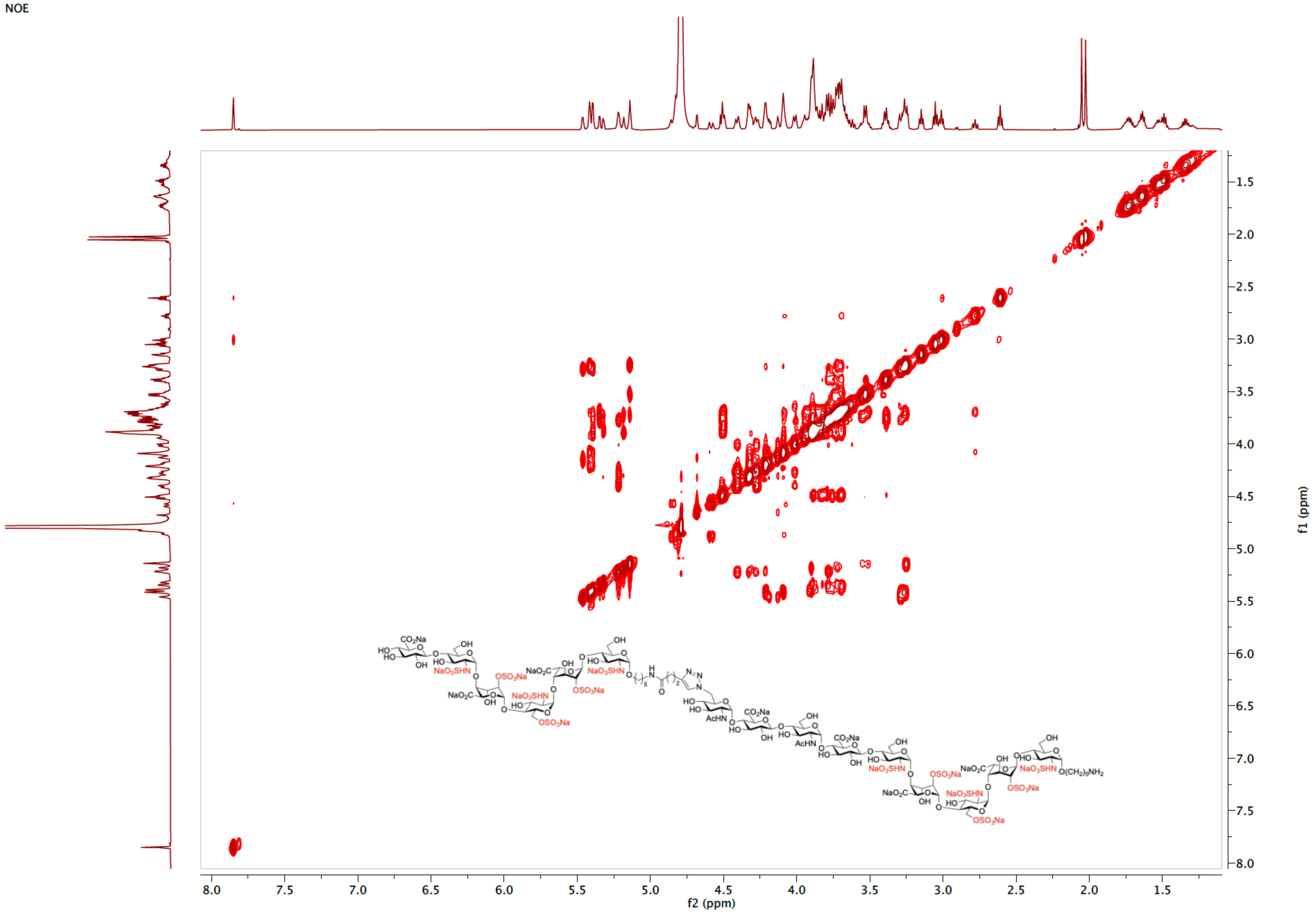

ESI-MS (negative) of **10**

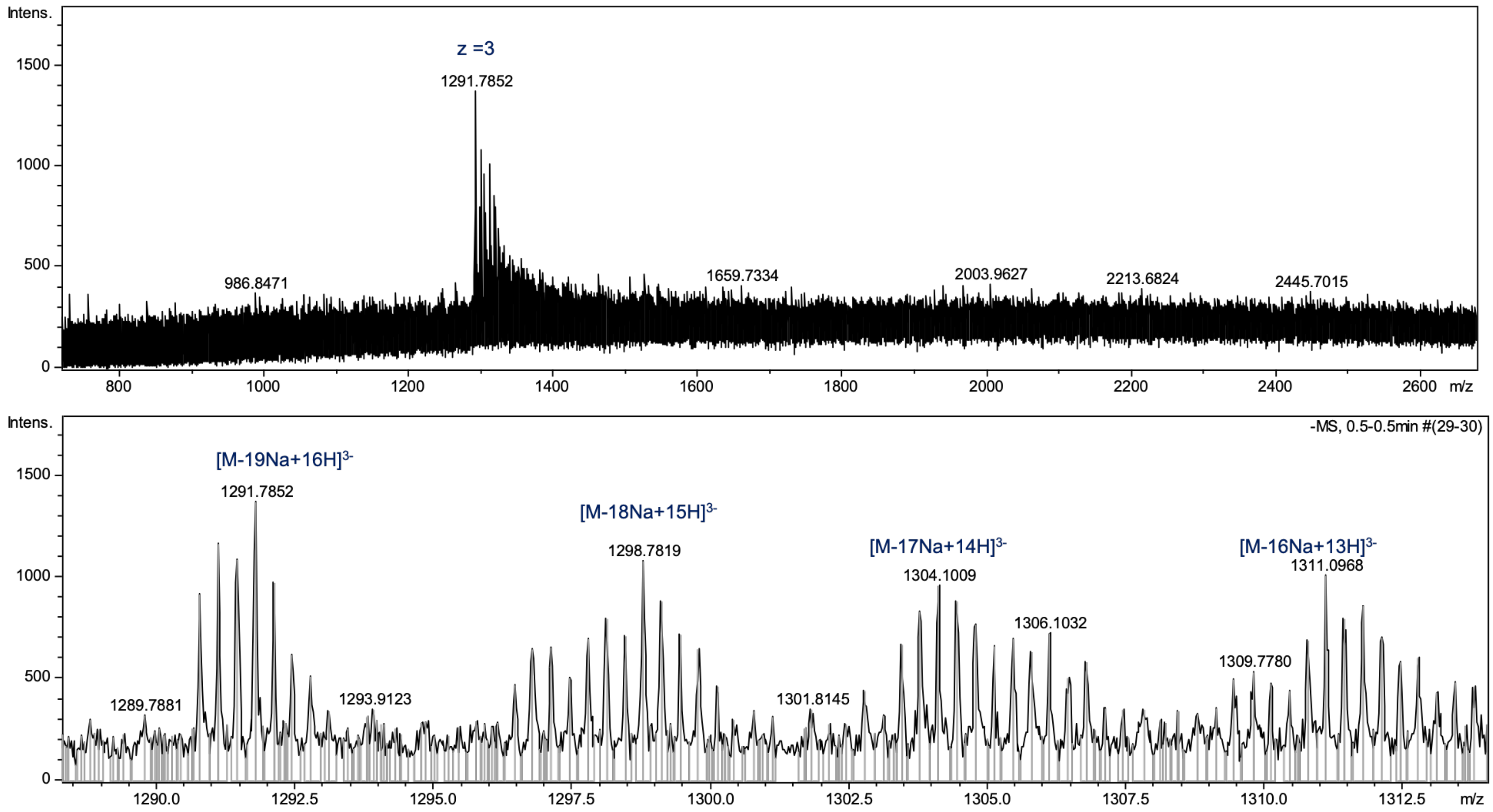

$^{1}H$ NMR spectrum of **11** (600 MHz, $D_2O$)

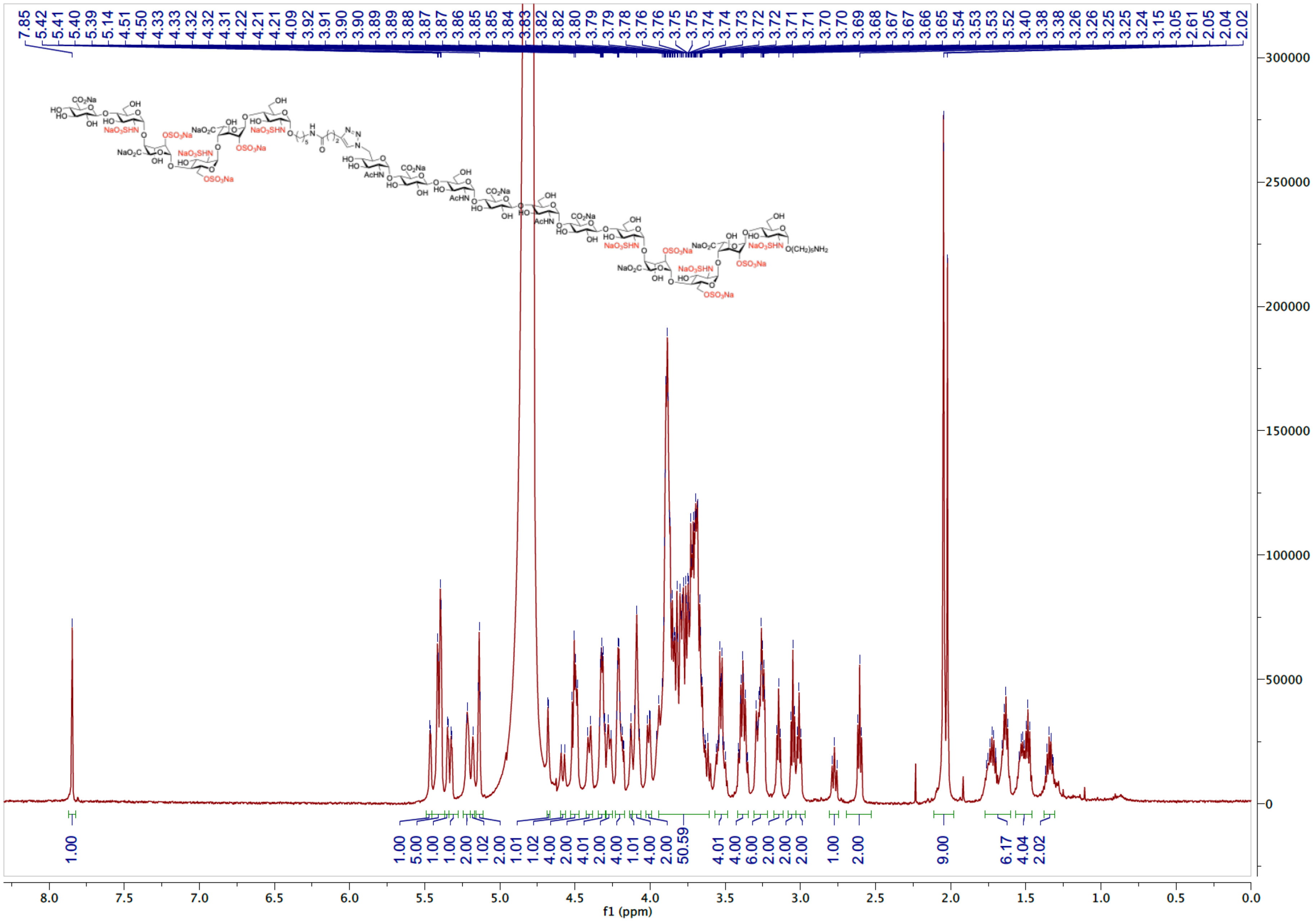

HSQC spectrum of **11** ($D_2O$)

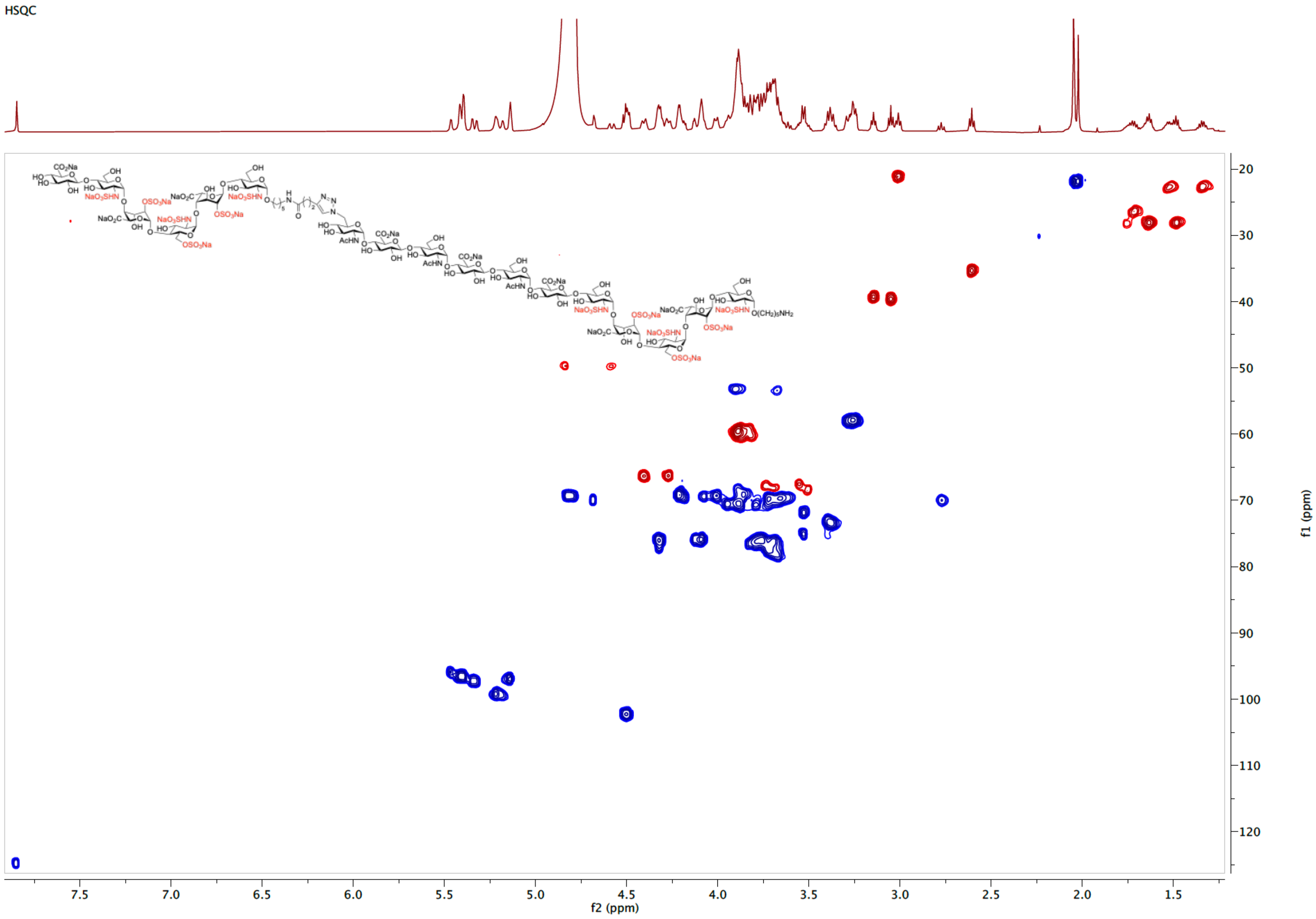

COSY spectrum of **11** ($D_2O$)

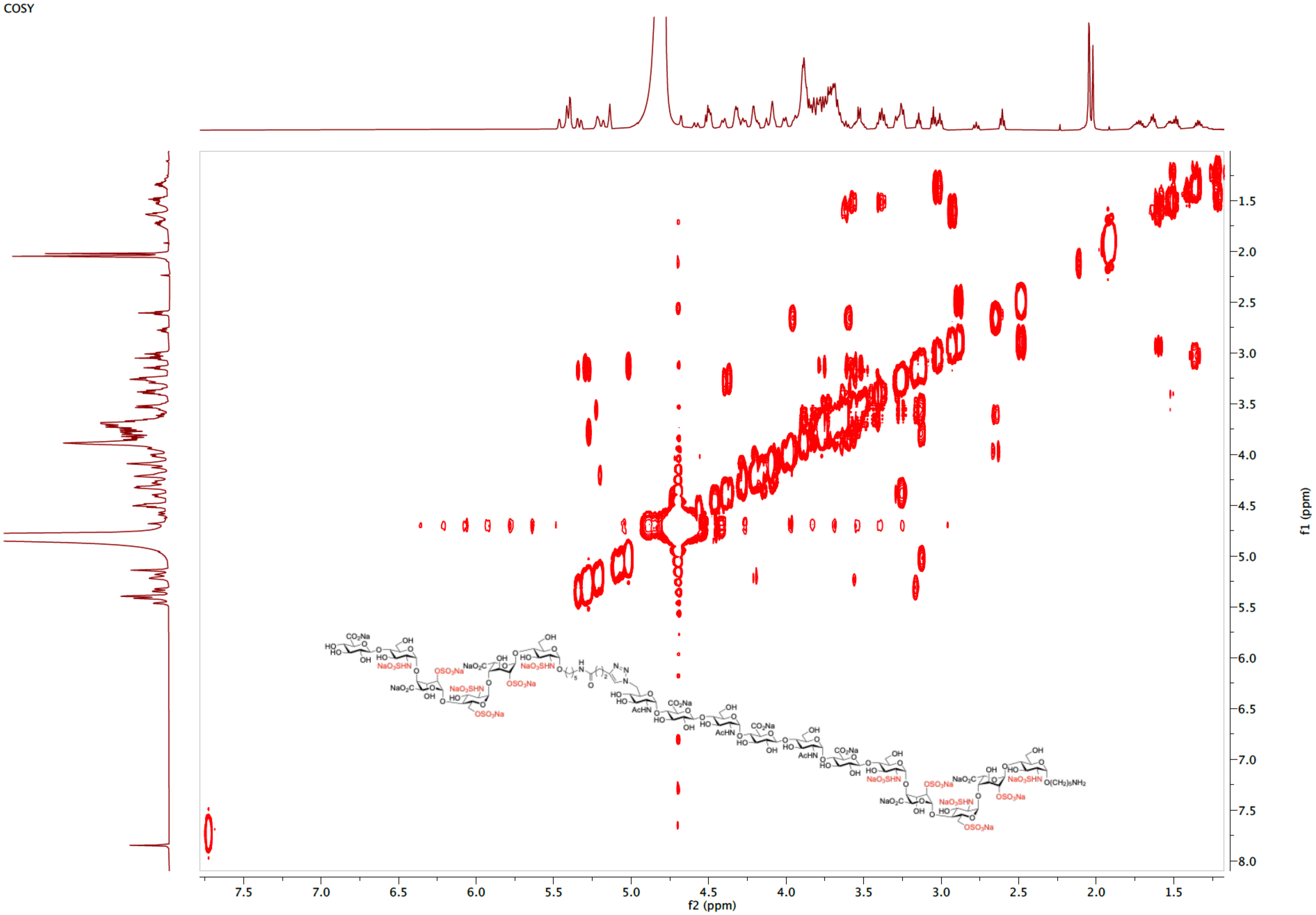

TOCSY spectrum of **11** ($D_2O$)

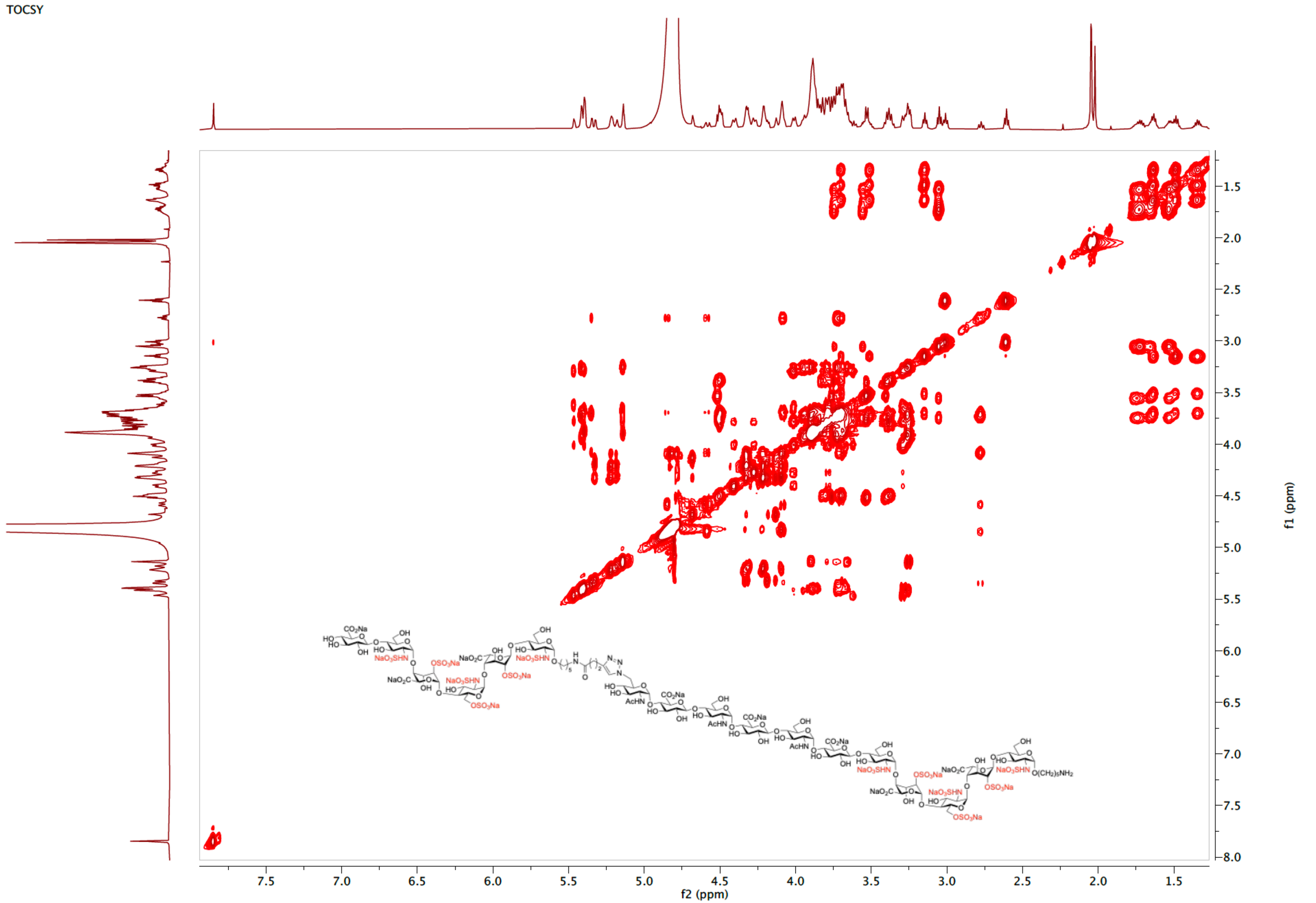

NOESY spectrum of **11** ($D_2O$)

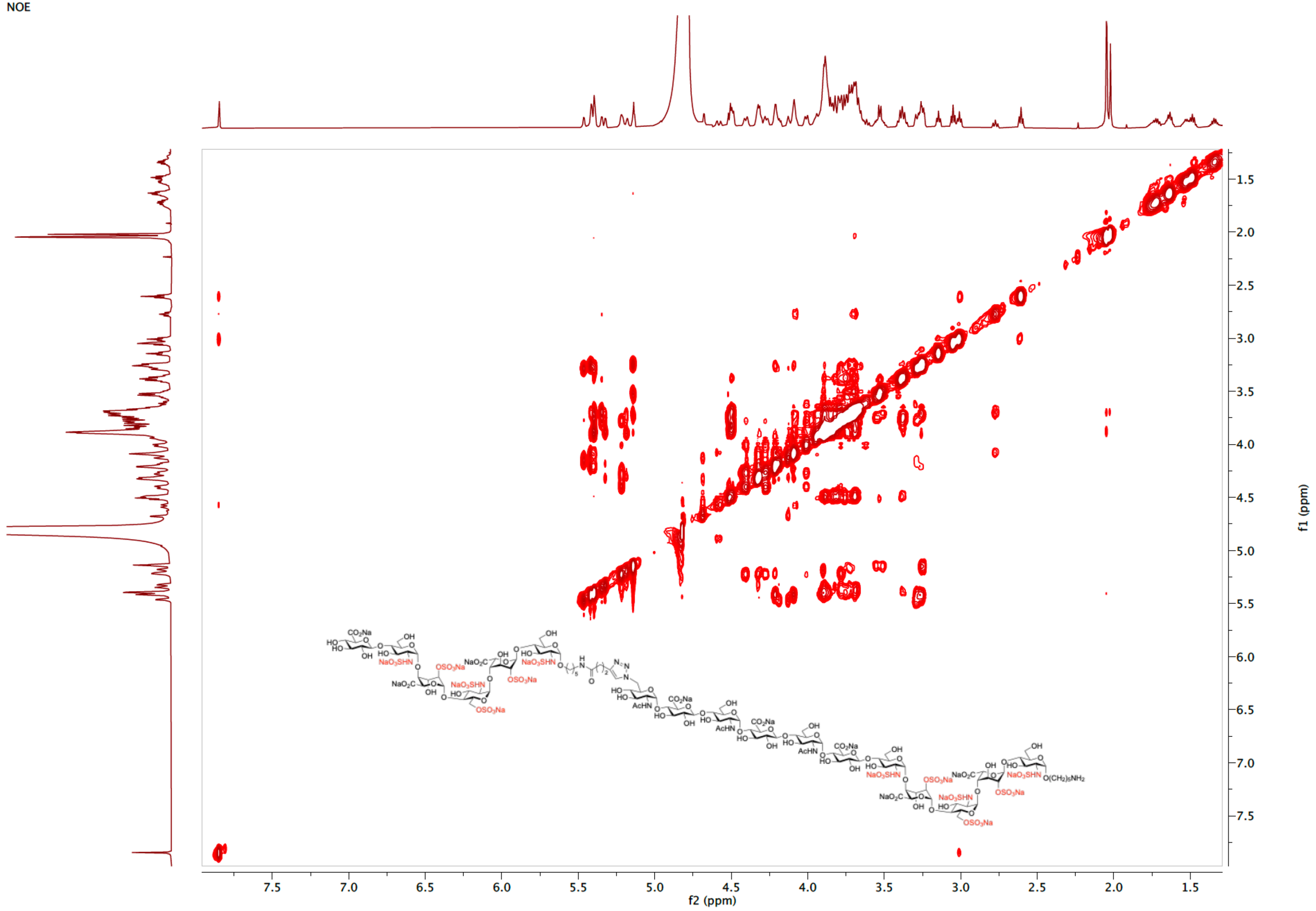

ESI-MS (negative) of **11**

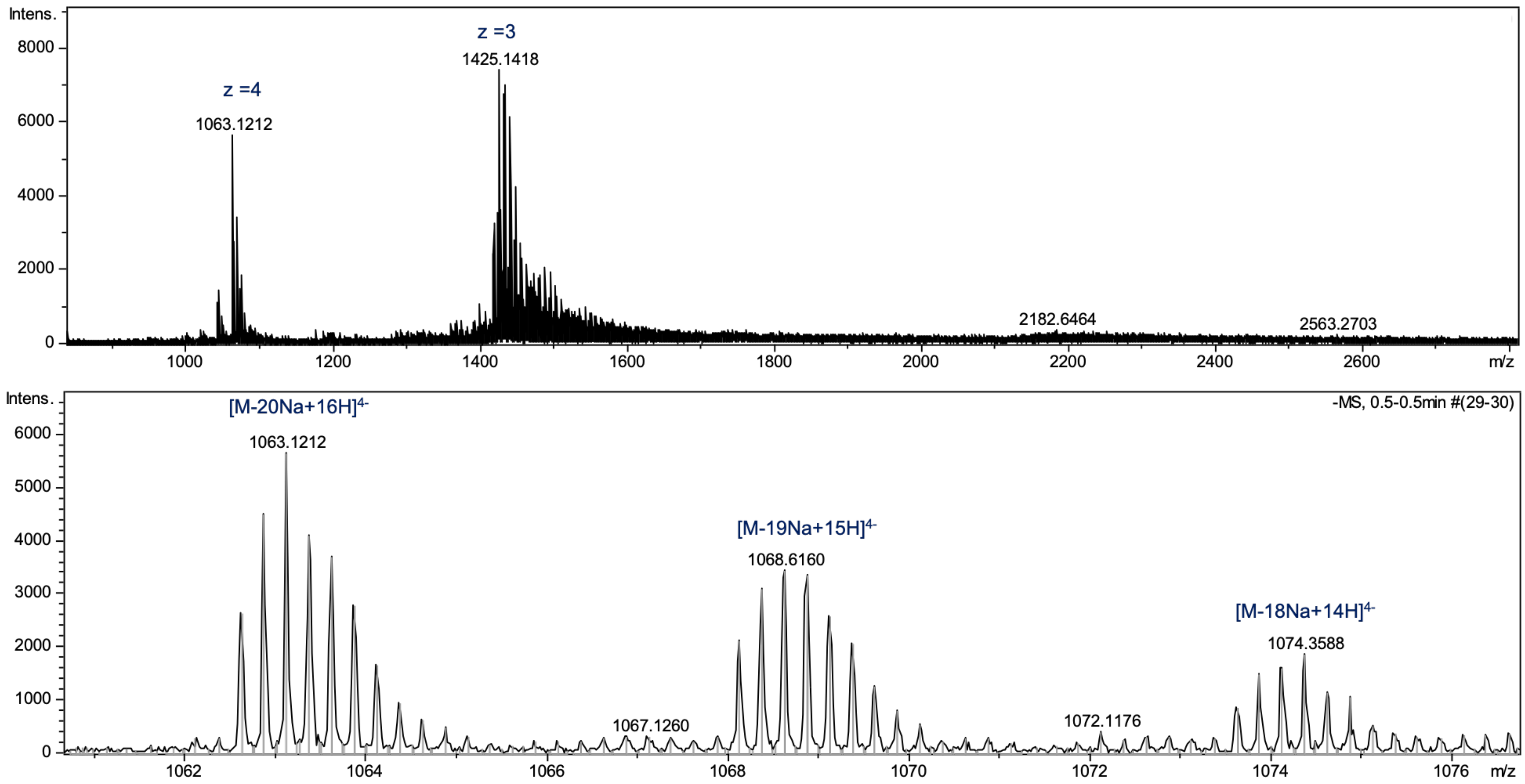

[1]H NMR spectrum of **12** (600 MHz, $D_2O$)

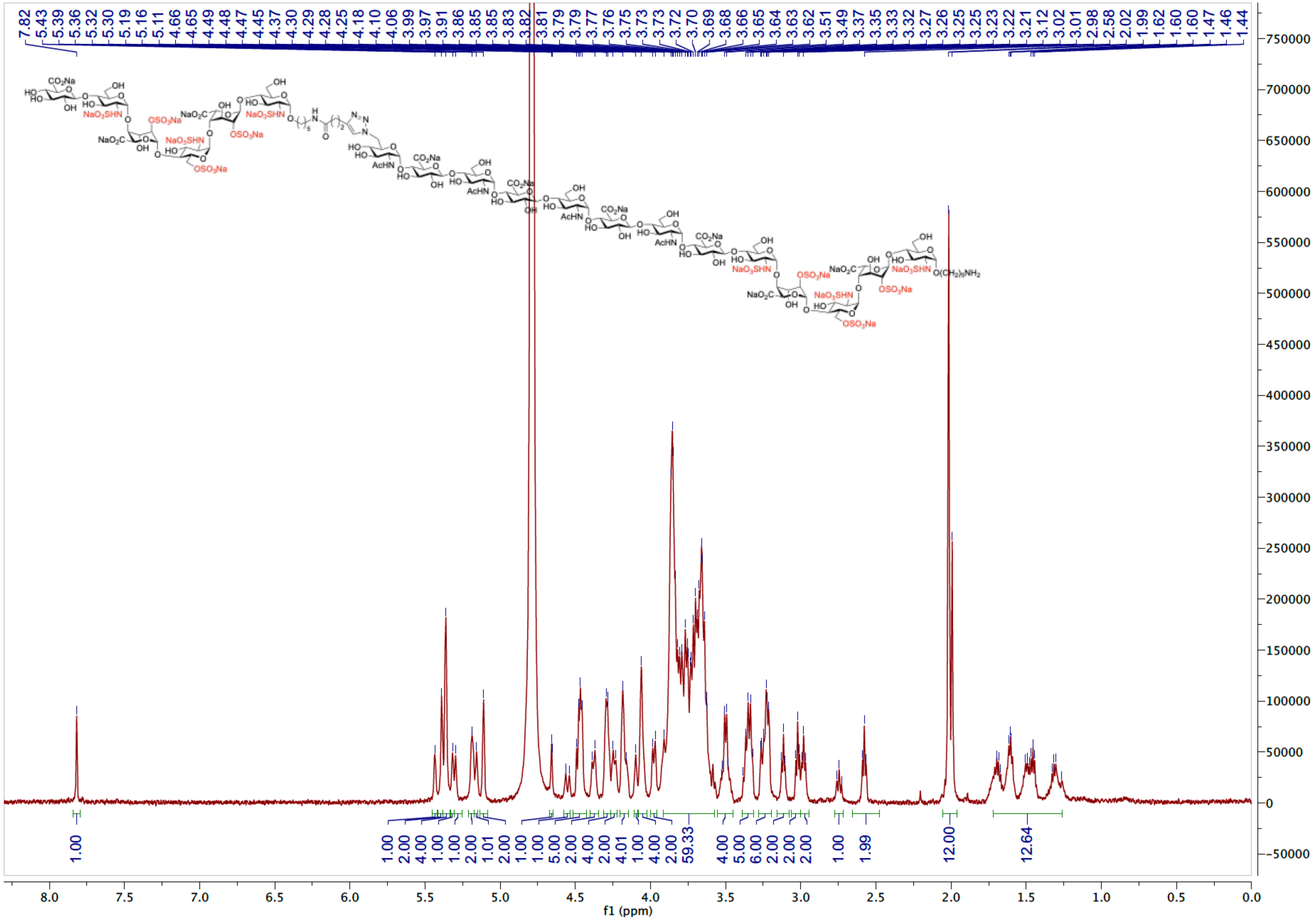

HSQC spectrum of **12** ($D_2O$)

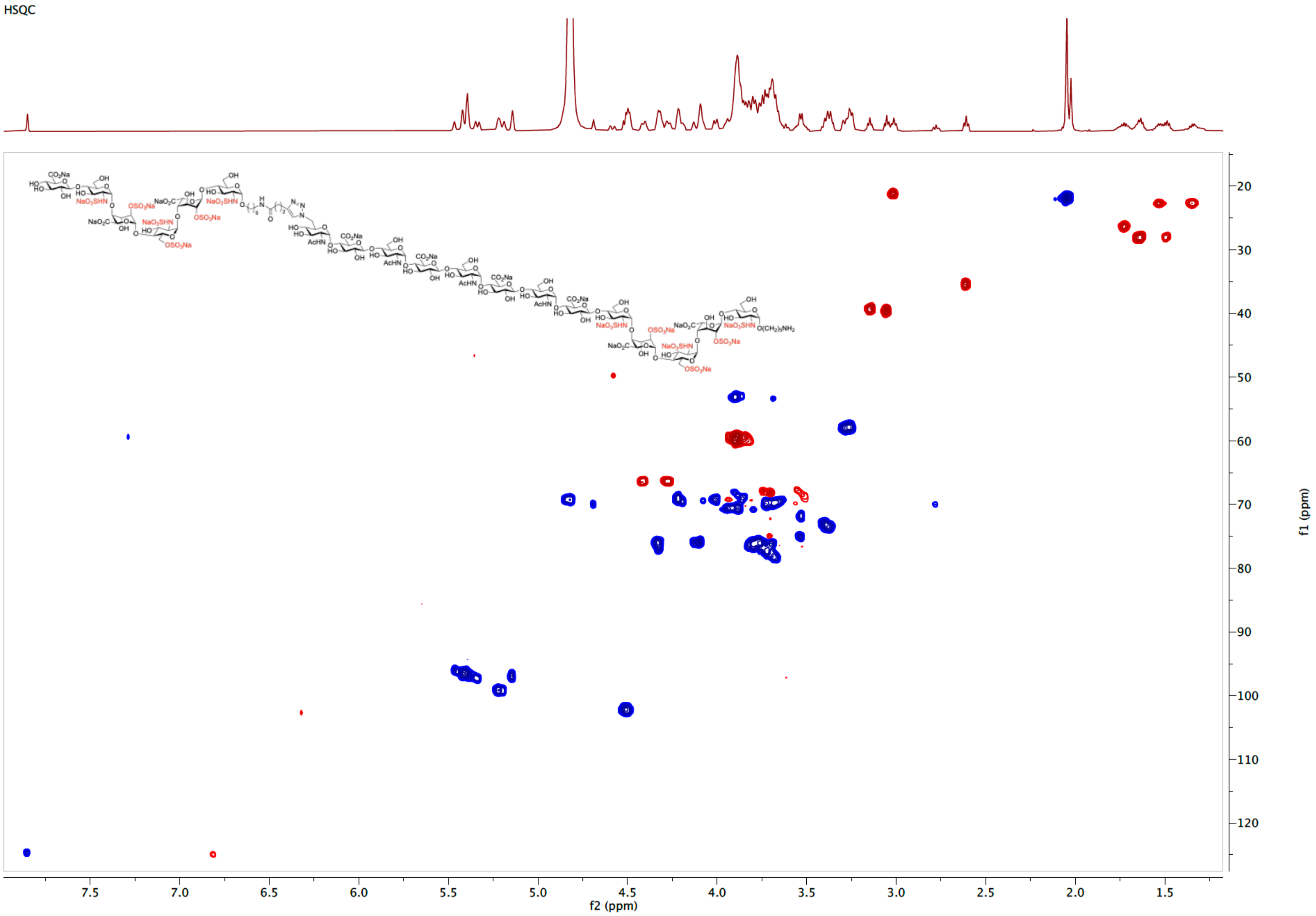

COSY spectrum of **12** ($D_2O$)

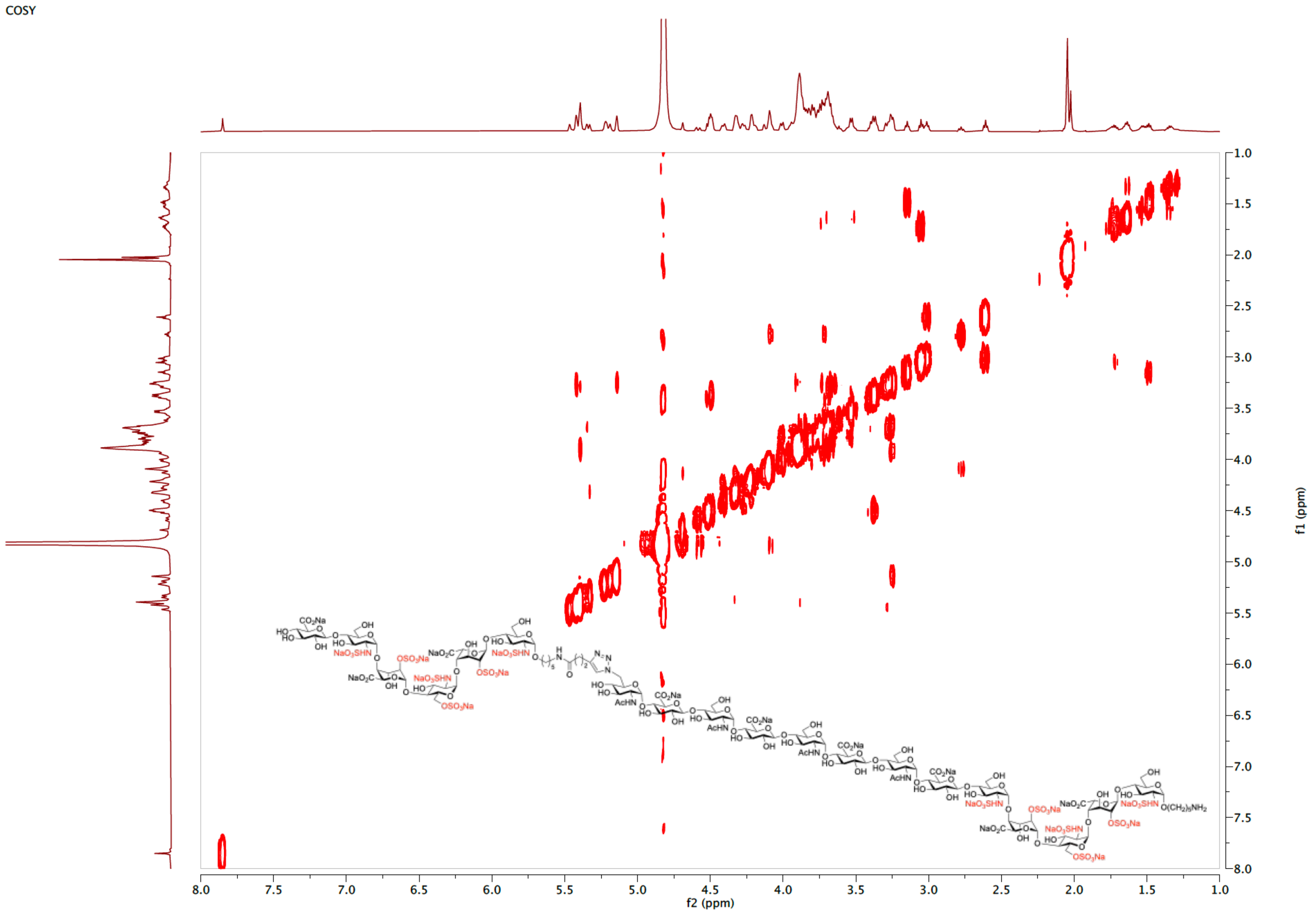

TOCSY spectrum of **12** ($D_2O$)

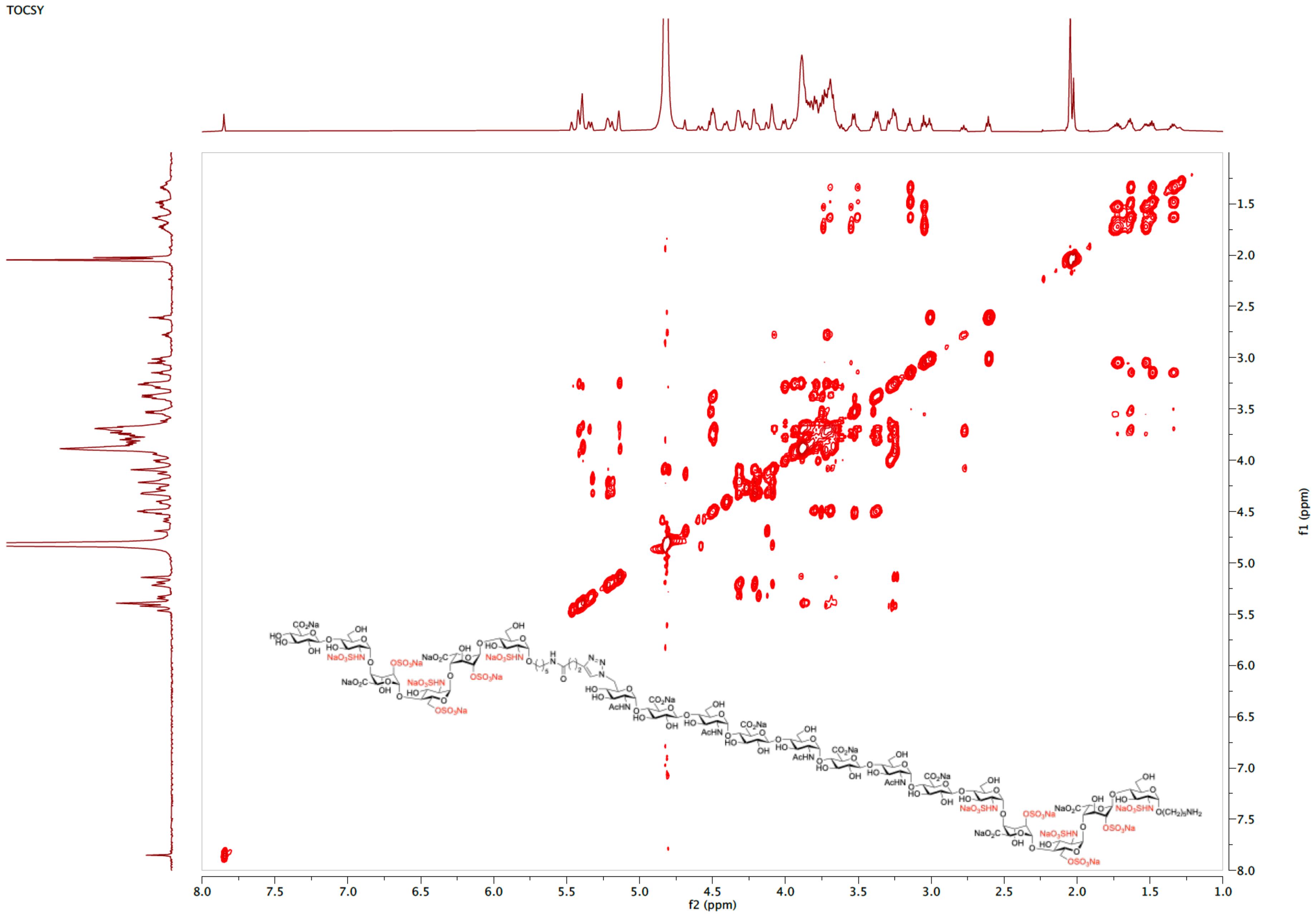

NOESY spectrum of **12** ($D_2O$)

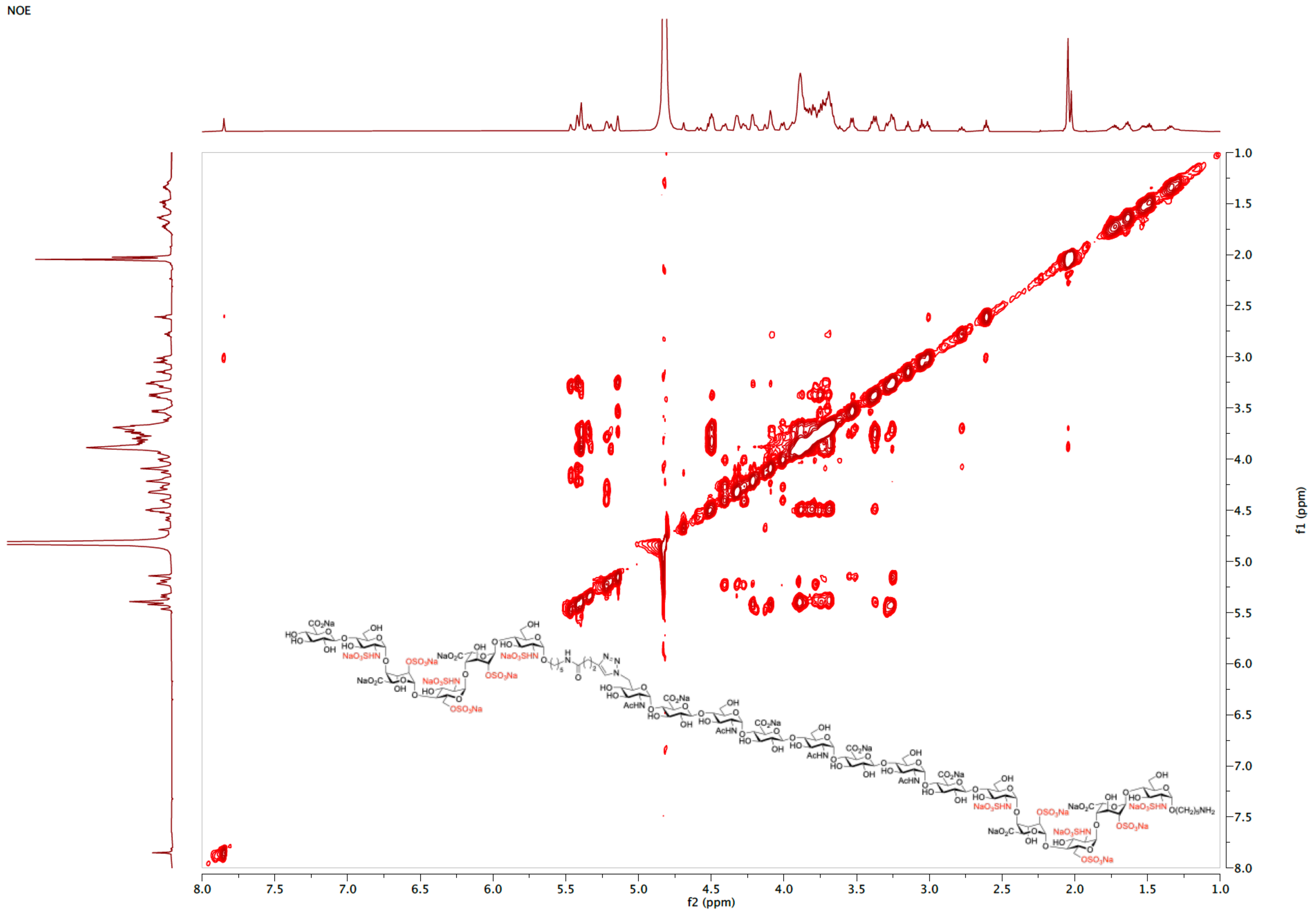

ESI-MS (negative) of **12**

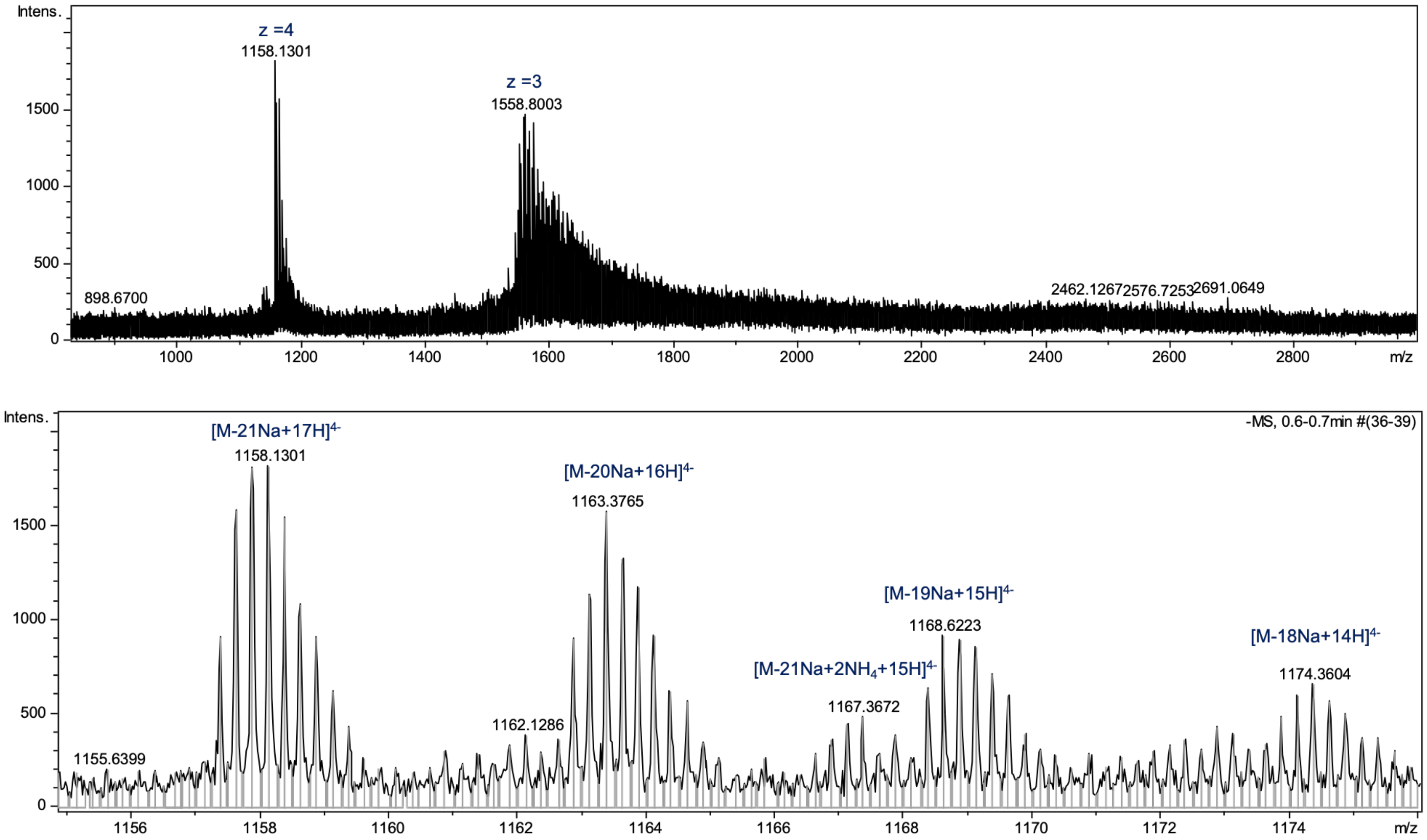

[1]H NMR spectrum of **13** (600 MHz, $D_2O$)

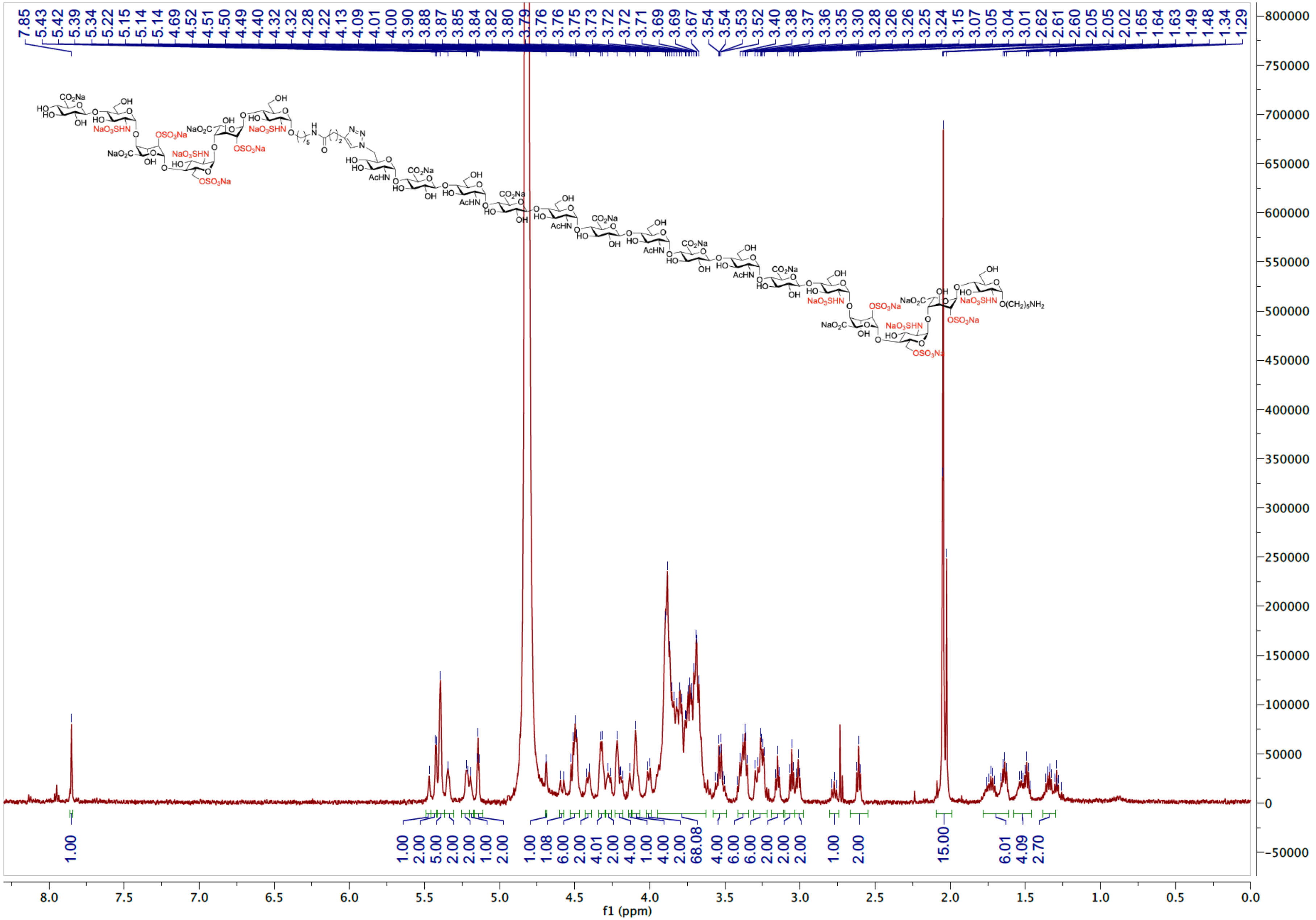

HSQC spectrum of **13** ($D_2O$)

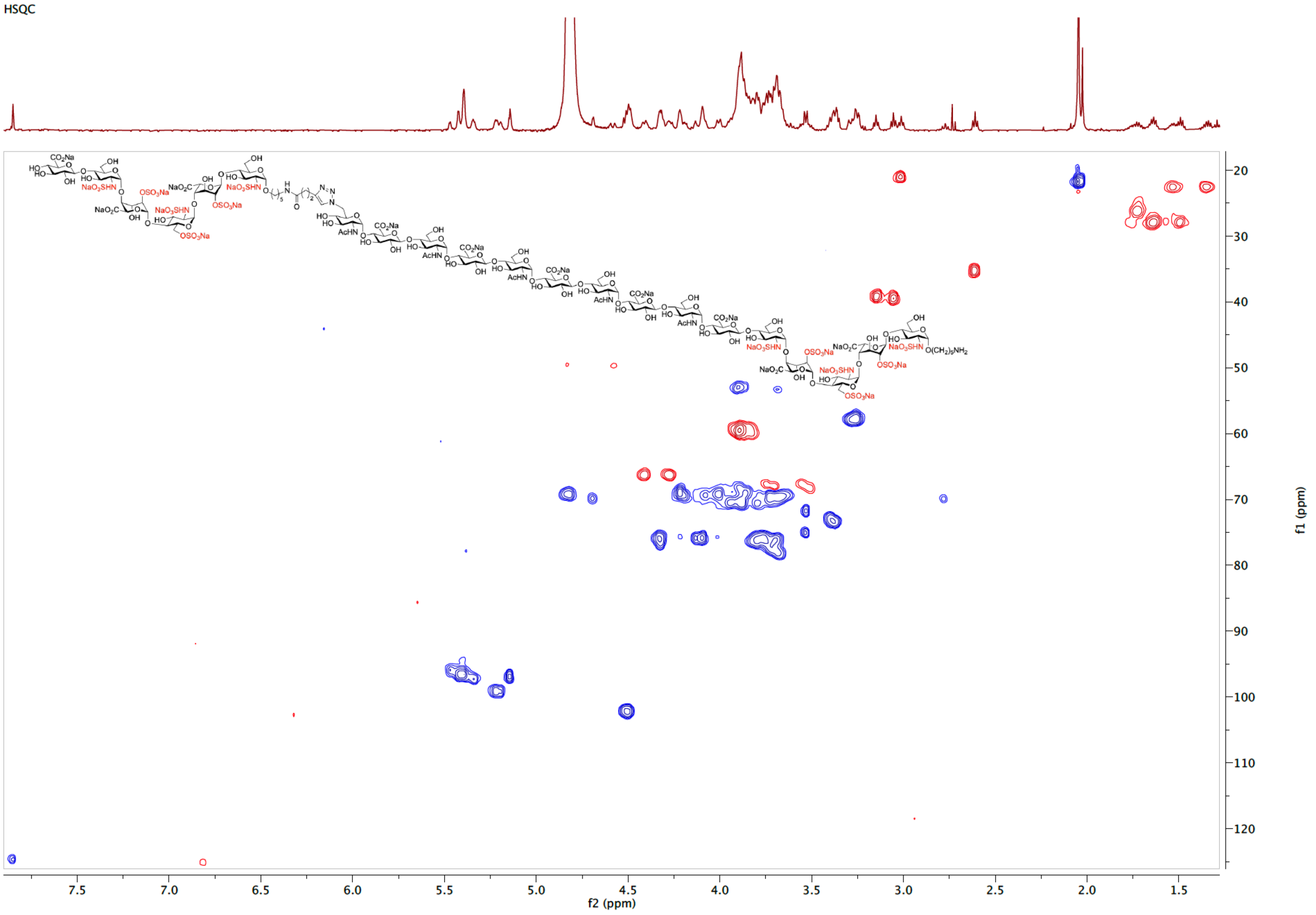

TOCSY spectrum of **13** ($D_2O$)

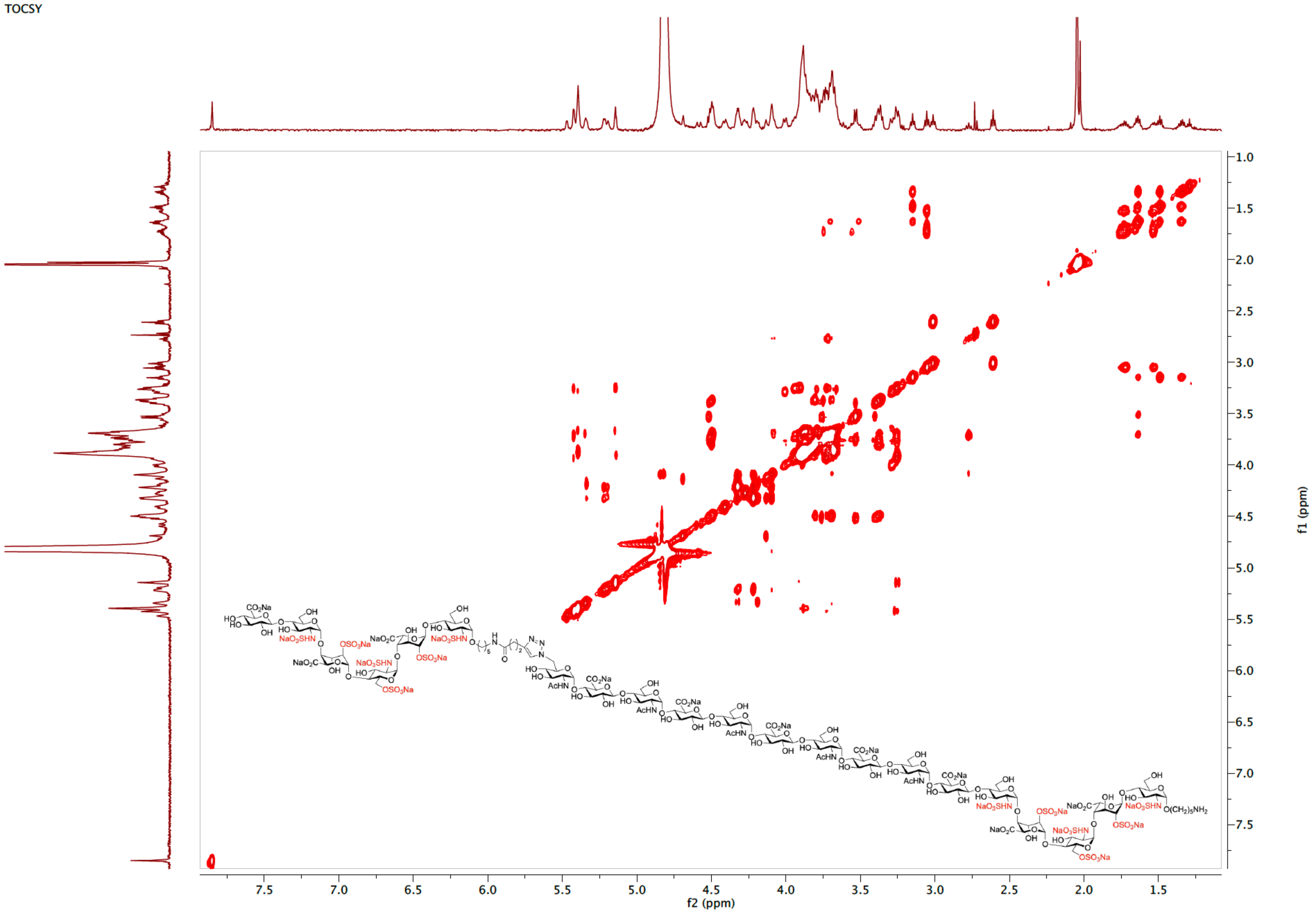

ESI-MS (negative) of **13**

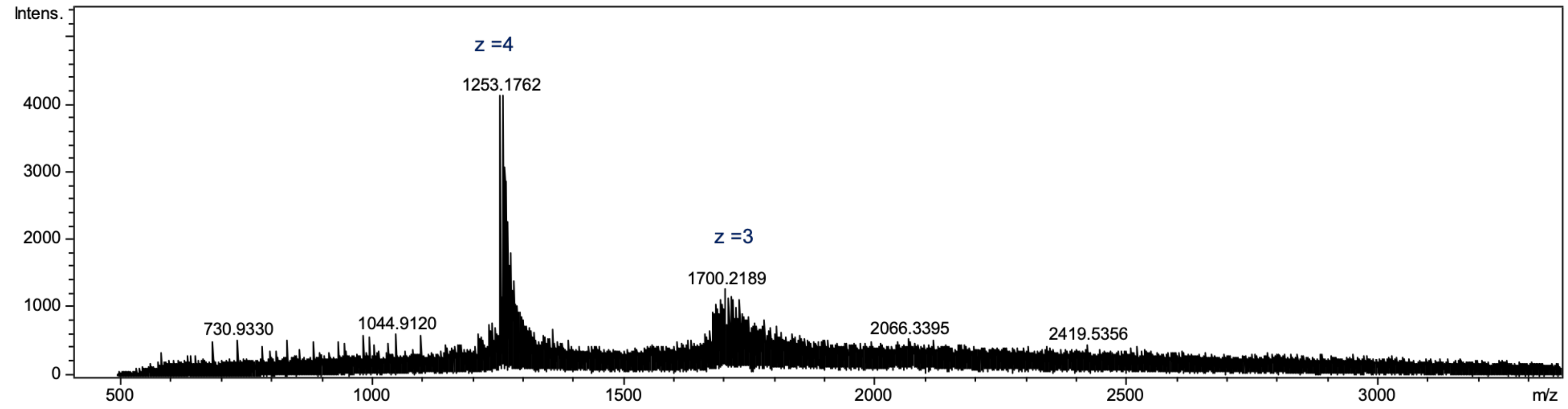

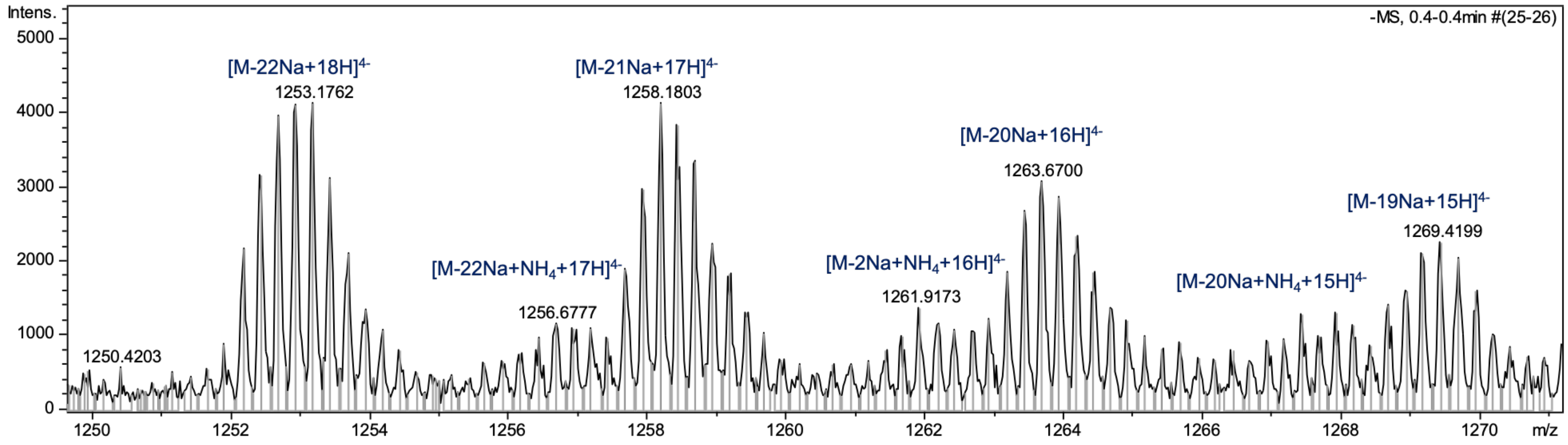

$^{1}$H NMR spectrum of **14** (600 MHz, $D_2O$)

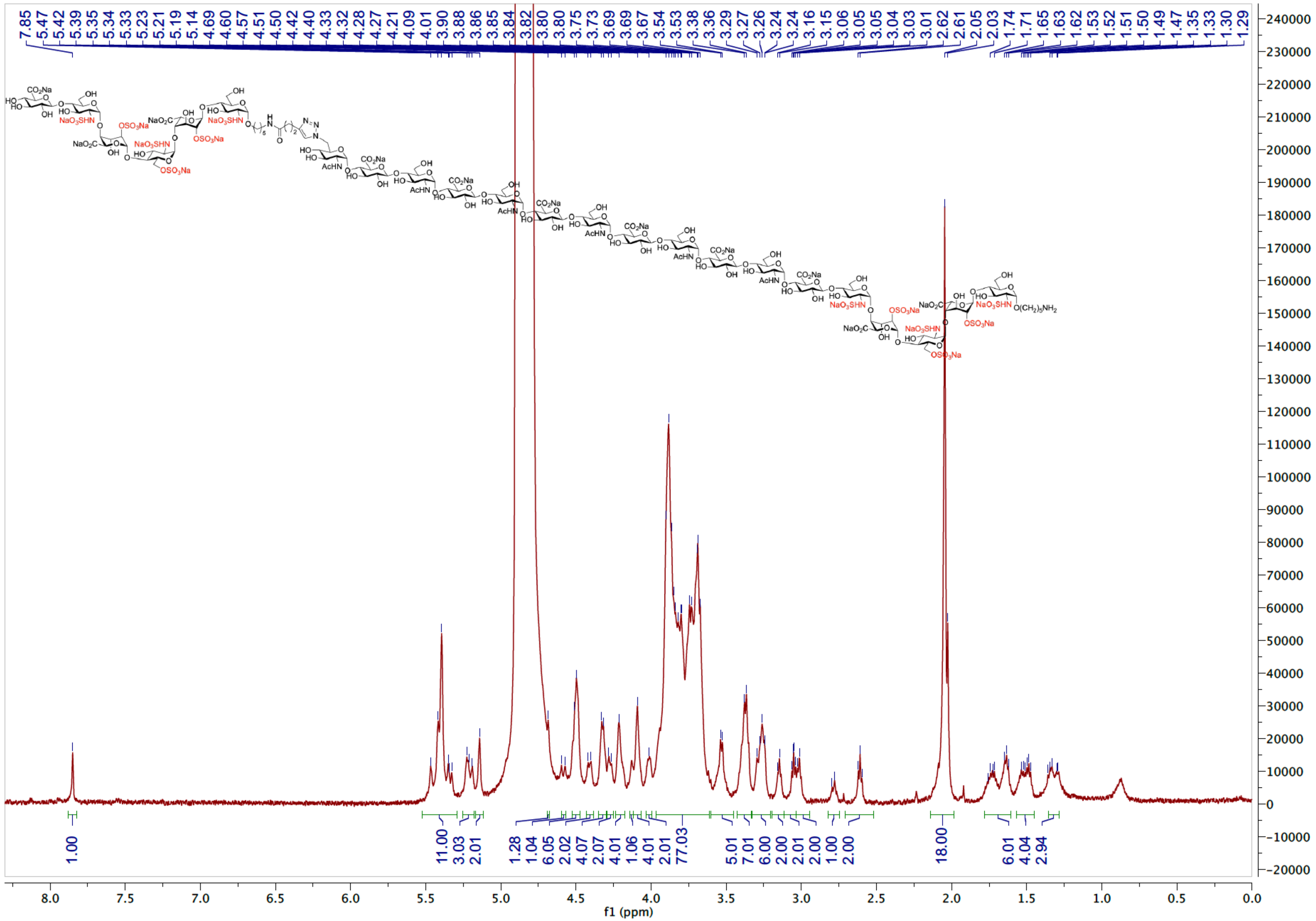

HSQC spectrum of **14** ($D_2O$)

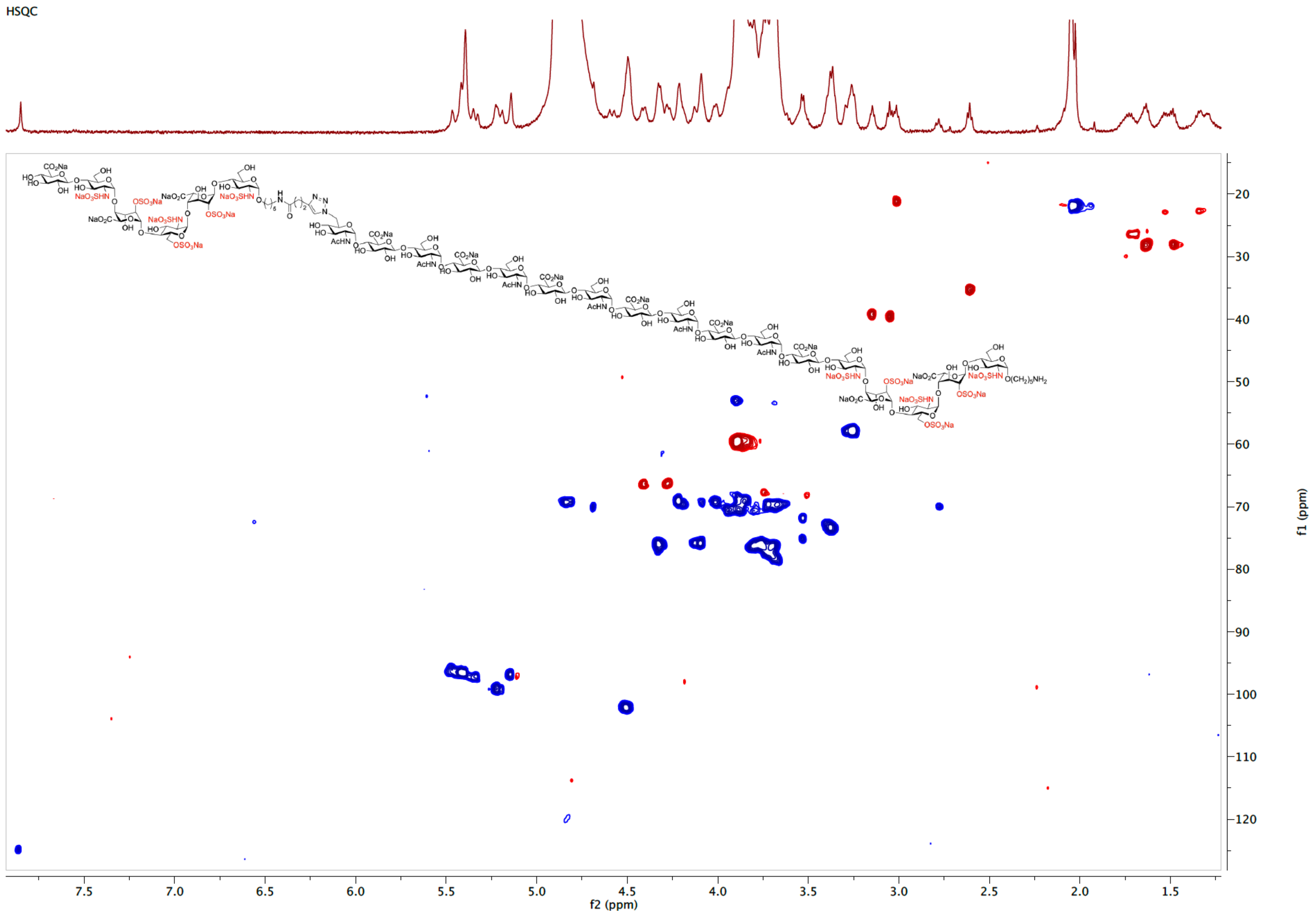

TOCSY spectrum of **14** ($D_2O$)

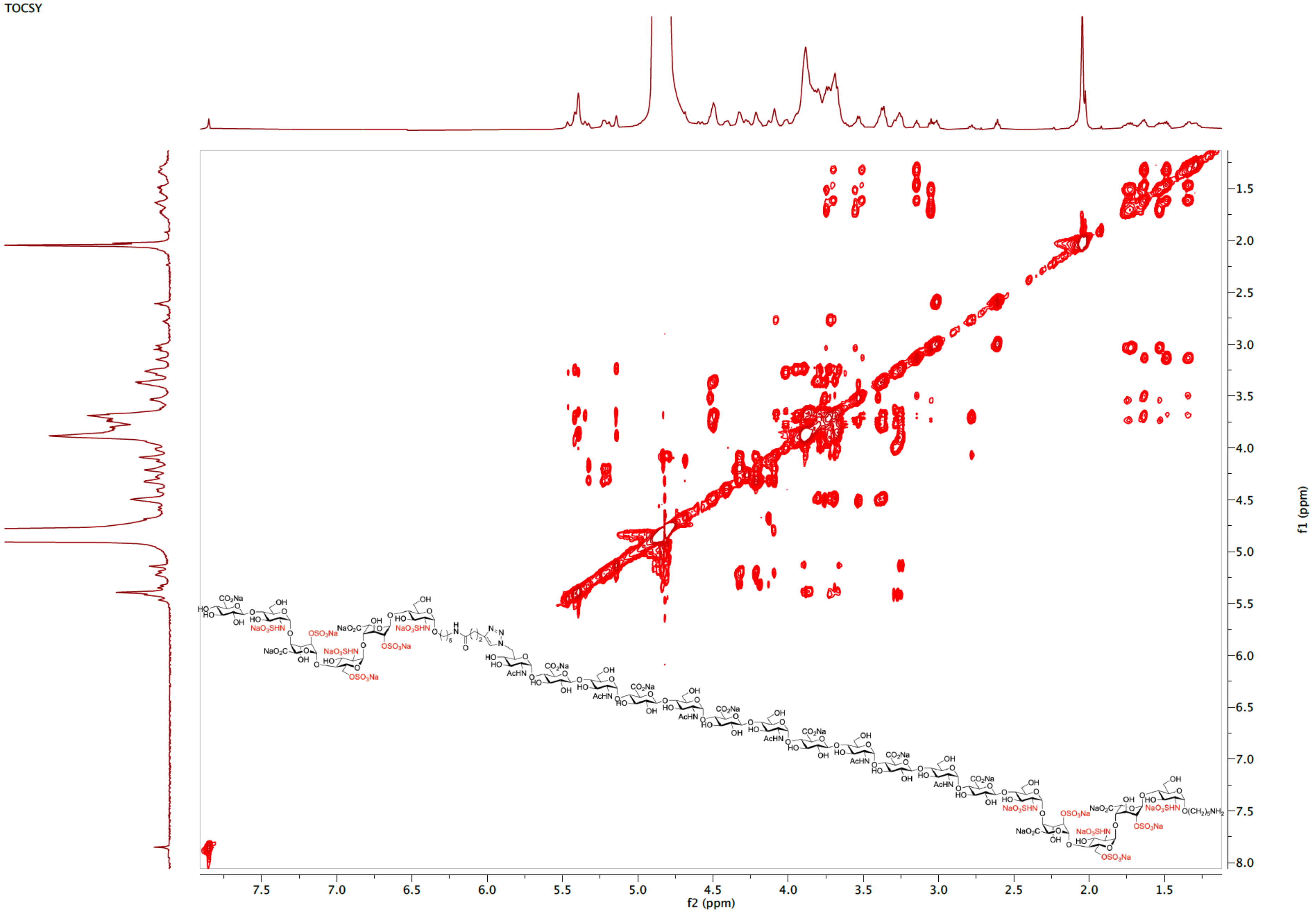

ESI-MS (negative) of **14**

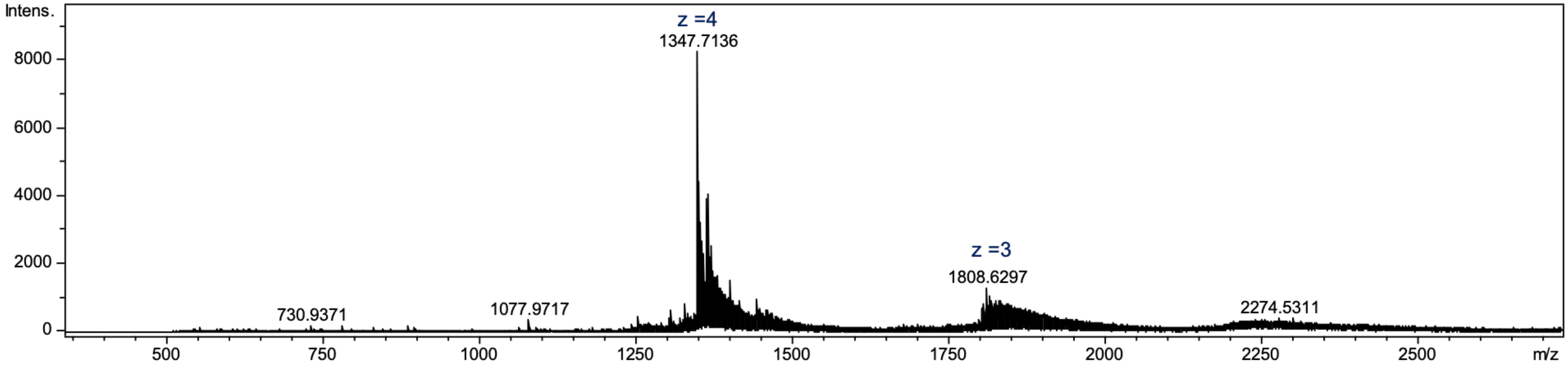

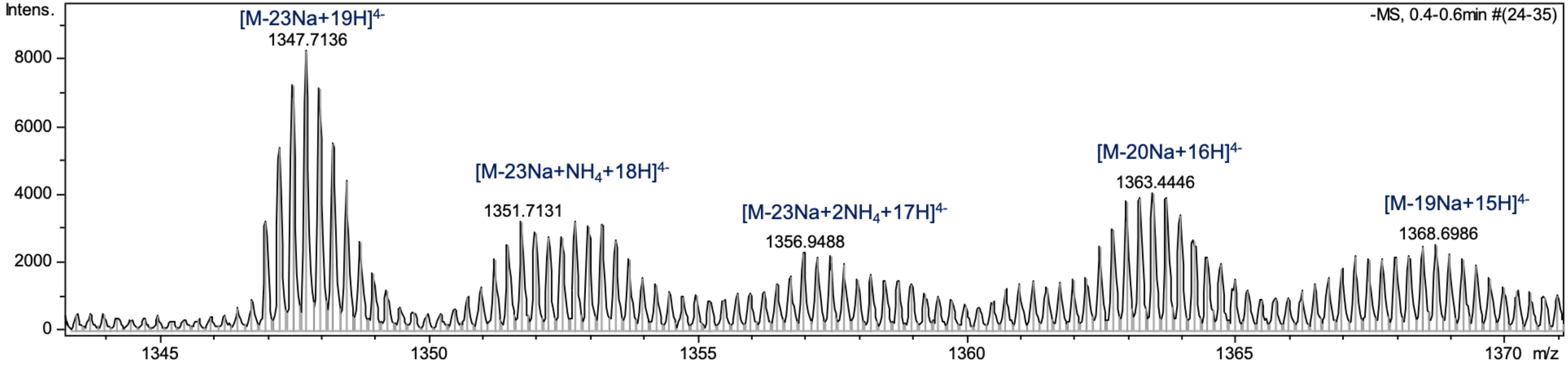

[1]H NMR spectrum of **31** (600 MHz, $D_2O$)

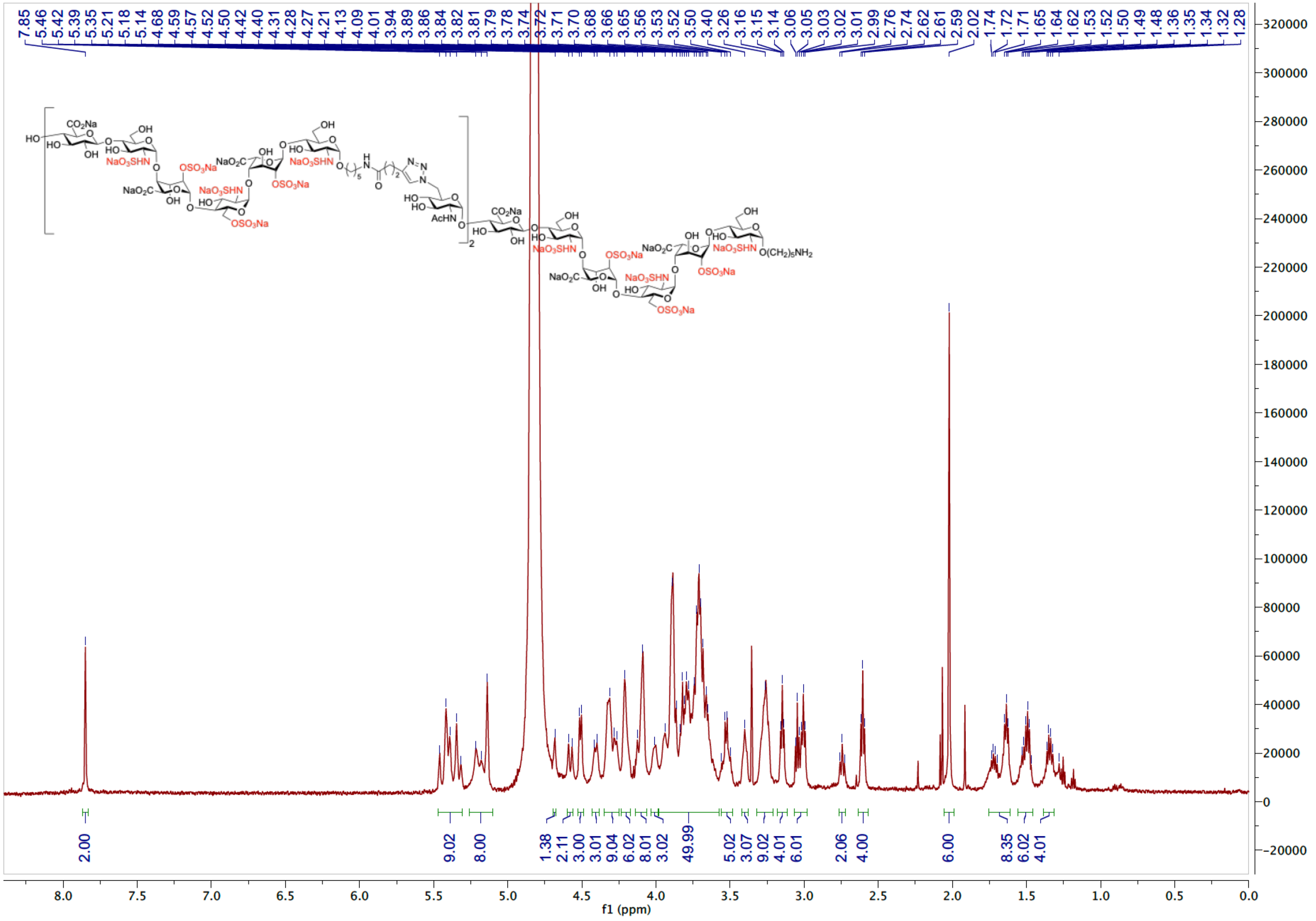

HSQC spectrum of **31** ($D_2O$)

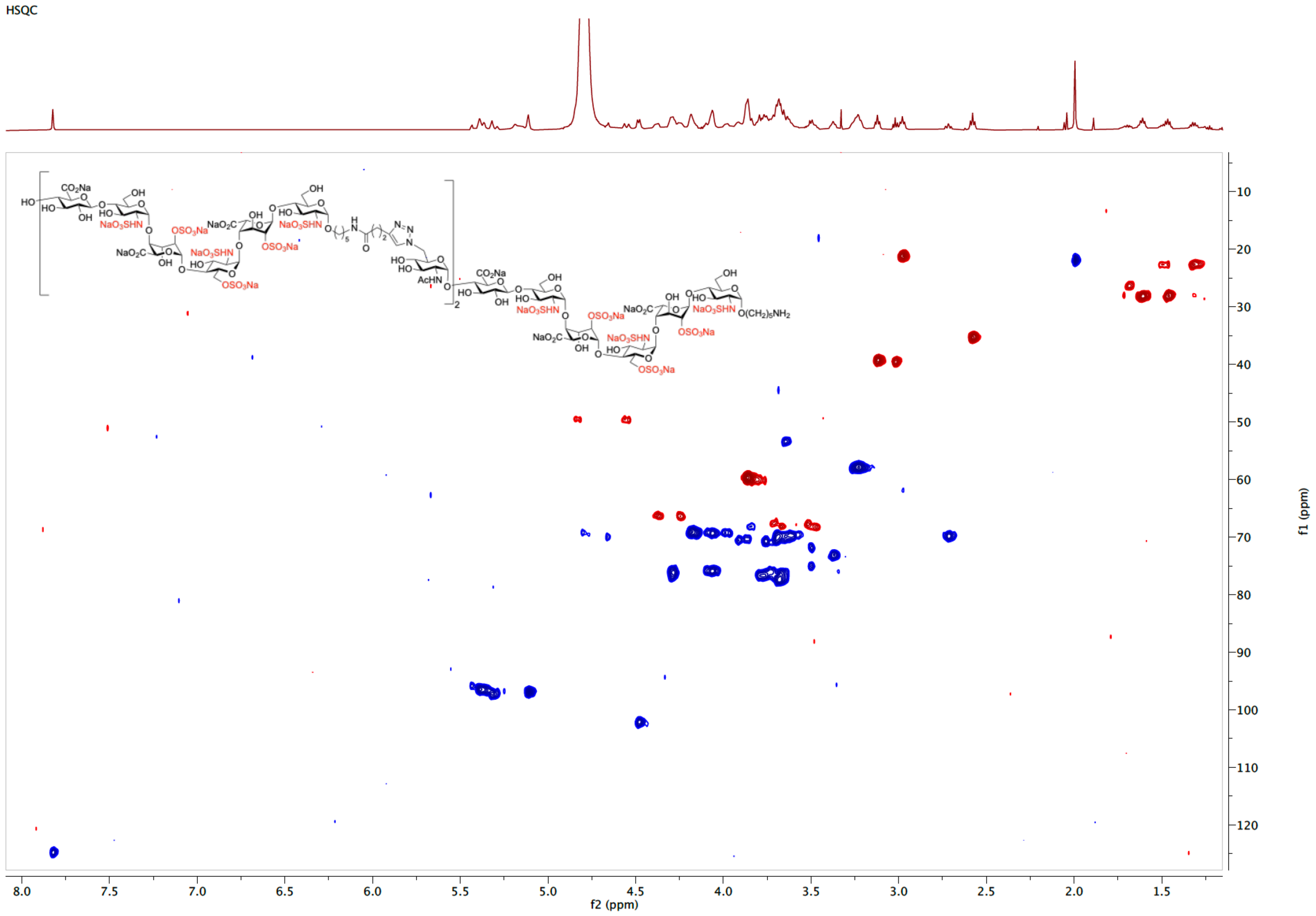

ESI-MS (negative) of **31**

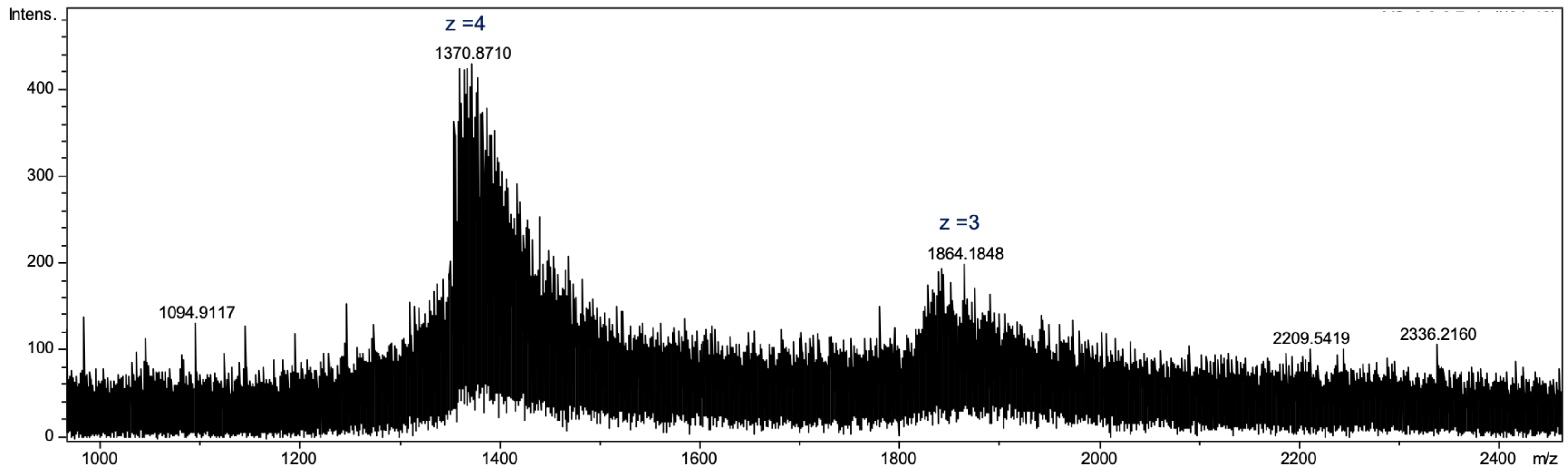

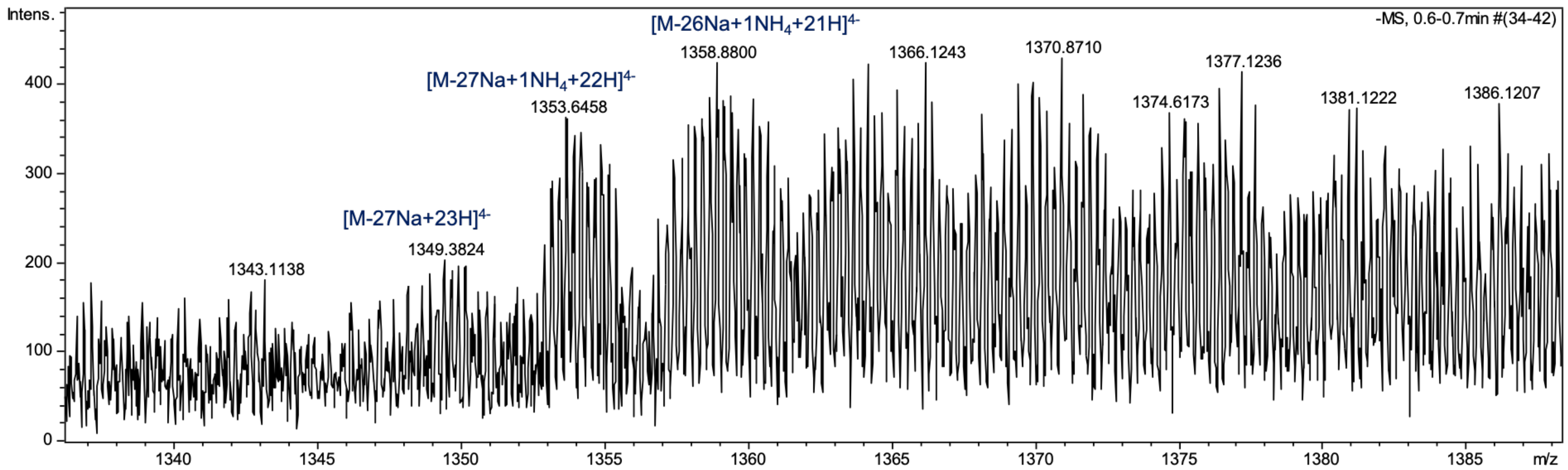

[1]H NMR spectrum of **32** (600 MHz, $D_2O$)

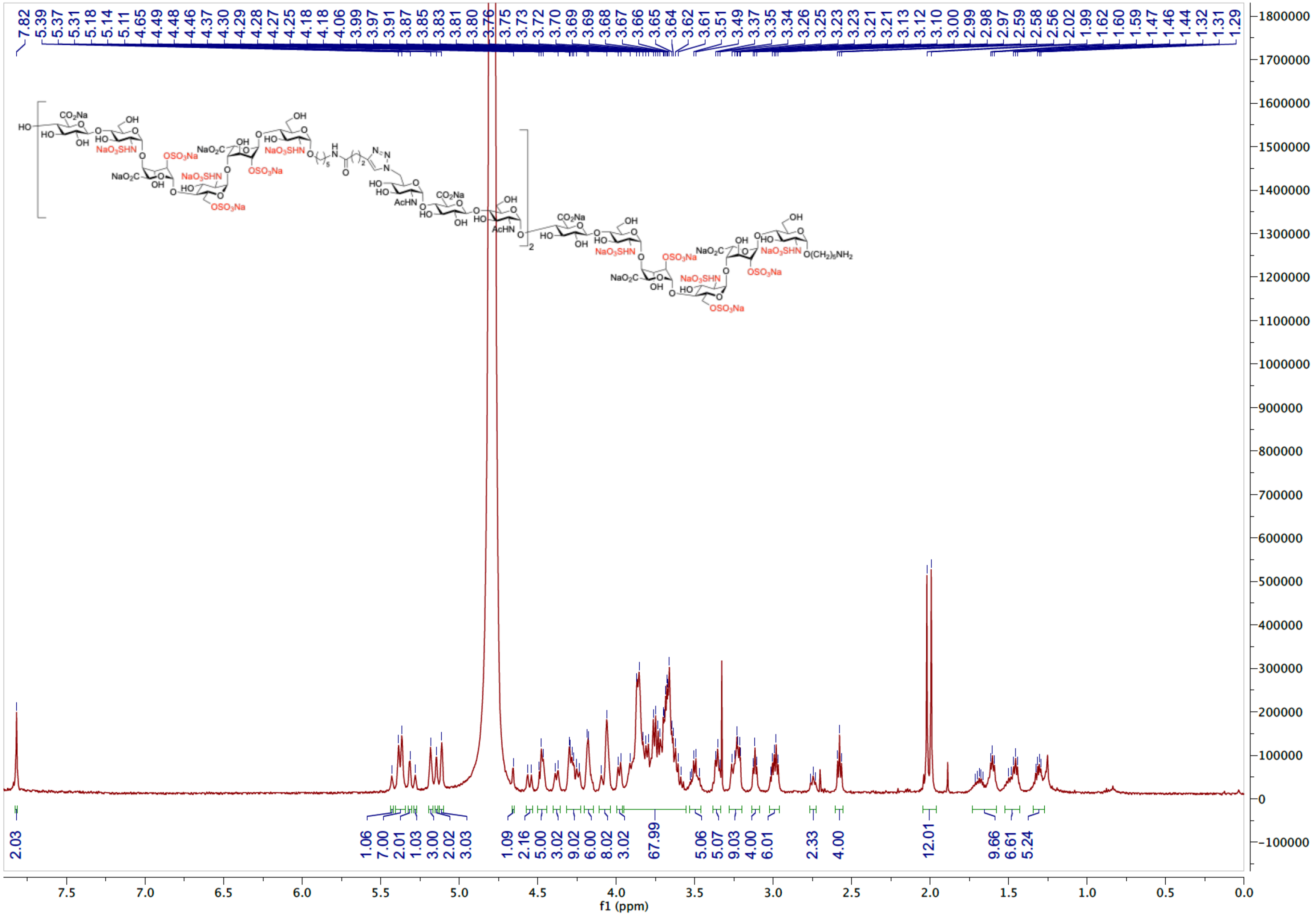

HSQC spectrum of **32** ($D_2O$)

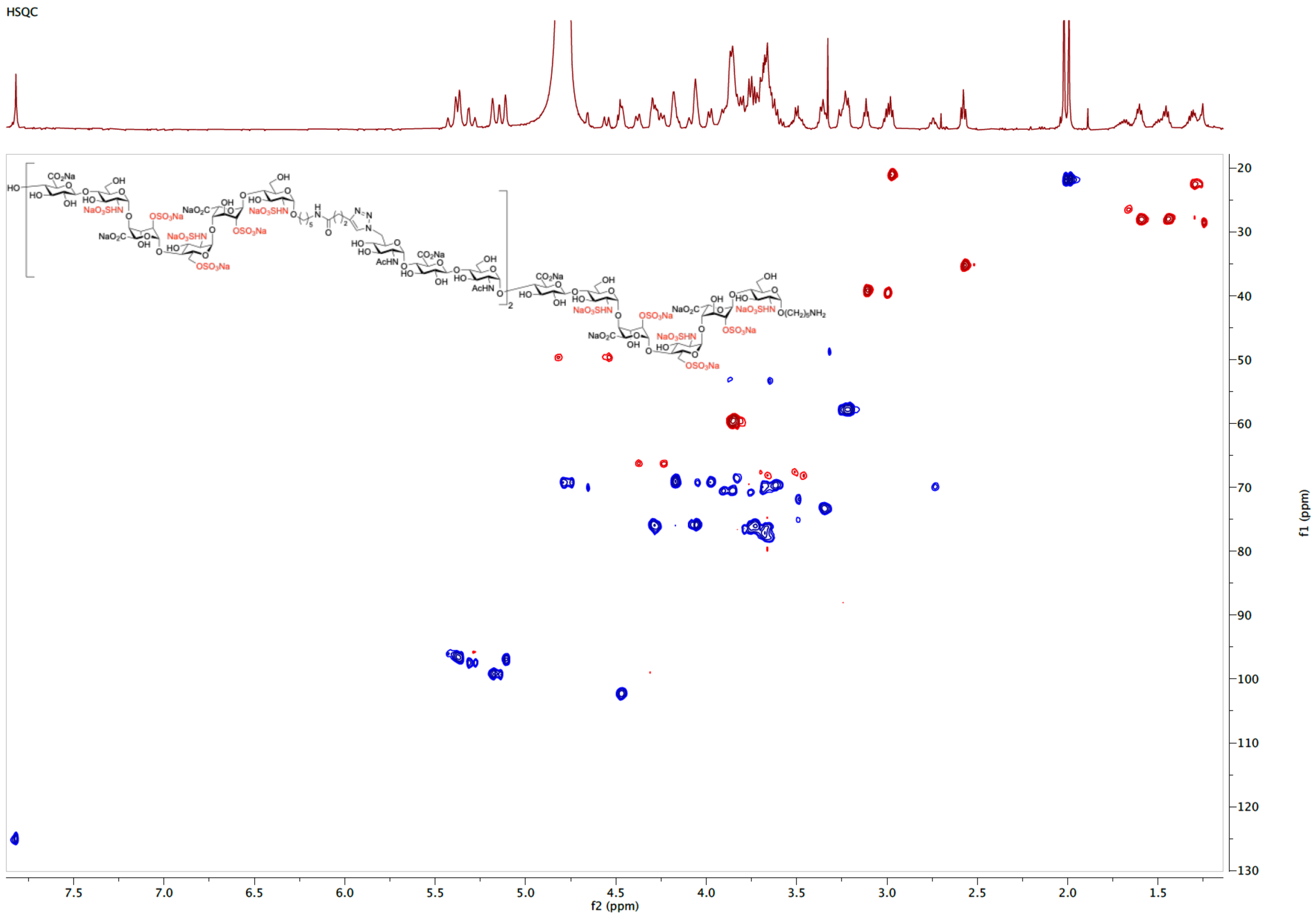

ESI-MS (negative) of **32**

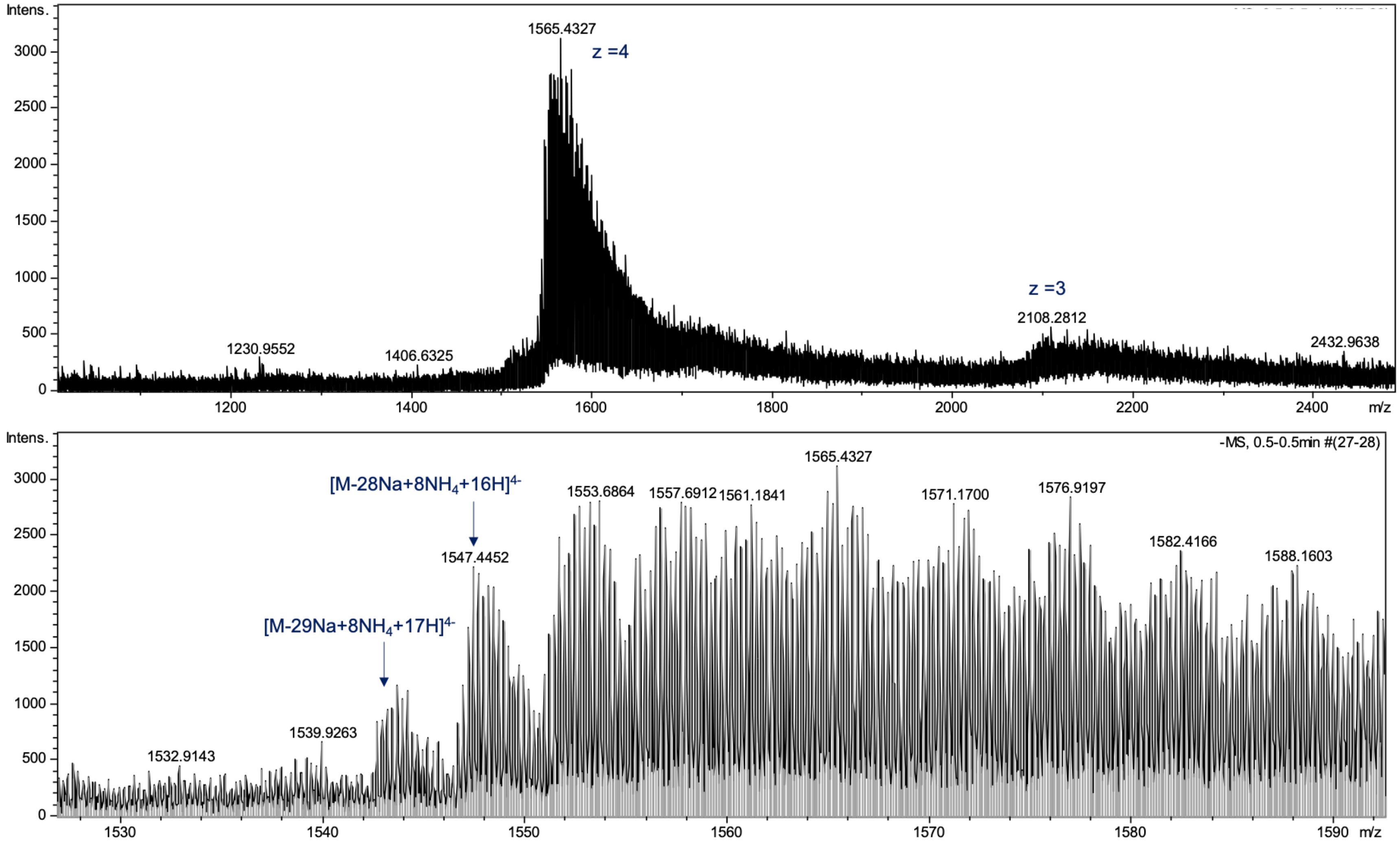

[1]H NMR spectrum of **33** (600 MHz, $D_2O$)

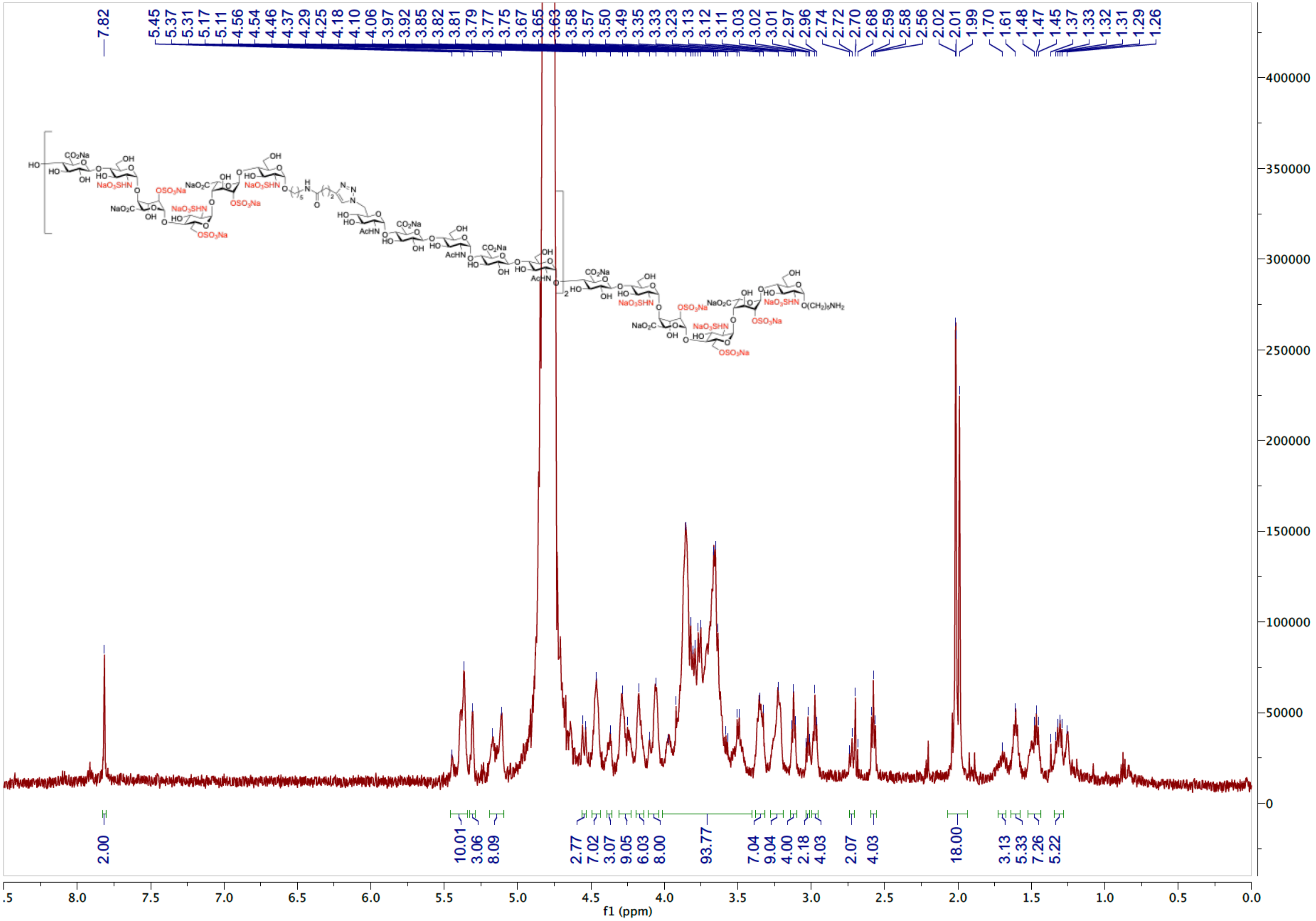

HSQC spectrum of **33** ($D_2O$)

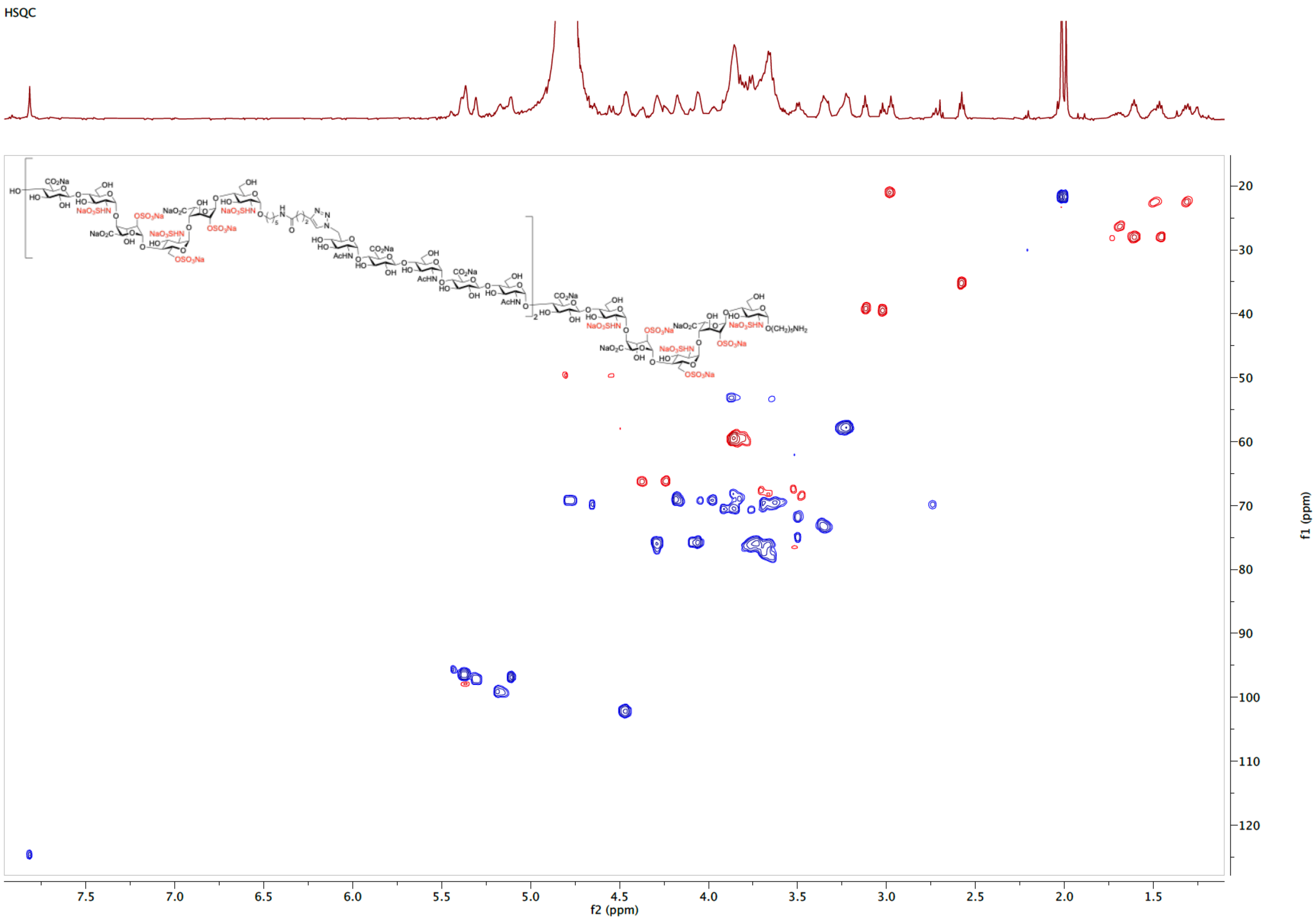

ESI-MS (negative) of **33**

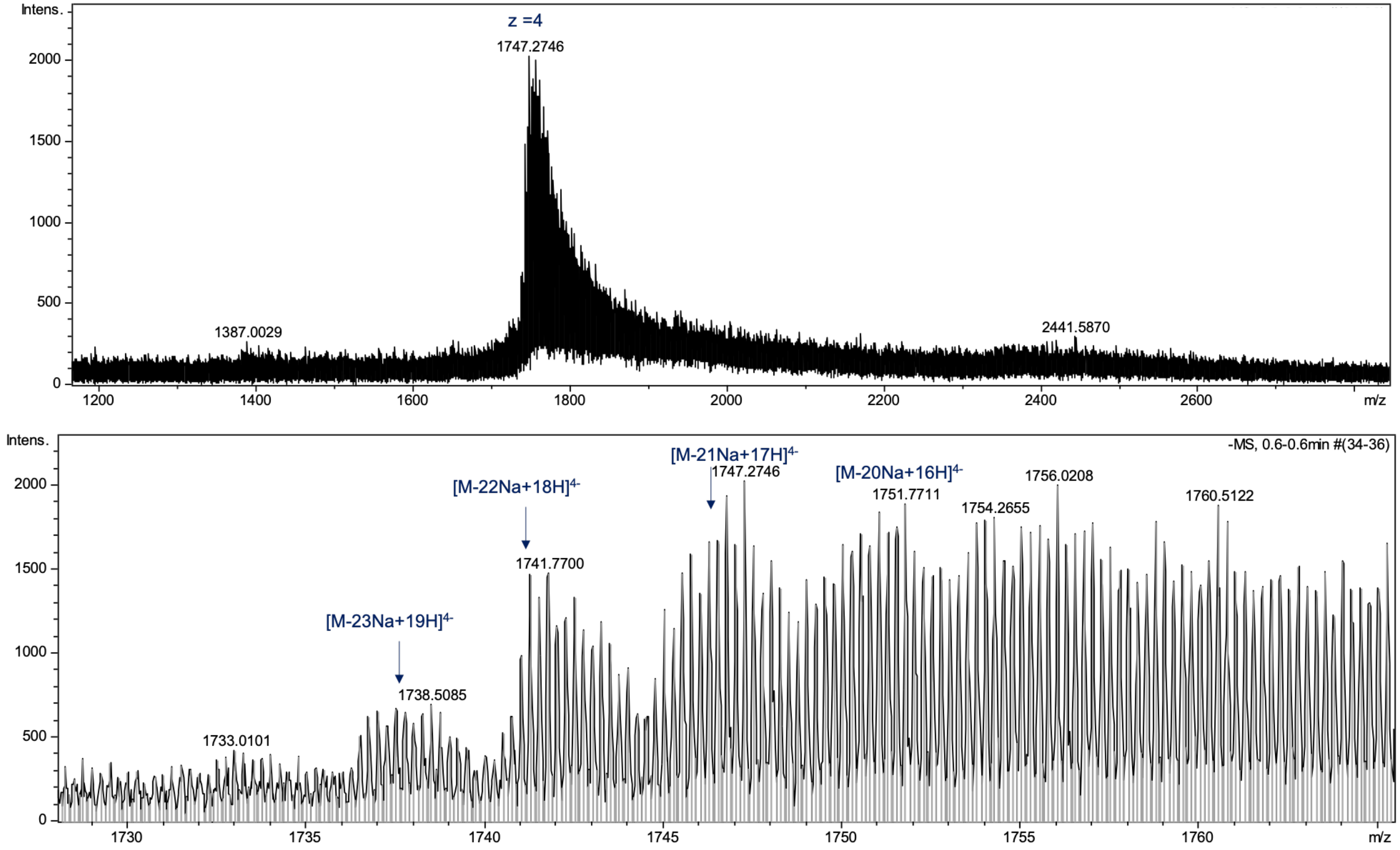
